# Interpretable Aging Signatures in Human Retinal Cell Types Revealed by Single-Cell RNA Sequencing and Sparse Logistic Regression

**DOI:** 10.1101/2025.05.29.656930

**Authors:** Luning Yang, Sen Lin, Yiwen Tao, Qi Pan, Tengda Cai, Yunyan Ye, Jianhui Liu, Yang Zhou, Yongqing Shao, Quanyong Yi, Zen Huat Lu, Lie Chen, Gareth McKay, Richard Rankin, Fan Li, Weihua Meng

## Abstract

**Purpose:** To characterize cell type specific transcriptional changes during human retinal aging and develop machine learning model for cellular age discrimination in a Chinese cohort.

**Design:** Cross-sectional, laboratory-based observational study.

**Participants:** Eighteen unfrozen retinas from 12 Chinese donors (9 young, 34-55y; 9 old, 68-92 y).

**Methods:** Single-cell RNA sequencing (10x, v3.1) generated 223612 cells, batch-corrected with scVI; age-related signatures were defined by intersecting single-cell and pseudo-bulk differentially expressed genes, then cell-type-specific panels were rank-ordered with L1-regularised logistic regression plus recursive feature elimination and interpreted through hallmark-pathway enrichment and transcription-factor regulon mapping.

**Main Outcome Measures:** Age-related cellular composition shifts; cell-type-specific differentially expressed genes; machine-learning classifier accuracy and feature rankings; transcription factor regulon activity changes.

**Results:** Eleven major retinal cell populations were identified. Aging showed declining rod-to-cone ratios, reduced bipolar cell proportions among interneurons, and increased astrocyte abundance. Müller glial cells exhibited the most pronounced transcriptional changes, followed by bipolar cells and rods. Machine-learning classifiers achieved 80-96% accuracy across cell types (microglia 96%, horizontal cells 93%, bipolar cells 91%, cones 90%, rods 89%). Shared aging signatures included mitochondrial dysfunction and inflammatory activation. Cell specific vulnerabilities emerged: mitochondria-centric stress in rods/bipolar cells, proteostasis-retinoid metabolism in cones, and structural-RNA maintenance in horizontal cells.

**Conclusions:** This study provides the first machine learning derived, cell-type specific aging signatures for human retina in a Chinese cohort, revealing both conserved molecular hallmarks and distinctive cellular vulnerabilities that inform targeted therapeutic strategies for retinal aging.

## Introduction

Aging is a fundamental biological process characterized by progressive functional and structural decline across multiple organ systems, including the retina ^1, 2^. Age-related retinal deterioration disrupts visual function, impairing acuity and quality of life, and increases vulnerability to retinal diseases such as age-related macular degeneration (AMD) and diabetic retinopathy (DR) ^3–5^. Understanding the mechanisms underlying retinal aging is essential for developing strategies to preserve vision in aging populations ^6, 7^.

Advances in high-resolution transcriptomic technologies, particularly single-cell RNA sequencing (scRNA-seq), have significantly enhanced our understanding of retinal aging by enabling detailed profiling of cellular heterogeneity and molecular dynamics ^8–10^. Recent scRNA-seq studies have revealed progressive transcriptional changes across retinal cell types during aging. In 2020, human and macaque analyses identified a foveal-to-peripheral aging gradient and *MYO9A*-negative rod vulnerability ^9^. By 2025, mouse studies uncovered transcriptionally distinct rod subpopulations with differential aging responses and bipolar cell (BC) shifts linked to scotopic vision decline ^8^. However, considerations regarding population diversity remain. Large-scale human retina transcriptomic datasets predominantly feature samples from European ancestry populations ^11, 12^, resulting in limited representation of other populations. While initiatives like the Human Cell Atlas have mapped retinal development across 122 donors, only 5 Asian donors were included in its most comprehensive dataset to date ^13^. Furthermore, smaller studies often lack adequate representation of younger age groups, potentially missing early molecular changes that preceded structural or functional deficits. Ethnic variations in retinal characteristics—such as macular pigment density and retinal layer thickness—necessitate diverse cohorts for generalizable insights ^14–16^.

To address these limitations, we characterize the age-dependent transcriptional landscape of the human retina using scRNA-seq in a cohort of 18 Chinese donors aged 34-92 years. Unique strengths of our dataset are the inclusion of early-age samples providing insights into initial stages of retinal aging, ethnic-specific characterization addressing representation gaps in Chinese populations. This study aims to define age-dependent changes in retinal cellular composition, identify transcriptional signatures of retinal aging, and elucidate underlying regulatory networks. This study integrates conventional differential expression analysis with L1-regularized logistic regression and recursive feature elimination to identify cell-type-specific aging signatures that transcend traditional statistical thresholds.

## Methods

### Retinal Sample Collection

Human retinas were collected from living and cadaveric donors under protocols approved by the ethics committees of the University of Nottingham Ningbo China, the Ningbo Lihuili Hospital, and the Ningbo Eye Hospital. Living donor samples were processed within 10 min post-enucleation, and cadaveric samples within 6 h postmortem. Retinal specimens were collected without regional dissection to ensure comprehensive coverage.

### Donor Demographics

We analyzed 18 retinal samples from 12 donors of Han Chinese ethnicity, stratified into young (34-55 years, n=9) and old (68-92 years, n=9) groups. Eight samples were from individuals with DR, evenly distributed across age groups. Table 1 provides donor details, including disease status and reasons for enucleation or death.

**Table 1.** Retinal sample information.

| Sample | Donor ID | Batch | Tissue Source | Eye‡ | Sex§ | Age(y) | Diabetic Status* | Eye Removal/Death Reason | Hospital† | Group |
| --- | --- | --- | --- | --- | --- | --- | --- | --- | --- | --- |
| 1 | D1 | B01 | Live | L | M | 76 | DM | Corneal staphyloma | L.H. | Old |
| 2 | D2 | B02 | Live | L | M | 55 | nonDM | Phthisis bulbi | L.H. | Young |
| 3 | D3 | B03 | Live | R | M | 71 | nonDM | Corneal ulcer | Y.H. | Old |
| 4 | D4 | B04 | Live | L | F | 50 | nonDM | Phthisis bulbi | Y.H. | Young |
| 5 | D5 | B04 | Live | R | F | 68 | nonDM | Corneal perforation | Y.H. | Old |
| 6 | D6 | B05 | Cadaver | L | M | 34 | nonDM | Cerebral hemorrhage | Y.H. | Young |
| 7 | D7 | B06 | Cadaver | L | F | 92 | nonDM | Natural death | Y.H. | Old |
| 8 | D7 | B06 | Cadaver | R | F | 92 | nonDM | Natural death | Y.H. | Old |
| 9 | D8 | B07 | Cadaver | L | F | 55 | DM | Traumatic brain injury | Y.H. | Young |
| 10 | D8 | B07 | Cadaver | R | F | 55 | DM | Traumatic brain injury | Y.H. | Young |
| 11 | D9 | B08 | Cadaver | L | M | 55 | DR | Brainstem hemorrhage | Y.H. | Young |
| 12 | D9 | B08 | Cadaver | R | M | 55 | DR | Brainstem hemorrhage | Y.H. | Young |
| 13 | D10 | B09 | Cadaver | L | M | 35 | DR | Traumatic shock | Y.H. | Young |
| 14 | D10 | B09 | Cadaver | R | M | 35 | DR | Traumatic shock | Y.H. | Young |
| 15 | D11 | B10 | Cadaver | L | M | 84 | DR | Prostate cancer | Y.H. | Old |
| 16 | D11 | B10 | Cadaver | R | M | 84 | DR | Prostate cancer | Y.H. | Old |
| 17 | D12 | B11 | Cadaver | L | M | 69 | DR | Lung cancer | Y.H. | Old |
| 18 | D12 | B11 | Cadaver | R | M | 69 | DR | Lung cancer | Y.H. | Old |

\*DM, diabetes mellitus without diabetic retinopathy DR; diabetic retinopathy; nonDM, non-diabetic retinopathy nor diabetic. Because the tissues were obtained intra-operatively or post-mortem and detailed fundus imaging records were unavailable, the exact stage or subtype of diabetic retinopathy (e.g., NPDR vs PDR) could not be ascertained for the DR cases.
†L.H., The Ningbo Lihuili Hospital; Y.H., The Ningbo Eye Hospital.
‡ L, left; R, right.
§M, male; F, female.

### Retinal Tissue Dissociation

Retinal tissues were dissociated using the LK003150-1bx Dissociation Kit (Worthington Biochemical Corporation). Tissues were minced into fragments and incubated at 37°C for 30 min with gentle agitation. Dissociation was performed using a gentleMACS C tube, followed by filtration through a 70 µm cell strainer. Red blood cells were lysed using Red Blood Cell Lysis Solution (Miltenyi Biotec), and viable cells were counted using AO/PI fluorescent dye with CountStar analysis. Cell suspensions with viability >85% were resuspended in 0.04% BSA/PBS at 700–1200 cells µL^-1^.

### scRNA-seq library preparation

Single-cell suspensions were processed using the Chromium Single Cell 3’ Library and Gel Bead Kit v3.1 (10x Genomics). Approximately 10,000 cells per sample were loaded onto the Chromium Controller to generate gel beads in emulsion (GEMs). Reverse transcription within GEMs produced barcoded cDNA, followed by amplification, fragmentation, and adapter ligation. Libraries were sequenced (Illumina NovaSeq, 150 bp paired-end), and quality was assessed with FastQC.

### scRNA-seq Data Processing and Analysis

Raw sequencing data were processed using Cell Ranger v8.0.1 (10x Genomics) for demultiplexing, alignment to the human reference genome (GRCh38-2020-A), barcode filtering, and UMI quantification. Processed sequencing data have been deposited in the Genome Sequence Archive for Human (GSA-Human; accession HRA014708) at the National Genomics Data Center, China National Center for Bioinformation / Beijing Institute of Genomics, Chinese Academy of Sciences, and are accessible via the GSA family portal ^17, 18^. Quality control was performed using a dynamic thresholding approach based on median absolute deviation statistics to adaptively filter cells. Retained cells expressed ≥200 genes, had UMI counts >500, and <15% mitochondrial transcripts. Doublets were identified and removed using DoubletFinder v2.0.4 ^19^. Integrated analysis of scRNA-seq datasets was performed using Scanpy v1.10.2 with batch-effect correction via scVI-tools v0.17.1 ^20, 21^. Highly variable genes were selected for dimensionality reduction using latent space embedding. A k-nearest neighbor graph was constructed for Leiden clustering at multiple resolutions to identify major retinal cell types. Marker gene expression was used for cell type annotation across clusters based on corresponding marker genes.

### Rod Contamination Filtering

To minimize technical artifacts from rod contamination in non-rod populations, we implemented a systematic filtering approach. Rod-specific expression scores were calculated for each cell using canonical photoreceptor markers (*RHO*, *PDE6B*, *GNAT1*) via Scanpy’s score_genes function. Optimal thresholds were determined by identifying local minima in kernel density estimation plots of the score distribution. For non-rod population analyses, cells exceeding the algorithm-determined threshold were excluded, while rod-specific analyses retained only cells above threshold.

### Age-dependent Differential Expression Analysis

Age-dependent transcriptional changes across retinal cell populations were examined using complementary single-cell differential expression (SC-DE) and pseudobulk differential expression (PB-DE) approaches. For SC-DE, cells were grouped by cell type and age category (young vs. old), with differentially expressed genes (DEGs) identified using likelihood-ratio tests in Seurat v5.1.0 (FDR < 0.05, |log□FC| > 0.25) ^22^. For PB-DE, expression counts were aggregated by cluster-sample combinations and analyzed with DESeq2 v1.42.1 (design: ∼ batch + age_group) ^23^. Batch denotes the sample processing/acquisition date; because both eyes from each donor were processed on the same date, each donor’s samples were assigned to a single batch. Following normalization and dispersion estimation, significance was determined using Wald tests with apeglm shrinkage (adjusted P < 0.05, with |log□FC| thresholds ≥0). Results identified three categories of age-associated genes: exclusively SC-DE, exclusively PB-DE, and high-confidence DEGs from both methods. We also performed a complementary donor-level sensitivity analysis using aggregated left/right eye data with additional covariates (design: ∼ sex + DM_status + age_group) as a conservative benchmark, where DM_status encodes non diabatic versus the combined diabetic and diabetic retinopathy groups.

### Pathway Enrichment Analysis

Pathway enrichment was conducted using the irGSEA v3.3.2 ^24^, integrating scoring methods, including AUCell, UCell, singscore, ssgsea, JASMINE, and viper. Hallmark gene sets from MSigDB ^25^ were used as reference pathways. Enrichment scores were calculated for each cell type and age group. Results were aggregated using robust rank aggregation to obtain consensus enrichment scores. Significantly enriched pathways were defined as those with an adjusted P < 0.05.

### Multi□Stage Machine Learning (ML) Based Key Genes Selection

A multi□stage, rigorously controlled ML framework to identify age□stage marker genes in each retinal cell type was implemented. Firstly, for each old□versus□young comparison, we curated the top 200 genes by absolute log□□fold change from DEGs and extracted their expression profiles for z□score normalization. The normalized matrix was then stratified into 70 % training and 30 % testing subsets, with all features mean□centered and variance□scaled to conform to linear□model prerequisites. Then, L1□regularized logistic regression was applied, optimizing the penalty hyperparameter C (0.01–10) via 5□fold cross-validation grid search to enforce model sparsity. Building upon this tuned model, recursive feature elimination with internal cross□validation (RFECV) was employed, systematically removing the least contributory genes until maximal cross□validated performance was attained. To validate the robustness of these key genes, we repeated RFECV across 20 bootstrap replicates, retaining only those genes selected in over 50 % of runs. This 50% threshold is commonly adopted in stability selection frameworks^26, 27^ to balance stringent feature reproducibility with the retention of biologically relevant signals. Model performance on the test set was evaluated by precision, recall, and F1□scores, complemented by ROC curves and AUC metrics for each class. By integrating L1 penalization, RFECV, and stability selection, this ML based framework yielded an important and stable gene signature for age□stage classification.

### Transcription Factor Regulatory Network Inference

Transcription factor (TF) regulatory networks were inferred using pySCENIC (v1.2.2) ^28^ to identify key transcriptional regulators across retinal cell populations during aging. For computational efficiency, 5,000 cells were randomly sampled from each dataset and processed into cell type-specific subsets for interneurons and glial cells. Co-expression modules were identified with GRNBoost2 with official list of human TFs (https://resources.aertslab.org/cistarget/tf_lists/), followed by cis-regulatory motif analysis using RcisTarget with hg38 motif rankings (10kb up/downstream of transcription start sites). Regulon activity scores were computed using AUCell and binarized to determine active regulons in individual cells. Regulon specificity scores were calculated to identify condition-specific transcriptional regulators.

### Gene Set Scoring Analysis in Aging Retinal Cell Types

Gene set scoring was performed to evaluate pathway activity across retinal cell types during aging using the UCell package (v2.6.2) ^29^. Predefined gene sets, including cellular senescence (SenMayo) ^30^ and genes related to polyamine metabolism (see Supplementary Table S1). Statistical comparisons of pathway scores between young and old age groups were conducted using Wilcoxon rank-sum tests, with significance defined at P<0.05.

### Definition of Age-Related Signatures

In this study, we define “age-related signatures” as comprehensive molecular profiles that integrate findings from both differential expression analysis and machine learning approaches to capture age-associated changes in retinal cell populations. These signatures comprise three key components: (1) proportion shifts reflecting changes in cell type composition with age, (2) DE-signatures representing genes consistently identified as differentially expressed in both single-cell and pseudobulk analyses, and (3) ML-signatures consisting of genes with stable predictive importance for age classification, as determined by L1-regularized logistic regression with recursive feature elimination and bootstrap validation. The final age-related signatures represent the integration of these complementary analytical approaches, ensuring both statistical significance and functional relevance for age discrimination across retinal cell types. The end-to-end workflow is summarized in Fig. 1a.

**Figure 1.**
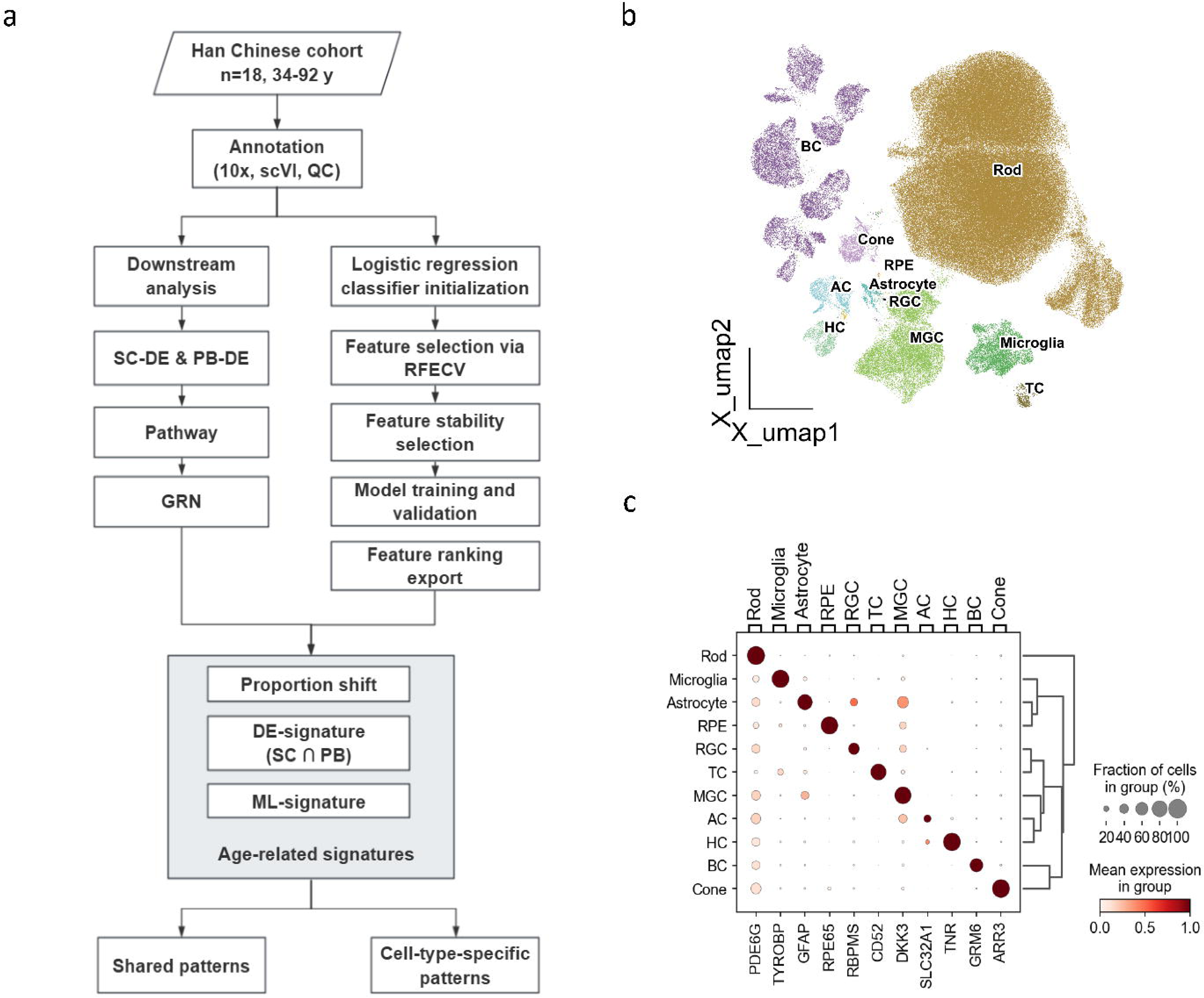
**(a)** Analytical workflow for identifying age-related transcriptional signatures in human retinal cells. **(b)** UMAP visualization of 11 major retinal cell populations annotated from 18 samples. Cell populations are color-coded by annotated cell types. **(c)** Dot plot showing marker genes’ expression for major retina cell type. Dot size represents the percentage of cells expressing each marker, while color intensity indicates average expression level.

## Results

### Age-Related Shifts in Retinal Cellular Composition

We performed scRNA-seq analysis of 18 retinal samples from young and older adults. Unsupervised clustering identified 11 major retinal cell populations (Figs. 1b, 1c) with the summary table in Supplementary Table S2. For aging analysis, we focused on glial cells, photoreceptors, and interneurons, excluding T cells (variable proportions) and retinal pigment epithelium (RPEs)/ retinal ganglion cells (low cell counts). Quantitative assessment revealed distinct age-related compositional changes. The rod-to-cone ratio showed a decreasing trend with advancing age (Fig. 2a). Among interneurons, the relative proportion of BCs shows a downward trend with increasing age (Fig. 2b). Within glial populations, astrocytes exhibited the most pronounced age-related changes, with their relative proportion showing a clear upward trend with advancing age, while Müller glial cells (MGCs) consistently represented the majority of glial cells across age groups (Figs. 2c-e).

**Figure 2.**
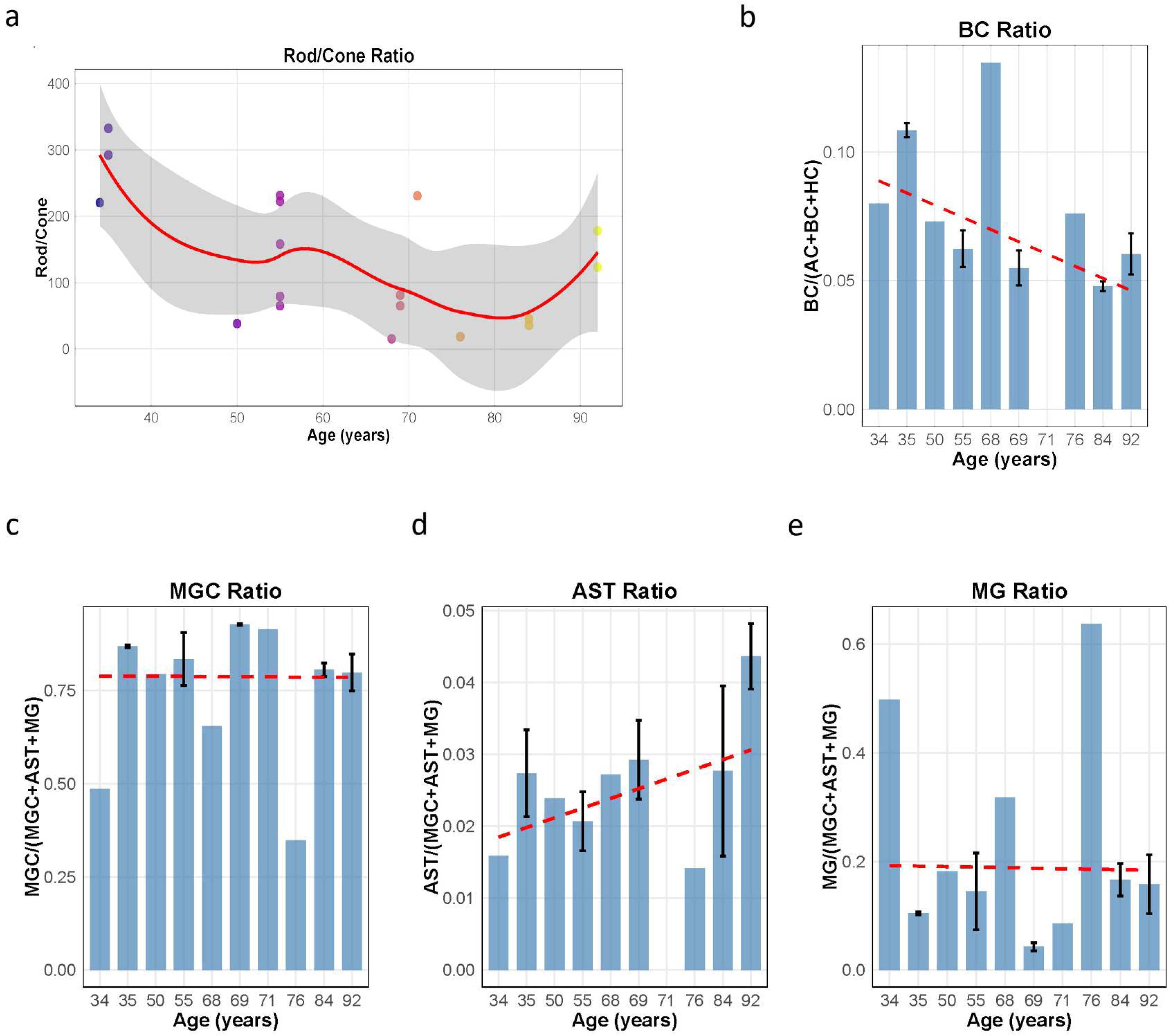
**(a)** Scatter plot showing the age-dependent decline in rod-to-cone photoreceptor ratio across individual samples. Each point represents an individual sample. **(b)** Age-related changes in bipolar cell proportion relative to total interneurons (BC /(AC+BC+HC)), showing a downward trend with advancing age. **(c)** Age-related changes in Müller glial cells proportion relative to total glial cell populations (astrocytes, microglia, and Müller glial cells). **(d)** Age-related changes in astrocytes proportion relative to total glial cell populations. **(e)** Age-related changes in microglia proportion relative to total glial cell population. AC: amacrine cell; BC: bipolar cell; HC: horizontal cell; MGC: Müller glial cell; RGC: retinal ganglion cell; RPE: retinal pigment epithelium; TC: T cell; MG: Microglia; AST: Astrocyte

Further analysis of identified BC subtypes (Figs. 3a, 3b) revealed distinct age-related patterns (Fig. 3c) with summary shown in Supplementary Table S3. Rod BCs (RB) consistently represented the largest proportion among BC subtypes (median 26%), with an outlier at age 50 (68.85%). Among diffuse BCs (DB), DB2 and DB3b were the most abundant, with DB2 showing higher proportions in middle-aged samples (∼12.88% at age 69) and lower values in older samples (∼6.57% at age 92). DB4a exhibited a gradual decline with age (14.47% at age 34 to ∼8.97% at age 69), while DB4b remained consistently below 2%. Midget BCs showed divergent patterns: Invaginating midget BCs (proportions remained relatively stable across ages (∼10-16%), while flat midget BCs showed greater variability, peaking at 15.79% at age 55 and declining to ∼5% in older samples.

**Figure 3.**
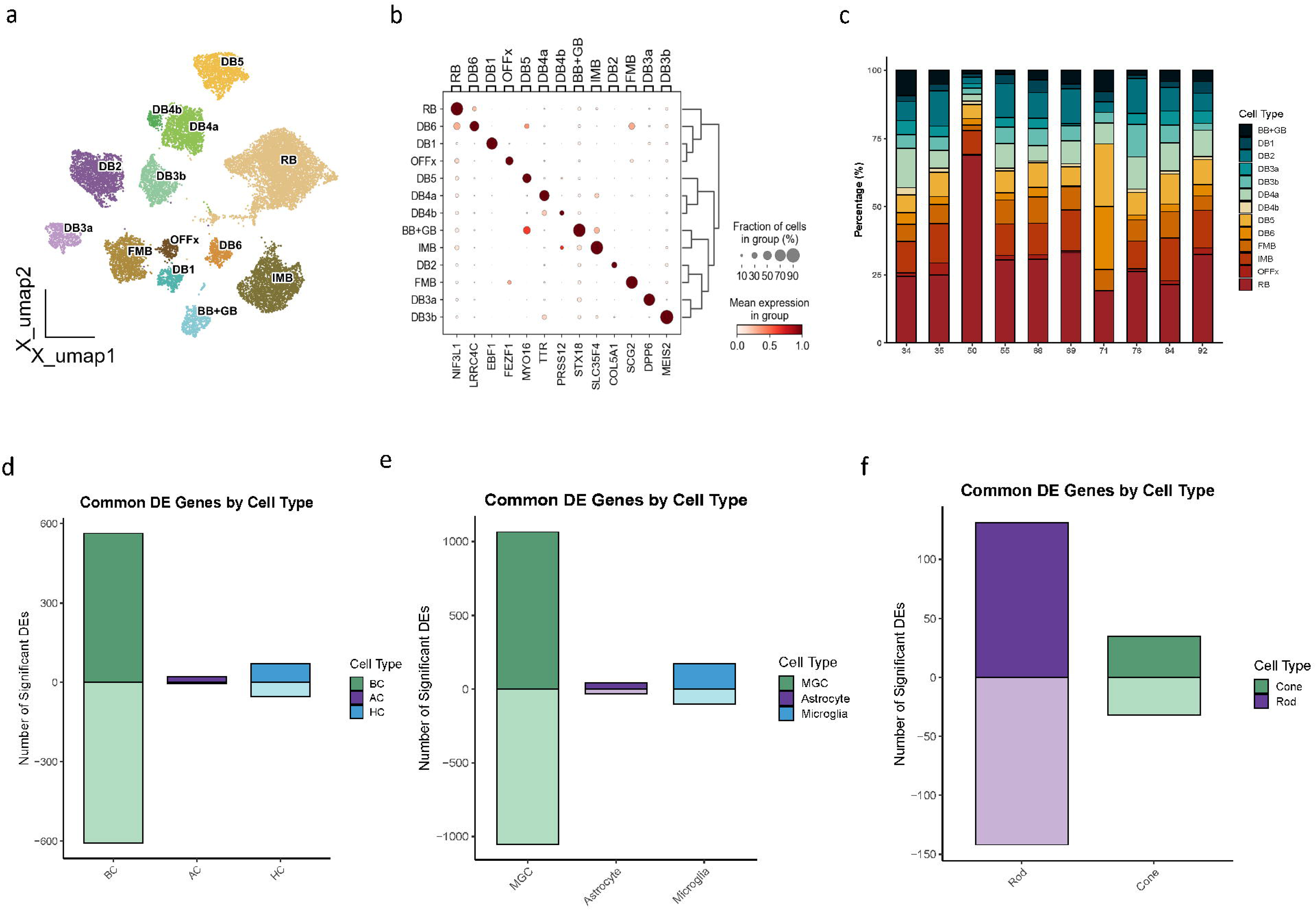
**(a)** UMAP visualization of bipolar cell subtypes. Distinct transcriptional clusters correspond to major bipolar cell subtypes, including rod bipolar cells (RB), ON diffuse bipolar cells (DB1–DB3a), OFF diffuse bipolar cells (DB3b– DB4), and midget bipolar cells—invaginating (IMB) and flat (FMB) types. **(b)** Dot plot showing marker genes’ expression for bipolar cell subtypes. Dot size represents the percentage of cells expressing each marker gene, while color intensity indicates the average expression level. **(c)** Stacked bar chart depicting the relative proportions of bipolar cell subtypes across samples arranged by donor age. **(d, e, f)** Bar plot quantifying age-associated differentially expressed genes (DEGs) across retinal interneuron populations for (d) interneurons, (e) glia and (f) photoreceptors identified by integrated single-cell (SC-DE) and pseudobulk (PB-DE) analyses. Upward bars (positive values) represent genes significantly upregulated in older retinas, while downward bars (negative values) indicate genes significantly downregulated with age.

### Comprehensive Profiling of Aging Retina Uncovers Cell□Type□Specific DEGs and Shared Hallmarks

Having characterized the age-related shifts in retinal cellular composition, we next aimed to delineate the underlying molecular alterations by performing a comprehensive transcriptional analysis across major retinal cell types. Transcriptional profiling revealed cell type-specific differential gene expression patterns across retinal cells with aging (Supplementary Table S4-S6). As a conservative benchmark, we also summarized expression at the donor level with covariate adjustment for sex and disease status, yielding fewer per cell-type DEGs (Supplementary Table S7). BCs exhibited the most substantial transcriptional changes among interneurons with 1,171 DEGs (563 upregulated, 608 downregulated), compared to amacrine cells (ACs) (26 DEGs) and horizontal cells (HCs) (124 DEGs) (Fig. 3d). Among these, 96.7% of BC DEGs were cell type-specific, compared to 53.8% in ACs and 71.8% in HCs, with 278 high-confidence genes detected by both analysis methods. Pathway analysis identified shared enrichment in hypoxia-related pathways across all aging interneurons, with glycolysis upregulated in aging AC and HC (Fig. 4a). Gene symbols and their corresponding full standardized names are provided in Supplementary Table S8. BCs demonstrated pronounced inflammatory activation (*CCL2*, *HLA-DRA*), gliosis (*GFAP*), and metabolic shifts toward glycolysis (*PFKFB4*, *PDK1*), along with extensive mitochondrial gene downregulation. ACs showed upregulation of angiogenic signaling (*VEGFA*) and mitochondrial quality control genes (*BNIP3*), while HCs exhibited unique enrichment in TGF-beta signaling and stress response genes (*EGR1*, *TMSB10*). Despite cell type-specific responses, conserved aging hallmarks emerged across all interneurons: mitochondrial dysfunction, immune activation, and synaptic weakening.

**Figure 4.**
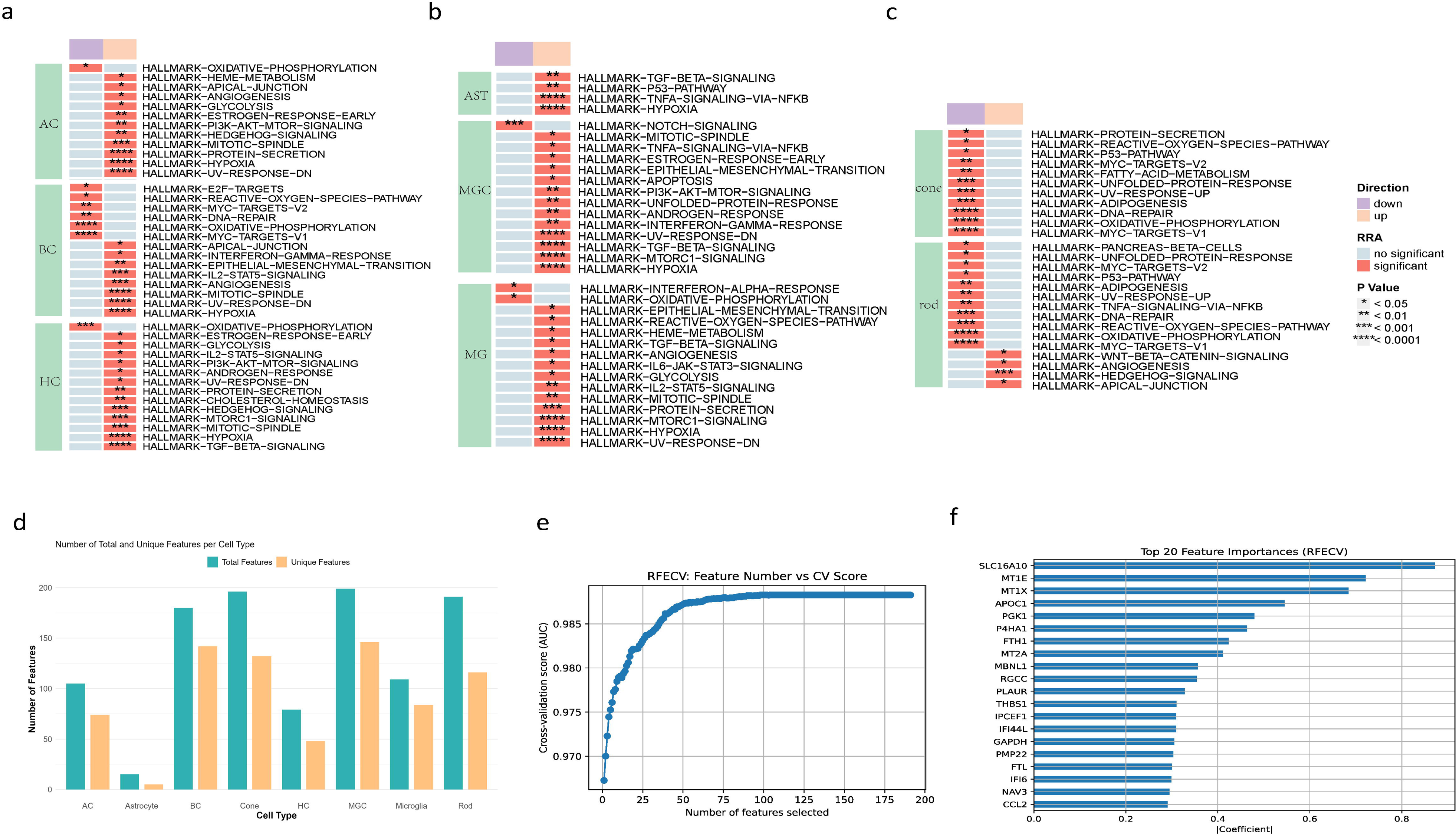
**(a, b, c)** Heatmap showing enriched pathways across retinal cell types of (a) interneurons, (b) glia and (c) photoreceptors based on irGSEA analysis. Color intensity indicates robust rank aggregation score, with red representing upregulation in old group and blue representing downregulation in old group. Significance levels: *P < 0.05, **P < 0.01, ***P < 0.001, ****P < 0.0001. **(d)** Bar plot showing the number of machine learning-identified discriminatory features across different retinal cell types. Green bars represent the total number of ML features, while yellow bars represent cell-type-specific features unique to each cell type. **(e)** Recursive feature elimination with cross-validation (RFECV) plot showing the relationship between feature number and cross-validation accuracy scores for the aging classifier of microglia. **(f)** Horizontal bar chart displaying the top 20 feature importance scores for microglia age classification, ranked by their contribution to distinguishing young from old groups. Features represent genes with the highest discriminatory power for microglia aging.

Among glial cells, MGCs exhibited the most pronounced alterations (2,117 DEGs), followed by microglia (276 DEGs) and astrocytes (74 DEGs) (Fig. 3e). Cell type specificity was highest in MGCs (94.2%), compared to microglia (68.5%) and astrocytes (43.2%). Core genes upregulated across all glial types included *FTH1* (iron sequestration), *LDHA* (glycolysis), and *SLC3A2* (amino acid transport), while *MT-ND6* (mitochondrial function) was consistently downregulated. Pathway analysis confirmed shared enrichment in Hypoxia, TNFA signaling via NF-kB, mTORC1 signaling, and EMT pathways (Fig. 4b). Each glial subtype demonstrated distinctive transcriptional signatures: MGCs uniquely displayed extensive inflammatory signaling (*CXCL1/2*) and matrix remodeling (*SERPINE1, LOX*); astrocytes showed induction of stress-response factors (*DNAJB1, DDIT3*) with concurrent downregulation of neuronal-supportive genes (*GRIA1, AQP4*). Comparative analysis with retinal interneurons revealed shared aging mechanisms but distinct effector pathways: glial cells prominently involved antigen-presentation, whereas interneurons primarily expressed chemokines.

Photoreceptors exhibited subtype-specific aging responses, with rods showing substantially greater transcriptional changes (273 DEGs) than cones (67 DEGs) (Fig. 3f). Cell type-specificity analysis revealed that 91.9% of rod DEGs were unique to rods versus 67.2% for cones. Both photoreceptor types shared upregulation of stress response genes (*CRYAB, MT2A*) and chaperones (*BAG3*), while commonly downregulating mitochondrial genes (*MT-ND6*) and cytoskeletal components (*KIF2A*). Integrated pathway analysis confirmed enrichment of oxidative stress response and mitochondrial dysfunction pathways in both cell types (Fig. 4c). Rods uniquely exhibited downregulation of TNFA-signaling-via-NFKB pathways alongside significant upregulation of WNT-β-catenin signaling, supported by increased expression of *WNT5B*, *WNT11* and *FZD3*. Rod-specific DEGs included upregulated stress response genes (*S100A6*, *NLN*) and downregulated phototransduction-related genes (*ELOVL4*, *CRB1*). Cones selectively downregulated fatty acid metabolism and protein secretion pathways, with substantial decreases in membrane transport components. Compared to glial cells, photoreceptors exhibited similar response patterns but with lower magnitude of transcriptional changes, underscoring their specialized functional vulnerability during aging.

### Machine Learning Identifies Robust Transcriptional Features of Retinal Aging

In the previous section, we performed DE and enrichment analysis to characterize aging-related changes within major retinal cell types. To further identify the robust transcriptional discriminators of cellular age, we applied a ML framework using a L1□regularized logistic regression with cross-validation. This approach ranked genes based on their importance in classification, providing a prioritized list of robust age-defining features (Supplementary Table S9). Complete model weights were provided in Supplementary Table S10 and bootstrap selection frequencies across all 20 replicates for each cell type were detailed in Supplementary Table S11, demonstrating the stability of selected features. Compared to traditional feature analysis approaches, the ML model excelled at identifying subtle yet highly consistent changes indicative of aging, including features with modest fold changes that might be overlooked in standard statistical tests, particularly in regulatory elements and less abundant transcripts that nevertheless serve as powerful age discriminators. Our ML models achieved high classification accuracy across all cell types, ranging from 80% in astrocytes to 96% in microglia, with an average accuracy of 89%, demonstrating strong classification power of the identified features (Table 2). The number of ML-identified discriminatory features varied considerably between cell types, with patterns that partially mirrored our DEG analysis but revealed important distinctions. Astrocytes showed the fewest ML features (15), consistent with their limited DEGs (74), while MGCs had the most (199), aligning with their extensive DEG count (2,117). Bipolar cells (180), rods (191), and cones (196) all exhibited substantial ML features, indicating strong age-related transcriptional signatures (Fig. 4d). Notably, cones presented a striking divergence between metrics-few DEGs (67) but abundant ML features (196), and this pattern was also observed in ACs (105 ML features versus 26 DEGs).

**Table 2.** Performance Metrics of Cell Age Classification Models Across Retinal Cell Types.

| Cell Type‡ | Accuracy* | F1-Score† | Precision† | Recall† | AUC |
| --- | --- | --- | --- | --- | --- |
| Microglia | 0.96 | 0.95 | 0.95 | 0.95 | 0.98 |
| HC | 0.93 | 0.93 | 0.93 | 0.93 | 0.97 |
| BC | 0.91 | 0.91 | 0.91 | 0.90 | 0.96 |
| Cone | 0.90 | 0.88 | 0.89 | 0.87 | 0.94 |
| Rod | 0.89 | 0.89 | 0.89 | 0.89 | 0.95 |
| MGC | 0.89 | 0.89 | 0.89 | 0.89 | 0.95 |
| AC | 0.85 | 0.83 | 0.84 | 0.82 | 0.93 |
| Astrocyte | 0.80 | 0.79 | 0.79 | 0.79 | 0.87 |

\*Accuracy represents the overall proportion of correctly classified cells. †F1-score, precision, and recall are reported as macro-averages across young and old classes. ‡ HC: horizontal cell; BC: bipolar cell; MGC: Müller glial cell; AC: amacrine cell.

The ML analysis consistently pinpointed genes central to cell-type-specific functions and critical homeostatic pathways as top age classifiers. Firstly, analysis of transcriptional features recurring in three or more retinal cell types, complement and reinforce the shared patterns of aging that discriminate between young and old groups (Supplementary Table S12). Mitochondrial dysfunction emerges as a dominant theme, with *MT-ND6* showing the highest frequency. Proteostasis decline is evident through frequent chaperones such as *CRYAB* and *HSPA1A*, and the ubiquitin system via *UBC*, indicating pan-retinal stress responses to protein misfolding. Oxidative stress responses are widespread, marked by *GPX3* and metallothioneins like *MT2A*, while low-grade inflammation is supported by immune genes including *HLA-A* and *S100A10*. Structural integrity is also affected, with cytoskeletal changes via *TUBA1B*, synaptic reorganization via *MAGI2*, and extracellular matrix remodeling via *P4HA1*. Beyond these established hallmarks, novel players include transcriptional regulators *BHLHE40*, linked to hypoxia and circadian disruption, and *ERO1A*. Additionally, non-coding RNAs like *FTX* and *SNHG14* point to epigenetic regulation, while *NCL* (nucleolin) hints at nucleolar stress, and *TF* (transferrin) indicates disrupted iron homeostasis. These findings reinforce classic aging themes while highlighting novel regulatory and metabolic vulnerabilities in the aging retina.

### Cell-Type Specific ML Features Reveal Distinct Aging Vulnerabilities

Our ML model of microglia demonstrated a high degree of accuracy (96%) in distinguishing between young and old cells (Table 2). This high performance, compared to other retinal cell types (which averaged 89% accuracy). Our recursive feature elimination with cross-validation approach identified the optimal feature set for classification (Fig. 4e), ensuring model performance while minimizing overfitting. Inspection of the top 20 importance ranks (Fig. 4f) reveals the dominant molecular programmes of Metal–redox buffering, accompanied by Inflammation-linked matrix remodeling. A smaller third set reflects metabolic and structural rewiring. Metallothioneins (*MT1E*, *MT1X*, *MT2A*) and ferritin subunits (*FTH1*, *FTL*) occupied five positions among the top-ranked features, revealing a concerted response to metal dysregulation as a defining hallmark of aged microglia. This metal-redox buffering signature has been previously noted in photoreceptors but is now specifically pinpointed as a key distinguishing feature in aged microglia cells, potentially indicating a tractable pathway for modulating glial contributions to retinal senescence.

For astrocytes, classification accuracy was more modest at 80%, reflecting a focused 15-gene signature related to acute-phase and protease regulation (Fig. 5a). Haptoglobin (*HP*) and serine-protease inhibitor *SLPI* were among the top-ranked features.

**Figure 5.**
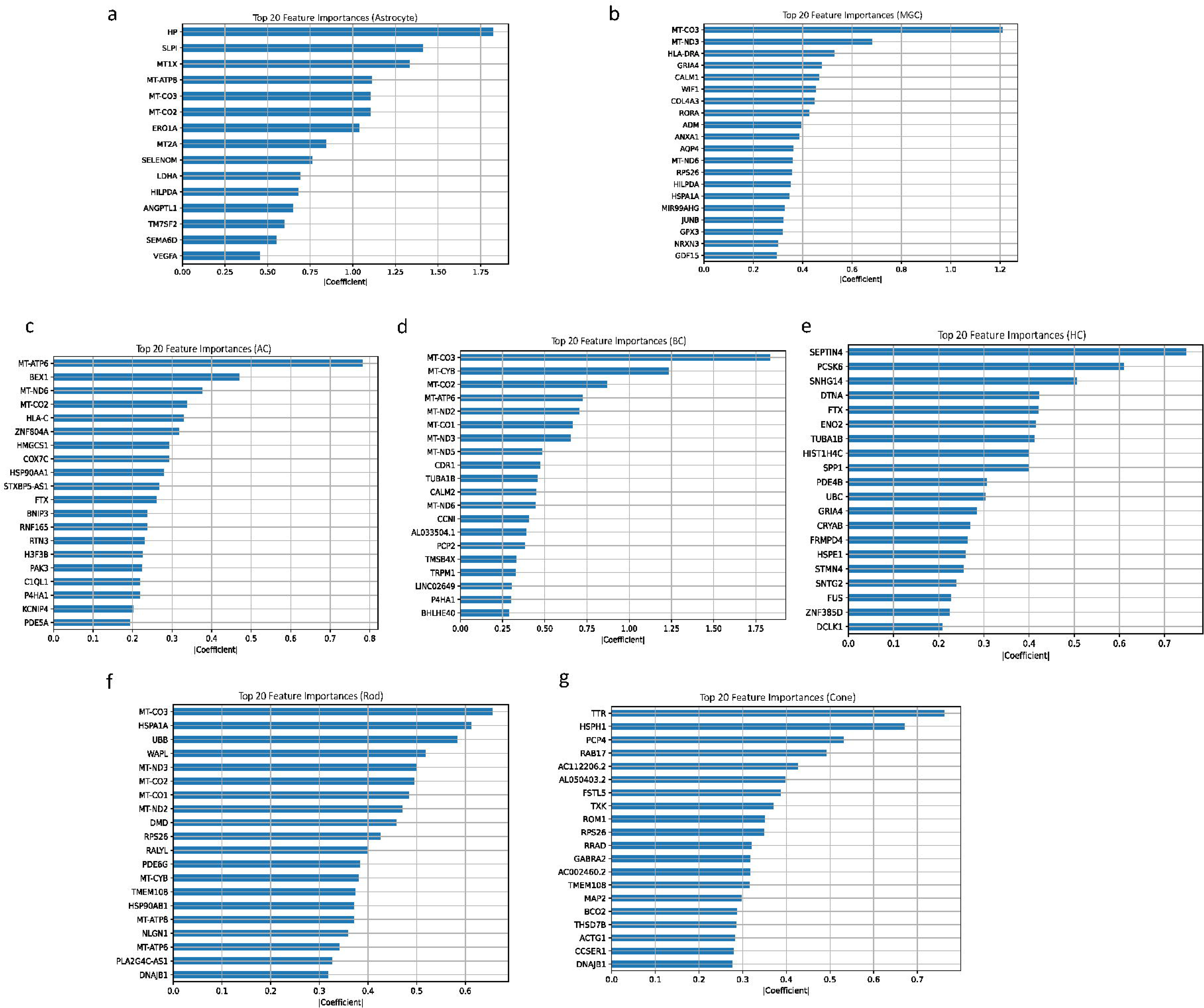
Horizontal bar charts showing the top 20 feature importance scores for age classification across different retinal cell types: (a) astrocytes, (b) Müller glial cells (MGCs), (c) amacrine cells (ACs), (d) bipolar cells (BCs), (e) horizontal cells (HCs), (f) rod photoreceptors, and (g) cone photoreceptors.

The MGC classifier achieved 89% accuracy in five-fold cross-validation (Fig. 5b). Analysis of the highest-weight features reveals three interconnected aging programs: mitochondrial distress (evidenced by *MT-CO3*, *MT-ND6*, and *MT-ND3*), matrix remodeling (featuring *COL4A3*), and a distinctive Ca²□/Wnt-lipid signaling network. This third program combines Ca²□ sensor *CALM1*, Wnt pathway inhibitor *WIF1*, and lipid/hypoxia response genes (*RORA*, *HILPDA*, and *ADM*). Our BC classifier performed robustly at 91% accuracy, while AC reached 85% accuracy and HC achieved 93% accuracy (Fig. 5c-e). In interneuron ACs and BCs, mitochondrial distress dominates-notably, eight of BC’s top ten features are mitochondrial transcripts (*MT-CO3*, *MT-CYB, MT-CO2, MT-ATP6, MT-ND2, MT-CO1*, *MT-ND3, MT-ND5*), pointing to respiratory chain impairment as the principal discriminator of BC aging. ACs similarly prioritize mitochondrial genes (*MT-ATP6, MT-ND6, MT-CO2*) but additionally elevate the mitophagy trigger *BNIP3*. While MGCs display only modest mitochondrial signal, their Ca²□ machinery integrates with tissue-modulatory networks. Each cell type shows distinct proteostasis-lipid-immune signatures: ACs feature chaperone *HSP90AA1*, ER-reticulon *RTN3*, lipid enzyme *HMGCS1*, and *MHC-I HLA-C*; BCs include circadian/hypoxia regulator *BHLHE40*; and MGCs highlight matrix-gliosis gene *COL4A3* and vasopeptide *ADM*. HCs break the pattern entirely-their top features are cytoskeletal and synaptic scaffolds (*SEPTIN4, TUBA1B, STMN4, DCLK1*), alongside a unique RNA-chaperone module (*SNHG14, FTX, CRYAB, HSPE1, FUS*) and signaling modulators (*PDE4B, GRIA4*). This analysis reveals that aging manifests as energy failure in AC/BC, metabolic-paracrine rewiring in MGC, and structural-RNA maintenance in HC.

Rods (89% accuracy, AUC = 0.95) are dominated by a dual axis of mitochondrial respiratory-chain failure-six MT-encoded OXPHOS transcripts (e.g., *MT-CO3, MT-ND3, MT-CO2, MT-CO1, MT-ND2, MT-CYB*) together with a broad heat-shock/ubiquitin response (*HSPA1A, UBB, HSP90AB1, DNAJB1*), indicating that aging rods depend on sustained ATP supply and intensive protein-quality control (Fig. 5f). Cones deliver comparable performance (90% accuracy, AUC = 0.96) yet place no mitochondrial genes within their top 20 features (Fig. 5g); instead, they emphasise chaperone-ribosome stress (*HSPH1, DNAJB1, RPS26*) and outer-segment membrane/retinoid maintenance (*TTR, ROM1, BCO2, THSD7B*). Thus, aging photoreceptors converge on proteostasis yet diverge into mitochondria-centric (rods) versus membrane-metabolite (cones) adaptations. Unlike inner neurons, they couple protein surveillance directly to phototransduction architecture, highlighting distinct therapeutic entry points for preserving scotopic versus photopic vision in the aging retina.

### Regulatory Network Alterations with Aging Human Retina

SCENIC analysis revealed both shared and cell type-specific TF regulatory patterns in aging retinal interneurons (Fig. S1a; see Supplementary Table S13). Across all 3 interneuron types, stress-responsive TFs *BHLHE40* and *MXI1* showed consistently increased regulon activity, indicating a conserved hypoxia-driven metabolic stress response. Notably, *BHLHE40* was also identified as one of the top aging features by BCs ML model, underscoring its importance in interneuron aging. In aging interneurons, *CHD1* regulon activity increased, whereas *NEUROD1* and ATF-family regulon activities decreased. Each interneuron type displayed distinctive regulatory signatures. ACs exhibited unique *PROX1* regulon activation (reduced in BCs) alongside elevated *NFIA* and *TCF4* regulons. BCs showed pronounced inflammatory signatures through increased *NFKB1* regulon activity, aligning with upregulated inflammatory genes (*CCL2*, *HLA-DRA*) from our differential expression analysis. HCs displayed distinctive *IRF1* regulon activation and compromised identity maintenance through substantial decreases in *ONECUT1* activity.

Gene set scoring analysis revealed significant decreases in polyamine metabolism pathway activity in aged cells across all interneuron types (Fig. S1b). Conversely, cellular senescence gene set (SenMayo) activity increased significantly in all three cell types (Fig. S1c).

SCENIC analysis revealed conserved activation of stress-responsive TFs across all glial cell types during aging (Fig. S2a; see Supplementary Table S14). *XBP1*, *JUND*, *FOSL2*, *MAFF*, *BHLHE40*, and *BHLHE41* showed increased regulon activity in astrocytes (obvious elevation), microglia (moderate elevation), and MGC (variable elevation), highlighting conserved regulatory mechanisms related to cellular stress, inflammation, and metabolic reprogramming. Each glial subtype exhibited distinctive regulatory signatures. MGC uniquely showed significant *ZXDA*(-) regulon activity reduction, suggesting altered chromatin remodeling. Microglia specifically displayed increased regulon activity of hypoxia-responsive (*MITF*, *MXI1*) and metabolic TFs (*FOSL1*), alongside reduced regulon activity of immune-related regulons (*RFXANK*). Astrocytes demonstrated substantial induction of stress-response (*CEBPB*, *HOXA5*) and lipid metabolism (*SREBF1*) regulons. While both interneurons and glia showed increased stress-responsive TF regulon activity with age, glial cells exhibited more pronounced differential activity, particularly with strong *ZBTB21* regulon activation in aged MGCs and astrocytes, and consistent *MAFF* elevation across all glial cells. In addition to these changes, TF analysis revealed distinct regulatory profiles in rods and cones, detailed in Supplementary Table S15. Gene set scoring analysis revealed differential metabolic resilience among glial subtypes. Polyamine metabolism gene set activity decreased significantly only in aging MGC, while microglia and astrocytes maintained stable activity (Fig. S2b). The SenMayo cellular senescence gene set showed robust increased activity across all glial types, particularly in astrocytes and MGC, exceeding the magnitude observed in interneurons and suggesting a more prominent role for glial senescence in retinal aging (Fig. S2c).

## Discussion

Although scRNA-seq transcriptomic studies have explored retinal aging at various biological levels, compared to AMD, the physiological aging of human retina has received less attention, despite being the foundation for late-onset retinal diseases. Current single-cell retinal aging studies have been constrained by limited sample sizes, incomplete age representation across the adult lifespan, and predominant focus on cohorts of European ancestry. Moreover, while ML models have been widely adopted for feature selection and predictive analysis in various biological domains ^31–34^, their application in human retinal aging transcriptomic research remains limited. Most previous studies on human retinal aging have predominantly utilized conventional statistical approaches to identify age-associated transcriptional changes^9, 10, 35^. Here, we generated scRNA-seq profiles from 18 unfrozen retinas of a genetically homogeneous Chinese cohort spanning 34–92 years. By coupling conventional differential-expression analysis with a cross-validated machine-learning framework, we delineate conserved molecular hallmarks and expose sharp cell-type-specific vulnerabilities that refine the molecular narrative of human retinal aging.

A key finding of our study was the progressive age-related reduction in the rod-to-cone ratio, suggesting selective rod vulnerability. This aligns with classic histological and clinical observations showing substantial rod loss by late adulthood ^36, 37^. Rods inherently possess higher metabolic demands due to extensive outer segment renewal and their reliance on aerobic glycolysis, making them particularly susceptible to metabolic stress, mitochondrial dysfunction, and oxidative damage ^38, 39^. Our transcriptomic analysis provided direct molecular evidence supporting this hypothesis, demonstrating substantial dysregulation of mitochondrial genes and oxidative phosphorylation pathways specifically in rod. Quantitatively, rods exhibited significantly more mitochondrial-related DEGs compared to cones. Additionally, marked inflammatory activation and upregulation of stress-response genes further highlight rods’ pronounced susceptibility to aging stressors ^8^. In contrast, cones demonstrate fewer transcriptional alterations, consistent with their known slower degenerative kinetics and greater resilience ^40, 41^. Interestingly, cones selectively upregulated certain immune-related genes, possibly reflecting adaptive responses rather than direct degeneration. Based on their relatively stable transcriptomic profiles, cones may adopt distinct adaptive responses to mitigate the impact of aging-induced cellular stress.

Among retinal interneurons, BCs exhibited the most significant age-related transcriptional changes, prominently characterized by inflammatory activation and pronounced mitochondrial dysfunction. Animal studies have shown BC dendritic remodeling and altered synaptic ribbon architecture during aging ^42^, suggesting a heightened sensitivity of BCs to aging stress. HCs and ACs showed comparatively fewer transcriptional changes. Despite also exhibiting hallmarks of oxidative stress and synaptic remodeling, these interneurons may maintain greater intrinsic stability or resilience.

We also highlighted significant glial involvement in retinal aging, particularly MGCs, which displayed profound inflammatory responses and extracellular matrix remodeling. These findings support prior evidence that reactive gliosis in MGCs, initially protective, can exacerbate inflammation and vascular dysregulation upon chronic activation ^43, 44^. Astrocytes, although fewer in number, similarly exhibited stress-related activation. Enhanced astrocytic stress responses could reshape local inflammatory and metabolic environments, influencing neuronal survival and retinal homeostasis.

In contrast to traditional retinal aging studies that rely mainly on DE analysis to identify age-associated genes^45^, ML approaches provide alternative methodologies that can extract informative signatures from noisy, high-dimensional data and alleviate certain statistical model assumptions^46^. While DE methods excel at detecting large-magnitude transcriptional changes, they frequently overlook subtle yet consistent alterations in regulatory elements and low-abundance transcripts that may be functionally significant. Traditional analyses rank genes by statistical significance, while machine learning provides feature importance rankings based on predictive power and biological relevance. Furthermore, while general aging gene sets like SenMayo provide valuable insights into conserved senescence pathways ^30^, they may not fully capture the cell-type-specific nuances or the broader spectrum of non-senescence-related aging within a complex, specialized tissue such as the human retina. Our ML-ranking approach addresses this gap by identifying transcripts with the highest classification power, highlighting features that maintain consistent age-associated patterns even when their absolute expression changes fall below statistical thresholds for differential expression. This complementary perspective helps bridge the gap between statistical significance and biological relevance in understanding retinal aging. These include metallothioneins, mitochondrial genes, proteostasis regulators, interferon-stimulated genes, chaperones, and long non-coding RNAs, many of which were not among the top DEGs but still emerged as robust indicators of aging. Importantly, the ML features also delineated cell type–resolved patterns: BCs, ACs, and rods shared a pronounced mitochondria-centric axis, whereas cones showed a retinoid metabolism, and proteostasis-focused profile. MGCs exhibited a coupled interferon-response and collagen signature, while HCs were marked by cytoskeletal and RNA-maintenance changes. To systematically connect our molecular findings with their functional implications, we have synthesized the cell-type-specific aging signatures, their associated hallmarks, potential retinal consequences, and translational applications in a comprehensive summary (Table 3). By capturing these multifaceted and cell-specific signals beyond conventional DE analysis, the ML-driven approach provides a more precise, systems-level understanding of retinal aging and could inform more targeted therapeutic strategies. While biologically plausible, these links are hypothesis-generating given our cross-sectional, transcriptomic design and will require future functional validation.

**Table 3.**
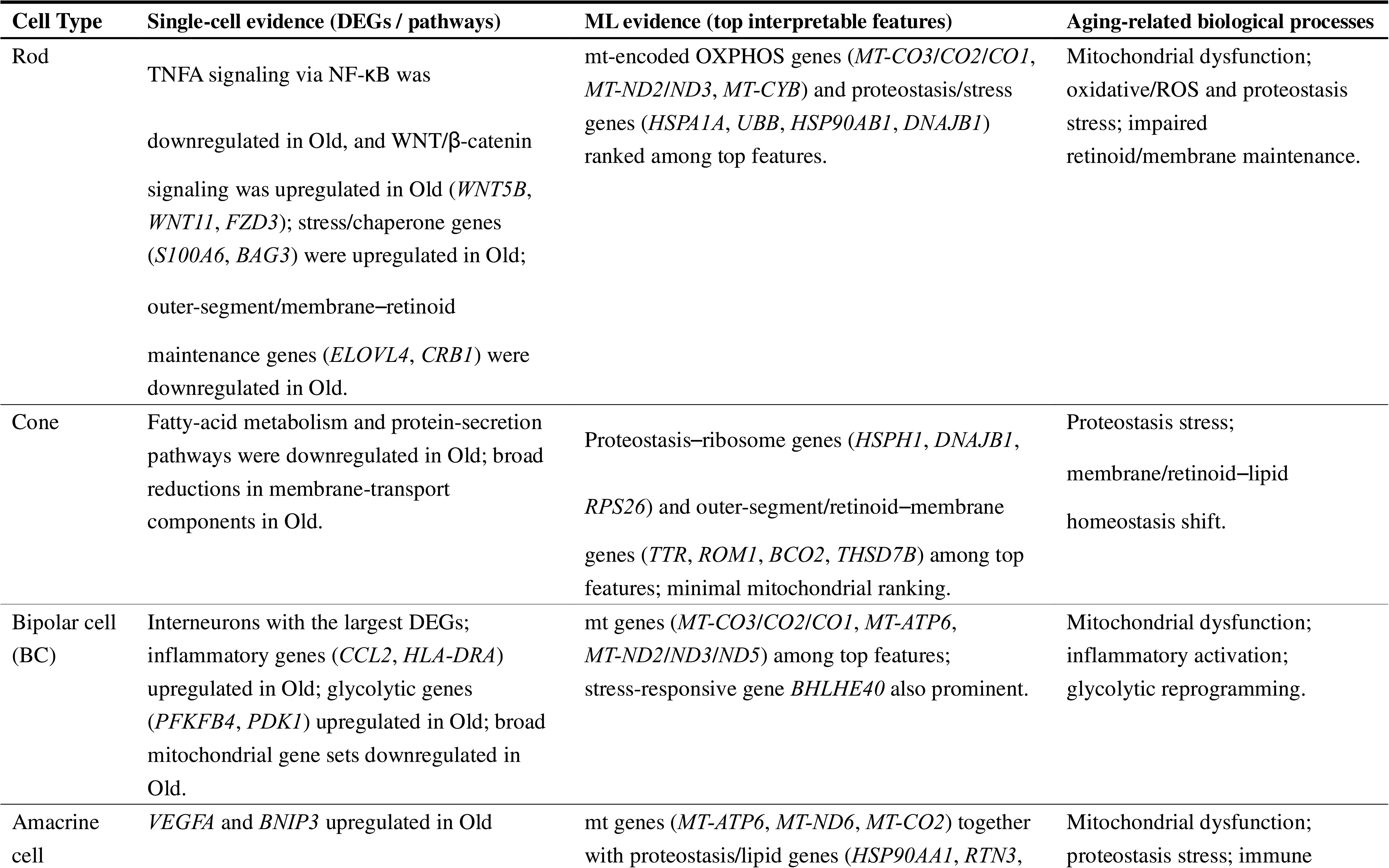
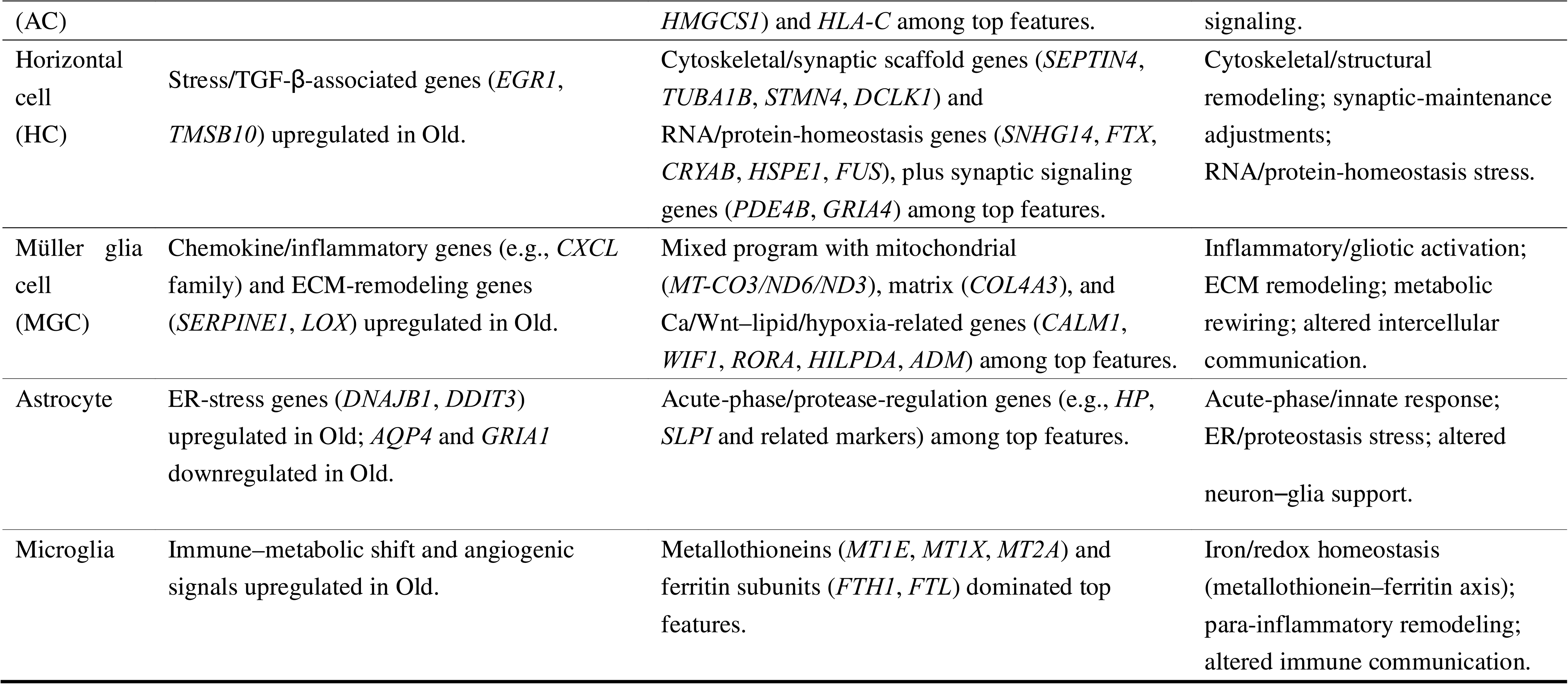
Cell-type aging signatures mapped to aging-related biological processes, integrating single-cell and ML evidence.

To address potential ethnic diversity limitations, we compared our Han Chinese cohort with the predominantly European ancestry human retina cell atlas (HRCA) age model (135 samples, 57 donors, ages 10-91)^13^. The atlas reported age-associated activation of complement/coagulation, adaptive immunity, steroid biosynthesis, and calcium import, alongside suppression of ribosomal translation and mitochondrial gene expression. Our findings largely align: mitochondrial down-regulation in rods, bipolar, and amacrine cells, with immune activation in BCs and MGCs, supporting cross-cohort robustness of core aging hallmarks. However, some signals were more prominent in our cohort, rod WNT/β-catenin upregulation, microglial metallothionein-ferritin programs, and cone retinoid/lipid shifts, potentially reflecting population-specific differences alongside methodological factors. We interpret these differences cautiously as cohort- or methodology-dependent rather than definitive ancestry effects, given that HRCA modeled age continuously and analyzed scRNA-seq and single-nucleus RNA sequencing (snRNA-seq) separately, whereas we used discrete old-versus-young comparisons with scRNA-seq only. Taken together, the findings are compatible with conserved aging programs while indicating additional, possibly cohort-enriched signals, underscoring the utility of including diverse populations.

This study has several limitations. The relatively small sample size in certain age brackets and the low RPE cell yield restrict our ability to capture RPE-specific aging changes. Proteomic and functional assays will be necessary to confirm these findings. Moreover, the ML models, while rigorously cross-validated, were trained on the same dataset used for feature discovery. A key limitation of this study is the absence of external validation using independent human retinal scRNA-seq datasets. We attempted to apply our classifiers to the HRCA retinal data; however, differences in sequencing platforms, sample preparation workflows, and donor demographics resulted in incomplete overlap of critical marker genes, preventing reliable evaluation. Future efforts will focus on harmonizing datasets across studies or generating additional cohorts to enable rigorous external validation. Such work will facilitate assessment of cohort-specific variations in classifier performance and aging-associated transcriptional signatures, thereby enhancing the generalizability, interpretability, and translational relevance of our machine-learning classifiers. Particularly, we plan to leverage advanced computational methods, such as transfer learning and domain adaptation, to bridge technical differences across platforms and integrate diverse datasets into our ML framework. These approaches will also help disentangle ancestry-related signals from cohorts and methodological confounders, enabling a more nuanced understanding of conserved versus population-specific aging mechanisms in the human retina.

In summary, we systematically characterized early-age transcriptional landscapes associated with human retinal aging within an ethnically homogeneous Chinese cohort. Our approach combined differential-expression statistics with a sparsity-driven machine-learning pipeline to identify robust aging signatures. This study addresses gaps previously overlooked in existing literature, particularly the focus on older starting ages and predominantly European ancestry representation. By generating the first machine-learning-derived, cell-type-specific aging gene lists for the human retina, our findings provide insights into conserved molecular hallmarks alongside distinctive cell-type-specific vulnerabilities relevant for developing targeted therapeutic interventions aimed at mitigating age-related retinal degeneration processes across diverse populations.

## Statements and Declarations

## Consent to Publish

All authors have consent for publication.

## Authors’ Contributions

LY and SL drafted the paper and performed the analysis. YT, QP, TC contributed to data formatting and correction. YY, JL, YZ, YS, QY, ZL, LC, GM, and RR provided comments on the paper. FL, and WM organized the project and provided comments. Collaborators from Queen’s University Belfast and Universiti Brunei Darussalam provided academic guidance and editorial suggestions only; they did not participate in data analysis, sample handling, or access to raw data. All substantive research activities were completed within Mainland China.

## Data and Code availability

The single-cell RNA sequencing data generated in this study and custom code for the multi-stage machine learning pipeline have been deposited in Figshare with the DOI: 10.6084/m9.figshare.28757048 and GSA-Human (accession ID: HRA014708).

## Human subjects Ethics statement

Human subjects were included in this study. This study was conducted at the University of Nottingham Ningbo China, Lihuili Hospital affiliated with Ningbo University, and the Affiliated Ningbo Eye Hospital of Wenzhou Medical University. The Institutional Review Boards/Ethics Committees of the participating institutions reviewed and approved the study protocol. Informed consent was obtained from all participants or their legal representatives prior to tissue collection. All study procedures adhered to the tenets of the Declaration of Helsinki.

## Abbreviations and Acronyms

AC: amacrine cell
AMD: age-related macular degeneration
AO/PI: acridine orange/propidium iodide
AUC: area under the curve
BC: bipolar cell
BSA: bovine serum albumin
DB: diffuse bipolar cell
DEGs: differentially expressed genes
DM: diabetes mellitus
DR: diabetic retinopathy
ECM: extracellular matrix
FDR: false discovery rate
FMB: flat midget bipolar cell
GEMs: gel beads in emulsion
HC: horizontal cell
HRCA: human retina cell atlas
IMB: invaginating midget bipolar cell
logFC: log fold change
MGC: Müller glial cell
ML: machine learning
PB-DE: pseudobulk differential expression
RB: rod bipolar cell
RFECV: recursive feature elimination with cross-validation
RGC: retinal ganglion cell
ROC: receiver operating characteristic
RPE: retinal pigment epithelium
SC-DE: single-cell differential expression
scRNA-seq: single-cell RNA sequencing
snRNA-seq: single-nucleus RNA sequencing
TC: T cell
TF: transcription factor
UMI: unique molecular identifier;
UMAP: uniform manifold approximation and projection

