## Supplementary material for "Interpretable Aging Signatures in Human Retinal Cell Types Revealed by Single-Cell RNA Sequencing and Sparse Logistic Regression": Figure S1

a

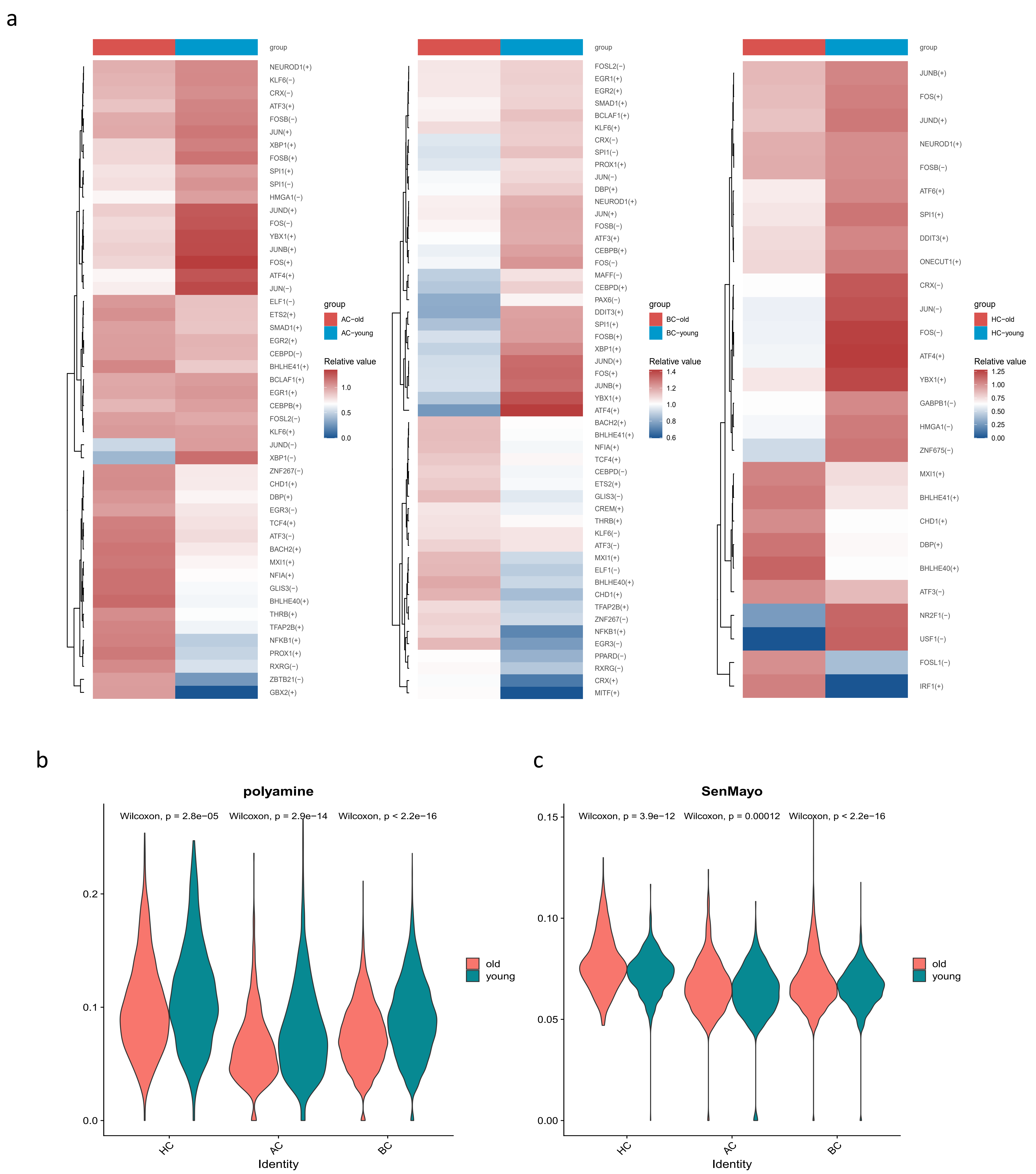


Figure S1. (a) Heatmap showing differential activity of transcription factor (TF) regulons between young and old groups across retinal interneuron types (amacrine cell, bipolar cell, and horizontal cell) identified by pySCENIC analysis. Each column represents an age group (old or young) within a specific cell type, while rows represent individual TF regulons. Red indicates increased regulon activity in old retinas compared to young, while blue indicates decreased activity. TFs are hierarchically clustered based on similarity in their activity patterns across cell types. (b) Box plots comparing polyamine metabolism pathway scores between young and old groups with P calculated by Wilcoxon rank-sum tests for each retinal interneuron. (c) Box plots comparing cellular senescence gene set (SenMayo) activity scores between young and old groups with P calculated by Wilcoxon rank-sum tests for each retinal interneuron.
