## Supplementary material for "Interpretable Aging Signatures in Human Retinal Cell Types Revealed by Single-Cell RNA Sequencing and Sparse Logistic Regression": Table S1

Table S1: Gene sets used for scoring analyses.

| GeneName | GeneSet |
|---|---|
| ACVR1B | SenMayo |
| ANG | SenMayo |
| ANGPT1 | SenMayo |
| ANGPTL4 | SenMayo |
| AREG | SenMayo |
| AXL | SenMayo |
| BEX3 | SenMayo |
| BMP2 | SenMayo |
| BMP6 | SenMayo |
| C3 | SenMayo |
| CCL1 | SenMayo |
| CCL13 | SenMayo |
| CCL16 | SenMayo |
| CCL2 | SenMayo |
| CCL20 | SenMayo |
| CCL24 | SenMayo |
| CCL26 | SenMayo |
| CCL3 | SenMayo |
| CCL3L1 | SenMayo |
| CCL4 | SenMayo |
| CCL5 | SenMayo |
| CCL7 | SenMayo |
| CCL8 | SenMayo |
| CD55 | SenMayo |
| CD9 | SenMayo |
| CSF1 | SenMayo |
| CSF2 | SenMayo |
| CSF2RB | SenMayo |
| CST4 | SenMayo |
| CTNNB1 | SenMayo |
| CTSB | SenMayo |
| CXCL1 | SenMayo |
| CXCL10 | SenMayo |
| CXCL12 | SenMayo |
| CXCL16 | SenMayo |
| CXCL2 | SenMayo |
| CXCL3 | SenMayo |
| CXCL8 | SenMayo |
| CXCR2 | SenMayo |
| DKK1 | SenMayo |
| EDN1 | SenMayo |
| EGF | SenMayo |
| EGFR | SenMayo |
| EREG | SenMayo |
| ESM1 | SenMayo |
| ETS2 | SenMayo |
| FAS | SenMayo |
| FGF1 | SenMayo |
| FGF2 | SenMayo |

| | |
|---|---|
| FGF7 | SenMayo |
| GDF15 | SenMayo |
| GEM | SenMayo |
| GMFG | SenMayo |
| HGF | SenMayo |
| HMGB1 | SenMayo |
| ICAM1 | SenMayo |
| ICAM3 | SenMayo |
| IGF1 | SenMayo |
| IGFBP1 | SenMayo |
| IGFBP2 | SenMayo |
| IGFBP3 | SenMayo |
| IGFBP4 | SenMayo |
| IGFBP5 | SenMayo |
| IGFBP6 | SenMayo |
| IGFBP7 | SenMayo |
| IL10 | SenMayo |
| IL13 | SenMayo |
| IL15 | SenMayo |
| IL18 | SenMayo |
| IL1A | SenMayo |
| IL1B | SenMayo |
| IL2 | SenMayo |
| IL32 | SenMayo |
| IL6 | SenMayo |
| IL6ST | SenMayo |
| IL7 | SenMayo |
| INHA | SenMayo |
| IQGAP2 | SenMayo |
| ITGA2 | SenMayo |
| ITPKA | SenMayo |
| JUN | SenMayo |
| KITLG | SenMayo |
| LCP1 | SenMayo |
| MIF | SenMayo |
| MMP1 | SenMayo |
| MMP10 | SenMayo |
| MMP12 | SenMayo |
| MMP13 | SenMayo |
| MMP14 | SenMayo |
| MMP2 | SenMayo |
| MMP3 | SenMayo |
| MMP9 | SenMayo |
| NAP1L4 | SenMayo |
| NRG1 | SenMayo |
| PAPPA | SenMayo |
| PECAM1 | SenMayo |
| PGF | SenMayo |
| PIGF | SenMayo |
| PLAT | SenMayo |

| | |
|---|---|
| PLAU | SenMayo |
| PLAUR | SenMayo |
| PTBP1 | SenMayo |
| PTGER2 | SenMayo |
| PTGES | SenMayo |
| RPS6KA5 | SenMayo |
| SCAMP4 | SenMayo |
| SELPLG | SenMayo |
| SEMA3F | SenMayo |
| SERPINB4 | SenMayo |
| SERPINE1 | SenMayo |
| SERPINE2 | SenMayo |
| SPP1 | SenMayo |
| SPX | SenMayo |
| TIMP2 | SenMayo |
| TNF | SenMayo |
| TNFRSF10C | SenMayo |
| TNFRSF11B | SenMayo |
| TNFRSF1A | SenMayo |
| TNFRSF1B | SenMayo |
| TUBGCP2 | SenMayo |
| VEGFA | SenMayo |
| VEGFC | SenMayo |
| VGF | SenMayo |
| WNT16 | SenMayo |
| WNT2 | SenMayo |
| PSMB1 | polyamine |
| PSMC4 | polyamine |
| PSMA4 | polyamine |
| PSME4 | polyamine |
| PSMC5 | polyamine |
| SMOX | polyamine |
| PSME1 | polyamine |
| PSMD5 | polyamine |
| PSMD8 | polyamine |
| PSMC6 | polyamine |
| PSMA3 | polyamine |
| PSMC1 | polyamine |
| PSMB5 | polyamine |
| PSMA6 | polyamine |
| PSME2 | polyamine |
| PSMA7 | polyamine |
| PSMD10 | polyamine |
| SMS | polyamine |
| PSMD7 | polyamine |
| OAZ1 | polyamine |
| PSMA2 | polyamine |
| PSMD3 | polyamine |
| PSMD11 | polyamine |
| PSMD9 | polyamine |

| | |
|---|---|
| PSMD14 | polyamine |
| ODC1 | polyamine |
| SRM | polyamine |
| AGMAT | polyamine |
| AMD1 | polyamine |
| PSMF1 | polyamine |
| PSMB2 | polyamine |
| SEM1 | polyamine |
| PSMA1 | polyamine |
| SAT1 | polyamine |
| PSME3 | polyamine |
| PSMB7 | polyamine |
| PSMB6 | polyamine |
| AZIN2 | polyamine |
| PSMA5 | polyamine |
| OAZ3 | polyamine |
| PAOX | polyamine |
| PSMA8 | polyamine |
| AZIN1 | polyamine |
| PSMD4 | polyamine |
| PSMB4 | polyamine |
| PSMC2 | polyamine |
| PSMD6 | polyamine |
| PSMC3 | polyamine |
| PSMD1 | polyamine |
| PSMD2 | polyamine |
| OAZ2 | polyamine |
| NQO1 | polyamine |
| PSMD13 | polyamine |
| PSMD12 | polyamine |
| PSMB8 | polyamine |
| PSMB10 | polyamine |
| PSMB11 | polyamine |
| PSMB9 | polyamine |
| PSMB3 | polyamine |
