## Supplementary material for "Interpretable Aging Signatures in Human Retinal Cell Types Revealed by Single-Cell RNA Sequencing and Sparse Logistic Regression": Table S2

Table S2: Composition of major retinal cell types across the 18 samples.

| Sample | Count_MGC | Percent_MGC | Count_Microglia | Percent_Microglia | Count_Cone |
|---|---|---|---|---|---|
| 1 | 1794 | 13.07008597 | 3284 | 23.92539706 | 276 |
| 2 | 123 | 1.305593886 | 86 | 0.912854262 | 55 |
| 3 | 183 | 4.150601043 | 17 | 0.38557496 | 18 |
| 4 | 931 | 16.69955157 | 213 | 3.820627803 | 79 |
| 5 | 1615 | 30.65679575 | 783 | 14.86332574 | 121 |
| 6 | 918 | 7.495101241 | 939 | 7.666557805 | 43 |
| 7 | 933 | 8.110222531 | 114 | 0.990959666 | 69 |
| 8 | 805 | 5.647141354 | 228 | 1.599438793 | 65 |
| 9 | 1330 | 10.45761912 | 32 | 0.251611889 | 131 |
| 10 | 1571 | 11.0416081 | 56 | 0.393590104 | 178 |
| 11 | 736 | 5.776626638 | 166 | 1.302880465 | 47 |
| 12 | 1039 | 6.044563384 | 98 | 0.570132061 | 67 |
| 13 | 570 | 3.570533701 | 67 | 0.419694312 | 43 |
| 14 | 531 | 3.73338958 | 65 | 0.457006257 | 44 |
| 15 | 298 | 1.75738633 | 74 | 0.436397948 | 337 |
| 16 | 459 | 2.661023827 | 76 | 0.440605252 | 419 |
| 17 | 1053 | 9.681868334 | 57 | 0.524089739 | 97 |
| 18 | 1258 | 8.352699024 | 48 | 0.318703937 | 173 |

| Percent_Cone | Count_BC | Percent_BC | Count_Rod | Percent_Rod | Count_TC | Percent_TC |
|---|---|---|---|---|---|---|
| 2.010782457 | 2269 | 16.53067172 | 5132 | 37.38889698 | 602 | 4.385837097 |
| 0.583802144 | 380 | 4.033542087 | 8697 | 92.31504087 | 42 | 0.445812546 |
| 0.40825584 | 26 | 0.58970288 | 4154 | 94.2163756 | 5 | 0.1134044 |
| 1.417040359 | 1101 | 19.74887892 | 3015 | 54.08071749 | 73 | 1.30941704 |
| 2.296886864 | 581 | 11.02885345 | 1864 | 35.38344723 | 19 | 0.360668185 |
| 0.351077727 | 553 | 4.515022861 | 9485 | 77.44121489 | 23 | 0.187785761 |
| 0.599791377 | 1531 | 13.30841446 | 8509 | 73.96557719 | 0 | 0 |
| 0.455980358 | 1232 | 8.64258155 | 11575 | 81.1995791 | 0 | 0 |
| 1.030036169 | 767 | 6.030822456 | 10373 | 81.56156628 | 3 | 0.023588615 |
| 1.251054259 | 717 | 5.03935901 | 11583 | 81.40989598 | 3 | 0.021085184 |
| 0.368887842 | 740 | 5.808021348 | 10874 | 85.34651911 | 2 | 0.015697355 |
| 0.389784164 | 815 | 4.741404387 | 14900 | 86.683344 | 3 | 0.017453022 |
| 0.269356051 | 754 | 4.723127036 | 14293 | 89.53269857 | 0 | 0 |
| 0.309358082 | 560 | 3.93728468 | 12862 | 90.43099206 | 0 | 0 |
| 1.987379843 | 795 | 4.688329304 | 15290 | 90.16925164 | 7 | 0.041280887 |
| 2.429126326 | 1073 | 6.220650472 | 15024 | 87.10070149 | 3 | 0.017392313 |
| 0.891872012 | 1443 | 13.26774549 | 7882 | 72.47149687 | 0 | 0 |
| 1.148662107 | 1918 | 12.73487816 | 11274 | 74.85558728 | 0 | 0 |

| Count_Astrocyte | Percent_Astrocyte | Count_AC | Percent_AC | Count_HC | Percent_HC | Count_RPE |
|---|---|---|---|---|---|---|
| 73 | 0.531837389 | 88 | 0.641119044 | 194 | 1.413376075 | 12 |
| 4 | 0.042458338 | 19 | 0.201677104 | 15 | 0.159218767 | 0 |
| 0 | 0 | 6 | 0.13608528 | 0 | 0 | 0 |
| 28 | 0.502242152 | 42 | 0.753363229 | 90 | 1.614349776 | 0 |
| 67 | 1.271829916 | 106 | 2.012148823 | 107 | 2.031131359 | 4 |
| 30 | 0.244937949 | 183 | 1.494121489 | 64 | 0.522534291 | 0 |
| 53 | 0.460709318 | 187 | 1.625521558 | 95 | 0.825799722 | 0 |
| 42 | 0.294633462 | 199 | 1.396001403 | 105 | 0.736583655 | 0 |
| 16 | 0.125805944 | 3 | 0.023588615 | 51 | 0.401006448 | 5 |
| 21 | 0.147596289 | 24 | 0.168681473 | 52 | 0.365476525 | 15 |
| 26 | 0.204065615 | 76 | 0.59649949 | 61 | 0.478769327 | 0 |
| 38 | 0.221071616 | 123 | 0.715573914 | 80 | 0.465413928 | 0 |
| 22 | 0.137810073 | 101 | 0.632673515 | 107 | 0.670258081 | 0 |
| 13 | 0.091401251 | 66 | 0.464037123 | 74 | 0.520284047 | 0 |
| 6 | 0.035383617 | 103 | 0.607418765 | 47 | 0.27717167 | 0 |
| 22 | 0.127543626 | 111 | 0.643515566 | 57 | 0.330453939 | 0 |
| 27 | 0.248253034 | 218 | 2.004413387 | 84 | 0.772342773 | 0 |
| 47 | 0.312064272 | 180 | 1.195139765 | 138 | 0.91627382 | 0 |

| Percent_RPE | Count_RGC | Percent_RGC |
|---|---|---|
| 0.087425324 | 2 | 0.014570887 |
| 0 | 0 | 0 |
| 0 | 0 | 0 |
| 0 | 3 | 0.053811659 |
| 0.075930144 | 1 | 0.018982536 |
| 0 | 10 | 0.081645983 |
| 0 | 13 | 0.113004172 |
| 0 | 4 | 0.02806033 |
| 0.039314358 | 7 | 0.055040101 |
| 0.105425921 | 8 | 0.056227158 |
| 0 | 13 | 0.102032807 |
| 0 | 26 | 0.151259526 |
| 0 | 7 | 0.043848659 |
| 0 | 8 | 0.056246924 |
| 0 | 0 | 0 |
| 0 | 5 | 0.028987188 |
| 0 | 15 | 0.137918352 |
| 0 | 25 | 0.165991634 |
