## Supplementary material for "Interpretable Aging Signatures in Human Retinal Cell Types Revealed by Single-Cell RNA Sequencing and Sparse Logistic Regression": Table S3

Table S3: Composition of bipolar cell subtypes across the 18 samples.

| Sample | Count_IMB | Percent_IMB | Count_DB3b | Percent_DB3b | Count_DB2 | Percent_DB2 |
|---|---|---|---|---|---|---|
| 1 | 232 | 10.22476862 | 271 | 11.94358748 | 290 | 12.78096078 |
| 2 | 31 | 8.157894737 | 21 | 5.526315789 | 53 | 13.94736842 |
| 3 | 0 | 0 | 1 | 3.846153846 | 1 | 3.846153846 |
| 4 | 95 | 8.628519528 | 24 | 2.179836512 | 25 | 2.270663034 |
| 5 | 67 | 11.53184165 | 36 | 6.196213425 | 55 | 9.466437177 |
| 6 | 64 | 11.57323689 | 28 | 5.063291139 | 39 | 7.05244123 |
| 7 | 205 | 13.38994121 | 61 | 3.984323971 | 101 | 6.596995428 |
| 8 | 174 | 14.12337662 | 14 | 1.136363636 | 81 | 6.574675325 |
| 9 | 54 | 7.04041721 | 52 | 6.779661017 | 82 | 10.69100391 |
| 10 | 79 | 11.0181311 | 44 | 6.136680614 | 103 | 14.36541144 |
| 11 | 116 | 15.67567568 | 47 | 6.351351351 | 87 | 11.75675676 |
| 12 | 131 | 16.07361963 | 42 | 5.153374233 | 96 | 11.7791411 |
| 13 | 108 | 14.32360743 | 27 | 3.580901857 | 87 | 11.53846154 |
| 14 | 81 | 14.46428571 | 35 | 6.25 | 80 | 14.28571429 |
| 15 | 135 | 16.98113208 | 46 | 5.786163522 | 58 | 7.295597484 |
| 16 | 154 | 14.35228332 | 78 | 7.269338304 | 106 | 9.878844362 |
| 17 | 217 | 15.03811504 | 85 | 5.890505891 | 179 | 12.4047124 |
| 18 | 285 | 14.85922836 | 100 | 5.213764338 | 247 | 12.87799791 |

| Count_RB | Percent_RB | Count_DB4b | Percent_DB4b | Count_DB4a | Percent_DB4a | Count_DB3a |
|---|---|---|---|---|---|---|
| 594 | 26.17893345 | 29 | 1.278096078 | 268 | 11.81137065 | 92 |
| 92 | 24.21052632 | 7 | 1.842105263 | 29 | 7.631578947 | 22 |
| 5 | 19.23076923 | 0 | 0 | 2 | 7.692307692 | 0 |
| 758 | 68.84650318 | 14 | 1.271571299 | 28 | 2.543142598 | 19 |
| 179 | 30.80895009 | 4 | 0.688468158 | 33 | 5.679862306 | 22 |
| 135 | 24.41229656 | 14 | 2.53164557 | 80 | 14.46654611 | 28 |
| 520 | 33.96472894 | 20 | 1.306335728 | 146 | 9.536250816 | 60 |
| 381 | 30.92532468 | 13 | 1.055194805 | 119 | 9.659090909 | 62 |
| 420 | 54.75880052 | 5 | 0.651890482 | 33 | 4.302477184 | 4 |
| 200 | 27.89400279 | 10 | 1.394700139 | 78 | 10.87866109 | 21 |
| 189 | 25.54054054 | 3 | 0.405405405 | 84 | 11.35135135 | 34 |
| 165 | 20.24539877 | 7 | 0.858895706 | 89 | 10.9202454 | 32 |
| 189 | 25.066313 | 22 | 2.917771883 | 56 | 7.427055703 | 19 |
| 140 | 25 | 2 | 0.357142857 | 31 | 5.535714286 | 30 |
| 190 | 23.89937107 | 9 | 1.132075472 | 87 | 10.94339623 | 37 |
| 205 | 19.10531221 | 14 | 1.304753029 | 103 | 9.599254427 | 60 |
| 504 | 34.92723493 | 18 | 1.247401247 | 111 | 7.692307692 | 11 |
| 601 | 31.33472367 | 20 | 1.042752868 | 172 | 8.967674661 | 19 |

| Percent_DB3a | Count_FMB | Percent_FMB | Count_DB1 | Percent_DB1 | Count_DB5 | Percent_DB5 |
|---|---|---|---|---|---|---|
| 4.054649625 | 177 | 7.800793301 | 27 | 1.18995152 | 187 | 8.241516086 |
| 5.789473684 | 60 | 15.78947368 | 26 | 6.842105263 | 26 | 6.842105263 |
| 0 | 2 | 7.692307692 | 1 | 3.846153846 | 6 | 23.07692308 |
| 1.725703906 | 22 | 1.99818347 | 15 | 1.36239782 | 56 | 5.086285195 |
| 3.786574871 | 56 | 9.638554217 | 27 | 4.647160069 | 52 | 8.950086059 |
| 5.063291139 | 35 | 6.329113924 | 12 | 2.169981917 | 37 | 6.690777577 |
| 3.919007185 | 82 | 5.355976486 | 56 | 3.657740039 | 139 | 9.079033312 |
| 5.032467532 | 65 | 5.275974026 | 59 | 4.788961039 | 115 | 9.334415584 |
| 0.521512386 | 28 | 3.650586701 | 7 | 0.912646675 | 65 | 8.474576271 |
| 2.928870293 | 37 | 5.160390516 | 18 | 2.510460251 | 65 | 9.065550907 |
| 4.594594595 | 67 | 9.054054054 | 15 | 2.027027027 | 55 | 7.432432432 |
| 3.926380368 | 82 | 10.06134969 | 35 | 4.294478528 | 65 | 7.975460123 |
| 2.519893899 | 50 | 6.631299735 | 13 | 1.724137931 | 61 | 8.090185676 |
| 5.357142857 | 42 | 7.5 | 18 | 3.214285714 | 54 | 9.642857143 |
| 4.65408805 | 70 | 8.805031447 | 14 | 1.761006289 | 92 | 11.57232704 |
| 5.591798695 | 113 | 10.53122088 | 29 | 2.702702703 | 112 | 10.43802423 |
| 0.762300762 | 122 | 8.454608455 | 31 | 2.148302148 | 82 | 5.682605683 |
| 0.990615224 | 164 | 8.550573514 | 31 | 1.616266945 | 160 | 8.342022941 |

| Count_BB+GB | Percent_BB+GB | Count_OFFx | Percent_OFFx | Count_DB6 | Percent_DB6 |
|---|---|---|---|---|---|
| 39 | 1.718818863 | 23 | 1.013662406 | 40 | 1.762891141 |
| 4 | 1.052631579 | 2 | 0.526315789 | 7 | 1.842105263 |
| 2 | 7.692307692 | 0 | 0 | 6 | 23.07692308 |
| 13 | 1.180744777 | 5 | 0.454132607 | 27 | 2.452316076 |
| 20 | 3.442340792 | 9 | 1.549053356 | 21 | 3.614457831 |
| 51 | 9.222423146 | 7 | 1.265822785 | 23 | 4.159132007 |
| 65 | 4.245591117 | 33 | 2.155453952 | 43 | 2.808621816 |
| 49 | 3.977272727 | 34 | 2.75974026 | 66 | 5.357142857 |
| 4 | 0.521512386 | 3 | 0.391134289 | 10 | 1.303780965 |
| 17 | 2.370990237 | 14 | 1.952580195 | 31 | 4.323570432 |
| 7 | 0.945945946 | 18 | 2.432432432 | 18 | 2.432432432 |
| 21 | 2.576687117 | 21 | 2.576687117 | 29 | 3.558282209 |
| 59 | 7.824933687 | 34 | 4.50928382 | 29 | 3.846153846 |
| 13 | 2.321428571 | 23 | 4.107142857 | 11 | 1.964285714 |
| 27 | 3.396226415 | 8 | 1.006289308 | 22 | 2.767295597 |
| 51 | 4.753028891 | 18 | 1.677539609 | 30 | 2.795899348 |
| 71 | 4.92030492 | 5 | 0.346500347 | 7 | 0.485100485 |
| 95 | 4.953076121 | 20 | 1.042752868 | 4 | 0.208550574 |
