## Supplementary material for "Interpretable Aging Signatures in Human Retinal Cell Types Revealed by Single-Cell RNA Sequencing and Sparse Logistic Regression": Table S4

Table S4: Differentially expressed genes identified through both single-cell differential expression and pseudobulk differential expression analyses across 3 retinal interneuron populations: amacrine cells, horizontal cells, and bipolar cells.

| gene | CellType | Direction | sc_avg_log2FC | pb_avg_log2FC | sc_p_val_adj | pb_p_val_adj |
|---|---|---|---|---|---|---|
| VEGFA | AC | old | 1.699981236 | 2.105961581 | 0.006815899 | 0.002312581 |
| BNIP3 | AC | old | 1.426958 | 1.930216713 | 1.14E-06 | 0.00048982 |
| AC097534.2 | AC | old | 1.423342888 | 1.653097889 | 0.019947656 | 0.037533055 |
| P4HA1 | AC | old | 1.397900494 | 1.776551291 | 8.69E-07 | 0.000700434 |
| RNF165 | AC | old | 1.131392452 | 1.365965401 | 0.012282929 | 0.023680354 |
| PGK1 | AC | old | 1.129651355 | 1.362255644 | 9.81E-05 | 0.011170354 |
| GPI | AC | old | 1.052677223 | 1.269869407 | 0.043109388 | 0.013456438 |
| HLA-C | AC | old | 0.884163564 | 1.689802466 | 0.002711751 | 0.000172771 |
| HLA-A | AC | old | 0.674256961 | 1.379792609 | 0.014982641 | 0.002589486 |
| TMSB10 | AC | old | 0.627196302 | 2.938562522 | 3.17E-07 | 2.86E-06 |
| SSBP3 | AC | old | 0.56971027 | 1.775870688 | 0.001961467 | 0.007274303 |
| CANX | AC | old | 0.524717438 | 1.046546113 | 0.015244252 | 0.008012543 |
| NAP1L3 | AC | old | 0.510365444 | 1.661762522 | 0.002706676 | 0.000848153 |
| PGRMC1 | AC | old | 0.379956782 | 1.752960468 | 0.007391447 | 2.15E-05 |
| UBE2B | AC | old | 0.344720026 | 1.720911223 | 0.01406851 | 0.000221632 |
| ACTG1 | AC | old | 0.312653862 | 1.118219716 | 8.27E-05 | 0.008903125 |
| MAP1B | AC | old | 0.304347676 | 1.22765505 | 0.040466856 | 0.015092452 |
| MARCKS | AC | old | 0.248233873 | 1.918094683 | 0.001055381 | 9.81E-06 |
| ACTB | AC | old | 0.208856488 | 1.754156199 | 4.15E-05 | 3.43E-05 |
| DPYSL2 | AC | old | 0.201769036 | 1.196261523 | 0.016822051 | 0.002654803 |
| PNMA2 | AC | old | 0.201612846 | 1.927143579 | 0.008698294 | 0.001349536 |
| MT-CO2 | AC | young | -0.240085204 | -1.075962419 | 0.02771934 | 0.011775488 |
| SFPQ | AC | young | -0.275969814 | -1.281173429 | 0.033932964 | 0.002987623 |
| AC002463.1 | AC | young | -0.569816163 | -1.711792845 | 0.000939959 | 0.000273748 |
| ZNF804A | AC | young | -0.982385832 | -2.454321253 | 4.78E-07 | 1.10E-06 |
| CNTN5 | AC | young | -1.075181115 | -2.641513608 | 0.028030523 | 9.03E-15 |
| CCL2 | BC | old | 4.025378087 | 2.025505096 | 1.24E-09 | 0.000220372 |
| HLA-DRA | BC | old | 3.185557587 | 1.199917425 | 1.63E-06 | 0.032835313 |
| S100A6 | BC | old | 3.124517958 | 1.886230595 | 2.56E-14 | 0.000113587 |
| ID1 | BC | old | 2.713420477 | 1.452757715 | 2.25E-05 | 0.004395024 |
| LINC02649 | BC | old | 2.495717292 | 3.01332857 | 1.73E-46 | 1.03E-10 |
| IFITM3 | BC | old | 2.357237869 | 1.759896611 | 1.19E-18 | 0.00040775 |
| ATG9B | BC | old | 2.311632012 | 1.671579091 | 3.58E-10 | 0.031432923 |
| NPVF | BC | old | 2.266628654 | 1.201522146 | 2.40E-35 | 0.040227493 |
| IFITM2 | BC | old | 2.264963761 | 1.683723115 | 1.31E-11 | 0.001949078 |
| GFAP | BC | old | 2.192849885 | 1.8618918 | 7.56E-22 | 5.77E-05 |
| S100A10 | BC | old | 2.159415027 | 1.408935532 | 0.000100739 | 0.029256281 |
| TMEM255A | BC | old | 1.881458399 | 1.403812155 | 3.73E-11 | 0.002248317 |
| HK2 | BC | old | 1.878352254 | 1.558055003 | 2.76E-19 | 0.000948056 |
| LGALS3 | BC | old | 1.76032401 | 1.830079488 | 2.68E-21 | 3.56E-05 |
| GPX3 | BC | old | 1.613317975 | 2.015187128 | 2.08E-31 | 3.94E-08 |
| SYCP2L | BC | old | 1.558083868 | 1.79948511 | 5.98E-07 | 0.000760477 |
| AC005040.2 | BC | old | 1.51017485 | 2.881892691 | 1.33E-25 | 2.42E-12 |
| IGFBP6 | BC | old | 1.496495896 | 1.045361735 | 0.006410514 | 0.025883102 |
| PFKFB4 | BC | old | 1.468596474 | 1.88751705 | 1.42E-45 | 2.27E-06 |
| PDK1 | BC | old | 1.428076809 | 1.525067806 | 2.83E-42 | 4.83E-05 |
| BAG3 | BC | old | 1.413356922 | 2.195432834 | 4.59E-23 | 4.41E-09 |
| ID3 | BC | old | 1.385175229 | 1.394904554 | 6.62E-06 | 0.000710702 |
| BHLHE40 | BC | old | 1.377080072 | 1.912344416 | 2.29E-79 | 1.65E-12 |
| LDLRAD4 | BC | old | 1.367757024 | 1.978031351 | 3.40E-21 | 6.05E-08 |
| DTNA | BC | old | 1.344101042 | 2.434801264 | 1.63E-39 | 1.34E-15 |
| LINC02224 | BC | old | 1.343495377 | 1.620361662 | 3.91E-05 | 0.000179478 |
| FRZB | BC | old | 1.339804065 | 1.154074188 | 1.54E-05 | 0.032835313 |

| | | | | | | |
|---|---|---|---|---|---|---|
| SAMD4A | BC | old | 1.339800877 | 1.410800058 | 9.64E-09 | 0.002516105 |
| CRYAB | BC | old | 1.337452151 | 3.000200622 | 1.20E-80 | 4.18E-23 |
| AK4 | BC | old | 1.303839462 | 1.571751667 | 4.70E-40 | 1.99E-06 |
| CEBPD | BC | old | 1.29940691 | 1.168594116 | 7.14E-06 | 0.031963922 |
| PARD6G-AS1 | BC | old | 1.299117445 | 1.439414756 | 6.52E-10 | 0.005484132 |
| MT1G | BC | old | 1.258672816 | 2.305468813 | 4.50E-08 | 2.66E-08 |
| RAPGEF4 | BC | old | 1.246415961 | 1.829349036 | 9.88E-23 | 7.07E-06 |
| ARHGAP10 | BC | old | 1.21956429 | 1.337540538 | 1.37E-07 | 0.008038524 |
| MT3 | BC | old | 1.19320272 | 1.123598699 | 6.31E-05 | 0.022967086 |
| P4HA1 | BC | old | 1.192908326 | 1.621201167 | 3.55E-61 | 5.74E-09 |
| PPP1R3C | BC | old | 1.184363713 | 1.404015356 | 5.94E-15 | 0.000174225 |
| PFKFB3 | BC | old | 1.141537732 | 1.256899714 | 1.79E-13 | 0.007573061 |
| STAC2 | BC | old | 1.114955343 | 1.180144806 | 0.004693884 | 0.046084179 |
| PPFIA4 | BC | old | 1.089453017 | 1.245909681 | 1.57E-07 | 0.040872164 |
| EGLN3 | BC | old | 1.078105583 | 1.562549561 | 7.11E-19 | 1.04E-07 |
| PCSK5 | BC | old | 1.077437172 | 1.245496853 | 0.000280455 | 0.009156612 |
| PLOD2 | BC | old | 1.067822818 | 1.403796834 | 6.92E-31 | 1.12E-06 |
| NDRG1 | BC | old | 1.027852791 | 1.438530499 | 4.85E-19 | 9.20E-05 |
| GLUL | BC | old | 1.021691252 | 0.9301776 | 0.001871802 | 0.035589993 |
| TEAD3 | BC | old | 1.017586262 | 0.927143668 | 0.040655636 | 0.037209141 |
| GHR | BC | old | 1.013711931 | 1.196761547 | 0.044232238 | 0.017258232 |
| PFKP | BC | old | 0.990353011 | 1.252943178 | 1.80E-15 | 0.000426224 |
| NR4A3 | BC | old | 0.989322685 | 0.968375635 | 1.79E-05 | 0.016035886 |
| AC097534.2 | BC | old | 0.980504726 | 1.019180551 | 2.36E-15 | 0.003046716 |
| C4orf47 | BC | old | 0.977036604 | 1.452618572 | 6.04E-18 | 7.27E-05 |
| RAB31 | BC | old | 0.96584142 | 1.316124807 | 0.008053717 | 0.031958551 |
| PLOD1 | BC | old | 0.963819064 | 1.088548279 | 0.000550369 | 0.044608954 |
| GSN | BC | old | 0.963162441 | 1.08816897 | 0.002493149 | 0.041449145 |
| NTRK2 | BC | old | 0.961139324 | 0.77172898 | 0.012335113 | 0.046750252 |
| HTR5A | BC | old | 0.954352567 | 1.288942946 | 0.000326836 | 0.049237773 |
| MEIS2 | BC | old | 0.948562424 | 1.47897236 | 1.56E-14 | 3.50E-05 |
| B4GALNT4 | BC | old | 0.940724349 | 1.202088664 | 0.000150777 | 0.016269282 |
| ANKRD37 | BC | old | 0.922264863 | 1.439098212 | 2.62E-18 | 1.12E-07 |
| DENND2A | BC | old | 0.910821593 | 1.359125383 | 1.18E-05 | 0.001694259 |
| MT1M | BC | old | 0.897989926 | 1.256949243 | 1.43E-05 | 0.010811731 |
| NPAS2 | BC | old | 0.891621334 | 1.836825596 | 1.30E-19 | 1.10E-06 |
| TRPM3 | BC | old | 0.88934887 | 1.09498952 | 0.004576832 | 0.001329802 |
| SH3GL3 | BC | old | 0.86921481 | 1.92963431 | 6.93E-44 | 7.15E-09 |
| KDM3A | BC | old | 0.864457595 | 1.333523889 | 6.36E-26 | 0.000167684 |
| HLA-DQB1 | BC | old | 0.855540392 | 1.586773566 | 1.52E-11 | 1.99E-05 |
| SEPTIN6 | BC | old | 0.85419169 | 1.280680642 | 2.46E-05 | 0.002261217 |
| PDK3 | BC | old | 0.850214182 | 1.289598226 | 8.89E-16 | 0.000332334 |
| EFNA5 | BC | old | 0.849381871 | 1.291660939 | 0.007370107 | 0.006318682 |
| MCTP1 | BC | old | 0.848547552 | 1.145137203 | 0.005158676 | 0.021259515 |
| MMP2 | BC | old | 0.848520185 | 0.800461748 | 8.41E-07 | 0.032514862 |
| ERO1A | BC | old | 0.848381419 | 1.44309024 | 2.26E-30 | 1.24E-06 |
| C8orf34 | BC | old | 0.847715412 | 1.484777924 | 0.034884951 | 0.000909259 |
| GRIK2 | BC | old | 0.845102005 | 1.142798175 | 9.34E-05 | 0.001741821 |
| PAM | BC | old | 0.8365206 | 1.035009164 | 0.004667887 | 0.011413715 |
| FHOD3 | BC | old | 0.818391696 | 1.411991241 | 6.56E-06 | 0.001110147 |
| SEPTIN9 | BC | old | 0.808456778 | 1.002017616 | 0.003434222 | 0.022967086 |
| SLC2A3 | BC | old | 0.805087712 | 1.462875808 | 5.03E-26 | 2.80E-05 |
| P4HA2 | BC | old | 0.802489575 | 1.573491107 | 2.40E-19 | 6.17E-05 |
| AP001972.3 | BC | old | 0.800526301 | 1.595190599 | 6.26E-11 | 3.24E-05 |

| | | | | | | |
|---|---|---|---|---|---|---|
| SORBS1 | BC | old | 0.786012564 | 1.252191271 | 9.32E-11 | 0.000153633 |
| MBOAT2 | BC | old | 0.78127623 | 1.289198119 | 8.16E-16 | 0.000183021 |
| PDZD4 | BC | old | 0.779628612 | 1.456163566 | 1.68E-07 | 0.000860548 |
| GADD45B | BC | old | 0.775660526 | 1.584294379 | 1.07E-10 | 3.58E-05 |
| KAZN | BC | old | 0.774891798 | 1.280701268 | 9.33E-13 | 0.002261217 |
| ARHGEF10 | BC | old | 0.771430393 | 1.157923682 | 3.42E-05 | 0.02849206 |
| NR4A2 | BC | old | 0.767164247 | 1.804887867 | 3.13E-17 | 7.06E-07 |
| ITPR2 | BC | old | 0.763556887 | 1.482098685 | 1.88E-11 | 0.000367086 |
| CCNG2 | BC | old | 0.762106701 | 1.280802958 | 6.75E-39 | 2.58E-06 |
| MLLT3 | BC | old | 0.75703575 | 1.351841381 | 2.85E-11 | 0.00010435 |
| DNAH17 | BC | old | 0.749977711 | 1.114770471 | 3.22E-13 | 0.000499133 |
| HLA-A | BC | old | 0.746145373 | 0.752236001 | 1.12E-08 | 0.029271903 |
| ESYT2 | BC | old | 0.742779949 | 1.135133841 | 5.74E-13 | 0.000312435 |
| STEAP2 | BC | old | 0.740862848 | 1.351535867 | 3.50E-09 | 0.000658162 |
| CYP27A1 | BC | old | 0.735581783 | 0.911623007 | 0.000417766 | 0.027083976 |
| YEATS2 | BC | old | 0.733408153 | 1.369354117 | 4.00E-30 | 2.42E-07 |
| FAM124A | BC | old | 0.733382746 | 0.968291168 | 7.18E-05 | 0.026703033 |
| ADAM19 | BC | old | 0.726919337 | 1.041187419 | 1.62E-11 | 0.00036447 |
| ARFGEF3 | BC | old | 0.724151457 | 1.205652989 | 5.55E-19 | 0.000256031 |
| SHISA5 | BC | old | 0.723490505 | 1.064473517 | 9.31E-05 | 0.008369491 |
| ULK3 | BC | old | 0.723451103 | 1.171423465 | 3.35E-10 | 0.000442308 |
| ZFYVE28 | BC | old | 0.716644128 | 1.294421822 | 1.94E-11 | 0.000387936 |
| RNF24 | BC | old | 0.716188064 | 1.141685603 | 1.69E-09 | 0.007114407 |
| PDLIM5 | BC | old | 0.715182151 | 1.327082037 | 1.19E-15 | 0.001657989 |
| RNF165 | BC | old | 0.714501843 | 1.055010347 | 4.03E-08 | 0.008404551 |
| NEAT1 | BC | old | 0.709955672 | 0.957706988 | 6.86E-11 | 0.000276228 |
| PDE2A | BC | old | 0.706548025 | 1.40083169 | 0.000170618 | 0.002760984 |
| FAM138D | BC | old | 0.703559793 | 1.243679128 | 0.000377921 | 0.003755984 |
| LRP2BP | BC | old | 0.697121534 | 0.956121897 | 1.51E-06 | 0.001985705 |
| ST6GALNAC3 | BC | old | 0.694881367 | 1.239569539 | 4.87E-07 | 0.000203274 |
| HIF1A | BC | old | 0.693329735 | 0.611014964 | 8.20E-08 | 0.046426518 |
| OGA | BC | old | 0.69029798 | 0.816564136 | 1.65E-22 | 0.001048239 |
| CDC14B | BC | old | 0.688332203 | 1.248255469 | 1.37E-07 | 0.001781682 |
| EGR1 | BC | old | 0.684255604 | 0.843863766 | 4.41E-06 | 0.003054306 |
| CORO1C | BC | old | 0.678952256 | 1.120937478 | 0.005179935 | 0.014957927 |
| RNF144A | BC | old | 0.676789765 | 1.337562875 | 2.93E-08 | 0.00059168 |
| CARMIL1 | BC | old | 0.675989024 | 0.824278529 | 4.14E-09 | 0.023381643 |
| C20orf194 | BC | old | 0.666435422 | 0.788660193 | 1.58E-05 | 0.013691136 |
| CPEB3 | BC | old | 0.661773374 | 0.952914376 | 3.70E-10 | 0.000362809 |
| PRTG | BC | old | 0.661452235 | 1.450762267 | 2.27E-06 | 6.66E-05 |
| PDLIM3 | BC | old | 0.6549785 | 1.334441664 | 0.000154694 | 0.00189325 |
| NTRK3 | BC | old | 0.651591332 | 0.990565903 | 0.00058869 | 0.032248932 |
| GBE1 | BC | old | 0.647304405 | 0.942999803 | 7.05E-08 | 0.001949078 |
| PPP1R15A | BC | old | 0.642568436 | 1.698040556 | 4.68E-17 | 2.00E-06 |
| GRK3 | BC | old | 0.642339367 | 1.389820661 | 3.72E-09 | 0.000252093 |
| KDM4C | BC | old | 0.640275508 | 1.005496543 | 1.71E-21 | 4.18E-05 |
| SETD5 | BC | old | 0.638171832 | 0.769695332 | 7.29E-09 | 0.001322873 |
| TMEM179 | BC | old | 0.637987029 | 1.378100815 | 1.14E-13 | 1.94E-05 |
| NOS1AP | BC | old | 0.635396837 | 1.271121176 | 0.001613658 | 0.004835209 |
| CDK19 | BC | old | 0.634189687 | 1.073750765 | 2.94E-12 | 0.00094327 |
| TENM4 | BC | old | 0.633170366 | 1.580693858 | 2.13E-13 | 2.90E-05 |
| SLCO4A1 | BC | old | 0.630751825 | 1.234491555 | 6.47E-07 | 0.002830218 |
| ZDHHC22 | BC | old | 0.630131051 | 1.128279868 | 8.00E-06 | 0.007904327 |
| XKR6 | BC | old | 0.629776196 | 0.855796079 | 3.13E-14 | 0.00059168 |

| | | | | | | |
|---|---|---|---|---|---|---|
| HS3ST4 | BC | old | 0.62939946 | 1.013335633 | 1.59E-08 | 0.008038524 |
| MAST1 | BC | old | 0.625355758 | 1.145202768 | 1.52E-06 | 0.004312919 |
| AGAP1 | BC | old | 0.623821318 | 0.704779838 | 6.95E-23 | 0.000322815 |
| NUP93 | BC | old | 0.622582006 | 1.279583833 | 3.09E-11 | 2.77E-05 |
| CXXC5 | BC | old | 0.622496555 | 1.083464027 | 7.35E-19 | 6.07E-05 |
| APP | BC | old | 0.621478068 | 1.834365173 | 3.89E-30 | 3.61E-09 |
| DPYSL4 | BC | old | 0.621437675 | 1.162369074 | 1.38E-08 | 0.001640391 |
| TENM3 | BC | old | 0.619744361 | 0.727939809 | 4.99E-07 | 0.027083976 |
| JPH4 | BC | old | 0.617890005 | 1.036480713 | 2.46E-05 | 0.003157433 |
| RAPGEF6 | BC | old | 0.616648935 | 0.787941673 | 5.56E-09 | 0.004799978 |
| ZCCHC2 | BC | old | 0.61662018 | 1.141525765 | 4.85E-14 | 0.000313293 |
| ZBTB16 | BC | old | 0.616407251 | 1.291000187 | 1.01E-07 | 0.000611393 |
| KDM4B | BC | old | 0.615978375 | 1.118911511 | 2.15E-16 | 0.000265562 |
| C11orf80 | BC | old | 0.615934836 | 0.960472701 | 0.023106986 | 0.022547489 |
| CPD | BC | old | 0.615199056 | 1.61228353 | 8.38E-44 | 1.04E-08 |
| CHD7 | BC | old | 0.614572295 | 1.30629898 | 1.34E-10 | 0.000171379 |
| C1orf21 | BC | old | 0.609772709 | 1.339463662 | 1.56E-09 | 9.60E-05 |
| CACNA1D | BC | old | 0.606983136 | 0.808750345 | 4.18E-08 | 0.007354567 |
| ST5 | BC | old | 0.604108681 | 1.101470652 | 0.022134157 | 0.00456954 |
| SEMA5B | BC | old | 0.602736398 | 1.274977732 | 1.65E-05 | 0.001360401 |
| MANBA | BC | old | 0.599407776 | 1.126248487 | 5.54E-13 | 0.000760477 |
| NFIA | BC | old | 0.595887732 | 1.298230402 | 7.55E-16 | 1.94E-05 |
| DIXDC1 | BC | old | 0.591826158 | 1.34928404 | 3.73E-17 | 0.000267231 |
| ANKRD13A | BC | old | 0.580862832 | 1.459079523 | 2.02E-23 | 1.42E-05 |
| RBPJ | BC | old | 0.580705732 | 0.961535737 | 8.32E-23 | 0.000163079 |
| RGPD5 | BC | old | 0.58014354 | 1.152070998 | 1.10E-05 | 0.012811461 |
| DIP2C | BC | old | 0.580110072 | 0.929082693 | 1.66E-09 | 0.001219564 |
| FARP1 | BC | old | 0.574858928 | 0.903709708 | 0.007995552 | 0.047267574 |
| RRAGD | BC | old | 0.57481568 | 1.168981094 | 7.80E-14 | 0.000750629 |
| SLC2A1 | BC | old | 0.574509319 | 1.127786131 | 1.69E-06 | 0.016619703 |
| MMP16 | BC | old | 0.572668033 | 1.604080524 | 2.37E-15 | 4.57E-06 |
| SLC15A4 | BC | old | 0.571544275 | 0.930753524 | 0.002649004 | 0.020264681 |
| FMN2 | BC | old | 0.569792262 | 0.673446019 | 1.12E-05 | 0.027848148 |
| ZNF516 | BC | old | 0.568342506 | 1.265735294 | 1.55E-11 | 4.80E-05 |
| ELMO1 | BC | old | 0.562684123 | 0.865561276 | 0.00055134 | 0.020625979 |
| INSIG2 | BC | old | 0.562043739 | 0.844851032 | 6.18E-06 | 0.028870195 |
| LRRN2 | BC | old | 0.558292427 | 0.875283509 | 0.013391071 | 0.02239856 |
| PRKAA2 | BC | old | 0.555084136 | 0.76538084 | 0.000166911 | 0.023128565 |
| CCNI | BC | old | 0.552477833 | 0.990115251 | 1.30E-37 | 4.06E-06 |
| ASAP2 | BC | old | 0.552378089 | 0.664813729 | 0.034189251 | 0.044742836 |
| JUN | BC | old | 0.552014578 | 1.74098336 | 2.43E-15 | 1.64E-10 |
| KCNIP4 | BC | old | 0.547021588 | 0.766391248 | 8.09E-16 | 0.003125467 |
| EPS15 | BC | old | 0.546865911 | 0.999933116 | 3.11E-07 | 0.004549525 |
| PIAS2 | BC | old | 0.544467147 | 0.918678527 | 1.21E-10 | 0.002377376 |
| TRAF3 | BC | old | 0.54327573 | 1.222912117 | 1.59E-23 | 2.48E-05 |
| XPNPEP1 | BC | old | 0.543071425 | 1.01229864 | 0.000197691 | 0.021173714 |
| EML6 | BC | old | 0.538932253 | 1.524980021 | 4.13E-24 | 2.70E-06 |
| CNN3 | BC | old | 0.537212072 | 0.929246792 | 0.00023532 | 0.019974896 |
| CYFIP2 | BC | old | 0.535971975 | 0.667472327 | 0.000850766 | 0.032514862 |
| SFMBT2 | BC | old | 0.534895776 | 0.973615372 | 0.001705221 | 0.006961346 |
| RIMKLB | BC | old | 0.534478727 | 1.151734757 | 8.41E-19 | 0.000322963 |
| CACNA1A | BC | old | 0.531579263 | 1.982965772 | 1.77E-23 | 1.03E-10 |
| MALT1 | BC | old | 0.529720293 | 1.065332461 | 9.30E-05 | 0.002429219 |
| DAPK1 | BC | old | 0.528610554 | 0.892867337 | 1.23E-11 | 0.000698506 |

| | | | | | | |
|---|---|---|---|---|---|---|
| IL1RAPL1 | BC | old | 0.527961298 | 1.290023048 | 0.025902799 | 0.000420902 |
| KIZ | BC | old | 0.52763972 | 0.84023441 | 0.000756769 | 0.010479446 |
| RAP1GAP2 | BC | old | 0.526391529 | 0.997372375 | 5.88E-06 | 0.009733948 |
| TRIO | BC | old | 0.522981701 | 0.801996464 | 1.58E-08 | 0.000789515 |
| DOCK7 | BC | old | 0.522935223 | 1.02971767 | 8.37E-16 | 0.004395024 |
| MAP2K5 | BC | old | 0.522623385 | 0.791282878 | 4.58E-06 | 0.022553497 |
| XPO6 | BC | old | 0.522072152 | 0.76452475 | 0.006329507 | 0.030843706 |
| TMEM145 | BC | old | 0.52197514 | 0.847183307 | 0.001229934 | 0.026801465 |
| DAAM1 | BC | old | 0.521257888 | 0.699379066 | 0.00364228 | 0.007631428 |
| DOCK4 | BC | old | 0.52092939 | 0.707904432 | 2.82E-05 | 0.045810166 |
| TSHZ2 | BC | old | 0.519689714 | 1.072654593 | 0.022431202 | 0.007824804 |
| TUBB2B | BC | old | 0.519373766 | 0.907319634 | 9.70E-06 | 0.013024511 |
| TTLL5 | BC | old | 0.518627967 | 0.96052126 | 1.16E-06 | 0.010158569 |
| MTCL1 | BC | old | 0.51805953 | 0.728907404 | 9.81E-09 | 0.004609547 |
| AASS | BC | old | 0.518040662 | 0.788379281 | 0.000886408 | 0.046811107 |
| PLXNB2 | BC | old | 0.517006829 | 1.134329927 | 5.91E-07 | 0.01419075 |
| ASTN1 | BC | old | 0.516669588 | 1.480801076 | 1.02E-16 | 2.93E-05 |
| NRG2 | BC | old | 0.514157853 | 1.246378809 | 3.10E-10 | 0.00025058 |
| PWWP3A | BC | old | 0.513242952 | 1.181255821 | 1.27E-21 | 1.08E-05 |
| LSAMP | BC | old | 0.513117824 | 0.658123537 | 5.36E-06 | 0.049022729 |
| LINC01695 | BC | old | 0.510273187 | 1.379042254 | 0.000584687 | 1.08E-05 |
| SFXN1 | BC | old | 0.50779572 | 0.744375864 | 1.11E-05 | 0.014284876 |
| RFLNB | BC | old | 0.507133937 | 0.941078017 | 0.001357007 | 0.017253428 |
| STARD9 | BC | old | 0.505936572 | 1.032511154 | 0.000513693 | 0.027773554 |
| RHBDL3 | BC | old | 0.499941221 | 1.033182014 | 0.015541304 | 0.008884759 |
| BRWD3 | BC | old | 0.4986952 | 0.957371494 | 5.05E-11 | 0.002775061 |
| ADCY5 | BC | old | 0.498176339 | 1.115684534 | 4.17E-11 | 0.003956231 |
| FBXL14 | BC | old | 0.497782658 | 1.783466826 | 9.86E-15 | 3.32E-07 |
| RLF | BC | old | 0.496935419 | 0.824524999 | 3.78E-08 | 0.009216127 |
| UBR5 | BC | old | 0.493070461 | 0.861536886 | 2.96E-13 | 0.000725302 |
| ASAP1 | BC | old | 0.492725164 | 0.84782618 | 0.000245314 | 0.012089232 |
| KCNJ2 | BC | old | 0.492666721 | 1.411192559 | 1.83E-09 | 4.33E-05 |
| SMARCA2 | BC | old | 0.491858704 | 0.650274512 | 2.48E-08 | 0.01176303 |
| PAK3 | BC | old | 0.491706825 | 0.932492251 | 2.30E-11 | 0.000144955 |
| CRY2 | BC | old | 0.491210263 | 1.547673677 | 1.48E-52 | 2.88E-11 |
| TMEM132C | BC | old | 0.491090311 | 0.709348239 | 1.49E-06 | 0.005541252 |
| IER5 | BC | old | 0.490883616 | 1.334150912 | 0.000170647 | 0.000207364 |
| EIF4E1B | BC | old | 0.490237542 | 0.936021351 | 0.009123206 | 0.011231135 |
| CBLB | BC | old | 0.486706683 | 0.559079706 | 4.75E-06 | 0.032814268 |
| PTP4A3 | BC | old | 0.486020765 | 0.669150481 | 0.000224172 | 0.033581941 |
| ARHGAP26 | BC | old | 0.483604347 | 1.122897146 | 6.23E-05 | 0.002764556 |
| CLASP1 | BC | old | 0.482510328 | 0.605618091 | 2.52E-05 | 0.027490322 |
| DIP2B | BC | old | 0.481874979 | 0.973146412 | 2.53E-18 | 0.000293693 |
| BTAF1 | BC | old | 0.480280575 | 0.962553395 | 6.65E-10 | 0.002654457 |
| KMT2E | BC | old | 0.479769151 | 0.844228488 | 3.23E-38 | 6.26E-05 |
| PMEPA1 | BC | old | 0.478527767 | 1.166242076 | 4.14E-09 | 0.000272416 |
| AHCYL2 | BC | old | 0.478037654 | 0.824141181 | 1.42E-06 | 0.002764556 |
| DNM3 | BC | old | 0.476303637 | 1.107895477 | 2.23E-18 | 0.000120711 |
| TTC21B | BC | old | 0.476162794 | 1.063996653 | 2.27E-09 | 0.000910306 |
| PKP1 | BC | old | 0.472562335 | 0.965599186 | 0.015545163 | 0.036950907 |
| PPP6R3 | BC | old | 0.47089786 | 0.563661167 | 0.044484882 | 0.039314932 |
| GGA2 | BC | old | 0.470535456 | 1.131168381 | 7.52E-25 | 1.79E-05 |
| APBA2 | BC | old | 0.46963831 | 0.96268635 | 0.000298627 | 0.015792052 |
| PHC2 | BC | old | 0.468801135 | 0.585853336 | 0.043911068 | 0.046956119 |

| | | | | | | |
|---|---|---|---|---|---|---|
| TFB1M | BC | old | 0.468638731 | 0.738629489 | 0.010778106 | 0.044651851 |
| POGZ | BC | old | 0.467794064 | 0.569441765 | 0.014053231 | 0.046811107 |
| KLHL3 | BC | old | 0.467580372 | 0.959328404 | 0.001081696 | 0.032940926 |
| ANKS1B | BC | old | 0.467384989 | 1.582323087 | 6.99E-32 | 2.94E-11 |
| LIMCH1 | BC | old | 0.465671891 | 1.413384887 | 3.85E-29 | 1.17E-05 |
| OTX2 | BC | old | 0.464475517 | 0.895698015 | 6.79E-23 | 1.59E-06 |
| FRMPD4 | BC | old | 0.464127733 | 1.015664067 | 0.000520284 | 0.017530794 |
| DYNC1I1 | BC | old | 0.463987745 | 0.790945317 | 4.03E-07 | 0.004366224 |
| RERE | BC | old | 0.46146382 | 0.869096028 | 7.49E-37 | 1.71E-05 |
| ZNF292 | BC | old | 0.460434335 | 0.645771694 | 7.12E-18 | 0.004366224 |
| SCN8A | BC | old | 0.457793782 | 0.905977724 | 1.09E-08 | 0.001876982 |
| LRCH2 | BC | old | 0.455076547 | 0.899326523 | 0.001438469 | 0.030749186 |
| NUP58 | BC | old | 0.455038129 | 0.697899585 | 7.24E-05 | 0.045119914 |
| DGKH | BC | old | 0.45481038 | 1.301395299 | 3.06E-11 | 0.000226061 |
| GTF2IRD1 | BC | old | 0.451176861 | 0.93297161 | 5.42E-13 | 0.00125216 |
| BHLHE41 | BC | old | 0.449590546 | 1.004272989 | 9.60E-36 | 2.45E-06 |
| MTSS1 | BC | old | 0.449564727 | 1.086863274 | 0.015858235 | 0.003708215 |
| CELF5 | BC | old | 0.449271791 | 0.945304063 | 6.75E-17 | 0.000850024 |
| PICALM | BC | old | 0.448822225 | 0.730905385 | 3.40E-05 | 0.012260515 |
| UBR3 | BC | old | 0.447787906 | 0.702367152 | 8.25E-09 | 0.004156789 |
| SEC24B | BC | old | 0.44732385 | 0.884349455 | 1.29E-09 | 0.003435133 |
| SRGAP2 | BC | old | 0.446806972 | 0.852373367 | 2.19E-05 | 0.017074106 |
| KMT2C | BC | old | 0.446450296 | 0.751746811 | 5.22E-18 | 0.001990916 |
| MAP4K5 | BC | old | 0.446437712 | 0.731349501 | 1.46E-11 | 0.005955661 |
| ELF2 | BC | old | 0.445673734 | 0.62634816 | 0.048690551 | 0.03121573 |
| LHX3 | BC | old | 0.445658778 | 1.147116816 | 0.000138839 | 0.001282389 |
| TFDP2 | BC | old | 0.445583606 | 0.631448163 | 0.000572092 | 0.049311751 |
| MLLT6 | BC | old | 0.445462305 | 0.675799867 | 6.76E-06 | 0.027254532 |
| WDR37 | BC | old | 0.444168799 | 0.765134676 | 9.50E-05 | 0.013875128 |
| SMYD2 | BC | old | 0.443833454 | 0.86239783 | 0.000382157 | 0.032204573 |
| BCAT1 | BC | old | 0.442886836 | 1.20566505 | 2.58E-30 | 2.90E-05 |
| SCYL2 | BC | old | 0.441451315 | 0.889130607 | 3.61E-06 | 0.009816449 |
| CAMK2G | BC | old | 0.441388287 | 1.044859156 | 1.22E-08 | 0.001826956 |
| CLASP2 | BC | old | 0.441249549 | 0.887324943 | 2.36E-44 | 2.87E-06 |
| PPP2R5C | BC | old | 0.440423856 | 0.913657575 | 0.0002257 | 0.002573378 |
| FIGN | BC | old | 0.439041282 | 0.83495067 | 1.13E-06 | 0.024156707 |
| CDC27 | BC | old | 0.438492394 | 0.713122758 | 0.000715151 | 0.042231307 |
| SCN2A | BC | old | 0.435478775 | 0.828698717 | 1.91E-10 | 0.0037628 |
| NBAS | BC | old | 0.43529237 | 0.736278474 | 3.32E-08 | 0.004395024 |
| PDE6G | BC | old | 0.43521034 | 1.135069513 | 0.009246747 | 0.000710125 |
| AC112493.1 | BC | old | 0.433997528 | 1.363570519 | 2.04E-05 | 0.000120814 |
| PER1 | BC | old | 0.433909816 | 1.348103896 | 7.49E-25 | 1.46E-05 |
| PHC3 | BC | old | 0.43380788 | 0.761745827 | 2.26E-10 | 0.003694357 |
| TTBK2 | BC | old | 0.433256439 | 0.763183604 | 9.50E-05 | 0.011428381 |
| KIAA1211 | BC | old | 0.432492034 | 0.876878972 | 6.48E-05 | 0.027789799 |
| TACC2 | BC | old | 0.43226392 | 1.016936449 | 2.66E-07 | 0.007120468 |
| BIRC6 | BC | old | 0.432220267 | 0.77280313 | 3.47E-16 | 0.000864074 |
| LONRF1 | BC | old | 0.431019375 | 1.079596837 | 7.40E-08 | 0.000419751 |
| DDHD2 | BC | old | 0.43060676 | 0.791096479 | 0.000125134 | 0.031607699 |
| SLC12A2 | BC | old | 0.430174795 | 1.221457552 | 6.74E-22 | 3.80E-05 |
| DPYSL5 | BC | old | 0.427930935 | 0.988867303 | 0.001321011 | 0.003908617 |
| RIMBP2 | BC | old | 0.427557107 | 0.74953057 | 0.001563544 | 0.041013686 |
| RBM33 | BC | old | 0.427421772 | 0.838540721 | 3.80E-06 | 0.004136791 |
| KIF1B | BC | old | 0.426575286 | 0.604647818 | 3.90E-11 | 0.006603807 |

| | | | | | | |
|---|---|---|---|---|---|---|
| USP28 | BC | old | 0.426270865 | 0.852418829 | 2.07E-05 | 0.023393563 |
| CAMSAP2 | BC | old | 0.425607726 | 0.612044678 | 0.003650992 | 0.028870195 |
| PPP3CC | BC | old | 0.421606397 | 1.033636549 | 8.29E-07 | 0.001457401 |
| ATP11A | BC | old | 0.418632437 | 0.832708948 | 0.000408636 | 0.014179896 |
| PATJ | BC | old | 0.415778654 | 0.672721806 | 0.023742878 | 0.023195151 |
| MLLT10 | BC | old | 0.412061551 | 0.6831793 | 1.80E-08 | 0.020132026 |
| EML4 | BC | old | 0.411765062 | 0.894699253 | 5.42E-08 | 0.00408277 |
| AFDN | BC | old | 0.409013011 | 0.895165144 | 2.43E-11 | 0.000449635 |
| TSHZ3 | BC | old | 0.407609534 | 0.78516739 | 0.002806744 | 0.032411791 |
| CCDC93 | BC | old | 0.406856934 | 0.968157312 | 5.38E-06 | 0.0037628 |
| FLT1 | BC | old | 0.406770381 | 1.02115072 | 0.001893555 | 0.004722725 |
| MAP7 | BC | old | 0.40563375 | 0.456953919 | 1.65E-18 | 0.022967086 |
| TEAD1 | BC | old | 0.404554907 | 1.315207719 | 1.62E-22 | 1.08E-05 |
| CABIN1 | BC | old | 0.402585487 | 0.863827399 | 2.18E-08 | 0.007865476 |
| PCNX1 | BC | old | 0.400980786 | 0.620064592 | 0.002156875 | 0.02829238 |
| GNAQ | BC | old | 0.40074764 | 0.833077657 | 9.93E-08 | 0.00332892 |
| MXI1 | BC | old | 0.400266236 | 0.713478801 | 1.03E-08 | 0.016515757 |
| SMCHD1 | BC | old | 0.400018546 | 0.764230348 | 5.12E-12 | 0.007180343 |
| PER3 | BC | old | 0.399150117 | 0.744163137 | 1.13E-28 | 0.000398607 |
| MAPK1IP1L | BC | old | 0.397745161 | 0.939010864 | 3.76E-08 | 0.013366188 |
| PTPRZ1 | BC | old | 0.397192542 | 0.954862066 | 8.28E-05 | 0.009791547 |
| VPS13A | BC | old | 0.396498474 | 0.889818638 | 2.21E-09 | 0.001650055 |
| PUM2 | BC | old | 0.396097112 | 0.602932388 | 8.54E-18 | 0.017530794 |
| SPRY4-AS1 | BC | old | 0.395976162 | 1.69442407 | 7.10E-15 | 6.63E-09 |
| LRBA | BC | old | 0.395855271 | 0.58374701 | 0.035742583 | 0.031611827 |
| MGAT5 | BC | old | 0.395173877 | 0.806405196 | 5.02E-10 | 0.005174149 |
| EPHA3 | BC | old | 0.39505833 | 0.755063497 | 5.83E-05 | 0.038847719 |
| DENND4C | BC | old | 0.394105012 | 1.14306717 | 1.77E-19 | 0.000117552 |
| NRG1 | BC | old | 0.393254047 | 1.395020293 | 1.70E-37 | 6.68E-08 |
| CACNA2D1 | BC | old | 0.391940767 | 1.094279262 | 2.50E-33 | 0.000251117 |
| WNK2 | BC | old | 0.39165329 | 0.992607268 | 8.05E-12 | 0.000350068 |
| USP24 | BC | old | 0.389636551 | 0.922381938 | 0.000262508 | 0.007225683 |
| HELZ | BC | old | 0.389589438 | 0.716465 | 2.14E-06 | 0.007677335 |
| PLPPR5 | BC | old | 0.388706131 | 1.152480445 | 1.41E-05 | 0.000535689 |
| GNB1 | BC | old | 0.387859486 | 0.817682548 | 0.0410773 | 0.031218231 |
| SH3BP5 | BC | old | 0.386838176 | 0.952655671 | 6.07E-05 | 0.005932971 |
| PAG1 | BC | old | 0.38636564 | 0.800862091 | 1.62E-05 | 0.011821854 |
| PRICKLE1 | BC | old | 0.384129598 | 0.998118793 | 1.13E-06 | 0.003987119 |
| LRP8 | BC | old | 0.383692053 | 0.787772453 | 0.001746461 | 0.040782162 |
| FAM126B | BC | old | 0.383489745 | 0.793338263 | 2.96E-10 | 0.006868835 |
| USP34 | BC | old | 0.38190598 | 0.707479018 | 4.31E-15 | 0.002815116 |
| RAPGEF2 | BC | old | 0.379042045 | 1.037564567 | 2.77E-09 | 0.000371148 |
| TPST1 | BC | old | 0.378683655 | 1.026447994 | 4.63E-08 | 0.000813939 |
| FAM66D | BC | old | 0.378674643 | 1.110536805 | 0.00815094 | 0.006267341 |
| PLCL2 | BC | old | 0.378555235 | 1.143005319 | 0.002572083 | 0.001195906 |
| SLC24A2 | BC | old | 0.378550958 | 0.946213029 | 4.55E-20 | 0.005872857 |
| POM121C | BC | old | 0.377261648 | 0.760059318 | 0.000237645 | 0.042478281 |
| MAST4 | BC | old | 0.376219656 | 1.467026773 | 1.22E-11 | 4.88E-06 |
| XKR4 | BC | old | 0.375242832 | 1.714154726 | 7.50E-12 | 3.32E-08 |
| GPCPD1 | BC | old | 0.374745401 | 1.230556148 | 5.60E-26 | 1.90E-05 |
| AC106897.1 | BC | old | 0.372316145 | 1.162117934 | 5.81E-05 | 0.002856248 |
| MAP4K3 | BC | old | 0.370596293 | 0.761101457 | 6.75E-10 | 0.006709816 |
| FEZF2 | BC | old | 0.369965285 | 1.518614381 | 9.98E-13 | 5.50E-07 |
| DGKD | BC | old | 0.369871869 | 1.335452384 | 1.97E-15 | 9.24E-06 |

| | | | | | | |
|---|---|---|---|---|---|---|
| MYO9A | BC | old | 0.369847147 | 0.83776984 | 1.84E-10 | 0.000760477 |
| YBX3 | BC | old | 0.369452457 | 1.028860872 | 0.000133046 | 0.008475616 |
| SORBS2 | BC | old | 0.369016288 | 1.21996175 | 2.69E-23 | 6.20E-07 |
| MYO16 | BC | old | 0.36874458 | 1.36838573 | 0.011737432 | 0.000210417 |
| HERC1 | BC | old | 0.368478732 | 0.651605027 | 0.002399213 | 0.01180245 |
| MAP1B | BC | old | 0.36817773 | 0.633222252 | 0.005704423 | 0.023683395 |
| ZNF536 | BC | old | 0.366105428 | 1.231713385 | 1.72E-27 | 5.51E-06 |
| WAC | BC | old | 0.365667399 | 0.589241374 | 3.33E-09 | 0.016715443 |
| CAMK4 | BC | old | 0.365024646 | 1.1966402 | 7.07E-29 | 6.74E-06 |
| ARHGAP12 | BC | old | 0.362787274 | 0.879810244 | 0.001065585 | 0.028410132 |
| ARHGEF10L | BC | old | 0.362739889 | 0.872358549 | 4.78E-05 | 0.021019888 |
| DDIT4 | BC | old | 0.362700432 | 0.980382521 | 1.88E-11 | 0.000874953 |
| ZNF532 | BC | old | 0.360691279 | 0.731250777 | 0.000194865 | 0.010782418 |
| IQSEC1 | BC | old | 0.360624768 | 0.641229963 | 8.05E-05 | 0.033712186 |
| EEPD1 | BC | old | 0.360038309 | 1.284602543 | 6.32E-09 | 0.000582606 |
| FADS2 | BC | old | 0.35987018 | 0.983671987 | 0.000371044 | 0.014557716 |
| L1CAM | BC | old | 0.359352995 | 0.849941125 | 0.001405727 | 0.028870195 |
| PDZRN3 | BC | old | 0.358953276 | 0.898401549 | 3.45E-05 | 0.020381222 |
| MYO6 | BC | old | 0.358690453 | 0.695996068 | 0.000105694 | 0.014179896 |
| IGSF3 | BC | old | 0.358492506 | 1.461876997 | 2.53E-08 | 1.42E-05 |
| HIVEP3 | BC | old | 0.35830942 | 0.875230476 | 7.90E-08 | 0.010149082 |
| ADARB1 | BC | old | 0.358249161 | 0.738428852 | 5.90E-07 | 0.015799624 |
| TRIM9 | BC | old | 0.358218257 | 0.805648766 | 0.033045845 | 0.041764577 |
| ATP2C1 | BC | old | 0.355914077 | 0.99747884 | 1.03E-11 | 0.000120711 |
| PGAP1 | BC | old | 0.354973552 | 0.812059678 | 1.91E-05 | 0.009358763 |
| ADAM9 | BC | old | 0.354070042 | 1.178744611 | 2.52E-10 | 6.00E-05 |
| DNER | BC | old | 0.353626926 | 0.717706009 | 0.002337312 | 0.023096391 |
| ELOVL5 | BC | old | 0.353537532 | 1.103901569 | 4.31E-13 | 0.001320179 |
| MTF2 | BC | old | 0.351488774 | 0.771117578 | 0.00210188 | 0.018293939 |
| FAM171A1 | BC | old | 0.351273722 | 0.942110061 | 0.001389944 | 0.02139855 |
| HMBOX1 | BC | old | 0.350942877 | 0.672807253 | 0.020587794 | 0.040422057 |
| GPRIN3 | BC | old | 0.348982275 | 1.296578619 | 6.18E-05 | 0.000650302 |
| PDS5B | BC | old | 0.348122235 | 0.795295873 | 1.24E-09 | 0.002178158 |
| CCDC88A | BC | old | 0.343337392 | 0.549823161 | 4.18E-05 | 0.033126118 |
| CYGB | BC | old | 0.34113896 | 0.860406139 | 0.000353626 | 0.013486832 |
| LGR4 | BC | old | 0.340468048 | 1.216987236 | 1.07E-10 | 0.000151396 |
| AL392023.2 | BC | old | 0.339544168 | 0.890159845 | 0.001625677 | 0.004050937 |
| PRKCB | BC | old | 0.339451395 | 1.281781719 | 1.92E-27 | 4.35E-05 |
| SNCB | BC | old | 0.338990162 | 0.809719831 | 3.46E-12 | 0.012629265 |
| SLC7A11 | BC | old | 0.335613142 | 1.008061042 | 0.018171192 | 0.008857769 |
| MAPT | BC | old | 0.335315108 | 0.523627922 | 1.75E-05 | 0.039442776 |
| CERS6 | BC | old | 0.334373395 | 0.827003119 | 0.000724645 | 0.003445836 |
| SGCD | BC | old | 0.332206769 | 1.203464461 | 1.69E-11 | 0.000361079 |
| SIPA1L2 | BC | old | 0.331936992 | 1.139399346 | 2.83E-09 | 0.000220372 |
| TANC2 | BC | old | 0.331493159 | 0.984751928 | 1.08E-23 | 0.000398607 |
| UBE2H | BC | old | 0.331232962 | 0.57177868 | 0.004113108 | 0.032170607 |
| NIM1K | BC | old | 0.331067754 | 0.870043317 | 0.004247732 | 0.034137129 |
| PTCH1 | BC | old | 0.329759358 | 0.748677698 | 0.040528003 | 0.049744617 |
| SCN3A | BC | old | 0.329689062 | 1.404759687 | 6.48E-14 | 0.000258004 |
| C3orf70 | BC | old | 0.329570074 | 1.175461027 | 8.00E-12 | 0.000138096 |
| CAMKK1 | BC | old | 0.326596634 | 0.901390135 | 1.02E-05 | 0.010782418 |
| GRM1 | BC | old | 0.326562594 | 0.906957609 | 0.026255949 | 0.015790617 |
| ABR | BC | old | 0.326553981 | 0.731097026 | 0.000446509 | 0.019094084 |
| WDR41 | BC | old | 0.325994668 | 0.599372767 | 0.006387419 | 0.041289946 |

| | | | | | | |
|---|---|---|---|---|---|---|
| HS3ST2 | BC | old | 0.32426052 | 1.074825468 | 0.025271813 | 0.001634157 |
| EPB41L5 | BC | old | 0.324194667 | 0.830938875 | 3.18E-09 | 0.003006852 |
| ATAD2B | BC | old | 0.323327486 | 0.857952428 | 0.00051955 | 0.003726714 |
| TMTC1 | BC | old | 0.322358058 | 1.435147618 | 7.10E-08 | 0.000117583 |
| REPS2 | BC | old | 0.322322737 | 1.54436815 | 5.58E-31 | 2.08E-07 |
| SLC6A6 | BC | old | 0.320096729 | 1.338786395 | 1.87E-32 | 3.32E-08 |
| CRMP1 | BC | old | 0.318062694 | 0.850340439 | 4.91E-09 | 0.000610444 |
| PEAK1 | BC | old | 0.317809236 | 1.119492469 | 2.92E-11 | 0.000150442 |
| SLC24A3 | BC | old | 0.315532562 | 1.496923649 | 5.44E-11 | 8.17E-07 |
| ARNTL | BC | old | 0.314754728 | 0.790353981 | 0.00248365 | 0.014025801 |
| DLG1 | BC | old | 0.313506084 | 0.801496769 | 1.81E-06 | 0.008899849 |
| NCKAP1 | BC | old | 0.313307436 | 0.713414241 | 2.04E-08 | 0.007631428 |
| VGLL4 | BC | old | 0.313058168 | 0.857290557 | 2.46E-06 | 0.004835209 |
| ACKR1 | BC | old | 0.312856978 | 0.864826682 | 0.006752277 | 0.018900184 |
| ST8SIA1 | BC | old | 0.312382084 | 1.126626221 | 4.14E-05 | 0.001457969 |
| PSME4 | BC | old | 0.31188236 | 0.777395675 | 0.030739352 | 0.024717717 |
| WSB1 | BC | old | 0.31161377 | 0.905181317 | 3.95E-14 | 0.000775252 |
| SLC2A13 | BC | old | 0.30886638 | 0.977892843 | 5.26E-05 | 0.010899232 |
| ADGRB1 | BC | old | 0.308620342 | 1.054880919 | 1.13E-09 | 0.019307489 |
| DIPK2A | BC | old | 0.308276527 | 0.834701807 | 0.000621836 | 0.026039447 |
| HSPA6 | BC | old | 0.305695688 | 1.23750705 | 6.24E-37 | 1.04E-05 |
| PREPL | BC | old | 0.305129501 | 0.57956492 | 2.29E-09 | 0.020012914 |
| STK35 | BC | old | 0.305049419 | 0.954403763 | 5.18E-19 | 0.000210417 |
| SCAI | BC | old | 0.304742187 | 0.874616498 | 3.74E-06 | 0.004213865 |
| NBEAL1 | BC | old | 0.30383789 | 0.737650168 | 0.002615217 | 0.026468633 |
| ARHGAP32 | BC | old | 0.303774056 | 0.647603227 | 0.007589711 | 0.039752416 |
| SLC1A2 | BC | old | 0.303212627 | 0.745273375 | 0.00034527 | 0.00318021 |
| LYST | BC | old | 0.302419223 | 0.906495206 | 1.38E-14 | 0.00040775 |
| ASCC3 | BC | old | 0.301900085 | 0.674830706 | 7.03E-06 | 0.021651471 |
| PRRC2B | BC | old | 0.30129238 | 0.621831149 | 2.14E-09 | 0.022964018 |
| ABHD3 | BC | old | 0.300582032 | 1.878014585 | 1.22E-19 | 2.26E-09 |
| OSBPL6 | BC | old | 0.300491323 | 0.649280814 | 0.003477991 | 0.009539423 |
| HOOK1 | BC | old | 0.300420985 | 0.908324883 | 1.36E-19 | 0.000232994 |
| ERBIN | BC | old | 0.299926074 | 1.055120842 | 0.000509243 | 0.004273384 |
| NRIP1 | BC | old | 0.298594814 | 0.866468975 | 8.92E-08 | 0.005349975 |
| GRIK1 | BC | old | 0.298024525 | 1.150328171 | 2.45E-25 | 3.46E-05 |
| AC007389.1 | BC | old | 0.297679613 | 1.679188032 | 1.18E-11 | 2.00E-08 |
| DOCK9 | BC | old | 0.297492521 | 1.202084851 | 2.60E-19 | 2.43E-05 |
| HOMER1 | BC | old | 0.294052011 | 0.941641264 | 2.57E-06 | 0.003750594 |
| PHLPP2 | BC | old | 0.292704085 | 0.764449004 | 0.000755464 | 0.014529461 |
| NFIX | BC | old | 0.292505835 | 1.044582962 | 0.000937301 | 0.011018108 |
| ARHGAP44 | BC | old | 0.2914895 | 0.950522617 | 0.015426476 | 0.02037768 |
| PRICKLE2 | BC | old | 0.290417663 | 0.765008503 | 3.46E-07 | 0.004018948 |
| ADD2 | BC | old | 0.289008578 | 0.839788904 | 1.80E-06 | 0.019469289 |
| IDS | BC | old | 0.288187546 | 0.629894365 | 1.31E-08 | 0.01228984 |
| FAM234B | BC | old | 0.288091882 | 1.160225682 | 1.19E-11 | 0.000441181 |
| FSD1L | BC | old | 0.287382603 | 0.969082626 | 2.52E-06 | 0.002451106 |
| DACH2 | BC | old | 0.286911954 | 1.251652142 | 0.039186123 | 4.54E-05 |
| IRF2BPL | BC | old | 0.285343076 | 0.918670608 | 1.48E-09 | 0.002500475 |
| ARHGAP29 | BC | old | 0.28468024 | 1.256844032 | 8.77E-08 | 0.000117583 |
| TNRC6C | BC | old | 0.284527116 | 0.632755417 | 1.20E-06 | 0.014284876 |
| CDK17 | BC | old | 0.283536472 | 0.745146324 | 0.000219278 | 0.049065413 |
| PHLPP1 | BC | old | 0.28179163 | 1.299436285 | 8.04E-20 | 2.27E-06 |
| PLEKHD1 | BC | old | 0.279670529 | 0.765515437 | 0.013237866 | 0.020012914 |

| | | | | | | |
|---|---|---|---|---|---|---|
| CRYBG3 | BC | old | 0.279544502 | 1.131200374 | 1.62E-34 | 2.10E-06 |
| PER2 | BC | old | 0.279221837 | 1.092618001 | 1.55E-11 | 0.003152381 |
| JARID2 | BC | old | 0.279142919 | 0.755867837 | 1.48E-13 | 0.001770058 |
| PPP1CB | BC | old | 0.278660114 | 0.702657142 | 1.18E-12 | 0.004131457 |
| TLK1 | BC | old | 0.278620504 | 0.918757312 | 8.01E-13 | 0.002855305 |
| ANK2 | BC | old | 0.278161797 | 0.770724437 | 3.12E-31 | 0.000644071 |
| MARK3 | BC | old | 0.277099064 | 0.656466136 | 0.001642145 | 0.023195151 |
| ASPH | BC | old | 0.276867268 | 0.513356603 | 6.04E-06 | 0.047732387 |
| SPIN1 | BC | old | 0.275802646 | 0.730356076 | 4.59E-07 | 0.008404551 |
| HECTD2 | BC | old | 0.275028169 | 1.225480963 | 9.22E-19 | 2.80E-05 |
| MYO7A | BC | old | 0.275009741 | 0.974227792 | 0.003714568 | 0.027083976 |
| ESRRG | BC | old | 0.274945034 | 1.053216417 | 3.11E-09 | 0.001507509 |
| AGAP3 | BC | old | 0.272422668 | 0.811976302 | 0.000109143 | 0.023683897 |
| FNIP1 | BC | old | 0.270808984 | 0.710547643 | 0.000763205 | 0.021450479 |
| LBH | BC | old | 0.270460835 | 1.112153601 | 4.24E-08 | 0.000488503 |
| KIF5C | BC | old | 0.270081969 | 0.574537081 | 1.24E-06 | 0.032994896 |
| GAB1 | BC | old | 0.269862613 | 1.053345473 | 8.38E-06 | 0.005870248 |
| BMPR1A | BC | old | 0.268255527 | 0.660313794 | 0.022815582 | 0.047663708 |
| PPFIA2 | BC | old | 0.267289634 | 0.578160664 | 1.12E-15 | 0.030345113 |
| MAPRE2 | BC | old | 0.267169956 | 0.733723909 | 1.26E-19 | 0.000401056 |
| SH3GL2 | BC | old | 0.265118123 | 0.752628713 | 5.74E-19 | 0.000634085 |
| ZFR | BC | old | 0.262774878 | 0.679738447 | 1.62E-06 | 0.017556626 |
| MPRIP-AS1 | BC | old | 0.261203726 | 0.744484144 | 0.047712773 | 0.038068178 |
| ZDHHC2 | BC | old | 0.260979803 | 0.524978048 | 5.63E-06 | 0.0437833 |
| ZFHX4 | BC | old | 0.260723028 | 0.862053119 | 6.07E-07 | 0.026166746 |
| DNAJA4 | BC | old | 0.255869789 | 1.15157004 | 3.58E-09 | 0.00071698 |
| KCNH8 | BC | old | 0.254274872 | 1.004088537 | 0.004733068 | 0.00413318 |
| DOCK10 | BC | old | 0.252126466 | 1.09913628 | 0.015161029 | 0.001527204 |
| GNG2 | BC | old | 0.251439585 | 1.180596435 | 4.59E-08 | 0.000336495 |
| TNRC6A | BC | old | 0.248041088 | 0.677419746 | 1.02E-11 | 0.003040775 |
| TMEM178B | BC | old | 0.247467822 | 0.752715413 | 3.41E-05 | 0.004171898 |
| ARL4C | BC | old | 0.247162884 | 1.081638885 | 9.92E-10 | 0.000350068 |
| SLC7A5 | BC | old | 0.24706313 | 0.925199645 | 0.000650201 | 0.026293225 |
| PCBP4 | BC | old | 0.246365537 | 0.629442971 | 5.51E-11 | 0.008322991 |
| SLC44A2 | BC | old | 0.244759828 | 0.926233138 | 7.05E-08 | 0.012089232 |
| GPSM1 | BC | old | 0.243121622 | 0.939436904 | 2.87E-06 | 0.012512945 |
| ZPBP | BC | old | 0.241857509 | 0.834183182 | 1.54E-12 | 0.002967754 |
| TRIM14 | BC | old | 0.241678934 | 1.265545415 | 0.001170646 | 0.001016429 |
| MTUS1 | BC | old | 0.235471482 | 0.941881262 | 2.92E-20 | 1.08E-05 |
| SLC44A1 | BC | old | 0.23505492 | 0.983704024 | 0.000929906 | 0.009346911 |
| B3GLCT | BC | old | 0.235048426 | 2.049971027 | 2.66E-26 | 8.66E-13 |
| KCNK3 | BC | old | 0.231927804 | 1.18589485 | 9.77E-06 | 0.000535689 |
| RALGAPA1 | BC | old | 0.229490735 | 0.565947732 | 0.001000663 | 0.021964435 |
| DSCAM | BC | old | 0.228921245 | 1.05076855 | 1.48E-06 | 0.00014382 |
| SLC4A7 | BC | old | 0.226536863 | 1.167402025 | 7.66E-13 | 0.00032656 |
| GUCY1B1 | BC | old | 0.225382229 | 0.623414605 | 0.000978832 | 0.015957263 |
| JUND | BC | old | 0.225338434 | 0.727535101 | 0.000763001 | 0.003066056 |
| CACNA1G | BC | old | 0.225109789 | 0.830273279 | 2.96E-07 | 0.00596251 |
| C11orf96 | BC | old | 0.222516812 | 1.103762137 | 9.04E-08 | 0.000801438 |
| TGFBR1 | BC | old | 0.222136473 | 0.825398504 | 0.035996377 | 0.029271903 |
| OSBPL8 | BC | old | 0.217549536 | 1.054946169 | 4.25E-18 | 4.73E-05 |
| CRY1 | BC | old | 0.217420489 | 0.893885159 | 4.61E-07 | 0.001650055 |
| LSS | BC | old | 0.216710564 | 0.844722721 | 0.024467325 | 0.048944496 |
| NKRF | BC | old | 0.215606252 | 1.084795056 | 7.81E-07 | 0.003025361 |

| | | | | | | |
|---|---|---|---|---|---|---|
| TAFA4 | BC | old | 0.215521275 | 1.405591911 | 7.11E-14 | 1.67E-07 |
| PDE1C | BC | old | 0.210923614 | 1.09725435 | 2.17E-13 | 1.28E-06 |
| CENPC | BC | old | 0.2070387 | 0.658039356 | 0.000250183 | 0.022967086 |
| SCN1A | BC | old | 0.204663442 | 0.736702112 | 1.74E-05 | 0.029173545 |
| AP3B1 | BC | old | 0.203813463 | 0.857872497 | 0.000143835 | 0.004564947 |
| TRAK1 | BC | old | 0.200357832 | 0.806425864 | 0.000314123 | 0.013249496 |
| CDC14A | BC | old | 0.19833193 | 1.100272338 | 3.43E-07 | 0.00062281 |
| UBC | BC | old | 0.195061601 | 1.092574749 | 9.31E-05 | 5.24E-07 |
| MLLT1 | BC | old | 0.193555825 | 0.740291329 | 0.001933124 | 0.044742836 |
| SREBF2 | BC | old | 0.193483601 | 1.951598743 | 1.80E-50 | 2.38E-11 |
| WDR17 | BC | old | 0.191896627 | 0.647057329 | 0.000108048 | 0.039611935 |
| ABTB2 | BC | old | 0.189539236 | 0.839518344 | 0.000400337 | 0.034567073 |
| NUDT4 | BC | old | 0.188927721 | 0.921191557 | 0.000109265 | 0.003353623 |
| ATP11B | BC | old | 0.188122321 | 0.932575508 | 1.39E-12 | 0.002087004 |
| CTBP2 | BC | old | 0.185958369 | 0.547300914 | 2.26E-07 | 0.039170473 |
| HECTD4 | BC | old | 0.184407047 | 0.664108081 | 2.61E-05 | 0.020836455 |
| RIMS1 | BC | old | 0.178175235 | 0.692803715 | 1.67E-08 | 0.009313265 |
| DNAJC5 | BC | old | 0.173160437 | 0.860964201 | 3.79E-08 | 0.006448942 |
| HLF | BC | old | 0.171988058 | 0.624373491 | 1.36E-07 | 0.005175485 |
| SLITRK6 | BC | old | 0.1695896 | 1.317310216 | 1.06E-10 | 6.35E-05 |
| NOL4L | BC | old | 0.168417703 | 0.716919862 | 7.36E-05 | 0.040758533 |
| GALNT13 | BC | old | 0.166433893 | 0.75707183 | 0.000653749 | 0.0065948 |
| DUSP8 | BC | old | 0.16461167 | 0.768806146 | 1.49E-06 | 0.010811731 |
| HCN1 | BC | old | 0.163618726 | 0.97608334 | 1.42E-08 | 0.000868388 |
| TENM1 | BC | old | 0.161568347 | 0.939196458 | 0.009379436 | 0.010285904 |
| PCLO | BC | old | 0.16101862 | 0.559522867 | 3.83E-11 | 0.030794696 |
| HMX1 | BC | old | 0.155794737 | 1.348395373 | 1.36E-08 | 6.35E-05 |
| LRFN5 | BC | old | 0.153401591 | 1.375217101 | 1.54E-07 | 0.000293693 |
| HSPA4L | BC | old | 0.151390784 | 1.577161521 | 3.41E-49 | 6.63E-11 |
| BCO2 | BC | old | 0.148425585 | 1.447589847 | 3.24E-13 | 6.39E-05 |
| LINC01896 | BC | old | 0.148396805 | 1.46585412 | 1.73E-05 | 3.89E-06 |
| PRMT9 | BC | old | 0.146338032 | 0.847444899 | 0.013511118 | 0.016856042 |
| STXBP1 | BC | old | 0.144899876 | 0.717593408 | 4.86E-11 | 0.001923265 |
| DST | BC | old | 0.142111681 | 0.727996711 | 1.80E-35 | 0.001871931 |
| FGD4 | BC | old | 0.137798213 | 1.043486045 | 3.91E-15 | 0.000973645 |
| OXR1 | BC | old | 0.136211233 | 0.768789945 | 4.63E-15 | 0.008494616 |
| SPATS2L | BC | old | 0.135434037 | 1.175494362 | 2.92E-07 | 0.000648058 |
| AAK1 | BC | old | 0.135300802 | 0.785357396 | 5.30E-12 | 0.000764414 |
| RHOBTB1 | BC | old | 0.123267195 | 0.977207186 | 0.006215112 | 0.00498985 |
| RNMT | BC | old | 0.122474468 | 0.520970545 | 0.000758135 | 0.049475015 |
| HIPK1 | BC | old | 0.12068474 | 0.648010551 | 0.00209777 | 0.034777414 |
| MBP | BC | old | 0.120618283 | 0.694282812 | 8.38E-05 | 0.011096755 |
| LRRC8B | BC | old | 0.118628658 | 0.830942376 | 0.006127567 | 0.012110564 |
| HSPA1A | BC | old | 0.116946939 | 2.812202052 | 2.72E-157 | 6.51E-26 |
| CLK1 | BC | old | 0.112545457 | 1.191479162 | 8.95E-16 | 5.42E-06 |
| CNTN1 | BC | old | 0.112011311 | 0.804064514 | 3.05E-13 | 0.000252093 |
| AMPD2 | BC | old | 0.106523984 | 1.181214946 | 1.40E-12 | 0.00058093 |
| FAM78A | BC | old | 0.103995697 | 0.830458355 | 0.003304762 | 0.008226183 |
| P2RY1 | BC | old | 0.101499393 | 0.939828617 | 0.000661216 | 0.008172315 |
| STARD4 | BC | old | 0.100408173 | 1.766284306 | 1.26E-39 | 2.66E-08 |
| MMADHC | BC | young | -0.108393453 | -0.732966696 | 9.96E-13 | 0.041826779 |
| BANF1 | BC | young | -0.114852833 | -0.704226587 | 3.77E-25 | 0.021214611 |
| HNRNPA3 | BC | young | -0.115207668 | -0.765296584 | 4.06E-39 | 0.000336495 |
| NSRP1 | BC | young | -0.116996884 | -0.767407765 | 2.65E-16 | 0.020085911 |

| | | | | | | |
|---|---|---|---|---|---|---|
| HNRNPM | BC | young | -0.117982966 | -0.884098873 | 1.64E-24 | 0.00032656 |
| KLHDC2 | BC | young | -0.119351769 | -0.713316365 | 1.56E-13 | 0.041799412 |
| COMMD10 | BC | young | -0.119632975 | -0.817399805 | 8.18E-22 | 0.001814231 |
| CLTB | BC | young | -0.123105241 | -0.721340584 | 2.88E-21 | 0.002968822 |
| BTF3 | BC | young | -0.126312642 | -0.70703814 | 1.29E-32 | 0.004395024 |
| YWHAE | BC | young | -0.128419136 | -0.756872739 | 2.01E-39 | 0.001507509 |
| PDZRN4 | BC | young | -0.133567105 | -1.116543868 | 2.36E-38 | 3.64E-05 |
| RPA2 | BC | young | -0.135285698 | -0.916672051 | 3.83E-17 | 0.01176303 |
| PSMB1 | BC | young | -0.136387903 | -0.902114901 | 2.56E-37 | 0.000179478 |
| CALM2 | BC | young | -0.140210362 | -1.041430047 | 1.57E-58 | 3.78E-08 |
| SRP9 | BC | young | -0.141325523 | -0.697114557 | 3.92E-18 | 0.015055429 |
| RPS28 | BC | young | -0.14299141 | -0.583836349 | 3.46E-19 | 0.028007327 |
| RPL18 | BC | young | -0.143128001 | -0.533830549 | 1.05E-21 | 0.049485573 |
| FUS | BC | young | -0.145320763 | -1.079468876 | 1.21E-32 | 1.33E-05 |
| ERH | BC | young | -0.151064754 | -0.925182411 | 1.36E-44 | 0.000163797 |
| PITHD1 | BC | young | -0.151347572 | -0.843574926 | 1.86E-13 | 0.012671652 |
| LYRM4 | BC | young | -0.151853836 | -0.879600885 | 4.51E-14 | 0.008989303 |
| SKP1 | BC | young | -0.152203481 | -0.740782384 | 1.00E-28 | 0.000634283 |
| TMED2 | BC | young | -0.153528735 | -0.878403969 | 1.62E-12 | 0.017530794 |
| GRM5 | BC | young | -0.15527822 | -0.855409448 | 2.70E-48 | 0.000496845 |
| RASGEF1B | BC | young | -0.157596593 | -1.050460453 | 2.46E-33 | 1.50E-05 |
| DGCR8 | BC | young | -0.157972822 | -0.900523789 | 2.95E-06 | 0.036246836 |
| ATP5PB | BC | young | -0.158487748 | -0.87705303 | 8.67E-27 | 0.002564088 |
| EMC4 | BC | young | -0.16040206 | -0.796780544 | 1.94E-15 | 0.023515031 |
| ATP6V1E1 | BC | young | -0.160839899 | -0.947044795 | 2.13E-22 | 0.002975228 |
| STOML2 | BC | young | -0.16346908 | -1.12454581 | 2.65E-19 | 0.001049814 |
| PDCD4 | BC | young | -0.163483903 | -0.678767179 | 1.57E-25 | 0.005811595 |
| AC010478.1 | BC | young | -0.16443443 | -1.092308161 | 6.51E-52 | 1.18E-05 |
| KCNH5 | BC | young | -0.166057516 | -1.005955241 | 3.96E-61 | 4.59E-05 |
| LAMTOR1 | BC | young | -0.166602438 | -0.798323025 | 8.18E-21 | 0.021942222 |
| STX3 | BC | young | -0.166841046 | -0.580328283 | 9.43E-17 | 0.014687934 |
| PSMB3 | BC | young | -0.167802508 | -1.017300898 | 9.53E-37 | 0.000117583 |
| CD27-AS1 | BC | young | -0.167911131 | -0.81356183 | 8.20E-13 | 0.02037768 |
| MED4 | BC | young | -0.169025236 | -0.822593264 | 4.08E-20 | 0.007875325 |
| PSMD6 | BC | young | -0.170015608 | -0.842987306 | 2.96E-13 | 0.031774049 |
| FN3KRP | BC | young | -0.171772545 | -0.855164583 | 4.36E-06 | 0.038312335 |
| YBX1 | BC | young | -0.173268688 | -0.927164749 | 2.01E-52 | 1.81E-05 |
| SNRPN | BC | young | -0.174785157 | -0.609587777 | 9.34E-15 | 0.042933805 |
| PARP1 | BC | young | -0.176204615 | -1.106657014 | 2.03E-27 | 0.00090509 |
| PUF60 | BC | young | -0.176980735 | -0.864402646 | 1.74E-09 | 0.028170567 |
| RANBP1 | BC | young | -0.179404362 | -1.050552536 | 2.03E-30 | 0.000191172 |
| ABCA1 | BC | young | -0.179888317 | -0.932947042 | 4.23E-13 | 0.026166746 |
| RTCB | BC | young | -0.180020334 | -0.780831001 | 3.19E-10 | 0.028870195 |
| ALYREF | BC | young | -0.181073382 | -0.88825407 | 1.50E-08 | 0.03438519 |
| RBX1 | BC | young | -0.182696167 | -0.623354499 | 1.31E-16 | 0.038334526 |
| ARPC1A | BC | young | -0.184145054 | -0.786746495 | 1.57E-11 | 0.023367332 |
| MGAT4C | BC | young | -0.184458371 | -0.943563364 | 4.74E-25 | 0.002933594 |
| TSR2 | BC | young | -0.184615599 | -0.954658662 | 5.28E-26 | 0.00332387 |
| CDH12 | BC | young | -0.187715507 | -0.906946124 | 2.91E-75 | 0.003725148 |
| DNAJC17 | BC | young | -0.188154266 | -0.886226511 | 2.22E-08 | 0.029721631 |
| TAX1BP1 | BC | young | -0.188267978 | -0.687588227 | 3.00E-37 | 0.035061695 |
| SDHB | BC | young | -0.188407815 | -0.701480381 | 3.19E-10 | 0.049187569 |
| CNTN5 | BC | young | -0.188744276 | -1.104779264 | 2.35E-56 | 0.000115339 |
| RHOA | BC | young | -0.190667282 | -0.707531467 | 1.16E-18 | 0.02037768 |

| | | | | | | |
|---|---|---|---|---|---|---|
| OAZ1 | BC | young | -0.191276745 | -0.630904002 | 2.35E-28 | 0.015792052 |
| RNH1 | BC | young | -0.193685724 | -0.853572113 | 2.06E-18 | 0.013366188 |
| TMEM176B | BC | young | -0.194983103 | -0.718922974 | 4.32E-20 | 0.002464593 |
| TXN2 | BC | young | -0.195717514 | -0.72814801 | 7.82E-15 | 0.030414434 |
| HMGN3 | BC | young | -0.196537731 | -0.672142273 | 2.31E-24 | 0.013266506 |
| TMEM176A | BC | young | -0.201006732 | -0.789249444 | 2.97E-21 | 0.017695627 |
| CCT3 | BC | young | -0.204052488 | -0.927827637 | 1.23E-31 | 0.002429219 |
| ANAPC13 | BC | young | -0.204629902 | -0.743609116 | 8.05E-10 | 0.039383866 |
| SPTSSA | BC | young | -0.207354978 | -0.72360115 | 1.49E-11 | 0.03942677 |
| GET1 | BC | young | -0.20971602 | -0.860604539 | 1.62E-23 | 0.004008681 |
| SUMO2 | BC | young | -0.209791456 | -0.914689898 | 6.67E-46 | 7.44E-06 |
| DNAJA2 | BC | young | -0.214866621 | -0.795579693 | 1.44E-20 | 0.014249129 |
| PIGH | BC | young | -0.215687404 | -0.772173276 | 4.53E-10 | 0.038755152 |
| GNB3 | BC | young | -0.21589044 | -1.117405359 | 3.97E-63 | 1.22E-05 |
| NLGN4X | BC | young | -0.221187061 | -0.934267293 | 2.04E-37 | 0.000667753 |
| VSTM2B | BC | young | -0.225195015 | -1.104593792 | 3.17E-32 | 0.000710702 |
| SNHG32 | BC | young | -0.225819314 | -0.739677242 | 8.39E-16 | 0.020381222 |
| GABARAPL2 | BC | young | -0.227730699 | -0.608559389 | 2.23E-27 | 0.020940331 |
| DDC | BC | young | -0.227803223 | -1.461790729 | 3.29E-37 | 2.38E-06 |
| ACP1 | BC | young | -0.230242585 | -0.698218175 | 1.03E-21 | 0.037247304 |
| NCBP2 | BC | young | -0.23040771 | -0.90395711 | 1.60E-18 | 0.006778613 |
| AC113383.1 | BC | young | -0.231090751 | -0.837997295 | 4.82E-19 | 0.004131457 |
| MGARP | BC | young | -0.233109017 | -0.811164588 | 1.29E-25 | 0.002063178 |
| MGST3 | BC | young | -0.233976874 | -0.755831139 | 6.35E-29 | 0.004407292 |
| CNPY3 | BC | young | -0.236135268 | -0.806863365 | 2.65E-16 | 0.02849206 |
| RTRAF | BC | young | -0.23709137 | -0.618390985 | 2.06E-11 | 0.041459644 |
| CALY | BC | young | -0.238672791 | -0.861550615 | 4.12E-25 | 0.027851543 |
| CCT8 | BC | young | -0.240035699 | -0.675490369 | 2.28E-21 | 0.045596927 |
| OARD1 | BC | young | -0.241102853 | -0.753019737 | 4.73E-13 | 0.015792052 |
| PCMT1 | BC | young | -0.242395333 | -0.696924465 | 3.36E-17 | 0.032810883 |
| ZNF33B | BC | young | -0.242705314 | -0.870700232 | 3.55E-17 | 0.007313882 |
| HDDC2 | BC | young | -0.242748753 | -0.653536915 | 6.11E-09 | 0.040782162 |
| ATP5MG | BC | young | -0.243018168 | -0.562190204 | 2.14E-21 | 0.026957967 |
| REEP5 | BC | young | -0.244783273 | -0.741454645 | 2.75E-19 | 0.008498782 |
| C12orf73 | BC | young | -0.244822507 | -0.906315468 | 7.66E-09 | 0.02037768 |
| MTERF4 | BC | young | -0.246622651 | -0.790607834 | 1.00E-11 | 0.030003307 |
| EAPP | BC | young | -0.247311428 | -0.663586362 | 9.73E-18 | 0.028819182 |
| SARS | BC | young | -0.247781113 | -1.065743163 | 8.29E-22 | 0.000540532 |
| STMP1 | BC | young | -0.248708188 | -0.721585616 | 3.16E-17 | 0.012629265 |
| LINC02275 | BC | young | -0.249197332 | -0.692945559 | 5.78E-23 | 0.038112551 |
| SNAP25-AS1 | BC | young | -0.252791594 | -0.844598928 | 1.23E-48 | 0.000350068 |
| VDAC2 | BC | young | -0.254697697 | -0.765572824 | 1.99E-33 | 0.005257692 |
| TEX264 | BC | young | -0.255150641 | -1.121900087 | 1.94E-10 | 0.007918975 |
| COX7A2L | BC | young | -0.256126422 | -0.655961687 | 3.01E-23 | 0.023949766 |
| TBCB | BC | young | -0.257624865 | -1.156085736 | 2.32E-46 | 2.96E-06 |
| RSRP1 | BC | young | -0.257801391 | -0.831319334 | 1.07E-17 | 0.006397225 |
| HNRNPA1 | BC | young | -0.257974221 | -0.760516445 | 3.12E-25 | 0.007560475 |
| ATRAID | BC | young | -0.258034479 | -0.902118284 | 1.17E-14 | 0.003498905 |
| PPP1R10 | BC | young | -0.261250114 | -0.692337776 | 2.23E-19 | 0.032835313 |
| TMEM14B | BC | young | -0.262083849 | -0.792615815 | 4.23E-31 | 0.00058093 |
| SPCS1 | BC | young | -0.265446107 | -0.994155523 | 1.05E-46 | 1.84E-07 |
| PCP4L1 | BC | young | -0.265958607 | -0.95386003 | 2.07E-24 | 0.000416916 |
| IK | BC | young | -0.266710652 | -0.964983061 | 2.05E-26 | 0.000860337 |
| ELOC | BC | young | -0.268827449 | -0.637664189 | 2.98E-16 | 0.039188347 |

| | | | | | | |
|---|---|---|---|---|---|---|
| PIN4 | BC | young | -0.269506403 | -0.829348637 | 2.08E-14 | 0.017280252 |
| PSMC3 | BC | young | -0.269744244 | -0.979392221 | 8.31E-10 | 0.022675005 |
| TMSB4X | BC | young | -0.270910631 | -1.210592572 | 2.17E-66 | 3.00E-07 |
| PPIA | BC | young | -0.271288364 | -0.776450819 | 1.83E-39 | 0.001985705 |
| OSGEP | BC | young | -0.271489551 | -0.778094908 | 1.51E-08 | 0.044020574 |
| SSU72 | BC | young | -0.274141643 | -0.922196325 | 2.99E-19 | 0.002764556 |
| RPL23 | BC | young | -0.274929709 | -0.645219478 | 4.01E-17 | 0.01018575 |
| TMCO1 | BC | young | -0.275403757 | -0.693100799 | 1.88E-18 | 0.020177168 |
| GTF2B | BC | young | -0.281755733 | -0.893891991 | 3.33E-13 | 0.016560152 |
| UPF3A | BC | young | -0.282466048 | -0.734032465 | 1.56E-16 | 0.016279839 |
| PDAP1 | BC | young | -0.282694833 | -0.746617715 | 6.61E-17 | 0.02910359 |
| DPY30 | BC | young | -0.28427977 | -1.079738709 | 8.42E-21 | 0.001156108 |
| PAIP2 | BC | young | -0.285486128 | -0.84724613 | 3.35E-36 | 0.000243624 |
| GAS7 | BC | young | -0.286335776 | -0.794494725 | 1.61E-13 | 0.017280252 |
| IER3IP1 | BC | young | -0.286757638 | -0.861183493 | 8.02E-23 | 0.002562273 |
| LHFPL3-AS1 | BC | young | -0.287310683 | -0.961753721 | 1.05E-11 | 0.032835313 |
| SUCLA2 | BC | young | -0.287704088 | -0.819433144 | 5.24E-17 | 0.011304366 |
| CCDC12 | BC | young | -0.288388869 | -0.754036577 | 1.66E-10 | 0.043743726 |
| HNRNPA2B1 | BC | young | -0.289071953 | -0.883893022 | 3.94E-55 | 1.14E-06 |
| CFAP298 | BC | young | -0.291308714 | -1.022892871 | 8.29E-16 | 0.005524905 |
| RPL27 | BC | young | -0.292690884 | -0.624776646 | 4.49E-21 | 0.023323315 |
| C16orf74 | BC | young | -0.292926987 | -0.839761787 | 6.11E-33 | 0.02180131 |
| TMEM230 | BC | young | -0.294496113 | -0.769501511 | 6.45E-21 | 0.003025361 |
| CADM2-AS1 | BC | young | -0.294529466 | -1.002165666 | 3.96E-10 | 0.013151231 |
| GNG10 | BC | young | -0.294945679 | -0.904843644 | 6.65E-09 | 0.025156609 |
| PPM1N | BC | young | -0.296714496 | -0.984444222 | 3.25E-10 | 0.040661759 |
| ATP5PO | BC | young | -0.297917398 | -0.677021852 | 3.15E-21 | 0.034419531 |
| PRDX3 | BC | young | -0.298273546 | -0.769411754 | 1.84E-14 | 0.027984856 |
| DSTN | BC | young | -0.299562561 | -0.788053421 | 5.55E-25 | 0.003904924 |
| DEGS1 | BC | young | -0.29959741 | -1.018142364 | 5.93E-13 | 0.013291929 |
| PPP4C | BC | young | -0.300414066 | -0.918638148 | 2.74E-12 | 0.015809151 |
| SRPRA | BC | young | -0.300611101 | -0.948574892 | 4.35E-11 | 0.032810883 |
| PSMB5 | BC | young | -0.30073811 | -1.055997781 | 5.08E-29 | 0.000479775 |
| HIRIP3 | BC | young | -0.305618169 | -1.239471307 | 5.19E-20 | 0.00071739 |
| TXN | BC | young | -0.306383237 | -0.934817063 | 1.77E-35 | 0.00186259 |
| SET | BC | young | -0.306901991 | -1.000297881 | 4.61E-44 | 1.76E-05 |
| SUMO1 | BC | young | -0.306912736 | -0.615798094 | 1.03E-19 | 0.027365394 |
| COX5A | BC | young | -0.306926643 | -0.984846997 | 6.74E-42 | 0.000354504 |
| RPS11 | BC | young | -0.309882908 | -0.724630689 | 5.63E-26 | 0.003110924 |
| AC091946.1 | BC | young | -0.310667946 | -1.028158339 | 8.55E-22 | 0.0037628 |
| HTATSF1 | BC | young | -0.311084068 | -0.900278231 | 1.54E-13 | 0.031432923 |
| PPP1R7 | BC | young | -0.312252618 | -0.787640893 | 3.80E-13 | 0.040758533 |
| ORMDL1 | BC | young | -0.314582842 | -0.777081822 | 7.78E-21 | 0.003278533 |
| AC117944.1 | BC | young | -0.316455603 | -1.338132152 | 3.77E-37 | 9.57E-05 |
| MRPS21 | BC | young | -0.317244493 | -0.849838981 | 4.08E-19 | 0.003900932 |
| PHF5A | BC | young | -0.319039289 | -0.812492347 | 7.44E-10 | 0.045810166 |
| SPCS2 | BC | young | -0.319538 | -0.700609251 | 1.59E-23 | 0.014025801 |
| BECN1 | BC | young | -0.319797697 | -0.977522167 | 7.01E-09 | 0.02281155 |
| VAMP2 | BC | young | -0.321542819 | -0.578680843 | 2.24E-14 | 0.015706528 |
| NME3 | BC | young | -0.322115616 | -0.764619074 | 3.12E-14 | 0.012101308 |
| APEX1 | BC | young | -0.32483018 | -1.163728699 | 4.52E-33 | 3.58E-05 |
| GSTP1 | BC | young | -0.326109998 | -0.759737733 | 1.19E-20 | 0.001732404 |
| TMED9 | BC | young | -0.326754533 | -1.098272002 | 4.32E-23 | 0.001781682 |
| ATP5MC3 | BC | young | -0.326901771 | -0.93755196 | 2.17E-39 | 0.000120711 |

| | | | | | | |
|---|---|---|---|---|---|---|
| TRIR | BC | young | -0.327162845 | -0.92872591 | 1.13E-28 | 0.000710702 |
| SEMA3E | BC | young | -0.327250024 | -1.228719467 | 3.29E-58 | 2.49E-07 |
| CAMK2N1 | BC | young | -0.327486549 | -0.730841172 | 1.21E-20 | 0.009991884 |
| PPM1G | BC | young | -0.327621488 | -0.938433736 | 2.18E-08 | 0.032032536 |
| GNG13 | BC | young | -0.328248973 | -1.171946873 | 4.05E-65 | 8.28E-07 |
| SNRPE | BC | young | -0.32907643 | -0.88513957 | 1.91E-28 | 0.001741821 |
| LAMTOR5 | BC | young | -0.32972878 | -0.649445841 | 3.58E-18 | 0.016297068 |
| CMPK1 | BC | young | -0.330455439 | -1.093609975 | 2.27E-19 | 0.001611629 |
| EMC3 | BC | young | -0.331127032 | -1.009537922 | 8.45E-18 | 0.004358922 |
| CYC1 | BC | young | -0.333509514 | -0.875708307 | 3.42E-19 | 0.009914772 |
| MRPS28 | BC | young | -0.339422913 | -0.950986977 | 6.68E-19 | 0.003547346 |
| AC022469.2 | BC | young | -0.339717211 | -1.248592205 | 2.74E-12 | 0.021769236 |
| ATP6V1G1 | BC | young | -0.339816695 | -0.808620776 | 3.64E-36 | 0.000293693 |
| B3GNT2 | BC | young | -0.341818533 | -0.931003047 | 3.55E-22 | 0.02502784 |
| ERLIN2 | BC | young | -0.342929467 | -0.918414995 | 2.23E-11 | 0.028741875 |
| IFI6 | BC | young | -0.343385561 | -1.272263068 | 4.78E-18 | 0.000271946 |
| WDR83OS | BC | young | -0.343618367 | -0.980607972 | 9.66E-25 | 0.001048787 |
| COA5 | BC | young | -0.345808957 | -0.782558414 | 9.21E-14 | 0.031218231 |
| SELENOF | BC | young | -0.348192794 | -0.749445803 | 1.14E-24 | 0.008475616 |
| CCDC136 | BC | young | -0.348708712 | -1.029835496 | 3.97E-48 | 0.000208398 |
| RAN | BC | young | -0.349529585 | -0.721524041 | 6.43E-24 | 0.006333418 |
| SNRPG | BC | young | -0.349799313 | -0.817413906 | 5.55E-29 | 0.00151112 |
| SNRPD2 | BC | young | -0.352806299 | -0.761947532 | 3.53E-25 | 0.017613045 |
| DCTN3 | BC | young | -0.354517813 | -0.85375005 | 1.94E-25 | 0.003478204 |
| RPS17 | BC | young | -0.355042213 | -1.034363014 | 1.56E-34 | 5.34E-05 |
| SRP14 | BC | young | -0.355108443 | -0.957425407 | 4.11E-53 | 6.05E-06 |
| NUDT16L1 | BC | young | -0.355479966 | -1.072205106 | 1.87E-15 | 0.003833194 |
| ABCA10 | BC | young | -0.355968153 | -0.819736354 | 7.82E-09 | 0.034245555 |
| MPC1 | BC | young | -0.358436344 | -0.619737881 | 2.47E-18 | 0.030307939 |
| SSB | BC | young | -0.35907475 | -0.809130009 | 2.63E-29 | 0.001535357 |
| SCG3 | BC | young | -0.359734374 | -0.718783914 | 1.44E-14 | 0.020277236 |
| HINT1 | BC | young | -0.359797648 | -0.906725675 | 3.49E-42 | 3.58E-05 |
| PFDN2 | BC | young | -0.360098903 | -0.652995199 | 2.73E-18 | 0.030307939 |
| TIMM17A | BC | young | -0.360382149 | -0.884442726 | 7.02E-14 | 0.023594612 |
| AP002370.2 | BC | young | -0.360817608 | -1.22394669 | 2.04E-28 | 0.000208398 |
| PDE5A | BC | young | -0.36249918 | -0.826020907 | 1.30E-22 | 0.006028526 |
| NAA20 | BC | young | -0.362881654 | -1.071496849 | 5.67E-31 | 0.000252093 |
| SNRPF | BC | young | -0.362935521 | -0.734187089 | 2.06E-18 | 0.011504486 |
| SLC25A24 | BC | young | -0.363624578 | -0.926578088 | 1.41E-12 | 0.027494786 |
| AL138799.4 | BC | young | -0.365083251 | -1.496478966 | 1.08E-25 | 6.39E-05 |
| AC011824.2 | BC | young | -0.368100838 | -1.206601369 | 4.84E-24 | 0.001657989 |
| DYNLRB1 | BC | young | -0.368160273 | -0.822826136 | 8.15E-30 | 0.003914105 |
| HAGH | BC | young | -0.368166829 | -0.938074244 | 2.68E-20 | 0.003046977 |
| PSMC4 | BC | young | -0.369334417 | -1.181788723 | 2.19E-29 | 0.000566553 |
| HLTF | BC | young | -0.371965766 | -0.593961636 | 5.40E-17 | 0.038403795 |
| MT-CO3 | BC | young | -0.372771341 | -1.209690487 | 1.79E-103 | 2.86E-07 |
| COX4I1 | BC | young | -0.373722694 | -1.060653292 | 1.05E-72 | 3.60E-05 |
| SOD1 | BC | young | -0.375110795 | -0.755738011 | 2.46E-43 | 0.003415433 |
| TBCA | BC | young | -0.375110891 | -0.803493705 | 2.09E-37 | 0.000188955 |
| AC093159.1 | BC | young | -0.375277685 | -1.097190394 | 8.03E-30 | 0.002667519 |
| CHCHD2 | BC | young | -0.377091977 | -1.047683456 | 3.64E-54 | 3.71E-06 |
| DNAJC8 | BC | young | -0.377107014 | -0.74728363 | 6.39E-23 | 0.01120523 |
| ZNF32 | BC | young | -0.377789399 | -0.831502741 | 1.53E-08 | 0.026293225 |
| COX16 | BC | young | -0.377832643 | -0.766781924 | 4.36E-22 | 0.015528913 |

| | | | | | | |
|---|---|---|---|---|---|---|
| EIF5A | BC | young | -0.378736055 | -1.148755386 | 1.63E-27 | 0.000251117 |
| RNF5 | BC | young | -0.378994924 | -0.863804877 | 1.31E-13 | 0.027083976 |
| NPM1 | BC | young | -0.379440156 | -0.70413534 | 1.36E-28 | 0.012387307 |
| PDRG1 | BC | young | -0.379641719 | -1.017613024 | 1.74E-07 | 0.032892787 |
| PSMA3 | BC | young | -0.379696372 | -0.953897634 | 4.08E-29 | 0.001404295 |
| CCDC144A | BC | young | -0.379717017 | -0.887977316 | 7.25E-22 | 0.032204573 |
| MIS18A | BC | young | -0.379883508 | -1.108283275 | 6.91E-11 | 0.006538521 |
| CYCS | BC | young | -0.380229914 | -0.682266478 | 2.84E-18 | 0.014911774 |
| SUPT4H1 | BC | young | -0.380799298 | -1.004572444 | 2.62E-12 | 0.016279839 |
| ATP5F1C | BC | young | -0.38118426 | -1.04166605 | 7.19E-50 | 3.53E-05 |
| ATP6V0B | BC | young | -0.381505191 | -1.184423364 | 3.51E-59 | 2.66E-08 |
| SPAG7 | BC | young | -0.381791478 | -0.985675295 | 2.25E-19 | 0.008038524 |
| UQCRQ | BC | young | -0.38379854 | -0.695943443 | 1.07E-23 | 0.029869717 |
| PSMD8 | BC | young | -0.383930081 | -0.965325376 | 1.47E-29 | 0.000772051 |
| COMT | BC | young | -0.385331605 | -0.738357823 | 5.17E-15 | 0.034615055 |
| PPIB | BC | young | -0.388068946 | -0.794382916 | 1.47E-17 | 0.04327506 |
| NTNG1 | BC | young | -0.389341337 | -1.175383696 | 1.79E-47 | 2.46E-05 |
| AC119868.2 | BC | young | -0.390645572 | -1.387600755 | 1.94E-81 | 1.47E-09 |
| BORCS7 | BC | young | -0.390927201 | -0.965910019 | 1.10E-23 | 0.001758509 |
| RPH3A | BC | young | -0.391147731 | -1.141699576 | 3.85E-29 | 0.001760972 |
| DMAC1 | BC | young | -0.391165452 | -0.862940506 | 1.15E-12 | 0.02122497 |
| NEDD8 | BC | young | -0.391846389 | -0.913634435 | 8.08E-35 | 0.001107148 |
| SF3B6 | BC | young | -0.392128921 | -0.830484997 | 1.46E-29 | 0.001856516 |
| TOMM6 | BC | young | -0.392831672 | -0.835650918 | 4.56E-20 | 0.00408277 |
| SLC25A4 | BC | young | -0.395164582 | -0.822697554 | 1.89E-26 | 0.000600024 |
| AP000857.2 | BC | young | -0.396265356 | -1.184783406 | 1.13E-36 | 0.000220372 |
| PPIL1 | BC | young | -0.397347682 | -0.957530263 | 4.73E-12 | 0.033383002 |
| TOMM22 | BC | young | -0.397409588 | -1.035335628 | 6.19E-18 | 0.00762866 |
| NAPG | BC | young | -0.399315191 | -0.75229034 | 6.10E-15 | 0.033432118 |
| POMP | BC | young | -0.399484886 | -0.618971346 | 7.59E-15 | 0.026484292 |
| DYNLL2 | BC | young | -0.399645575 | -0.792840266 | 3.28E-13 | 0.018423839 |
| POLR3F | BC | young | -0.400857707 | -1.10563029 | 4.76E-16 | 0.006187664 |
| ESF1 | BC | young | -0.401135988 | -0.759822505 | 2.99E-11 | 0.023393563 |
| PSMA2 | BC | young | -0.40341604 | -1.006653062 | 1.32E-31 | 0.000535689 |
| SERPINI1 | BC | young | -0.404121749 | -1.028550459 | 4.12E-55 | 0.001891735 |
| TRNP1 | BC | young | -0.406654025 | -1.017108776 | 2.08E-57 | 0.002989716 |
| UQCRFS1 | BC | young | -0.407870746 | -1.016496016 | 3.44E-25 | 0.0003945 |
| UBTF | BC | young | -0.410213967 | -0.912823935 | 2.21E-09 | 0.026153747 |
| NORAD | BC | young | -0.410466815 | -0.704782541 | 1.74E-16 | 0.013270283 |
| PEBP1 | BC | young | -0.41099063 | -0.882391159 | 2.66E-44 | 0.000237334 |
| CC2D2A | BC | young | -0.41190153 | -1.042321835 | 1.04E-30 | 0.000566553 |
| SNRPD1 | BC | young | -0.412431808 | -0.853542909 | 2.34E-27 | 0.003831971 |
| AC002463.1 | BC | young | -0.4135492 | -0.789703636 | 4.16E-19 | 0.013614796 |
| AC018695.9 | BC | young | -0.41386142 | -0.950980819 | 1.97E-12 | 0.040335812 |
| ACTR6 | BC | young | -0.414318147 | -0.729669661 | 2.03E-16 | 0.049475015 |
| MT-CO1 | BC | young | -0.41500435 | -0.827223855 | 6.55E-52 | 0.000347667 |
| LINC01266 | BC | young | -0.415197794 | -1.006397108 | 2.49E-18 | 0.001872516 |
| NUTF2 | BC | young | -0.416462939 | -0.857667558 | 4.06E-08 | 0.044742836 |
| VPS29 | BC | young | -0.417800168 | -0.981437123 | 4.39E-29 | 0.0005705 |
| ENO4 | BC | young | -0.420309986 | -0.947271735 | 7.92E-14 | 0.024290959 |
| BUD23 | BC | young | -0.423480574 | -0.988041755 | 2.51E-15 | 0.009185675 |
| PSMA5 | BC | young | -0.424021115 | -1.0308222 | 5.51E-21 | 0.002464593 |
| ABHD14A | BC | young | -0.424420622 | -0.969890195 | 1.87E-25 | 0.001516308 |
| PDCD5 | BC | young | -0.426896069 | -0.773759173 | 4.20E-21 | 0.013270283 |

| | | | | | | |
|---|---|---|---|---|---|---|
| PFDN4 | BC | young | -0.427125171 | -0.968823414 | 2.31E-19 | 0.009267992 |
| COX6A1 | BC | young | -0.427137339 | -0.756925227 | 1.59E-34 | 0.012089232 |
| HMOX2 | BC | young | -0.427144016 | -1.063660441 | 9.96E-11 | 0.019094084 |
| GTF2H5 | BC | young | -0.428506735 | -0.824350696 | 1.79E-21 | 0.007883636 |
| SF3B5 | BC | young | -0.428622809 | -1.231174425 | 2.90E-36 | 6.85E-05 |
| SEC61B | BC | young | -0.428644106 | -0.671515977 | 1.43E-20 | 0.026801465 |
| SRSF9 | BC | young | -0.428853645 | -1.035887083 | 3.20E-42 | 5.04E-05 |
| NDUFV3 | BC | young | -0.429519657 | -0.790864783 | 3.13E-16 | 0.02753026 |
| C1QBP | BC | young | -0.430036023 | -1.313539027 | 7.03E-48 | 2.54E-05 |
| MYL12B | BC | young | -0.431865778 | -1.015646218 | 2.08E-49 | 0.000418351 |
| SYF2 | BC | young | -0.43379072 | -0.782391334 | 7.85E-16 | 0.029369512 |
| PRDX2 | BC | young | -0.434128932 | -0.845712603 | 3.04E-36 | 0.002933594 |
| NDUFV2 | BC | young | -0.435144758 | -0.676023186 | 1.49E-25 | 0.023323315 |
| MRPL28 | BC | young | -0.435361506 | -0.93832993 | 7.25E-12 | 0.047663708 |
| ZFHX2 | BC | young | -0.435396239 | -1.154130147 | 2.13E-38 | 8.19E-05 |
| PDHA1 | BC | young | -0.436814303 | -1.205530016 | 8.25E-25 | 0.000256318 |
| CHMP5 | BC | young | -0.437148336 | -1.018289101 | 8.10E-47 | 0.000482787 |
| H2AFZ | BC | young | -0.438370845 | -0.7468126 | 5.36E-32 | 0.002031195 |
| DUT | BC | young | -0.439898726 | -0.797991819 | 5.51E-17 | 0.010018515 |
| ARF5 | BC | young | -0.440863286 | -0.894108457 | 3.32E-16 | 0.017258232 |
| ZBTB20-AS5 | BC | young | -0.44128578 | -0.901764145 | 3.34E-17 | 0.014825396 |
| PNKD | BC | young | -0.441451701 | -1.016730926 | 7.92E-32 | 0.000760477 |
| ELOB | BC | young | -0.441853493 | -0.813758261 | 5.34E-25 | 0.004406514 |
| MFAP1 | BC | young | -0.442307945 | -1.08634168 | 1.25E-23 | 0.002861577 |
| MRPL20 | BC | young | -0.442709019 | -0.896077197 | 1.61E-28 | 0.000663823 |
| MRPS7 | BC | young | -0.447496587 | -0.9135253 | 2.05E-15 | 0.018423839 |
| EIF4G2 | BC | young | -0.44906797 | -0.831703971 | 1.09E-23 | 0.001831394 |
| AL354740.1 | BC | young | -0.449693188 | -0.984856973 | 9.24E-10 | 0.02100854 |
| DDX49 | BC | young | -0.450203115 | -1.134323601 | 2.97E-08 | 0.028870195 |
| TRMT112 | BC | young | -0.450493812 | -0.713148084 | 2.19E-28 | 0.010899232 |
| ATP5PF | BC | young | -0.450576234 | -0.816644664 | 1.00E-40 | 0.000566553 |
| MIEN1 | BC | young | -0.451283171 | -0.987768672 | 9.23E-12 | 0.009940571 |
| HIST3H2A | BC | young | -0.451508111 | -1.506242945 | 2.99E-18 | 6.61E-05 |
| DYNLL1 | BC | young | -0.451976153 | -0.658655111 | 2.90E-25 | 0.018622502 |
| PSMC1 | BC | young | -0.452279524 | -0.969102922 | 1.29E-13 | 0.017552357 |
| NDUFA5 | BC | young | -0.452464132 | -0.642128753 | 1.69E-22 | 0.009700108 |
| CCT7 | BC | young | -0.452600389 | -0.919273789 | 1.54E-17 | 0.005608105 |
| CCT6A | BC | young | -0.45280208 | -1.087927736 | 1.48E-34 | 0.000611393 |
| COX5B | BC | young | -0.45368894 | -0.734028303 | 2.59E-32 | 0.009327523 |
| EIF3I | BC | young | -0.453883015 | -1.014392906 | 2.80E-17 | 0.011304366 |
| H3F3A | BC | young | -0.455129299 | -1.244510955 | 1.22E-81 | 2.21E-12 |
| COX7B | BC | young | -0.45563885 | -0.641842377 | 1.03E-16 | 0.036867906 |
| MT-ATP6 | BC | young | -0.457910285 | -0.921945715 | 4.69E-81 | 0.001165087 |
| FUNDC2 | BC | young | -0.460772978 | -0.800185102 | 4.30E-15 | 0.024290959 |
| LSM7 | BC | young | -0.463635212 | -0.942186351 | 1.30E-19 | 0.010084954 |
| NCL | BC | young | -0.464017382 | -1.573064179 | 7.92E-89 | 9.14E-10 |
| ATP5IF1 | BC | young | -0.465057482 | -0.969464073 | 1.14E-46 | 0.000411352 |
| DDX24 | BC | young | -0.465889795 | -0.846183811 | 1.44E-29 | 0.005524905 |
| SMDT1 | BC | young | -0.466121436 | -0.944621335 | 1.05E-31 | 0.001439498 |
| UBE2S | BC | young | -0.466203633 | -0.805991397 | 2.43E-16 | 0.024113665 |
| TMLHE-AS1 | BC | young | -0.46720131 | -1.01298687 | 6.78E-10 | 0.008953307 |
| CLTA | BC | young | -0.467894068 | -0.592204828 | 1.14E-18 | 0.042958545 |
| RPS19BP1 | BC | young | -0.469914118 | -1.126130381 | 7.47E-49 | 2.77E-05 |
| ERLEC1 | BC | young | -0.473581064 | -0.769604995 | 7.24E-12 | 0.031918805 |

| | | | | | | |
|---|---|---|---|---|---|---|
| LSM4 | BC | young | -0.47406395 | -1.13564626 | 5.58E-50 | 5.47E-05 |
| FSIP2 | BC | young | -0.474737531 | -0.973726749 | 5.68E-12 | 0.042906369 |
| NHP2 | BC | young | -0.474857569 | -0.934462154 | 1.48E-21 | 0.027083976 |
| TMA7 | BC | young | -0.477551447 | -0.634208209 | 7.24E-21 | 0.032035534 |
| TALDO1 | BC | young | -0.478076237 | -0.860023556 | 9.01E-15 | 0.017542598 |
| FIS1 | BC | young | -0.478700891 | -1.256350358 | 1.39E-38 | 0.000163797 |
| CISD1 | BC | young | -0.479269671 | -0.860439988 | 2.44E-23 | 0.004477516 |
| COX7A1 | BC | young | -0.479377662 | -0.695109079 | 2.59E-05 | 0.040661759 |
| ANAPC15 | BC | young | -0.48049563 | -1.230505717 | 8.68E-22 | 0.000363452 |
| EID2 | BC | young | -0.481354843 | -1.158417088 | 3.95E-19 | 0.009804635 |
| POLR2I | BC | young | -0.483242133 | -0.758206998 | 1.59E-13 | 0.041223278 |
| RWDD1 | BC | young | -0.483529884 | -0.728296151 | 2.59E-24 | 0.011447508 |
| PARK7 | BC | young | -0.48403771 | -0.988163035 | 3.49E-52 | 0.000125863 |
| TRAPPC4 | BC | young | -0.48572365 | -1.051475349 | 1.33E-21 | 0.003086538 |
| C1orf122 | BC | young | -0.487449542 | -0.748029546 | 4.08E-17 | 0.023393563 |
| STUB1 | BC | young | -0.488363928 | -0.926187179 | 3.78E-23 | 0.007274664 |
| GPN3 | BC | young | -0.488869547 | -0.816179791 | 1.28E-13 | 0.038755152 |
| AC090531.1 | BC | young | -0.489251354 | -1.145419978 | 5.64E-13 | 0.007296131 |
| SMIM26 | BC | young | -0.48993641 | -0.647944554 | 1.70E-15 | 0.038650049 |
| EIF2S2 | BC | young | -0.490689532 | -0.733118465 | 1.46E-13 | 0.044915962 |
| TIMM17B | BC | young | -0.491803206 | -1.031627983 | 1.66E-23 | 0.003415433 |
| PYURF | BC | young | -0.491872415 | -0.870972005 | 4.90E-14 | 0.02407532 |
| TCEAL3 | BC | young | -0.492040684 | -0.830874225 | 2.50E-24 | 0.017258232 |
| UQCR11 | BC | young | -0.492935427 | -0.751644926 | 1.06E-22 | 0.011001776 |
| AP001825.1 | BC | young | -0.494408046 | -1.271717397 | 6.32E-79 | 8.74E-08 |
| C12orf65 | BC | young | -0.498140121 | -0.954340775 | 3.01E-11 | 0.020042554 |
| NRN1L | BC | young | -0.498710573 | -1.409779017 | 2.37E-46 | 2.11E-06 |
| LEO1 | BC | young | -0.501061274 | -0.993785694 | 1.36E-16 | 0.00616198 |
| APOBEC2 | BC | young | -0.501882061 | -1.734598548 | 7.53E-54 | 6.47E-06 |
| ATP5MPL | BC | young | -0.502093375 | -0.68564909 | 2.06E-21 | 0.024832063 |
| SELENOS | BC | young | -0.502371759 | -0.993483114 | 1.15E-17 | 0.009816449 |
| COX14 | BC | young | -0.502972916 | -0.856260359 | 1.17E-22 | 0.005377288 |
| TSFM | BC | young | -0.504763731 | -0.95661874 | 3.24E-12 | 0.049199834 |
| PNO1 | BC | young | -0.504914009 | -0.989180375 | 1.39E-12 | 0.023367332 |
| TCEAL2 | BC | young | -0.505910804 | -0.784769368 | 8.61E-24 | 0.016015728 |
| NDUFA6 | BC | young | -0.510528122 | -0.782771288 | 3.57E-17 | 0.029256281 |
| MT-ND4 | BC | young | -0.512228833 | -0.697408284 | 3.97E-35 | 0.005524905 |
| DNAJC19 | BC | young | -0.512370948 | -1.053716045 | 1.71E-24 | 0.003006852 |
| MRPS23 | BC | young | -0.512419753 | -1.094777911 | 2.18E-21 | 0.002377376 |
| PCNA | BC | young | -0.513673826 | -1.253098599 | 6.95E-16 | 0.000540532 |
| LRTM1 | BC | young | -0.514413845 | -1.605927692 | 4.49E-88 | 1.86E-14 |
| NDUFS3 | BC | young | -0.514967481 | -1.04441524 | 2.49E-22 | 0.006870714 |
| AP2S1 | BC | young | -0.515208554 | -0.873172742 | 4.02E-14 | 0.021173714 |
| AL513164.1 | BC | young | -0.515358897 | -1.074109118 | 5.23E-35 | 0.005382825 |
| SLC24A1 | BC | young | -0.515948942 | -1.358548981 | 1.70E-25 | 0.000272575 |
| MRPL13 | BC | young | -0.517723283 | -0.8502031 | 6.83E-15 | 0.022675005 |
| CHMP2A | BC | young | -0.518435964 | -0.756177329 | 3.42E-20 | 0.043860434 |
| SIVA1 | BC | young | -0.519549912 | -1.06908823 | 6.69E-22 | 0.002169185 |
| PHOSPHO2 | BC | young | -0.519634047 | -1.101229718 | 6.06E-07 | 0.0345489 |
| GET3 | BC | young | -0.522630027 | -1.233207848 | 5.78E-15 | 0.0028547 |
| SLC25A5 | BC | young | -0.524493945 | -0.995412206 | 1.06E-20 | 0.00056928 |
| MICOS13 | BC | young | -0.524863535 | -0.848343659 | 1.49E-10 | 0.040758533 |
| DCTPP1 | BC | young | -0.527578347 | -0.930530896 | 3.05E-11 | 0.044742836 |
| AC011306.1 | BC | young | -0.527721646 | -1.134039531 | 1.61E-18 | 0.007560475 |

| | | | | | | |
|---|---|---|---|---|---|---|
| UBL5 | BC | young | -0.528014373 | -0.886338299 | 3.07E-45 | 0.000535689 |
| MICOS10 | BC | young | -0.529898315 | -0.966105815 | 1.03E-39 | 8.15E-05 |
| IL1RAP | BC | young | -0.531576331 | -1.105641038 | 2.71E-28 | 0.001985705 |
| B3GALT2 | BC | young | -0.532967725 | -1.049530863 | 1.66E-27 | 0.002164841 |
| SMIM8 | BC | young | -0.535156898 | -0.997420375 | 3.46E-13 | 0.017838076 |
| MIS12 | BC | young | -0.536825693 | -1.113256058 | 6.45E-14 | 0.011425992 |
| UQCR10 | BC | young | -0.536995208 | -0.769710007 | 5.94E-31 | 0.001891735 |
| CAMK2B | BC | young | -0.537566318 | -0.867422228 | 5.44E-28 | 0.000387936 |
| TMEM160 | BC | young | -0.53806845 | -1.17085602 | 2.48E-17 | 0.001219564 |
| VBP1 | BC | young | -0.538491492 | -0.930667106 | 4.11E-33 | 0.002855305 |
| PSMC5 | BC | young | -0.539473915 | -0.905452062 | 4.85E-15 | 0.013798582 |
| ATP5F1D | BC | young | -0.540617372 | -0.806154734 | 4.93E-20 | 0.017132961 |
| SLIRP | BC | young | -0.540858023 | -0.838721576 | 1.01E-31 | 0.003435133 |
| HPF1 | BC | young | -0.541929972 | -0.776156117 | 1.99E-15 | 0.028654397 |
| CNPY2 | BC | young | -0.545502385 | -1.03576669 | 2.66E-26 | 0.002464351 |
| MRPL47 | BC | young | -0.546952656 | -0.938480132 | 5.79E-13 | 0.022909921 |
| TXNDC17 | BC | young | -0.549295385 | -1.141406483 | 1.31E-11 | 0.006528368 |
| UFC1 | BC | young | -0.552601866 | -1.013714782 | 1.03E-28 | 0.001880015 |
| ZCRB1 | BC | young | -0.553483406 | -0.864051335 | 1.40E-21 | 0.00332387 |
| TMEM147 | BC | young | -0.553780871 | -1.013457907 | 1.24E-13 | 0.012671652 |
| ECHS1 | BC | young | -0.554569664 | -1.201842757 | 3.75E-27 | 0.000667753 |
| NDUFB8 | BC | young | -0.556306978 | -0.737734587 | 4.94E-28 | 0.013355087 |
| AL445526.1 | BC | young | -0.557107239 | -0.905275394 | 2.47E-08 | 0.031611827 |
| C18orf32 | BC | young | -0.557387273 | -0.894891265 | 6.70E-47 | 0.000990475 |
| VPS28 | BC | young | -0.558106676 | -0.981416865 | 2.20E-21 | 0.004080632 |
| COX7A2 | BC | young | -0.558385268 | -0.840907177 | 2.37E-43 | 0.000637712 |
| PSMA4 | BC | young | -0.558524196 | -1.360309792 | 2.00E-30 | 1.46E-05 |
| COA3 | BC | young | -0.560247305 | -1.023530025 | 5.47E-20 | 0.004406514 |
| RD3 | BC | young | -0.563908821 | -0.722264167 | 2.32E-12 | 0.042360159 |
| BEX1 | BC | young | -0.563920229 | -0.83692091 | 5.07E-39 | 0.005016224 |
| EBAG9 | BC | young | -0.564693914 | -0.922934464 | 4.10E-15 | 0.017258232 |
| NDUFA7 | BC | young | -0.565312544 | -0.841914395 | 2.00E-29 | 0.004395024 |
| POP5 | BC | young | -0.565953267 | -1.219576206 | 7.93E-11 | 0.006845964 |
| SAP18 | BC | young | -0.567356292 | -1.056315461 | 4.39E-47 | 0.00027742 |
| MXRA7 | BC | young | -0.567751507 | -0.885896988 | 9.25E-30 | 0.002464593 |
| NDUFB10 | BC | young | -0.569572478 | -0.926335956 | 2.13E-22 | 0.004363341 |
| ARL16 | BC | young | -0.570044243 | -1.091302643 | 4.40E-16 | 0.026840855 |
| NDUFC2 | BC | young | -0.57047405 | -0.971014003 | 6.13E-50 | 0.000350422 |
| GCSH | BC | young | -0.571846632 | -0.973378163 | 9.34E-19 | 0.008229166 |
| MRPS26 | BC | young | -0.572075023 | -1.116385114 | 2.06E-24 | 0.002968467 |
| FKBP3 | BC | young | -0.572157195 | -1.260061372 | 1.61E-47 | 2.11E-06 |
| CWC15 | BC | young | -0.572771655 | -0.702235954 | 2.23E-16 | 0.036739875 |
| UQCC2 | BC | young | -0.573375922 | -0.857936832 | 2.49E-21 | 0.012387307 |
| GLRX5 | BC | young | -0.574835172 | -0.834965536 | 7.79E-11 | 0.047569725 |
| NDUFS5 | BC | young | -0.575889169 | -0.961077327 | 2.74E-39 | 0.000208867 |
| COX6C | BC | young | -0.57878192 | -0.800695083 | 5.16E-35 | 0.000916149 |
| MZT1 | BC | young | -0.579105059 | -0.844885056 | 7.67E-16 | 0.026293225 |
| AC023136.1 | BC | young | -0.579218287 | -1.047530083 | 8.15E-16 | 0.024907956 |
| HINT2 | BC | young | -0.582579387 | -1.025352167 | 1.38E-12 | 0.013032849 |
| TMEM14C | BC | young | -0.583228118 | -0.839673719 | 5.16E-23 | 0.006872708 |
| COX6B1 | BC | young | -0.585230018 | -0.751805297 | 1.59E-25 | 0.008494616 |
| C15orf61 | BC | young | -0.586404016 | -0.757168085 | 8.05E-16 | 0.016310291 |
| MAGOH | BC | young | -0.588495425 | -1.025478667 | 1.08E-24 | 0.0032417 |
| PIGBOS1 | BC | young | -0.588886228 | -1.063121967 | 9.50E-07 | 0.045867482 |

| | | | | | | |
|---|---|---|---|---|---|---|
| NOP56 | BC | young | -0.589820452 | -1.070317433 | 2.17E-08 | 0.020645157 |
| TCEAL6 | BC | young | -0.589925984 | -0.953113756 | 1.54E-30 | 0.004609547 |
| EEF1E1 | BC | young | -0.589999272 | -0.879648037 | 9.09E-16 | 0.025371483 |
| ANOS1 | BC | young | -0.590371967 | -1.193914239 | 9.25E-64 | 0.000131245 |
| NDUFA8 | BC | young | -0.590726432 | -0.982137292 | 1.10E-26 | 0.002144985 |
| POP4 | BC | young | -0.590996799 | -1.251676733 | 6.43E-20 | 0.004058237 |
| ANAPC11 | BC | young | -0.59135415 | -0.861640787 | 2.36E-21 | 0.006103661 |
| VMO1 | BC | young | -0.591554443 | -1.397282872 | 1.22E-17 | 0.006529114 |
| THUMPD1 | BC | young | -0.592435627 | -1.01815126 | 4.73E-15 | 0.007121419 |
| KCNMA1-AS1 | BC | young | -0.593616924 | -1.216601652 | 1.28E-23 | 0.000860337 |
| PSMA7 | BC | young | -0.597830242 | -0.901520223 | 7.74E-32 | 0.000564866 |
| AC104041.1 | BC | young | -0.602475244 | -1.285673404 | 6.58E-22 | 0.000918634 |
| HRAS | BC | young | -0.602486408 | -1.213176477 | 1.16E-17 | 0.002323982 |
| LSM1 | BC | young | -0.602631313 | -0.951646934 | 7.45E-16 | 0.015152009 |
| MRFAP1 | BC | young | -0.603967477 | -0.823441117 | 2.59E-31 | 0.017280252 |
| TAFA3 | BC | young | -0.606283341 | -1.357877515 | 4.36E-35 | 0.000485092 |
| ANKRD39 | BC | young | -0.606537048 | -1.017041098 | 6.62E-11 | 0.02577769 |
| KRT10 | BC | young | -0.607386079 | -0.699759282 | 4.31E-21 | 0.031607699 |
| NDUFA4 | BC | young | -0.607780274 | -0.644583726 | 4.78E-27 | 0.00680415 |
| C14orf119 | BC | young | -0.609285224 | -0.923339606 | 4.42E-16 | 0.01916146 |
| GRHPR | BC | young | -0.609358009 | -0.837889288 | 1.23E-15 | 0.040330345 |
| EIF3K | BC | young | -0.609442349 | -0.925670335 | 4.51E-20 | 0.002212166 |
| POLR2K | BC | young | -0.609827519 | -1.129632154 | 1.27E-40 | 5.98E-05 |
| LINC02470 | BC | young | -0.611072816 | -1.12086256 | 0.000853628 | 0.049146147 |
| GTF2A2 | BC | young | -0.611428249 | -0.789403913 | 1.65E-21 | 0.02384533 |
| NDUFS8 | BC | young | -0.611860017 | -0.850725468 | 2.90E-19 | 0.026957967 |
| SBDS | BC | young | -0.612537682 | -0.907474141 | 1.11E-27 | 0.003547346 |
| NAA38 | BC | young | -0.613542269 | -0.686743869 | 7.83E-18 | 0.01916146 |
| NQO1 | BC | young | -0.613963997 | -1.796211344 | 2.16E-56 | 8.29E-07 |
| LINC02055 | BC | young | -0.614279332 | -1.850199274 | 5.37E-64 | 4.94E-09 |
| ATXN7L3B | BC | young | -0.614520716 | -0.813703486 | 8.60E-20 | 0.011096755 |
| ISCA2 | BC | young | -0.617082406 | -1.110936455 | 3.27E-17 | 0.005975743 |
| HNRNPAB | BC | young | -0.618065822 | -1.478571834 | 2.97E-24 | 6.57E-05 |
| AC009975.2 | BC | young | -0.618967756 | -1.702379251 | 2.15E-28 | 4.49E-05 |
| SNRNP25 | BC | young | -0.619471741 | -0.997139618 | 4.24E-10 | 0.024162659 |
| LARP7 | BC | young | -0.621474061 | -1.107936413 | 1.96E-23 | 0.001483118 |
| SELENOW | BC | young | -0.623973743 | -0.7744965 | 1.45E-32 | 0.003986635 |
| PIN1 | BC | young | -0.62427276 | -1.023472432 | 6.62E-25 | 0.004409745 |
| C2orf83 | BC | young | -0.624469349 | -1.733165336 | 7.35E-14 | 0.000300741 |
| MRPL11 | BC | young | -0.624951696 | -1.180686752 | 2.83E-11 | 0.006847008 |
| TPRKB | BC | young | -0.628525492 | -0.895887176 | 3.88E-10 | 0.035997457 |
| PRDX5 | BC | young | -0.629368281 | -0.919042014 | 1.10E-35 | 0.004722725 |
| YEATS4 | BC | young | -0.631500351 | -0.924728446 | 8.93E-13 | 0.035134781 |
| CCDC173 | BC | young | -0.632337883 | -0.937091555 | 2.78E-14 | 0.037247304 |
| TOMM5 | BC | young | -0.632859593 | -0.998281498 | 8.16E-25 | 0.00408277 |
| COX8A | BC | young | -0.63472271 | -0.957326992 | 4.49E-54 | 0.001832773 |
| C17orf75 | BC | young | -0.634858765 | -1.072108496 | 1.02E-17 | 0.004409745 |
| RAB5IF | BC | young | -0.635414584 | -1.110703548 | 6.35E-22 | 0.001223917 |
| CKLF | BC | young | -0.638786171 | -1.507482536 | 5.90E-21 | 0.000286125 |
| SNF8 | BC | young | -0.640010138 | -1.008057529 | 9.55E-15 | 0.009700108 |
| PRELID1 | BC | young | -0.641888836 | -1.23399674 | 1.45E-28 | 0.000103062 |
| RNASEK | BC | young | -0.642144826 | -1.004288743 | 3.06E-39 | 0.000321945 |
| CALM1 | BC | young | -0.642282115 | -0.883164266 | 2.31E-48 | 0.010491115 |
| NIF3L1 | BC | young | -0.647901792 | -1.116822881 | 1.01E-59 | 0.027814667 |

| | | | | | | |
|---|---|---|---|---|---|---|
| MRPL58 | BC | young | -0.650243079 | -1.275288931 | 5.24E-20 | 0.001094758 |
| KRTCAP2 | BC | young | -0.651283203 | -0.973657316 | 3.86E-23 | 0.004469866 |
| NAGK | BC | young | -0.651885867 | -0.979855107 | 2.30E-11 | 0.027083976 |
| PIGP | BC | young | -0.652275845 | -0.872914988 | 1.73E-17 | 0.018423839 |
| SLC25A11 | BC | young | -0.653042432 | -1.359589523 | 1.76E-22 | 0.000326381 |
| NDUFB4 | BC | young | -0.65333411 | -0.993914614 | 2.24E-48 | 0.000117583 |
| SEC11C | BC | young | -0.653929011 | -0.784657081 | 1.04E-20 | 0.021651471 |
| ETFB | BC | young | -0.654371209 | -1.120506494 | 2.05E-30 | 0.000297598 |
| ATP5MC1 | BC | young | -0.657071409 | -1.540937372 | 1.68E-68 | 9.30E-09 |
| POLR2J | BC | young | -0.657675374 | -0.924692079 | 6.06E-12 | 0.021964435 |
| BAG1 | BC | young | -0.659386637 | -0.76968525 | 6.78E-10 | 0.019911568 |
| ATP5MF | BC | young | -0.664394925 | -0.76680401 | 2.96E-18 | 0.016224624 |
| LRRC24 | BC | young | -0.669040327 | -1.369550001 | 1.16E-25 | 0.001141643 |
| EID1 | BC | young | -0.67079087 | -1.264145081 | 4.10E-75 | 3.09E-09 |
| SNHG30 | BC | young | -0.67148243 | -1.055142515 | 1.55E-12 | 0.005908123 |
| PDHB | BC | young | -0.672329102 | -1.164826423 | 6.23E-14 | 0.006448942 |
| JTB | BC | young | -0.673250065 | -1.032614351 | 7.65E-23 | 0.002270971 |
| MDP1 | BC | young | -0.675967517 | -1.168257765 | 1.47E-15 | 0.005020937 |
| NOL7 | BC | young | -0.678483344 | -0.860017646 | 4.45E-27 | 0.004494761 |
| POLR3K | BC | young | -0.679030544 | -1.253486194 | 1.53E-14 | 0.003488453 |
| ATP5PD | BC | young | -0.679296641 | -0.806605076 | 1.25E-25 | 0.003783869 |
| AC016042.1 | BC | young | -0.680252459 | -0.803721554 | 7.34E-19 | 0.040397978 |
| NDUFB9 | BC | young | -0.681479238 | -1.119148097 | 1.01E-43 | 0.000275867 |
| AC087855.1 | BC | young | -0.68362571 | -1.064607504 | 6.41E-40 | 0.000498649 |
| SCAND1 | BC | young | -0.686822744 | -1.053451611 | 1.01E-23 | 0.00358454 |
| NDUFB3 | BC | young | -0.696493519 | -0.805661233 | 1.27E-22 | 0.010865954 |
| APRT | BC | young | -0.69775627 | -0.939840777 | 1.16E-09 | 0.037617984 |
| COPS9 | BC | young | -0.700422794 | -1.129508111 | 8.54E-41 | 0.000106522 |
| GSTO1 | BC | young | -0.702169066 | -1.335197892 | 3.91E-35 | 2.80E-05 |
| NDUFAB1 | BC | young | -0.70288404 | -1.300771616 | 6.15E-68 | 5.39E-07 |
| CCT5 | BC | young | -0.702940149 | -1.088808581 | 5.80E-32 | 0.001923265 |
| NDUFA2 | BC | young | -0.705066434 | -1.031630796 | 1.73E-37 | 0.000609642 |
| MRPL21 | BC | young | -0.705846068 | -1.145171688 | 4.48E-25 | 0.000714706 |
| AURKAIP1 | BC | young | -0.712614239 | -0.905076331 | 1.62E-27 | 0.003018598 |
| PGP | BC | young | -0.714462601 | -1.20542301 | 7.27E-21 | 0.001053415 |
| FMC1 | BC | young | -0.716425658 | -1.105079142 | 8.68E-35 | 6.25E-05 |
| NDUFS6 | BC | young | -0.717221535 | -1.261226941 | 7.62E-43 | 5.31E-06 |
| BLOC1S1 | BC | young | -0.720265479 | -0.873927774 | 2.30E-14 | 0.01120523 |
| MRPL33 | BC | young | -0.724806871 | -0.689252972 | 2.24E-19 | 0.013561746 |
| NDUFA1 | BC | young | -0.725609659 | -0.825464149 | 1.41E-33 | 0.001659626 |
| MRPS34 | BC | young | -0.730337291 | -1.274497741 | 2.93E-20 | 0.001291251 |
| AC068051.1 | BC | young | -0.733199915 | -1.060314417 | 1.05E-21 | 0.00368145 |
| GADD45GIP1 | BC | young | -0.734925721 | -0.873818105 | 5.73E-14 | 0.02604969 |
| ROMO1 | BC | young | -0.736578858 | -0.990910392 | 9.40E-24 | 0.007110341 |
| ATP5ME | BC | young | -0.738254713 | -0.724466137 | 1.07E-18 | 0.040227493 |
| MRPL27 | BC | young | -0.74093723 | -1.205245478 | 5.21E-17 | 0.001361289 |
| MRPL32 | BC | young | -0.741470628 | -1.139416356 | 1.31E-19 | 0.005073069 |
| HIGD2A | BC | young | -0.74230759 | -0.906577487 | 4.86E-17 | 0.014841625 |
| NDUFAF3 | BC | young | -0.748364103 | -1.133802208 | 6.51E-28 | 0.00065258 |
| EMC6 | BC | young | -0.748788685 | -1.225065944 | 1.76E-24 | 0.000587706 |
| HELLPAR | BC | young | -0.750875938 | -1.238897443 | 3.41E-25 | 0.000225814 |
| NDUFB7 | BC | young | -0.75181519 | -1.003794772 | 1.78E-41 | 0.000440359 |
| GTF3C6 | BC | young | -0.752223782 | -1.106027506 | 3.93E-20 | 0.002440217 |
| PSMB6 | BC | young | -0.755850191 | -1.436837179 | 5.03E-46 | 9.49E-07 |

| | | | | | | |
|---|---|---|---|---|---|---|
| NME1 | BC | young | -0.760518132 | -1.151986906 | 1.12E-51 | 1.90E-05 |
| OTUD6B-AS1 | BC | young | -0.762111011 | -0.680560117 | 3.10E-16 | 0.035117354 |
| COX17 | BC | young | -0.764440567 | -0.79624961 | 7.84E-32 | 0.00236126 |
| PDCL3 | BC | young | -0.767730899 | -1.503010839 | 1.29E-21 | 0.000328152 |
| NDUFB6 | BC | young | -0.773748436 | -0.996021731 | 1.09E-41 | 0.000252845 |
| ALKBH7 | BC | young | -0.773896757 | -0.871309586 | 7.36E-21 | 0.013853233 |
| MRPL34 | BC | young | -0.778262538 | -0.811420548 | 1.10E-13 | 0.037034887 |
| NDUFB11 | BC | young | -0.779631053 | -1.065520804 | 1.27E-33 | 0.000310978 |
| SRSF3 | BC | young | -0.783291507 | -0.758223855 | 4.42E-29 | 0.026316469 |
| AC091938.1 | BC | young | -0.789198021 | -1.7620027 | 1.34E-90 | 6.63E-09 |
| POLR2L | BC | young | -0.789293938 | -1.167962731 | 6.10E-31 | 0.000119953 |
| MRPL15 | BC | young | -0.790174545 | -1.007496482 | 1.37E-08 | 0.030924707 |
| BOLA3 | BC | young | -0.791036418 | -0.903195622 | 4.33E-21 | 0.011778053 |
| MRPL54 | BC | young | -0.793670357 | -1.156044571 | 8.49E-29 | 0.000586384 |
| NPIPB2 | BC | young | -0.799093527 | -1.456754297 | 2.62E-12 | 0.002855305 |
| AL589693.1 | BC | young | -0.80072798 | -0.979562795 | 1.50E-05 | 0.013567242 |
| EFCAB2 | BC | young | -0.811367888 | -1.190075746 | 6.25E-14 | 0.001404833 |
| MRPS33 | BC | young | -0.816975094 | -0.89532881 | 1.05E-12 | 0.035598014 |
| NCBP2AS2 | BC | young | -0.818958733 | -1.187472471 | 1.03E-15 | 0.005550972 |
| PSG8 | BC | young | -0.819855261 | -1.178794643 | 1.83E-23 | 0.005541252 |
| MRPL57 | BC | young | -0.820008866 | -1.316301913 | 2.88E-48 | 9.48E-06 |
| C1QTNF7 | BC | young | -0.824669026 | -1.026345798 | 2.20E-21 | 0.005758053 |
| TRAPPC5 | BC | young | -0.831988078 | -0.912530011 | 1.19E-22 | 0.00814129 |
| KRT222 | BC | young | -0.832584488 | -1.229601551 | 1.22E-51 | 0.00045799 |
| MRPS36 | BC | young | -0.833275678 | -0.993850951 | 7.56E-22 | 0.014249129 |
| MT-ND1 | BC | young | -0.833314608 | -0.758710161 | 5.96E-56 | 0.002189101 |
| PHB | BC | young | -0.835020594 | -1.119124137 | 5.29E-18 | 0.01120523 |
| CD320 | BC | young | -0.837719511 | -1.28072098 | 6.18E-11 | 0.005165816 |
| CHN2 | BC | young | -0.842479202 | -1.098322067 | 1.05E-15 | 0.019147959 |
| DNAJC15 | BC | young | -0.85250204 | -0.943448434 | 3.31E-16 | 0.015926282 |
| BX664615.2 | BC | young | -0.853046388 | -1.001474776 | 8.28E-21 | 0.007815093 |
| POP7 | BC | young | -0.85605476 | -1.049295137 | 1.07E-12 | 0.044771253 |
| AC104117.3 | BC | young | -0.856235705 | -1.229504253 | 3.23E-45 | 0.000446135 |
| PGLS | BC | young | -0.864579522 | -1.070749211 | 3.65E-07 | 0.032835313 |
| NDUFB1 | BC | young | -0.886564915 | -1.061283022 | 5.55E-41 | 0.001251396 |
| CXCL14 | BC | young | -0.892322589 | -1.088289496 | 3.25E-40 | 0.003506614 |
| HIST1H2AC | BC | young | -0.898648149 | -1.02766566 | 7.09E-27 | 0.012089232 |
| NDUFA11 | BC | young | -0.907650519 | -0.97380034 | 6.00E-22 | 0.001343314 |
| MT-CYB | BC | young | -0.924686188 | -1.04026706 | 1.63E-63 | 3.95E-05 |
| NDUFB2 | BC | young | -0.930152936 | -0.86485391 | 5.69E-31 | 0.000498198 |
| MT-CO2 | BC | young | -0.960854294 | -1.685962086 | 8.62E-164 | 5.12E-15 |
| TTYH1 | BC | young | -0.999149569 | -1.723030183 | 8.99E-49 | 7.44E-06 |
| HIST1H4C | BC | young | -1.009469128 | -0.804647198 | 6.50E-19 | 0.007412586 |
| GRM5-AS1 | BC | young | -1.018036528 | -1.540648616 | 9.17E-26 | 0.001402337 |
| PCP2 | BC | young | -1.046032779 | -1.61867726 | 4.50E-69 | 3.64E-05 |
| BX255923.1 | BC | young | -1.04937545 | -1.141021182 | 1.13E-06 | 0.025500342 |
| MT-ND2 | BC | young | -1.051523615 | -0.905743747 | 2.60E-57 | 0.000990475 |
| MT-ND3 | BC | young | -1.053556553 | -1.691980464 | 8.40E-158 | 9.17E-15 |
| LINC02315 | BC | young | -1.058177535 | -1.16920228 | 7.09E-09 | 0.001326485 |
| FAM153A | BC | young | -1.102923476 | -2.145101236 | 0.000347991 | 0.040661759 |
| REC114 | BC | young | -1.10887549 | -1.038123059 | 1.49E-14 | 0.029687764 |
| RIT2 | BC | young | -1.163549646 | -1.056076241 | 4.05E-12 | 0.019489786 |
| TDRG1 | BC | young | -1.178373845 | -1.184858616 | 1.09E-31 | 0.013042501 |
| FAM138C | BC | young | -1.19596266 | -2.063285897 | 6.38E-32 | 0.00143305 |

| | | | | | | |
|---|---|---|---|---|---|---|
| AC092155.1 | BC | young | -1.20214503 | -1.859405949 | 3.01E-53 | 4.17E-07 |
| MT-ATP8 | BC | young | -1.203433235 | -0.953280346 | 4.17E-52 | 2.09E-05 |
| MED11 | BC | young | -1.205137783 | -1.339301335 | 1.52E-12 | 0.016560152 |
| MT-ND5 | BC | young | -1.228867053 | -1.233931771 | 2.23E-97 | 6.05E-08 |
| ABCA12 | BC | young | -1.395498901 | -1.073177953 | 2.63E-23 | 0.009464599 |
| PGAM2 | BC | young | -1.410653401 | -1.443333457 | 4.36E-14 | 0.002015075 |
| ATP6V0D2 | BC | young | -1.496725258 | -1.384160037 | 1.46E-07 | 0.032819382 |
| AL627171.2 | BC | young | -1.580307171 | -1.091574823 | 6.02E-20 | 0.029417089 |
| CDR1 | BC | young | -1.858196583 | -1.930289779 | 3.19E-83 | 2.30E-08 |
| MT-ND6 | BC | young | -2.271707299 | -2.479011362 | 4.83E-213 | 1.36E-17 |
| EGR1 | HC | old | 2.470307757 | 1.274488253 | 0.001387217 | 0.002141236 |
| ADARB2 | HC | old | 2.299753194 | 1.812001617 | 0.008307407 | 0.007672862 |
| MT2A | HC | old | 2.217421621 | 1.631915846 | 6.23E-05 | 0.000149814 |
| DTNA | HC | old | 2.126727007 | 2.12054242 | 1.69E-06 | 3.76E-05 |
| CRYAB | HC | old | 2.061411299 | 2.2986941 | 2.22E-10 | 2.85E-07 |
| TMSB10 | HC | old | 1.998649754 | 1.610143547 | 2.69E-08 | 7.99E-05 |
| HSPA6 | HC | old | 1.950559972 | 0.62060056 | 9.59E-14 | 0.004727617 |
| BHLHE40 | HC | old | 1.856581502 | 2.03446556 | 2.72E-08 | 0.00013574 |
| XKR4 | HC | old | 1.785821028 | 1.313532003 | 0.002350121 | 0.049838945 |
| DCLK1 | HC | old | 1.549748928 | 1.318427034 | 0.016030902 | 0.049396255 |
| PPP1R3C | HC | old | 1.5200214 | 1.55021965 | 0.023934395 | 0.018786425 |
| MARCKSL1 | HC | old | 1.50213979 | 1.476205012 | 0.000728858 | 0.015483496 |
| GPX3 | HC | old | 1.464771446 | 1.019006587 | 5.74E-06 | 0.044668162 |
| ERO1A | HC | old | 1.4056335 | 1.364275293 | 0.008576076 | 0.025037151 |
| DDIT4 | HC | old | 1.404583893 | 1.888288886 | 0.040167917 | 0.000574562 |
| CCNG2 | HC | old | 1.402609368 | 1.477448689 | 0.000266251 | 0.019262919 |
| 1-Mar | HC | old | 1.365235821 | 2.319234907 | 2.65E-08 | 1.03E-08 |
| FAM162A | HC | old | 1.279893569 | 1.566968073 | 6.67E-05 | 2.54E-06 |
| HMGCS1 | HC | old | 1.151668405 | 2.912105567 | 5.10E-24 | 9.00E-20 |
| STMN4 | HC | old | 1.110047975 | 1.402081698 | 3.59E-07 | 0.00031102 |
| DNAJB4 | HC | old | 1.08583726 | 1.244596079 | 0.000130755 | 0.025024948 |
| HSPB1 | HC | old | 1.054197051 | 2.863241266 | 6.29E-26 | 2.26E-21 |
| GADD45B | HC | old | 0.958203489 | 1.759709504 | 3.12E-13 | 1.16E-05 |
| JUNB | HC | old | 0.948618623 | 2.257232562 | 9.20E-12 | 1.04E-10 |
| STAT3 | HC | old | 0.919849796 | 1.347776814 | 7.06E-07 | 0.000825527 |
| S100A10 | HC | old | 0.861444784 | 1.300331064 | 0.012492223 | 0.001339043 |
| ENO2 | HC | old | 0.852566954 | 1.156349221 | 3.52E-05 | 4.26E-05 |
| WSB1 | HC | old | 0.852446078 | 1.50961243 | 9.70E-05 | 5.87E-05 |
| LDHA | HC | old | 0.83658946 | 0.870072271 | 0.047439154 | 0.012279248 |
| MARCKS | HC | old | 0.78954565 | 1.047403381 | 7.54E-05 | 0.005983985 |
| UBC | HC | old | 0.777445581 | 2.292315305 | 6.72E-18 | 1.08E-26 |
| SAT1 | HC | old | 0.691473508 | 1.539460924 | 9.32E-06 | 6.43E-06 |
| SCD | HC | old | 0.687275364 | 1.970215068 | 1.12E-09 | 3.16E-07 |
| NRG3 | HC | old | 0.665007626 | 1.701424297 | 2.16E-05 | 0.000482631 |
| NSG1 | HC | old | 0.652405681 | 1.320111571 | 2.89E-08 | 0.001875976 |
| BNIP3 | HC | old | 0.620420399 | 0.965526464 | 0.003942532 | 0.000535305 |
| WDR37 | HC | old | 0.619824874 | 1.319175053 | 0.000634813 | 0.012698147 |
| DDIT3 | HC | old | 0.60719878 | 1.148593149 | 0.01146713 | 0.002998137 |
| C4orf3 | HC | old | 0.581275981 | 0.929624191 | 0.03513368 | 0.012462598 |
| CPE | HC | old | 0.579810809 | 1.382766426 | 5.80E-05 | 7.03E-05 |
| ELOVL5 | HC | old | 0.546875465 | 1.333016901 | 0.006187284 | 0.005037822 |
| PGAM1 | HC | old | 0.530382322 | 1.05568322 | 1.96E-05 | 0.002156802 |
| MSMO1 | HC | old | 0.507852601 | 2.142894269 | 1.02E-24 | 3.37E-12 |
| HMGCR | HC | old | 0.50774298 | 1.055932772 | 8.82E-05 | 0.020270305 |

| | | | | | | |
|---|---|---|---|---|---|---|
| TOMM20 | HC | old | 0.477891851 | 1.266873244 | 0.000116648 | 0.000278141 |
| HPCAL1 | HC | old | 0.466960879 | 1.017804707 | 0.020072978 | 0.009583879 |
| FDFT1 | HC | old | 0.450966786 | 1.302165023 | 8.02E-11 | 0.000245034 |
| SARAF | HC | old | 0.447607162 | 0.938260974 | 0.000115935 | 0.008114874 |
| HSP90AB1 | HC | old | 0.442978167 | 0.941713199 | 0.002034233 | 0.002923326 |
| INSIG1 | HC | old | 0.40407784 | 1.612526909 | 8.90E-10 | 3.30E-06 |
| JUND | HC | old | 0.398744334 | 1.540626869 | 1.37E-10 | 2.85E-07 |
| IDI1 | HC | old | 0.360319208 | 1.248284823 | 3.06E-06 | 0.00031102 |
| MIR7-3HG | HC | old | 0.35472482 | 1.552888433 | 1.42E-06 | 0.000135104 |
| MAPRE2 | HC | old | 0.351339438 | 1.048723352 | 0.003297336 | 0.001483101 |
| PPP1CB | HC | old | 0.341354568 | 0.952878304 | 0.000179726 | 0.002640492 |
| MORF4L2 | HC | old | 0.334002501 | 0.929590171 | 0.019695541 | 0.010219726 |
| JUN | HC | old | 0.298071895 | 1.614874525 | 0.021273956 | 4.32E-06 |
| RHOB | HC | old | 0.285530638 | 1.281958668 | 6.12E-08 | 0.00052904 |
| ZNF10 | HC | old | 0.278244451 | 1.449454758 | 0.002518919 | 8.70E-05 |
| RPL10 | HC | old | 0.265929706 | 1.22240557 | 0.004654133 | 4.21E-07 |
| IER2 | HC | old | 0.261106484 | 0.830058045 | 0.014495651 | 0.011420286 |
| HSPA4L | HC | old | 0.259397501 | 1.201893256 | 0.00431716 | 0.009855031 |
| MT-ND4 | HC | old | 0.253838858 | 0.73576135 | 3.27E-07 | 0.02019095 |
| RPL26 | HC | old | 0.226952413 | 1.114737756 | 0.03302458 | 1.35E-05 |
| HSPH1 | HC | old | 0.220974362 | 2.24210754 | 6.72E-17 | 2.59E-15 |
| UBE2B | HC | old | 0.214670512 | 1.118896188 | 1.77E-05 | 0.000985255 |
| RPL17 | HC | old | 0.210855656 | 1.014012229 | 0.003162586 | 0.000223184 |
| DNAJB1 | HC | old | 0.196487931 | 2.300711437 | 4.28E-20 | 4.40E-15 |
| MTUS1 | HC | old | 0.186715554 | 1.011821252 | 0.000238281 | 0.000278141 |
| TCEAL7 | HC | old | 0.182463412 | 1.380735295 | 1.05E-06 | 2.29E-05 |
| MIR181A2HG | HC | young | -0.11766374 | -0.918554143 | 1.86E-13 | 0.003038073 |
| ADD2 | HC | young | -0.128215566 | -0.913233037 | 0.000257437 | 0.011354696 |
| LINC01184 | HC | young | -0.143668887 | -0.787504723 | 2.47E-05 | 0.048896929 |
| PAK5 | HC | young | -0.145579094 | -0.84021812 | 3.07E-05 | 0.006619175 |
| SEMA5A | HC | young | -0.145746365 | -1.022494419 | 1.74E-05 | 0.0004135 |
| MAGI2 | HC | young | -0.14858099 | -1.145419857 | 9.31E-15 | 2.01E-06 |
| MAGI1 | HC | young | -0.15433579 | -0.82497971 | 2.07E-05 | 0.000386166 |
| PROX1 | HC | young | -0.198683707 | -1.358379765 | 1.45E-19 | 1.23E-08 |
| ONECUT1 | HC | young | -0.217083807 | -0.809196891 | 9.37E-11 | 0.001219516 |
| AC092691.1 | HC | young | -0.235676493 | -1.10401955 | 4.21E-10 | 5.17E-06 |
| PLA2R1 | HC | young | -0.264250939 | -1.570768416 | 3.69E-09 | 2.52E-05 |
| DAB1 | HC | young | -0.302894508 | -1.110154286 | 5.59E-08 | 5.00E-05 |
| CRADD | HC | young | -0.309098542 | -0.978111853 | 1.15E-07 | 0.009067254 |
| ZEB2 | HC | young | -0.319712539 | -1.041599037 | 2.89E-13 | 3.39E-05 |
| GRIP1 | HC | young | -0.33958225 | -1.215381223 | 4.05E-10 | 2.01E-06 |
| FRMD3 | HC | young | -0.341455137 | -0.834939306 | 0.035138306 | 0.041577347 |
| RORB | HC | young | -0.342657209 | -1.286901876 | 8.20E-33 | 1.74E-09 |
| SLC4A5 | HC | young | -0.344495744 | -0.815883734 | 0.006579249 | 0.039697317 |
| EPM2A | HC | young | -0.357383324 | -1.164366395 | 1.18E-05 | 0.002588288 |
| ARHGAP24 | HC | young | -0.369377448 | -0.976543046 | 4.81E-05 | 0.001336541 |
| GRIA4 | HC | young | -0.370322096 | -1.175682287 | 1.19E-15 | 8.09E-09 |
| ANK2 | HC | young | -0.382953798 | -0.8986104 | 6.62E-06 | 9.42E-05 |
| ZNF385D | HC | young | -0.400432355 | -3.029189701 | 4.13E-37 | 2.43E-35 |
| NRG1 | HC | young | -0.41515826 | -1.070329461 | 9.72E-13 | 8.43E-05 |
| FUS | HC | young | -0.423344686 | -1.206465218 | 1.96E-07 | 0.002444896 |
| TMEFF2 | HC | young | -0.447271298 | -0.868528661 | 0.000259518 | 0.029340464 |
| HDAC9 | HC | young | -0.456104485 | -0.874571817 | 0.000681999 | 0.012279248 |
| DOCK3 | HC | young | -0.45653294 | -0.829901502 | 0.000278676 | 0.033527615 |

| | | | | | | |
|---|---|---|---|---|---|---|
| LHFPL6 | HC | young | -0.457120672 | -0.750853571 | 1.98E-08 | 0.028016109 |
| NDST3 | HC | young | -0.492395161 | -1.130244329 | 2.79E-06 | 0.049396255 |
| ATF7IP | HC | young | -0.495848728 | -1.050662233 | 0.019765359 | 0.01308764 |
| SESTD1 | HC | young | -0.504518691 | -0.816175634 | 8.51E-05 | 0.005252465 |
| KCNH7 | HC | young | -0.515896038 | -1.368890359 | 8.40E-09 | 4.21E-07 |
| PARD3B | HC | young | -0.555592068 | -1.329546873 | 1.57E-06 | 0.001158265 |
| DACH1 | HC | young | -0.576168127 | -0.92157719 | 0.000349915 | 0.001807601 |
| PTPRR | HC | young | -0.585030249 | -0.842821535 | 0.003516774 | 0.046577638 |
| PPM1E | HC | young | -0.587863548 | -0.999256718 | 2.85E-08 | 0.001380483 |
| NCKAP5 | HC | young | -0.596194274 | -1.533238733 | 0.000284229 | 0.000160766 |
| SEMA6A | HC | young | -0.601731759 | -1.527441957 | 4.34E-16 | 2.01E-06 |
| ADCY2 | HC | young | -0.638608515 | -1.220149767 | 6.69E-09 | 5.87E-05 |
| PRR16 | HC | young | -0.64135034 | -1.488702486 | 1.46E-06 | 4.69E-05 |
| MID1 | HC | young | -0.662043916 | -0.937029231 | 0.001905956 | 0.035657442 |
| FHIT | HC | young | -0.667447459 | -1.18153641 | 1.96E-05 | 0.000993637 |
| KHDRBS2 | HC | young | -0.671845723 | -1.083582158 | 0.018399269 | 0.005311003 |
| KCNJ3 | HC | young | -0.672547167 | -1.269996891 | 6.75E-12 | 1.03E-05 |
| GALNT13 | HC | young | -0.687182827 | -1.348517302 | 2.42E-12 | 8.93E-08 |
| FRMPD4 | HC | young | -0.696073187 | -1.012088275 | 8.20E-07 | 0.001384714 |
| PDE4B | HC | young | -0.748420289 | -0.767808812 | 3.96E-06 | 0.008000105 |
| TAPT1-AS1 | HC | young | -0.816788814 | -1.516816771 | 1.96E-08 | 0.001847559 |
| AC093765.3 | HC | young | -0.877153364 | -1.591042446 | 6.22E-09 | 7.13E-05 |
| SEMA6A-AS1 | HC | young | -0.902590722 | -1.214458744 | 0.003318019 | 0.018509984 |
| CRPPA | HC | young | -0.978211636 | -1.388477087 | 6.58E-05 | 0.015878467 |
| SGCD | HC | young | -1.072439118 | -1.948451151 | 7.10E-12 | 1.93E-09 |
| AC093765.2 | HC | young | -1.835028672 | -1.89760259 | 9.74E-11 | 2.29E-05 |
