## Supplementary material for "Interpretable Aging Signatures in Human Retinal Cell Types Revealed by Single-Cell RNA Sequencing and Sparse Logistic Regression": Table S5

Table S5: Differentially expressed genes identified through both single-cell differential expression and pseudobulk differential expression analyses across 3 retinal glial populations: Müller glial cells, astrocytes, and microglia.

| gene | CellType | Direction | sc_avg_log2FC | pb_avg_log2FC | sc_p_val_adj | pb_p_val_adj |
|---|---|---|---|---|---|---|
| HP | Astrocyte | old | 9.297749822 | 5.170743953 | 0.002764205 | 0.000406032 |
| GDF15 | Astrocyte | old | 5.610288557 | 3.561625004 | 0.028561731 | 0.002987172 |
| SLPI | Astrocyte | old | 5.466733639 | 4.004437412 | 2.07E-06 | 7.66E-09 |
| ZNF331 | Astrocyte | old | 4.222428779 | 4.685631969 | 0.000113152 | 7.46E-09 |
| RARRES1 | Astrocyte | old | 3.156426544 | 2.510590816 | 0.013395853 | 7.85E-05 |
| NNMT | Astrocyte | old | 2.989707602 | 2.753916092 | 0.000265419 | 0.00564269 |
| C11orf96 | Astrocyte | old | 2.910219992 | 2.441253186 | 0.000474774 | 0.027648534 |
| SERPINE1 | Astrocyte | old | 2.68744521 | 2.238374651 | 1.09E-08 | 0.002860663 |
| SLC19A2 | Astrocyte | old | 2.624404304 | 3.19923871 | 2.18E-05 | 0.000575549 |
| ADM | Astrocyte | old | 2.496877947 | 2.853995389 | 0.005988639 | 0.003523848 |
| NAMPT | Astrocyte | old | 2.171698947 | 2.39431368 | 0.004277391 | 0.001699342 |
| HES4 | Astrocyte | old | 2.052026117 | 2.292916265 | 0.008836074 | 0.000367484 |
| ICAM1 | Astrocyte | old | 1.733122782 | 2.078302042 | 0.039612458 | 0.005934909 |
| DNAJB1 | Astrocyte | old | 1.724200581 | 2.675247936 | 0.001466429 | 3.04E-05 |
| VASN | Astrocyte | old | 1.683468746 | 2.316951591 | 0.04252101 | 0.002815104 |
| DDIT3 | Astrocyte | old | 1.409220011 | 2.975935847 | 1.26E-06 | 6.06E-05 |
| PPP1R15A | Astrocyte | old | 1.40808929 | 2.178232768 | 0.010445355 | 0.004522458 |
| AKAP12 | Astrocyte | old | 1.314826033 | 1.903947782 | 9.52E-05 | 0.000598682 |
| LDHA | Astrocyte | old | 1.292380631 | 2.097981139 | 0.000839841 | 0.00031321 |
| WTAP | Astrocyte | old | 1.252792531 | 1.998058127 | 0.012912676 | 0.002301163 |
| MT1X | Astrocyte | old | 1.20932464 | 1.908139849 | 7.82E-07 | 0.002038431 |
| FSTL3 | Astrocyte | old | 1.20267129 | 1.955030595 | 0.009162166 | 0.003578333 |
| CEBPB | Astrocyte | old | 1.137058845 | 2.340012086 | 0.002178525 | 0.000116133 |
| GADD45B | Astrocyte | old | 0.897927849 | 2.69824898 | 6.28E-05 | 1.21E-06 |
| DUSP1 | Astrocyte | old | 0.819908888 | 2.307573464 | 0.004391897 | 0.000809532 |
| IGFBP6 | Astrocyte | old | 0.791008535 | 1.430297182 | 0.002281406 | 0.008094905 |
| FTH1 | Astrocyte | old | 0.786065104 | 1.37272467 | 0.012117585 | 0.03518766 |
| BHLHE41 | Astrocyte | old | 0.678495417 | 1.618283186 | 0.010224511 | 0.001384526 |
| RPS3 | Astrocyte | old | 0.588878045 | 1.337070103 | 0.001974601 | 0.005398962 |
| SNHG16 | Astrocyte | old | 0.573276282 | 1.963757477 | 0.012708023 | 0.007161113 |
| RPS27A | Astrocyte | old | 0.571712673 | 1.453428424 | 0.022023461 | 0.002975475 |
| HSP90AA1 | Astrocyte | old | 0.561692436 | 1.386772197 | 8.14E-05 | 0.004487627 |
| IGFBP2 | Astrocyte | old | 0.541824602 | 1.791080559 | 0.001702083 | 7.85E-05 |
| RPS18 | Astrocyte | old | 0.507927704 | 1.35756852 | 0.002167044 | 0.002339561 |
| RPS28 | Astrocyte | old | 0.495209267 | 1.386878477 | 0.000114751 | 0.002344889 |
| RPS13 | Astrocyte | old | 0.460355105 | 1.421774237 | 0.028442872 | 0.002068238 |
| SLC3A2 | Astrocyte | old | 0.430478419 | 1.75031412 | 0.004078763 | 0.001118888 |
| RPL41 | Astrocyte | old | 0.41508829 | 1.531651385 | 0.044725694 | 0.000832892 |
| RPS9 | Astrocyte | old | 0.383503595 | 1.612523282 | 0.031365528 | 0.000406032 |
| EIF1 | Astrocyte | old | 0.372800194 | 1.300874359 | 0.01338109 | 0.002699926 |
| RPL23A | Astrocyte | old | 0.104652489 | 1.756128804 | 0.001973246 | 5.70E-05 |
| PARD3B | Astrocyte | young | -0.325244564 | -2.011417969 | 0.001006745 | 0.000575549 |
| DAAM1 | Astrocyte | young | -0.355104158 | -1.690854217 | 0.035768614 | 0.005934909 |
| TUBA1A | Astrocyte | young | -0.384827319 | -2.537542627 | 4.05E-09 | 1.77E-07 |
| FIGN | Astrocyte | young | -0.393280703 | -2.637292736 | 0.000488085 | 0.00027959 |
| DDX5 | Astrocyte | young | -0.409074001 | -1.072233722 | 5.50E-06 | 0.035799508 |
| UNC5C | Astrocyte | young | -0.440556108 | -2.262794692 | 0.014697562 | 3.89E-05 |
| SOX2 | Astrocyte | young | -0.550567899 | -2.157195163 | 0.000143535 | 0.000264345 |
| NR2F1 | Astrocyte | young | -0.557260004 | -1.84171928 | 0.008621532 | 0.000451388 |
| SLCO1C1 | Astrocyte | young | -0.598718762 | -2.176899899 | 0.047876746 | 0.000451388 |
| DACH1 | Astrocyte | young | -0.617716166 | -2.143031045 | 0.02498842 | 0.002033544 |
| GRIA1 | Astrocyte | young | -0.624191031 | -2.382657605 | 0.03648664 | 2.65E-05 |
| SOX2-OT | Astrocyte | young | -0.687874648 | -1.517433851 | 0.01994628 | 0.005595086 |

| | | | | | | |
|---|---|---|---|---|---|---|
| BICD1 | Astrocyte | young | -0.699081642 | -2.650571676 | 0.000415382 | 4.34E-07 |
| SLC20A2 | Astrocyte | young | -0.767639349 | -1.586865231 | 0.049278266 | 0.014540831 |
| ITGB8 | Astrocyte | young | -0.778066964 | -1.908103442 | 0.011385047 | 0.002646255 |
| RHOJ | Astrocyte | young | -0.783179857 | -2.311514441 | 0.001098223 | 0.001118888 |
| DCLK1 | Astrocyte | young | -0.852999893 | -2.005635352 | 0.007885729 | 2.84E-06 |
| KLHDC8A | Astrocyte | young | -0.861072141 | -2.671575325 | 0.000497732 | 0.010038994 |
| LRRTM4 | Astrocyte | young | -0.943048567 | -2.366219568 | 0.001017076 | 0.011533099 |
| DCLK2 | Astrocyte | young | -0.960054842 | -2.182500591 | 0.002803983 | 9.94E-05 |
| ANGPT1 | Astrocyte | young | -1.040151663 | -2.656612438 | 0.001144517 | 0.00027959 |
| SPON1 | Astrocyte | young | -1.048921926 | -2.504825285 | 2.04E-05 | 0.000575549 |
| ID4 | Astrocyte | young | -1.053177248 | -1.572135219 | 0.017244692 | 0.025898689 |
| AQP4 | Astrocyte | young | -1.09730771 | -2.359046251 | 0.00465567 | 0.00023674 |
| ZIC1 | Astrocyte | young | -1.098009971 | -1.690364469 | 0.04050945 | 0.000896909 |
| SEMA6A | Astrocyte | young | -1.197558402 | -1.999562247 | 0.002924626 | 0.00108215 |
| PDE3A | Astrocyte | young | -1.278001252 | -2.597924974 | 0.013739439 | 0.00032608 |
| ACSL6 | Astrocyte | young | -1.357567417 | -2.222232712 | 0.001042634 | 0.009379515 |
| MT-ND6 | Astrocyte | young | -1.585124852 | -2.425051925 | 1.54E-08 | 0.000575549 |
| ANGPTL1 | Astrocyte | young | -1.652662881 | -2.675186096 | 2.88E-06 | 1.77E-07 |
| SEMA6D | Astrocyte | young | -1.709791282 | -2.230983066 | 0.004875616 | 0.01153514 |
| MGAT4C | Astrocyte | young | -1.777166625 | -2.598343619 | 0.004877128 | 0.00271508 |
| AL008633.1 | Astrocyte | young | -1.820157734 | -2.71021559 | 0.020865876 | 0.001086745 |
| C11orf96 | MGC | old | 5.923122905 | 2.565212427 | 5.23E-177 | 7.17E-08 |
| SPOCD1 | MGC | old | 5.727065046 | 3.573452209 | 2.97E-31 | 2.77E-06 |
| GDF15 | MGC | old | 5.573145887 | 6.687088877 | 2.56E-212 | 2.23E-61 |
| SLCO4A1-AS1 | MGC | old | 5.343100843 | 5.52495239 | 9.34E-121 | 6.35E-24 |
| ESM1 | MGC | old | 5.246813371 | 2.476191383 | 3.10E-07 | 0.03255599 |
| LINC00906 | MGC | old | 5.164883543 | 2.630379262 | 0.000654824 | 0.030617562 |
| GGT5 | MGC | old | 4.84159184 | 2.818179966 | 1.93E-38 | 2.53E-06 |
| CHI3L2 | MGC | old | 4.838403809 | 3.050195171 | 2.03E-64 | 2.66E-17 |
| SLPI | MGC | old | 4.765399856 | 3.308660766 | 2.11E-23 | 9.61E-10 |
| KLK4 | MGC | old | 4.614078227 | 2.760494991 | 4.05E-16 | 0.000330025 |
| STRA6 | MGC | old | 4.526838124 | 3.039812821 | 1.64E-29 | 6.25E-05 |
| HMOX1 | MGC | old | 4.487788818 | 2.812414935 | 4.02E-35 | 2.08E-15 |
| CXCL1 | MGC | old | 4.455712068 | 3.326342837 | 5.98E-34 | 5.27E-10 |
| SDS | MGC | old | 4.122146106 | 5.986735931 | 8.23E-104 | 1.09E-23 |
| SERPINE2 | MGC | old | 4.098700166 | 2.093180964 | 0.004742761 | 0.018175199 |
| PHLDA2 | MGC | old | 4.05585474 | 2.473025734 | 4.97E-14 | 0.000557389 |
| BDNF | MGC | old | 4.00026625 | 2.743885671 | 1.85E-23 | 1.32E-12 |
| AC009313.1 | MGC | old | 3.981511703 | 2.26936496 | 2.20E-10 | 0.000573694 |
| ANXA1 | MGC | old | 3.97863152 | 3.261275324 | 5.17E-115 | 2.80E-19 |
| RASD2 | MGC | old | 3.966832881 | 2.586873573 | 8.99E-21 | 0.000682381 |
| AC023194.3 | MGC | old | 3.950206735 | 2.746954334 | 3.30E-37 | 3.74E-05 |
| IGFBP3 | MGC | old | 3.934817029 | 2.581313121 | 3.72E-36 | 9.18E-12 |
| C3 | MGC | old | 3.926503487 | 2.591024461 | 2.51E-62 | 1.35E-13 |
| ITGB3 | MGC | old | 3.780040803 | 2.840326997 | 3.43E-50 | 7.79E-05 |
| AC004264.1 | MGC | old | 3.772622567 | 4.36336597 | 8.02E-107 | 2.76E-30 |
| SERPINE1 | MGC | old | 3.712240015 | 1.944078195 | 4.23E-128 | 0.000863566 |
| THBS1 | MGC | old | 3.700003119 | 2.054487446 | 4.30E-10 | 0.035661955 |
| IGFBP4 | MGC | old | 3.608492618 | 3.951398917 | 1.78E-84 | 1.01E-16 |
| CH25H | MGC | old | 3.597891641 | 3.634463877 | 1.75E-76 | 2.29E-16 |
| ENTHD1 | MGC | old | 3.583170066 | 2.17567871 | 1.19E-08 | 0.00241652 |
| CLDN1 | MGC | old | 3.534483753 | 3.170149982 | 3.79E-93 | 1.89E-13 |
| ISG20 | MGC | old | 3.520665372 | 3.168302696 | 1.28E-62 | 1.57E-07 |
| CXCL2 | MGC | old | 3.502317407 | 1.713658858 | 5.02E-140 | 7.30E-06 |

| | | | | | | |
|---|---|---|---|---|---|---|
| FIBCD1 | MGC | old | 3.448096139 | 2.470380541 | 5.74E-41 | 1.93E-05 |
| HECW1 | MGC | old | 3.394058862 | 1.980165239 | 1.40E-08 | 0.009116776 |
| CXCL8 | MGC | old | 3.303162058 | 0.536935608 | 0.000111873 | 0.024664835 |
| C15orf48 | MGC | old | 3.251928053 | 2.785040882 | 1.30E-16 | 5.35E-05 |
| TNFRSF18 | MGC | old | 3.249600255 | 2.075458003 | 1.61E-17 | 0.013163068 |
| CD68 | MGC | old | 3.247370055 | 2.679818371 | 4.26E-32 | 6.20E-07 |
| KRT19 | MGC | old | 3.186702615 | 3.024619597 | 8.28E-31 | 1.05E-05 |
| MCTP2 | MGC | old | 3.181987014 | 1.891513697 | 4.61E-07 | 0.004990223 |
| CHI3L1 | MGC | old | 3.12483409 | 1.723933298 | 3.08E-23 | 4.70E-08 |
| CCDC85A | MGC | old | 3.12466648 | 2.126420268 | 5.49E-05 | 0.014300018 |
| AC017002.5 | MGC | old | 3.123113021 | 2.257795389 | 5.10E-12 | 0.000699879 |
| CLEC2B | MGC | old | 3.117432169 | 2.478320936 | 1.05E-27 | 7.84E-09 |
| MYOF | MGC | old | 3.110314178 | 1.911578803 | 2.81E-05 | 0.014104606 |
| CNN2 | MGC | old | 3.110101896 | 3.139501145 | 1.93E-60 | 8.79E-09 |
| GNG2 | MGC | old | 3.091540305 | 2.799651554 | 1.56E-44 | 1.32E-07 |
| RRAD | MGC | old | 3.06198416 | 5.020496795 | 1.73E-108 | 1.80E-35 |
| IL32 | MGC | old | 3.050428932 | 2.702146601 | 3.06E-10 | 1.85E-05 |
| CYTOR | MGC | old | 3.047371594 | 2.434166411 | 3.20E-58 | 4.80E-10 |
| AEBP1 | MGC | old | 3.021870616 | 1.892387709 | 2.41E-27 | 0.000180733 |
| RARRES1 | MGC | old | 2.975231268 | 1.827714522 | 9.06E-232 | 2.15E-07 |
| HLA-DQB1 | MGC | old | 2.968360933 | 2.381851845 | 3.67E-33 | 5.19E-05 |
| S100A11 | MGC | old | 2.960962145 | 2.177852726 | 6.43E-91 | 1.46E-09 |
| CXCL12 | MGC | old | 2.948257114 | 1.812557894 | 0.000171198 | 0.008379828 |
| HLA-DRA | MGC | old | 2.92239827 | 1.87287491 | 4.11E-208 | 1.30E-05 |
| ADM | MGC | old | 2.914473902 | 2.608115561 | 4.49E-111 | 2.43E-11 |
| NPPC | MGC | old | 2.912310463 | 4.23873361 | 1.24E-61 | 6.40E-22 |
| FGFBP2 | MGC | old | 2.901845816 | 0.674403672 | 3.23E-177 | 0.025306748 |
| TUBB3 | MGC | old | 2.885856409 | 2.239737319 | 9.11E-47 | 1.46E-10 |
| TUBB6 | MGC | old | 2.882039571 | 2.503898469 | 6.16E-66 | 6.18E-08 |
| EGR2 | MGC | old | 2.880249047 | 3.128812115 | 1.58E-63 | 9.98E-21 |
| PRSS23 | MGC | old | 2.86289282 | 2.873021056 | 2.60E-90 | 4.13E-17 |
| SERINC2 | MGC | old | 2.857389953 | 2.195068429 | 1.03E-07 | 0.015605826 |
| GALNT12 | MGC | old | 2.852630226 | 2.317171149 | 1.37E-16 | 0.002052456 |
| NIBAN1 | MGC | old | 2.77373007 | 2.384140034 | 3.58E-24 | 2.79E-09 |
| KCNK15 | MGC | old | 2.758144248 | 2.480715924 | 1.72E-35 | 2.52E-08 |
| CD74 | MGC | old | 2.757856309 | 2.884662767 | 1.25E-163 | 4.69E-34 |
| RGCC | MGC | old | 2.646617373 | 2.050922574 | 5.63E-12 | 0.003340269 |
| ANGPTL4 | MGC | old | 2.627966648 | 2.01054539 | 1.53E-15 | 1.98E-06 |
| PDLIM1 | MGC | old | 2.624624705 | 2.688192921 | 2.69E-39 | 5.58E-10 |
| LOX | MGC | old | 2.606179828 | 1.962375225 | 8.25E-08 | 0.001675211 |
| CASP1 | MGC | old | 2.600672568 | 1.235514729 | 0.000114142 | 0.011963893 |
| CXCL3 | MGC | old | 2.580423291 | 1.060964724 | 1.02E-99 | 2.27E-05 |
| CHRNA3 | MGC | old | 2.522697325 | 1.693316766 | 6.12E-18 | 0.000461082 |
| GBP2 | MGC | old | 2.510293259 | 2.057815087 | 2.15E-68 | 2.07E-08 |
| DUSP4 | MGC | old | 2.509581249 | 1.582280784 | 5.30E-12 | 0.012959299 |
| G0S2 | MGC | old | 2.476240079 | 4.540256542 | 2.35E-85 | 3.57E-41 |
| BIRC3 | MGC | old | 2.472619698 | 2.541389697 | 1.15E-83 | 8.34E-13 |
| HLA-DPA1 | MGC | old | 2.464705627 | 2.002198221 | 3.52E-33 | 2.74E-08 |
| CA9 | MGC | old | 2.435951582 | 2.278830958 | 4.67E-34 | 2.56E-05 |
| SPSB1 | MGC | old | 2.419398431 | 3.604861897 | 3.90E-110 | 7.79E-24 |
| GPRC5A | MGC | old | 2.410680574 | 4.058145785 | 6.62E-143 | 2.03E-20 |
| CXCL14 | MGC | old | 2.385033754 | 2.610596944 | 2.71E-13 | 1.45E-08 |
| ZNF331 | MGC | old | 2.379643626 | 2.78876896 | 6.50E-114 | 9.85E-12 |
| HILPDA | MGC | old | 2.374187493 | 3.419047969 | 9.79E-96 | 4.62E-30 |

| | | | | | | |
|---|---|---|---|---|---|---|
| DIO3 | MGC | old | 2.334262 | 2.711645791 | 8.48E-37 | 2.01E-13 |
| OSMR | MGC | old | 2.318243728 | 1.879948179 | 2.60E-31 | 9.56E-06 |
| ARHGAP8 | MGC | old | 2.311110855 | 1.975223803 | 0.00081037 | 0.029104791 |
| SPNS2 | MGC | old | 2.246220446 | 2.155027376 | 5.27E-11 | 0.001946816 |
| NPY1R | MGC | old | 2.242244297 | 1.680081869 | 6.77E-05 | 0.009232062 |
| SOD2 | MGC | old | 2.238665953 | 3.793370101 | 7.83E-293 | 3.42E-91 |
| ERO1A | MGC | old | 2.229761868 | 2.465658933 | 1.29E-116 | 2.29E-24 |
| RASGEF1B | MGC | old | 2.226662245 | 1.575217128 | 7.59E-09 | 0.000277058 |
| SERPINA3 | MGC | old | 2.199277011 | 1.680594261 | 1.01E-115 | 0.003439737 |
| PVT1 | MGC | old | 2.197014757 | 2.189500102 | 6.90E-30 | 0.000116889 |
| OSMR-AS1 | MGC | old | 2.191141605 | 1.330754021 | 4.87E-10 | 0.025309217 |
| TNFRSF11B | MGC | old | 2.17961073 | 1.691050213 | 5.61E-12 | 0.000150666 |
| ARFGEF3 | MGC | old | 2.148572177 | 1.836684746 | 3.79E-13 | 0.001322891 |
| FOSL1 | MGC | old | 2.145285064 | 2.625684387 | 4.91E-52 | 5.35E-10 |
| LMO2 | MGC | old | 2.134582121 | 1.306292477 | 0.000445919 | 0.007417314 |
| ACTN1 | MGC | old | 2.126053132 | 2.295526837 | 9.50E-36 | 3.59E-08 |
| ANXA2 | MGC | old | 2.122494683 | 1.860108297 | 5.13E-82 | 4.00E-10 |
| HLA-DRB1 | MGC | old | 2.11788453 | 2.702557609 | 2.19E-127 | 1.74E-16 |
| LINC01115 | MGC | old | 2.116786908 | 2.030897594 | 2.60E-06 | 0.00274158 |
| AHNAK2 | MGC | old | 2.113035343 | 1.968122665 | 1.89E-21 | 0.000175924 |
| FAM177B | MGC | old | 2.104324345 | 2.071158501 | 4.25E-14 | 0.002116385 |
| FAM138D | MGC | old | 2.09356203 | 2.107595178 | 0.033596314 | 0.01728241 |
| TMSB10 | MGC | old | 2.092380194 | 1.797505428 | 3.47E-90 | 5.42E-09 |
| NPTX1 | MGC | old | 2.092188319 | 2.565149934 | 5.37E-21 | 2.41E-05 |
| KRT7 | MGC | old | 2.069803711 | 1.705364866 | 6.52E-06 | 0.013804756 |
| AL592528.1 | MGC | old | 2.043823897 | 2.095180256 | 4.91E-18 | 1.89E-05 |
| AC020892.2 | MGC | old | 2.031118322 | 2.650110994 | 0.017367242 | 2.59E-06 |
| CFB | MGC | old | 2.014433141 | 2.43985994 | 3.96E-75 | 1.36E-15 |
| DGKD | MGC | old | 2.007272155 | 2.165511616 | 4.79E-44 | 2.86E-15 |
| FOSL2 | MGC | old | 1.989477113 | 2.981328983 | 2.72E-112 | 1.41E-14 |
| ARL4C | MGC | old | 1.988480658 | 1.869468529 | 1.26E-25 | 3.59E-10 |
| S100A2 | MGC | old | 1.988401728 | 2.005013807 | 3.51E-17 | 4.95E-06 |
| EPB41L3 | MGC | old | 1.975866202 | 1.440446942 | 0.005488164 | 0.025669261 |
| ACTN2 | MGC | old | 1.975428376 | 2.134673897 | 4.72E-09 | 7.12E-05 |
| SLC39A14 | MGC | old | 1.966882072 | 2.875010701 | 3.37E-87 | 2.58E-22 |
| NMB | MGC | old | 1.953200669 | 3.397113416 | 9.65E-50 | 1.11E-26 |
| RIMBP2 | MGC | old | 1.951678134 | 2.282271646 | 6.75E-38 | 2.14E-10 |
| PLAUR | MGC | old | 1.934861819 | 2.639206301 | 4.61E-21 | 8.27E-09 |
| AC108134.2 | MGC | old | 1.933044868 | 1.862571488 | 0.004724042 | 0.022603374 |
| DUSP5 | MGC | old | 1.925961405 | 2.053314445 | 2.65E-19 | 1.10E-06 |
| S100A10 | MGC | old | 1.920264519 | 2.086920202 | 4.56E-99 | 3.11E-15 |
| CHST15 | MGC | old | 1.911041143 | 1.646920311 | 4.92E-10 | 0.011744486 |
| LST1 | MGC | old | 1.907558635 | 1.546911731 | 4.73E-16 | 0.001518831 |
| PPP1R3B | MGC | old | 1.902806694 | 2.218648985 | 4.42E-33 | 1.32E-07 |
| PLK3 | MGC | old | 1.875519817 | 2.795642786 | 1.04E-60 | 3.77E-11 |
| NAMPT | MGC | old | 1.86336416 | 3.257944958 | 3.83E-204 | 8.75E-27 |
| ICAM1 | MGC | old | 1.841802752 | 2.668832952 | 1.93E-68 | 4.10E-24 |
| CD83 | MGC | old | 1.837038777 | 2.251305901 | 5.83E-23 | 4.99E-10 |
| LEFTY2 | MGC | old | 1.823637296 | 2.068404142 | 2.51E-21 | 1.58E-05 |
| LUNAR1 | MGC | old | 1.81499442 | 1.70817947 | 3.24E-08 | 0.001737017 |
| GPRIN3 | MGC | old | 1.813652291 | 1.646452727 | 1.04E-10 | 0.000472641 |
| SLC16A6 | MGC | old | 1.804244131 | 2.785268242 | 6.63E-49 | 1.61E-15 |
| A1BG | MGC | old | 1.794474741 | 2.334544352 | 2.33E-32 | 1.43E-06 |
| SCG2 | MGC | old | 1.793161543 | 1.565547064 | 9.45E-09 | 0.000204981 |

| | | | | | | |
|---|---|---|---|---|---|---|
| EYA2 | MGC | old | 1.792405298 | 1.50604209 | 8.27E-08 | 0.013434164 |
| AHR | MGC | old | 1.784944218 | 1.748012882 | 2.36E-16 | 0.000372886 |
| SGK1 | MGC | old | 1.783262261 | 1.918094425 | 5.53E-08 | 1.94E-06 |
| TIMP1 | MGC | old | 1.782469753 | 0.996776889 | 4.92E-18 | 0.000567095 |
| SERPINF1 | MGC | old | 1.768179643 | 1.727448335 | 0.00028555 | 8.32E-05 |
| LINC02019 | MGC | old | 1.765425656 | 1.956434939 | 4.61E-11 | 0.002093301 |
| CLCF1 | MGC | old | 1.762199771 | 2.213000626 | 2.74E-36 | 1.93E-08 |
| INHBA | MGC | old | 1.75411722 | 4.238128855 | 9.35E-193 | 2.89E-63 |
| CNN1 | MGC | old | 1.752733695 | 1.597454966 | 0.012122641 | 0.019690555 |
| NTSR1 | MGC | old | 1.749704007 | 1.624321734 | 0.000819012 | 0.024421788 |
| SH3BP4 | MGC | old | 1.749678411 | 1.905786449 | 2.58E-14 | 0.000539131 |
| AC084346.1 | MGC | old | 1.745120088 | 1.502989964 | 5.42E-06 | 0.043537577 |
| IRAK2 | MGC | old | 1.743483058 | 3.024950085 | 2.39E-83 | 1.01E-15 |
| NRP1 | MGC | old | 1.739767139 | 1.255568434 | 0.001852554 | 0.00897408 |
| CLDN11 | MGC | old | 1.729440806 | 1.276129924 | 4.47E-25 | 0.007433029 |
| SPINT1-AS1 | MGC | old | 1.722432195 | 1.755181074 | 1.34E-07 | 0.02119736 |
| CITED1 | MGC | old | 1.721126299 | 1.183352818 | 5.76E-12 | 0.001702188 |
| IGFBP5 | MGC | old | 1.71825801 | 0.82582675 | 0.000170009 | 0.034635519 |
| H19 | MGC | old | 1.718026302 | 2.400469824 | 2.57E-31 | 4.71E-08 |
| LDLRAD3 | MGC | old | 1.711962576 | 2.103439909 | 4.23E-43 | 2.34E-10 |
| CD44 | MGC | old | 1.710065809 | 2.184748864 | 4.90E-181 | 9.28E-28 |
| TMEFF1 | MGC | old | 1.706177347 | 1.958532263 | 3.64E-14 | 0.000162131 |
| LINC01503 | MGC | old | 1.701296577 | 2.347712567 | 2.29E-34 | 1.24E-07 |
| LUCAT1 | MGC | old | 1.699288672 | 2.187644296 | 2.30E-29 | 4.13E-19 |
| ARID3A | MGC | old | 1.691763231 | 2.228573323 | 9.12E-06 | 0.000118372 |
| LIPG | MGC | old | 1.682004453 | 1.825835171 | 7.85E-20 | 0.001367849 |
| LIF | MGC | old | 1.681803979 | 2.200459022 | 2.67E-34 | 3.51E-08 |
| TTR | MGC | old | 1.664667358 | 2.770144758 | 1.88E-05 | 9.53E-08 |
| SQOR | MGC | old | 1.66354092 | 1.909498223 | 6.76E-32 | 9.35E-08 |
| PAWR | MGC | old | 1.660402916 | 2.377920195 | 2.41E-26 | 2.15E-08 |
| TCIM | MGC | old | 1.653272018 | 1.858958415 | 5.66E-11 | 6.54E-06 |
| CTSK | MGC | old | 1.648209686 | 1.779818817 | 0.001236092 | 0.001613921 |
| GNG12 | MGC | old | 1.647383862 | 1.296184919 | 5.00E-06 | 0.010215635 |
| ARC | MGC | old | 1.618750601 | 2.261098352 | 6.27E-10 | 1.79E-06 |
| AC009549.1 | MGC | old | 1.614269238 | 2.273470541 | 4.05E-16 | 3.86E-05 |
| HSPA6 | MGC | old | 1.613797494 | 4.864469573 | 4.78E-112 | 2.64E-35 |
| ANKRD33B | MGC | old | 1.61157796 | 1.939927556 | 5.97E-28 | 1.37E-07 |
| CRABP2 | MGC | old | 1.604948834 | 1.252119462 | 0.000242279 | 0.021198024 |
| FKBP7 | MGC | old | 1.603840601 | 2.713665381 | 2.78E-53 | 1.46E-13 |
| UCHL1 | MGC | old | 1.603217547 | 0.929903426 | 3.34E-06 | 0.010681579 |
| CBLC | MGC | old | 1.60095233 | 1.373563142 | 6.46E-07 | 0.036154012 |
| CASP4 | MGC | old | 1.591674854 | 1.355041809 | 5.16E-11 | 0.005348874 |
| SH3PXD2B | MGC | old | 1.587957638 | 2.174956731 | 2.33E-45 | 3.46E-11 |
| PCSK6 | MGC | old | 1.585110331 | 1.831828523 | 4.80E-21 | 2.68E-07 |
| SDC4 | MGC | old | 1.583721505 | 3.104257937 | 1.19E-130 | 1.65E-29 |
| BCL3 | MGC | old | 1.581478127 | 1.59218648 | 1.21E-07 | 0.006242437 |
| GAB2 | MGC | old | 1.569710149 | 2.172830673 | 1.07E-28 | 2.38E-08 |
| EMP1 | MGC | old | 1.565783412 | 1.830589996 | 4.16E-29 | 5.92E-07 |
| INAFM1 | MGC | old | 1.565397435 | 1.884798672 | 8.22E-24 | 2.54E-06 |
| MYC | MGC | old | 1.565360258 | 2.123737188 | 1.02E-15 | 3.63E-05 |
| STOM | MGC | old | 1.563155398 | 2.466847019 | 1.37E-73 | 1.87E-17 |
| ADGRG1 | MGC | old | 1.562515757 | 1.704655512 | 9.32E-30 | 2.06E-06 |
| ITGA3 | MGC | old | 1.559259569 | 1.484960007 | 9.14E-07 | 0.002101758 |
| SCD | MGC | old | 1.552948833 | 2.334243103 | 1.23E-93 | 7.59E-16 |

| | | | | | | |
|---|---|---|---|---|---|---|
| SHISA8 | MGC | old | 1.551845306 | 1.657394599 | 0.000496257 | 0.036956411 |
| HACD1 | MGC | old | 1.546422529 | 1.864900515 | 3.37E-20 | 8.28E-06 |
| FSTL3 | MGC | old | 1.546421104 | 2.409750435 | 9.97E-44 | 5.50E-11 |
| SLC10A4 | MGC | old | 1.521929427 | 0.975046746 | 4.17E-07 | 0.037191811 |
| HMGA1 | MGC | old | 1.5102086 | 1.772723179 | 2.83E-55 | 6.81E-12 |
| GFRA2 | MGC | old | 1.502179379 | 2.744338078 | 4.17E-38 | 3.97E-10 |
| RHOU | MGC | old | 1.492335683 | 2.485215927 | 1.54E-119 | 9.22E-13 |
| FKBP5 | MGC | old | 1.487135923 | 1.732975746 | 4.63E-14 | 1.65E-08 |
| PIM1 | MGC | old | 1.482967052 | 4.108148942 | 1.00E-110 | 1.62E-39 |
| MAMLD1 | MGC | old | 1.476487534 | 2.575730407 | 4.34E-42 | 2.93E-11 |
| RAB20 | MGC | old | 1.473324333 | 2.132784869 | 6.75E-34 | 1.99E-06 |
| TNFAIP3 | MGC | old | 1.469796682 | 2.716702747 | 2.14E-53 | 7.04E-15 |
| GBE1 | MGC | old | 1.464886439 | 1.21770747 | 1.06E-12 | 0.004783247 |
| PLEKHG4B | MGC | old | 1.460242636 | 1.386970994 | 7.90E-15 | 0.00237745 |
| C1QL1 | MGC | old | 1.458589459 | 1.873923773 | 9.16E-43 | 5.84E-07 |
| NSG1 | MGC | old | 1.456930449 | 1.519455292 | 7.24E-10 | 0.00275694 |
| ATP6V0D2 | MGC | old | 1.446647749 | 3.045556798 | 2.15E-80 | 9.04E-15 |
| SGMS2 | MGC | old | 1.439864133 | 1.797014472 | 7.63E-19 | 1.31E-05 |
| PRSS56 | MGC | old | 1.4386032 | 2.831408793 | 2.81E-67 | 7.28E-18 |
| NPAS2 | MGC | old | 1.430432331 | 1.166252677 | 1.38E-09 | 0.001807813 |
| TCERG1L | MGC | old | 1.427231433 | 2.131525375 | 1.36E-45 | 1.57E-06 |
| DIP2C | MGC | old | 1.42651823 | 1.64305321 | 1.02E-31 | 1.78E-18 |
| FAS | MGC | old | 1.419210003 | 1.888790473 | 2.00E-37 | 2.88E-11 |
| AC145124.1 | MGC | old | 1.417613747 | 1.466942052 | 7.79E-07 | 0.000980432 |
| WARS | MGC | old | 1.415909095 | 1.423115904 | 4.92E-05 | 6.08E-06 |
| ELF3 | MGC | old | 1.414213164 | 1.849565535 | 4.02E-12 | 0.000393712 |
| SCG5 | MGC | old | 1.40711356 | 1.186795424 | 8.26E-06 | 0.017681982 |
| TNFRSF12A | MGC | old | 1.402585488 | 1.527186532 | 4.99E-74 | 1.51E-06 |
| CBARP | MGC | old | 1.401929041 | 2.517933633 | 9.77E-25 | 2.61E-08 |
| SLC7A5 | MGC | old | 1.401734813 | 2.710172472 | 7.80E-99 | 1.79E-15 |
| S100A6 | MGC | old | 1.401292204 | 2.144745386 | 9.33E-118 | 1.15E-23 |
| SLC35F2 | MGC | old | 1.389635493 | 1.347747704 | 0.000331274 | 0.030493581 |
| TMSB4X | MGC | old | 1.388226855 | 1.044003889 | 3.98E-57 | 0.001510025 |
| NEBL | MGC | old | 1.388001193 | 1.23960062 | 0.000220002 | 0.023785013 |
| EGR3 | MGC | old | 1.387549761 | 2.430634678 | 2.10E-16 | 1.41E-07 |
| MAST1 | MGC | old | 1.386669963 | 2.011245139 | 5.88E-09 | 8.35E-06 |
| SLC19A2 | MGC | old | 1.38602648 | 3.278972966 | 1.32E-106 | 5.36E-28 |
| ESYT2 | MGC | old | 1.381840967 | 2.388459928 | 2.78E-53 | 2.05E-18 |
| CYBA | MGC | old | 1.378279004 | 1.449972969 | 3.68E-06 | 0.000142793 |
| C1R | MGC | old | 1.372545092 | 2.415439378 | 1.53E-73 | 2.38E-17 |
| TNXB | MGC | old | 1.372156113 | 1.969674876 | 9.49E-21 | 8.18E-06 |
| IER5L | MGC | old | 1.360723262 | 2.652672686 | 2.69E-127 | 8.38E-22 |
| TFRC | MGC | old | 1.359400725 | 2.794111016 | 4.48E-120 | 3.02E-13 |
| HJV | MGC | old | 1.356779138 | 2.860473316 | 1.26E-09 | 1.65E-12 |
| CHST11 | MGC | old | 1.354277433 | 1.54432388 | 3.76E-08 | 0.000141835 |
| SLC29A1 | MGC | old | 1.353842906 | 2.236807311 | 5.30E-21 | 2.83E-07 |
| NEAT1 | MGC | old | 1.346912955 | 1.832106363 | 3.34E-07 | 1.33E-33 |
| NRP2 | MGC | old | 1.337596355 | 2.496784525 | 4.39E-62 | 1.11E-12 |
| ELL2 | MGC | old | 1.334182114 | 2.271345676 | 6.17E-32 | 1.15E-12 |
| NFKB2 | MGC | old | 1.333533978 | 1.447432365 | 1.37E-08 | 0.007966871 |
| AC068234.2 | MGC | old | 1.328826794 | 1.85814296 | 2.15E-17 | 0.000175627 |
| A4GALT | MGC | old | 1.327863144 | 1.370950311 | 1.55E-06 | 0.011649958 |
| YBX3 | MGC | old | 1.327309592 | 2.282367361 | 4.86E-153 | 1.75E-26 |
| SLC5A3 | MGC | old | 1.32444153 | 1.666751844 | 2.65E-14 | 6.66E-05 |

| | | | | | | |
|---|---|---|---|---|---|---|
| RELB | MGC | old | 1.32431442 | 2.855924015 | 4.71E-86 | 5.72E-18 |
| EFNA1 | MGC | old | 1.319202506 | 2.571699361 | 1.46E-48 | 6.73E-18 |
| AC099489.1 | MGC | old | 1.309736201 | 1.459728093 | 0.007496189 | 0.004076806 |
| IL6R | MGC | old | 1.306897902 | 2.11557692 | 3.81E-23 | 7.33E-09 |
| PDLIM4 | MGC | old | 1.30227902 | 2.186659048 | 1.95E-118 | 1.45E-18 |
| MSANTD3 | MGC | old | 1.299973548 | 2.287945859 | 9.25E-52 | 1.94E-11 |
| DEPP1 | MGC | old | 1.292719161 | 2.826734606 | 1.37E-42 | 1.03E-28 |
| ZC3H12A | MGC | old | 1.275686745 | 1.306018197 | 2.44E-06 | 0.006716162 |
| UPP1 | MGC | old | 1.269182696 | 2.406347755 | 1.58E-44 | 1.27E-12 |
| TSPAN9 | MGC | old | 1.267297495 | 1.826326383 | 2.56E-19 | 1.30E-07 |
| EMP2 | MGC | old | 1.266076076 | 1.301458182 | 6.75E-05 | 0.003495204 |
| ARHGAP29-AS1 | MGC | old | 1.262288471 | 1.647001075 | 1.71E-17 | 0.000404833 |
| SMIM3 | MGC | old | 1.250169578 | 1.427992309 | 3.94E-26 | 2.67E-06 |
| TPM4 | MGC | old | 1.248316156 | 1.651440655 | 2.63E-23 | 4.54E-05 |
| C2CD2 | MGC | old | 1.244564079 | 1.745917844 | 5.50E-28 | 2.01E-05 |
| AC061992.2 | MGC | old | 1.244235486 | 1.275906058 | 3.78E-06 | 0.012127036 |
| ANKDD1A | MGC | old | 1.239185487 | 1.630362213 | 1.80E-27 | 5.02E-13 |
| SLC13A3 | MGC | old | 1.238292538 | 1.759173486 | 4.72E-11 | 0.000466305 |
| NFKBIE | MGC | old | 1.235301426 | 1.946523089 | 5.43E-30 | 3.86E-06 |
| IER3 | MGC | old | 1.229527738 | 3.045421017 | 7.97E-116 | 2.88E-22 |
| GBP1 | MGC | old | 1.228351565 | 2.121449883 | 7.77E-27 | 2.65E-10 |
| LINC01358 | MGC | old | 1.228230508 | 1.154719707 | 0.000432412 | 0.021511438 |
| AC007032.1 | MGC | old | 1.227465685 | 2.818537793 | 2.88E-49 | 3.62E-12 |
| MAST2 | MGC | old | 1.227059426 | 1.590317209 | 8.89E-14 | 4.17E-06 |
| SERPING1 | MGC | old | 1.224633633 | 2.300485288 | 1.06E-161 | 8.40E-25 |
| CDKN2B | MGC | old | 1.219475698 | 1.455195341 | 1.91E-12 | 0.000497947 |
| HLA-DMA | MGC | old | 1.218212361 | 1.631701457 | 2.18E-24 | 1.35E-06 |
| CDK18 | MGC | old | 1.217299642 | 1.291274855 | 2.43E-13 | 0.000966179 |
| OSGIN1 | MGC | old | 1.21639308 | 1.310783863 | 0.008866879 | 0.014814401 |
| AC016831.1 | MGC | old | 1.21435068 | 2.247751797 | 2.16E-21 | 4.24E-06 |
| CHMP1B | MGC | old | 1.214069253 | 3.331001429 | 1.27E-114 | 7.54E-50 |
| C2 | MGC | old | 1.213033 | 1.410231951 | 0.039189996 | 0.002766691 |
| ENDOD1 | MGC | old | 1.209916318 | 1.172784594 | 0.000508702 | 0.017754496 |
| WWTR1 | MGC | old | 1.209350582 | 1.006539025 | 0.001892579 | 0.000180663 |
| AL157394.3 | MGC | old | 1.209285466 | 1.300425764 | 0.003068359 | 0.045193229 |
| HES2 | MGC | old | 1.204944673 | 0.987420333 | 5.51E-06 | 0.018603688 |
| KIF18A | MGC | old | 1.190922379 | 1.31824719 | 0.000547493 | 0.006601105 |
| HLA-A | MGC | old | 1.188171024 | 2.331870226 | 4.05E-194 | 1.17E-39 |
| AGPAT4 | MGC | old | 1.183185587 | 1.730663693 | 2.01E-30 | 7.73E-08 |
| SLC2A3 | MGC | old | 1.173570997 | 1.886400653 | 1.20E-51 | 4.32E-09 |
| ASS1 | MGC | old | 1.173154004 | 1.838425694 | 1.11E-17 | 3.80E-05 |
| CPEB1 | MGC | old | 1.164684624 | 1.619094121 | 1.36E-09 | 7.98E-06 |
| TNFRSF10D | MGC | old | 1.164406456 | 1.174814917 | 0.000895787 | 0.036438567 |
| TSPAN2 | MGC | old | 1.162367388 | 1.473758359 | 7.83E-10 | 0.008522781 |
| AP003086.1 | MGC | old | 1.159341769 | 1.846558647 | 1.74E-18 | 3.25E-06 |
| IRF1 | MGC | old | 1.157771852 | 2.6273088 | 2.21E-51 | 1.49E-18 |
| CRACR2B | MGC | old | 1.156404861 | 1.879251782 | 2.26E-24 | 2.59E-08 |
| KLF4 | MGC | old | 1.155406131 | 1.210340316 | 0.001137754 | 0.011329656 |
| IGF1R | MGC | old | 1.148256125 | 1.32849836 | 6.02E-15 | 9.97E-09 |
| RPS6KA2 | MGC | old | 1.147385744 | 1.478290262 | 1.56E-15 | 9.86E-09 |
| GFAP | MGC | old | 1.147230943 | 2.229532346 | 1.26E-172 | 1.14E-17 |
| C1S | MGC | old | 1.143055082 | 1.817257639 | 2.14E-35 | 1.21E-09 |
| YIPF7 | MGC | old | 1.142061375 | 1.631193147 | 1.23E-07 | 0.00734924 |
| UBASH3B | MGC | old | 1.141430171 | 1.881815302 | 9.50E-11 | 7.46E-05 |

| | | | | | | |
|---|---|---|---|---|---|---|
| STBD1 | MGC | old | 1.13371175 | 1.319769318 | 2.18E-09 | 0.010535971 |
| ACVR1 | MGC | old | 1.132945344 | 1.288670957 | 2.39E-07 | 0.000183306 |
| DYRK3 | MGC | old | 1.131530406 | 2.811092159 | 1.16E-74 | 1.45E-17 |
| KDM7A | MGC | old | 1.13120428 | 1.920880348 | 7.51E-27 | 5.84E-10 |
| LINC00958 | MGC | old | 1.130624688 | 1.663836515 | 4.21E-10 | 0.00048205 |
| CLIC4 | MGC | old | 1.127938586 | 1.51956369 | 2.54E-29 | 3.53E-09 |
| FHL2 | MGC | old | 1.125298629 | 2.070565618 | 2.40E-14 | 7.35E-06 |
| ME3 | MGC | old | 1.125246065 | 1.444014291 | 1.06E-11 | 4.93E-06 |
| MDK | MGC | old | 1.125047207 | 1.324813148 | 1.86E-40 | 8.72E-05 |
| PTTG1 | MGC | old | 1.123201812 | 1.57597129 | 8.29E-07 | 0.000231199 |
| RRAGD | MGC | old | 1.118987527 | 2.011698246 | 1.08E-50 | 2.68E-15 |
| BAG3 | MGC | old | 1.113517139 | 3.329123067 | 3.19E-85 | 1.85E-33 |
| SLC26A2 | MGC | old | 1.112203289 | 1.016371828 | 0.02180025 | 0.001125173 |
| HSPA5 | MGC | old | 1.108766391 | 1.726083101 | 2.90E-61 | 1.38E-24 |
| AL354794.1 | MGC | old | 1.103078348 | 1.663392276 | 1.75E-07 | 1.32E-05 |
| NXN | MGC | old | 1.101490384 | 2.403254402 | 2.94E-50 | 3.14E-17 |
| AC016831.5 | MGC | old | 1.092206338 | 1.643695478 | 9.37E-06 | 0.002487446 |
| PDLIM3 | MGC | old | 1.090131287 | 2.468607066 | 9.97E-153 | 2.81E-35 |
| P2RY6 | MGC | old | 1.087851196 | 1.638190768 | 1.34E-07 | 0.001095368 |
| PERP | MGC | old | 1.0877485 | 1.671766937 | 3.55E-53 | 2.37E-11 |
| LXN | MGC | old | 1.08575393 | 2.067939199 | 1.26E-32 | 1.85E-13 |
| SERPINA5 | MGC | old | 1.083692828 | 1.519784979 | 1.19E-16 | 0.00312551 |
| MYADM | MGC | old | 1.079169895 | 2.061612808 | 1.15E-24 | 7.84E-07 |
| IGFBP7 | MGC | old | 1.078943292 | 1.723284486 | 7.07E-59 | 3.79E-08 |
| JAM2 | MGC | old | 1.078904402 | 0.971185439 | 0.031162175 | 0.049890294 |
| TMEM265 | MGC | old | 1.077207858 | 2.67714651 | 9.60E-40 | 3.02E-12 |
| MLLT11 | MGC | old | 1.074591964 | 1.458928864 | 1.18E-08 | 9.02E-06 |
| SPARC | MGC | old | 1.074399195 | 1.220782445 | 2.19E-52 | 0.00019228 |
| SLC43A2 | MGC | old | 1.06753682 | 2.028032976 | 1.17E-27 | 3.20E-08 |
| WTAP | MGC | old | 1.067052442 | 2.770639008 | 7.93E-159 | 3.11E-59 |
| HBEGF | MGC | old | 1.065019754 | 3.049783346 | 8.90E-81 | 2.73E-22 |
| LMCD1 | MGC | old | 1.061544179 | 2.796356742 | 9.30E-131 | 4.81E-40 |
| TUBA1C | MGC | old | 1.059048159 | 1.601133852 | 4.36E-16 | 2.12E-06 |
| DUSP2 | MGC | old | 1.057079138 | 2.705498753 | 2.10E-19 | 1.92E-05 |
| SPRED3 | MGC | old | 1.054957177 | 1.351349949 | 0.000620202 | 0.014540309 |
| MAP2K3 | MGC | old | 1.053633354 | 2.373834035 | 2.19E-72 | 8.60E-15 |
| ENO2 | MGC | old | 1.052310567 | 1.014105369 | 4.52E-07 | 0.000339115 |
| YOD1 | MGC | old | 1.05091716 | 1.286031241 | 2.15E-06 | 0.007044314 |
| TPST1 | MGC | old | 1.049705381 | 1.860253325 | 3.03E-53 | 5.25E-13 |
| CD276 | MGC | old | 1.04962356 | 1.388256426 | 2.63E-11 | 0.001315465 |
| PARD6G-AS1 | MGC | old | 1.047922807 | 2.440629169 | 8.19E-18 | 1.21E-09 |
| AC005081.1 | MGC | old | 1.041708682 | 1.574247985 | 0.002169431 | 0.011992164 |
| TRH | MGC | old | 1.03453593 | 2.381969778 | 3.03E-94 | 2.37E-11 |
| COTL1 | MGC | old | 1.031672414 | 2.167809594 | 1.01E-91 | 1.44E-22 |
| PTP4A3 | MGC | old | 1.031348636 | 1.353874537 | 8.88E-23 | 2.97E-06 |
| IGFBP6 | MGC | old | 1.030924562 | 1.76491268 | 1.41E-57 | 2.02E-08 |
| AL358075.2 | MGC | old | 1.026808425 | 1.927856309 | 6.86E-07 | 1.20E-06 |
| GEM | MGC | old | 1.026772038 | 3.403003784 | 3.17E-227 | 1.41E-45 |
| PLEKHO1 | MGC | old | 1.025166722 | 2.673224215 | 1.23E-68 | 5.41E-17 |
| SAPCD1 | MGC | old | 1.024332151 | 1.116743176 | 0.006621443 | 0.041383584 |
| PDK3 | MGC | old | 1.023093843 | 1.142402923 | 8.36E-07 | 3.70E-05 |
| B2M | MGC | old | 1.020347996 | 1.523490419 | 9.06E-133 | 1.01E-11 |
| NOL3 | MGC | old | 1.020279782 | 0.752738462 | 3.84E-10 | 0.046784437 |
| PMP22 | MGC | old | 1.020156029 | 2.238987961 | 1.18E-129 | 6.09E-25 |

| | | | | | | |
|---|---|---|---|---|---|---|
| ARID5B | MGC | old | 1.013047411 | 2.041444494 | 2.76E-36 | 8.29E-18 |
| RTKN2 | MGC | old | 1.010587983 | 1.350212626 | 7.14E-09 | 0.004179621 |
| INSIG1 | MGC | old | 1.009992822 | 3.388516809 | 1.78E-132 | 5.16E-55 |
| MBP | MGC | old | 1.009655272 | 1.088322044 | 4.73E-07 | 0.001606563 |
| AL133453.1 | MGC | old | 1.009164481 | 1.11987672 | 9.86E-08 | 0.01216858 |
| TENT5B | MGC | old | 1.005711848 | 2.620934029 | 5.84E-44 | 1.15E-10 |
| KSR1 | MGC | old | 1.000326424 | 1.894073949 | 6.49E-22 | 4.45E-13 |
| ZNF57 | MGC | old | 0.994105534 | 1.105238916 | 5.20E-07 | 0.016927719 |
| ATP13A3 | MGC | old | 0.991368313 | 1.720022587 | 1.35E-27 | 3.21E-09 |
| XBP1 | MGC | old | 0.989496364 | 2.13671512 | 4.75E-79 | 1.99E-17 |
| RAPGEF1 | MGC | old | 0.989374998 | 1.400903048 | 7.21E-14 | 1.69E-05 |
| TTC39C-AS1 | MGC | old | 0.986996788 | 1.220286583 | 1.71E-05 | 0.009945612 |
| PLK2 | MGC | old | 0.985883551 | 1.492635919 | 0.000113146 | 0.000179565 |
| BHLHE41 | MGC | old | 0.985731132 | 2.081882027 | 2.24E-125 | 5.42E-27 |
| HLA-F | MGC | old | 0.983086347 | 1.753591186 | 1.26E-35 | 5.68E-07 |
| PIM3 | MGC | old | 0.980626734 | 2.614197046 | 6.45E-50 | 3.72E-17 |
| CARD19 | MGC | old | 0.979937113 | 2.008310353 | 3.43E-36 | 5.78E-12 |
| SERPINH1 | MGC | old | 0.979553937 | 2.415749811 | 8.83E-29 | 4.93E-13 |
| AK5 | MGC | old | 0.975019407 | 1.317736053 | 0.03474546 | 0.014531596 |
| KCNN3 | MGC | old | 0.969040979 | 2.032172679 | 2.38E-23 | 1.45E-07 |
| KRT80 | MGC | old | 0.965354473 | 0.945819384 | 9.70E-07 | 0.011853324 |
| CLIP2 | MGC | old | 0.964819474 | 1.305440766 | 1.81E-06 | 0.000542105 |
| CHAC1 | MGC | old | 0.962962505 | 1.417697643 | 0.046340114 | 0.004657114 |
| YARS | MGC | old | 0.956984796 | 1.089466584 | 0.000614729 | 0.003454034 |
| SLC45A4 | MGC | old | 0.954065449 | 1.358873812 | 0.000405522 | 0.003909823 |
| TMEM123 | MGC | old | 0.949858266 | 0.890334824 | 3.02E-05 | 0.011218955 |
| CORO1C | MGC | old | 0.947617854 | 1.502185926 | 2.38E-20 | 4.39E-05 |
| ZBTB21 | MGC | old | 0.945434637 | 2.637991536 | 1.53E-110 | 1.16E-21 |
| AC018682.1 | MGC | old | 0.94371927 | 2.371583125 | 1.14E-27 | 6.86E-14 |
| MRPS6 | MGC | old | 0.942811437 | 1.451008268 | 8.41E-35 | 2.06E-08 |
| AC021242.2 | MGC | old | 0.941898483 | 1.119436039 | 0.035684602 | 0.023636392 |
| NUAK2 | MGC | old | 0.937547179 | 2.350002305 | 5.76E-29 | 1.28E-08 |
| TRIO | MGC | old | 0.93478873 | 1.228303211 | 2.96E-08 | 2.04E-07 |
| ELOVL7 | MGC | old | 0.93474737 | 1.861792834 | 3.32E-15 | 5.48E-08 |
| HLA-B | MGC | old | 0.93348138 | 1.665215675 | 4.10E-85 | 7.08E-21 |
| CAST | MGC | old | 0.93257436 | 1.855535245 | 1.26E-70 | 1.26E-18 |
| TGIF1 | MGC | old | 0.924278053 | 2.793074434 | 4.69E-80 | 2.07E-26 |
| TUBA4A | MGC | old | 0.923639568 | 1.304228156 | 0.007326909 | 1.41E-05 |
| LHFPL6 | MGC | old | 0.921025641 | 1.085953778 | 3.94E-09 | 0.001010942 |
| ARL4D | MGC | old | 0.920634981 | 2.835214583 | 3.08E-137 | 4.94E-26 |
| ADAM17 | MGC | old | 0.918878731 | 1.306512504 | 3.02E-10 | 1.52E-07 |
| KCNE4 | MGC | old | 0.918668697 | 2.643453402 | 2.96E-33 | 1.95E-12 |
| PCBP3 | MGC | old | 0.915381154 | 1.325746459 | 5.21E-20 | 7.17E-10 |
| SLC7A1 | MGC | old | 0.909444983 | 1.999003201 | 5.36E-26 | 7.27E-08 |
| PVR | MGC | old | 0.909219998 | 1.330882937 | 0.000353004 | 0.018428008 |
| LYST | MGC | old | 0.908534341 | 1.285266893 | 1.55E-30 | 1.31E-05 |
| COL4A1 | MGC | old | 0.902220548 | 1.615257612 | 1.70E-10 | 3.56E-05 |
| MORN4 | MGC | old | 0.902195911 | 1.793824354 | 1.04E-31 | 6.62E-09 |
| PLP2 | MGC | old | 0.901205372 | 1.158548472 | 1.21E-11 | 0.000456747 |
| S100A4 | MGC | old | 0.900066615 | 1.423963888 | 0.000108142 | 1.67E-05 |
| STEAP1B | MGC | old | 0.898716501 | 1.146730682 | 6.31E-06 | 0.003865588 |
| TRAK2 | MGC | old | 0.897404375 | 1.935998393 | 2.29E-33 | 4.06E-13 |
| LINC02731 | MGC | old | 0.893979979 | 1.784588516 | 3.15E-10 | 2.73E-05 |
| LINC01128 | MGC | old | 0.891691096 | 1.754056294 | 1.26E-15 | 1.81E-06 |

| | | | | | | |
|---|---|---|---|---|---|---|
| LGALS3 | MGC | old | 0.88882838 | 1.965508641 | 7.35E-193 | 4.46E-21 |
| SYTL4 | MGC | old | 0.887755787 | 1.543119626 | 0.030660817 | 0.000361504 |
| KXD1 | MGC | old | 0.884692459 | 1.176923362 | 1.37E-05 | 0.001198992 |
| AC104695.4 | MGC | old | 0.881672331 | 1.593352234 | 0.02122068 | 0.00832813 |
| GYPC | MGC | old | 0.880753226 | 1.231371252 | 0.001278113 | 0.015545031 |
| S100A16 | MGC | old | 0.878590311 | 2.282022441 | 1.24E-102 | 6.03E-22 |
| AC092287.1 | MGC | old | 0.874969531 | 1.35835429 | 9.33E-06 | 0.009388299 |
| HIVEP2 | MGC | old | 0.87063563 | 1.269092178 | 2.74E-11 | 0.003972089 |
| PPIF | MGC | old | 0.869397983 | 2.858251357 | 2.39E-59 | 4.81E-22 |
| RAMP1 | MGC | old | 0.866814631 | 1.485990986 | 3.41E-08 | 0.000231537 |
| PFKP | MGC | old | 0.866650253 | 1.482897284 | 2.78E-42 | 1.29E-07 |
| SYNJ2 | MGC | old | 0.862831382 | 1.747700163 | 3.55E-16 | 3.06E-06 |
| CERT1 | MGC | old | 0.862387026 | 1.324907539 | 1.59E-19 | 4.15E-09 |
| GAS5 | MGC | old | 0.861358321 | 0.714085244 | 5.59E-06 | 0.010088089 |
| CCDC107 | MGC | old | 0.857087007 | 2.507295061 | 6.53E-150 | 2.26E-25 |
| TAGLN2 | MGC | old | 0.856907975 | 1.205087212 | 4.41E-43 | 5.93E-08 |
| ZYX | MGC | old | 0.855965053 | 1.187477008 | 3.11E-09 | 0.000371248 |
| NAV2-AS3 | MGC | old | 0.853630007 | 1.178289508 | 1.15E-06 | 0.01739738 |
| SLCO4A1 | MGC | old | 0.852098371 | 2.011257 | 1.69E-09 | 1.00E-05 |
| CSF1 | MGC | old | 0.851040969 | 1.87965151 | 4.30E-22 | 1.62E-07 |
| ADRB2 | MGC | old | 0.847593671 | 1.597026815 | 0.033661808 | 0.019006172 |
| WEE1 | MGC | old | 0.845627271 | 1.702333975 | 3.38E-28 | 3.46E-10 |
| FAM210A | MGC | old | 0.84540409 | 1.10800634 | 1.36E-07 | 0.000125996 |
| MDN1 | MGC | old | 0.841161277 | 1.396623113 | 1.50E-08 | 2.72E-06 |
| GPCPD1 | MGC | old | 0.838867723 | 1.197137684 | 2.56E-13 | 7.68E-05 |
| TRPM7 | MGC | old | 0.83655334 | 1.274507818 | 1.88E-07 | 3.39E-06 |
| RHOQ | MGC | old | 0.835967232 | 1.644970122 | 2.11E-55 | 1.37E-12 |
| P4HA1 | MGC | old | 0.834561029 | 1.054648284 | 6.49E-06 | 0.001315465 |
| EEPD1 | MGC | old | 0.829827611 | 1.416532554 | 1.66E-05 | 0.000545399 |
| PGK1 | MGC | old | 0.829422513 | 0.730323768 | 1.28E-12 | 0.016348526 |
| NEDD4L | MGC | old | 0.82711824 | 1.154693597 | 5.75E-06 | 2.20E-07 |
| SIAH1 | MGC | old | 0.82620197 | 1.613927702 | 1.28E-39 | 2.67E-08 |
| PCLO | MGC | old | 0.824492057 | 1.179151582 | 5.60E-11 | 4.73E-05 |
| B4GALT5 | MGC | old | 0.823197978 | 1.564606016 | 2.76E-17 | 8.73E-09 |
| SPAG9 | MGC | old | 0.823099431 | 1.765353268 | 4.65E-36 | 1.14E-16 |
| RETREG1 | MGC | old | 0.819527007 | 1.008236596 | 1.44E-06 | 0.001867785 |
| STARD4 | MGC | old | 0.818854079 | 1.190656053 | 1.04E-15 | 0.001871114 |
| NOCT | MGC | old | 0.817951166 | 1.378711404 | 4.93E-08 | 0.000175097 |
| AHI1 | MGC | old | 0.817110294 | 0.933912459 | 2.80E-08 | 2.87E-06 |
| CACNA2D2 | MGC | old | 0.816780901 | 1.516844425 | 3.87E-07 | 0.000974925 |
| MRTFA | MGC | old | 0.810498574 | 1.009372021 | 0.000968291 | 0.000417819 |
| TAGLN3 | MGC | old | 0.80935714 | 1.514331493 | 3.17E-07 | 0.000277961 |
| MDM2 | MGC | old | 0.808899885 | 1.289261824 | 1.50E-12 | 2.15E-07 |
| AL162457.2 | MGC | old | 0.806432534 | 1.362602283 | 1.41E-08 | 0.005311036 |
| DGCR6 | MGC | old | 0.804768986 | 1.945839419 | 5.50E-24 | 2.39E-07 |
| MAT2A | MGC | old | 0.801079563 | 1.951764046 | 3.58E-52 | 1.43E-11 |
| C17orf67 | MGC | old | 0.799470156 | 1.432969525 | 5.22E-08 | 0.000211633 |
| VIPR2 | MGC | old | 0.797761733 | 1.847996275 | 5.36E-29 | 2.35E-08 |
| DIP2B | MGC | old | 0.792054465 | 1.103058298 | 5.76E-06 | 6.48E-06 |
| RFX2 | MGC | old | 0.791814753 | 1.33815146 | 3.83E-06 | 2.86E-05 |
| DBF4 | MGC | old | 0.791031028 | 1.663097771 | 2.53E-08 | 2.57E-05 |
| SEMA3B | MGC | old | 0.788112973 | 2.715305801 | 5.47E-54 | 3.77E-18 |
| AC002378.1 | MGC | old | 0.785437765 | 1.914801776 | 9.55E-15 | 4.66E-05 |
| NPM3 | MGC | old | 0.785417779 | 1.062764227 | 0.003960419 | 0.019006172 |

| | | | | | | |
|---|---|---|---|---|---|---|
| AQP9 | MGC | old | 0.78439871 | 1.407056278 | 0.003253539 | 0.00068277 |
| UGDH | MGC | old | 0.784239222 | 1.524866832 | 4.16E-14 | 2.03E-05 |
| LMNA | MGC | old | 0.784237233 | 1.460489295 | 9.30E-30 | 5.06E-09 |
| HMGCS1 | MGC | old | 0.783560532 | 1.966350162 | 9.91E-22 | 1.95E-17 |
| STK40 | MGC | old | 0.783399211 | 1.546646489 | 1.64E-11 | 1.89E-06 |
| ID1 | MGC | old | 0.783360148 | 5.509368084 | 5.56E-193 | 5.41E-66 |
| BAIAP2 | MGC | old | 0.782797362 | 2.987323404 | 6.10E-96 | 7.22E-25 |
| DKK1 | MGC | old | 0.7817668 | 2.182180159 | 0.000125704 | 4.34E-07 |
| SDCBP | MGC | old | 0.779774195 | 1.922214002 | 1.29E-83 | 6.94E-26 |
| DIRAS1 | MGC | old | 0.775620177 | 1.221661523 | 0.000544776 | 0.021761681 |
| OPTN | MGC | old | 0.774857189 | 0.925635555 | 4.76E-06 | 0.000976593 |
| RCL1 | MGC | old | 0.772193306 | 1.103447755 | 0.002037107 | 0.001068037 |
| MYO1E | MGC | old | 0.771868666 | 1.30906419 | 2.13E-13 | 7.75E-07 |
| DOT1L | MGC | old | 0.770771471 | 1.41951383 | 1.64E-06 | 0.002622848 |
| LITAF | MGC | old | 0.767131174 | 2.410385634 | 4.60E-183 | 2.00E-38 |
| SOD3 | MGC | old | 0.76287724 | 1.236924447 | 4.01E-06 | 0.00141331 |
| BOC | MGC | old | 0.762661475 | 1.213635036 | 3.32E-09 | 8.45E-05 |
| PFKFB2 | MGC | old | 0.761862914 | 1.401898078 | 0.000202813 | 0.000582059 |
| RPLP0 | MGC | old | 0.760840871 | 0.651319687 | 1.07E-12 | 0.013000934 |
| FBXO32 | MGC | old | 0.759233559 | 2.759568338 | 1.39E-123 | 2.35E-27 |
| DNM3 | MGC | old | 0.757065726 | 0.906107179 | 0.037076678 | 0.003154187 |
| SIK1B | MGC | old | 0.755888929 | 1.944655952 | 7.49E-06 | 0.000177473 |
| PPP1R1A | MGC | old | 0.755282137 | 1.21309901 | 0.001337917 | 0.008487984 |
| HSPH1 | MGC | old | 0.753127676 | 2.541202157 | 1.60E-64 | 1.89E-19 |
| LGALS1 | MGC | old | 0.752084243 | 1.053624779 | 2.10E-38 | 7.15E-05 |
| AC096564.1 | MGC | old | 0.751320046 | 1.212040574 | 0.018518853 | 0.018454858 |
| PATL1 | MGC | old | 0.745091129 | 1.58987647 | 3.65E-19 | 3.17E-06 |
| USP36 | MGC | old | 0.743505947 | 2.278534785 | 2.38E-35 | 6.96E-14 |
| RAB1A | MGC | old | 0.742106517 | 1.804135346 | 3.99E-103 | 1.05E-16 |
| KLF6 | MGC | old | 0.739816002 | 1.410421587 | 1.02E-16 | 1.31E-07 |
| ITSN2 | MGC | old | 0.734468843 | 1.237659223 | 6.59E-10 | 8.39E-07 |
| PIM2 | MGC | old | 0.733696478 | 1.936129848 | 4.89E-06 | 0.000784314 |
| VAMP5 | MGC | old | 0.731995711 | 1.399100552 | 3.71E-29 | 4.49E-06 |
| SEC24D | MGC | old | 0.726244088 | 1.003680595 | 0.021105831 | 0.014551469 |
| AC009041.1 | MGC | old | 0.725921452 | 1.335373738 | 5.17E-14 | 0.000902886 |
| ZNF433 | MGC | old | 0.725911897 | 1.666274629 | 1.29E-21 | 6.02E-07 |
| SOCS3 | MGC | old | 0.724155909 | 1.762853547 | 4.70E-18 | 3.45E-09 |
| AC009053.2 | MGC | old | 0.723631635 | 2.340936033 | 1.04E-28 | 1.88E-08 |
| COL6A2 | MGC | old | 0.723365362 | 2.098428394 | 3.63E-16 | 8.10E-06 |
| RFLNB | MGC | old | 0.723098168 | 1.751413223 | 6.21E-09 | 0.000144019 |
| CTSL | MGC | old | 0.722833348 | 1.922026937 | 5.18E-30 | 3.39E-24 |
| VASN | MGC | old | 0.721579298 | 3.769306052 | 2.28E-85 | 1.01E-37 |
| CLEC18B | MGC | old | 0.72155194 | 1.012782596 | 5.09E-05 | 0.035424041 |
| LINC-PINT | MGC | old | 0.721101257 | 1.979195557 | 9.52E-32 | 4.06E-13 |
| PPP3CC | MGC | old | 0.717944226 | 0.894869209 | 0.004007722 | 2.28E-05 |
| SLC20A1 | MGC | old | 0.717856439 | 2.137500543 | 4.63E-41 | 2.38E-14 |
| TNFSF9 | MGC | old | 0.716691458 | 1.798199065 | 0.000121503 | 0.000127133 |
| P4HA2 | MGC | old | 0.715514704 | 1.075670921 | 5.47E-05 | 0.00238473 |
| AKAP12 | MGC | old | 0.714100578 | 1.080270885 | 1.26E-34 | 2.85E-05 |
| FNIP1 | MGC | old | 0.712967069 | 1.079639948 | 4.97E-11 | 9.35E-05 |
| RNASE4 | MGC | old | 0.712953502 | 1.125109253 | 0.000578223 | 0.000773601 |
| IL1R1 | MGC | old | 0.711838832 | 1.221409674 | 7.37E-21 | 2.56E-06 |
| SLC3A2 | MGC | old | 0.710787408 | 2.167968279 | 9.85E-143 | 2.98E-40 |
| SLC25A37 | MGC | old | 0.708941229 | 1.228413 | 4.96E-06 | 1.02E-07 |

| | | | | | | |
|---|---|---|---|---|---|---|
| IER5 | MGC | old | 0.707744395 | 1.452471294 | 1.15E-09 | 3.08E-06 |
| ABHD3 | MGC | old | 0.705857651 | 1.729441163 | 1.78E-11 | 2.31E-07 |
| IFRD1 | MGC | old | 0.705117704 | 1.078640399 | 0.000325567 | 2.17E-06 |
| VMP1 | MGC | old | 0.703624905 | 1.834761078 | 1.69E-28 | 1.01E-16 |
| CDC42EP3 | MGC | old | 0.70283322 | 2.226353206 | 2.45E-88 | 1.66E-14 |
| MAOB | MGC | old | 0.702671131 | 0.828002287 | 0.001910566 | 0.003039799 |
| ZNF330 | MGC | old | 0.702433058 | 1.931672759 | 8.21E-62 | 4.55E-15 |
| AC055733.2 | MGC | old | 0.702237148 | 1.72242456 | 1.01E-11 | 1.54E-08 |
| PCP4 | MGC | old | 0.702177704 | 1.711444217 | 7.30E-16 | 1.03E-06 |
| SERPINB9 | MGC | old | 0.699928773 | 2.203751066 | 3.20E-23 | 4.59E-14 |
| EIF4EBP1 | MGC | old | 0.693258132 | 1.540962471 | 1.62E-16 | 3.55E-06 |
| POLD4 | MGC | old | 0.692168896 | 1.555896836 | 7.04E-19 | 2.05E-11 |
| BIN3 | MGC | old | 0.690271847 | 1.983084263 | 3.01E-42 | 2.58E-15 |
| RYBP | MGC | old | 0.687588061 | 1.304356216 | 2.32E-19 | 9.42E-07 |
| SLC2A1 | MGC | old | 0.686578222 | 1.975637558 | 2.79E-89 | 6.37E-26 |
| ATF3 | MGC | old | 0.685102129 | 3.180096635 | 2.27E-154 | 3.15E-21 |
| CSNK1G1 | MGC | old | 0.685054396 | 1.224310365 | 2.39E-07 | 1.21E-06 |
| UAP1 | MGC | old | 0.683486578 | 1.286410815 | 8.26E-09 | 0.00028158 |
| MOSPD1 | MGC | old | 0.682999883 | 1.146392426 | 0.000142146 | 0.001806622 |
| BX470102.2 | MGC | old | 0.682793268 | 1.412615481 | 2.16E-11 | 0.000211918 |
| STX4 | MGC | old | 0.677287565 | 1.797699233 | 8.87E-30 | 1.09E-09 |
| ARSG | MGC | old | 0.674534978 | 1.035387061 | 0.045738984 | 0.003734911 |
| GADD45G | MGC | old | 0.674517207 | 1.253943985 | 0.005619203 | 6.99E-05 |
| LRRC8B | MGC | old | 0.673519518 | 1.280187157 | 0.000205558 | 0.000306003 |
| CBFB | MGC | old | 0.672302427 | 0.942819901 | 0.011933168 | 0.000465402 |
| HSPA9 | MGC | old | 0.669205835 | 1.323232597 | 7.92E-26 | 1.46E-10 |
| CXXC5 | MGC | old | 0.668682403 | 0.904241479 | 4.88E-11 | 0.000344388 |
| TTTY14 | MGC | old | 0.666404882 | 2.24506173 | 0.000734229 | 0.002920918 |
| AC083870.1 | MGC | old | 0.663009205 | 1.362953222 | 0.000287314 | 3.10E-06 |
| TPM2 | MGC | old | 0.662589716 | 1.461442175 | 7.34E-10 | 0.000622925 |
| KLF10 | MGC | old | 0.660145793 | 2.92122763 | 1.69E-81 | 5.15E-20 |
| SLC6A6 | MGC | old | 0.659466185 | 1.693955878 | 1.97E-16 | 3.20E-08 |
| TSKU | MGC | old | 0.659215774 | 1.443896489 | 3.79E-06 | 0.000531145 |
| ABR | MGC | old | 0.657133412 | 1.179468173 | 2.79E-06 | 7.43E-06 |
| CDIP1 | MGC | old | 0.65709822 | 1.627065099 | 5.64E-21 | 3.22E-07 |
| AC058791.1 | MGC | old | 0.656465907 | 1.867912249 | 6.86E-18 | 1.73E-08 |
| RPF2 | MGC | old | 0.653312608 | 1.421400417 | 3.09E-34 | 6.19E-06 |
| SQSTM1 | MGC | old | 0.650959751 | 1.573089881 | 2.52E-84 | 1.40E-14 |
| IFITM2 | MGC | old | 0.650816814 | 1.986532778 | 2.02E-166 | 2.59E-09 |
| SAMD11 | MGC | old | 0.64931027 | 1.499512084 | 2.50E-16 | 3.39E-06 |
| BACE2 | MGC | old | 0.646860867 | 1.39076031 | 1.90E-07 | 0.001315465 |
| GCGR | MGC | old | 0.645709022 | 0.941818423 | 5.48E-06 | 0.038340624 |
| MID1 | MGC | old | 0.644956785 | 2.085034109 | 2.12E-35 | 4.15E-30 |
| GLA | MGC | old | 0.644071842 | 2.42370289 | 2.56E-31 | 6.86E-18 |
| CDCA4 | MGC | old | 0.643716182 | 1.419303161 | 0.000370442 | 0.002073585 |
| AMDHD2 | MGC | old | 0.643341539 | 1.784521662 | 3.65E-14 | 2.49E-05 |
| PUSL1 | MGC | old | 0.640683444 | 1.470715771 | 9.78E-07 | 0.001147143 |
| RAP1GAP | MGC | old | 0.638759637 | 2.055146781 | 8.08E-21 | 1.59E-08 |
| RAB32 | MGC | old | 0.638038026 | 1.79267299 | 1.01E-36 | 6.81E-10 |
| MSMO1 | MGC | old | 0.638001602 | 1.796369266 | 6.32E-30 | 6.74E-12 |
| PAXBP1 | MGC | old | 0.632502516 | 1.204570444 | 3.43E-10 | 4.12E-07 |
| LINC02669 | MGC | old | 0.630474838 | 1.431768397 | 0.00010904 | 0.007395678 |
| ANXA11 | MGC | old | 0.62962225 | 1.044172976 | 7.92E-06 | 0.000814591 |
| GABARAPL1 | MGC | old | 0.629357271 | 1.892684483 | 1.20E-57 | 5.53E-21 |

| | | | | | | |
|---|---|---|---|---|---|---|
| DUSP15 | MGC | old | 0.629171346 | 1.394698265 | 5.27E-11 | 0.002608642 |
| MAP4K5 | MGC | old | 0.62724699 | 1.258110429 | 8.75E-14 | 2.11E-10 |
| TNFRSF10B | MGC | old | 0.627192934 | 1.150756159 | 0.001124458 | 0.012177312 |
| MYH9 | MGC | old | 0.627057093 | 1.750602982 | 4.42E-29 | 3.93E-09 |
| CTNNB1 | MGC | old | 0.626380015 | 1.552850095 | 2.12E-32 | 4.34E-08 |
| BTG3 | MGC | old | 0.625266484 | 1.754820191 | 6.31E-53 | 1.48E-10 |
| MTURN | MGC | old | 0.623619153 | 1.158104724 | 5.36E-06 | 0.003367607 |
| BCL2L1 | MGC | old | 0.622791124 | 1.181835303 | 2.00E-07 | 0.001015512 |
| SH3RF2 | MGC | old | 0.622509936 | 1.259624954 | 0.004661177 | 0.004265131 |
| CCL2 | MGC | old | 0.621411261 | 2.822096318 | 2.31E-109 | 1.95E-17 |
| CST3 | MGC | old | 0.618784859 | 1.098426656 | 1.12E-31 | 2.45E-05 |
| SUPV3L1 | MGC | old | 0.616491439 | 1.560541443 | 3.33E-18 | 2.70E-07 |
| SOCS1 | MGC | old | 0.616050942 | 1.684074893 | 3.62E-15 | 1.35E-06 |
| ZNF620 | MGC | old | 0.614517518 | 1.203522915 | 2.83E-07 | 0.005125841 |
| SREBF2-AS1 | MGC | old | 0.6141984 | 2.498284632 | 1.33E-20 | 2.97E-13 |
| FN3K | MGC | old | 0.614005618 | 1.854322142 | 6.64E-22 | 8.49E-07 |
| SINHCAF | MGC | old | 0.613890577 | 1.382445111 | 3.89E-06 | 0.000316903 |
| DIRAS2 | MGC | old | 0.613529308 | 1.50552002 | 1.62E-19 | 4.70E-07 |
| ATP6V1H | MGC | old | 0.613226817 | 1.070900313 | 2.80E-11 | 3.50E-06 |
| MYL9 | MGC | old | 0.612030912 | 1.674094721 | 4.44E-73 | 6.24E-16 |
| MFSD12 | MGC | old | 0.60865058 | 1.503124566 | 5.94E-11 | 8.56E-05 |
| HSPA4L | MGC | old | 0.607657857 | 1.235107494 | 3.71E-09 | 1.67E-06 |
| VCL | MGC | old | 0.60704286 | 1.520157411 | 1.25E-20 | 9.07E-11 |
| MAFF | MGC | old | 0.606825753 | 3.544951358 | 1.96E-216 | 3.98E-27 |
| FAM162A | MGC | old | 0.606001534 | 1.317052089 | 2.63E-42 | 7.84E-06 |
| PLEKHM1 | MGC | old | 0.605010563 | 1.208400337 | 2.77E-05 | 0.000902886 |
| COL5A1 | MGC | old | 0.604974444 | 1.519141792 | 1.11E-21 | 7.54E-07 |
| AC016831.7 | MGC | old | 0.604052974 | 1.439104546 | 6.70E-11 | 3.99E-06 |
| WBP2 | MGC | old | 0.603566128 | 1.508342095 | 2.00E-32 | 1.01E-12 |
| USPL1 | MGC | old | 0.603023944 | 1.464951957 | 1.57E-11 | 1.69E-07 |
| KIAA0355 | MGC | old | 0.602375592 | 1.284874553 | 2.13E-09 | 0.000115549 |
| MKNK2 | MGC | old | 0.601349016 | 2.691201019 | 4.17E-115 | 4.22E-28 |
| CALD1 | MGC | old | 0.600442481 | 1.402276322 | 4.26E-27 | 1.93E-14 |
| LIMK2 | MGC | old | 0.598549141 | 1.474668936 | 1.04E-11 | 1.91E-06 |
| VPS18 | MGC | old | 0.598200636 | 1.322794855 | 3.52E-06 | 0.001662613 |
| ACTR3 | MGC | old | 0.597942877 | 0.927198125 | 2.06E-14 | 2.52E-05 |
| PIEZO1 | MGC | old | 0.597673702 | 1.965564272 | 7.10E-18 | 1.35E-06 |
| HLA-DPB1 | MGC | old | 0.59763644 | 1.568808139 | 5.30E-32 | 5.02E-11 |
| DNAJA4 | MGC | old | 0.597239471 | 1.488526748 | 2.47E-06 | 8.36E-05 |
| CD55 | MGC | old | 0.596118373 | 1.06167405 | 0.019603294 | 0.008812188 |
| UBALD1 | MGC | old | 0.595174165 | 1.242547061 | 0.000111823 | 0.00096847 |
| RNF114 | MGC | old | 0.594747227 | 1.428992335 | 4.52E-26 | 7.56E-08 |
| TUBA1B | MGC | old | 0.591955609 | 1.179352544 | 8.22E-23 | 6.59E-14 |
| BCL2 | MGC | old | 0.589830321 | 1.409381437 | 1.90E-19 | 1.45E-07 |
| TBC1D7 | MGC | old | 0.589688572 | 1.636302072 | 1.46E-31 | 3.96E-08 |
| RPS26 | MGC | old | 0.588941938 | 1.601433129 | 5.75E-110 | 1.56E-24 |
| HLA-E | MGC | old | 0.585716376 | 1.410681087 | 6.07E-36 | 7.30E-11 |
| PALM2-AKAP2 | MGC | old | 0.585405017 | 1.729116543 | 6.69E-14 | 1.43E-05 |
| SLC66A2 | MGC | old | 0.58471764 | 1.36459595 | 1.83E-12 | 9.27E-07 |
| ZFAND2A | MGC | old | 0.58370564 | 1.425363405 | 2.36E-14 | 2.08E-05 |
| GTF2IRD1 | MGC | old | 0.583630263 | 1.940737183 | 6.10E-14 | 4.29E-15 |
| WNT5B | MGC | old | 0.581763706 | 1.195802358 | 1.74E-06 | 1.01E-09 |
| HLA-C | MGC | old | 0.581472866 | 1.624409738 | 2.99E-96 | 1.68E-26 |
| PAK1 | MGC | old | 0.580094921 | 1.009135385 | 0.000220493 | 0.000252542 |

| | | | | | | |
|---|---|---|---|---|---|---|
| SLBP | MGC | old | 0.579033421 | 1.787910129 | 1.14E-29 | 6.23E-11 |
| DNAJB1 | MGC | old | 0.578987466 | 3.341115635 | 2.15E-131 | 6.80E-35 |
| WDFY1 | MGC | old | 0.57762337 | 1.035090218 | 0.013064127 | 0.00283074 |
| ZFHX3 | MGC | old | 0.576643636 | 1.632135356 | 2.78E-18 | 1.34E-07 |
| NINJ1 | MGC | old | 0.573601828 | 1.467369611 | 4.17E-34 | 3.43E-07 |
| FKBP4 | MGC | old | 0.572257622 | 1.80509593 | 1.43E-33 | 1.79E-11 |
| SNHG16 | MGC | old | 0.57061391 | 2.275470823 | 6.55E-124 | 2.64E-35 |
| RNMT | MGC | old | 0.570341219 | 1.366911646 | 1.16E-25 | 2.17E-12 |
| STRADB | MGC | old | 0.569080101 | 0.978493718 | 3.65E-07 | 0.00681446 |
| PMEPA1 | MGC | old | 0.568328578 | 2.256804676 | 1.80E-143 | 3.23E-32 |
| ANKRD13A | MGC | old | 0.566431298 | 1.220867396 | 6.69E-05 | 0.000487253 |
| TMEM38B | MGC | old | 0.562756767 | 1.6033028 | 4.75E-54 | 2.07E-07 |
| SNHG12 | MGC | old | 0.560517707 | 1.281564341 | 8.92E-05 | 0.000354158 |
| SDSL | MGC | old | 0.560126486 | 1.606380638 | 1.44E-44 | 4.01E-08 |
| STK32A | MGC | old | 0.55932334 | 1.146240003 | 2.52E-11 | 0.002473779 |
| WFDC2 | MGC | old | 0.558918143 | 0.791367946 | 5.37E-21 | 0.010902216 |
| EEF1A1 | MGC | old | 0.557006263 | 1.069366376 | 2.20E-70 | 2.75E-10 |
| RAB31 | MGC | old | 0.556789221 | 1.098738534 | 0.000604355 | 0.000150945 |
| ARHGEF4 | MGC | old | 0.553600728 | 1.006597627 | 1.65E-05 | 0.014481684 |
| RXRA | MGC | old | 0.553156777 | 1.352787009 | 2.85E-07 | 1.06E-05 |
| PHC2 | MGC | old | 0.552023815 | 1.074055597 | 9.31E-08 | 4.62E-07 |
| PLEKHG3 | MGC | old | 0.551907285 | 2.019246359 | 1.51E-29 | 4.35E-08 |
| GFPT2 | MGC | old | 0.551009752 | 1.623428692 | 6.29E-09 | 5.26E-05 |
| NSDHL | MGC | old | 0.54957681 | 1.525584381 | 3.51E-08 | 0.000229625 |
| AXL | MGC | old | 0.548358064 | 1.319536643 | 0.000125431 | 2.32E-06 |
| MAD2L2 | MGC | old | 0.547216418 | 1.45483573 | 3.12E-12 | 2.54E-05 |
| PLIN2 | MGC | old | 0.546625847 | 1.811122704 | 8.69E-19 | 6.59E-15 |
| SLC38A1 | MGC | old | 0.546405928 | 1.210633864 | 1.56E-26 | 5.34E-07 |
| RAB40B | MGC | old | 0.54604013 | 1.290285597 | 2.37E-08 | 0.000334529 |
| NAV1 | MGC | old | 0.545638262 | 0.933486418 | 0.03611463 | 0.004332113 |
| GPRC5C | MGC | old | 0.544960767 | 2.230621513 | 2.59E-37 | 3.73E-11 |
| SNHG5 | MGC | old | 0.542490818 | 1.512505704 | 4.30E-41 | 5.12E-16 |
| IFITM3 | MGC | old | 0.542233241 | 1.940337194 | 6.12E-197 | 3.77E-11 |
| MET | MGC | old | 0.541438639 | 1.179233431 | 1.37E-07 | 0.000501568 |
| PPP1R15A | MGC | old | 0.54133798 | 2.912049538 | 2.23E-129 | 5.36E-28 |
| CYLD | MGC | old | 0.540036073 | 1.014010059 | 0.000171857 | 8.64E-05 |
| CRYAB | MGC | old | 0.539584975 | 2.998487 | 1.03E-304 | 3.27E-31 |
| NFKBIA | MGC | old | 0.538480802 | 3.028713671 | 2.48E-93 | 3.45E-35 |
| DHRS3 | MGC | old | 0.537696898 | 2.557200853 | 1.11E-83 | 4.42E-15 |
| KDELR2 | MGC | old | 0.534262836 | 1.259920566 | 2.08E-18 | 2.44E-05 |
| KDM6B | MGC | old | 0.529486173 | 1.438592747 | 4.65E-05 | 3.55E-05 |
| SERTAD1 | MGC | old | 0.528928681 | 2.623418499 | 4.09E-82 | 2.34E-21 |
| LURAP1L | MGC | old | 0.52755709 | 1.215232488 | 0.025271297 | 0.000639541 |
| ZFAND5 | MGC | old | 0.526028441 | 1.227617716 | 4.23E-28 | 1.67E-07 |
| FGF2 | MGC | old | 0.526022578 | 1.149791773 | 7.35E-05 | 0.00151806 |
| HSPA1A | MGC | old | 0.525379346 | 3.982667716 | 5.14E-167 | 8.74E-55 |
| PDXK | MGC | old | 0.524490676 | 1.295153811 | 3.58E-15 | 8.81E-06 |
| HEXA | MGC | old | 0.524268613 | 1.275446991 | 1.23E-15 | 1.85E-07 |
| PPME1 | MGC | old | 0.519873427 | 1.077231716 | 7.08E-10 | 7.52E-06 |
| ZBTB1 | MGC | old | 0.518833551 | 1.288300186 | 8.66E-09 | 7.94E-06 |
| PDPN | MGC | old | 0.518712778 | 1.603491001 | 7.51E-61 | 6.14E-11 |
| IMPDH2 | MGC | old | 0.518689248 | 0.990142934 | 2.32E-05 | 0.003677486 |
| PSME4 | MGC | old | 0.517966595 | 1.037944078 | 1.07E-05 | 5.56E-06 |
| LRP2 | MGC | old | 0.517357338 | 1.806006631 | 1.35E-12 | 2.21E-16 |

| | | | | | | |
|---|---|---|---|---|---|---|
| ATP6V1C1 | MGC | old | 0.515950761 | 1.21855825 | 2.22E-18 | 1.16E-06 |
| AL035427.1 | MGC | old | 0.514246405 | 1.644430768 | 2.44E-07 | 4.39E-05 |
| GNA13 | MGC | old | 0.513832855 | 0.928689522 | 0.001721792 | 0.000745827 |
| AC016716.2 | MGC | old | 0.512141993 | 1.41049481 | 0.012810592 | 4.22E-05 |
| PAX8-AS1 | MGC | old | 0.512115372 | 1.215360867 | 4.70E-05 | 0.004319962 |
| CLIC1 | MGC | old | 0.511330136 | 1.055855055 | 5.39E-23 | 4.22E-07 |
| NPC2 | MGC | old | 0.510776762 | 0.557323666 | 0.004052748 | 0.01580603 |
| AL022323.4 | MGC | old | 0.509461453 | 1.527496379 | 2.76E-05 | 0.000383717 |
| MT2A | MGC | old | 0.509131903 | 2.839132171 | 5.63E-143 | 1.46E-28 |
| PPTC7 | MGC | old | 0.503486308 | 1.375355675 | 1.04E-06 | 0.00120937 |
| TPM1 | MGC | old | 0.501665707 | 1.176896369 | 3.98E-12 | 1.49E-05 |
| DCDC2 | MGC | old | 0.50149072 | 0.99570561 | 0.000460615 | 0.006442783 |
| PXYLP1 | MGC | old | 0.50143228 | 1.201676403 | 6.38E-08 | 7.59E-05 |
| AC026202.2 | MGC | old | 0.500646658 | 1.055938016 | 0.002146052 | 0.002954115 |
| CCNI | MGC | old | 0.499195333 | 0.804615204 | 1.34E-08 | 2.60E-06 |
| HSPB1 | MGC | old | 0.498308961 | 2.670764594 | 2.85E-87 | 1.65E-22 |
| GLIPR2 | MGC | old | 0.498006839 | 0.840775912 | 0.000492703 | 0.006547924 |
| LDLR | MGC | old | 0.49675975 | 2.687242075 | 2.79E-34 | 4.83E-12 |
| NEURL3 | MGC | old | 0.496341403 | 1.06618245 | 5.43E-09 | 0.000971708 |
| CARHSP1 | MGC | old | 0.495933515 | 1.093120422 | 8.33E-09 | 0.005430928 |
| AC011379.1 | MGC | old | 0.495186337 | 1.474685298 | 5.79E-06 | 0.001568316 |
| LARP4 | MGC | old | 0.4946089 | 0.834106311 | 0.045860496 | 0.002213291 |
| NFIL3 | MGC | old | 0.494127079 | 2.095890626 | 3.91E-44 | 1.01E-16 |
| NCK1 | MGC | old | 0.493355843 | 0.915122837 | 9.89E-05 | 0.000335803 |
| SH3BGRL3 | MGC | old | 0.493328182 | 0.895632267 | 2.56E-14 | 0.001017174 |
| TNFRSF14 | MGC | old | 0.49227664 | 1.073954595 | 0.017112014 | 0.01192974 |
| MANF | MGC | old | 0.491668127 | 1.141394204 | 1.67E-26 | 2.76E-06 |
| SLC25A25 | MGC | old | 0.490400663 | 0.977127935 | 0.030393835 | 0.01345117 |
| PRKAG2 | MGC | old | 0.488536484 | 1.385303057 | 4.08E-09 | 1.49E-06 |
| RAB7A | MGC | old | 0.488361601 | 1.357520109 | 2.11E-47 | 1.51E-11 |
| HPRT1 | MGC | old | 0.488306354 | 0.989674525 | 2.57E-07 | 7.21E-05 |
| ARRDC3 | MGC | old | 0.487044514 | 1.128026212 | 1.07E-26 | 7.46E-05 |
| HSF1 | MGC | old | 0.484844623 | 1.173251047 | 0.000458624 | 0.000467198 |
| STARD10 | MGC | old | 0.484729537 | 1.150312916 | 0.000128479 | 0.000444154 |
| FTL | MGC | old | 0.482457221 | 0.840384365 | 1.12E-33 | 0.000122875 |
| KDM7A-DT | MGC | old | 0.481823055 | 1.536646928 | 6.26E-09 | 0.000588429 |
| MIF | MGC | old | 0.481465021 | 0.9848621 | 2.18E-55 | 2.59E-06 |
| RRP1 | MGC | old | 0.480890358 | 0.930909263 | 0.010486907 | 0.022166821 |
| APOL2 | MGC | old | 0.478623488 | 1.148247222 | 0.020207651 | 0.003021696 |
| FNBP4 | MGC | old | 0.478080974 | 1.022422806 | 2.13E-06 | 1.67E-05 |
| ULBP2 | MGC | old | 0.47505633 | 0.940942579 | 2.47E-11 | 0.002300377 |
| RHOG | MGC | old | 0.474514809 | 0.931533516 | 0.003628089 | 0.025029409 |
| DDIT3 | MGC | old | 0.473135366 | 2.713707927 | 3.00E-193 | 2.64E-35 |
| TOPBP1 | MGC | old | 0.472471129 | 1.12330456 | 7.66E-06 | 0.002693973 |
| ARHGAP29 | MGC | old | 0.472040948 | 0.894573769 | 6.11E-06 | 3.21E-07 |
| MAPK1IP1L | MGC | old | 0.471352398 | 1.292725332 | 6.29E-23 | 7.11E-09 |
| FAM131C | MGC | old | 0.467231478 | 1.32049965 | 5.07E-06 | 0.000193941 |
| CEBPB | MGC | old | 0.465555036 | 2.536430007 | 1.11E-177 | 1.92E-35 |
| DENND5A | MGC | old | 0.46516 | 0.970570385 | 0.005320306 | 7.64E-05 |
| SLC47A2 | MGC | old | 0.46216656 | 0.836530793 | 0.001830783 | 0.030089649 |
| VSIR | MGC | old | 0.461662757 | 1.562604396 | 3.72E-06 | 8.23E-06 |
| CHD1 | MGC | old | 0.461483164 | 1.107402386 | 4.58E-06 | 4.62E-07 |
| FTH1 | MGC | old | 0.460799325 | 0.999382578 | 4.45E-72 | 1.97E-07 |
| SLC6A8 | MGC | old | 0.459706085 | 1.228375072 | 0.003345453 | 0.007253897 |

| | | | | | | |
|---|---|---|---|---|---|---|
| RPS2 | MGC | old | 0.457756972 | 1.043224435 | 5.07E-72 | 1.60E-06 |
| FERMT2 | MGC | old | 0.457585964 | 1.304373728 | 2.48E-19 | 2.62E-08 |
| CAPG | MGC | old | 0.455906677 | 1.432944698 | 1.23E-20 | 1.81E-07 |
| AK2 | MGC | old | 0.455802703 | 1.00920087 | 6.33E-05 | 0.000713959 |
| DAZAP1 | MGC | old | 0.455174135 | 1.133472848 | 3.12E-06 | 0.000315243 |
| S100B | MGC | old | 0.455130209 | 1.63397424 | 4.40E-45 | 6.95E-07 |
| CDS2 | MGC | old | 0.455083531 | 1.103995156 | 7.20E-05 | 0.000133406 |
| RSL1D1 | MGC | old | 0.45413273 | 0.847517456 | 1.14E-07 | 0.000751037 |
| GOLGA7B | MGC | old | 0.451457376 | 1.495119262 | 4.78E-05 | 0.002307154 |
| DNAJB9 | MGC | old | 0.450772075 | 1.240526227 | 9.30E-22 | 2.79E-10 |
| CACYBP | MGC | old | 0.447384284 | 1.573839525 | 1.61E-21 | 2.01E-14 |
| RAB11FIP3 | MGC | old | 0.446270923 | 1.299388786 | 1.36E-07 | 5.82E-05 |
| NR4A3 | MGC | old | 0.444470671 | 2.536624383 | 7.86E-63 | 1.21E-11 |
| EMP3 | MGC | old | 0.441741984 | 1.134151789 | 1.43E-32 | 5.47E-08 |
| DYNLL1 | MGC | old | 0.441724523 | 0.962491779 | 7.29E-24 | 1.60E-06 |
| KIF1A | MGC | old | 0.441385094 | 0.934746959 | 0.000117702 | 0.000240495 |
| IGF2R | MGC | old | 0.440205902 | 0.928172058 | 0.040868884 | 0.00102621 |
| SERGEF | MGC | old | 0.439320669 | 0.934025263 | 0.010416947 | 7.80E-05 |
| RND3 | MGC | old | 0.438018712 | 1.831972633 | 2.82E-29 | 7.23E-10 |
| PLA2G4C | MGC | old | 0.436063823 | 1.271103231 | 5.08E-21 | 4.35E-13 |
| CD63 | MGC | old | 0.433925763 | 1.126601526 | 2.73E-48 | 6.06E-09 |
| SPHK1 | MGC | old | 0.431926509 | 2.95616353 | 9.36E-148 | 5.53E-39 |
| CTTN | MGC | old | 0.431793205 | 0.953154636 | 0.001276565 | 8.77E-05 |
| NUDT16 | MGC | old | 0.431573657 | 1.524394819 | 1.94E-26 | 4.00E-06 |
| ADSSL1 | MGC | old | 0.431523615 | 1.092310581 | 1.39E-16 | 0.000101349 |
| UBA2 | MGC | old | 0.431259798 | 0.835666375 | 2.15E-05 | 0.00051619 |
| VAT1 | MGC | old | 0.429688214 | 1.4202685 | 7.11E-18 | 6.84E-05 |
| PTP4A2 | MGC | old | 0.428526923 | 0.95700711 | 2.61E-17 | 1.47E-05 |
| RABGGTB | MGC | old | 0.427976123 | 1.032856295 | 4.38E-13 | 0.000392046 |
| WSB1 | MGC | old | 0.427911687 | 1.303567087 | 4.97E-05 | 1.47E-17 |
| ANKRD9 | MGC | old | 0.427034554 | 1.448616523 | 4.37E-07 | 0.000477074 |
| LINC00592 | MGC | old | 0.426867179 | 0.862995248 | 0.00044322 | 0.018146633 |
| PLA2G12A | MGC | old | 0.425675789 | 0.879595777 | 3.88E-05 | 0.000199126 |
| C9orf16 | MGC | old | 0.425658928 | 1.34387264 | 2.47E-44 | 2.27E-09 |
| TUBB | MGC | old | 0.423865077 | 1.059127574 | 3.95E-10 | 1.96E-07 |
| ITPKB | MGC | old | 0.419708425 | 1.16811666 | 0.000177621 | 0.014378957 |
| BNIP3L | MGC | old | 0.418293677 | 0.984158571 | 5.14E-21 | 1.08E-06 |
| ATP1B3 | MGC | old | 0.417817389 | 1.521714229 | 1.59E-19 | 7.08E-12 |
| HSPA1B | MGC | old | 0.41777914 | 3.889398124 | 1.48E-142 | 5.67E-42 |
| BCAR1 | MGC | old | 0.41591274 | 2.191793003 | 1.00E-21 | 4.44E-10 |
| PNO1 | MGC | old | 0.415674878 | 0.841576319 | 0.025518232 | 0.009150315 |
| PPP1CB | MGC | old | 0.414850071 | 1.403319061 | 1.17E-45 | 5.13E-15 |
| LDHA | MGC | old | 0.414755784 | 1.146301024 | 2.42E-62 | 5.53E-07 |
| ZUP1 | MGC | old | 0.414624352 | 1.161544398 | 2.77E-06 | 0.000157895 |
| CTSD | MGC | old | 0.413853249 | 1.071740579 | 5.67E-06 | 2.90E-07 |
| EMD | MGC | old | 0.410660202 | 1.371259293 | 5.60E-22 | 3.07E-09 |
| PHACTR1 | MGC | old | 0.409603682 | 1.183563319 | 0.000255133 | 2.95E-08 |
| MAFG | MGC | old | 0.409458741 | 1.226681743 | 7.70E-10 | 0.000184404 |
| ELOVL5 | MGC | old | 0.409103997 | 2.273786249 | 7.75E-63 | 2.75E-24 |
| GRAMD1A | MGC | old | 0.409097737 | 1.304934252 | 7.63E-05 | 0.000171206 |
| CNOT9 | MGC | old | 0.406483587 | 1.14768142 | 2.54E-10 | 0.00014996 |
| ZMIZ1 | MGC | old | 0.405392054 | 1.186762757 | 2.42E-07 | 4.73E-05 |
| MVD | MGC | old | 0.403889319 | 2.130676999 | 2.10E-16 | 2.70E-12 |
| SEPHS2 | MGC | old | 0.403647353 | 1.129353864 | 1.41E-05 | 5.38E-05 |

| | | | | | | |
|---|---|---|---|---|---|---|
| AC005920.2 | MGC | old | 0.400382614 | 2.073062491 | 8.32E-18 | 6.78E-09 |
| LARP6 | MGC | old | 0.398414811 | 1.180624656 | 8.18E-10 | 9.35E-05 |
| CSRNP1 | MGC | old | 0.397502241 | 2.773232585 | 3.21E-40 | 1.23E-10 |
| NCS1 | MGC | old | 0.396862693 | 2.51283799 | 2.76E-79 | 5.01E-22 |
| SPG21 | MGC | old | 0.396737185 | 1.007618302 | 3.94E-06 | 0.000169429 |
| STK11 | MGC | old | 0.394068848 | 1.35448036 | 1.57E-06 | 1.77E-05 |
| AC109460.1 | MGC | old | 0.393577446 | 0.536867089 | 8.98E-05 | 0.030581036 |
| CHPF | MGC | old | 0.393380113 | 1.404396321 | 1.27E-18 | 3.89E-08 |
| HSD17B7 | MGC | old | 0.389100383 | 1.1553816 | 0.000456173 | 6.86E-05 |
| NEU1 | MGC | old | 0.388902888 | 1.451403181 | 7.71E-29 | 1.86E-07 |
| RPL4 | MGC | old | 0.388758145 | 0.860751353 | 1.29E-16 | 0.000601989 |
| DUSP3 | MGC | old | 0.385823101 | 1.490858726 | 1.39E-14 | 5.87E-06 |
| LRRC75A | MGC | old | 0.385239648 | 1.301773979 | 6.21E-12 | 0.001567212 |
| CLEC18A | MGC | old | 0.384833345 | 1.17978046 | 1.84E-07 | 0.00566372 |
| PLEKHB1 | MGC | old | 0.383821921 | 0.943745682 | 4.88E-07 | 3.98E-06 |
| NRBF2 | MGC | old | 0.383716582 | 1.21524928 | 3.59E-05 | 0.000114234 |
| EIF4A1 | MGC | old | 0.383566912 | 1.70983138 | 3.87E-76 | 6.67E-20 |
| MIR3681HG | MGC | old | 0.381519486 | 2.497853728 | 0.000380917 | 5.58E-14 |
| TMEM176B | MGC | old | 0.379677557 | 0.982516656 | 2.13E-18 | 0.001882743 |
| UBE2D3 | MGC | old | 0.378522974 | 1.064933667 | 1.26E-22 | 6.52E-09 |
| AC006994.2 | MGC | old | 0.377509205 | 1.630657491 | 3.86E-07 | 3.18E-07 |
| NQO2 | MGC | old | 0.376387131 | 1.309113028 | 2.56E-07 | 1.27E-08 |
| SCRG1 | MGC | old | 0.376347102 | 1.400691492 | 3.35E-58 | 1.88E-07 |
| PRNP | MGC | old | 0.376283379 | 1.258059031 | 4.93E-22 | 2.90E-13 |
| DUSP10 | MGC | old | 0.374372179 | 1.514886292 | 6.69E-18 | 6.00E-10 |
| LSM10 | MGC | old | 0.373071283 | 1.806875125 | 3.07E-66 | 6.24E-16 |
| TPT1 | MGC | old | 0.371895807 | 1.094024436 | 1.10E-68 | 3.18E-07 |
| KLHL24 | MGC | old | 0.370931434 | 1.009910802 | 0.003420871 | 1.51E-05 |
| CDC42EP1 | MGC | old | 0.370438812 | 1.63188002 | 2.81E-17 | 3.00E-07 |
| WDR1 | MGC | old | 0.369280879 | 1.154502858 | 5.79E-18 | 5.65E-05 |
| CRY1 | MGC | old | 0.368927065 | 1.25463307 | 3.67E-19 | 5.95E-06 |
| C1orf43 | MGC | old | 0.368188525 | 1.238368023 | 2.76E-29 | 9.19E-07 |
| MORF4L2 | MGC | old | 0.367960594 | 1.072170487 | 1.19E-30 | 6.10E-09 |
| UBC | MGC | old | 0.364735974 | 2.466512581 | 2.73E-221 | 5.97E-44 |
| SPP1 | MGC | old | 0.362101669 | 0.704886845 | 2.12E-08 | 0.001164483 |
| AFAP1 | MGC | old | 0.362079956 | 1.544800604 | 1.04E-27 | 1.88E-09 |
| NME2 | MGC | old | 0.361996553 | 1.022132454 | 7.94E-30 | 1.73E-07 |
| ZNF410 | MGC | old | 0.361905015 | 1.015853131 | 0.001250022 | 0.00024087 |
| HPS5 | MGC | old | 0.360962687 | 1.024793667 | 0.00074662 | 0.000558942 |
| MMP14 | MGC | old | 0.359714567 | 1.229159509 | 5.63E-05 | 0.000905473 |
| SMAD3 | MGC | old | 0.358718133 | 1.539687548 | 5.45E-20 | 1.88E-07 |
| EPB41L4A-AS1 | MGC | old | 0.353739821 | 1.162764699 | 1.44E-12 | 0.000149829 |
| HSD17B14 | MGC | old | 0.351619308 | 1.933563021 | 1.55E-46 | 7.10E-12 |
| FAM122A | MGC | old | 0.35131447 | 1.684324032 | 1.39E-28 | 2.66E-10 |
| NAF1 | MGC | old | 0.351024099 | 0.953068996 | 0.011713597 | 0.000638706 |
| SREBF2 | MGC | old | 0.349839381 | 0.975462574 | 0.00753705 | 0.000863247 |
| CYSTM1 | MGC | old | 0.346201685 | 1.592757141 | 2.66E-106 | 1.84E-23 |
| TRIB2 | MGC | old | 0.345906653 | 0.987223658 | 8.23E-09 | 0.002600201 |
| OOEP | MGC | old | 0.345821723 | 0.952802765 | 1.23E-06 | 0.008636921 |
| ZNF10 | MGC | old | 0.345564814 | 1.830669146 | 1.48E-27 | 1.86E-15 |
| NNAT | MGC | old | 0.345297491 | 1.556065834 | 1.42E-18 | 6.40E-12 |
| EPAS1 | MGC | old | 0.344073217 | 1.120118986 | 4.59E-14 | 1.70E-07 |
| MAPRE1 | MGC | old | 0.343392633 | 1.433186742 | 4.62E-24 | 4.15E-09 |
| SLC25A36 | MGC | old | 0.342676672 | 0.979751686 | 1.01E-06 | 3.24E-06 |

| | | | | | | |
|---|---|---|---|---|---|---|
| PHLDA3 | MGC | old | 0.342658815 | 1.646049897 | 5.60E-39 | 2.47E-09 |
| C1orf54 | MGC | old | 0.33674884 | 1.197202885 | 2.38E-25 | 6.80E-05 |
| GNL3 | MGC | old | 0.336140795 | 1.190151933 | 1.27E-06 | 5.76E-05 |
| GNAL | MGC | old | 0.33588758 | 2.031898 | 4.70E-08 | 3.62E-07 |
| PIP4P1 | MGC | old | 0.335707575 | 1.926922563 | 9.02E-34 | 3.81E-13 |
| ZFAS1 | MGC | old | 0.333911718 | 0.838929881 | 1.74E-23 | 0.000831461 |
| INHBA-AS1 | MGC | old | 0.333606625 | 1.264566118 | 4.22E-06 | 0.002335063 |
| HSP90AB1 | MGC | old | 0.333273318 | 0.665088155 | 1.03E-07 | 0.000786561 |
| DAPK3 | MGC | old | 0.331839617 | 1.747495196 | 8.70E-19 | 5.45E-07 |
| NEK6 | MGC | old | 0.331824889 | 1.355267823 | 2.00E-32 | 2.63E-09 |
| NAPA | MGC | old | 0.33132847 | 1.408246189 | 7.71E-29 | 4.98E-09 |
| BHLHE40 | MGC | old | 0.330711427 | 2.632121222 | 2.03E-141 | 6.00E-18 |
| PHF1 | MGC | old | 0.329230503 | 0.982552084 | 0.007799109 | 0.003750639 |
| MVK | MGC | old | 0.329011074 | 1.587024767 | 1.40E-08 | 0.000132128 |
| RPL13A | MGC | old | 0.327628972 | 0.643791613 | 1.43E-24 | 0.015862073 |
| MTHFD1L | MGC | old | 0.327498504 | 0.929585072 | 0.009875407 | 0.003980084 |
| RAB18 | MGC | old | 0.326372415 | 0.750049112 | 1.04E-06 | 0.001036241 |
| COA7 | MGC | old | 0.32529741 | 1.02011031 | 0.000130456 | 0.030629937 |
| CCL28 | MGC | old | 0.324758216 | 1.189083646 | 1.45E-11 | 0.000350388 |
| ZNRD1 | MGC | old | 0.324526401 | 1.489409904 | 2.72E-29 | 1.75E-07 |
| RELT | MGC | old | 0.321404311 | 1.500995065 | 1.65E-16 | 9.55E-05 |
| PLXNA4 | MGC | old | 0.319723424 | 0.929578531 | 0.048938524 | 0.004801266 |
| MLF1 | MGC | old | 0.317878882 | 1.117856062 | 1.30E-11 | 3.05E-06 |
| TBC1D10A | MGC | old | 0.31577993 | 1.198497014 | 0.003696509 | 0.002718387 |
| EI24 | MGC | old | 0.312810556 | 0.920852073 | 0.027467759 | 0.000183941 |
| RBM38 | MGC | old | 0.310749337 | 1.16600398 | 0.002415858 | 0.004921449 |
| FKBP1A | MGC | old | 0.309788022 | 0.862461823 | 0.001097499 | 0.002546458 |
| TKT | MGC | old | 0.306458723 | 0.695663243 | 0.000391109 | 0.003755328 |
| CHKA | MGC | old | 0.301395123 | 0.942040616 | 0.011815415 | 8.64E-05 |
| SHB | MGC | old | 0.300806893 | 1.018901533 | 0.00257433 | 0.004163403 |
| EIF3H | MGC | old | 0.300635989 | 0.693258389 | 7.88E-05 | 0.001435764 |
| NTMT1 | MGC | old | 0.299990288 | 1.54798606 | 1.78E-30 | 3.99E-08 |
| SNHG8 | MGC | old | 0.299850224 | 1.451524006 | 5.57E-47 | 3.27E-11 |
| UBALD2 | MGC | old | 0.297703962 | 1.169157718 | 7.44E-06 | 4.99E-05 |
| EIF1B | MGC | old | 0.29656539 | 1.653553161 | 9.20E-52 | 8.16E-17 |
| TIMP2 | MGC | old | 0.296022985 | 1.231058241 | 2.75E-16 | 5.84E-07 |
| ARL8B | MGC | old | 0.295723241 | 0.841550531 | 7.65E-05 | 0.000352739 |
| PAPOLA | MGC | old | 0.295505848 | 0.757085892 | 3.61E-07 | 0.000185137 |
| CAMK2N1 | MGC | old | 0.294039539 | 1.12905429 | 1.69E-10 | 3.41E-05 |
| ZNF738 | MGC | old | 0.293786998 | 1.061503319 | 0.001051867 | 0.008045557 |
| PAPSS2 | MGC | old | 0.293249223 | 1.585672561 | 0.000119069 | 2.75E-05 |
| CXorf40B | MGC | old | 0.291447651 | 1.082675375 | 0.036456679 | 0.01192974 |
| PEMT | MGC | old | 0.287333827 | 1.55491693 | 1.58E-31 | 2.63E-09 |
| TGFB1I1 | MGC | old | 0.286282194 | 1.358891786 | 2.42E-06 | 0.002067998 |
| OSER1 | MGC | old | 0.285924537 | 1.461679672 | 2.16E-31 | 3.30E-07 |
| GLUL | MGC | old | 0.284999824 | 1.153180331 | 9.05E-36 | 1.67E-15 |
| TRPM4 | MGC | old | 0.284588163 | 1.351335994 | 0.000308466 | 0.002395594 |
| TUFT1 | MGC | old | 0.283534903 | 1.336675822 | 1.81E-05 | 0.004353503 |
| COPS2 | MGC | old | 0.281776723 | 0.752098483 | 0.037919771 | 0.001597985 |
| EIF3F | MGC | old | 0.281528884 | 0.808998212 | 1.79E-06 | 0.000681601 |
| HNRNPLL | MGC | old | 0.281257111 | 0.920148457 | 0.001159051 | 0.002819023 |
| LIMS1 | MGC | old | 0.279107802 | 0.865947251 | 0.000515173 | 5.96E-07 |
| DEXI | MGC | old | 0.278180404 | 1.636248895 | 1.12E-22 | 1.24E-08 |
| ACTB | MGC | old | 0.276977506 | 1.89742879 | 1.72E-146 | 2.68E-17 |

| | | | | | | |
|---|---|---|---|---|---|---|
| SERPINB6 | MGC | old | 0.275788913 | 1.076315074 | 4.85E-16 | 3.76E-08 |
| AGFG1 | MGC | old | 0.274866925 | 0.856766579 | 0.00706508 | 0.000263985 |
| ZNF93 | MGC | old | 0.271415886 | 1.537745639 | 2.50E-06 | 0.000438692 |
| PEX13 | MGC | old | 0.270636588 | 1.260575607 | 6.01E-14 | 2.10E-06 |
| AC036108.1 | MGC | old | 0.269660686 | 1.769487223 | 1.01E-09 | 6.08E-05 |
| BLOC1S6 | MGC | old | 0.269113034 | 0.844485288 | 0.004336884 | 0.000911216 |
| MAPKAPK3 | MGC | old | 0.268994945 | 1.140226586 | 0.00164159 | 0.003104688 |
| YWHAG | MGC | old | 0.268985004 | 0.767001704 | 0.0372368 | 0.002226407 |
| ASAH1 | MGC | old | 0.268367639 | 1.154395523 | 4.27E-48 | 2.83E-06 |
| CYGB | MGC | old | 0.268078178 | 1.097113721 | 0.000222581 | 0.002795806 |
| TC2N | MGC | old | 0.266856621 | 1.155980079 | 0.003559852 | 0.003229632 |
| BMP7 | MGC | old | 0.266402502 | 1.017752897 | 0.000849887 | 0.00026272 |
| PRR13 | MGC | old | 0.265271808 | 0.764567986 | 0.001738163 | 0.00090938 |
| LAMA1 | MGC | old | 0.264150799 | 1.604026677 | 1.39E-14 | 3.03E-10 |
| DUSP8 | MGC | old | 0.264044353 | 1.465799819 | 6.18E-07 | 1.34E-05 |
| PPP2R1B | MGC | old | 0.262166674 | 1.217925762 | 1.62E-06 | 8.43E-05 |
| GAPDH | MGC | old | 0.26178947 | 1.286627201 | 2.55E-83 | 8.14E-12 |
| IAH1 | MGC | old | 0.261511252 | 1.088246308 | 3.70E-14 | 4.23E-06 |
| MAPK6 | MGC | old | 0.261326862 | 0.92932115 | 1.41E-06 | 0.000671921 |
| RAB11A | MGC | old | 0.260743184 | 0.710998529 | 0.001973538 | 0.000250355 |
| NPC1 | MGC | old | 0.26047278 | 1.057017028 | 2.90E-05 | 1.67E-06 |
| EDN2 | MGC | old | 0.260294485 | 1.915203903 | 3.05E-10 | 7.80E-06 |
| TMEM176A | MGC | old | 0.259337389 | 0.743632442 | 0.00029389 | 0.006192379 |
| GRINA | MGC | old | 0.256770921 | 1.068502539 | 3.89E-08 | 6.97E-06 |
| ECE1 | MGC | old | 0.256521353 | 0.925930998 | 3.25E-05 | 1.21E-05 |
| FXYD6 | MGC | old | 0.25536774 | 1.172119058 | 3.45E-20 | 2.14E-07 |
| DYRK4 | MGC | old | 0.251753987 | 0.924744942 | 0.049971749 | 0.00125943 |
| ADORA2B | MGC | old | 0.251532739 | 1.265261412 | 6.89E-12 | 2.12E-05 |
| EIF4EBP3 | MGC | old | 0.251373621 | 0.923051604 | 0.02779529 | 0.01229489 |
| SNX10 | MGC | old | 0.245788791 | 1.021804559 | 9.99E-31 | 2.38E-07 |
| RPL22L1 | MGC | old | 0.245213822 | 2.063474408 | 9.97E-83 | 1.15E-19 |
| CLN6 | MGC | old | 0.244930599 | 1.199595878 | 0.032900316 | 0.002681634 |
| LINC02762 | MGC | old | 0.244595207 | 1.89390408 | 3.31E-65 | 7.88E-11 |
| HSPE1 | MGC | old | 0.244262537 | 1.990471667 | 4.42E-97 | 1.72E-22 |
| MAP3K8 | MGC | old | 0.243386731 | 1.576652104 | 2.37E-10 | 1.46E-07 |
| TOM1 | MGC | old | 0.243136374 | 1.134744356 | 3.49E-05 | 0.000686315 |
| TPI1 | MGC | old | 0.242333192 | 1.310043137 | 3.55E-76 | 9.39E-10 |
| MMP24OS | MGC | old | 0.239328708 | 1.107733553 | 1.90E-11 | 1.67E-06 |
| REL | MGC | old | 0.237233347 | 1.865843703 | 6.64E-20 | 2.00E-12 |
| ODC1 | MGC | old | 0.236685365 | 0.955759851 | 3.04E-11 | 4.97E-05 |
| STAM | MGC | old | 0.235937919 | 1.017908777 | 6.57E-06 | 0.000351348 |
| PNPLA8 | MGC | old | 0.234255302 | 1.009511359 | 3.73E-09 | 1.15E-05 |
| DEDD2 | MGC | old | 0.232821874 | 1.215881608 | 1.14E-05 | 0.000733425 |
| UBE2B | MGC | old | 0.232684634 | 1.218728373 | 4.46E-43 | 1.33E-14 |
| OSGIN2 | MGC | old | 0.231006277 | 1.191947754 | 9.65E-14 | 0.000158617 |
| LINC00513 | MGC | old | 0.230666617 | 1.681710574 | 2.51E-25 | 7.09E-14 |
| RHBDD2 | MGC | old | 0.229459381 | 1.093319749 | 1.57E-06 | 3.65E-05 |
| PSD4 | MGC | old | 0.229261917 | 1.10473846 | 0.01484588 | 0.007115436 |
| ACTG1 | MGC | old | 0.227539621 | 1.717291603 | 1.07E-122 | 1.00E-22 |
| AC139149.1 | MGC | old | 0.226630069 | 1.462556716 | 1.62E-05 | 0.000623828 |
| STAT3 | MGC | old | 0.225571879 | 0.903358089 | 3.44E-11 | 0.000227441 |
| CEBPD | MGC | old | 0.223608946 | 2.0605482 | 4.99E-143 | 2.27E-19 |
| WSB2 | MGC | old | 0.220571233 | 1.193354202 | 5.58E-12 | 0.000167206 |
| GTF2H2 | MGC | old | 0.219166751 | 1.037432665 | 0.007124869 | 0.004173233 |

| | | | | | | |
|---|---|---|---|---|---|---|
| TUBB4B | MGC | old | 0.216098961 | 1.043353282 | 1.64E-11 | 9.56E-09 |
| SLC25A6 | MGC | old | 0.215855178 | 0.929949802 | 2.60E-18 | 4.49E-06 |
| HCFC1R1 | MGC | old | 0.21541303 | 0.871377875 | 0.000498056 | 0.001062703 |
| PFN1 | MGC | old | 0.213234643 | 0.704310397 | 0.001000398 | 0.000616549 |
| ZNF433-AS1 | MGC | old | 0.21235002 | 1.49825082 | 9.94E-15 | 3.23E-10 |
| SIAH2 | MGC | old | 0.212101077 | 1.550962865 | 3.85E-09 | 1.59E-05 |
| FLVCR2 | MGC | old | 0.210390984 | 1.22660916 | 0.001470362 | 0.007875839 |
| SPATA2L | MGC | old | 0.209820154 | 1.249220898 | 0.022937541 | 0.004081351 |
| GLRX2 | MGC | old | 0.209555259 | 1.201174962 | 5.80E-13 | 3.63E-05 |
| SKIL | MGC | old | 0.209219174 | 0.863245594 | 0.044766249 | 0.003946647 |
| CAV1 | MGC | old | 0.208880263 | 0.861799203 | 1.16E-18 | 3.91E-05 |
| SLC35F1 | MGC | old | 0.208309939 | 0.705053438 | 3.52E-11 | 0.003544247 |
| CLK1 | MGC | old | 0.204398511 | 1.19339732 | 4.11E-10 | 3.04E-06 |
| CALR | MGC | old | 0.202379487 | 0.770085734 | 1.02E-10 | 1.33E-05 |
| FOXK2 | MGC | old | 0.202243552 | 1.148857778 | 0.00233572 | 1.53E-05 |
| UBE2L6 | MGC | old | 0.201260046 | 0.98727225 | 1.71E-06 | 0.000646222 |
| NPVF | MGC | old | 0.200821891 | 1.401556345 | 1.43E-52 | 2.17E-06 |
| VAPA | MGC | old | 0.198755465 | 0.978767838 | 3.17E-21 | 1.60E-07 |
| FGFR1 | MGC | old | 0.19775108 | 1.364815262 | 5.13E-34 | 1.93E-10 |
| TRA2A | MGC | old | 0.197729201 | 0.946303972 | 1.19E-05 | 2.23E-06 |
| PGAM1 | MGC | old | 0.196975963 | 0.988125454 | 3.99E-19 | 6.10E-07 |
| RPS8 | MGC | old | 0.196804794 | 0.718794144 | 2.71E-28 | 0.002013749 |
| HMGB2 | MGC | old | 0.196263404 | 1.464011422 | 1.13E-47 | 1.73E-08 |
| MNT | MGC | old | 0.195382812 | 1.59262206 | 1.46E-10 | 7.74E-05 |
| FAM89B | MGC | old | 0.194646616 | 1.098746546 | 0.025503654 | 0.000734199 |
| COPZ1 | MGC | old | 0.192867311 | 1.274925215 | 2.80E-28 | 4.44E-08 |
| CAB39L | MGC | old | 0.19257337 | 1.094688046 | 3.46E-15 | 3.19E-05 |
| JOSD1 | MGC | old | 0.192106055 | 1.697159263 | 2.11E-32 | 3.98E-08 |
| ZNF385A | MGC | old | 0.190647895 | 1.186110598 | 3.47E-08 | 9.48E-07 |
| CMC4 | MGC | old | 0.187644123 | 1.02469886 | 1.28E-06 | 0.000390335 |
| SLC6A12 | MGC | old | 0.18625222 | 1.375139071 | 5.30E-13 | 0.000183941 |
| PRKAG2-AS1 | MGC | old | 0.186119666 | 1.404006502 | 5.99E-10 | 0.000110231 |
| ETF1 | MGC | old | 0.185013326 | 0.979108261 | 5.63E-11 | 0.00047922 |
| PSMB9 | MGC | old | 0.18423383 | 1.270144618 | 1.47E-17 | 6.20E-06 |
| ZNF778 | MGC | old | 0.183707099 | 1.351523508 | 2.24E-09 | 0.000816017 |
| TPD52L1 | MGC | old | 0.18227746 | 1.400593131 | 3.02E-06 | 0.00065619 |
| HSPA2 | MGC | old | 0.182125371 | 1.513269445 | 1.53E-09 | 0.000108316 |
| PXDC1 | MGC | old | 0.181957758 | 1.244359099 | 6.61E-09 | 0.00054536 |
| RIDA | MGC | old | 0.179969496 | 0.893280031 | 2.03E-06 | 0.001954755 |
| AC091181.2 | MGC | old | 0.179553269 | 1.539501386 | 0.000143785 | 0.00102003 |
| CYCS | MGC | old | 0.175902892 | 1.731493762 | 5.54E-50 | 1.47E-11 |
| CD151 | MGC | old | 0.174768183 | 0.907262331 | 1.87E-11 | 0.00061583 |
| FAM241B | MGC | old | 0.173975431 | 0.974301426 | 4.86E-06 | 0.000597915 |
| OXTR | MGC | old | 0.172781726 | 1.042663469 | 0.001001525 | 0.003459372 |
| EIF4H | MGC | old | 0.171958986 | 0.818263749 | 0.000569067 | 0.020408114 |
| SNHG29 | MGC | old | 0.171786863 | 1.03258553 | 1.91E-48 | 4.85E-07 |
| POLR2H | MGC | old | 0.170428419 | 1.242470846 | 2.34E-14 | 5.70E-07 |
| AC103591.3 | MGC | old | 0.169700376 | 2.400808652 | 1.40E-07 | 3.36E-08 |
| FLOT1 | MGC | old | 0.168978543 | 0.902838189 | 1.25E-06 | 0.00017019 |
| ZNF442 | MGC | old | 0.167806373 | 1.330305675 | 3.78E-11 | 0.001999374 |
| RPL28 | MGC | old | 0.167678982 | 0.708725036 | 6.27E-29 | 0.00440299 |
| AIMP2 | MGC | old | 0.165838684 | 1.380712431 | 0.00194933 | 1.44E-06 |
| ITPRIP | MGC | old | 0.164705039 | 1.274069717 | 2.62E-05 | 0.000773304 |
| MAP1LC3B | MGC | old | 0.162977591 | 0.870633612 | 1.54E-12 | 2.35E-07 |

| | | | | | | |
|---|---|---|---|---|---|---|
| HSP90AA1 | MGC | old | 0.162656038 | 1.678068909 | 3.37E-70 | 7.28E-19 |
| CYP51A1 | MGC | old | 0.161998807 | 0.706413284 | 0.015985265 | 0.018852496 |
| VWA1 | MGC | old | 0.160669588 | 1.077813264 | 8.69E-06 | 0.000297191 |
| TIMM10B | MGC | old | 0.159275093 | 0.807128552 | 0.017274681 | 0.012467274 |
| RPL26 | MGC | old | 0.158805331 | 0.838581435 | 5.76E-32 | 3.00E-05 |
| CDKN1A | MGC | old | 0.157973137 | 1.836877912 | 1.75E-47 | 1.20E-14 |
| TP53RK | MGC | old | 0.157196874 | 1.015859474 | 0.000925621 | 0.005451211 |
| RPL12 | MGC | old | 0.156853735 | 0.743934236 | 1.39E-18 | 0.000373036 |
| AFF4 | MGC | old | 0.155639488 | 1.143175577 | 1.89E-13 | 6.45E-10 |
| PAK1IP1 | MGC | old | 0.155410046 | 0.935371321 | 0.005963473 | 0.004211284 |
| CREM | MGC | old | 0.154040129 | 1.064860192 | 6.48E-09 | 1.97E-07 |
| RPL5 | MGC | old | 0.148875509 | 0.684467091 | 5.80E-22 | 0.002262829 |
| POLR2E | MGC | old | 0.148501615 | 0.938845722 | 0.000253382 | 0.00046234 |
| TSPAN3 | MGC | old | 0.148245436 | 0.687276696 | 1.75E-10 | 0.000564638 |
| RNF181 | MGC | old | 0.1477591 | 1.105670049 | 2.81E-22 | 1.94E-08 |
| ELMSAN1 | MGC | old | 0.146018634 | 1.82546918 | 3.42E-23 | 4.64E-12 |
| ARF1 | MGC | old | 0.14422788 | 0.763946727 | 0.001978687 | 3.01E-05 |
| RACK1 | MGC | old | 0.143642887 | 0.807568623 | 3.37E-27 | 0.000538698 |
| NUCB1 | MGC | old | 0.142350667 | 0.840043533 | 0.044456274 | 0.001781283 |
| FDFT1 | MGC | old | 0.142103639 | 0.995352421 | 0.002290985 | 6.17E-05 |
| DNAJB11 | MGC | old | 0.141322063 | 1.09497337 | 1.21E-12 | 1.74E-06 |
| HOOK2 | MGC | old | 0.139402489 | 1.639957045 | 2.93E-12 | 9.89E-14 |
| LSM12 | MGC | old | 0.139233785 | 1.053203524 | 1.62E-13 | 0.000113007 |
| CDC42SE1 | MGC | old | 0.139042363 | 1.26417175 | 8.93E-14 | 2.18E-05 |
| VPS26A | MGC | old | 0.138863309 | 0.840455756 | 5.71E-09 | 0.000837555 |
| TIPARP | MGC | old | 0.138361696 | 1.380327881 | 4.45E-13 | 4.05E-08 |
| CLDN6 | MGC | old | 0.137324007 | 0.512573831 | 0.043980863 | 0.038465142 |
| EMC9 | MGC | old | 0.136472332 | 1.025983916 | 6.85E-05 | 0.002251226 |
| UQCRC2 | MGC | old | 0.136432429 | 0.780544838 | 0.018454426 | 0.000609467 |
| CD59 | MGC | old | 0.136303449 | 1.052309261 | 1.70E-35 | 7.24E-07 |
| CLN5 | MGC | old | 0.135888039 | 0.852329391 | 3.68E-05 | 5.98E-05 |
| PEF1 | MGC | old | 0.135477241 | 1.070055911 | 1.69E-07 | 0.000321109 |
| TATDN3 | MGC | old | 0.135098761 | 0.962264272 | 0.002044606 | 0.002257877 |
| EIF1 | MGC | old | 0.134737844 | 1.39671914 | 1.56E-109 | 2.53E-16 |
| MPV17 | MGC | old | 0.133945771 | 0.878169722 | 1.92E-07 | 0.001732253 |
| GGCX | MGC | old | 0.131714962 | 1.157876806 | 3.99E-06 | 0.000183941 |
| CCNL1 | MGC | old | 0.130642483 | 1.109598805 | 1.19E-15 | 2.19E-06 |
| PTRH2 | MGC | old | 0.128949986 | 1.215499735 | 7.73E-13 | 4.08E-06 |
| SSPN | MGC | old | 0.128818729 | 0.984889896 | 0.013814618 | 0.000129771 |
| RPS12 | MGC | old | 0.127475765 | 0.944305199 | 1.05E-41 | 1.05E-05 |
| GPX4 | MGC | old | 0.127015956 | 1.060473605 | 6.42E-46 | 2.83E-05 |
| C8orf88 | MGC | old | 0.1261116 | 1.143485212 | 3.11E-06 | 0.001728835 |
| JMJD6 | MGC | old | 0.121572642 | 1.526287713 | 2.35E-14 | 1.34E-07 |
| DHX36 | MGC | old | 0.121196466 | 0.90154224 | 8.66E-07 | 5.21E-05 |
| ATP6V1D | MGC | old | 0.120104542 | 1.047814718 | 1.76E-27 | 1.71E-05 |
| RPS20 | MGC | old | 0.119318394 | 0.493454157 | 3.05E-11 | 0.047892579 |
| PDE6G | MGC | old | 0.11844869 | 1.388068417 | 0.036696296 | 2.88E-08 |
| TMUB1 | MGC | old | 0.11840568 | 1.259668707 | 1.02E-05 | 7.66E-05 |
| APBB3 | MGC | old | 0.115604724 | 1.365704095 | 9.83E-07 | 0.000253228 |
| LINC00623 | MGC | old | 0.115126491 | 0.78406886 | 0.005384133 | 0.013916069 |
| GLMP | MGC | old | 0.115042328 | 1.29036463 | 2.84E-09 | 1.48E-05 |
| MAP1LC3A | MGC | old | 0.113960196 | 1.064846803 | 4.68E-13 | 6.52E-07 |
| CLDND1 | MGC | old | 0.112951405 | 0.841706879 | 2.33E-06 | 0.001564629 |
| C4orf3 | MGC | old | 0.112682772 | 0.98873017 | 1.60E-31 | 8.33E-07 |

| | | | | | | |
|---|---|---|---|---|---|---|
| GPX3 | MGC | old | 0.111427487 | 2.214268935 | 4.90E-200 | 2.41E-24 |
| HNRNPH1 | MGC | old | 0.110547784 | 1.180653142 | 5.11E-18 | 4.65E-06 |
| UPK3BL1 | MGC | old | 0.107353211 | 1.110568294 | 0.002531014 | 0.001125173 |
| RPS4X | MGC | old | 0.104482612 | 0.785526078 | 1.05E-31 | 8.43E-06 |
| DYNLT3 | MGC | old | 0.101681826 | 0.793175687 | 0.000224454 | 0.000251058 |
| TWISTNB | MGC | old | 0.101366938 | 1.572637104 | 1.84E-13 | 2.52E-07 |
| MSH3 | MGC | young | -0.101695904 | -1.143025638 | 3.65E-31 | 4.49E-06 |
| AL589935.1 | MGC | young | -0.101742557 | -1.038012036 | 7.31E-09 | 0.006677247 |
| HS3ST1 | MGC | young | -0.103066317 | -1.202135872 | 4.10E-20 | 0.00012326 |
| MFSD6 | MGC | young | -0.104132107 | -0.892873764 | 1.69E-13 | 0.007954742 |
| SLC25A12 | MGC | young | -0.104759887 | -0.608989742 | 4.91E-17 | 0.017063798 |
| CHL1 | MGC | young | -0.10672629 | -0.648349216 | 4.29E-52 | 0.005348874 |
| SOSTDC1 | MGC | young | -0.106873144 | -1.529634737 | 5.26E-08 | 0.002615331 |
| ABI3BP | MGC | young | -0.106995906 | -2.106674324 | 1.42E-201 | 9.26E-31 |
| CBX6 | MGC | young | -0.107056463 | -0.628691681 | 3.41E-08 | 0.045283908 |
| GULP1 | MGC | young | -0.107271574 | -0.955221362 | 2.86E-80 | 1.85E-08 |
| LINC02006 | MGC | young | -0.107786197 | -1.298473099 | 4.87E-12 | 0.005373815 |
| NETO2 | MGC | young | -0.107788792 | -1.293770077 | 4.57E-43 | 6.30E-07 |
| MICU2 | MGC | young | -0.107808777 | -0.483159969 | 1.13E-20 | 0.045103001 |
| DCLK2 | MGC | young | -0.107894307 | -1.001588574 | 1.61E-59 | 1.11E-06 |
| FPGT | MGC | young | -0.108849995 | -1.144133827 | 1.20E-31 | 1.74E-05 |
| TCHP | MGC | young | -0.108876231 | -1.103758467 | 5.55E-09 | 0.001738043 |
| VPS9D1 | MGC | young | -0.109043176 | -1.182839036 | 5.04E-06 | 0.012327316 |
| ARHGAP28 | MGC | young | -0.109266955 | -1.144580277 | 4.84E-07 | 0.015769782 |
| HHAT | MGC | young | -0.1092913 | -0.980917844 | 5.80E-24 | 0.000481015 |
| EPB41L2 | MGC | young | -0.109974106 | -1.083843884 | 3.03E-157 | 2.46E-13 |
| GFRA1 | MGC | young | -0.110113628 | -0.97919665 | 4.08E-11 | 0.003022782 |
| PLAGL1 | MGC | young | -0.110211827 | -1.08002398 | 8.63E-08 | 0.042690692 |
| TUBGCP2 | MGC | young | -0.110326504 | -1.006703626 | 4.42E-27 | 0.000225844 |
| AC104596.1 | MGC | young | -0.110409804 | -1.202082983 | 2.53E-37 | 1.88E-06 |
| RAD51B | MGC | young | -0.110416041 | -0.742575186 | 1.19E-33 | 0.000552829 |
| LNPEP | MGC | young | -0.110809805 | -1.101212025 | 4.53E-45 | 3.40E-06 |
| ZNF280D | MGC | young | -0.112356621 | -0.717902393 | 7.18E-24 | 0.002955632 |
| ZNF229 | MGC | young | -0.112642029 | -0.970673104 | 1.64E-11 | 0.020991464 |
| ZNF704 | MGC | young | -0.112677202 | -1.403887297 | 4.16E-54 | 2.17E-08 |
| ATP2B2 | MGC | young | -0.113109598 | -1.019362974 | 1.08E-11 | 0.006629929 |
| SPAG16 | MGC | young | -0.114056042 | -0.623023562 | 1.61E-32 | 0.008093268 |
| ZNF678 | MGC | young | -0.114244416 | -0.768236149 | 2.29E-14 | 0.02732673 |
| EIF4E3 | MGC | young | -0.114367855 | -0.72967463 | 1.22E-17 | 0.024957686 |
| SCGB2B2 | MGC | young | -0.115490688 | -1.347289854 | 2.42E-19 | 1.77E-05 |
| ORMDL3 | MGC | young | -0.116286638 | -0.923693507 | 3.06E-14 | 0.016416123 |
| CHD4 | MGC | young | -0.116821776 | -0.810336427 | 5.77E-22 | 0.003436521 |
| LRRC6 | MGC | young | -0.116993309 | -0.818634137 | 4.35E-07 | 0.037892047 |
| AC025159.1 | MGC | young | -0.117121927 | -0.824174039 | 6.55E-24 | 0.001996056 |
| RNF170 | MGC | young | -0.117923424 | -0.886072781 | 4.47E-19 | 0.001621058 |
| ACP6 | MGC | young | -0.118212072 | -1.052022194 | 1.94E-15 | 0.011450232 |
| LEKR1 | MGC | young | -0.118708645 | -1.103920834 | 2.31E-14 | 0.002167887 |
| RAB12 | MGC | young | -0.118976073 | -0.92192345 | 2.19E-19 | 0.002409508 |
| AL645568.1 | MGC | young | -0.119331087 | -1.372878532 | 5.64E-23 | 4.35E-06 |
| NFE2L1 | MGC | young | -0.119351795 | -0.70371394 | 2.48E-11 | 0.028609577 |
| AC008056.1 | MGC | young | -0.119733618 | -1.808959661 | 3.00E-63 | 2.06E-11 |
| SLC10A7 | MGC | young | -0.120462527 | -1.632471323 | 3.28E-26 | 2.40E-05 |
| CRYBG3 | MGC | young | -0.120621645 | -0.741225601 | 7.51E-31 | 0.01338627 |
| PUS7L | MGC | young | -0.120858422 | -0.974403019 | 7.95E-26 | 0.000139802 |

| | | | | | | |
|---|---|---|---|---|---|---|
| JAKMIP2 | MGC | young | -0.121158942 | -1.144606506 | 1.13E-31 | 0.000770145 |
| AC008969.1 | MGC | young | -0.121220682 | -1.069256969 | 1.62E-16 | 0.004706757 |
| CEMIP2 | MGC | young | -0.121294747 | -0.691716481 | 1.12E-49 | 0.000306441 |
| ZC3H13 | MGC | young | -0.121744169 | -0.524578266 | 9.37E-13 | 0.039231112 |
| ENPP5 | MGC | young | -0.122120593 | -0.733924978 | 1.28E-31 | 0.000994131 |
| TRIM52 | MGC | young | -0.122226026 | -0.856008538 | 5.17E-10 | 0.016180546 |
| FANCM | MGC | young | -0.12318964 | -0.93151618 | 1.14E-05 | 0.029939176 |
| ZC4H2 | MGC | young | -0.123306461 | -1.309162638 | 9.43E-20 | 0.000568874 |
| SMC1A | MGC | young | -0.12351872 | -0.834511016 | 6.02E-17 | 0.007830622 |
| FAM228B | MGC | young | -0.124837516 | -0.663174454 | 5.81E-25 | 0.003172576 |
| ATP2B1 | MGC | young | -0.124849205 | -1.018616807 | 1.29E-67 | 0.000128466 |
| TNRC18 | MGC | young | -0.125091188 | -0.862077279 | 1.07E-18 | 0.001612721 |
| ABCA3 | MGC | young | -0.125171906 | -0.98508313 | 0.000131875 | 0.030757493 |
| TAPT1-AS1 | MGC | young | -0.125270568 | -1.234559964 | 2.56E-46 | 6.59E-07 |
| TIA1 | MGC | young | -0.125482708 | -0.627734822 | 3.64E-18 | 0.025062572 |
| STPG2 | MGC | young | -0.127824607 | -2.156386269 | 4.98E-54 | 7.70E-13 |
| PAX6 | MGC | young | -0.128032547 | -1.552294731 | 3.00E-116 | 5.15E-17 |
| LINC01004 | MGC | young | -0.128123272 | -0.928536032 | 3.73E-25 | 0.00030276 |
| CUBN | MGC | young | -0.129032249 | -0.701420352 | 3.20E-15 | 0.023060736 |
| ERMP1 | MGC | young | -0.130989214 | -1.104212219 | 2.70E-07 | 0.008093268 |
| STAC3 | MGC | young | -0.13121404 | -0.959158529 | 0.000173806 | 0.016990867 |
| ADD1 | MGC | young | -0.131532985 | -1.164995138 | 3.56E-94 | 3.18E-10 |
| GUCY1A2 | MGC | young | -0.131587668 | -1.885710364 | 1.51E-66 | 7.33E-09 |
| IL33 | MGC | young | -0.131643795 | -1.16925038 | 1.11E-30 | 0.005890382 |
| SLC44A5 | MGC | young | -0.131865202 | -1.296444191 | 2.25E-07 | 0.004257302 |
| ZNF594 | MGC | young | -0.1337319 | -0.962829431 | 2.67E-12 | 0.031789964 |
| EWSR1 | MGC | young | -0.134092399 | -0.562077538 | 6.86E-16 | 0.017489788 |
| RAD51-AS1 | MGC | young | -0.134159211 | -0.966980462 | 2.55E-12 | 0.010001122 |
| CFAP54 | MGC | young | -0.135807775 | -1.819325189 | 2.29E-42 | 1.08E-10 |
| ADGRG2 | MGC | young | -0.135819623 | -1.64271168 | 7.70E-16 | 1.64E-05 |
| DPP8 | MGC | young | -0.136506543 | -0.878123985 | 8.22E-18 | 0.004784363 |
| BAALC-AS1 | MGC | young | -0.137024109 | -1.041562976 | 1.21E-16 | 0.013053993 |
| TM9SF2 | MGC | young | -0.13769032 | -0.772602298 | 3.68E-30 | 0.000408899 |
| SEMA6D | MGC | young | -0.138611002 | -1.843932955 | 2.91E-40 | 9.00E-09 |
| NT5C2 | MGC | young | -0.13991058 | -0.969430732 | 1.01E-83 | 2.75E-06 |
| SERBP1 | MGC | young | -0.139955326 | -0.522417596 | 1.40E-25 | 0.030932429 |
| CASP8AP2 | MGC | young | -0.140199216 | -0.820477942 | 3.79E-10 | 0.015650805 |
| PRDM11 | MGC | young | -0.140219481 | -1.221903637 | 2.50E-06 | 0.008826404 |
| PHLPP1 | MGC | young | -0.140253968 | -1.147468102 | 1.01E-92 | 1.36E-09 |
| ARHGAP6 | MGC | young | -0.140365103 | -1.608094437 | 1.28E-26 | 4.49E-06 |
| FBXO3 | MGC | young | -0.140925997 | -0.651508758 | 1.98E-19 | 0.011800919 |
| AP001767.3 | MGC | young | -0.142911009 | -0.982041285 | 6.43E-11 | 0.036203675 |
| TRMT2B | MGC | young | -0.143895476 | -1.175430929 | 4.52E-05 | 0.009822288 |
| AC007098.1 | MGC | young | -0.144538858 | -1.278246254 | 1.72E-20 | 0.001226229 |
| RFTN1 | MGC | young | -0.144637934 | -0.946781672 | 5.79E-26 | 0.000725627 |
| NEIL1 | MGC | young | -0.145137717 | -1.02685485 | 2.78E-06 | 0.046168855 |
| FUT10 | MGC | young | -0.145771434 | -0.784729243 | 5.60E-39 | 0.005780816 |
| CC2D2B | MGC | young | -0.146533109 | -1.850409296 | 1.07E-22 | 8.82E-06 |
| TMEM80 | MGC | young | -0.146661472 | -1.091953487 | 2.94E-10 | 0.002016193 |
| AC068051.1 | MGC | young | -0.147856134 | -1.379002538 | 1.06E-09 | 0.003156275 |
| KCTD8 | MGC | young | -0.14907435 | -0.824914124 | 8.97E-07 | 0.035128617 |
| CCDC30 | MGC | young | -0.149285386 | -1.018830438 | 6.12E-15 | 0.008412086 |
| FLT1 | MGC | young | -0.150116476 | -0.478360323 | 1.91E-63 | 0.005599645 |
| TECPR2 | MGC | young | -0.150448804 | -0.879741839 | 2.60E-14 | 0.005614354 |

| | | | | | | |
|---|---|---|---|---|---|---|
| GTF2I | MGC | young | -0.151882198 | -0.92049379 | 1.36E-77 | 9.49E-07 |
| LRRC4C | MGC | young | -0.152606383 | -1.244634542 | 2.99E-127 | 8.27E-11 |
| CROT | MGC | young | -0.153629844 | -0.889598787 | 5.67E-27 | 0.009433921 |
| DCLK1 | MGC | young | -0.154333304 | -0.648784421 | 5.90E-59 | 0.001513025 |
| AC007262.2 | MGC | young | -0.154681649 | -1.012814791 | 2.08E-06 | 0.023560685 |
| RIC3 | MGC | young | -0.155005651 | -0.545992051 | 2.15E-18 | 0.033501179 |
| LINC00240 | MGC | young | -0.155363535 | -1.360686867 | 3.85E-22 | 0.000332056 |
| NKAIN3 | MGC | young | -0.156614304 | -1.235545027 | 0.000113749 | 0.013374984 |
| MT-ND4 | MGC | young | -0.15715886 | -0.682754293 | 4.93E-45 | 0.001140898 |
| HCFC1 | MGC | young | -0.158051335 | -0.873747258 | 6.68E-05 | 0.045022913 |
| KDM3B | MGC | young | -0.158991945 | -1.186549795 | 3.98E-36 | 1.15E-06 |
| AC103923.1 | MGC | young | -0.160442777 | -1.24930306 | 1.73E-10 | 0.006767094 |
| DYRK1B | MGC | young | -0.160889043 | -0.902239792 | 0.003422749 | 0.036117803 |
| ITIH5 | MGC | young | -0.161222884 | -1.002563834 | 7.78E-09 | 0.009405395 |
| SPAST | MGC | young | -0.161414724 | -0.913037494 | 4.47E-14 | 0.009232062 |
| FBXO9 | MGC | young | -0.163243213 | -0.988364602 | 3.78E-36 | 0.000116412 |
| C2orf92 | MGC | young | -0.163813591 | -1.016376693 | 1.28E-12 | 0.012718305 |
| DOCK9 | MGC | young | -0.164211941 | -0.900237253 | 3.91E-29 | 0.002887003 |
| RARB | MGC | young | -0.164595205 | -1.039024348 | 4.92E-58 | 7.19E-05 |
| RSRP1 | MGC | young | -0.166201005 | -0.45892811 | 1.92E-32 | 0.021175724 |
| AC114971.1 | MGC | young | -0.166466152 | -1.352283591 | 8.20E-69 | 1.20E-06 |
| OBSL1 | MGC | young | -0.166691684 | -0.580726943 | 1.37E-09 | 0.047286868 |
| SDCBP2-AS1 | MGC | young | -0.16712932 | -1.069011866 | 3.52E-15 | 0.004336357 |
| NFIA | MGC | young | -0.167909371 | -1.325477868 | 3.58E-145 | 1.69E-10 |
| ZNF334 | MGC | young | -0.168059607 | -1.168698678 | 2.22E-33 | 0.000247971 |
| WAC-AS1 | MGC | young | -0.168727676 | -0.856308973 | 3.17E-16 | 0.008180479 |
| ARHGAP21 | MGC | young | -0.169071492 | -1.333057037 | 3.23E-129 | 1.51E-11 |
| CXXC1 | MGC | young | -0.170407862 | -0.876247302 | 0.006566071 | 0.048921481 |
| BRD7 | MGC | young | -0.171077293 | -0.600769953 | 1.94E-15 | 0.026885826 |
| NEDD9 | MGC | young | -0.171376068 | -0.62934139 | 2.45E-19 | 0.006826745 |
| KCNH8 | MGC | young | -0.171746119 | -1.401773263 | 4.15E-21 | 2.16E-06 |
| CCDC136 | MGC | young | -0.171748279 | -1.321211772 | 2.30E-20 | 0.000213742 |
| SPATC1 | MGC | young | -0.172192515 | -1.281130743 | 0.000156455 | 0.010685809 |
| FAM184B | MGC | young | -0.172607979 | -0.814163374 | 2.84E-17 | 0.008158505 |
| MEGF10 | MGC | young | -0.173619326 | -1.704679148 | 3.66E-104 | 3.82E-13 |
| SLC8A1-AS1 | MGC | young | -0.17500569 | -1.52752393 | 1.19E-29 | 2.22E-06 |
| LINC01182 | MGC | young | -0.175388658 | -1.103776573 | 8.32E-11 | 0.010310156 |
| ZNF420 | MGC | young | -0.175434591 | -1.177985465 | 1.82E-18 | 0.000424451 |
| SGCZ | MGC | young | -0.175844133 | -1.345568093 | 8.87E-25 | 0.004567066 |
| DIO2 | MGC | young | -0.176242953 | -1.56770025 | 4.72E-90 | 6.00E-10 |
| LRP6 | MGC | young | -0.177989458 | -1.205391314 | 1.02E-34 | 5.19E-05 |
| UBA6-AS1 | MGC | young | -0.179075466 | -0.880653097 | 1.87E-27 | 0.000289842 |
| COQ10A | MGC | young | -0.179672975 | -0.769520779 | 1.00E-06 | 0.036438567 |
| ANGEL1 | MGC | young | -0.179991054 | -0.885260175 | 4.24E-06 | 0.039138962 |
| LINC02340 | MGC | young | -0.180067733 | -1.292232676 | 8.13E-46 | 2.95E-05 |
| CASTOR2 | MGC | young | -0.180219507 | -0.834936431 | 5.49E-07 | 0.019919774 |
| UNG | MGC | young | -0.180649982 | -0.85760034 | 1.19E-08 | 0.039743121 |
| PLEKHG1 | MGC | young | -0.180717876 | -1.081247149 | 3.10E-51 | 1.77E-05 |
| CASP2 | MGC | young | -0.181762003 | -1.053763708 | 1.55E-07 | 0.014795371 |
| SORBS2 | MGC | young | -0.18212275 | -1.275393372 | 1.15E-163 | 3.21E-12 |
| NR3C2 | MGC | young | -0.18238846 | -1.340554837 | 6.57E-52 | 2.17E-08 |
| PEBP4 | MGC | young | -0.182653476 | -1.574103247 | 1.07E-62 | 1.62E-08 |
| DDX58 | MGC | young | -0.182769896 | -0.822178462 | 2.33E-19 | 0.008897773 |
| C2orf88 | MGC | young | -0.182774083 | -0.978037374 | 3.67E-10 | 0.012493709 |

| | | | | | | |
|---|---|---|---|---|---|---|
| LINC00685 | MGC | young | -0.182846422 | -1.196026072 | 0.000176411 | 0.025492161 |
| ITGB8 | MGC | young | -0.183105974 | -0.871660734 | 6.59E-36 | 0.007282832 |
| SPEF2 | MGC | young | -0.183132692 | -1.039880319 | 8.24E-23 | 0.000567736 |
| ZNF214 | MGC | young | -0.183350392 | -1.045189969 | 7.99E-11 | 0.009151671 |
| TNS2 | MGC | young | -0.183483796 | -1.102181552 | 1.01E-11 | 0.002567847 |
| SLC30A10 | MGC | young | -0.183658809 | -1.287962568 | 0.00192294 | 0.013332946 |
| AC130650.1 | MGC | young | -0.18371436 | -1.262707816 | 0.000120244 | 0.009470137 |
| C9orf85 | MGC | young | -0.184099743 | -0.717586053 | 8.89E-16 | 0.017820851 |
| SMARCC2 | MGC | young | -0.184188829 | -0.803758367 | 1.24E-16 | 0.010703513 |
| SLC24A3 | MGC | young | -0.185011759 | -1.096079801 | 6.96E-13 | 0.017740196 |
| IFIH1 | MGC | young | -0.185308123 | -1.076161705 | 2.45E-11 | 0.003774763 |
| MFSD4B | MGC | young | -0.185330377 | -0.669238835 | 9.71E-33 | 0.049891872 |
| BCL2L11 | MGC | young | -0.186588882 | -0.74891436 | 0.001524359 | 0.039330833 |
| THBS4 | MGC | young | -0.186857298 | -1.24167 | 3.75E-24 | 0.000145454 |
| AP002495.1 | MGC | young | -0.186884353 | -1.224107958 | 3.05E-23 | 6.55E-05 |
| CSRNP3 | MGC | young | -0.187544025 | -1.77481209 | 1.15E-56 | 1.67E-13 |
| MT-ATP6 | MGC | young | -0.188050211 | -0.96065374 | 2.77E-91 | 0.000282862 |
| ELP4 | MGC | young | -0.188341212 | -0.954664774 | 4.22E-61 | 1.46E-05 |
| CEP162 | MGC | young | -0.188903064 | -0.871210804 | 6.53E-16 | 0.00681303 |
| THAP7-AS1 | MGC | young | -0.18941563 | -1.241102576 | 3.58E-11 | 0.002732747 |
| CCDC7 | MGC | young | -0.189645731 | -1.784289986 | 2.10E-51 | 1.77E-11 |
| TCAIM | MGC | young | -0.190863764 | -0.630266784 | 6.34E-10 | 0.034426508 |
| SPTLC3 | MGC | young | -0.19181186 | -1.153586376 | 3.67E-20 | 0.000390678 |
| MTAP | MGC | young | -0.192232221 | -1.274470983 | 2.69E-35 | 4.00E-06 |
| IGSF10 | MGC | young | -0.192306444 | -1.71098602 | 2.30E-42 | 4.79E-09 |
| PDCL | MGC | young | -0.19289212 | -0.757502532 | 3.75E-14 | 0.013418049 |
| BRI3BP | MGC | young | -0.193109978 | -0.897473565 | 5.35E-09 | 0.021102577 |
| NACC2 | MGC | young | -0.193649738 | -0.846880388 | 7.41E-11 | 0.024588588 |
| MANEA-DT | MGC | young | -0.193690231 | -0.930072901 | 1.16E-05 | 0.02764115 |
| CFTR | MGC | young | -0.193926754 | -1.136536064 | 2.33E-05 | 0.003572352 |
| NCKAP5 | MGC | young | -0.194097741 | -1.333155159 | 1.37E-102 | 9.20E-09 |
| CHRDL1 | MGC | young | -0.194698114 | -0.652018555 | 7.72E-40 | 0.009248193 |
| MALRD1 | MGC | young | -0.196265246 | -1.124825569 | 1.30E-12 | 0.007276094 |
| AC007193.2 | MGC | young | -0.19704361 | -1.466365767 | 1.33E-05 | 0.001171017 |
| NCL | MGC | young | -0.197395615 | -0.801914318 | 4.34E-63 | 2.85E-05 |
| MDM1 | MGC | young | -0.197528286 | -0.898501036 | 2.27E-16 | 0.005709666 |
| AL590652.1 | MGC | young | -0.198449304 | -1.199129416 | 1.10E-06 | 0.003487092 |
| CDC14A | MGC | young | -0.198915515 | -0.636685004 | 4.88E-25 | 0.003266134 |
| LINC01515 | MGC | young | -0.199096687 | -2.091434198 | 6.69E-62 | 2.84E-09 |
| CTNNA3 | MGC | young | -0.199133381 | -1.670515281 | 2.74E-15 | 1.73E-05 |
| SP1 | MGC | young | -0.199145564 | -1.08182716 | 5.62E-24 | 3.31E-05 |
| P2RX7 | MGC | young | -0.200067607 | -1.218893756 | 0.00063938 | 0.014817289 |
| TMEM184B | MGC | young | -0.20029077 | -0.773408191 | 2.77E-18 | 0.003120018 |
| ZNF362 | MGC | young | -0.200417273 | -0.947123892 | 2.55E-08 | 0.022651545 |
| RALB | MGC | young | -0.200729135 | -0.91584432 | 6.99E-12 | 0.013271229 |
| TMEM47 | MGC | young | -0.200968132 | -0.743615866 | 9.95E-39 | 0.037056906 |
| CHROMR | MGC | young | -0.200981274 | -1.131084601 | 8.00E-10 | 0.005292904 |
| ARHGEF26-AS1 | MGC | young | -0.201623613 | -1.274741182 | 3.29E-73 | 1.33E-11 |
| GAB1 | MGC | young | -0.201742991 | -0.639724239 | 8.85E-27 | 0.003894856 |
| COL16A1 | MGC | young | -0.202774543 | -1.189571525 | 4.66E-22 | 0.001095042 |
| AC092691.1 | MGC | young | -0.203758747 | -1.391594161 | 9.80E-97 | 5.00E-20 |
| GPR174 | MGC | young | -0.203878485 | -1.721733136 | 3.49E-13 | 0.000819876 |
| NR2F2 | MGC | young | -0.205099243 | -1.381280951 | 3.64E-12 | 0.000899918 |
| GALNT1 | MGC | young | -0.205112671 | -0.632005578 | 2.49E-35 | 0.005721561 |

| | | | | | | |
|---|---|---|---|---|---|---|
| PTCH2 | MGC | young | -0.205157709 | -0.970418462 | 0.000212639 | 0.025369112 |
| CCDC66 | MGC | young | -0.205169543 | -0.563212069 | 4.66E-18 | 0.021362174 |
| ZNF658 | MGC | young | -0.205382007 | -1.329219291 | 3.01E-13 | 0.003698613 |
| USP1 | MGC | young | -0.206294651 | -0.632537393 | 9.34E-14 | 0.03723296 |
| LINC01535 | MGC | young | -0.20718157 | -0.860717132 | 1.71E-17 | 0.007412253 |
| CCDC125 | MGC | young | -0.207515922 | -1.045376726 | 4.61E-14 | 0.009276964 |
| KAT6B | MGC | young | -0.207916256 | -1.119436284 | 4.29E-64 | 5.65E-07 |
| PARP1 | MGC | young | -0.208020184 | -1.061325815 | 8.97E-32 | 8.55E-05 |
| ALDH1L1-AS2 | MGC | young | -0.208445068 | -1.349532315 | 2.46E-06 | 0.000912859 |
| SCAI | MGC | young | -0.208865777 | -1.141320678 | 8.04E-51 | 4.98E-07 |
| MIR181A2HG | MGC | young | -0.209705702 | -1.378222081 | 9.52E-44 | 6.06E-11 |
| ATP8B1 | MGC | young | -0.209917156 | -1.198170237 | 2.13E-12 | 0.001619652 |
| DNAH6 | MGC | young | -0.210298855 | -0.84802834 | 4.05E-13 | 0.013793727 |
| ETV5 | MGC | young | -0.210649831 | -0.985534386 | 8.94E-34 | 0.000548723 |
| EPCAM | MGC | young | -0.211004121 | -2.416586884 | 1.80E-101 | 1.48E-14 |
| NLGN4X | MGC | young | -0.211240247 | -1.172923751 | 4.36E-50 | 7.10E-06 |
| DTWD2 | MGC | young | -0.212389521 | -1.312571132 | 8.80E-23 | 0.000341003 |
| MSRA | MGC | young | -0.212893145 | -0.719090638 | 1.43E-13 | 0.03646966 |
| AC027117.1 | MGC | young | -0.213283153 | -1.421903638 | 0.000382458 | 0.005764991 |
| KIAA1217 | MGC | young | -0.214391645 | -1.766561559 | 5.33E-195 | 1.33E-14 |
| FREM1 | MGC | young | -0.214465723 | -2.147419715 | 3.10E-46 | 2.38E-11 |
| AL392086.3 | MGC | young | -0.21461878 | -2.025801385 | 6.18E-79 | 6.70E-09 |
| BCHE | MGC | young | -0.215261784 | -0.634692471 | 4.26E-19 | 0.018058495 |
| ARID4A | MGC | young | -0.215280733 | -1.031212314 | 6.88E-40 | 5.02E-06 |
| PAFAH1B2 | MGC | young | -0.216323722 | -0.654283918 | 6.70E-18 | 0.014592561 |
| RFTN2 | MGC | young | -0.216400654 | -1.207432232 | 2.05E-54 | 9.39E-07 |
| ZHX1 | MGC | young | -0.218160325 | -0.60552389 | 6.80E-21 | 0.023581388 |
| GALNT3 | MGC | young | -0.219192004 | -1.154219949 | 4.91E-35 | 0.001357523 |
| LINC02610 | MGC | young | -0.21950802 | -1.239579584 | 9.27E-09 | 0.007932168 |
| MTBP | MGC | young | -0.221358129 | -0.978881811 | 5.11E-05 | 0.036688842 |
| AL355612.1 | MGC | young | -0.221944042 | -1.765434146 | 3.39E-21 | 0.00027334 |
| ABHD2 | MGC | young | -0.22228358 | -0.83424954 | 2.91E-33 | 0.001157804 |
| MAP2 | MGC | young | -0.222285184 | -1.010960475 | 2.54E-110 | 1.55E-07 |
| FUT8 | MGC | young | -0.223367021 | -2.085456441 | 2.61E-150 | 2.73E-25 |
| GLRB | MGC | young | -0.224467773 | -0.938080753 | 4.23E-13 | 0.009716783 |
| RAVER2 | MGC | young | -0.225664552 | -1.460989957 | 1.62E-54 | 5.94E-07 |
| UTRN | MGC | young | -0.22640512 | -3.299907417 | 9.57E-125 | 4.92E-22 |
| ACADM | MGC | young | -0.226470361 | -0.911899318 | 2.11E-38 | 0.000247959 |
| AC012613.1 | MGC | young | -0.227097026 | -1.057265424 | 3.78E-05 | 0.007987803 |
| CLVS2 | MGC | young | -0.227686394 | -1.760898092 | 2.76E-82 | 2.04E-13 |
| FREM2 | MGC | young | -0.228026799 | -2.596357244 | 1.41E-32 | 6.26E-07 |
| FANCL | MGC | young | -0.228497911 | -0.711934542 | 1.82E-24 | 0.006927035 |
| METAP1 | MGC | young | -0.229219237 | -0.886610387 | 6.38E-13 | 0.022321291 |
| NBPF1 | MGC | young | -0.229699207 | -0.869644328 | 8.43E-11 | 0.030629937 |
| PIK3R3 | MGC | young | -0.229911334 | -1.688154048 | 2.27E-79 | 1.01E-15 |
| MT-ND1 | MGC | young | -0.230605875 | -0.817220385 | 1.34E-65 | 0.000458401 |
| PRIM2 | MGC | young | -0.230792668 | -0.736692787 | 7.67E-06 | 0.042066887 |
| GPM6A | MGC | young | -0.230950862 | -1.039057329 | 1.15E-134 | 1.29E-08 |
| AC011416.4 | MGC | young | -0.233170075 | -1.400076669 | 3.54E-18 | 5.44E-05 |
| GABPB1-AS1 | MGC | young | -0.233984615 | -0.865620509 | 2.13E-28 | 0.001871114 |
| PXMP4 | MGC | young | -0.235078096 | -0.964073248 | 3.53E-08 | 0.018524247 |
| FIBIN | MGC | young | -0.235266025 | -1.642147318 | 4.76E-20 | 3.44E-05 |
| C9orf64 | MGC | young | -0.235378886 | -1.298432105 | 4.55E-10 | 0.002430511 |
| DDX46 | MGC | young | -0.235694822 | -0.654622868 | 4.17E-20 | 0.019006172 |

| | | | | | | |
|---|---|---|---|---|---|---|
| HNRNPM | MGC | young | -0.235700052 | -0.671360724 | 1.69E-28 | 0.001698774 |
| ZNF605 | MGC | young | -0.23697479 | -1.128200485 | 7.99E-12 | 0.005363017 |
| HIST3H2A | MGC | young | -0.237107844 | -1.020360014 | 7.25E-10 | 0.00158067 |
| GRIN2A | MGC | young | -0.237255669 | -1.311044815 | 1.32E-25 | 6.50E-06 |
| ANTXR1 | MGC | young | -0.237521213 | -1.117668819 | 1.57E-25 | 0.000208969 |
| AGGF1 | MGC | young | -0.237717666 | -0.66539318 | 1.89E-12 | 0.036521488 |
| CTSO | MGC | young | -0.238251503 | -1.127189165 | 3.24E-15 | 0.002621761 |
| ANKRD30BL | MGC | young | -0.239301094 | -1.283595389 | 1.15E-31 | 6.29E-05 |
| KDM4A | MGC | young | -0.23975701 | -1.299283999 | 1.90E-26 | 5.05E-05 |
| SUCLG2-AS1 | MGC | young | -0.240064363 | -1.324692633 | 7.28E-33 | 1.66E-05 |
| NCAPD2 | MGC | young | -0.240665563 | -1.594015788 | 2.47E-16 | 7.64E-06 |
| DGLUCY | MGC | young | -0.241063845 | -0.980602433 | 2.52E-16 | 0.002415484 |
| NRBP2 | MGC | young | -0.241375324 | -1.011371851 | 2.27E-14 | 0.006168144 |
| AC061958.1 | MGC | young | -0.241573097 | -1.426221362 | 0.000189805 | 0.003294063 |
| AC008269.1 | MGC | young | -0.24223169 | -1.757325959 | 1.99E-11 | 1.53E-05 |
| AC110296.1 | MGC | young | -0.242619953 | -1.004921379 | 8.30E-17 | 0.009603385 |
| PRPF6 | MGC | young | -0.242842111 | -0.58214985 | 1.10E-16 | 0.049622342 |
| AC005498.3 | MGC | young | -0.243000835 | -1.212844072 | 6.79E-09 | 0.005545689 |
| AC024558.2 | MGC | young | -0.243333206 | -1.194751128 | 0.003481457 | 0.024509399 |
| AC084198.2 | MGC | young | -0.243749496 | -1.702577338 | 1.04E-48 | 1.58E-08 |
| TSGA10 | MGC | young | -0.244291664 | -0.879139783 | 1.37E-18 | 0.000822267 |
| AC025442.2 | MGC | young | -0.244365721 | -1.382569177 | 1.10E-16 | 0.003895393 |
| NT5DC1 | MGC | young | -0.244479259 | -0.661777812 | 9.12E-12 | 0.022950308 |
| FLRT2 | MGC | young | -0.244520153 | -1.23664734 | 4.79E-71 | 3.84E-07 |
| TRIM73 | MGC | young | -0.245047442 | -1.194715357 | 1.92E-19 | 0.006442783 |
| DZIP3 | MGC | young | -0.24643923 | -0.957967235 | 5.02E-38 | 0.001026945 |
| VPS50 | MGC | young | -0.246591538 | -0.958382067 | 5.11E-19 | 0.003535039 |
| AL161668.3 | MGC | young | -0.246923386 | -1.512435234 | 5.34E-25 | 0.000178327 |
| LDB1 | MGC | young | -0.247125315 | -0.815180254 | 5.76E-08 | 0.033444197 |
| SOX5 | MGC | young | -0.247808387 | -2.132243211 | 6.16E-156 | 7.76E-20 |
| AMER2 | MGC | young | -0.24826433 | -2.00959167 | 1.03E-146 | 7.00E-25 |
| PAUPAR | MGC | young | -0.250169519 | -2.026720017 | 1.43E-61 | 2.73E-21 |
| DHX38 | MGC | young | -0.25078318 | -0.879680453 | 8.10E-06 | 0.049803285 |
| RIMS1 | MGC | young | -0.251980098 | -2.19412051 | 7.33E-48 | 1.38E-14 |
| PREX1 | MGC | young | -0.253223363 | -0.776329056 | 1.18E-14 | 0.011295071 |
| GSAP | MGC | young | -0.25331646 | -1.171217422 | 8.23E-36 | 1.49E-07 |
| PCMTD1 | MGC | young | -0.25445256 | -0.611421639 | 2.75E-61 | 0.006841407 |
| DNAH12 | MGC | young | -0.254700986 | -1.135342827 | 1.63E-09 | 0.003372609 |
| SH3BP2 | MGC | young | -0.255766137 | -0.968116437 | 6.75E-16 | 0.006557305 |
| AL137009.1 | MGC | young | -0.256891345 | -1.111739472 | 1.56E-12 | 0.020408114 |
| RNF180 | MGC | young | -0.257446382 | -1.207074783 | 6.09E-52 | 1.09E-06 |
| KCTD1 | MGC | young | -0.258418799 | -1.273192982 | 2.44E-33 | 1.06E-05 |
| CLHC1 | MGC | young | -0.258452701 | -0.678810792 | 1.25E-10 | 0.048473282 |
| PCSK2 | MGC | young | -0.258503895 | -1.555341807 | 2.50E-111 | 1.22E-17 |
| PRC1-AS1 | MGC | young | -0.258792411 | -1.184597082 | 0.002621419 | 0.013562316 |
| JCAD | MGC | young | -0.258901343 | -1.069700309 | 6.09E-11 | 0.000550822 |
| PLCH1 | MGC | young | -0.259081395 | -1.650634272 | 2.06E-77 | 5.97E-14 |
| SNRNP200 | MGC | young | -0.260167033 | -0.924637587 | 1.31E-34 | 5.15E-05 |
| AC024588.1 | MGC | young | -0.261559373 | -2.478598755 | 1.20E-16 | 1.25E-07 |
| OBI1-AS1 | MGC | young | -0.26168245 | -1.184932567 | 0.002949833 | 0.020867505 |
| MAP6 | MGC | young | -0.2633593 | -0.834148774 | 7.72E-07 | 0.029910512 |
| AC021055.1 | MGC | young | -0.263581255 | -0.935392407 | 6.64E-12 | 0.009837833 |
| EPHB1 | MGC | young | -0.263640762 | -1.742022947 | 2.34E-65 | 4.06E-13 |
| FAM111A | MGC | young | -0.263780984 | -0.801101141 | 2.85E-25 | 0.004900262 |

| | | | | | | |
|---|---|---|---|---|---|---|
| CSPG5 | MGC | young | -0.263867374 | -1.092607711 | 2.50E-28 | 0.000528759 |
| MT-CO2 | MGC | young | -0.264501084 | -1.5881894 | 9.38E-205 | 1.52E-12 |
| KREMEN1 | MGC | young | -0.264786192 | -1.312398352 | 5.83E-22 | 5.13E-05 |
| IVD | MGC | young | -0.26529944 | -0.698273228 | 1.25E-09 | 0.030581036 |
| AGL | MGC | young | -0.266642432 | -1.098056877 | 7.65E-86 | 3.22E-06 |
| NCK2 | MGC | young | -0.267299509 | -1.276876464 | 6.01E-45 | 2.78E-06 |
| AC124854.1 | MGC | young | -0.267342358 | -1.72715697 | 3.67E-41 | 5.06E-09 |
| TEC | MGC | young | -0.267366602 | -1.095399161 | 2.26E-47 | 1.04E-05 |
| CC2D2A | MGC | young | -0.268531148 | -0.646681353 | 5.31E-13 | 0.023721577 |
| MYO6 | MGC | young | -0.269390715 | -1.090690878 | 2.75E-123 | 4.26E-11 |
| SOX6 | MGC | young | -0.269400715 | -1.823903336 | 8.21E-163 | 1.33E-19 |
| RBM45 | MGC | young | -0.269613404 | -1.014007716 | 1.08E-11 | 0.006638 |
| TYW5 | MGC | young | -0.269746598 | -0.592630195 | 2.38E-18 | 0.029292601 |
| STPG2-AS1 | MGC | young | -0.269862175 | -1.229384612 | 3.39E-13 | 0.009559076 |
| CLSTN1 | MGC | young | -0.271464726 | -0.753437676 | 3.55E-36 | 0.00027484 |
| RGL1 | MGC | young | -0.272142076 | -1.086988351 | 8.22E-73 | 5.84E-07 |
| AL157400.3 | MGC | young | -0.272471135 | -1.424183237 | 7.06E-06 | 0.003094146 |
| AL160272.1 | MGC | young | -0.272972476 | -1.366737359 | 5.56E-18 | 0.002380465 |
| DLG3 | MGC | young | -0.273378165 | -0.897616861 | 1.61E-08 | 0.023501427 |
| PDE11A | MGC | young | -0.273616528 | -0.82255056 | 9.64E-08 | 0.045501041 |
| FAM227B | MGC | young | -0.273691926 | -1.100755684 | 6.18E-35 | 2.12E-05 |
| ABCA8 | MGC | young | -0.274720817 | -1.53488582 | 3.97E-94 | 7.19E-09 |
| FAT4 | MGC | young | -0.275635212 | -1.526436501 | 8.28E-17 | 0.000409674 |
| IRAK1BP1 | MGC | young | -0.276394507 | -0.921887768 | 2.44E-23 | 0.001391414 |
| PSTPIP2 | MGC | young | -0.27672663 | -1.112564815 | 3.23E-46 | 4.75E-06 |
| CLIC5 | MGC | young | -0.277489734 | -1.627088265 | 6.35E-49 | 1.11E-10 |
| CPA6 | MGC | young | -0.277733957 | -1.545614752 | 7.80E-138 | 3.77E-13 |
| TRIM41 | MGC | young | -0.278147658 | -0.848327948 | 4.18E-05 | 0.046582647 |
| FAP | MGC | young | -0.278625985 | -1.584435355 | 3.77E-97 | 1.32E-09 |
| CELSR2 | MGC | young | -0.278949336 | -1.107313027 | 2.06E-15 | 0.002498904 |
| NAIP | MGC | young | -0.279233293 | -1.32910304 | 1.00E-07 | 0.002378778 |
| ADAMTS12 | MGC | young | -0.279525635 | -1.592249695 | 3.82E-32 | 9.96E-06 |
| THY1 | MGC | young | -0.279703174 | -2.191346251 | 3.89E-43 | 3.50E-09 |
| CTTNBP2 | MGC | young | -0.279793008 | -1.261483272 | 1.16E-99 | 3.70E-10 |
| WDR92 | MGC | young | -0.280610459 | -1.085215802 | 6.45E-13 | 0.005979452 |
| TUBGCP5 | MGC | young | -0.28262613 | -1.283846977 | 7.78E-18 | 0.000468287 |
| TSR2 | MGC | young | -0.283473489 | -0.669426152 | 5.57E-16 | 0.028903761 |
| CLVS1 | MGC | young | -0.284750196 | -2.354899751 | 2.69E-77 | 6.06E-20 |
| AL158195.1 | MGC | young | -0.284881374 | -0.958940682 | 2.28E-05 | 0.04922014 |
| ADAMTS6 | MGC | young | -0.285055964 | -2.54297338 | 1.29E-59 | 2.15E-14 |
| KLHL8 | MGC | young | -0.285754482 | -1.91432602 | 1.74E-173 | 5.45E-22 |
| ALDH9A1 | MGC | young | -0.28598666 | -0.712438443 | 1.10E-29 | 0.013562316 |
| AC073050.1 | MGC | young | -0.286203528 | -0.786046073 | 1.17E-17 | 0.042356025 |
| HNRNPUL2 | MGC | young | -0.286478525 | -1.079270174 | 1.79E-25 | 0.000252521 |
| AL158055.1 | MGC | young | -0.286953424 | -1.359191481 | 2.98E-11 | 0.009222405 |
| RFXAP | MGC | young | -0.288594247 | -1.560672555 | 1.10E-18 | 1.80E-05 |
| DGKH | MGC | young | -0.290055706 | -0.721929986 | 2.43E-19 | 0.049266235 |
| MARVELD1 | MGC | young | -0.290284511 | -1.318994789 | 2.66E-05 | 0.007645744 |
| MDFIC | MGC | young | -0.291745353 | -0.650428156 | 2.15E-49 | 0.005024023 |
| RTCA-AS1 | MGC | young | -0.292496514 | -0.91080834 | 2.37E-16 | 0.019714817 |
| PTPN20 | MGC | young | -0.294073867 | -1.935872283 | 5.71E-33 | 7.77E-08 |
| MTRF1 | MGC | young | -0.294951781 | -0.907081364 | 2.52E-11 | 0.040678771 |
| NRXN1 | MGC | young | -0.295903744 | -0.922002572 | 8.24E-05 | 0.035099028 |
| AL591242.1 | MGC | young | -0.29649005 | -1.446228423 | 3.21E-05 | 0.007505453 |

| | | | | | | |
|---|---|---|---|---|---|---|
| LYPLAL1-DT | MGC | young | -0.296963584 | -1.244457193 | 7.07E-11 | 0.003909904 |
| POC1B-AS1 | MGC | young | -0.297138331 | -1.013972799 | 1.80E-07 | 0.016365045 |
| ZMPSTE24 | MGC | young | -0.297417155 | -0.837750184 | 7.79E-21 | 0.021102017 |
| ADGRL3 | MGC | young | -0.297711486 | -1.933568523 | 1.74E-178 | 1.54E-19 |
| ZNF233 | MGC | young | -0.299250272 | -1.045270689 | 2.71E-06 | 0.026234757 |
| MEP1B | MGC | young | -0.299633522 | -1.271043459 | 0.000178312 | 0.017372627 |
| AC063944.1 | MGC | young | -0.302086594 | -2.034000476 | 3.64E-34 | 2.03E-07 |
| AL132857.1 | MGC | young | -0.302659949 | -1.154752117 | 3.75E-07 | 0.006415517 |
| PROX1 | MGC | young | -0.303673794 | -1.281439818 | 7.64E-38 | 1.74E-05 |
| AKR1C1 | MGC | young | -0.304435358 | -1.030580507 | 1.01E-14 | 0.000976338 |
| ZNF197 | MGC | young | -0.306068358 | -1.055348783 | 1.14E-27 | 0.003281581 |
| ZNF883 | MGC | young | -0.306822081 | -1.094947971 | 4.24E-23 | 0.00088312 |
| LINC00910 | MGC | young | -0.307505098 | -1.071166581 | 0.002659538 | 0.021438679 |
| PIGS | MGC | young | -0.30770813 | -0.837458801 | 5.07E-08 | 0.027281426 |
| KLF7 | MGC | young | -0.307870696 | -1.117403535 | 6.12E-27 | 0.00089397 |
| RAB30-DT | MGC | young | -0.307955165 | -0.520204602 | 1.80E-22 | 0.041340405 |
| CALM2 | MGC | young | -0.30870269 | -0.45319247 | 9.40E-61 | 0.046526952 |
| C4orf19 | MGC | young | -0.309392241 | -1.162528212 | 1.68E-28 | 2.38E-05 |
| SFXN5 | MGC | young | -0.309701199 | -1.7440275 | 5.72E-101 | 2.80E-17 |
| KIF4A | MGC | young | -0.310037546 | -0.983901971 | 0.000224516 | 0.028878779 |
| DSE | MGC | young | -0.311266398 | -0.64077789 | 1.60E-14 | 0.015420658 |
| SCD5 | MGC | young | -0.311928207 | -0.68234963 | 9.22E-43 | 0.010660347 |
| SRGAP3 | MGC | young | -0.313339723 | -1.325298947 | 3.43E-104 | 3.38E-13 |
| AC007432.1 | MGC | young | -0.313678982 | -1.411318928 | 1.37E-11 | 0.00160667 |
| SMC6 | MGC | young | -0.315079079 | -0.99853129 | 9.31E-28 | 0.000433828 |
| LRP5 | MGC | young | -0.316131977 | -0.956165406 | 1.45E-06 | 0.010757264 |
| AC020659.1 | MGC | young | -0.316959683 | -0.903522106 | 5.60E-08 | 0.012209971 |
| IK | MGC | young | -0.317514508 | -0.748755283 | 2.84E-30 | 0.005484927 |
| CASC19 | MGC | young | -0.317545239 | -1.066357062 | 3.12E-05 | 0.046392781 |
| ZNF687 | MGC | young | -0.317587264 | -1.269538485 | 0.02726116 | 0.026527835 |
| NXPE3 | MGC | young | -0.318186449 | -0.716772115 | 2.26E-14 | 0.016037612 |
| DDX60 | MGC | young | -0.318643202 | -1.150542506 | 8.60E-13 | 0.00317989 |
| EHMT2 | MGC | young | -0.320005664 | -1.351251696 | 4.79E-12 | 0.001665504 |
| SHROOM4 | MGC | young | -0.320097498 | -1.933945746 | 7.16E-46 | 7.18E-11 |
| NOXA1 | MGC | young | -0.320307664 | -0.826976399 | 2.38E-07 | 0.044288725 |
| LINC02021 | MGC | young | -0.321085391 | -1.151408034 | 0.000359357 | 0.025369112 |
| SLC13A4 | MGC | young | -0.32151158 | -1.097324057 | 4.65E-11 | 0.006477921 |
| MGAT4A | MGC | young | -0.321547882 | -0.896272014 | 7.67E-17 | 0.013150679 |
| ZNF253 | MGC | young | -0.321667088 | -0.985844694 | 3.30E-17 | 0.010795386 |
| GOPC | MGC | young | -0.321767409 | -0.582819896 | 7.50E-12 | 0.041936394 |
| LNPK | MGC | young | -0.322117601 | -0.888307621 | 1.08E-56 | 0.00079292 |
| AC040168.1 | MGC | young | -0.322349641 | -1.778060993 | 2.15E-19 | 9.71E-06 |
| SLC6A1-AS1 | MGC | young | -0.323296067 | -1.434148482 | 2.00E-05 | 0.021542372 |
| EIF2AK2 | MGC | young | -0.324328309 | -0.761522315 | 6.68E-20 | 0.013163068 |
| GPR156 | MGC | young | -0.324732209 | -2.057566669 | 1.39E-20 | 1.74E-06 |
| MIR181A1HG | MGC | young | -0.324953029 | -1.902382571 | 2.14E-65 | 9.49E-12 |
| ACSL6 | MGC | young | -0.325012356 | -1.541535648 | 4.14E-30 | 1.84E-07 |
| SNAI3 | MGC | young | -0.325278185 | -1.362776212 | 0.009654363 | 0.015930746 |
| C15orf40 | MGC | young | -0.325313745 | -0.836839638 | 1.93E-21 | 0.002107244 |
| SLC4A8 | MGC | young | -0.325700084 | -1.853711779 | 4.31E-20 | 2.41E-07 |
| ARHGEF40 | MGC | young | -0.326154836 | -0.780531368 | 1.80E-07 | 0.026606033 |
| LYPD6 | MGC | young | -0.326477362 | -1.219208841 | 0.000204784 | 0.004614777 |
| MYCBP2-AS1 | MGC | young | -0.326595306 | -0.815504432 | 8.87E-12 | 0.047716379 |
| AC009495.3 | MGC | young | -0.327325941 | -1.755742538 | 4.09E-33 | 3.54E-07 |

| | | | | | | |
|---|---|---|---|---|---|---|
| ARL6IP6 | MGC | young | -0.327476227 | -0.652898483 | 1.55E-20 | 0.041861157 |
| DDIT4L | MGC | young | -0.327630991 | -1.380315676 | 4.94E-16 | 0.001477913 |
| AL445430.2 | MGC | young | -0.32788651 | -1.301591678 | 0.046182398 | 0.02300932 |
| WSCD1 | MGC | young | -0.328965099 | -1.257283384 | 2.72E-05 | 0.009380718 |
| DUSP16 | MGC | young | -0.330213832 | -1.000838414 | 1.79E-69 | 3.79E-08 |
| CASKIN1 | MGC | young | -0.330412924 | -1.5343146 | 2.85E-07 | 0.000301091 |
| LRP4-AS1 | MGC | young | -0.330826308 | -2.082787947 | 2.76E-29 | 2.66E-06 |
| PIP4K2B | MGC | young | -0.331110851 | -0.997902745 | 1.00E-08 | 0.016485196 |
| GLI3 | MGC | young | -0.331302392 | -1.714895501 | 3.21E-42 | 1.11E-12 |
| CHD7 | MGC | young | -0.331916998 | -0.822477453 | 9.84E-52 | 0.000156683 |
| AMIGO2 | MGC | young | -0.332074258 | -0.981158489 | 5.43E-07 | 0.016976032 |
| REEP1 | MGC | young | -0.332122411 | -1.083412325 | 1.00E-12 | 0.002487446 |
| KRT40 | MGC | young | -0.33282647 | -2.654355337 | 1.07E-26 | 3.62E-10 |
| UPF3A | MGC | young | -0.332999301 | -0.510355878 | 2.97E-20 | 0.048358173 |
| NAV3 | MGC | young | -0.333108221 | -2.505453856 | 4.44E-232 | 3.92E-37 |
| CTBP1-DT | MGC | young | -0.333798215 | -1.041680529 | 0.000422473 | 0.017928927 |
| AP002026.1 | MGC | young | -0.334066173 | -1.544230448 | 1.23E-13 | 0.000548723 |
| GPR180 | MGC | young | -0.334897531 | -1.007308001 | 1.06E-15 | 0.003582128 |
| DPYSL5 | MGC | young | -0.335879637 | -1.043741974 | 3.77E-17 | 0.001269898 |
| PCAT6 | MGC | young | -0.33616503 | -0.691180089 | 8.97E-10 | 0.016805404 |
| GOLIM4 | MGC | young | -0.337197133 | -0.70288508 | 1.13E-62 | 0.001394748 |
| TET1 | MGC | young | -0.337399918 | -1.100639813 | 2.05E-23 | 0.000134203 |
| MPPED2 | MGC | young | -0.337585818 | -1.041952259 | 3.09E-61 | 6.20E-07 |
| AC090709.1 | MGC | young | -0.338036908 | -1.016164511 | 7.99E-07 | 0.047834125 |
| P2RY1 | MGC | young | -0.338711402 | -1.522338322 | 3.19E-58 | 6.26E-07 |
| SPINK2 | MGC | young | -0.339442234 | -1.488678385 | 1.26E-25 | 0.003340582 |
| CHST9 | MGC | young | -0.340164709 | -1.948058367 | 1.16E-146 | 1.61E-18 |
| AP000866.1 | MGC | young | -0.340778417 | -0.880283245 | 0.00576171 | 0.047034967 |
| HP1BP3 | MGC | young | -0.341283211 | -0.477319529 | 1.39E-31 | 0.042521338 |
| MEF2C-AS2 | MGC | young | -0.341544822 | -1.029037862 | 2.83E-09 | 0.003022782 |
| TLR4 | MGC | young | -0.341688092 | -1.683348917 | 8.42E-35 | 3.01E-05 |
| CDKN1B | MGC | young | -0.341993689 | -1.169466981 | 2.76E-68 | 1.35E-06 |
| CMYA5 | MGC | young | -0.343438668 | -1.308495431 | 1.98E-38 | 0.000205651 |
| PRKDC | MGC | young | -0.344057133 | -0.722904708 | 3.77E-37 | 0.000845924 |
| LINC01748 | MGC | young | -0.345714197 | -3.655149464 | 9.07E-72 | 1.53E-23 |
| AC009899.1 | MGC | young | -0.346674466 | -1.376372158 | 1.46E-06 | 0.010203521 |
| BRD8 | MGC | young | -0.346964866 | -1.196893338 | 7.21E-34 | 1.04E-05 |
| RAB11FIP4 | MGC | young | -0.347540408 | -0.83169829 | 3.07E-13 | 0.029070646 |
| PSPH | MGC | young | -0.351968763 | -0.793056343 | 6.77E-12 | 0.015415396 |
| SESN3 | MGC | young | -0.352440383 | -1.21974505 | 1.23E-83 | 5.45E-10 |
| NSUN3 | MGC | young | -0.355650621 | -0.695159203 | 3.80E-14 | 0.028903761 |
| CDC42EP4 | MGC | young | -0.355960904 | -0.816232249 | 1.48E-51 | 8.89E-06 |
| ADORA1 | MGC | young | -0.356808522 | -0.842469137 | 1.78E-08 | 0.028900806 |
| VSIG10 | MGC | young | -0.358343281 | -1.003469178 | 2.47E-17 | 0.000497678 |
| PREX2 | MGC | young | -0.358629697 | -2.720930435 | 1.71E-179 | 3.45E-24 |
| MINDY4B | MGC | young | -0.35896674 | -1.559761634 | 2.66E-11 | 0.000864694 |
| IL12RB2 | MGC | young | -0.359311559 | -1.24738841 | 0.002080923 | 0.022200239 |
| PRR12 | MGC | young | -0.359915621 | -1.352217825 | 2.38E-10 | 0.00036053 |
| CKAP2 | MGC | young | -0.360048068 | -1.467356963 | 9.41E-20 | 3.40E-05 |
| AC097662.1 | MGC | young | -0.360334311 | -1.794431292 | 9.10E-43 | 2.59E-07 |
| CLRN1-AS1 | MGC | young | -0.363127143 | -3.413594589 | 7.76E-75 | 1.05E-22 |
| BCL2L14 | MGC | young | -0.363667763 | -1.288564855 | 2.72E-16 | 0.000383717 |
| GLIPR1L1 | MGC | young | -0.365620655 | -1.447542583 | 2.88E-11 | 0.00089397 |
| PDE5A | MGC | young | -0.365626772 | -1.132234783 | 3.45E-61 | 1.05E-05 |

| | | | | | | |
|---|---|---|---|---|---|---|
| KCNH1 | MGC | young | -0.36584617 | -1.218034219 | 0.000943714 | 0.008158505 |
| SMIM8 | MGC | young | -0.365873267 | -0.852233223 | 5.10E-25 | 0.001469543 |
| CNPY4 | MGC | young | -0.36598058 | -0.774611329 | 1.56E-13 | 0.018395202 |
| TRIL | MGC | young | -0.366592843 | -1.408497562 | 0.015560768 | 0.004548696 |
| ZIC4 | MGC | young | -0.366693454 | -0.89506062 | 9.75E-10 | 0.028512137 |
| AC096576.3 | MGC | young | -0.367362286 | -1.263922242 | 3.67E-07 | 0.00847397 |
| AC044781.1 | MGC | young | -0.369266985 | -1.381718105 | 4.88E-06 | 0.003132275 |
| MOCOS | MGC | young | -0.369615302 | -1.289406355 | 8.08E-19 | 0.000974925 |
| ZNF618 | MGC | young | -0.370704867 | -1.910846383 | 4.92E-26 | 7.66E-08 |
| TFPI | MGC | young | -0.370871008 | -0.858224224 | 2.42E-27 | 0.003268974 |
| CTDSP2 | MGC | young | -0.370936419 | -1.006041441 | 7.39E-17 | 0.000620141 |
| HEPN1 | MGC | young | -0.37220747 | -1.347913357 | 7.57E-09 | 0.001927073 |
| GAS2 | MGC | young | -0.373019286 | -1.223377512 | 1.72E-09 | 0.0028819 |
| ZKSCAN7-AS1 | MGC | young | -0.373330922 | -0.877564826 | 1.44E-24 | 0.008093268 |
| GPSM2 | MGC | young | -0.373819595 | -1.090455952 | 5.49E-16 | 0.003982251 |
| BRMS1L | MGC | young | -0.376362384 | -0.777984804 | 6.56E-07 | 0.018255535 |
| CDK5 | MGC | young | -0.377139374 | -0.996623512 | 5.05E-06 | 0.026612966 |
| ZFYVE21 | MGC | young | -0.37736045 | -0.589087357 | 2.07E-17 | 0.023629978 |
| TRIQK | MGC | young | -0.378015609 | -0.643243659 | 8.21E-23 | 0.02590718 |
| ZNF350-AS1 | MGC | young | -0.378370902 | -0.731163715 | 1.61E-09 | 0.035754661 |
| SCRN1 | MGC | young | -0.378929433 | -0.987516695 | 3.98E-42 | 6.84E-05 |
| AC064807.1 | MGC | young | -0.379328461 | -1.307716496 | 2.02E-14 | 0.000182356 |
| ADGRA2 | MGC | young | -0.380830716 | -0.876628472 | 1.15E-25 | 0.006009209 |
| ZNF30 | MGC | young | -0.380933955 | -1.342649001 | 8.47E-14 | 0.004130079 |
| FAN1 | MGC | young | -0.381900229 | -0.705316609 | 3.10E-16 | 0.020002447 |
| MIA3 | MGC | young | -0.382187545 | -0.635510401 | 2.71E-17 | 0.021586458 |
| SIX3 | MGC | young | -0.382462019 | -1.016858405 | 1.20E-71 | 2.53E-08 |
| ABHD12B | MGC | young | -0.382902528 | -0.51173234 | 1.78E-32 | 0.022997413 |
| AC114316.1 | MGC | young | -0.383262745 | -1.938260703 | 2.90E-23 | 2.17E-07 |
| METTL25 | MGC | young | -0.383363951 | -1.398017125 | 1.83E-19 | 5.91E-05 |
| TMEM220 | MGC | young | -0.384761688 | -1.02967348 | 5.73E-28 | 1.32E-05 |
| ANAPC5 | MGC | young | -0.385248516 | -0.934789589 | 8.24E-33 | 0.000889169 |
| CDH2 | MGC | young | -0.386390258 | -1.382186331 | 2.52E-176 | 6.81E-12 |
| CCND2 | MGC | young | -0.386664615 | -0.719425446 | 2.69E-49 | 0.000472502 |
| SMAD5 | MGC | young | -0.386925988 | -0.550289976 | 3.30E-18 | 0.047404943 |
| ERCC4 | MGC | young | -0.387757633 | -0.876155163 | 2.45E-17 | 0.003734918 |
| GPAA1 | MGC | young | -0.389185802 | -0.928389751 | 1.44E-19 | 0.001467237 |
| RHPN2 | MGC | young | -0.389539039 | -1.00644886 | 3.28E-37 | 5.25E-06 |
| STON1-GTF2A1L | MGC | young | -0.392144452 | -1.285375936 | 1.80E-10 | 0.005037671 |
| COG2 | MGC | young | -0.392216097 | -1.418059139 | 1.00E-34 | 3.67E-06 |
| TTC7A | MGC | young | -0.392361704 | -1.252704967 | 4.15E-31 | 1.51E-05 |
| SACS | MGC | young | -0.393162285 | -0.893124071 | 7.45E-20 | 0.007433029 |
| PDE7B | MGC | young | -0.393989007 | -1.519852674 | 3.85E-82 | 4.91E-10 |
| AC093827.4 | MGC | young | -0.393989111 | -1.009006209 | 9.84E-06 | 0.02520456 |
| DUBR | MGC | young | -0.39441915 | -1.254951024 | 2.71E-24 | 5.87E-05 |
| DACH1 | MGC | young | -0.394681559 | -2.340961185 | 7.77E-249 | 4.21E-32 |
| AC107398.3 | MGC | young | -0.394881296 | -1.535593789 | 7.82E-12 | 0.000256942 |
| MEI4 | MGC | young | -0.397582327 | -1.887390171 | 2.80E-14 | 3.91E-05 |
| ZDHHC21 | MGC | young | -0.397850571 | -0.748515029 | 1.34E-48 | 0.001722691 |
| PTPRG | MGC | young | -0.397884366 | -1.562880382 | 1.24E-89 | 2.53E-09 |
| FSTL1 | MGC | young | -0.398322195 | -0.804169087 | 1.41E-45 | 0.001726869 |
| AC010260.1 | MGC | young | -0.399329734 | -1.545831483 | 2.24E-24 | 0.000440542 |
| EBAG9 | MGC | young | -0.400109562 | -0.571254529 | 5.88E-15 | 0.046445901 |
| RYR1 | MGC | young | -0.400232297 | -1.387763303 | 3.81E-19 | 1.23E-05 |

| | | | | | | |
|---|---|---|---|---|---|---|
| TARS | MGC | young | -0.401973709 | -1.141922702 | 1.00E-64 | 5.93E-07 |
| SORL1 | MGC | young | -0.402069357 | -0.763311363 | 8.21E-10 | 0.02450121 |
| EPHA5-AS1 | MGC | young | -0.402093751 | -3.15400501 | 1.18E-67 | 2.73E-16 |
| ATAD2 | MGC | young | -0.403572666 | -1.14189421 | 4.01E-11 | 0.000899346 |
| KTN1 | MGC | young | -0.404286533 | -0.594130054 | 1.71E-55 | 0.002989706 |
| CPPED1 | MGC | young | -0.404767646 | -0.845684314 | 4.01E-09 | 0.043741512 |
| GRID2 | MGC | young | -0.404819547 | -2.805299732 | 3.48E-257 | 8.65E-33 |
| HEPH | MGC | young | -0.405542764 | -1.361548041 | 1.72E-35 | 9.48E-07 |
| GRIN2C | MGC | young | -0.40594825 | -1.011893014 | 8.90E-06 | 0.025141368 |
| DOC2B | MGC | young | -0.407773397 | -0.905655303 | 2.47E-19 | 0.001022885 |
| SASS6 | MGC | young | -0.408999217 | -0.832144407 | 3.09E-09 | 0.015019941 |
| NOSTRIN | MGC | young | -0.411475345 | -1.468443497 | 1.66E-11 | 0.001328859 |
| DPH5 | MGC | young | -0.411668651 | -0.708482289 | 1.09E-13 | 0.026944234 |
| XPOT | MGC | young | -0.411993904 | -1.175053511 | 7.50E-59 | 1.26E-06 |
| TSN | MGC | young | -0.412118272 | -0.646525633 | 2.26E-21 | 0.01878627 |
| AC091646.1 | MGC | young | -0.412488956 | -2.270481678 | 2.12E-16 | 1.84E-07 |
| NECTIN3 | MGC | young | -0.412835411 | -0.932028188 | 8.00E-30 | 0.000438595 |
| FBXO15 | MGC | young | -0.412981756 | -1.209031105 | 4.22E-08 | 0.003408728 |
| GPT2 | MGC | young | -0.412984469 | -0.870443422 | 5.42E-10 | 0.020986735 |
| AP000787.1 | MGC | young | -0.413226502 | -0.948745978 | 2.52E-17 | 0.00241576 |
| ANOS1 | MGC | young | -0.414410589 | -1.631038238 | 1.86E-109 | 5.51E-10 |
| MYO3A | MGC | young | -0.414479602 | -0.704903751 | 4.85E-74 | 0.001927073 |
| AKAP11 | MGC | young | -0.416731643 | -1.065052206 | 3.69E-67 | 2.21E-07 |
| LRATD1 | MGC | young | -0.416788969 | -1.424064627 | 2.43E-14 | 0.002068365 |
| AC092436.3 | MGC | young | -0.417022869 | -1.081299167 | 1.92E-08 | 0.025574541 |
| AC023509.6 | MGC | young | -0.417373411 | -1.125140318 | 3.11E-19 | 0.000250881 |
| TBC1D9 | MGC | young | -0.423160779 | -0.601679947 | 2.47E-26 | 0.008719651 |
| TMEM221 | MGC | young | -0.425731568 | -1.175914998 | 3.58E-18 | 0.000158617 |
| AC132153.1 | MGC | young | -0.425732447 | -1.128089872 | 4.45E-10 | 0.024095973 |
| UBA1 | MGC | young | -0.428926948 | -0.748541044 | 4.12E-10 | 0.046412678 |
| NR2F1 | MGC | young | -0.429033998 | -0.807336997 | 5.30E-51 | 0.000556289 |
| CEP97 | MGC | young | -0.429296548 | -0.771056259 | 1.32E-12 | 0.008379828 |
| TMEM202-AS1 | MGC | young | -0.429510318 | -1.270698264 | 3.77E-09 | 0.001787539 |
| HTR1F | MGC | young | -0.430131356 | -1.321126434 | 6.62E-23 | 7.57E-05 |
| DPH6-DT | MGC | young | -0.433004273 | -1.611748891 | 1.77E-16 | 5.54E-05 |
| GABRB1 | MGC | young | -0.433567049 | -1.326789283 | 7.87E-89 | 1.08E-13 |
| DDX25 | MGC | young | -0.43368256 | -0.840193661 | 4.25E-05 | 0.040437106 |
| CYP2J2 | MGC | young | -0.4349453 | -0.993178993 | 8.12E-20 | 0.005536463 |
| AC018742.1 | MGC | young | -0.436762287 | -1.137898052 | 6.98E-05 | 0.018223517 |
| KY | MGC | young | -0.436794414 | -1.396279903 | 7.92E-08 | 0.002580866 |
| AC009975.1 | MGC | young | -0.436904523 | -1.797314988 | 1.00E-16 | 0.000105961 |
| KCNE3 | MGC | young | -0.437482076 | -1.053389745 | 5.04E-11 | 0.008380777 |
| AL590560.3 | MGC | young | -0.437597002 | -1.356234969 | 1.57E-08 | 0.010633167 |
| LYPLAL1 | MGC | young | -0.438889042 | -0.725693433 | 2.07E-50 | 0.000387073 |
| MLC1 | MGC | young | -0.43890284 | -1.581545584 | 1.08E-20 | 0.000309859 |
| LINC00298 | MGC | young | -0.439520723 | -2.144900782 | 9.54E-17 | 7.53E-05 |
| PIK3R2 | MGC | young | -0.44021626 | -0.988261003 | 7.44E-08 | 0.020836025 |
| SFT2D3 | MGC | young | -0.440657468 | -0.828638583 | 4.70E-11 | 0.036981728 |
| MADCAM1 | MGC | young | -0.443280813 | -1.739228332 | 2.17E-07 | 0.000671266 |
| RESF1 | MGC | young | -0.444470539 | -0.75532768 | 5.09E-12 | 0.032355476 |
| DAAM2 | MGC | young | -0.444996659 | -1.177546298 | 9.38E-07 | 0.025796796 |
| P2RY14 | MGC | young | -0.445477403 | -1.282123379 | 3.26E-57 | 1.38E-06 |
| MNDA | MGC | young | -0.445631012 | -1.240831723 | 2.31E-89 | 9.36E-13 |
| SLC6A1 | MGC | young | -0.447827586 | -1.092039134 | 1.54E-28 | 0.000201786 |

| | | | | | | |
|---|---|---|---|---|---|---|
| C12orf56 | MGC | young | -0.448898664 | -1.304013046 | 5.84E-40 | 2.97E-07 |
| VGLL3 | MGC | young | -0.449443901 | -3.487735181 | 1.95E-67 | 5.60E-18 |
| SHC3 | MGC | young | -0.449586363 | -0.971836506 | 2.93E-06 | 0.022185722 |
| RHOJ | MGC | young | -0.449899865 | -1.969467027 | 7.23E-86 | 3.82E-16 |
| LINC00886 | MGC | young | -0.449997773 | -1.217170724 | 2.29E-13 | 0.006614943 |
| NPIPB2 | MGC | young | -0.450070917 | -2.028636988 | 9.94E-26 | 2.04E-07 |
| CYP39A1 | MGC | young | -0.450999864 | -0.835660502 | 2.15E-17 | 0.008792525 |
| ZNF33B | MGC | young | -0.45118955 | -1.665921593 | 9.96E-37 | 3.27E-07 |
| HECW2 | MGC | young | -0.451743041 | -2.511509152 | 5.29E-126 | 3.35E-25 |
| AC002451.1 | MGC | young | -0.452994711 | -1.133647288 | 3.11E-20 | 0.000448246 |
| DDX23 | MGC | young | -0.453724076 | -1.215252756 | 2.06E-20 | 0.000283259 |
| AC008966.1 | MGC | young | -0.454754313 | -1.08623063 | 4.72E-06 | 0.016460941 |
| GSTM2 | MGC | young | -0.455369128 | -1.685652621 | 1.64E-30 | 2.83E-07 |
| KRT222 | MGC | young | -0.456493796 | -1.103149705 | 1.92E-24 | 0.000249086 |
| AP001605.1 | MGC | young | -0.456719615 | -1.342900473 | 2.13E-07 | 0.017683177 |
| LRRK2 | MGC | young | -0.458006632 | -1.521253776 | 1.15E-23 | 7.15E-05 |
| SHOC1 | MGC | young | -0.46092415 | -1.07408064 | 6.56E-16 | 0.003408366 |
| TOB2 | MGC | young | -0.461631229 | -0.730166788 | 1.10E-15 | 0.007104434 |
| PRTFDC1 | MGC | young | -0.462579448 | -1.091445611 | 1.63E-40 | 1.74E-07 |
| ICK | MGC | young | -0.463010285 | -1.066010976 | 1.52E-16 | 0.001743394 |
| CAAP1 | MGC | young | -0.463615118 | -0.625637831 | 1.19E-16 | 0.022589 |
| SOWAHA | MGC | young | -0.464779213 | -1.795548617 | 1.29E-23 | 3.38E-08 |
| OSGEPL1 | MGC | young | -0.466576729 | -0.948862028 | 2.62E-12 | 0.010902216 |
| RWDD3 | MGC | young | -0.466595718 | -0.599948585 | 1.42E-16 | 0.035157601 |
| AC004594.1 | MGC | young | -0.466648287 | -1.369849353 | 2.13E-19 | 6.55E-05 |
| GRIA1 | MGC | young | -0.467286374 | -1.93406033 | 1.94E-129 | 1.70E-21 |
| PTPRZ1 | MGC | young | -0.468051285 | -1.731531825 | 1.70E-121 | 1.63E-12 |
| RCBTB2 | MGC | young | -0.468719614 | -0.81018519 | 4.19E-15 | 0.005363017 |
| SUOX | MGC | young | -0.469934299 | -1.26676249 | 5.06E-07 | 0.002220681 |
| ADH5 | MGC | young | -0.469972549 | -0.751558072 | 7.10E-40 | 0.00034812 |
| RIOX2 | MGC | young | -0.470030705 | -0.892469146 | 5.35E-12 | 0.020976885 |
| AC099063.4 | MGC | young | -0.470417785 | -1.634103187 | 1.35E-17 | 0.000304463 |
| CCDC80 | MGC | young | -0.47088839 | -1.123651207 | 1.91E-36 | 4.40E-05 |
| NUDT3 | MGC | young | -0.47150993 | -0.872833913 | 1.22E-21 | 0.006188321 |
| TACR1 | MGC | young | -0.471609165 | -0.985940106 | 0.026306611 | 0.049266235 |
| AP001372.2 | MGC | young | -0.472381661 | -1.34854624 | 1.23E-13 | 0.000915387 |
| CNTN1 | MGC | young | -0.474208134 | -2.258136404 | 5.57E-110 | 3.28E-24 |
| SCRN3 | MGC | young | -0.476496306 | -0.960228521 | 9.64E-21 | 0.000537139 |
| TFDP1 | MGC | young | -0.478568445 | -0.976521412 | 2.78E-10 | 0.009151671 |
| GSPT2 | MGC | young | -0.479233293 | -0.640928547 | 2.09E-14 | 0.032171971 |
| ZNF429 | MGC | young | -0.479371735 | -1.027236466 | 3.08E-22 | 0.00020649 |
| HIRIP3 | MGC | young | -0.480080027 | -0.993694186 | 3.56E-08 | 0.009061452 |
| MAB21L1 | MGC | young | -0.48170449 | -1.229333331 | 3.84E-83 | 2.73E-10 |
| UBTF | MGC | young | -0.482469453 | -1.081143642 | 3.94E-20 | 0.001772377 |
| PXDNL | MGC | young | -0.48295043 | -1.940447957 | 4.77E-52 | 8.81E-11 |
| BX890604.2 | MGC | young | -0.48309221 | -1.092644713 | 0.002442247 | 0.0434859 |
| AL035701.1 | MGC | young | -0.484364785 | -0.896483101 | 0.020054898 | 0.014262458 |
| PNMA8B | MGC | young | -0.484581854 | -1.374570282 | 0.000275146 | 0.017828124 |
| CDNF | MGC | young | -0.485868542 | -0.961981737 | 1.50E-06 | 0.02872715 |
| SLC13A5 | MGC | young | -0.487332117 | -1.320784262 | 1.25E-11 | 0.001662405 |
| AL049775.1 | MGC | young | -0.487951116 | -1.08252274 | 0.000537727 | 0.029768218 |
| AC104850.2 | MGC | young | -0.488197201 | -0.96009949 | 0.000969558 | 0.025141368 |
| AC026780.2 | MGC | young | -0.488343174 | -1.53616086 | 0.000262549 | 0.026816954 |
| LINC02328 | MGC | young | -0.488547206 | -2.423730895 | 9.33E-38 | 3.42E-11 |

| | | | | | | |
|---|---|---|---|---|---|---|
| SPON1 | MGC | young | -0.491449469 | -1.697403747 | 1.69E-174 | 4.60E-16 |
| GUCY1A1 | MGC | young | -0.492590532 | -1.265041638 | 1.91E-24 | 0.005433681 |
| AC008462.1 | MGC | young | -0.49487972 | -1.05927605 | 1.23E-05 | 0.023688495 |
| CDH10 | MGC | young | -0.495304084 | -1.265852604 | 0.002293183 | 0.026645723 |
| IGSF9B | MGC | young | -0.495689278 | -0.854929226 | 1.21E-13 | 0.006178628 |
| AC137810.1 | MGC | young | -0.497183443 | -1.30148482 | 1.08E-06 | 0.016874278 |
| AL049874.3 | MGC | young | -0.497523288 | -1.105304057 | 7.63E-19 | 0.003580148 |
| PHYHIPL | MGC | young | -0.499332256 | -1.562070017 | 8.94E-132 | 1.80E-22 |
| AC099792.1 | MGC | young | -0.501845503 | -2.233439249 | 1.24E-72 | 3.15E-10 |
| GPR171 | MGC | young | -0.503954691 | -2.084200836 | 1.79E-14 | 2.20E-05 |
| ZBTB14 | MGC | young | -0.504372944 | -1.46288095 | 4.32E-14 | 0.000551221 |
| HAUS6 | MGC | young | -0.506324403 | -0.83396574 | 3.41E-09 | 0.035661955 |
| SEPTIN8 | MGC | young | -0.506827022 | -1.342515877 | 1.38E-55 | 1.13E-10 |
| OGFRL1 | MGC | young | -0.507293747 | -0.447977236 | 9.32E-23 | 0.038812454 |
| AC012404.1 | MGC | young | -0.508323875 | -1.857356626 | 8.65E-57 | 1.90E-12 |
| MT-ND5 | MGC | young | -0.509352509 | -1.691408765 | 2.16E-200 | 5.97E-14 |
| AC092802.1 | MGC | young | -0.509565296 | -1.086590471 | 1.73E-09 | 0.015894733 |
| MT-ND2 | MGC | young | -0.510668018 | -1.121893503 | 7.75E-102 | 5.62E-08 |
| RORB | MGC | young | -0.511384488 | -1.508220842 | 9.31E-207 | 1.32E-08 |
| AL592295.3 | MGC | young | -0.511775657 | -1.057875384 | 0.003112832 | 0.047716379 |
| ZNF660 | MGC | young | -0.512839958 | -0.779982468 | 4.38E-10 | 0.044542472 |
| ZMYM3 | MGC | young | -0.51336248 | -0.833473984 | 0.005841282 | 0.044664665 |
| SLC25A21 | MGC | young | -0.514806632 | -1.246039155 | 3.33E-14 | 0.001916913 |
| TEX261 | MGC | young | -0.516915468 | -0.793217788 | 1.75E-08 | 0.047873938 |
| COCH | MGC | young | -0.518007764 | -2.713612654 | 7.05E-41 | 8.49E-12 |
| NOTCH1 | MGC | young | -0.51823146 | -1.636384945 | 4.77E-26 | 1.00E-08 |
| AKAP7 | MGC | young | -0.519118899 | -1.357122959 | 5.68E-46 | 3.60E-08 |
| MSI1 | MGC | young | -0.520391815 | -1.057272441 | 8.28E-27 | 2.52E-06 |
| TMLHE-AS1 | MGC | young | -0.522142711 | -1.348920132 | 2.49E-09 | 0.008607855 |
| AL133415.1 | MGC | young | -0.522182259 | -1.403511026 | 0.000933479 | 0.020004425 |
| MAGEF1 | MGC | young | -0.522855678 | -0.770677747 | 2.47E-22 | 0.00344457 |
| CFAP70 | MGC | young | -0.525478258 | -1.606134036 | 2.59E-39 | 1.76E-07 |
| AC002429.2 | MGC | young | -0.526458435 | -1.04896094 | 9.99E-16 | 0.002798261 |
| CRYGD | MGC | young | -0.526474178 | -2.352644153 | 5.55E-35 | 1.62E-07 |
| NDP | MGC | young | -0.527467726 | -1.02385364 | 2.19E-30 | 1.94E-05 |
| PIGM | MGC | young | -0.527985161 | -0.79675577 | 9.02E-13 | 0.010483861 |
| AL136419.3 | MGC | young | -0.528191081 | -1.361892603 | 2.71E-10 | 0.00577603 |
| KLHL3 | MGC | young | -0.530305642 | -1.374451911 | 2.73E-34 | 1.21E-07 |
| NUF2 | MGC | young | -0.532521615 | -1.365256114 | 2.29E-11 | 0.003263903 |
| MIR99AHG | MGC | young | -0.532596472 | -1.494662638 | 2.59E-150 | 6.87E-15 |
| ALDH3A2 | MGC | young | -0.533428331 | -1.300484695 | 1.23E-76 | 1.33E-07 |
| LINC02250 | MGC | young | -0.535545165 | -1.246910499 | 3.78E-08 | 0.0151649 |
| RNF32 | MGC | young | -0.535580443 | -0.937207993 | 5.75E-05 | 0.022928798 |
| AC117834.1 | MGC | young | -0.535734683 | -1.360316093 | 1.17E-13 | 0.001234315 |
| AC092969.1 | MGC | young | -0.535788624 | -1.69343376 | 4.46E-23 | 8.10E-05 |
| ADAL | MGC | young | -0.538731 | -0.836794649 | 1.92E-10 | 0.022517228 |
| NORAD | MGC | young | -0.539948492 | -0.654955373 | 5.71E-48 | 0.007390391 |
| CCDC191 | MGC | young | -0.542176747 | -0.870483773 | 4.05E-23 | 0.006729233 |
| SOX2 | MGC | young | -0.542581177 | -1.412427911 | 3.34E-89 | 2.33E-15 |
| EPHA6 | MGC | young | -0.542929988 | -2.641210049 | 1.52E-74 | 1.93E-22 |
| COX18 | MGC | young | -0.544364802 | -1.102386012 | 2.87E-14 | 0.008543647 |
| NDRG4 | MGC | young | -0.545035798 | -0.971724331 | 1.63E-30 | 0.000192345 |
| PGM2 | MGC | young | -0.546312686 | -1.058037532 | 4.01E-64 | 1.32E-05 |
| AP001172.1 | MGC | young | -0.546315687 | -1.916915776 | 1.51E-33 | 2.32E-06 |

| | | | | | | |
|---|---|---|---|---|---|---|
| DTD2 | MGC | young | -0.54670146 | -0.922796781 | 3.08E-09 | 0.014653194 |
| TSBP1 | MGC | young | -0.546900602 | -1.32584054 | 4.16E-15 | 0.003697481 |
| NWD1 | MGC | young | -0.547148969 | -1.174589626 | 1.63E-06 | 0.02854424 |
| SSX2IP | MGC | young | -0.547240379 | -1.220598959 | 1.83E-89 | 3.86E-08 |
| AC016590.1 | MGC | young | -0.547288266 | -0.992272764 | 4.35E-07 | 0.040151073 |
| NRXN2 | MGC | young | -0.548125818 | -0.785724545 | 1.47E-07 | 0.019469352 |
| OARD1 | MGC | young | -0.54876264 | -0.967643367 | 5.25E-35 | 0.000217841 |
| ANKRD18B | MGC | young | -0.549064498 | -1.017489762 | 1.06E-05 | 0.021548464 |
| KIAA0100 | MGC | young | -0.549444929 | -0.885297631 | 5.34E-08 | 0.037429208 |
| TAFA1 | MGC | young | -0.550168684 | -1.477725249 | 8.81E-13 | 0.00120559 |
| SELENOP | MGC | young | -0.550639233 | -0.752286521 | 9.81E-52 | 0.003344527 |
| TTPA | MGC | young | -0.551913997 | -1.642261041 | 1.21E-22 | 5.52E-06 |
| AC005162.3 | MGC | young | -0.552231158 | -2.561651058 | 8.06E-17 | 0.000610006 |
| SGO2 | MGC | young | -0.554754803 | -1.443585342 | 2.15E-07 | 0.004228804 |
| CFAP298 | MGC | young | -0.555766773 | -0.605034187 | 9.34E-31 | 0.003021696 |
| C17orf75 | MGC | young | -0.556124062 | -0.688399449 | 9.84E-11 | 0.04173799 |
| ZBED3-AS1 | MGC | young | -0.556315675 | -1.230333408 | 1.14E-12 | 0.006477921 |
| CTNNBIP1 | MGC | young | -0.557224673 | -0.552652424 | 2.76E-25 | 0.016112818 |
| DENND11 | MGC | young | -0.557318993 | -2.40885515 | 2.05E-107 | 4.35E-23 |
| GAS1RR | MGC | young | -0.55762754 | -1.788051179 | 3.22E-24 | 3.76E-06 |
| TST | MGC | young | -0.558122426 | -0.933379058 | 1.09E-13 | 0.004797999 |
| GRAMD2B | MGC | young | -0.558674649 | -1.426797573 | 1.81E-43 | 2.00E-07 |
| PALMD | MGC | young | -0.558821079 | -0.791111799 | 1.23E-33 | 0.000797719 |
| CYP4V2 | MGC | young | -0.559240751 | -1.27115397 | 1.38E-90 | 2.57E-12 |
| AC211433.1 | MGC | young | -0.559750762 | -0.983782533 | 7.40E-11 | 0.037190481 |
| LTN1 | MGC | young | -0.560844923 | -0.579887052 | 2.91E-17 | 0.035966056 |
| ABCG1 | MGC | young | -0.562805299 | -1.120466634 | 3.84E-29 | 0.000332917 |
| ANKS4B | MGC | young | -0.565825406 | -1.263167002 | 0.011921359 | 0.047034967 |
| AC008945.2 | MGC | young | -0.566927961 | -1.202453959 | 1.25E-07 | 0.043742047 |
| AL138828.1 | MGC | young | -0.567378595 | -0.75257961 | 7.31E-13 | 0.038025587 |
| AQP4-AS1 | MGC | young | -0.56908806 | -2.033793594 | 1.19E-134 | 4.18E-17 |
| FAM217B | MGC | young | -0.569853799 | -1.463197638 | 3.90E-18 | 8.85E-05 |
| AC092944.1 | MGC | young | -0.571600388 | -0.900671836 | 7.98E-13 | 0.016280883 |
| AC093535.1 | MGC | young | -0.577134912 | -1.19584391 | 2.81E-08 | 0.021160104 |
| PCYOX1 | MGC | young | -0.577959707 | -0.865072889 | 4.36E-27 | 0.00049273 |
| ABCA9 | MGC | young | -0.579399762 | -1.428564345 | 2.37E-19 | 2.88E-05 |
| TTC32 | MGC | young | -0.579423349 | -1.102933752 | 3.60E-39 | 5.13E-06 |
| AC108472.1 | MGC | young | -0.58216112 | -1.630030932 | 1.11E-23 | 0.000310372 |
| LINC01833 | MGC | young | -0.584157126 | -0.845059239 | 6.13E-53 | 0.000749974 |
| ARHGEF37 | MGC | young | -0.584887931 | -1.210475195 | 1.47E-34 | 5.45E-07 |
| DYNLL2 | MGC | young | -0.585714861 | -0.962180531 | 2.02E-29 | 0.000162344 |
| MSANTD4 | MGC | young | -0.58740356 | -0.754986048 | 1.34E-52 | 0.00130813 |
| ITGA2 | MGC | young | -0.587467952 | -0.867845646 | 2.24E-21 | 0.014035247 |
| LINC02060 | MGC | young | -0.589517717 | -2.005374641 | 3.32E-13 | 3.85E-06 |
| LINC00923 | MGC | young | -0.593171632 | -1.108180177 | 3.78E-11 | 0.020836025 |
| TIMM21 | MGC | young | -0.593548767 | -0.857379785 | 1.50E-19 | 0.030303322 |
| TRDN-AS1 | MGC | young | -0.593837433 | -1.725938033 | 6.37E-66 | 5.75E-09 |
| RAX | MGC | young | -0.594589315 | -0.518262479 | 2.50E-44 | 0.007807846 |
| AC116903.2 | MGC | young | -0.596843866 | -1.275927128 | 0.001806653 | 0.03364497 |
| ADAMTS19 | MGC | young | -0.597069665 | -0.878018483 | 1.04E-15 | 0.017075025 |
| DDX59-AS1 | MGC | young | -0.599576594 | -1.053980356 | 2.38E-21 | 0.044674096 |
| C9orf129 | MGC | young | -0.599970931 | -1.361362691 | 4.98E-05 | 0.017519738 |
| NCKAP5-AS2 | MGC | young | -0.601055576 | -1.171631591 | 5.95E-05 | 0.032743981 |
| CFAP58 | MGC | young | -0.602565623 | -1.316213833 | 0.002286705 | 0.037931296 |

| | | | | | | |
|---|---|---|---|---|---|---|
| FGD5-AS1 | MGC | young | -0.60402209 | -0.635921019 | 8.19E-21 | 0.023581388 |
| AXIN2 | MGC | young | -0.604335026 | -1.152191429 | 3.61E-07 | 0.025574541 |
| MYO16 | MGC | young | -0.606840463 | -2.063715961 | 4.44E-32 | 6.20E-07 |
| HLF | MGC | young | -0.607511211 | -1.091900558 | 1.13E-44 | 2.70E-05 |
| KLHL14 | MGC | young | -0.607690615 | -2.264702157 | 1.06E-64 | 7.36E-11 |
| AVIL | MGC | young | -0.607914156 | -2.313785378 | 4.40E-26 | 4.04E-09 |
| LINC01659 | MGC | young | -0.608771234 | -1.720402201 | 1.55E-06 | 0.000681601 |
| AL133257.1 | MGC | young | -0.609577529 | -2.079807623 | 1.18E-66 | 2.80E-13 |
| TSPYL5 | MGC | young | -0.609675853 | -0.960890947 | 3.06E-08 | 0.038817288 |
| EPHA7 | MGC | young | -0.609942451 | -1.990434748 | 4.29E-08 | 0.000111629 |
| AC012085.2 | MGC | young | -0.610272406 | -0.775970205 | 4.95E-05 | 0.045397693 |
| AL136366.1 | MGC | young | -0.611550915 | -1.574095499 | 3.44E-56 | 3.25E-09 |
| AC035140.1 | MGC | young | -0.611789533 | -1.282558441 | 0.00143952 | 0.014572594 |
| TRDN | MGC | young | -0.612204543 | -0.702316881 | 1.69E-61 | 0.004640088 |
| AC108047.1 | MGC | young | -0.613724532 | -1.076776265 | 7.33E-07 | 0.031878316 |
| MKLN1-AS | MGC | young | -0.614573864 | -0.97548324 | 1.48E-06 | 0.024368095 |
| GRIK3 | MGC | young | -0.615587633 | -1.350830524 | 5.09E-32 | 5.88E-05 |
| SFTPD | MGC | young | -0.615997937 | -1.435484474 | 5.96E-08 | 0.007768047 |
| VSX2 | MGC | young | -0.617966881 | -2.124852558 | 8.44E-75 | 6.45E-21 |
| MT-ND4L | MGC | young | -0.619200144 | -1.209064157 | 8.94E-125 | 9.48E-10 |
| MYCN | MGC | young | -0.619588992 | -1.229110571 | 5.27E-06 | 0.013220585 |
| NKD1 | MGC | young | -0.620253047 | -2.149179047 | 4.68E-20 | 1.64E-05 |
| AC010978.1 | MGC | young | -0.620417144 | -1.417747423 | 0.008142903 | 0.046208669 |
| ABAT | MGC | young | -0.620934388 | -1.241932196 | 8.34E-29 | 4.14E-06 |
| AC009495.1 | MGC | young | -0.621455823 | -2.321976115 | 4.56E-26 | 3.78E-08 |
| SHMT1 | MGC | young | -0.622101016 | -0.612485824 | 5.25E-11 | 0.029344267 |
| PKP2 | MGC | young | -0.622590044 | -1.308245797 | 6.42E-16 | 0.000314147 |
| POLG2 | MGC | young | -0.626064395 | -0.933093092 | 0.00231198 | 0.03517229 |
| C14orf39 | MGC | young | -0.626825152 | -1.003108714 | 1.89E-38 | 0.000134831 |
| CLDN10-AS1 | MGC | young | -0.626849071 | -1.631084167 | 0.000451818 | 0.021050589 |
| FAM13C | MGC | young | -0.628600212 | -1.475381746 | 2.31E-21 | 0.000183941 |
| SLC15A2 | MGC | young | -0.629149153 | -1.046670969 | 3.55E-29 | 2.18E-05 |
| PRIM1 | MGC | young | -0.629785823 | -0.976834909 | 2.46E-08 | 0.02668604 |
| EFCAB2 | MGC | young | -0.631106537 | -0.589434139 | 1.31E-10 | 0.044877364 |
| KDR | MGC | young | -0.63131078 | -1.764184243 | 1.42E-153 | 1.74E-16 |
| CYS1 | MGC | young | -0.631765775 | -1.015168086 | 1.28E-11 | 0.005297412 |
| RAB9B | MGC | young | -0.632740023 | -1.286343327 | 9.58E-30 | 2.81E-05 |
| AC096719.1 | MGC | young | -0.632816012 | -1.201580468 | 6.54E-17 | 0.001740595 |
| MGP | MGC | young | -0.632845183 | -2.396243282 | 7.69E-47 | 1.48E-15 |
| DAG1 | MGC | young | -0.633844354 | -0.905180459 | 1.65E-19 | 0.003783166 |
| LINC00299 | MGC | young | -0.634525207 | -2.614593457 | 2.95E-65 | 1.11E-14 |
| LINC02388 | MGC | young | -0.634626485 | -1.215791406 | 2.76E-06 | 0.019253327 |
| CALM1 | MGC | young | -0.636326347 | -1.003767627 | 4.69E-99 | 1.97E-05 |
| KMT5A | MGC | young | -0.636916749 | -0.700453348 | 7.10E-08 | 0.038785866 |
| RASSF4 | MGC | young | -0.638519682 | -1.044006044 | 6.32E-78 | 7.57E-07 |
| SH3TC2-DT | MGC | young | -0.640714685 | -1.533521266 | 4.91E-11 | 0.000229253 |
| FBP1 | MGC | young | -0.641648944 | -1.269331818 | 3.34E-11 | 0.010710446 |
| PTCD2 | MGC | young | -0.641684297 | -0.930085946 | 1.76E-06 | 0.048743737 |
| PAX2 | MGC | young | -0.643387979 | -2.084673644 | 2.96E-18 | 7.66E-05 |
| SLC9A3-AS1 | MGC | young | -0.644626921 | -1.466525244 | 5.90E-06 | 0.00689095 |
| AC104532.2 | MGC | young | -0.645792248 | -1.378756917 | 0.004688348 | 0.017072852 |
| TMEM37 | MGC | young | -0.646919986 | -0.669055487 | 1.16E-19 | 0.005640198 |
| GCNT2 | MGC | young | -0.647889596 | -1.795488327 | 1.09E-16 | 1.77E-05 |
| MCFD2 | MGC | young | -0.648148169 | -0.938659013 | 1.42E-64 | 0.000227563 |

| | | | | | | |
|---|---|---|---|---|---|---|
| AC131571.1 | MGC | young | -0.648511176 | -1.732603649 | 2.14E-44 | 2.97E-08 |
| SLC16A2 | MGC | young | -0.649630219 | -1.122820536 | 5.29E-12 | 0.004900931 |
| AKR1C3 | MGC | young | -0.653103919 | -0.988183705 | 1.18E-60 | 9.34E-06 |
| SULF1 | MGC | young | -0.654173778 | -1.54977742 | 4.15E-165 | 1.56E-06 |
| FGL2 | MGC | young | -0.654748476 | -1.781206528 | 8.46E-15 | 0.000155469 |
| RASSF9 | MGC | young | -0.657284679 | -0.772683874 | 1.74E-27 | 0.01398476 |
| SAMD9L | MGC | young | -0.657354884 | -0.949300269 | 6.99E-20 | 0.023453174 |
| PKHD1 | MGC | young | -0.657638206 | -1.812759445 | 3.68E-31 | 9.36E-06 |
| C8orf37-AS1 | MGC | young | -0.657885444 | -1.318134735 | 1.30E-10 | 0.015742371 |
| EPHB6 | MGC | young | -0.65860448 | -2.140847577 | 1.03E-96 | 2.75E-12 |
| LINC00461 | MGC | young | -0.661030849 | -1.37636002 | 3.87E-163 | 1.11E-18 |
| AC024257.1 | MGC | young | -0.661509941 | -1.631378272 | 0.000368975 | 0.013784784 |
| ENKUR | MGC | young | -0.661668728 | -1.13115456 | 1.11E-27 | 0.000432739 |
| AC097515.1 | MGC | young | -0.663203229 | -2.187887325 | 2.58E-19 | 4.73E-07 |
| IDI2-AS1 | MGC | young | -0.663663049 | -0.681014415 | 2.49E-12 | 0.046168855 |
| SAMD13 | MGC | young | -0.666094281 | -1.53930045 | 8.65E-10 | 0.000709207 |
| AC013287.1 | MGC | young | -0.666154022 | -1.711231105 | 5.48E-25 | 2.96E-05 |
| AC010974.2 | MGC | young | -0.667812306 | -1.234023207 | 5.28E-11 | 0.044438368 |
| MT-ATP8 | MGC | young | -0.667908742 | -1.746930209 | 2.01E-197 | 3.64E-12 |
| CRPPA | MGC | young | -0.669341091 | -1.083324508 | 4.74E-13 | 0.019919774 |
| METTL7A | MGC | young | -0.669616862 | -0.7293023 | 1.08E-15 | 0.02424482 |
| ILDR2 | MGC | young | -0.670469419 | -2.403866462 | 1.37E-186 | 8.29E-18 |
| NYNRIN | MGC | young | -0.67215279 | -1.182125019 | 3.18E-05 | 0.022200239 |
| COX19 | MGC | young | -0.673367039 | -0.692522421 | 2.65E-20 | 0.012980059 |
| ETV1 | MGC | young | -0.673805461 | -1.922020024 | 6.80E-60 | 2.53E-10 |
| C5orf30 | MGC | young | -0.675176803 | -1.418562585 | 1.14E-31 | 5.05E-08 |
| DUSP19 | MGC | young | -0.67792538 | -1.121276149 | 5.72E-10 | 0.005504676 |
| FADS2 | MGC | young | -0.677974359 | -0.681011221 | 2.00E-07 | 0.04490647 |
| DHRS4-AS1 | MGC | young | -0.678317203 | -0.935753968 | 1.32E-13 | 0.007446218 |
| POU3F2 | MGC | young | -0.678670424 | -1.286453432 | 3.93E-08 | 0.01192974 |
| POP5 | MGC | young | -0.681512217 | -0.63321319 | 1.38E-19 | 0.029548975 |
| FZD8 | MGC | young | -0.683499478 | -0.907287556 | 3.73E-23 | 0.009344126 |
| KLKB1 | MGC | young | -0.683789513 | -3.498632405 | 4.18E-107 | 9.12E-24 |
| GPR63 | MGC | young | -0.691938046 | -1.265322724 | 5.23E-05 | 0.035278009 |
| FSCN1 | MGC | young | -0.692473383 | -0.501210416 | 6.89E-08 | 0.042757129 |
| INSL6 | MGC | young | -0.69349728 | -1.299093565 | 1.19E-08 | 0.020412359 |
| SYT11 | MGC | young | -0.693881162 | -1.510313647 | 8.10E-136 | 3.56E-15 |
| MYO10 | MGC | young | -0.694954774 | -1.938284996 | 2.04E-154 | 8.38E-22 |
| JDP2 | MGC | young | -0.696422933 | -0.933986003 | 3.89E-15 | 0.000695341 |
| CCDC141 | MGC | young | -0.702475648 | -2.183059659 | 2.05E-26 | 4.54E-06 |
| ASPA | MGC | young | -0.704215116 | -1.183957459 | 0.001244968 | 0.03646966 |
| SMIM10 | MGC | young | -0.706713912 | -0.992920376 | 2.40E-17 | 0.013102134 |
| MT-ND3 | MGC | young | -0.707106465 | -1.240112124 | 4.36E-136 | 1.61E-09 |
| SLC39A12 | MGC | young | -0.71161119 | -1.213048284 | 2.42E-36 | 1.25E-07 |
| CASC6 | MGC | young | -0.713722184 | -3.446254238 | 3.49E-51 | 2.00E-08 |
| GLIDR | MGC | young | -0.714378099 | -1.16207866 | 1.21E-12 | 0.004434816 |
| RBM43 | MGC | young | -0.714407096 | -1.157982804 | 2.60E-29 | 0.000550438 |
| AP001604.1 | MGC | young | -0.714857019 | -1.517572292 | 8.08E-06 | 0.016498475 |
| DIPK1C | MGC | young | -0.71545685 | -1.368284495 | 5.28E-58 | 6.53E-09 |
| PLPPR1 | MGC | young | -0.718457495 | -2.257807837 | 5.62E-24 | 3.57E-09 |
| AC025887.2 | MGC | young | -0.718956001 | -1.609572763 | 6.53E-16 | 0.000676544 |
| IFIT1 | MGC | young | -0.71959095 | -1.330543773 | 1.12E-39 | 4.14E-08 |
| MGARP | MGC | young | -0.719649611 | -0.774028057 | 6.66E-45 | 0.003253735 |
| PCYT1B | MGC | young | -0.720640918 | -2.206404764 | 2.27E-135 | 7.34E-19 |

| | | | | | | |
|---|---|---|---|---|---|---|
| CCDC196 | MGC | young | -0.721998455 | -1.134314844 | 8.40E-06 | 0.029257662 |
| S1PR1 | MGC | young | -0.722115477 | -1.228799795 | 5.90E-32 | 0.000297248 |
| LANCL1-AS1 | MGC | young | -0.722407971 | -1.485416521 | 4.55E-10 | 0.006953564 |
| AC100793.2 | MGC | young | -0.732629477 | -1.25085709 | 0.000642256 | 0.041664485 |
| AC104041.1 | MGC | young | -0.732868724 | -2.29345263 | 3.43E-66 | 1.91E-09 |
| C1orf141 | MGC | young | -0.734329459 | -0.836911424 | 3.44E-12 | 0.023581388 |
| HVCN1 | MGC | young | -0.734933031 | -1.052725096 | 9.35E-11 | 0.007333536 |
| SLC9C2 | MGC | young | -0.735546882 | -1.830220039 | 2.85E-16 | 0.000165081 |
| CDH7 | MGC | young | -0.739853968 | -2.105458377 | 5.81E-45 | 4.76E-11 |
| GPAM | MGC | young | -0.740374645 | -1.492362822 | 2.66E-15 | 0.002597771 |
| FNDC5 | MGC | young | -0.742807829 | -0.908543919 | 2.81E-10 | 0.044692072 |
| CCDC8 | MGC | young | -0.743895542 | -1.648149962 | 4.62E-07 | 0.001502476 |
| ST6GAL2 | MGC | young | -0.745438045 | -1.349104232 | 1.02E-12 | 0.031711464 |
| CDK5R1 | MGC | young | -0.748452689 | -1.141854803 | 4.14E-05 | 0.011827657 |
| AL035670.1 | MGC | young | -0.749043913 | -2.141975925 | 2.06E-12 | 7.34E-05 |
| KCTD21-AS1 | MGC | young | -0.750430884 | -1.397392939 | 4.35E-07 | 0.028141778 |
| DACH2 | MGC | young | -0.750859645 | -1.848863525 | 4.80E-25 | 1.03E-08 |
| MRPS34 | MGC | young | -0.754119069 | -0.614593624 | 8.45E-15 | 0.033649018 |
| AL353660.1 | MGC | young | -0.756786474 | -1.65662768 | 1.66E-08 | 0.001580984 |
| SERPINI1 | MGC | young | -0.757601856 | -0.691646309 | 1.32E-21 | 0.018328843 |
| AC069224.1 | MGC | young | -0.757823549 | -1.200063166 | 4.19E-05 | 0.034154273 |
| FANCI | MGC | young | -0.758131447 | -0.72270007 | 4.68E-07 | 0.044248498 |
| B3GAT1 | MGC | young | -0.75988685 | -1.917665859 | 2.59E-60 | 2.69E-13 |
| HNMT | MGC | young | -0.760246008 | -1.125668184 | 9.58E-30 | 0.000181897 |
| KLHDC8A | MGC | young | -0.761741325 | -2.619194563 | 5.23E-97 | 1.20E-14 |
| ABCB11 | MGC | young | -0.762000161 | -1.512162496 | 7.39E-13 | 7.64E-05 |
| AC006115.2 | MGC | young | -0.76205434 | -1.590002129 | 1.18E-24 | 6.98E-07 |
| LINC01229 | MGC | young | -0.763551883 | -1.550051715 | 2.01E-10 | 0.00235342 |
| KHDRBS2 | MGC | young | -0.769478705 | -3.498282003 | 2.12E-272 | 1.10E-30 |
| NFIA-AS2 | MGC | young | -0.77015241 | -1.412664211 | 1.51E-14 | 0.0006489 |
| COL24A1 | MGC | young | -0.770503385 | -3.830943956 | 1.20E-140 | 1.55E-35 |
| AMACR | MGC | young | -0.770642887 | -2.61199016 | 2.09E-113 | 3.39E-24 |
| AC008555.4 | MGC | young | -0.771420441 | -1.666222868 | 7.86E-09 | 0.000276035 |
| PVALEF | MGC | young | -0.7718127 | -1.313124626 | 0.02762284 | 0.029758782 |
| ELP6 | MGC | young | -0.772489838 | -0.780348965 | 2.69E-18 | 0.005726267 |
| CLRN1 | MGC | young | -0.774757735 | -1.57009401 | 7.55E-137 | 2.09E-15 |
| MAF | MGC | young | -0.776681751 | -0.884772095 | 4.66E-46 | 0.00018618 |
| LINC00901 | MGC | young | -0.77874469 | -1.363346807 | 0.00055732 | 0.01565897 |
| AC104232.3 | MGC | young | -0.782411312 | -1.341222815 | 0.000243119 | 0.009603385 |
| AC025284.1 | MGC | young | -0.78329731 | -1.754351631 | 0.002982803 | 0.017321079 |
| NSMCE4A | MGC | young | -0.783810331 | -1.23887687 | 2.79E-24 | 6.54E-05 |
| AL139807.1 | MGC | young | -0.786357671 | -0.92006702 | 1.73E-07 | 0.017674811 |
| AC092924.2 | MGC | young | -0.787681443 | -2.29366035 | 2.04E-86 | 9.02E-17 |
| TRAM2-AS1 | MGC | young | -0.789228617 | -0.954370602 | 2.34E-11 | 0.031570635 |
| SPRY2 | MGC | young | -0.793381691 | -0.585874256 | 2.89E-40 | 0.023636294 |
| KDM4D | MGC | young | -0.794726261 | -1.179334376 | 0.001008897 | 0.028570507 |
| PRSS48 | MGC | young | -0.795136053 | -1.05790681 | 0.002238737 | 0.043194462 |
| AL078587.2 | MGC | young | -0.795356719 | -1.938127132 | 1.95E-35 | 3.28E-05 |
| AC091826.2 | MGC | young | -0.795435666 | -1.945607428 | 1.40E-32 | 1.08E-09 |
| AC004918.1 | MGC | young | -0.795914302 | -2.570766682 | 1.15E-39 | 1.19E-14 |
| CYP7B1 | MGC | young | -0.796249004 | -0.939382429 | 2.80E-12 | 0.046473343 |
| SULT1A1 | MGC | young | -0.80185882 | -0.810383904 | 4.67E-24 | 0.000974925 |
| TUSC7 | MGC | young | -0.80245987 | -1.514062199 | 2.01E-09 | 0.008969005 |
| SPRY1 | MGC | young | -0.803200735 | -1.787559356 | 1.26E-75 | 7.43E-15 |

| | | | | | | |
|---|---|---|---|---|---|---|
| IRX5 | MGC | young | -0.804498496 | -2.112240244 | 5.19E-37 | 1.32E-12 |
| LRIT3 | MGC | young | -0.804647007 | -0.94569851 | 9.59E-09 | 0.028499858 |
| LRP1 | MGC | young | -0.806343208 | -1.424101388 | 8.47E-90 | 4.06E-13 |
| CETN3 | MGC | young | -0.812025256 | -1.070039189 | 2.71E-55 | 1.06E-05 |
| AC108169.1 | MGC | young | -0.813406323 | -1.538834245 | 8.69E-05 | 0.017637275 |
| TRIM29 | MGC | young | -0.814364459 | -1.382226823 | 4.26E-15 | 0.008511153 |
| AKR7A2 | MGC | young | -0.816599385 | -0.488418597 | 3.40E-17 | 0.037205229 |
| HHIPL1 | MGC | young | -0.818046734 | -1.031957929 | 0.000365732 | 0.039318762 |
| FZD2 | MGC | young | -0.818447875 | -1.474036736 | 9.11E-05 | 0.002537885 |
| AC092691.3 | MGC | young | -0.820942285 | -1.579097818 | 1.16E-17 | 8.43E-05 |
| AL451123.1 | MGC | young | -0.823768863 | -1.0686288 | 3.98E-05 | 0.047882094 |
| KCNJ13 | MGC | young | -0.825737823 | -1.272875402 | 1.12E-10 | 6.72E-05 |
| LINC02774 | MGC | young | -0.827775263 | -1.65076146 | 1.79E-10 | 0.000369204 |
| C4orf50 | MGC | young | -0.830608875 | -2.044476353 | 2.69E-18 | 2.83E-05 |
| SOX9 | MGC | young | -0.832662031 | -0.850505122 | 3.42E-38 | 0.00016008 |
| AC063979.2 | MGC | young | -0.833147422 | -1.583616692 | 2.58E-08 | 0.002467306 |
| AC053513.1 | MGC | young | -0.833361772 | -1.395150393 | 2.52E-18 | 0.000247971 |
| TM7SF2 | MGC | young | -0.833775895 | -1.15732421 | 5.44E-34 | 0.000468287 |
| DHRS4L2 | MGC | young | -0.834590881 | -0.839532663 | 6.12E-18 | 0.00621958 |
| ADAMTS18 | MGC | young | -0.838426136 | -1.914645943 | 3.41E-63 | 7.58E-06 |
| CCND1 | MGC | young | -0.841042346 | -0.77020283 | 5.06E-20 | 0.009186193 |
| LIX1-AS1 | MGC | young | -0.844446884 | -1.651221062 | 3.70E-18 | 2.21E-05 |
| PEG10 | MGC | young | -0.846746861 | -0.840692334 | 4.73E-15 | 0.030332799 |
| CCNI2 | MGC | young | -0.847341067 | -1.381916233 | 7.46E-05 | 0.021751393 |
| TENM3-AS1 | MGC | young | -0.847850469 | -1.477663636 | 9.98E-26 | 4.46E-08 |
| ATXN7L3B | MGC | young | -0.850367364 | -0.802971695 | 1.18E-56 | 0.000601923 |
| ZBTB47 | MGC | young | -0.852975758 | -1.188160048 | 9.59E-08 | 0.014148493 |
| ECT2L | MGC | young | -0.854099222 | -2.102783585 | 2.04E-26 | 3.45E-07 |
| MYH15 | MGC | young | -0.854857533 | -1.805404834 | 1.09E-24 | 3.92E-06 |
| AC005746.2 | MGC | young | -0.859166115 | -1.314738287 | 2.72E-07 | 0.01861419 |
| TMEM121 | MGC | young | -0.860373547 | -2.036427892 | 3.29E-30 | 7.74E-08 |
| AC097537.1 | MGC | young | -0.868163049 | -1.651257034 | 7.26E-19 | 0.000831344 |
| FSIP2 | MGC | young | -0.88114272 | -1.828496144 | 5.41E-32 | 6.23E-06 |
| RGS7BP | MGC | young | -0.885618845 | -1.592297731 | 2.03E-16 | 0.00029424 |
| AC021086.1 | MGC | young | -0.886768073 | -1.368544482 | 5.30E-11 | 0.003054274 |
| GAS1 | MGC | young | -0.887242349 | -1.749691263 | 3.14E-41 | 2.07E-12 |
| CMPK2 | MGC | young | -0.89280415 | -1.521043224 | 0.000337425 | 0.006399921 |
| AC007091.1 | MGC | young | -0.89410996 | -1.94057975 | 0.027027098 | 0.010483861 |
| MCMDC2 | MGC | young | -0.898137011 | -1.150283311 | 0.010662371 | 0.04392668 |
| AC090138.1 | MGC | young | -0.902436906 | -1.566106493 | 1.00E-19 | 0.000129637 |
| AHCYL2 | MGC | young | -0.904447915 | -0.800454198 | 6.37E-46 | 0.000128466 |
| LINC00632 | MGC | young | -0.907657792 | -0.748615589 | 5.86E-43 | 0.000696842 |
| AC107204.1 | MGC | young | -0.912865852 | -1.59994012 | 7.88E-20 | 0.001322674 |
| AC005856.1 | MGC | young | -0.920858543 | -1.84672286 | 8.29E-23 | 0.000116412 |
| LINC02315 | MGC | young | -0.920964317 | -2.156359542 | 1.32E-20 | 5.87E-06 |
| AL157392.2 | MGC | young | -0.925304391 | -2.071180705 | 2.89E-09 | 0.000725627 |
| DTX4 | MGC | young | -0.927013094 | -0.964622 | 5.25E-10 | 0.002101175 |
| AC011246.1 | MGC | young | -0.933520045 | -1.505088872 | 0.000401066 | 0.019208336 |
| AF123462.1 | MGC | young | -0.935645184 | -2.784896207 | 5.83E-29 | 3.81E-11 |
| FZD5 | MGC | young | -0.942502512 | -1.505207704 | 4.68E-59 | 3.06E-10 |
| LINC01697 | MGC | young | -0.949045685 | -1.657536403 | 7.26E-51 | 0.000643607 |
| HCG11 | MGC | young | -0.949089709 | -1.473012894 | 1.02E-18 | 0.003096907 |
| AL354733.3 | MGC | young | -0.960490779 | -1.27198289 | 8.11E-12 | 0.01169528 |
| AC008957.2 | MGC | young | -0.976316016 | -1.518243639 | 4.27E-71 | 3.65E-06 |

| | | | | | | |
|---|---|---|---|---|---|---|
| CYHR1 | MGC | young | -0.978541351 | -0.689413642 | 8.35E-29 | 0.014104606 |
| HMGN5 | MGC | young | -0.98894898 | -1.79000544 | 5.00E-23 | 9.31E-09 |
| AC008957.1 | MGC | young | -0.998580102 | -1.722445073 | 3.70E-25 | 1.64E-05 |
| SIX6 | MGC | young | -0.999107759 | -0.537131577 | 2.33E-25 | 0.023581388 |
| AC009315.1 | MGC | young | -1.008976619 | -1.807029258 | 1.09E-08 | 0.003985722 |
| CCDC89 | MGC | young | -1.009669378 | -1.920199286 | 3.31E-11 | 0.000472299 |
| ABCA9-AS1 | MGC | young | -1.010510976 | -1.783334369 | 7.54E-12 | 0.000690552 |
| AC008277.1 | MGC | young | -1.015031421 | -1.695572259 | 5.86E-15 | 0.003926928 |
| AC078820.2 | MGC | young | -1.017967132 | -2.081590639 | 7.43E-26 | 6.40E-08 |
| AJ006995.1 | MGC | young | -1.020978321 | -1.397590149 | 5.03E-22 | 0.001412146 |
| AL355001.2 | MGC | young | -1.023507504 | -1.256716217 | 1.29E-07 | 0.006940195 |
| BBOX1 | MGC | young | -1.024143839 | -1.118989172 | 0.000724412 | 0.022589 |
| AC022146.2 | MGC | young | -1.02654109 | -1.337880802 | 2.16E-17 | 0.001301511 |
| DUSP27 | MGC | young | -1.037532587 | -2.019403583 | 0.000272823 | 0.003983888 |
| MSTN | MGC | young | -1.040332753 | -2.77966432 | 4.08E-45 | 2.52E-11 |
| AL445259.1 | MGC | young | -1.046222554 | -2.901087918 | 1.79E-35 | 2.16E-09 |
| FGF14-AS2 | MGC | young | -1.049803916 | -1.518339337 | 1.40E-21 | 8.54E-07 |
| MTTP | MGC | young | -1.050644531 | -2.214050845 | 1.58E-157 | 2.72E-24 |
| AC246817.2 | MGC | young | -1.057529533 | -2.044690142 | 3.14E-20 | 0.000224473 |
| SMAD6 | MGC | young | -1.060676031 | -1.326943087 | 3.41E-11 | 0.001750105 |
| CNMD | MGC | young | -1.064507021 | -0.968128223 | 9.31E-32 | 0.000351708 |
| HIST1H4C | MGC | young | -1.070420127 | -0.88084679 | 1.05E-57 | 4.77E-05 |
| AL138760.1 | MGC | young | -1.071298746 | -1.726390286 | 1.50E-11 | 0.001562288 |
| PDGFRA | MGC | young | -1.078661609 | -2.730199028 | 3.61E-25 | 2.56E-12 |
| RERG | MGC | young | -1.07872599 | -1.795689459 | 7.70E-15 | 8.64E-05 |
| KCNA2 | MGC | young | -1.079157227 | -2.786564558 | 7.54E-54 | 2.65E-16 |
| AQP4 | MGC | young | -1.080586497 | -1.336483871 | 4.89E-92 | 4.29E-11 |
| AC006160.1 | MGC | young | -1.091239862 | -1.178864999 | 1.09E-10 | 0.016937509 |
| LINC02389 | MGC | young | -1.09585776 | -3.454988164 | 3.84E-41 | 2.28E-11 |
| PDGFD | MGC | young | -1.105673389 | -2.244565684 | 2.87E-34 | 4.72E-07 |
| KCNJ10 | MGC | young | -1.118558362 | -2.225968207 | 8.70E-54 | 6.95E-15 |
| PTPN7 | MGC | young | -1.139056522 | -1.644076842 | 0.000134992 | 0.011305563 |
| CYP4F26P | MGC | young | -1.13913585 | -1.893853808 | 0.00414798 | 0.008889638 |
| TMEM71 | MGC | young | -1.147853701 | -1.732719891 | 9.50E-16 | 0.000970028 |
| P2RY12 | MGC | young | -1.162004281 | -1.898339177 | 1.74E-20 | 8.55E-05 |
| BX664615.2 | MGC | young | -1.163015357 | -1.482764054 | 3.69E-31 | 6.86E-05 |
| COL2A1 | MGC | young | -1.164568592 | -1.191939673 | 8.79E-16 | 0.000222153 |
| OAS1 | MGC | young | -1.166055492 | -1.273051689 | 2.16E-07 | 0.021188834 |
| AL360178.1 | MGC | young | -1.177181265 | -1.923259415 | 8.23E-23 | 4.15E-05 |
| AL390957.1 | MGC | young | -1.181241722 | -2.37421622 | 7.38E-110 | 1.55E-10 |
| AL160262.1 | MGC | young | -1.182642492 | -1.518690651 | 1.54E-05 | 0.026502824 |
| AC007513.1 | MGC | young | -1.184208778 | -2.808829833 | 2.76E-23 | 1.19E-08 |
| AC135895.1 | MGC | young | -1.190052106 | -2.285147377 | 6.45E-11 | 0.00252582 |
| AC245123.1 | MGC | young | -1.194351069 | -3.189820696 | 6.97E-27 | 3.40E-06 |
| AC083805.1 | MGC | young | -1.202139581 | -1.216264299 | 0.002346971 | 0.030155097 |
| HERC6 | MGC | young | -1.203211853 | -1.692858969 | 2.22E-18 | 0.001806669 |
| OMG | MGC | young | -1.204537784 | -2.261574555 | 5.75E-73 | 2.44E-19 |
| GRIA4 | MGC | young | -1.212988882 | -2.369469255 | 9.47E-160 | 4.74E-22 |
| DIAPH3 | MGC | young | -1.213574706 | -1.423351248 | 1.38E-05 | 0.003096907 |
| ANGPTL1 | MGC | young | -1.214553287 | -2.020031638 | 3.27E-133 | 4.73E-14 |
| LINC02231 | MGC | young | -1.219789838 | -3.134827704 | 1.40E-43 | 4.50E-12 |
| CCDC26 | MGC | young | -1.22101852 | -2.303647467 | 2.80E-09 | 0.00010754 |
| LINC00290 | MGC | young | -1.233182252 | -1.769027745 | 5.25E-55 | 1.07E-11 |
| LINC00707 | MGC | young | -1.25533519 | -1.713005287 | 2.93E-07 | 0.008192756 |

| | | | | | | |
|---|---|---|---|---|---|---|
| AC073571.1 | MGC | young | -1.270270368 | -2.030444431 | 8.30E-11 | 0.000237717 |
| LINC01625 | MGC | young | -1.272903757 | -1.352138542 | 2.18E-08 | 0.015902927 |
| AC245187.2 | MGC | young | -1.274212336 | -2.76347746 | 2.95E-24 | 2.08E-05 |
| AL139142.1 | MGC | young | -1.281398314 | -1.585435696 | 1.25E-06 | 0.014556241 |
| AL627171.2 | MGC | young | -1.282947004 | -0.698659832 | 7.30E-25 | 0.019949519 |
| LINC02696 | MGC | young | -1.286568332 | -1.094263252 | 1.09E-13 | 0.002890586 |
| ZPLD1 | MGC | young | -1.30534217 | -1.762166581 | 5.49E-36 | 1.29E-06 |
| MT-ND6 | MGC | young | -1.35079546 | -3.347999692 | 8.38506331312 | 8.32E-44 |
| PLA2G5 | MGC | young | -1.376920813 | -1.645321903 | 1.60E-08 | 0.000646222 |
| CNTN4-AS2 | MGC | young | -1.39940875 | -1.447275091 | 9.69E-07 | 0.017287044 |
| FAM153A | MGC | young | -1.402226827 | -2.683565711 | 1.95E-16 | 0.000438692 |
| XAF1 | MGC | young | -1.410380812 | -2.017879259 | 1.63E-21 | 0.001894209 |
| ZNF385D-AS1 | MGC | young | -1.450301484 | -1.630642402 | 0.018964942 | 0.033164375 |
| TNFRSF21 | MGC | young | -1.495056798 | -1.763530733 | 1.89E-38 | 0.001250587 |
| FAM181B | MGC | young | -1.500334414 | -1.482430035 | 2.96E-11 | 0.007320332 |
| AC111194.2 | MGC | young | -1.500675214 | -1.911503991 | 5.33E-12 | 0.000399479 |
| AC007106.2 | MGC | young | -1.532701389 | -2.234618347 | 3.81E-14 | 0.000269179 |
| RASSF1-AS1 | MGC | young | -1.559814263 | -1.586281013 | 0.000938701 | 0.017881512 |
| LINC01673 | MGC | young | -1.592442325 | -1.694790724 | 1.95E-10 | 0.013562316 |
| RGS18 | MGC | young | -1.724072612 | -2.393675616 | 2.26E-24 | 5.45E-05 |
| SRGAP3-AS4 | MGC | young | -1.753246451 | -1.392000069 | 0.005185356 | 0.030696325 |
| LINC02500 | MGC | young | -1.811706555 | -1.551057247 | 1.18E-24 | 7.19E-06 |
| LINC00113 | MGC | young | -1.818752838 | -1.612626549 | 2.44E-12 | 0.021362983 |
| CDR1 | MGC | young | -1.850399944 | -1.056302592 | 1.03E-16 | 0.003436441 |
| AL355347.1 | MGC | young | -2.011621271 | -3.527365228 | 2.66E-34 | 0.000103995 |
| HIST1H1E | MGC | young | -2.01976412 | -1.143299703 | 5.83E-21 | 0.002728525 |
| AC108516.1 | MGC | young | -2.065172643 | -2.696816215 | 1.63E-50 | 9.91E-15 |
| PMP2 | MGC | young | -2.197619703 | -1.774779887 | 6.69E-11 | 0.011372521 |
| AC023866.2 | MGC | young | -2.36427934 | -2.485114184 | 6.21E-06 | 0.005744214 |
| RGS21 | MGC | young | -2.42820279 | -2.754107926 | 1.09E-45 | 5.82E-10 |
| AC116407.1 | MGC | young | -3.010897318 | -1.525221137 | 1.74E-09 | 0.00012326 |
| AC093151.8 | MGC | young | -4.400441161 | -2.863013521 | 2.46E-27 | 0.000683682 |
| MT3 | Microgliaold | | 4.69365585 | 5.046116923 | 1.48E-39 | 5.27E-29 |
| MT1G | Microgliaold | | 4.374136623 | 1.359974149 | 1.43E-35 | 0.014656508 |
| MT1E | Microgliaold | | 4.191686264 | 3.355922583 | 7.15E-33 | 3.77E-16 |
| THBS1 | Microgliaold | | 3.953207739 | 3.070734235 | 1.15E-38 | 8.85E-06 |
| ADAM8 | Microgliaold | | 3.760159279 | 2.982199314 | 6.90E-49 | 3.29E-05 |
| SLC16A10 | Microgliaold | | 3.714418557 | 1.664289711 | 9.24E-47 | 0.002738448 |
| MT1X | Microgliaold | | 3.350121235 | 3.202129904 | 6.58E-40 | 8.65E-12 |
| HILPDA | Microgliaold | | 2.750441856 | 3.17128703 | 2.13E-23 | 2.24E-08 |
| PMP22 | Microgliaold | | 2.74354374 | 2.146232785 | 4.42E-36 | 0.000710755 |
| PLOD2 | Microgliaold | | 2.712860028 | 2.001314772 | 5.02E-11 | 0.009468108 |
| SDC2 | Microgliaold | | 2.701938279 | 1.913367008 | 2.20E-18 | 0.011721361 |
| MT1M | Microgliaold | | 2.672312927 | 1.553105923 | 0.000526218 | 0.032940913 |
| BNIP3 | Microgliaold | | 2.548348235 | 2.73910071 | 7.42E-70 | 1.11E-08 |
| RAB42 | Microgliaold | | 2.485226459 | 2.414013226 | 4.13E-24 | 0.000394183 |
| MT2A | Microgliaold | | 2.43656439 | 2.628879948 | 2.19E-93 | 3.78E-07 |
| ERO1A | Microgliaold | | 2.340979416 | 2.865265116 | 1.50E-77 | 1.27E-10 |
| TIMP1 | Microgliaold | | 2.149739113 | 0.972378035 | 4.77E-10 | 0.020160979 |
| SLC2A1 | Microgliaold | | 2.141193927 | 2.173654125 | 2.27E-25 | 0.000390653 |
| TNFRSF11B | Microgliaold | | 2.104054779 | 2.957405152 | 1.84E-25 | 4.85E-08 |
| PPP1R3C | Microgliaold | | 1.887834147 | 2.085551478 | 1.57E-12 | 0.000806027 |
| GFOD1 | Microgliaold | | 1.875947811 | 1.828299777 | 1.17E-13 | 0.03372426 |
| CRYAB | Microgliaold | | 1.867311711 | 1.821682346 | 6.42E-14 | 0.001229551 |

| | | | | | |
|---|---|---|---|---|---|
| NUPR1 | Microgliaold | 1.845935463 | 3.259606907 | 1.72E-101 | 3.30E-18 |
| SLC35F1 | Microgliaold | 1.831220393 | 2.348166984 | 1.04E-89 | 5.05E-07 |
| NDRG1 | Microgliaold | 1.778202094 | 1.477884952 | 9.76E-22 | 0.015684259 |
| CYP27A1 | Microgliaold | 1.766936336 | 1.280415589 | 1.82E-09 | 0.024694588 |
| SERPINH1 | Microgliaold | 1.672508525 | 3.142280253 | 1.44E-16 | 1.39E-09 |
| AC131944.1 | Microgliaold | 1.648389559 | 1.161513409 | 1.51E-09 | 0.001341545 |
| HIF1A-AS3 | Microgliaold | 1.634809588 | 1.262231405 | 4.82E-09 | 0.005248184 |
| GPNMB | Microgliaold | 1.627232507 | 1.840422277 | 1.49E-56 | 3.56E-08 |
| GPI | Microgliaold | 1.585670041 | 1.349920577 | 3.68E-07 | 0.033833975 |
| PIM1 | Microgliaold | 1.580193228 | 1.727652167 | 9.73E-17 | 0.000434657 |
| HSPA6 | Microgliaold | 1.567051845 | 2.347477207 | 4.05E-19 | 0.002091748 |
| PLTP | Microgliaold | 1.563533498 | 2.280245108 | 1.38E-20 | 8.93E-08 |
| ID3 | Microgliaold | 1.542454931 | 2.260690143 | 1.25E-13 | 7.03E-05 |
| AL133453.1 | Microgliaold | 1.498909585 | 1.362103097 | 1.40E-23 | 0.021873478 |
| SPP1 | Microgliaold | 1.493610574 | 1.307654729 | 1.98E-56 | 5.39E-06 |
| SLC2A3 | Microgliaold | 1.457484008 | 2.205698685 | 8.61E-39 | 1.57E-11 |
| PAM | Microgliaold | 1.438671067 | 1.669080845 | 2.25E-14 | 0.019426532 |
| FOSL2 | Microgliaold | 1.40315248 | 1.135881104 | 1.05E-11 | 0.047451454 |
| SNHG12 | Microgliaold | 1.359993881 | 2.67992535 | 3.02E-45 | 1.35E-09 |
| PGK1 | Microgliaold | 1.329591266 | 1.487638217 | 4.32E-37 | 9.76E-06 |
| CTSB | Microgliaold | 1.32252257 | 1.072566293 | 3.07E-10 | 0.000272749 |
| TGFBI | Microgliaold | 1.29115704 | 1.008372113 | 6.72E-05 | 0.042144647 |
| MXI1 | Microgliaold | 1.255035753 | 1.458715599 | 7.30E-11 | 0.038021383 |
| GLUL | Microgliaold | 1.254040153 | 1.626584748 | 1.92E-33 | 1.09E-06 |
| FABP5 | Microgliaold | 1.239422161 | 1.17068378 | 0.000765675 | 0.003550036 |
| FAM13A | Microgliaold | 1.236606269 | 1.683330096 | 0.000107787 | 0.002368834 |
| GBE1 | Microgliaold | 1.236451337 | 1.724851425 | 3.74E-19 | 0.000396021 |
| PLIN2 | Microgliaold | 1.221661924 | 2.95548422 | 4.24E-72 | 1.83E-14 |
| MT1F | Microgliaold | 1.206195659 | 1.610062331 | 5.10E-24 | 0.017800307 |
| C15orf48 | Microgliaold | 1.194519563 | 2.252379683 | 2.10E-45 | 1.36E-08 |
| RNASE4 | Microgliaold | 1.188230645 | 2.391857372 | 1.19E-38 | 6.61E-06 |
| KYNU | Microgliaold | 1.173072381 | 0.956107852 | 7.04E-08 | 0.025597542 |
| VMO1 | Microgliaold | 1.161437504 | 1.606008734 | 1.92E-24 | 7.44E-05 |
| MIF | Microgliaold | 1.139708941 | 2.10877669 | 8.66E-74 | 8.65E-12 |
| GOLIM4 | Microgliaold | 1.135674314 | 1.621658203 | 4.44E-13 | 0.006169484 |
| EMP2 | Microgliaold | 1.133814261 | 1.666949015 | 7.62E-09 | 0.005300428 |
| AUTS2 | Microgliaold | 1.102502021 | 1.619933397 | 2.05E-16 | 0.000716415 |
| CLIC2 | Microgliaold | 1.091421545 | 1.892776808 | 3.48E-13 | 0.000701024 |
| ENO1 | Microgliaold | 1.077819067 | 1.268620275 | 1.60E-27 | 0.0001199 |
| BNIP3L | Microgliaold | 1.072898914 | 1.802743582 | 5.27E-42 | 9.76E-06 |
| SMPDL3A | Microgliaold | 1.056395501 | 1.591151273 | 2.73E-14 | 0.011721361 |
| ZFAND2A | Microgliaold | 1.054495852 | 2.121356554 | 5.18E-08 | 3.39E-06 |
| FAM162A | Microgliaold | 1.047891879 | 2.545670237 | 4.41E-63 | 1.13E-11 |
| RGS16 | Microgliaold | 1.025192796 | 1.213726272 | 1.37E-10 | 0.016657562 |
| FCGR2B | Microgliaold | 1.007743553 | 1.592549888 | 3.91E-39 | 0.000112695 |
| CACYBP | Microgliaold | 0.997652079 | 1.793389077 | 3.10E-09 | 0.00020699 |
| TMEM176A | Microgliaold | 0.980856381 | 1.108053223 | 1.36E-07 | 0.018470788 |
| P4HA1 | Microgliaold | 0.96673104 | 2.219370307 | 3.32E-54 | 2.06E-09 |
| HK2 | Microgliaold | 0.966184717 | 1.719273338 | 3.35E-23 | 0.000110824 |
| ANG | Microgliaold | 0.964679364 | 2.164561125 | 1.74E-29 | 6.60E-05 |
| APP | Microgliaold | 0.960767772 | 1.521875477 | 0.001093627 | 0.005462267 |
| LGALS3 | Microgliaold | 0.930610796 | 1.00561552 | 1.33E-15 | 0.04116112 |
| UGCG | Microgliaold | 0.925567851 | 1.623220284 | 6.21E-15 | 5.52E-05 |
| AVPI1 | Microgliaold | 0.916312771 | 2.241603599 | 9.63E-11 | 0.013384232 |

| | | | | | |
|---|---|---|---|---|---|
| PRKAG2 | Microgliaold | 0.9074156 | 1.338213517 | 1.64E-06 | 0.025332768 |
| CA2 | Microgliaold | 0.900358433 | 1.404438403 | 0.003047442 | 0.019560202 |
| TMEM176B | Microgliaold | 0.899418027 | 1.221285313 | 2.13E-13 | 0.004035511 |
| MPZL1 | Microgliaold | 0.895951165 | 1.312961759 | 6.13E-07 | 0.033440149 |
| GAPDH | Microgliaold | 0.891174142 | 2.377425092 | 4.96E-91 | 3.70E-16 |
| SLC16A3 | Microgliaold | 0.878244793 | 1.15818463 | 8.78E-09 | 0.020141023 |
| HEBP2 | Microgliaold | 0.84729912 | 1.248611269 | 1.67E-08 | 0.039196869 |
| S100A10 | Microgliaold | 0.832409602 | 1.869885786 | 2.11E-65 | 9.91E-10 |
| UPP1 | Microgliaold | 0.810654498 | 1.872996585 | 2.67E-36 | 9.91E-05 |
| CSTB | Microgliaold | 0.808172991 | 2.225228373 | 4.18E-36 | 4.12E-13 |
| SEC61G | Microgliaold | 0.806548178 | 1.691563845 | 1.85E-36 | 7.08E-06 |
| LGALS1 | Microgliaold | 0.787721184 | 1.506626895 | 1.16E-33 | 3.78E-07 |
| FNDC3A | Microgliaold | 0.775226365 | 1.374091222 | 5.55E-21 | 0.014656508 |
| SYNGR2 | Microgliaold | 0.748062941 | 1.136943192 | 8.31E-06 | 0.030111122 |
| CYTH1 | Microgliaold | 0.747041762 | 1.131577493 | 1.19E-07 | 0.033138945 |
| EPB41L3 | Microgliaold | 0.745446415 | 1.224787272 | 6.20E-12 | 0.001892572 |
| GRINA | Microgliaold | 0.74430862 | 1.56278335 | 2.68E-16 | 0.002672744 |
| TPI1 | Microgliaold | 0.734539172 | 1.531545245 | 9.24E-40 | 1.04E-06 |
| ARL4C | Microgliaold | 0.709336768 | 1.902623135 | 5.27E-37 | 0.000525414 |
| SLA | Microgliaold | 0.699391499 | 1.185926527 | 0.002610733 | 0.012378513 |
| EIF4EBP1 | Microgliaold | 0.676840075 | 1.347237916 | 3.21E-18 | 0.011721361 |
| VKORC1 | Microgliaold | 0.674063677 | 1.887713711 | 2.39E-42 | 3.29E-05 |
| TMEM70 | Microgliaold | 0.652889622 | 0.953436592 | 2.69E-07 | 0.048873344 |
| PPP1CB | Microgliaold | 0.636826724 | 1.080687423 | 8.99E-09 | 0.024164143 |
| ARL6IP1 | Microgliaold | 0.609717014 | 0.886855453 | 2.61E-05 | 0.043468931 |
| HLA-A | Microgliaold | 0.590620915 | 1.300276653 | 1.00E-33 | 1.30E-05 |
| LDHA | Microgliaold | 0.582522469 | 2.07939763 | 4.23E-71 | 8.18E-10 |
| TNFRSF1B | Microgliaold | 0.579964424 | 1.337110465 | 7.65E-12 | 0.001419857 |
| ANKRD9 | Microgliaold | 0.57853949 | 1.371519877 | 5.01E-14 | 0.029539969 |
| EMD | Microgliaold | 0.577938106 | 1.334768286 | 2.85E-09 | 0.03103467 |
| SOD2 | Microgliaold | 0.576688825 | 1.958594016 | 1.16E-49 | 1.08E-12 |
| AC097534.2 | Microgliaold | 0.565503354 | 1.574391578 | 2.58E-06 | 0.027368027 |
| AC120193.1 | Microgliaold | 0.562034065 | 1.450348031 | 0.000334716 | 0.012907747 |
| SRGN | Microgliaold | 0.556818657 | 1.894624552 | 2.82E-56 | 1.62E-14 |
| FAM110B | Microgliaold | 0.546850283 | 1.377459343 | 0.000297343 | 0.03117155 |
| CAST | Microgliaold | 0.54369403 | 1.286502631 | 9.82E-08 | 0.003443806 |
| SLC11A1 | Microgliaold | 0.527358795 | 2.45269472 | 5.16E-63 | 2.70E-13 |
| CDKN1C | Microgliaold | 0.510081601 | 2.266649212 | 1.38E-22 | 0.000240479 |
| LIPN | Microgliaold | 0.493945131 | 1.135667059 | 2.31E-25 | 0.046630047 |
| TREM1 | Microgliaold | 0.491719845 | 1.523179654 | 5.34E-17 | 0.001587023 |
| MRPL44 | Microgliaold | 0.491645925 | 1.525377627 | 3.35E-13 | 0.018581429 |
| CD9 | Microgliaold | 0.489714747 | 2.101850801 | 7.93E-59 | 6.48E-08 |
| RAB13 | Microgliaold | 0.489531369 | 1.18143297 | 1.72E-07 | 0.032189431 |
| PGAM1 | Microgliaold | 0.481128276 | 1.071120772 | 1.60E-09 | 0.04116112 |
| CCL20 | Microgliaold | 0.476270455 | 0.981788022 | 3.89E-48 | 1.05E-05 |
| DHRS3 | Microgliaold | 0.475106585 | 1.291778443 | 2.41E-14 | 0.011166104 |
| C4orf3 | Microgliaold | 0.453951916 | 2.018949202 | 3.55E-52 | 3.08E-09 |
| IL18R1 | Microgliaold | 0.44935507 | 2.48510853 | 1.48E-13 | 0.001739326 |
| LY96 | Microgliaold | 0.431619604 | 1.139767328 | 3.51E-09 | 0.00309169 |
| NRIP3 | Microgliaold | 0.409996394 | 1.646416518 | 4.47E-07 | 0.002710829 |
| HAMP | Microgliaold | 0.407664937 | 1.497261385 | 6.02E-13 | 1.30E-05 |
| CXCL16 | Microgliaold | 0.392686675 | 1.447751177 | 1.38E-20 | 3.93E-05 |
| AQP9 | Microgliaold | 0.38053571 | 1.284558873 | 6.21E-25 | 0.00728659 |
| G0S2 | Microgliaold | 0.373574809 | 0.440991059 | 1.44E-90 | 2.27E-05 |

| | | | | | |
|---|---|---|---|---|---|
| FTH1 | Microgliaold | 0.372452782 | 3.195272666 | 1.48E-138 | 1.14E-31 |
| CARD19 | Microgliaold | 0.371032666 | 1.090533498 | 0.000129926 | 0.047622049 |
| SLC2A5 | Microgliaold | 0.353028484 | 1.266097502 | 1.89E-15 | 0.000869193 |
| IFI30 | Microgliaold | 0.3469799 | 2.080874312 | 6.42E-57 | 6.97E-11 |
| SELENOM | Microgliaold | 0.340042608 | 1.085865256 | 1.29E-08 | 0.026639162 |
| ANKRD37 | Microgliaold | 0.331714463 | 1.730757925 | 1.07E-06 | 3.77E-05 |
| PLP2 | Microgliaold | 0.329704252 | 1.251439664 | 2.15E-13 | 0.017413723 |
| SLC7A5 | Microgliaold | 0.326094552 | 1.588219499 | 2.03E-23 | 0.000922593 |
| SEC62 | Microgliaold | 0.313221699 | 1.084149033 | 1.96E-09 | 0.004605482 |
| VAPA | Microgliaold | 0.304592014 | 1.078686819 | 2.72E-08 | 0.011721361 |
| BLVRB | Microgliaold | 0.303514406 | 1.329164753 | 0.000117848 | 0.002385406 |
| HAVCR2 | Microgliaold | 0.295432288 | 1.001084367 | 0.001226285 | 0.016970113 |
| GUK1 | Microgliaold | 0.294087276 | 1.182818986 | 1.10E-09 | 0.003761616 |
| VEGFA | Microgliaold | 0.286015892 | 1.892051746 | 2.35E-36 | 1.24E-05 |
| SNHG5 | Microgliaold | 0.285335728 | 1.366421147 | 9.83E-19 | 0.000354029 |
| DYNLL1 | Microgliaold | 0.281663306 | 0.950585399 | 2.60E-06 | 0.017303746 |
| CD300A | Microgliaold | 0.280501522 | 1.294237587 | 2.49E-13 | 0.004735478 |
| FTL | Microgliaold | 0.280486801 | 1.522744836 | 7.36E-40 | 8.93E-08 |
| PRR13 | Microgliaold | 0.277194271 | 0.993808993 | 1.42E-05 | 0.044835882 |
| VIM | Microgliaold | 0.276815561 | 1.986123243 | 6.17E-79 | 3.01E-14 |
| MGAT1 | Microgliaold | 0.276309435 | 1.230952886 | 1.10E-14 | 0.001286498 |
| HLA-DQA2 | Microgliaold | 0.272114675 | 4.621955941 | 9.49E-118 | 1.14E-31 |
| CD55 | Microgliaold | 0.268448453 | 1.013122355 | 5.11E-11 | 0.044108143 |
| RNF181 | Microgliaold | 0.255393599 | 1.39648138 | 6.24E-18 | 0.001788796 |
| SOCS3 | Microgliaold | 0.251635035 | 1.923292245 | 1.32E-25 | 1.86E-07 |
| DEGS1 | Microgliaold | 0.251330469 | 1.033582179 | 1.82E-06 | 0.039264041 |
| A1BG | Microgliaold | 0.250882465 | 1.569893183 | 8.43E-18 | 0.001768167 |
| LAPTM4A | Microgliaold | 0.248454664 | 1.234550507 | 1.62E-19 | 0.002774256 |
| SLC3A2 | Microgliaold | 0.238486584 | 2.002193331 | 4.76E-50 | 1.78E-08 |
| ABL2 | Microgliaold | 0.231796937 | 0.876571648 | 0.001307072 | 0.047451454 |
| CD99 | Microgliaold | 0.227643365 | 1.205047698 | 2.39E-12 | 0.001852953 |
| NUDT1 | Microgliaold | 0.221671846 | 1.623444141 | 8.46E-22 | 0.000263617 |
| GSTO1 | Microgliaold | 0.217048985 | 1.406152088 | 5.41E-30 | 0.000299592 |
| ENPP4 | Microgliaold | 0.210590212 | 1.756754817 | 5.34E-15 | 0.001768703 |
| ATP6V1G1 | Microgliaold | 0.209285148 | 0.975238341 | 9.43E-06 | 0.012907747 |
| BZW1 | Microgliaold | 0.207021445 | 0.914519083 | 9.18E-05 | 0.046052433 |
| CARD16 | Microgliaold | 0.198868112 | 1.128336615 | 1.99E-07 | 0.024694588 |
| UBE2B | Microgliaold | 0.194811776 | 1.096645374 | 6.40E-07 | 0.005210116 |
| SMIM25 | Microgliaold | 0.172519982 | 1.143879387 | 9.67E-09 | 0.025773594 |
| SELENOS | Microgliaold | 0.171730564 | 1.067472096 | 2.48E-05 | 0.025773594 |
| METRNL | Microgliaold | 0.1640857 | 1.908423045 | 1.06E-41 | 8.94E-07 |
| GNA15 | Microgliaold | 0.156773406 | 1.198909135 | 0.000595599 | 0.047451454 |
| ALAS1 | Microgliaold | 0.144463886 | 1.302494402 | 1.32E-05 | 0.020140794 |
| LAPTM5 | Microgliaold | 0.136036784 | 1.310951773 | 7.62E-22 | 1.05E-05 |
| TCHH | Microgliayoung | -0.100492364 | -2.189689183 | 3.75E-10 | 0.000922593 |
| IFI16 | Microgliayoung | -0.164006103 | -1.024920707 | 5.54E-37 | 0.030579615 |
| SFMBT2 | Microgliayoung | -0.179441421 | -1.046896225 | 3.68E-31 | 0.01630057 |
| JDP2 | Microgliayoung | -0.19130776 | -1.508188123 | 5.74E-28 | 0.001419857 |
| CDK6 | Microgliayoung | -0.227121163 | -1.989322854 | 1.15E-58 | 4.68E-05 |
| CCNH | Microgliayoung | -0.249407355 | -1.093391677 | 1.94E-21 | 0.012849774 |
| DDX18 | Microgliayoung | -0.257884577 | -1.299812775 | 1.73E-14 | 0.021437534 |
| REV3L | Microgliayoung | -0.261419757 | -1.107050732 | 8.73E-14 | 0.02057463 |
| NSUN6 | Microgliayoung | -0.280448145 | -1.678336238 | 2.05E-17 | 0.000372461 |
| MIR646HG | Microgliayoung | -0.287297259 | -1.165315073 | 2.43E-10 | 0.010546905 |

| | | | | | |
|---|---|---|---|---|---|
| LINC02712 | Microgliayoung | -0.305552529 | -1.856162573 | 2.56E-12 | 0.009283631 |
| SHTN1 | Microgliayoung | -0.360484666 | -1.169972699 | 2.48E-29 | 0.032189431 |
| TUBA1B | Microgliayoung | -0.367618869 | -1.414888441 | 3.14E-37 | 8.85E-06 |
| LDLRAD4 | Microgliayoung | -0.371906191 | -1.346239213 | 3.60E-45 | 2.50E-05 |
| SAMD9L | Microgliayoung | -0.378253469 | -1.684960766 | 9.46E-28 | 0.011524795 |
| FPR3 | Microgliayoung | -0.378465862 | -1.425453551 | 2.65E-31 | 0.004735478 |
| FKBP5 | Microgliayoung | -0.378661057 | -1.104155655 | 1.13E-22 | 0.001860223 |
| SPTLC2 | Microgliayoung | -0.393334756 | -1.134450509 | 6.10E-18 | 0.033084913 |
| FCGR1A | Microgliayoung | -0.415429975 | -1.034672015 | 1.75E-28 | 0.033440149 |
| PPT1 | Microgliayoung | -0.418279409 | -0.988155674 | 4.40E-21 | 0.039870254 |
| CLEC12A | Microgliayoung | -0.447329773 | -1.406087961 | 2.07E-14 | 0.024694588 |
| STK38L | Microgliayoung | -0.453523388 | -1.47834901 | 3.31E-17 | 0.011223688 |
| MT-ND6 | Microgliayoung | -0.467100833 | -1.829227429 | 6.26E-71 | 0.000275554 |
| NLRP3 | Microgliayoung | -0.485048546 | -1.480645532 | 2.21E-15 | 0.008411947 |
| AL355881.1 | Microgliayoung | -0.494222284 | -2.105709517 | 3.15E-37 | 1.90E-06 |
| LPCAT2 | Microgliayoung | -0.509042943 | -1.64375465 | 4.09E-47 | 3.11E-06 |
| CYB5B | Microgliayoung | -0.510270806 | -1.721996494 | 2.09E-23 | 0.000275554 |
| XAF1 | Microgliayoung | -0.542181093 | -1.724200168 | 2.62E-34 | 0.022291597 |
| THADA | Microgliayoung | -0.5437375 | -1.460281537 | 1.40E-33 | 0.000806027 |
| AP001636.3 | Microgliayoung | -0.545854116 | -1.078541305 | 7.28E-12 | 0.048559188 |
| CCL4 | Microgliayoung | -0.578877376 | -1.477463086 | 2.71E-23 | 0.007149461 |
| CALM2 | Microgliayoung | -0.588318968 | -0.735082933 | 4.06E-33 | 0.042248619 |
| TEX14 | Microgliayoung | -0.592784574 | -1.232926602 | 3.38E-18 | 0.00026519 |
| SATB1-AS1 | Microgliayoung | -0.607121011 | -1.216451576 | 4.32E-26 | 0.016970113 |
| AC084871.1 | Microgliayoung | -0.633179973 | -1.355765458 | 6.91E-17 | 0.020063076 |
| AC106028.4 | Microgliayoung | -0.650015861 | -1.606148948 | 9.67E-09 | 0.01408013 |
| FCGR1B | Microgliayoung | -0.681645267 | -1.65241139 | 7.70E-26 | 0.011051442 |
| TCOF1 | Microgliayoung | -0.713982734 | -1.608583601 | 3.55E-14 | 0.009661605 |
| AC018754.1 | Microgliayoung | -0.719357734 | -1.251867572 | 1.93E-17 | 0.002417052 |
| CXorf21 | Microgliayoung | -0.71942808 | -1.668472363 | 2.18E-24 | 0.001101492 |
| CCL3 | Microgliayoung | -0.720292563 | -1.14156487 | 1.34E-20 | 0.008685281 |
| AADACL2-AS1 | Microgliayoung | -0.724632518 | -2.09358155 | 2.24E-12 | 0.01503229 |
| AC022217.3 | Microgliayoung | -0.774803278 | -1.324474867 | 2.50E-24 | 0.000322357 |
| PSME2 | Microgliayoung | -0.776692294 | -1.58570006 | 1.30E-29 | 0.003153682 |
| BTG2 | Microgliayoung | -0.791567436 | -1.26160057 | 3.27E-27 | 0.000564579 |
| KLHL6 | Microgliayoung | -0.80302097 | -1.304084931 | 1.47E-14 | 0.010490532 |
| FOSB | Microgliayoung | -0.815747062 | -1.680589819 | 3.85E-30 | 5.72E-07 |
| HLA-DRB5 | Microgliayoung | -0.827408292 | -3.664517878 | 2.34E-158 | 1.29E-31 |
| FOS | Microgliayoung | -0.832866259 | -0.924393779 | 9.55E-26 | 0.024768354 |
| CMSS1 | Microgliayoung | -0.862926594 | -1.48603997 | 1.07E-20 | 0.00227588 |
| AL138720.1 | Microgliayoung | -0.878859181 | -1.429747347 | 4.01E-08 | 0.012970581 |
| BLNK | Microgliayoung | -0.912104609 | -1.592052255 | 4.97E-39 | 0.002706774 |
| FLT1 | Microgliayoung | -0.919163887 | -1.820137729 | 8.63E-21 | 0.003397123 |
| TNFAIP8L3 | Microgliayoung | -0.935568407 | -2.171205584 | 1.50E-22 | 0.00316699 |
| CORO1A | Microgliayoung | -0.940159806 | -1.439335137 | 2.75E-22 | 0.002473549 |
| IER2 | Microgliayoung | -0.990856195 | -0.985171504 | 6.93E-19 | 0.005454448 |
| CH25H | Microgliayoung | -0.99733433 | -3.01471514 | 6.15E-33 | 3.20E-14 |
| RBKS | Microgliayoung | -1.016477987 | -1.238466732 | 1.13E-23 | 0.016229958 |
| FOXP2 | Microgliayoung | -1.02248433 | -1.564199057 | 2.88E-10 | 0.01075476 |
| IFI6 | Microgliayoung | -1.023809998 | -1.742151529 | 1.72E-29 | 0.035494151 |
| CCL4L2 | Microgliayoung | -1.039036316 | -2.491581156 | 1.08E-35 | 0.005846118 |
| AL031599.1 | Microgliayoung | -1.08513415 | -3.976844116 | 3.97E-59 | 1.02E-14 |
| AL499604.1 | Microgliayoung | -1.100082765 | -1.131479567 | 2.09E-18 | 0.034644371 |
| PALD1 | Microgliayoung | -1.131605106 | -2.235182993 | 7.48E-43 | 2.62E-07 |

| | | | | | |
|---|---|---|---|---|---|
| TTC33 | Microgliayoung | -1.166794702 | -1.173708605 | 6.02E-12 | 0.028262735 |
| NCK1-DT | Microgliayoung | -1.188487549 | -1.398834989 | 6.39E-17 | 0.019267864 |
| MYOM2 | Microgliayoung | -1.192866447 | -1.312099575 | 1.96E-05 | 0.037787766 |
| IFIT3 | Microgliayoung | -1.212824132 | -2.253546737 | 6.22E-21 | 0.000802091 |
| ILDR1 | Microgliayoung | -1.22526055 | -2.683597059 | 3.89E-32 | 5.26E-06 |
| PDE1C | Microgliayoung | -1.252561746 | -1.388343618 | 1.16E-28 | 0.025749699 |
| AL691403.1 | Microgliayoung | -1.290887981 | -1.895225382 | 1.92E-15 | 7.98E-06 |
| AF111167.1 | Microgliayoung | -1.298242733 | -1.628013967 | 5.93E-09 | 0.002561356 |
| AC022868.2 | Microgliayoung | -1.312425123 | -1.81041481 | 5.85E-23 | 0.003421155 |
| CCDC26 | Microgliayoung | -1.359490374 | -3.443613585 | 2.02E-72 | 3.48E-16 |
| SSPN | Microgliayoung | -1.395729535 | -1.774097393 | 7.69E-13 | 0.000117607 |
| C20orf27 | Microgliayoung | -1.410021275 | -1.642387668 | 3.01E-19 | 0.006925981 |
| LINC01480 | Microgliayoung | -1.448772979 | -1.686571363 | 1.31E-16 | 0.003239676 |
| AC022217.2 | Microgliayoung | -1.449467014 | -1.42483231 | 1.11E-10 | 0.04947952 |
| ENSA | Microgliayoung | -1.466258389 | -1.384351134 | 1.27E-31 | 0.001317596 |
| MIR4713HG | Microgliayoung | -1.491914514 | -2.055355061 | 6.75E-29 | 0.006719921 |
| IPCEF1 | Microgliayoung | -1.496663397 | -4.068750825 | 2.71E-82 | 1.38E-20 |
| HDAC9 | Microgliayoung | -1.512654987 | -1.74953918 | 2.55E-41 | 0.000148879 |
| ISG15 | Microgliayoung | -1.518908868 | -1.761293378 | 5.09E-18 | 0.016587149 |
| MIR3945HG | Microgliayoung | -1.552150767 | -1.484823218 | 2.25E-11 | 0.048619853 |
| GRID2 | Microgliayoung | -1.566475791 | -1.758000179 | 1.11E-12 | 0.023250679 |
| CX3CR1 | Microgliayoung | -1.605210003 | -3.621015873 | 9.57E-74 | 3.75E-17 |
| AC016745.1 | Microgliayoung | -1.615706352 | -1.605648531 | 4.43E-06 | 0.020305344 |
| POPDC2 | Microgliayoung | -1.639442107 | -1.596336183 | 1.36E-21 | 0.032189431 |
| IGSF6 | Microgliayoung | -1.681017619 | -2.099034442 | 1.83E-57 | 7.03E-05 |
| AC055854.1 | Microgliayoung | -1.70782845 | -1.95838808 | 3.49E-13 | 0.017033399 |
| SUSD3 | Microgliayoung | -1.753001674 | -1.8385203 | 6.40E-17 | 0.002473549 |
| CCDC200 | Microgliayoung | -1.767352581 | -1.746049394 | 4.60E-22 | 0.006277383 |
| AC007991.3 | Microgliayoung | -1.794971399 | -2.06569842 | 1.66E-05 | 0.000806027 |
| AL591518.1 | Microgliayoung | -1.920021046 | -1.909943024 | 4.44E-29 | 0.001599524 |
| P2RY13 | Microgliayoung | -1.972415054 | -2.175412683 | 1.71E-47 | 9.71E-08 |
| KCNIP1 | Microgliayoung | -2.246120476 | -1.879552152 | 4.39E-13 | 0.004113084 |
| AC073352.2 | Microgliayoung | -2.338853377 | -2.29512067 | 3.80E-26 | 0.000499007 |
| PLD4 | Microgliayoung | -2.393407604 | -2.031585935 | 1.95E-23 | 7.36E-05 |
| NAV3 | Microgliayoung | -2.470775549 | -3.391923256 | 6.67E-29 | 2.59E-10 |
| P2RY12 | Microgliayoung | -2.472626082 | -1.915573607 | 2.47E-21 | 0.007955638 |
| AC012150.1 | Microgliayoung | -2.654470746 | -2.050145489 | 8.53E-13 | 0.004809033 |
| CLEC9A | Microgliayoung | -3.070969122 | -2.121426623 | 0.00109694 | 0.006277383 |
