## Supplementary material for "Interpretable Aging Signatures in Human Retinal Cell Types Revealed by Single-Cell RNA Sequencing and Sparse Logistic Regression": Table S6

Table S6: Differentially expressed genes identified through both single-cell differential expression and pseudobulk differential expression analyses across 2 retinal photoreceptor populations: rods, and cones.

| gene | CellType | Direction | sc_avg_log2FC | pb_avg_log2FC | sc_p_val_adj | pb_p_val_adj |
|---|---|---|---|---|---|---|
| GABRA2 | Cone | old | 3.676539795 | 1.792551625 | 2.60E-06 | 0.008321657 |
| TMEM176A | Cone | old | 3.043292518 | 2.275540451 | 1.15E-11 | 3.09E-06 |
| UCMA | Cone | old | 2.942754144 | 1.380692497 | 2.18E-16 | 0.003460792 |
| PRUNE2 | Cone | old | 2.913612921 | 1.872331956 | 7.38E-06 | 1.96E-06 |
| MNDA | Cone | old | 2.428960814 | 1.58379621 | 0.001274139 | 0.007571795 |
| SBSPON | Cone | old | 1.75723583 | 1.236091122 | 1.63E-05 | 0.032489757 |
| SLC35F1 | Cone | old | 1.733859068 | 1.926706283 | 7.39E-18 | 0.000354133 |
| MT2A | Cone | old | 1.586546496 | 1.884059231 | 0.037533028 | 9.38E-11 |
| HLA-A | Cone | old | 1.253856694 | 1.04164889 | 3.48E-06 | 0.01603475 |
| CST3 | Cone | old | 1.168503862 | 1.825785384 | 2.16E-11 | 1.45E-09 |
| CRYAB | Cone | old | 1.079286182 | 2.384200841 | 0.003408236 | 1.34E-09 |
| RAB17 | Cone | old | 1.073216415 | 1.52826305 | 0.003137095 | 0.004791148 |
| EFNA5 | Cone | old | 0.942512415 | 1.443186868 | 0.006483899 | 0.00157838 |
| RPS26 | Cone | old | 0.932914406 | 1.249383139 | 3.59E-07 | 3.06E-05 |
| TTR | Cone | old | 0.919718007 | 2.149636154 | 1.69E-25 | 2.19E-11 |
| SLC22A17 | Cone | old | 0.889419333 | 1.166808749 | 0.010372123 | 0.022212687 |
| MBP | Cone | old | 0.814542607 | 1.617435934 | 1.71E-12 | 3.29E-05 |
| ACTB | Cone | old | 0.769294128 | 1.403018261 | 0.000320126 | 1.61E-09 |
| LRFN5 | Cone | old | 0.744365103 | 1.484494926 | 1.63E-06 | 0.000158363 |
| CCNO | Cone | old | 0.69410614 | 1.337845229 | 4.76E-05 | 0.002554116 |
| MESP1 | Cone | old | 0.693806324 | 0.991257374 | 0.001034613 | 0.01603475 |
| H1FX | Cone | old | 0.671401957 | 1.270934378 | 6.28E-05 | 9.14E-05 |
| KLRD1 | Cone | old | 0.653065148 | 1.376746404 | 0.00058044 | 0.001372059 |
| AC116903.2 | Cone | old | 0.628899498 | 1.577849328 | 9.94E-08 | 0.000634599 |
| AL118516.1 | Cone | old | 0.609602625 | 1.126604254 | 0.02143879 | 0.006599198 |
| ACTG1 | Cone | old | 0.510906442 | 0.929828065 | 0.005710744 | 0.000322482 |
| SDHAF3 | Cone | old | 0.504963119 | 0.777735474 | 0.015501711 | 0.042127279 |
| BCO2 | Cone | old | 0.46128034 | 1.067640838 | 8.52E-05 | 0.000326707 |
| LRRFIP1 | Cone | old | 0.428616052 | 1.215484723 | 3.16E-05 | 0.019534448 |
| HPRT1 | Cone | old | 0.395992212 | 0.759783272 | 0.000298517 | 0.040338222 |
| HNRNPH1 | Cone | old | 0.383460181 | 0.981892732 | 0.000194698 | 0.001370209 |
| HMGB2 | Cone | old | 0.303334909 | 1.051108496 | 1.73E-05 | 0.004914393 |
| BAG3 | Cone | old | 0.213538577 | 1.301655804 | 3.28E-05 | 0.000613263 |
| GLUL | Cone | old | 0.199491143 | 1.501237185 | 0.028848106 | 0.000127161 |
| TF | Cone | old | 0.146326066 | 2.395039546 | 0.00080396 | 1.36E-10 |
| MAGI2 | Cone | young | -0.124385975 | -0.856129039 | 2.64E-08 | 0.002581852 |
| GAS7 | Cone | young | -0.128442783 | -1.518181563 | 4.48E-09 | 1.04E-06 |
| AGBL4 | Cone | young | -0.130427951 | -0.803398006 | 0.000214014 | 0.009530598 |
| SNAP91 | Cone | young | -0.130975285 | -0.76197286 | 2.97E-05 | 0.019887787 |
| DLG2 | Cone | young | -0.162770165 | -0.927073378 | 0.000278694 | 0.000385886 |
| NCL | Cone | young | -0.188547865 | -0.96768946 | 5.06E-06 | 0.019776641 |
| KCNB1 | Cone | young | -0.192689623 | -0.713358782 | 0.013576852 | 0.023446181 |
| SAMD7 | Cone | young | -0.193485635 | -0.988297618 | 0.000490363 | 0.015242338 |
| PCAT1 | Cone | young | -0.219344756 | -1.043043372 | 9.08E-05 | 0.003873784 |
| KIAA0825 | Cone | young | -0.231729028 | -1.333379721 | 2.84E-11 | 8.94E-07 |
| KIF2A | Cone | young | -0.240243697 | -1.070014701 | 4.02E-09 | 5.89E-05 |
| PRKDC | Cone | young | -0.284468131 | -0.862492823 | 1.74E-05 | 0.023496564 |
| SGCD | Cone | young | -0.288151704 | -1.749273081 | 8.10E-11 | 1.72E-10 |
| SSX2IP | Cone | young | -0.305641946 | -1.132704664 | 3.18E-08 | 6.69E-06 |
| TMEM108 | Cone | young | -0.30733921 | -0.826134729 | 0.000922758 | 0.000297122 |
| MIR2052HG | Cone | young | -0.338773235 | -1.679506957 | 0.000727503 | 0.000125448 |
| MT-ND6 | Cone | young | -0.33998485 | -2.029172526 | 3.69E-20 | 1.34E-09 |
| LINC02343 | Cone | young | -0.355885207 | -1.685033795 | 9.63E-17 | 8.85E-08 |

| | | | | | | |
|---|---|---|---|---|---|---|
| RGS9 | Cone | young | -0.358474381 | -1.138318763 | 3.85E-09 | 4.16E-07 |
| MAP2 | Cone | young | -0.358907914 | -0.915164164 | 0.004507057 | 2.03E-05 |
| PDE6C | Cone | young | -0.37560067 | -0.954437137 | 0.001017435 | 0.011109345 |
| AC007349.2 | Cone | young | -0.381175161 | -1.513402472 | 5.10E-09 | 2.36E-06 |
| LINGO2 | Cone | young | -0.390341279 | -2.936283357 | 7.94E-22 | 4.93E-14 |
| WWOX | Cone | young | -0.433843859 | -1.803408187 | 1.32E-07 | 1.34E-09 |
| AC137770.1 | Cone | young | -0.474575423 | -1.280273085 | 0.001111096 | 0.000680209 |
| SLC4A8 | Cone | young | -0.493332047 | -1.160984478 | 0.012026024 | 0.010140934 |
| LINC00871 | Cone | young | -0.530066211 | -1.249767112 | 2.26E-07 | 0.000125448 |
| AC112206.2 | Cone | young | -0.550042746 | -1.31610121 | 1.62E-07 | 2.03E-08 |
| AC106798.1 | Cone | young | -0.792972357 | -1.228364764 | 0.001795032 | 0.000670367 |
| AL050403.2 | Cone | young | -0.94963311 | -1.653829665 | 0.000226719 | 1.74E-09 |
| ANTXR2 | Cone | young | -1.203441405 | -1.094749034 | 0.005237733 | 0.007128618 |
| RRAD | Cone | young | -1.817726889 | -0.294210719 | 4.17E-05 | 0.00115895 |
| BAG3 | Rod | old | 3.389610251 | 4.007505842 | 1.33E-40 | 3.28E-53 |
| RALYL | Rod | old | 3.335097142 | 1.873324113 | 6.57E-34 | 3.14E-06 |
| CCL2 | Rod | old | 3.139176751 | 2.003489679 | 0.019299926 | 2.24E-11 |
| WNT11 | Rod | old | 3.067925201 | 2.479107128 | 7.63E-06 | 8.94E-16 |
| FAM155A | Rod | old | 2.771477955 | 2.077177787 | 1.30E-36 | 3.68E-09 |
| CD44 | Rod | old | 2.726093873 | 1.507596681 | 1.73E-05 | 1.38E-09 |
| RPRM | Rod | old | 2.493891369 | 2.37983706 | 7.86E-09 | 3.45E-14 |
| PTPRM | Rod | old | 2.350493757 | 2.99257356 | 1.33E-27 | 5.58E-41 |
| P4HA2 | Rod | old | 2.298138298 | 2.370165465 | 2.74E-17 | 6.92E-21 |
| AC107419.1 | Rod | old | 2.239529257 | 1.726513444 | 3.09E-07 | 3.93E-08 |
| AC006994.2 | Rod | old | 2.137068108 | 1.675078403 | 0.021090973 | 6.69E-16 |
| MT3 | Rod | old | 1.982850563 | 1.134260699 | 1.26E-14 | 0.000461339 |
| LGALS3 | Rod | old | 1.894415939 | 1.502271747 | 4.23E-12 | 0.000682847 |
| AC097534.2 | Rod | old | 1.883484322 | 0.992720628 | 1.99E-06 | 2.37E-10 |
| CRYAB | Rod | old | 1.840019915 | 1.941938442 | 1.38E-48 | 1.34E-07 |
| CCN1 | Rod | old | 1.829912861 | 1.962181023 | 0.003589668 | 2.59E-07 |
| HSPA6 | Rod | old | 1.802335714 | 1.783803579 | 1.67E-57 | 4.77E-10 |
| P3H2 | Rod | old | 1.789604933 | 1.45315358 | 1.30E-11 | 3.38E-05 |
| MBP | Rod | old | 1.777376976 | 2.705389483 | 6.95E-24 | 3.34E-26 |
| TMEM176A | Rod | old | 1.7752528 | 1.434578592 | 5.50E-46 | 0.03460241 |
| BMPR1B | Rod | old | 1.73264605 | 1.256445496 | 5.70E-32 | 0.000942387 |
| PARP8 | Rod | old | 1.731846174 | 2.145675188 | 4.86E-05 | 1.56E-13 |
| NPVF | Rod | old | 1.726938855 | 1.128516706 | 2.27E-17 | 0.025115583 |
| HSPB1 | Rod | old | 1.713168254 | 3.169670245 | 1.40E-29 | 1.20E-26 |
| PTPRD | Rod | old | 1.624108079 | 0.831045857 | 2.94E-08 | 3.53E-09 |
| WNT5B | Rod | old | 1.618739844 | 1.811211102 | 0.000231895 | 8.55E-15 |
| MT2A | Rod | old | 1.532261586 | 0.887135802 | 6.81E-11 | 0.000147199 |
| AC007221.1 | Rod | old | 1.525718346 | 1.376221279 | 0.016933227 | 2.87E-12 |
| KITLG | Rod | old | 1.516239538 | 2.842415505 | 1.84E-13 | 2.95E-19 |
| TMEM176B | Rod | old | 1.494538829 | 1.339822279 | 1.48E-33 | 0.025023822 |
| SLC7A8 | Rod | old | 1.426127786 | 1.405072705 | 1.54E-06 | 8.19E-10 |
| GFAP | Rod | old | 1.413855263 | 1.50417403 | 0.015852719 | 6.70E-07 |
| SQOR | Rod | old | 1.34334132 | 1.39953197 | 0.002840846 | 1.75E-06 |
| CAP2 | Rod | old | 1.311201336 | 1.832431599 | 1.55E-13 | 7.79E-11 |
| EEF1A2 | Rod | old | 1.290926611 | 1.548976087 | 0.003831655 | 1.48E-07 |
| HECW1 | Rod | old | 1.277869209 | 1.079314193 | 0.001247344 | 0.000597185 |
| A2M | Rod | old | 1.228512499 | 1.236936174 | 4.57E-52 | 0.00145858 |
| FBXL14 | Rod | old | 1.21888859 | 2.063232637 | 9.44E-17 | 4.16E-24 |
| GPX3 | Rod | old | 1.199292247 | 1.233163333 | 1.35E-11 | 0.017678867 |
| AC012178.1 | Rod | old | 1.19400693 | 1.799100403 | 3.72E-11 | 8.38E-29 |

| | | | | | | |
|---|---|---|---|---|---|---|
| TRNP1 | Rod | old | 1.137568563 | 1.843049091 | 1.17E-54 | 3.74E-12 |
| PARD6G-AS1 | Rod | old | 1.12851608 | 2.2880064 | 4.30E-27 | 1.99E-34 |
| HSD17B7 | Rod | old | 1.114001144 | 1.539081333 | 2.03E-18 | 5.71E-20 |
| LTBP1 | Rod | old | 1.08908205 | 1.211321639 | 5.91E-18 | 4.69E-05 |
| PGK1 | Rod | old | 1.083296645 | 0.459463717 | 0.006744222 | 9.03E-05 |
| PARD3-AS1 | Rod | old | 1.048985912 | 1.50282237 | 0.015712755 | 9.03E-07 |
| NKRF | Rod | old | 1.020721338 | 1.586077822 | 6.80E-06 | 9.88E-12 |
| LRP2 | Rod | old | 0.999720499 | 1.47920341 | 3.38E-20 | 3.23E-32 |
| PDK3 | Rod | old | 0.986019748 | 1.196244097 | 0.000765408 | 6.28E-14 |
| SLC35F1 | Rod | old | 0.961946383 | 3.659943543 | 7.61E-73 | 7.62E-86 |
| AL356804.1 | Rod | old | 0.946631179 | 2.625588449 | 3.86E-42 | 6.01E-59 |
| APP | Rod | old | 0.939401389 | 1.933959875 | 3.64E-05 | 1.00E-11 |
| RAB39A | Rod | old | 0.937288321 | 1.808198001 | 1.27E-09 | 1.71E-14 |
| S100A10 | Rod | old | 0.923651208 | 1.286267712 | 2.35E-14 | 7.61E-12 |
| CERS6 | Rod | old | 0.853084883 | 1.606522878 | 0.009836882 | 2.41E-08 |
| PARD3 | Rod | old | 0.842742064 | 0.711751454 | 0.018502254 | 4.07E-08 |
| OLFM3 | Rod | old | 0.824411429 | 1.659066395 | 1.45E-17 | 2.28E-33 |
| ME3 | Rod | old | 0.800915403 | 1.589564991 | 1.80E-07 | 2.21E-13 |
| HSPA1A | Rod | old | 0.770741596 | 2.079600587 | 2.69E-170 | 4.07E-20 |
| C17orf67 | Rod | old | 0.762139313 | 1.28622981 | 0.000179257 | 1.23E-09 |
| HSPH1 | Rod | old | 0.758533015 | 1.577180059 | 4.97E-82 | 8.46E-10 |
| CYP39A1 | Rod | old | 0.755917536 | 1.382518699 | 0.000135942 | 1.24E-13 |
| AANAT | Rod | old | 0.7449532 | 1.692564082 | 5.42E-06 | 1.66E-11 |
| MTAP | Rod | old | 0.731087791 | 2.474817598 | 1.05E-07 | 1.67E-19 |
| TF | Rod | old | 0.727996054 | 1.020151729 | 3.16E-18 | 0.030996713 |
| ATF7IP2 | Rod | old | 0.708072117 | 1.059258696 | 0.000301774 | 7.44E-17 |
| TENT5A | Rod | old | 0.686230018 | 1.112449848 | 0.003490917 | 3.84E-11 |
| C3orf70 | Rod | old | 0.681721475 | 1.637818377 | 0.000483497 | 2.51E-11 |
| PRR16 | Rod | old | 0.669254339 | 2.952597198 | 1.19E-19 | 6.60E-34 |
| DYNC1I1 | Rod | old | 0.648975712 | 1.257516114 | 8.34E-15 | 1.80E-13 |
| FNDC3B | Rod | old | 0.647682375 | 0.841487326 | 0.018733213 | 1.52E-10 |
| EEF1A1 | Rod | old | 0.645767453 | 0.687986479 | 0.006279498 | 3.28E-08 |
| ID1 | Rod | old | 0.645335818 | 2.003649587 | 6.41E-07 | 1.05E-33 |
| FAM149A | Rod | old | 0.632295399 | 1.085768998 | 0.00888995 | 7.79E-16 |
| PDE6H | Rod | old | 0.628547073 | 0.669490749 | 5.55E-21 | 0.034692478 |
| ABHD3 | Rod | old | 0.624133604 | 1.32212392 | 7.12E-15 | 2.92E-08 |
| ID3 | Rod | old | 0.60946562 | 1.553404622 | 0.001705284 | 1.49E-08 |
| SNAP23 | Rod | old | 0.599019067 | 1.483892195 | 1.23E-06 | 4.24E-29 |
| RBP7 | Rod | old | 0.591467684 | 1.512355344 | 2.95E-12 | 6.29E-14 |
| HIBCH | Rod | old | 0.587594112 | 0.814275972 | 0.000939587 | 4.65E-13 |
| AL022068.1 | Rod | old | 0.583959536 | 1.276281542 | 3.85E-11 | 2.52E-12 |
| DIP2B | Rod | old | 0.581753526 | 1.096617648 | 3.91E-08 | 1.88E-15 |
| RPS26 | Rod | old | 0.557384892 | 1.393222177 | 8.07E-23 | 1.23E-23 |
| HSPA1B | Rod | old | 0.556284825 | 1.714530243 | 1.42E-132 | 5.05E-15 |
| AC068774.1 | Rod | old | 0.551024707 | 1.234438609 | 0.002629476 | 4.82E-09 |
| LINC02177 | Rod | old | 0.549728966 | 1.721781375 | 2.58E-07 | 8.03E-15 |
| DENND2C | Rod | old | 0.540865762 | 1.567970382 | 4.62E-07 | 1.58E-18 |
| C4orf19 | Rod | old | 0.538179032 | 1.394233425 | 0.018473278 | 6.99E-18 |
| NXN | Rod | old | 0.53622205 | 1.679230751 | 3.18E-08 | 2.42E-32 |
| GNAS | Rod | old | 0.507460846 | 0.831198147 | 5.81E-05 | 2.44E-09 |
| AP3B1 | Rod | old | 0.474789512 | 0.950415931 | 7.90E-05 | 9.63E-14 |
| SERPINF1 | Rod | old | 0.473967341 | 2.730024692 | 3.73E-20 | 2.07E-45 |
| EPB41L3 | Rod | old | 0.466532104 | 1.711931758 | 5.42E-35 | 1.63E-30 |
| OSBPL6 | Rod | old | 0.460606497 | 1.072057833 | 0.011952847 | 2.09E-10 |

| | | | | | | |
|---|---|---|---|---|---|---|
| HNRNPH1 | Rod | old | 0.446355975 | 1.051913694 | 6.72E-21 | 4.88E-13 |
| LIMS1 | Rod | old | 0.421738795 | 0.915451756 | 2.73E-08 | 1.19E-11 |
| GAPDH | Rod | old | 0.412612866 | 0.901215431 | 3.53E-09 | 3.12E-05 |
| PLCL2 | Rod | old | 0.389245036 | 2.975961143 | 8.49E-29 | 9.42E-41 |
| FAM138D | Rod | old | 0.386376136 | 2.987069035 | 8.30E-16 | 6.42E-33 |
| PPP4R1 | Rod | old | 0.361246394 | 1.064636758 | 0.001632586 | 8.87E-13 |
| DNAJA4 | Rod | old | 0.359333342 | 1.674798214 | 5.50E-17 | 8.12E-18 |
| PNRC1 | Rod | old | 0.357052471 | 0.70542246 | 0.000448444 | 2.09E-07 |
| EVI5 | Rod | old | 0.347573988 | 0.610897479 | 0.019926468 | 6.91E-06 |
| NELL2 | Rod | old | 0.332346935 | 0.836337883 | 1.20E-06 | 3.27E-14 |
| RPL4 | Rod | old | 0.33031853 | 0.778859847 | 6.22E-07 | 7.22E-06 |
| WBP2 | Rod | old | 0.32967124 | 1.027080715 | 3.50E-11 | 2.13E-11 |
| AP1S2 | Rod | old | 0.324128485 | 0.807130633 | 2.17E-12 | 5.49E-09 |
| CHORDC1 | Rod | old | 0.304674831 | 1.537336967 | 8.90E-27 | 1.79E-26 |
| ANKRD37 | Rod | old | 0.304256234 | 0.722044508 | 0.001374881 | 9.22E-07 |
| AC018362.1 | Rod | old | 0.290885851 | 1.642689744 | 1.21E-06 | 4.40E-27 |
| CACYBP | Rod | old | 0.286200342 | 2.038814444 | 1.96E-52 | 1.24E-46 |
| RSPO2 | Rod | old | 0.271484554 | 1.250656382 | 0.000193562 | 3.21E-06 |
| ZNF532 | Rod | old | 0.253786491 | 1.807604772 | 5.20E-25 | 6.82E-20 |
| RPL39 | Rod | old | 0.235670986 | 0.654923653 | 0.00180833 | 7.63E-06 |
| AHSA1 | Rod | old | 0.232472379 | 1.718196816 | 5.70E-33 | 6.03E-26 |
| PLEKHB1 | Rod | old | 0.222291858 | 0.753786974 | 0.000419836 | 9.49E-08 |
| PGAM1 | Rod | old | 0.213570127 | 0.694842863 | 5.49E-09 | 1.75E-06 |
| ZNF682 | Rod | old | 0.200519218 | 1.343897889 | 5.55E-08 | 2.26E-14 |
| DNAJB4 | Rod | old | 0.198868131 | 1.702341438 | 8.17E-25 | 8.93E-35 |
| PCLO | Rod | old | 0.19804496 | 0.957247611 | 2.22E-05 | 6.54E-21 |
| HSPD1 | Rod | old | 0.190929202 | 1.729372478 | 2.19E-38 | 5.26E-36 |
| RARB | Rod | old | 0.190083841 | 1.170784514 | 0.004059927 | 9.27E-11 |
| HEBP2 | Rod | old | 0.186175239 | 1.189399451 | 0.003066325 | 9.88E-12 |
| WDR61 | Rod | old | 0.184482668 | 1.295105186 | 3.37E-07 | 7.63E-21 |
| KLHL41 | Rod | old | 0.142935618 | 1.511127642 | 7.04E-05 | 1.68E-15 |
| METRNL | Rod | old | 0.138577941 | 3.123529536 | 7.33E-69 | 1.88E-39 |
| MESP2 | Rod | old | 0.137093313 | 0.894307735 | 0.003453302 | 1.20E-05 |
| HPRT1 | Rod | old | 0.133516044 | 1.242508616 | 1.16E-33 | 2.56E-26 |
| RPL22 | Rod | old | 0.128191516 | 0.642281174 | 4.72E-05 | 3.92E-09 |
| BCO2 | Rod | old | 0.113805174 | 1.485311786 | 1.84E-22 | 1.67E-46 |
| SNHG29 | Rod | old | 0.105729033 | 0.791917087 | 4.12E-06 | 5.49E-09 |
| AL049552.1 | Rod | young | -0.100879505 | -0.792906381 | 3.09E-05 | 0.001294523 |
| PNN | Rod | young | -0.101253121 | -0.376547528 | 3.86E-07 | 0.032398166 |
| TRIM73 | Rod | young | -0.107893694 | -0.894266126 | 0.002159482 | 0.000361361 |
| MT-ND5 | Rod | young | -0.111710766 | -0.650072093 | 1.99E-45 | 0.009157871 |
| ANKRD18A | Rod | young | -0.112161532 | -0.829423176 | 0.006424986 | 2.75E-06 |
| RGS9 | Rod | young | -0.112943699 | -0.595382032 | 2.00E-29 | 1.42E-07 |
| CCDC110 | Rod | young | -0.11303828 | -1.01876678 | 0.000745357 | 1.68E-05 |
| HMGN1 | Rod | young | -0.113545274 | -0.387661145 | 8.82E-32 | 0.003601292 |
| PGM5 | Rod | young | -0.113992388 | -0.995028551 | 0.003186509 | 0.000441299 |
| TMEM108 | Rod | young | -0.115341903 | -0.639355879 | 2.35E-23 | 1.57E-06 |
| POLR2A | Rod | young | -0.119191167 | -0.538469002 | 1.12E-09 | 3.50E-05 |
| RHOA | Rod | young | -0.120586682 | -0.344133091 | 0.000133026 | 0.014684754 |
| ABLIM3 | Rod | young | -0.121830553 | -0.679011685 | 2.47E-14 | 1.12E-08 |
| MPP4 | Rod | young | -0.122400866 | -0.480347982 | 9.56E-09 | 0.000649408 |
| SNX19 | Rod | young | -0.133860107 | -0.774624505 | 0.007003586 | 8.49E-06 |
| ANP32A | Rod | young | -0.138485176 | -0.332541876 | 0.048237322 | 0.009623399 |
| SUCO | Rod | young | -0.139833962 | -0.277262103 | 7.64E-05 | 0.039181833 |

| | | | | | | |
|---|---|---|---|---|---|---|
| IGSF9 | Rod | young | -0.139912952 | -0.477100917 | 2.41E-06 | 0.001777215 |
| AC084361.1 | Rod | young | -0.140564069 | -0.929659695 | 4.62E-12 | 0.001122788 |
| PCMTD2 | Rod | young | -0.141051501 | -0.292804619 | 0.004484102 | 0.014642071 |
| USP5 | Rod | young | -0.142155272 | -0.881798223 | 0.002481991 | 0.001203453 |
| POPDC2 | Rod | young | -0.142747992 | -1.300972414 | 0.005016889 | 6.56E-06 |
| NUDCD1 | Rod | young | -0.143234737 | -0.823227525 | 0.000248272 | 3.65E-05 |
| CFAP46 | Rod | young | -0.143594534 | -0.82747867 | 4.73E-05 | 1.77E-05 |
| AC109129.1 | Rod | young | -0.14494279 | -1.068495601 | 0.034527247 | 0.00141469 |
| AK9 | Rod | young | -0.147303541 | -0.730992504 | 0.000269496 | 9.73E-06 |
| CABYR | Rod | young | -0.147877381 | -0.613537279 | 0.000367448 | 0.027778158 |
| RUNX2 | Rod | young | -0.149645061 | -0.661467894 | 0.002086035 | 0.004031442 |
| AL592183.1 | Rod | young | -0.154292171 | -1.366448008 | 3.27E-07 | 8.45E-13 |
| TRIM44 | Rod | young | -0.157132131 | -0.370523991 | 0.001356118 | 0.015074315 |
| HIST1H2BC | Rod | young | -0.160469945 | -0.552058713 | 0.002967889 | 0.005163218 |
| MYEF2 | Rod | young | -0.162304542 | -0.586917742 | 3.28E-05 | 2.07E-05 |
| ABCA1 | Rod | young | -0.170138252 | -1.201049903 | 0.001994996 | 1.73E-06 |
| DLG4 | Rod | young | -0.171179773 | -0.318931145 | 0.046426149 | 0.032053903 |
| OSBP2 | Rod | young | -0.174842823 | -0.285812342 | 0.00019762 | 0.031070378 |
| MT-ND6 | Rod | young | -0.184934462 | -2.405837607 | 9.70E-85 | 3.22E-56 |
| AC012613.1 | Rod | young | -0.186703486 | -0.621848425 | 0.011438203 | 0.000212484 |
| ANAPC7 | Rod | young | -0.191852718 | -0.750204699 | 0.0006267 | 0.000450532 |
| HNRNPA3 | Rod | young | -0.193295723 | -0.475561262 | 1.73E-13 | 0.0002277 |
| GOLGB1 | Rod | young | -0.195465121 | -0.389017579 | 0.000239814 | 0.000726812 |
| PUS7L | Rod | young | -0.198389177 | -0.621419008 | 0.000747241 | 8.22E-06 |
| ATP6V0D2 | Rod | young | -0.199186769 | -0.536327079 | 8.23E-06 | 0.001012197 |
| AP000462.3 | Rod | young | -0.209038403 | -0.94440608 | 5.64E-09 | 5.84E-07 |
| ACSL6 | Rod | young | -0.211551755 | -0.598097819 | 0.000115317 | 1.77E-05 |
| KLHDC4 | Rod | young | -0.216530808 | -0.871723115 | 0.012113609 | 0.000174935 |
| EMC1 | Rod | young | -0.219933396 | -0.70198425 | 3.39E-06 | 6.21E-06 |
| KCNB1 | Rod | young | -0.224521592 | -0.555663953 | 3.84E-22 | 0.000170125 |
| TRMT13 | Rod | young | -0.226535785 | -0.555897245 | 0.000168983 | 0.00207958 |
| CNKSR2 | Rod | young | -0.234397803 | -0.695281877 | 1.41E-14 | 3.03E-06 |
| SMC1A | Rod | young | -0.238830226 | -0.592485874 | 9.45E-05 | 0.00021354 |
| PRKDC | Rod | young | -0.243052167 | -0.527408685 | 0.000728755 | 2.33E-05 |
| THUMPD3 | Rod | young | -0.249815974 | -0.520065391 | 0.000948622 | 0.002606058 |
| IMPG1 | Rod | young | -0.252091903 | -0.508768132 | 1.98E-15 | 0.000287574 |
| HNRNPD | Rod | young | -0.252287998 | -0.398089187 | 1.21E-10 | 0.0021729 |
| CCDC171 | Rod | young | -0.257051845 | -0.585042838 | 6.47E-06 | 9.17E-05 |
| UNC13C | Rod | young | -0.257203805 | -0.364752725 | 0.000371802 | 0.004851373 |
| SARS | Rod | young | -0.259035935 | -0.452500038 | 2.43E-05 | 0.010311467 |
| PPM1N | Rod | young | -0.260226026 | -0.50034476 | 0.006156463 | 0.009188506 |
| AC044810.2 | Rod | young | -0.266511107 | -0.415086778 | 7.85E-07 | 0.013254764 |
| AC099788.1 | Rod | young | -0.267118862 | -0.996017198 | 9.76E-05 | 3.54E-05 |
| EIF4G2 | Rod | young | -0.268161458 | -0.39677807 | 8.19E-06 | 0.001792206 |
| CC2D2B | Rod | young | -0.273175649 | -1.140269619 | 6.37E-05 | 2.75E-06 |
| ENO4 | Rod | young | -0.285212872 | -0.494361839 | 0.011211334 | 0.02584264 |
| AC096589.1 | Rod | young | -0.286272612 | -0.801434803 | 0.002343698 | 5.68E-05 |
| LARS | Rod | young | -0.298060659 | -0.459093411 | 2.33E-07 | 0.002615016 |
| MPV17 | Rod | young | -0.304113475 | -0.708796463 | 4.95E-06 | 0.002319892 |
| NDUFAF2 | Rod | young | -0.307647263 | -0.592170717 | 0.000193555 | 0.000265416 |
| GDAP1 | Rod | young | -0.309748061 | -0.452940232 | 2.18E-07 | 0.001019004 |
| LINC00871 | Rod | young | -0.318818081 | -0.57212131 | 4.79E-08 | 0.001377213 |
| NFKBIZ | Rod | young | -0.320859105 | -0.530758154 | 0.002954457 | 0.024776298 |
| SMC6 | Rod | young | -0.325245128 | -0.547325245 | 0.017242613 | 0.007865884 |

| | | | | | | |
|---|---|---|---|---|---|---|
| AP000462.2 | Rod | young | -0.329343065 | -1.278077614 | 1.01E-07 | 8.52E-06 |
| AC007952.4 | Rod | young | -0.329382718 | -1.243638664 | 0.000187878 | 4.95E-05 |
| NCBP1 | Rod | young | -0.33442249 | -0.434235043 | 0.042021878 | 0.022610545 |
| RORB | Rod | young | -0.334847651 | -0.474747975 | 0.019589134 | 0.000266784 |
| SRRM2 | Rod | young | -0.337743385 | -0.447267046 | 8.36E-08 | 0.003897877 |
| BCL2L13 | Rod | young | -0.33805339 | -0.409042515 | 0.000792552 | 0.002013563 |
| CDH12 | Rod | young | -0.339172579 | -0.354417563 | 2.34E-06 | 0.005161722 |
| RD3 | Rod | young | -0.341308098 | -0.538079095 | 2.77E-18 | 0.000147284 |
| KIF2A | Rod | young | -0.356100558 | -0.35911157 | 0.005557379 | 0.013026763 |
| AC130456.2 | Rod | young | -0.357738343 | -2.85610469 | 6.84E-16 | 4.45E-11 |
| RS1 | Rod | young | -0.36532128 | -0.315954055 | 1.73E-06 | 0.038615286 |
| XKR4 | Rod | young | -0.367553832 | -2.731215984 | 5.16E-08 | 2.92E-38 |
| COG2 | Rod | young | -0.368272856 | -0.419111314 | 0.032695262 | 0.044720157 |
| NORAD | Rod | young | -0.370195825 | -0.707642252 | 1.14E-12 | 5.65E-06 |
| AC010601.1 | Rod | young | -0.372378875 | -0.827947601 | 0.000546345 | 0.005952066 |
| ELSPBP1 | Rod | young | -0.383020513 | -0.856144005 | 1.57E-09 | 4.29E-08 |
| SLC1A7 | Rod | young | -0.391268088 | -0.382046054 | 0.016501469 | 0.004271593 |
| RBM3 | Rod | young | -0.398919119 | -0.411988262 | 0.000285225 | 0.028916956 |
| SLC25A24 | Rod | young | -0.399266398 | -0.811163097 | 3.44E-13 | 4.85E-09 |
| FAM53C | Rod | young | -0.409996447 | -0.910307037 | 0.00443038 | 9.54E-05 |
| SRFBP1 | Rod | young | -0.410829698 | -0.605115992 | 1.98E-06 | 0.000504228 |
| FRMPD1 | Rod | young | -0.415960086 | -0.368863626 | 1.72E-05 | 0.013137281 |
| AL050403.2 | Rod | young | -0.428880182 | -0.956539512 | 3.47E-10 | 6.00E-13 |
| AL157944.1 | Rod | young | -0.43537608 | -0.521205086 | 2.88E-06 | 8.37E-05 |
| MT-CO2 | Rod | young | -0.437188223 | -0.501319088 | 3.41E-42 | 0.001576358 |
| OSGEP | Rod | young | -0.446429643 | -0.441336056 | 0.041850487 | 0.022523131 |
| AL160408.3 | Rod | young | -0.447727384 | -0.831740046 | 0.038434569 | 0.003071408 |
| EMC1-AS1 | Rod | young | -0.456924907 | -0.588702528 | 0.005984491 | 0.001185642 |
| PRPF6 | Rod | young | -0.467332547 | -0.517454239 | 0.001704742 | 0.000707581 |
| AC055733.2 | Rod | young | -0.482471097 | -0.378113174 | 0.000174512 | 0.015155238 |
| HS3ST3B1 | Rod | young | -0.486676746 | -0.479429073 | 0.031770086 | 0.005557687 |
| SPARC | Rod | young | -0.487144969 | -0.935950973 | 0.004023472 | 3.18E-06 |
| AC087457.1 | Rod | young | -0.502165577 | -0.827521832 | 6.17E-06 | 2.73E-05 |
| NAGK | Rod | young | -0.503735745 | -0.492340197 | 0.028005617 | 0.020582804 |
| AL591242.1 | Rod | young | -0.505335843 | -0.802907112 | 0.029892957 | 0.01108796 |
| HNRNPA2B1 | Rod | young | -0.509626308 | -0.338733569 | 3.23E-16 | 0.006898461 |
| AL513327.1 | Rod | young | -0.539955703 | -0.48789702 | 0.00462447 | 0.020697187 |
| NCL | Rod | young | -0.55839191 | -0.818951708 | 8.01E-34 | 1.36E-09 |
| AC008040.1 | Rod | young | -0.562046249 | -1.011879707 | 0.004476015 | 2.44E-05 |
| DEK | Rod | young | -0.562361548 | -0.47782695 | 5.46E-05 | 0.001325871 |
| FAM161A | Rod | young | -0.566481871 | -0.452573358 | 4.07E-11 | 0.002471909 |
| SIVA1 | Rod | young | -0.571761263 | -0.400538201 | 0.046060588 | 0.045210656 |
| MT-ND2 | Rod | young | -0.590129209 | -0.688752124 | 2.16E-37 | 0.003530008 |
| HSPA14.1 | Rod | young | -0.601120107 | -0.918131559 | 0.013475217 | 0.000160987 |
| AC053513.1 | Rod | young | -0.602359553 | -0.550540647 | 2.71E-05 | 0.009650475 |
| DYNLL2 | Rod | young | -0.627517179 | -0.506720587 | 0.043703989 | 0.002574115 |
| AL392023.2 | Rod | young | -0.627679479 | -2.522532523 | 4.98E-28 | 1.97E-27 |
| SAMD7 | Rod | young | -0.684499129 | -0.526908114 | 4.26E-09 | 6.53E-05 |
| HNRNPDL | Rod | young | -0.693585642 | -0.516518706 | 1.12E-10 | 0.000197621 |
| AC008735.2 | Rod | young | -0.695517595 | -0.998246083 | 0.018928967 | 0.002482465 |
| LINC02275 | Rod | young | -0.710177355 | -0.676735741 | 2.11E-06 | 2.00E-05 |
| MIR2052HG | Rod | young | -0.714454124 | -1.925097323 | 3.07E-06 | 3.85E-12 |
| AL365295.1 | Rod | young | -0.745953313 | -1.502489849 | 0.000472664 | 5.31E-10 |
| TMEM108-AS1 | Rod | young | -0.794840924 | -0.987501226 | 0.015385632 | 0.005911937 |

| | | | | | | |
|---|---|---|---|---|---|---|
| ATP5MC1 | Rod | young | -0.816486884 | -0.424109571 | 0.003944932 | 0.008606083 |
| AL355835.1 | Rod | young | -0.843413836 | -1.98273846 | 7.64E-31 | 4.06E-18 |
| HNRNPAB | Rod | young | -0.875610878 | -0.534716331 | 0.000580119 | 0.004146963 |
| MRLN | Rod | young | -0.888095446 | -0.757261134 | 1.50E-11 | 3.70E-06 |
| LRRC4 | Rod | young | -0.895371085 | -1.069288046 | 3.83E-06 | 0.000125682 |
| BX664615.2 | Rod | young | -0.899602114 | -1.004349396 | 2.89E-05 | 1.41E-05 |
| IFI27 | Rod | young | -0.906047685 | -1.199231877 | 0.01235197 | 1.23E-08 |
| FAM138C | Rod | young | -0.907643395 | -1.646036047 | 0.000422757 | 0.000805781 |
| AC022217.3 | Rod | young | -0.920274615 | -0.897723759 | 0.002359602 | 9.36E-06 |
| PLA2G4C-AS1 | Rod | young | -0.935485485 | -1.005752672 | 2.12E-11 | 3.50E-09 |
| LINC01619 | Rod | young | -0.939432122 | -1.163176489 | 0.003298308 | 1.65E-05 |
| WWC2-AS1 | Rod | young | -1.482246349 | -0.849079591 | 8.96E-06 | 1.61E-06 |
| SPON1 | Rod | young | -1.495843687 | -1.242539938 | 0.047737759 | 1.33E-06 |
| TRIB3 | Rod | young | -1.588169477 | -1.203830932 | 0.002330762 | 4.23E-06 |
| AL445250.1 | Rod | young | -2.172665571 | -2.574682492 | 8.03E-12 | 6.10E-25 |
| LINC02516 | Rod | young | -2.254674846 | -2.27060966 | 1.31E-05 | 0.000724231 |
| AC093151.8 | Rod | young | -2.403338837 | -2.364563343 | 6.10E-10 | 0.000314805 |
