## Supplementary material for "Interpretable Aging Signatures in Human Retinal Cell Types Revealed by Single-Cell RNA Sequencing and Sparse Logistic Regression": Table S7

Table S7: Donor-level pseudobulk sensitivity analysis of age-associated differential expression (conservative model). This table reports per‑cell-type differential expression (DE) results from a donor-level pseudobulk model that aggregates counts from the left and right eyes of each donor and includes sex and diabetic status covariates. The design was ~ sex + DM_status + age_group, where DM_status contrasts nonDM vs the combined DM+DR groups and age_group contrasts Old vs Young. Donors serve as the biological replicate unit. Abbreviations: DE, differential expression; DM, diabetes mellitus (no diagnosed diabetic retinopathy); DR, diabetic retinopathy; nonDM, non-diabetic.

| gene | log2FoldChange | padj | Direction | CellType |
|---|---|---|---|---|
| INPP4B | 1.172020023 | 0.034004952 | old | AC |
| AGAP1 | 0.660225373 | 0.034004952 | old | AC |
| EFCAB2 | -1.185817706 | 0.01754828 | young | AC |
| KCNMB2 | -1.364205076 | 0.01754828 | young | AC |
| AC109466.1 | -1.762249503 | 0.000284641 | young | AC |
| ADM | 2.059262554 | 0.00697988 | old | Astrocyte |
| ANGPTL4 | 2.047255129 | 0.003757976 | old | Astrocyte |
| HILPDA | 2.037424362 | 0.003757976 | old | Astrocyte |
| GADD45A | 1.951280239 | 0.039207012 | old | Astrocyte |
| NNMT | 1.825286353 | 0.048690097 | old | Astrocyte |
| HSPA6 | 1.804178349 | 0.048377913 | old | Astrocyte |
| SELENOM | 1.485100679 | 0.049185508 | old | Astrocyte |
| NEAT1 | 1.263122911 | 0.043149152 | old | Astrocyte |
| YBX3 | 1.116769892 | 0.049185508 | old | Astrocyte |
| PGK1 | 1.011323431 | 0.028453455 | old | Astrocyte |
| SLC1A3 | -1.04707598 | 0.049185508 | young | Astrocyte |
| KLHL7 | -1.223876282 | 0.048377913 | young | Astrocyte |
| METRN | -1.344071232 | 0.040835391 | young | Astrocyte |
| ANGPTL1 | -1.695501659 | 2.36E-06 | young | Astrocyte |
| IFIT1 | -1.70359784 | 0.043149152 | young | Astrocyte |
| IFI44 | -1.717009374 | 0.043149152 | young | Astrocyte |
| AC079793.1 | -1.864200297 | 0.043149152 | young | Astrocyte |
| IFI44L | -2.026553831 | 0.000141575 | young | Astrocyte |
| ISG15 | -2.218118788 | 0.007911922 | young | Astrocyte |
| LINC02649 | 1.806801875 | 1.39E-07 | old | BC |
| LDLRAD4 | 1.293793082 | 5.92E-06 | old | BC |
| PTGDS | 1.172223989 | 0.001527741 | old | BC |
| PFKFB4 | 1.168152754 | 0.00376047 | old | BC |
| SYCP2L | 1.146764568 | 0.002563044 | old | BC |
| PDK1 | 1.129421438 | 0.002961804 | old | BC |
| RAPGEF4 | 1.106954827 | 0.001893064 | old | BC |
| DTNA | 1.083769881 | 0.001386511 | old | BC |
| PFKFB3 | 1.036590795 | 0.002961804 | old | BC |
| GPX3 | 1.008795276 | 0.00772741 | old | BC |
| PCSK5 | 0.990835736 | 0.002895764 | old | BC |
| AK4 | 0.980432375 | 0.009923637 | old | BC |
| S100A10 | 0.975574121 | 0.015482107 | old | BC |
| DNAH17 | 0.974769735 | 0.000299789 | old | BC |
| NPAS2 | 0.915939293 | 0.003094891 | old | BC |
| NDRG1 | 0.897124583 | 0.000740415 | old | BC |
| PLOD2 | 0.883393357 | 0.007167233 | old | BC |
| KDM3A | 0.877625993 | 0.011913685 | old | BC |
| FRZB | 0.877550019 | 0.011087279 | old | BC |
| IGFBP5 | 0.875337457 | 0.043577473 | old | BC |
| MLLT3 | 0.86769407 | 0.005418813 | old | BC |
| CAV1 | 0.858547426 | 0.027639094 | old | BC |
| SH3GL3 | 0.858287647 | 0.007550735 | old | BC |
| DPY19L1 | 0.856195976 | 0.005418813 | old | BC |
| MBOAT2 | 0.855070155 | 0.000129169 | old | BC |
| ZMIZ1-AS1 | 0.838252893 | 0.000112102 | old | BC |
| RFLNA | 0.835184775 | 0.003269694 | old | BC |
| ARHGAP10 | 0.826404442 | 0.025774372 | old | BC |
| LEMD1 | 0.813694778 | 0.003479301 | old | BC |

| | | | | |
|---|---|---|---|---|
| ITPR1 | 0.812279569 | 0.039223185 | old | BC |
| PAM | 0.806605152 | 0.000154775 | old | BC |
| C8orf34 | 0.801748079 | 0.04575009 | old | BC |
| SLC16A10 | 0.798528593 | 0.01905131 | old | BC |
| PDK3 | 0.787321705 | 0.010869247 | old | BC |
| NTRK2 | 0.782652318 | 0.01751365 | old | BC |
| PFKP | 0.781397877 | 0.019188594 | old | BC |
| EFNA5 | 0.779657352 | 0.033108028 | old | BC |
| CREB3L2 | 0.777085618 | 0.001015176 | old | BC |
| AC079760.1 | 0.773861168 | 0.022366982 | old | BC |
| AC027601.6 | 0.773108261 | 0.048090709 | old | BC |
| PXN | 0.762664278 | 0.006170899 | old | BC |
| FHOD3 | 0.758350625 | 0.030232644 | old | BC |
| GSN | 0.755341449 | 0.02761536 | old | BC |
| KDM4C | 0.74838473 | 0.001527741 | old | BC |
| MTERF1 | 0.743932946 | 0.005629377 | old | BC |
| TRPM3 | 0.74090701 | 0.039451019 | old | BC |
| CDK19 | 0.740054476 | 0.006024874 | old | BC |
| GRK3 | 0.731814977 | 0.008053747 | old | BC |
| KAZN | 0.728912258 | 0.011801698 | old | BC |
| RNF24 | 0.72733557 | 0.007683207 | old | BC |
| MEIS2 | 0.72626192 | 0.043575833 | old | BC |
| SMOC2 | 0.713589156 | 0.030232644 | old | BC |
| KDM4B | 0.711866485 | 0.00281983 | old | BC |
| AFF3 | 0.710693819 | 0.003952329 | old | BC |
| YEATS2 | 0.710400316 | 0.013602126 | old | BC |
| NTRK3 | 0.70959307 | 0.032786423 | old | BC |
| AP001972.3 | 0.701925553 | 0.039351655 | old | BC |
| SORBS1 | 0.687617552 | 0.010098804 | old | BC |
| ESYT2 | 0.686412625 | 0.001741148 | old | BC |
| ADARB2 | 0.686025701 | 0.045907084 | old | BC |
| SNX10 | 0.683499495 | 0.005468374 | old | BC |
| GPR137B | 0.682929736 | 0.011920145 | old | BC |
| IFT122 | 0.682657292 | 0.007136086 | old | BC |
| ABCC1 | 0.68222129 | 0.018769241 | old | BC |
| KDM2B-DT | 0.681899855 | 0.02778213 | old | BC |
| EGLN1 | 0.677850448 | 0.020796294 | old | BC |
| CACNA1D | 0.672719346 | 0.000680845 | old | BC |
| RNF165 | 0.67108631 | 0.019644286 | old | BC |
| COL4A2 | 0.670666448 | 0.046444481 | old | BC |
| PRTG | 0.66941547 | 0.044400173 | old | BC |
| PDZD4 | 0.669355033 | 0.046400736 | old | BC |
| ZNF516 | 0.669278781 | 0.00854587 | old | BC |
| FNTB | 0.667691033 | 0.017055621 | old | BC |
| NEAT1 | 0.664527695 | 0.013554406 | old | BC |
| AC007785.1 | 0.66303291 | 0.024636217 | old | BC |
| AGAP1 | 0.659870398 | 0.000120221 | old | BC |
| AC025809.1 | 0.658575689 | 0.020222544 | old | BC |
| ARFGEF3 | 0.65341016 | 0.02333436 | old | BC |
| CDC14B | 0.651253901 | 0.016018756 | old | BC |
| C11orf80 | 0.650941761 | 0.012193191 | old | BC |
| TFAP2E | 0.646259052 | 0.014203594 | old | BC |
| TNIK | 0.641814138 | 0.000172791 | old | BC |
| RNF217-AS1 | 0.640859965 | 0.018718475 | old | BC |

| | | | | |
|---|---|---|---|---|
| AL591519.1 | 0.635272581 | 0.0362505 | old | BC |
| DGCR9 | 0.634691032 | 0.008362082 | old | BC |
| MANBA | 0.633870832 | 0.018031171 | old | BC |
| CDHR3 | 0.633607502 | 0.02778213 | old | BC |
| ST6GALNAC3 | 0.631486045 | 0.032765075 | old | BC |
| OGA | 0.629188108 | 4.82E-05 | old | BC |
| BRWD3 | 0.625312487 | 0.011497988 | old | BC |
| GLCCI1 | 0.623957852 | 0.029157768 | old | BC |
| ANTXR2 | 0.623831668 | 0.002471225 | old | BC |
| GPI | 0.620528172 | 0.02615806 | old | BC |
| PTPN21 | 0.620204745 | 0.02778213 | old | BC |
| RBPJ | 0.617729196 | 0.009221837 | old | BC |
| SEPTIN6 | 0.617194683 | 0.043606945 | old | BC |
| TENM2 | 0.613564777 | 0.04575009 | old | BC |
| XPNPEP1 | 0.613489125 | 0.011012232 | old | BC |
| DIP2C | 0.612782977 | 0.000993859 | old | BC |
| TEX9 | 0.609244591 | 0.028992684 | old | BC |
| LIMS1 | 0.609116191 | 0.00095508 | old | BC |
| CPD | 0.608294755 | 0.040825699 | old | BC |
| SNX25 | 0.604608827 | 0.0041238 | old | BC |
| PIAS2 | 0.604291523 | 0.005023154 | old | BC |
| XKR6 | 0.603610261 | 0.003556621 | old | BC |
| PLEKHB1 | 0.602248436 | 0.032746846 | old | BC |
| ULK1 | 0.60062811 | 0.037180449 | old | BC |
| SNTB1 | 0.599590821 | 0.01999373 | old | BC |
| SLCO3A1 | 0.595702243 | 0.041001837 | old | BC |
| SFMBT2 | 0.592977233 | 0.026962102 | old | BC |
| TNNI3K | 0.584210487 | 0.013129109 | old | BC |
| RRAGD | 0.583671898 | 0.040028245 | old | BC |
| ZBTB16 | 0.580815238 | 0.0342766 | old | BC |
| EPS15 | 0.576497498 | 0.00475703 | old | BC |
| CPEB3 | 0.569809175 | 0.009449429 | old | BC |
| STK39 | 0.569068674 | 0.029933599 | old | BC |
| DOCK7 | 0.568389094 | 0.021168507 | old | BC |
| C20orf194 | 0.567278699 | 0.00772741 | old | BC |
| AC002460.2 | 0.566444588 | 0.04183146 | old | BC |
| KLF12 | 0.565502959 | 0.029570559 | old | BC |
| NRG2 | 0.564725306 | 0.025774372 | old | BC |
| SEL1L3 | 0.562486202 | 0.042342172 | old | BC |
| RORA | 0.562311149 | 0.008362082 | old | BC |
| ADCY1 | 0.559441669 | 0.003328804 | old | BC |
| UBR5 | 0.556614099 | 0.001169105 | old | BC |
| ARL15 | 0.554232793 | 0.000993859 | old | BC |
| ZNF248 | 0.551768188 | 0.010736555 | old | BC |
| RPS6KA2 | 0.549340304 | 0.002285319 | old | BC |
| SLC25A29 | 0.548204666 | 0.035906999 | old | BC |
| ADCY9 | 0.548150093 | 0.026561895 | old | BC |
| TPT1-AS1 | 0.547982312 | 0.001439926 | old | BC |
| HDAC7 | 0.547972653 | 0.042672187 | old | BC |
| LINC-PINT | 0.546457218 | 0.029697813 | old | BC |
| TTLL5 | 0.545514413 | 0.016649174 | old | BC |
| XPO6 | 0.545469221 | 0.001527741 | old | BC |
| MAP2K5 | 0.543534369 | 0.004118527 | old | BC |
| MALT1 | 0.537783987 | 0.025299434 | old | BC |

| | | | | |
|---|---|---|---|---|
| LRRFIP1 | 0.537279721 | 0.001043935 | old | BC |
| SLC15A4 | 0.535098421 | 0.031229604 | old | BC |
| DENND1A | 0.533145992 | 0.002727547 | old | BC |
| FMNL2 | 0.532909689 | 0.012900287 | old | BC |
| AL731577.2 | 0.529093881 | 0.032031834 | old | BC |
| DOCK4 | 0.529033727 | 0.006021858 | old | BC |
| PRKAA2 | 0.528935562 | 0.003328804 | old | BC |
| GGA2 | 0.527365338 | 0.033292615 | old | BC |
| APBB2 | 0.527217251 | 0.006066077 | old | BC |
| CPSF6 | 0.527067073 | 0.002160912 | old | BC |
| LRCH1 | 0.525942824 | 0.000908376 | old | BC |
| ARL6 | 0.525756161 | 0.025207239 | old | BC |
| ASAP2 | 0.524919414 | 0.003521136 | old | BC |
| CARMIL1 | 0.520722717 | 0.029933599 | old | BC |
| RAPGEF6 | 0.520708001 | 0.03082707 | old | BC |
| DAPK1 | 0.518848325 | 0.00786318 | old | BC |
| RALGAPA2 | 0.51835351 | 0.013033194 | old | BC |
| ZNF266 | 0.518018255 | 0.043558184 | old | BC |
| BTAF1 | 0.516467163 | 0.043302262 | old | BC |
| AEBP2 | 0.516135356 | 0.016018756 | old | BC |
| CACNA1C | 0.513436746 | 0.019964472 | old | BC |
| KIF13B | 0.511260975 | 0.020728579 | old | BC |
| ATP1B3 | 0.509732992 | 0.038654859 | old | BC |
| TMEM132C | 0.509147836 | 0.003979162 | old | BC |
| KMT2C | 0.503774385 | 0.002950019 | old | BC |
| TTC13 | 0.501349463 | 0.048090709 | old | BC |
| TRIO | 0.500590437 | 0.005331007 | old | BC |
| RIMBP2 | 0.499367535 | 0.022627224 | old | BC |
| FOCAD | 0.497811216 | 0.002160912 | old | BC |
| DNM3 | 0.495763139 | 0.02778213 | old | BC |
| ZSWIM5 | 0.495384129 | 0.019644286 | old | BC |
| SH3PXD2A | 0.494645536 | 0.026962102 | old | BC |
| PCCA | 0.492649403 | 0.003556621 | old | BC |
| RBM33 | 0.492488572 | 0.016729167 | old | BC |
| SLC25A37 | 0.488246886 | 0.010963537 | old | BC |
| MTCL1 | 0.488183703 | 0.005434775 | old | BC |
| CLASP1 | 0.487948117 | 0.001893064 | old | BC |
| KCNQ3 | 0.487919451 | 0.034717909 | old | BC |
| GRB10 | 0.487322021 | 0.013967815 | old | BC |
| RLF | 0.483895065 | 0.045760116 | old | BC |
| FMN2 | 0.483626731 | 0.030841312 | old | BC |
| PARP8 | 0.483407961 | 0.03701806 | old | BC |
| MED12L | 0.481659119 | 0.037508633 | old | BC |
| USP24 | 0.480540184 | 0.036444229 | old | BC |
| PTPRJ | 0.479135618 | 0.02778213 | old | BC |
| TTBK2 | 0.478852855 | 0.017579657 | old | BC |
| AC068587.4 | 0.478151103 | 0.027272363 | old | BC |
| HACE1 | 0.477125794 | 0.045794157 | old | BC |
| MDGA1 | 0.476006152 | 0.028248543 | old | BC |
| SETD5 | 0.475001021 | 0.008007975 | old | BC |
| OTX2-AS1 | 0.474441511 | 0.012970443 | old | BC |
| FBXO42 | 0.472087389 | 0.029837047 | old | BC |
| DAAM1 | 0.471982799 | 0.027143282 | old | BC |
| SSH2 | 0.471514068 | 0.007103552 | old | BC |

| | | | | |
|---|---|---|---|---|
| MAP3K5 | 0.46877853 | 0.04717699 | old | BC |
| SIPA1L3 | 0.467803518 | 0.017823005 | old | BC |
| ELF2 | 0.466146778 | 0.025774372 | old | BC |
| SLAIN1 | 0.465351552 | 0.041458943 | old | BC |
| ZCCHC14 | 0.462954933 | 0.019168335 | old | BC |
| EP400 | 0.462212545 | 0.003589566 | old | BC |
| PARN | 0.461859659 | 0.019749387 | old | BC |
| RERE | 0.461382499 | 0.008484459 | old | BC |
| PICALM | 0.460121262 | 0.009791753 | old | BC |
| PARD3 | 0.459993425 | 0.019644286 | old | BC |
| SEC24B | 0.459630438 | 0.019655224 | old | BC |
| NUP58 | 0.459547439 | 0.030408412 | old | BC |
| ARHGEF7 | 0.458562761 | 0.005027089 | old | BC |
| MGAT5 | 0.457043716 | 0.029567196 | old | BC |
| HDAC4 | 0.456318456 | 0.030841312 | old | BC |
| CLIP4 | 0.455445585 | 0.034706673 | old | BC |
| KDM7A | 0.454255301 | 0.04183146 | old | BC |
| AHCYL2 | 0.453333672 | 0.023866718 | old | BC |
| PUM2 | 0.451540537 | 0.01888407 | old | BC |
| NBAS | 0.45032113 | 0.014010639 | old | BC |
| METTL15 | 0.450304221 | 0.025752085 | old | BC |
| TAOK1 | 0.449495762 | 0.020776974 | old | BC |
| UBR3 | 0.449006996 | 0.018132831 | old | BC |
| MLLT10 | 0.447980906 | 0.027639094 | old | BC |
| DSCAML1 | 0.447596555 | 0.022988474 | old | BC |
| MAP4K5 | 0.446445868 | 0.015628459 | old | BC |
| BRWD1 | 0.446317346 | 0.005364869 | old | BC |
| USP34 | 0.445745857 | 0.01999373 | old | BC |
| DIP2A | 0.444827346 | 0.01911864 | old | BC |
| SCN2A | 0.443715187 | 0.014662328 | old | BC |
| PHF21A | 0.443397627 | 0.000993859 | old | BC |
| CDK8 | 0.44271736 | 0.005104031 | old | BC |
| EVL | 0.440117849 | 0.012265558 | old | BC |
| CBLB | 0.439800919 | 0.027143282 | old | BC |
| TSPAN9 | 0.438108348 | 0.025559958 | old | BC |
| ARID2 | 0.437389044 | 0.018541822 | old | BC |
| BIRC6 | 0.43421367 | 0.011278507 | old | BC |
| PHC3 | 0.433799595 | 0.048981449 | old | BC |
| TRERF1 | 0.433499869 | 0.014662328 | old | BC |
| ARMH3 | 0.433391412 | 0.022083389 | old | BC |
| CPNE1 | 0.432185534 | 0.049326547 | old | BC |
| SFXN1 | 0.430760784 | 0.025559958 | old | BC |
| SCN8A | 0.430527663 | 0.027272363 | old | BC |
| RAVER2 | 0.430341673 | 0.049604946 | old | BC |
| LARP4B | 0.430099595 | 0.028248543 | old | BC |
| LMCD1-AS1 | 0.429369087 | 0.027639094 | old | BC |
| KIZ | 0.428502336 | 0.027272363 | old | BC |
| AKAP10 | 0.426567866 | 0.025752085 | old | BC |
| LRCH3 | 0.426130003 | 0.036871487 | old | BC |
| TFDP2 | 0.425487033 | 0.006596211 | old | BC |
| TRPC1 | 0.423787664 | 0.029228688 | old | BC |
| USP49 | 0.423551812 | 0.012265558 | old | BC |
| FAM193A | 0.423446956 | 0.024639395 | old | BC |
| AFDN | 0.422506751 | 0.030763951 | old | BC |

| | | | | | |
|---|---|---|---|---|---|
| KATNAL1 | | 0.41847418 | 0.031489344 | old | BC |
| ZNF438 | | 0.417728472 | 0.012334557 | old | BC |
| DNAJC6 | | 0.416762987 | 0.007938121 | old | BC |
| PDS5A | | 0.416481131 | 0.014662328 | old | BC |
| PCNX1 | | 0.41558267 | 0.012015566 | old | BC |
| HERC4 | | 0.415522494 | 0.021606289 | old | BC |
| PPP1R12B | | 0.414256701 | 0.045258018 | old | BC |
| MAGI1 | | 0.414029671 | 0.034949064 | old | BC |
| SIPA1L1 | | 0.413938754 | 0.04717699 | old | BC |
| WDR59 | | 0.412873651 | 0.033731217 | old | BC |
| PPP6R3 | | 0.412192813 | 0.013033194 | old | BC |
| NF1 | | 0.412015833 | 0.001796975 | old | BC |
| IGF1R | | 0.411241496 | 0.026561895 | old | BC |
| ATXN1 | | 0.410458179 | 0.003965308 | old | BC |
| ERC1 | | 0.409418624 | 0.008344309 | old | BC |
| MON2 | | 0.408577903 | 0.048382294 | old | BC |
| PAN3 | | 0.408229762 | 0.002600096 | old | BC |
| MKLN1 | | 0.407032284 | 0.035753382 | old | BC |
| CBFA2T2 | | 0.406251606 | 0.014677083 | old | BC |
| LRBA | | 0.405644108 | 0.008007975 | old | BC |
| SMARCA2 | | 0.405297265 | 0.011686913 | old | BC |
| MSRB3 | | 0.404768415 | 0.006043277 | old | BC |
| MED13L | | 0.404404884 | 0.037067829 | old | BC |
| FCHSD2 | | 0.404189041 | 0.014077963 | old | BC |
| MAP4K3 | | 0.402258863 | 0.033267316 | old | BC |
| ARHGAP32 | | 0.399549707 | 0.029933599 | old | BC |
| KIF1B | | 0.399138219 | 0.02778213 | old | BC |
| HMBOX1 | | 0.398503436 | 0.037302018 | old | BC |
| ZNF451 | | 0.397459406 | 0.039199228 | old | BC |
| HDAC9 | | 0.395529641 | 0.01350432 | old | BC |
| TENT2 | | 0.394456785 | 0.013329769 | old | BC |
| GRID1 | | 0.394213983 | 0.03090144 | old | BC |
| KMT2E | | 0.392289821 | 0.034390354 | old | BC |
| DENND6A | | 0.391568924 | 0.041142471 | old | BC |
| CDC27 | | 0.390868407 | 0.038252033 | old | BC |
| POGZ | | 0.388865852 | 0.024572428 | old | BC |
| DYM | | 0.388807153 | 0.01905131 | old | BC |
| CEP85L | | 0.387471875 | 0.02778213 | old | BC |
| KIF13A | | 0.386254175 | 0.009065004 | old | BC |
| AGO4 | | 0.385996646 | 0.048326072 | old | BC |
| ADD3 | | 0.385533305 | 0.024378692 | old | BC |
| FRMD4A | | 0.385115629 | 0.043214929 | old | BC |
| QKI | | 0.383361053 | 0.032878857 | old | BC |
| TAF1 | | 0.383173498 | 0.042259512 | old | BC |
| ZMYM2 | | 0.382553856 | 0.031753539 | old | BC |
| USP3 | | 0.382036465 | 0.033427316 | old | BC |
| ARFGEF2 | | 0.381507939 | 0.034717909 | old | BC |
| | 6-Mar | 0.381426157 | 0.0450457 | old | BC |
| WDFY2 | | 0.380874748 | 0.046382575 | old | BC |
| VOPP1 | | 0.38012802 | 0.042956961 | old | BC |
| CADPS | | 0.378634512 | 0.031948746 | old | BC |
| AUH | | 0.377940459 | 0.01794918 | old | BC |
| SBF2 | | 0.377774715 | 0.048090709 | old | BC |
| SMARCC1 | | 0.377578867 | 0.030232644 | old | BC |

| | | | | |
|---|---|---|---|---|
| EIF4G3 | 0.37751339 | 0.042672187 | old | BC |
| NPEPPS | 0.377380972 | 0.016649174 | old | BC |
| ARID1B | 0.375984492 | 0.010051966 | old | BC |
| EPB41L3 | 0.374526616 | 0.032641337 | old | BC |
| PELI2 | 0.371686992 | 0.008003222 | old | BC |
| NKTR | 0.371153439 | 0.026689815 | old | BC |
| UBE2E2 | 0.369677623 | 0.03040265 | old | BC |
| ANKRD28 | 0.369019293 | 0.0450457 | old | BC |
| HERC1 | 0.365234134 | 0.014662328 | old | BC |
| SVIL | 0.365024875 | 0.048765422 | old | BC |
| POU2F1 | 0.36489055 | 0.04318227 | old | BC |
| CASK | 0.364770426 | 0.031363645 | old | BC |
| CAMK1D | 0.364667596 | 0.019536126 | old | BC |
| CLEC16A | 0.362217948 | 0.032355641 | old | BC |
| WDFY3 | 0.360532241 | 0.025207239 | old | BC |
| CDC42BPA | 0.360292555 | 0.016635049 | old | BC |
| SOS1 | 0.360280096 | 0.018769241 | old | BC |
| CELF2 | 0.360267538 | 0.037331366 | old | BC |
| LARGE1 | 0.359590209 | 0.025616868 | old | BC |
| AKT3 | 0.359377224 | 0.01786878 | old | BC |
| KLHL7 | 0.356538509 | 0.027272363 | old | BC |
| LRRC1 | 0.356037405 | 0.02104957 | old | BC |
| DENND5B | 0.354363432 | 0.028729951 | old | BC |
| RAB6A | 0.354015339 | 0.042878762 | old | BC |
| CLMN | 0.353094293 | 0.013163528 | old | BC |
| IQSEC1 | 0.352529277 | 0.033388405 | old | BC |
| SERGEF | 0.352515742 | 0.037894563 | old | BC |
| TBL1XR1 | 0.3504358 | 0.01361916 | old | BC |
| ACAP2 | 0.350266466 | 0.003556621 | old | BC |
| PPP2R5E | 0.350166934 | 0.015009037 | old | BC |
| NRCAM | 0.34472831 | 0.028217934 | old | BC |
| PTK2 | 0.344026696 | 0.009582184 | old | BC |
| KDM2A | 0.34304587 | 0.010736555 | old | BC |
| UBE3C | 0.341724294 | 0.029933599 | old | BC |
| SORCS2 | 0.341653105 | 0.038656286 | old | BC |
| BTBD9 | 0.340458485 | 0.033304332 | old | BC |
| LCOR | 0.337939862 | 0.044565596 | old | BC |
| EML5 | 0.33483495 | 0.045794157 | old | BC |
| SMAD2 | 0.32924529 | 0.019188594 | old | BC |
| COP1 | 0.319251584 | 0.036871487 | old | BC |
| FYN | 0.318413143 | 0.022627224 | old | BC |
| MCC | 0.315389052 | 0.013129109 | old | BC |
| TRIM33 | 0.311979705 | 0.039120247 | old | BC |
| CCDC88A | 0.310983038 | 0.036874092 | old | BC |
| TFCP2 | 0.309674611 | 0.045303891 | old | BC |
| SIK3 | 0.309356478 | 0.044789158 | old | BC |
| REV1 | 0.307237368 | 0.034047485 | old | BC |
| FTO | 0.290589602 | 0.032878857 | old | BC |
| CAMTA1 | 0.288468002 | 0.021006585 | old | BC |
| TNRC6B | 0.278631088 | 0.025341057 | old | BC |
| XIST | -0.122238381 | 0.041159604 | young | BC |
| PSMC6 | -0.316891166 | 0.046601162 | young | BC |
| SRP72 | -0.369702226 | 0.049071147 | young | BC |
| SLTM | -0.378617034 | 0.046601162 | young | BC |

| | | | | |
|---|---|---|---|---|
| PLGRKT | -0.38163202 | 0.046609962 | young | BC |
| EAPP | -0.387336556 | 0.043244464 | young | BC |
| UBE2L3 | -0.388378361 | 0.030841312 | young | BC |
| SRFBP1 | -0.388674476 | 0.017055621 | young | BC |
| PSIP1 | -0.395156239 | 0.026200181 | young | BC |
| SMC3 | -0.402439966 | 0.018769241 | young | BC |
| SMIM15 | -0.403809243 | 0.045588967 | young | BC |
| RSRC2 | -0.404716538 | 0.043558184 | young | BC |
| PCSK1N | -0.406699735 | 0.037548813 | young | BC |
| PSMB7 | -0.418512651 | 0.048596872 | young | BC |
| DDX3X | -0.418717335 | 0.03421855 | young | BC |
| FAM120AOS | -0.42032701 | 0.021152216 | young | BC |
| EIF2S1 | -0.420838906 | 0.043575833 | young | BC |
| ARL14EP | -0.422910404 | 0.010911005 | young | BC |
| HNRNPA2B1 | -0.425737661 | 0.032598113 | young | BC |
| SSB | -0.426716295 | 0.048090709 | young | BC |
| MED31 | -0.426784045 | 0.038020794 | young | BC |
| DEK | -0.426862339 | 0.037282638 | young | BC |
| PPARGC1B | -0.42913174 | 0.02973824 | young | BC |
| RTF2 | -0.43129852 | 0.030326028 | young | BC |
| CCDC112 | -0.434469035 | 0.030326028 | young | BC |
| SERBP1 | -0.438336197 | 0.0313162 | young | BC |
| GGCX | -0.439179502 | 0.046444481 | young | BC |
| CLUAP1 | -0.439721834 | 0.031708044 | young | BC |
| MRPS10 | -0.442201741 | 0.03421855 | young | BC |
| PWP1 | -0.442819476 | 0.026818994 | young | BC |
| SEC62 | -0.444691267 | 0.033267316 | young | BC |
| ISCA1 | -0.445539856 | 0.042490981 | young | BC |
| TIMM17A | -0.446395614 | 0.049925087 | young | BC |
| PPP1R7 | -0.448247535 | 0.035755632 | young | BC |
| PFDN1 | -0.448921153 | 0.026200181 | young | BC |
| DDX52 | -0.450623554 | 0.019168335 | young | BC |
| KIN | -0.450968757 | 0.049071147 | young | BC |
| SLC25A6 | -0.452134872 | 0.040920084 | young | BC |
| FRG1 | -0.452208891 | 0.020386674 | young | BC |
| MTHFS | -0.454030898 | 0.012281654 | young | BC |
| SUCLA2 | -0.454773521 | 0.025299434 | young | BC |
| FGD5-AS1 | -0.455696286 | 0.035067726 | young | BC |
| TCEAL4 | -0.457006487 | 0.025774372 | young | BC |
| ARPC3 | -0.458382782 | 0.044330628 | young | BC |
| SMIM19 | -0.460189648 | 0.030785579 | young | BC |
| ROGDI | -0.460304626 | 0.040207551 | young | BC |
| NARS | -0.461843767 | 0.00772741 | young | BC |
| SGCB | -0.463048856 | 0.046326437 | young | BC |
| CDC5L | -0.464087094 | 0.025752085 | young | BC |
| TMEM126B | -0.46735497 | 0.043389771 | young | BC |
| SOD1 | -0.472918568 | 0.03090144 | young | BC |
| CYCS | -0.473413565 | 0.040752239 | young | BC |
| SNRPC | -0.474397689 | 0.039357848 | young | BC |
| ASF1A | -0.475738985 | 0.048090709 | young | BC |
| TMEM205 | -0.47748789 | 0.045598454 | young | BC |
| NIFK | -0.477748586 | 0.04821636 | young | BC |
| MRPL1 | -0.477909974 | 0.013592356 | young | BC |
| DNAJC9 | -0.478601545 | 0.023672954 | young | BC |

| | | | | |
|---|---|---|---|---|
| SSR4 | -0.478690981 | 0.012453813 | young | BC |
| TFAM | -0.479066249 | 0.024882691 | young | BC |
| CGRRF1 | -0.479214071 | 0.021343015 | young | BC |
| TSPAN31 | -0.479429969 | 0.017519463 | young | BC |
| HMGB1 | -0.479558431 | 0.005418813 | young | BC |
| MMAB | -0.480856599 | 0.04958629 | young | BC |
| C14orf132 | -0.48163613 | 0.041204863 | young | BC |
| TBCA | -0.483716913 | 0.029933599 | young | BC |
| SNW1 | -0.484062943 | 0.007893838 | young | BC |
| PSMA3 | -0.484566936 | 0.031708044 | young | BC |
| RABAC1 | -0.486516771 | 0.046173276 | young | BC |
| POLR2F | -0.487817903 | 0.04575009 | young | BC |
| FAM174A | -0.490258765 | 0.013554406 | young | BC |
| CRELD2 | -0.491432838 | 0.048090709 | young | BC |
| RAN | -0.492705512 | 0.029697813 | young | BC |
| NDUFV3 | -0.492872715 | 0.025299434 | young | BC |
| CHMP3 | -0.493009732 | 0.006024874 | young | BC |
| JKAMP | -0.493691381 | 0.043302262 | young | BC |
| FAIM2 | -0.493797005 | 0.002014262 | young | BC |
| GSTP1 | -0.496695964 | 0.026173488 | young | BC |
| SCG3 | -0.497375473 | 0.020781984 | young | BC |
| MZF1 | -0.498430653 | 0.011017176 | young | BC |
| PFDN4 | -0.498774164 | 0.045258018 | young | BC |
| RPF1 | -0.49948781 | 0.039018783 | young | BC |
| LRPAP1 | -0.499712588 | 0.033375227 | young | BC |
| DYNLRB1 | -0.500336464 | 0.043214929 | young | BC |
| MPC2 | -0.500489626 | 0.023182921 | young | BC |
| MYL6 | -0.502076102 | 0.021006585 | young | BC |
| DAD1 | -0.502222366 | 0.039949417 | young | BC |
| SEC61B | -0.502260605 | 0.04717699 | young | BC |
| PLRG1 | -0.502497018 | 0.026689815 | young | BC |
| DMAC1 | -0.503394084 | 0.036010088 | young | BC |
| NDUFAF2 | -0.504673288 | 0.009066567 | young | BC |
| RFK | -0.505179572 | 0.001527741 | young | BC |
| BCAS2 | -0.506020371 | 0.017213701 | young | BC |
| CFAP20 | -0.506263317 | 0.04183146 | young | BC |
| DNAJC8 | -0.506496924 | 0.006043277 | young | BC |
| POMP | -0.506881705 | 0.026561895 | young | BC |
| C1D | -0.507112339 | 0.025341057 | young | BC |
| DUT | -0.50730444 | 0.008003222 | young | BC |
| NAPG | -0.508155874 | 0.015490029 | young | BC |
| C9orf78 | -0.509103322 | 0.022400069 | young | BC |
| ESF1 | -0.509424487 | 0.003927927 | young | BC |
| SNRPD1 | -0.511621026 | 0.044089255 | young | BC |
| HTATSF1 | -0.511621512 | 0.025294749 | young | BC |
| ATP5PF | -0.51318107 | 0.029933599 | young | BC |
| COX7B | -0.51400893 | 0.047145941 | young | BC |
| NAA20 | -0.514851646 | 0.040131863 | young | BC |
| NDUFA5 | -0.516154663 | 0.006678554 | young | BC |
| C6orf120 | -0.516420376 | 0.027272363 | young | BC |
| PNKD | -0.516656672 | 0.044643214 | young | BC |
| DNMT1 | -0.516829298 | 0.016018756 | young | BC |
| TMA7 | -0.517058873 | 0.02532099 | young | BC |
| METTL5 | -0.517297052 | 0.032598113 | young | BC |

| | | | | |
|---|---|---|---|---|
| C3orf14 | -0.517712453 | 0.018769241 | young | BC |
| PNPLA4 | -0.518423146 | 0.025589147 | young | BC |
| UBLCP1 | -0.518858938 | 0.017316294 | young | BC |
| MRPS21 | -0.520285302 | 0.036010088 | young | BC |
| PSMD8 | -0.521492907 | 0.044643214 | young | BC |
| DDX46 | -0.52150558 | 0.003328804 | young | BC |
| RNF7 | -0.522633557 | 0.032426365 | young | BC |
| RIDA | -0.522699549 | 0.028339189 | young | BC |
| BLOC1S2 | -0.523194153 | 0.043994234 | young | BC |
| MXRA7 | -0.526595393 | 0.039223185 | young | BC |
| LSM3 | -0.527259647 | 0.01542006 | young | BC |
| NDUFS7 | -0.527380733 | 0.044914818 | young | BC |
| KIF1BP | -0.527675722 | 0.01542006 | young | BC |
| NEDD8 | -0.528567409 | 0.026561895 | young | BC |
| VPS29 | -0.529007225 | 0.011278507 | young | BC |
| LINC00667 | -0.529308466 | 0.005418813 | young | BC |
| PAM16 | -0.529887494 | 0.048945739 | young | BC |
| FUNDC2 | -0.532202662 | 0.029933599 | young | BC |
| PSMD4 | -0.533033552 | 0.02456591 | young | BC |
| LINC02456 | -0.5345781 | 0.033304332 | young | BC |
| RUVBL1 | -0.535971683 | 0.041142471 | young | BC |
| ATP5F1C | -0.537497438 | 0.048596872 | young | BC |
| ZNF622 | -0.538659757 | 0.0450457 | young | BC |
| MZT1 | -0.54096849 | 0.026200181 | young | BC |
| COMMD8 | -0.541013589 | 0.026689815 | young | BC |
| TALDO1 | -0.542605671 | 0.032878857 | young | BC |
| B3GALT2 | -0.542883695 | 0.049270531 | young | BC |
| C19orf53 | -0.542889412 | 0.029958394 | young | BC |
| NPM1 | -0.54310433 | 0.030232644 | young | BC |
| NDUFV2 | -0.543739699 | 0.019188594 | young | BC |
| CLTA | -0.544932134 | 0.027639094 | young | BC |
| LINC01003 | -0.545319651 | 0.017055621 | young | BC |
| PSMD7 | -0.54550975 | 0.027683806 | young | BC |
| ZBTB24 | -0.545765159 | 0.007776203 | young | BC |
| SYF2 | -0.546305397 | 0.006678554 | young | BC |
| COPE | -0.546992624 | 0.004697837 | young | BC |
| MRPL40 | -0.547228395 | 0.0450457 | young | BC |
| SRA1 | -0.547526551 | 0.006678554 | young | BC |
| SMIM8 | -0.547814122 | 0.030491865 | young | BC |
| CCDC25 | -0.547935069 | 0.03095039 | young | BC |
| CHCHD2 | -0.549334392 | 0.043581319 | young | BC |
| PSMB4 | -0.549487874 | 0.032775465 | young | BC |
| TMEM242 | -0.549665661 | 0.013129109 | young | BC |
| VAMP2 | -0.550534107 | 0.03547219 | young | BC |
| PSMG1 | -0.551231083 | 0.010650469 | young | BC |
| TCEAL3 | -0.552939739 | 0.014662328 | young | BC |
| CENPX | -0.553170553 | 0.024572428 | young | BC |
| ERLEC1 | -0.553921448 | 0.017316294 | young | BC |
| MRPS18C | -0.554309524 | 0.008212474 | young | BC |
| NUDT16 | -0.554383452 | 0.02778213 | young | BC |
| LSM4 | -0.555620733 | 0.045042256 | young | BC |
| ARPC5L | -0.555864201 | 0.039223185 | young | BC |
| PSMA5 | -0.556096217 | 0.030837783 | young | BC |
| TMBIM4 | -0.556256923 | 0.006678554 | young | BC |

| | | | | |
|---|---|---|---|---|
| SYS1 | -0.557182383 | 0.017238727 | young | BC |
| JPT1 | -0.557316724 | 0.029933599 | young | BC |
| SMDT1 | -0.558823298 | 0.049377292 | young | BC |
| WDR74 | -0.560274909 | 0.027325672 | young | BC |
| ITGAE | -0.560464469 | 0.0450457 | young | BC |
| AK6 | -0.56072767 | 0.01521024 | young | BC |
| MRPL51 | -0.560857241 | 0.015073588 | young | BC |
| CCT7 | -0.560867662 | 0.032422398 | young | BC |
| NENF | -0.561414462 | 0.009060748 | young | BC |
| ATP5MD | -0.562838191 | 0.011506448 | young | BC |
| MAGEH1 | -0.563027013 | 0.021106681 | young | BC |
| CHMP5 | -0.563538699 | 0.03421855 | young | BC |
| SRSF7 | -0.563856115 | 0.012334557 | young | BC |
| COMMD4 | -0.56408669 | 0.037180449 | young | BC |
| AL589740.1 | -0.564095617 | 0.025207239 | young | BC |
| DCAF13 | -0.564251418 | 0.012015566 | young | BC |
| WASHC3 | -0.564572565 | 0.002160912 | young | BC |
| FAM136A | -0.564576297 | 0.026200181 | young | BC |
| GPATCH11 | -0.564923145 | 0.038346119 | young | BC |
| C12orf65 | -0.565270942 | 0.014662328 | young | BC |
| COX6A1 | -0.566292042 | 0.011978321 | young | BC |
| MFAP1 | -0.569005124 | 0.018769241 | young | BC |
| PSMA6 | -0.569017322 | 0.003253915 | young | BC |
| RP9 | -0.569669466 | 0.043575833 | young | BC |
| BEX4 | -0.569709463 | 0.03624405 | young | BC |
| UBTF | -0.570001216 | 0.043871411 | young | BC |
| SSNA1 | -0.570303264 | 0.016700364 | young | BC |
| SLC25A4 | -0.570621344 | 0.033906471 | young | BC |
| MPV17L | -0.570691472 | 0.034390354 | young | BC |
| ZNF428 | -0.571049768 | 0.005665991 | young | BC |
| SLIRP | -0.572901929 | 0.024636217 | young | BC |
| KCNMA1-AS1 | -0.573712835 | 0.045113065 | young | BC |
| PPCS | -0.573731541 | 0.02778213 | young | BC |
| ELP6 | -0.573809629 | 0.034370606 | young | BC |
| MLX | -0.573884549 | 0.032203906 | young | BC |
| MED7 | -0.573935514 | 0.032878857 | young | BC |
| IDH3B | -0.575211013 | 0.037790333 | young | BC |
| C12orf57 | -0.575255055 | 0.002285319 | young | BC |
| BUD23 | -0.575945561 | 0.034949064 | young | BC |
| TRAPPC2L | -0.576651573 | 0.02778213 | young | BC |
| ATP6V0B | -0.57700777 | 0.046809766 | young | BC |
| PDHA1 | -0.577393808 | 0.012320806 | young | BC |
| DCTPP1 | -0.577527402 | 0.038284428 | young | BC |
| MRPL13 | -0.578076215 | 0.013526043 | young | BC |
| POLR2I | -0.57868979 | 0.013129109 | young | BC |
| COX5B | -0.579250314 | 0.02854322 | young | BC |
| LLPH | -0.5794677 | 0.035126021 | young | BC |
| TSFM | -0.580131995 | 0.025299434 | young | BC |
| EIF5 | -0.580429776 | 0.009130819 | young | BC |
| NKAPL | -0.580750508 | 0.018232441 | young | BC |
| HAGH | -0.581088006 | 0.035329574 | young | BC |
| UCHL3 | -0.581421767 | 0.016712681 | young | BC |
| MYL12B | -0.581843967 | 0.013180802 | young | BC |
| MRPL20 | -0.582149563 | 0.020386674 | young | BC |

| | | | | |
|---|---|---|---|---|
| NUDT6 | -0.582288256 | 0.013602126 | young | BC |
| NDUFA6 | -0.583063961 | 0.007683207 | young | BC |
| NHP2 | -0.583466763 | 0.012244096 | young | BC |
| UQCR11 | -0.583598155 | 0.033108028 | young | BC |
| CISD1 | -0.583739016 | 0.005199497 | young | BC |
| ELOB | -0.58387742 | 0.0145941 | young | BC |
| MRPL14 | -0.584302221 | 0.032853826 | young | BC |
| RIIAD1 | -0.584733414 | 0.01888407 | young | BC |
| SET | -0.584834499 | 0.0134301 | young | BC |
| UBE2S | -0.585388128 | 0.01350432 | young | BC |
| RWDD1 | -0.586128202 | 0.001162661 | young | BC |
| UBXN1 | -0.586196227 | 0.037180449 | young | BC |
| MICOS10 | -0.587561906 | 0.032203906 | young | BC |
| CWC15 | -0.587584924 | 0.006021858 | young | BC |
| TIMM29 | -0.587891308 | 0.023257813 | young | BC |
| MED8 | -0.588774048 | 0.029570559 | young | BC |
| PCNA | -0.588901473 | 0.032031834 | young | BC |
| AK1 | -0.589084747 | 0.043159887 | young | BC |
| MRPL28 | -0.589300791 | 0.049836618 | young | BC |
| NDUFA12 | -0.589321816 | 0.013763973 | young | BC |
| NUP42 | -0.589854296 | 0.0052644 | young | BC |
| MIS12 | -0.589871823 | 0.022627224 | young | BC |
| RRP15 | -0.590273327 | 0.012015566 | young | BC |
| VBP1 | -0.591223954 | 0.008362082 | young | BC |
| GADD45A | -0.591298749 | 0.003349244 | young | BC |
| CCT2 | -0.591544756 | 0.006421049 | young | BC |
| PLEKHJ1 | -0.592624386 | 0.025207239 | young | BC |
| CCDC181 | -0.593565437 | 0.003556621 | young | BC |
| SELENOH | -0.594291449 | 0.009118887 | young | BC |
| CHMP2A | -0.594529879 | 0.008519127 | young | BC |
| MPHOSPH10 | -0.594803982 | 0.019964472 | young | BC |
| LACTB2 | -0.595124453 | 0.010581847 | young | BC |
| ATP5MPL | -0.595209016 | 0.012887152 | young | BC |
| UQCC3 | -0.595399725 | 0.008362082 | young | BC |
| PYURF | -0.595601582 | 0.005566328 | young | BC |
| AC007541.1 | -0.595620003 | 0.029933599 | young | BC |
| UQCR10 | -0.596382611 | 0.015009037 | young | BC |
| GSPT2 | -0.59886361 | 0.03697862 | young | BC |
| C16orf91 | -0.599119268 | 0.020115935 | young | BC |
| KRT10 | -0.599150401 | 0.023230681 | young | BC |
| DNAJC30 | -0.600531581 | 0.0450457 | young | BC |
| SF3B5 | -0.600628511 | 0.027272363 | young | BC |
| PNO1 | -0.601035896 | 0.016663663 | young | BC |
| MRPS22 | -0.601965441 | 0.008499835 | young | BC |
| MRPL55 | -0.602356713 | 0.014662328 | young | BC |
| TIMM17B | -0.604417053 | 0.022962832 | young | BC |
| LSM1 | -0.606549276 | 0.013554406 | young | BC |
| MICOS13 | -0.60698455 | 0.027272363 | young | BC |
| PRDX2 | -0.607175982 | 0.0041238 | young | BC |
| UNC50 | -0.607314181 | 0.017479511 | young | BC |
| C12orf10 | -0.609642627 | 0.022939411 | young | BC |
| TCEAL6 | -0.610396142 | 0.007136936 | young | BC |
| TXNDC17 | -0.61056431 | 0.043575833 | young | BC |
| C19orf81 | -0.611180337 | 0.032853826 | young | BC |

| | | | | |
|---|---|---|---|---|
| C15orf61 | -0.611957627 | 0.046809766 | young | BC |
| HPF1 | -0.612363232 | 0.002160912 | young | BC |
| MT-CYB | -0.612965986 | 0.048765422 | young | BC |
| PARK7 | -0.613416222 | 0.015482107 | young | BC |
| DUSP26 | -0.613637194 | 0.039018783 | young | BC |
| PET117 | -0.6141643 | 0.023230681 | young | BC |
| ATXN7L3B | -0.614608281 | 0.01355416 | young | BC |
| EID2 | -0.614832392 | 0.026047022 | young | BC |
| CCT6A | -0.615002625 | 0.019188594 | young | BC |
| AC104117.3 | -0.615454667 | 0.046601162 | young | BC |
| ABHD14A | -0.615873399 | 0.018044065 | young | BC |
| ANAPC15 | -0.617642548 | 0.014803268 | young | BC |
| FAM50A | -0.617901875 | 0.041328293 | young | BC |
| NIF3L1 | -0.617923552 | 0.045588967 | young | BC |
| TMEM70 | -0.61816656 | 0.003556621 | young | BC |
| EIF6 | -0.618578163 | 0.028339189 | young | BC |
| LAGE3 | -0.618695448 | 0.013554406 | young | BC |
| ZNHIT1 | -0.61885761 | 0.003349244 | young | BC |
| MRPL2 | -0.619022505 | 0.013373354 | young | BC |
| EEF1E1 | -0.619728485 | 0.005052674 | young | BC |
| CDKN2AIPNL | -0.619899164 | 0.002889412 | young | BC |
| PTPMT1 | -0.619986986 | 0.032598113 | young | BC |
| LEO1 | -0.620594479 | 0.003047145 | young | BC |
| ZCRB1 | -0.620751132 | 0.00089785 | young | BC |
| MRPS24 | -0.620855066 | 0.021168507 | young | BC |
| U2AF1L4 | -0.621503184 | 0.038748363 | young | BC |
| ANAPC11 | -0.622234763 | 0.007068148 | young | BC |
| TEX30 | -0.622235146 | 0.043302262 | young | BC |
| UBL5 | -0.622254771 | 0.025589147 | young | BC |
| NDUFA7 | -0.623018563 | 0.021143538 | young | BC |
| MRPS23 | -0.623166493 | 0.015886376 | young | BC |
| ZNF688 | -0.623532468 | 0.043302262 | young | BC |
| TMEM14C | -0.623610329 | 0.0196755 | young | BC |
| RD3 | -0.624468145 | 0.00034178 | young | BC |
| LSM10 | -0.624975734 | 0.011087279 | young | BC |
| LRTM1 | -0.625286906 | 0.025299434 | young | BC |
| C1orf122 | -0.62641203 | 0.007212871 | young | BC |
| CNPY2 | -0.626763979 | 0.012766737 | young | BC |
| FAM50B | -0.627262826 | 0.040752239 | young | BC |
| SNHG9 | -0.627324452 | 0.017560388 | young | BC |
| SELENOS | -0.627592846 | 0.011177763 | young | BC |
| LINC01750 | -0.627645049 | 0.032031834 | young | BC |
| ATP5IF1 | -0.62784974 | 0.019960191 | young | BC |
| EIF3I | -0.628211743 | 0.012281654 | young | BC |
| SIVA1 | -0.628369945 | 0.025752085 | young | BC |
| ATP5F1E | -0.628703973 | 0.00772218 | young | BC |
| BEX2 | -0.628747257 | 0.024378692 | young | BC |
| FAM32A | -0.629730037 | 0.012059792 | young | BC |
| GRHPR | -0.630107206 | 0.001862448 | young | BC |
| MRPL33 | -0.63045569 | 0.013533764 | young | BC |
| GET3 | -0.630514678 | 0.046444481 | young | BC |
| COX7A2 | -0.630603616 | 0.026010233 | young | BC |
| PSENEN | -0.630664068 | 0.014662328 | young | BC |
| PEBP1 | -0.631516907 | 0.025752085 | young | BC |

| | | | | |
|---|---|---|---|---|
| AP2S1 | -0.631867013 | 0.006024874 | young | BC |
| COX7C | -0.632252193 | 0.027150955 | young | BC |
| FIS1 | -0.632772491 | 0.010963537 | young | BC |
| CISD3 | -0.632917622 | 0.016653923 | young | BC |
| NAA38 | -0.633040512 | 0.002160912 | young | BC |
| UBB | -0.633218914 | 0.006965341 | young | BC |
| C8orf82 | -0.633510878 | 0.022866923 | young | BC |
| DDX24 | -0.633604937 | 0.004189796 | young | BC |
| GGCT | -0.633652445 | 0.01542006 | young | BC |
| TMEM160 | -0.633654697 | 0.030232644 | young | BC |
| C14orf119 | -0.633830462 | 0.012059792 | young | BC |
| DGCR6L | -0.634647207 | 0.012281654 | young | BC |
| PSMC1 | -0.634732635 | 0.023437767 | young | BC |
| RPL7L1 | -0.63515151 | 0.002665543 | young | BC |
| NDUFB8 | -0.635793293 | 0.01031208 | young | BC |
| MSRB2 | -0.637226634 | 0.012281654 | young | BC |
| TSR3 | -0.637245645 | 0.018971654 | young | BC |
| UFC1 | -0.637500074 | 0.003556621 | young | BC |
| LYSMD2 | -0.637687274 | 0.005023154 | young | BC |
| COPRS | -0.63779275 | 0.034706673 | young | BC |
| EBAG9 | -0.640389659 | 0.003815374 | young | BC |
| MRPS7 | -0.641715039 | 0.013163528 | young | BC |
| MAGOH | -0.641727833 | 0.011012232 | young | BC |
| MRPL47 | -0.641874314 | 0.00794191 | young | BC |
| POP5 | -0.642502614 | 0.041830056 | young | BC |
| PAXX | -0.642749777 | 0.001169105 | young | BC |
| EIF3K | -0.643324413 | 0.005468374 | young | BC |
| MRPL41 | -0.643501536 | 0.007550735 | young | BC |
| SLC25A5 | -0.643933524 | 0.0450457 | young | BC |
| TUSC1 | -0.644402164 | 0.001741148 | young | BC |
| R3HCC1 | -0.644530717 | 0.025774372 | young | BC |
| ZNF830 | -0.645107602 | 0.007943723 | young | BC |
| PET100 | -0.64539422 | 0.012281654 | young | BC |
| WDR83 | -0.645764154 | 0.030290832 | young | BC |
| PHAX | -0.646951691 | 0.002640247 | young | BC |
| STUB1 | -0.647022182 | 0.011460566 | young | BC |
| NOP10 | -0.648203898 | 0.038020794 | young | BC |
| YIF1A | -0.64838448 | 0.024875113 | young | BC |
| DNAJC19 | -0.648673422 | 0.005629377 | young | BC |
| DPM3 | -0.648788596 | 0.016143596 | young | BC |
| TPRKB | -0.649624632 | 0.004514214 | young | BC |
| TRPT1 | -0.650094988 | 0.037894563 | young | BC |
| AL137017.1 | -0.650223616 | 0.030330314 | young | BC |
| PEX16 | -0.650382984 | 0.040028245 | young | BC |
| COPS5 | -0.651547949 | 0.002285319 | young | BC |
| PFDN6 | -0.651678812 | 0.026634838 | young | BC |
| TOMM5 | -0.651788437 | 0.024194239 | young | BC |
| TRAPPC2B | -0.651850225 | 0.013129109 | young | BC |
| COX6B1 | -0.65299252 | 0.003453041 | young | BC |
| BUD31 | -0.655182623 | 0.002010236 | young | BC |
| GTF2A2 | -0.655654351 | 0.009433672 | young | BC |
| CCDC28B | -0.657289853 | 0.035637147 | young | BC |
| PSMC5 | -0.657316253 | 0.009433672 | young | BC |
| VPS28 | -0.657506184 | 0.006678554 | young | BC |

| | | | | |
|---|---|---|---|---|
| TXNDC9 | -0.657605064 | 0.004597172 | young | BC |
| YEATS4 | -0.657646113 | 0.006923753 | young | BC |
| AP000997.3 | -0.658420063 | 0.032775465 | young | BC |
| COX6C | -0.658466033 | 0.012256396 | young | BC |
| PIGP | -0.658487345 | 0.008655572 | young | BC |
| LINC00632 | -0.658836201 | 0.001564811 | young | BC |
| SNRNP25 | -0.65897885 | 0.016804207 | young | BC |
| GLRX5 | -0.659325458 | 0.006021858 | young | BC |
| FKBP3 | -0.659432199 | 0.001248225 | young | BC |
| ZNF511 | -0.659743551 | 0.009891541 | young | BC |
| CCDC43 | -0.66001523 | 0.00354876 | young | BC |
| ATP5F1D | -0.660353213 | 0.023945216 | young | BC |
| CLEC11A | -0.660601394 | 0.001169105 | young | BC |
| POLR2K | -0.660979785 | 0.007136936 | young | BC |
| SURF1 | -0.661579005 | 0.020728579 | young | BC |
| NDUFA1 | -0.661884068 | 0.005629377 | young | BC |
| COA3 | -0.662086986 | 0.00417699 | young | BC |
| TCEAL2 | -0.662670592 | 0.003269694 | young | BC |
| TPGS1 | -0.664866612 | 0.039437493 | young | BC |
| RDH14 | -0.666310302 | 0.000144994 | young | BC |
| JAGN1 | -0.6664467 | 0.00772741 | young | BC |
| C18orf32 | -0.666588043 | 0.001213774 | young | BC |
| THUMPD1 | -0.668605746 | 0.002122482 | young | BC |
| POP4 | -0.668956475 | 0.017159872 | young | BC |
| NDUFS3 | -0.670104034 | 0.013158526 | young | BC |
| RNF220 | -0.671112899 | 0.006678554 | young | BC |
| BNIP1 | -0.671671649 | 0.022627224 | young | BC |
| C18orf21 | -0.674016456 | 0.005468374 | young | BC |
| NDUFAF4 | -0.674082759 | 0.00492339 | young | BC |
| REX1BD | -0.674885456 | 0.001893064 | young | BC |
| NOL7 | -0.675225772 | 0.002160912 | young | BC |
| NDUFA8 | -0.675532788 | 0.001509784 | young | BC |
| CD2BP2 | -0.676365071 | 0.036049957 | young | BC |
| PBDC1 | -0.676374411 | 0.003415572 | young | BC |
| ZDHHC4 | -0.676462203 | 0.011497988 | young | BC |
| TIMM8B | -0.677223299 | 0.003253915 | young | BC |
| ISCA2 | -0.677267925 | 0.009118887 | young | BC |
| CAMK2B | -0.678195207 | 0.000114137 | young | BC |
| NDUFA13 | -0.678642413 | 0.000299789 | young | BC |
| ISCU | -0.679894263 | 0.013022041 | young | BC |
| ANKRD39 | -0.680210076 | 0.00786318 | young | BC |
| NOL12 | -0.680356147 | 0.00772741 | young | BC |
| SAP18 | -0.680859588 | 0.001347115 | young | BC |
| BLOC1S1 | -0.681181794 | 0.006319579 | young | BC |
| TMEM203 | -0.681333571 | 0.004463005 | young | BC |
| LAMTOR4 | -0.681431754 | 0.000129169 | young | BC |
| MRFAP1 | -0.682915231 | 0.000179866 | young | BC |
| PAQR9-AS1 | -0.683236758 | 0.006678554 | young | BC |
| SNRNP35 | -0.684506917 | 0.005052674 | young | BC |
| PIGBOS1 | -0.684560353 | 0.015482107 | young | BC |
| MYL12A | -0.684812801 | 0.004224833 | young | BC |
| CHCHD1 | -0.68547888 | 0.004546221 | young | BC |
| GCSH | -0.686497627 | 0.002010236 | young | BC |
| SMIM10L1 | -0.68767477 | 0.002160912 | young | BC |

| | | | | |
|---|---|---|---|---|
| RPL26L1 | -0.687835376 | 0.003952329 | young | BC |
| TRMT6 | -0.689569136 | 0.013129109 | young | BC |
| POLR3K | -0.690426337 | 0.024841725 | young | BC |
| NAGK | -0.691064634 | 0.005078024 | young | BC |
| HINT2 | -0.691579981 | 0.017091939 | young | BC |
| NT5C | -0.691988835 | 0.028961742 | young | BC |
| NDUFB4 | -0.692974481 | 0.00095508 | young | BC |
| NDUFB3 | -0.693842245 | 0.003556621 | young | BC |
| C17orf75 | -0.693920363 | 0.00251155 | young | BC |
| MRPL52 | -0.694593968 | 0.005468374 | young | BC |
| COX8A | -0.6973138 | 0.005052674 | young | BC |
| PSMA7 | -0.69825973 | 0.000201145 | young | BC |
| NDUFC2 | -0.698295001 | 0.002895764 | young | BC |
| RRS1 | -0.698303566 | 0.025299434 | young | BC |
| CST3 | -0.698984304 | 0.013163528 | young | BC |
| FMC1 | -0.69951277 | 0.015292341 | young | BC |
| RAB5IF | -0.699999063 | 0.003979162 | young | BC |
| AC005746.2 | -0.700206134 | 0.022239169 | young | BC |
| SBDS | -0.700652438 | 0.000120221 | young | BC |
| TMEM147 | -0.700658004 | 0.007218599 | young | BC |
| COX17 | -0.700728682 | 0.007212871 | young | BC |
| RD3L | -0.700748339 | 0.014077963 | young | BC |
| H3F3A | -0.702067048 | 0.002184547 | young | BC |
| CCDC124 | -0.702117376 | 0.007212871 | young | BC |
| TIMM13 | -0.702633549 | 0.002285319 | young | BC |
| NDUFS8 | -0.703264563 | 0.007028626 | young | BC |
| CYHR1 | -0.703740866 | 0.000114137 | young | BC |
| EBNA1BP2 | -0.704207047 | 0.00772218 | young | BC |
| TBCC | -0.704384725 | 0.00786318 | young | BC |
| UQCC2 | -0.704958708 | 0.00398317 | young | BC |
| COPS9 | -0.705777151 | 0.013129109 | young | BC |
| MRPS26 | -0.705986314 | 0.009891541 | young | BC |
| NDUFB10 | -0.706031264 | 0.007212871 | young | BC |
| ZNF593 | -0.706230453 | 0.038888456 | young | BC |
| COMMD5 | -0.706467253 | 0.025299434 | young | BC |
| POLR2J | -0.708449451 | 0.002874714 | young | BC |
| ECHS1 | -0.709385503 | 0.002041224 | young | BC |
| GSTO1 | -0.710108274 | 0.002895764 | young | BC |
| KRT222 | -0.710971597 | 0.008124663 | young | BC |
| BOLA3 | -0.711104456 | 0.013129109 | young | BC |
| MRPL11 | -0.711277297 | 0.014662328 | young | BC |
| THNSL2 | -0.711498722 | 0.009118887 | young | BC |
| NDUFS5 | -0.711528289 | 0.006319579 | young | BC |
| BEX1 | -0.713588916 | 0.00772741 | young | BC |
| MAPKAPK5-AS1 | -0.714548289 | 0.001466218 | young | BC |
| NCL | -0.714578119 | 0.011278507 | young | BC |
| LARP7 | -0.714800362 | 0.002010236 | young | BC |
| TRAPPC4 | -0.715166513 | 0.00674413 | young | BC |
| NOP56 | -0.716933325 | 0.009060748 | young | BC |
| BLOC1S4 | -0.717539879 | 2.26E-05 | young | BC |
| PGM5P4-AS1 | -0.71754253 | 0.04318227 | young | BC |
| CKLF | -0.718184622 | 0.032426365 | young | BC |
| SEC11C | -0.71840641 | 0.000129169 | young | BC |
| SDF2 | -0.718800097 | 0.001527741 | young | BC |

| | | | | |
|---|---|---|---|---|
| TAFA3 | -0.720628066 | 0.012607305 | young | BC |
| BAG1 | -0.721074945 | 0.00564882 | young | BC |
| CALM1 | -0.722420705 | 0.007938121 | young | BC |
| NDUFA4 | -0.722914466 | 0.012607305 | young | BC |
| TMEM215 | -0.723068247 | 0.005468374 | young | BC |
| PIN1 | -0.724309092 | 0.005468374 | young | BC |
| PAK1IP1 | -0.724489047 | 0.002895764 | young | BC |
| SNF8 | -0.724925235 | 0.006319579 | young | BC |
| SELENOW | -0.725828004 | 0.004489363 | young | BC |
| PRDX5 | -0.72599291 | 0.001502595 | young | BC |
| ATP5MC1 | -0.726897042 | 0.016147686 | young | BC |
| KRTCAP2 | -0.729594868 | 0.001527741 | young | BC |
| ATP5ME | -0.730837226 | 0.000167308 | young | BC |
| MDP1 | -0.732471718 | 0.015985366 | young | BC |
| TMEM251 | -0.732550347 | 0.002600096 | young | BC |
| OTUD6B-AS1 | -0.734501624 | 5.25E-05 | young | BC |
| ANTKMT | -0.734694923 | 0.001347115 | young | BC |
| ATP5MF | -0.735603833 | 0.000325995 | young | BC |
| SNHG30 | -0.735681584 | 0.002037441 | young | BC |
| APRT | -0.736423412 | 0.001527741 | young | BC |
| NDUFA3 | -0.736759838 | 0.00022566 | young | BC |
| TSTD1 | -0.736841997 | 0.006021858 | young | BC |
| MRPS12 | -0.73948175 | 0.002889412 | young | BC |
| ROMO1 | -0.740612627 | 0.002583794 | young | BC |
| ATP5PD | -0.740719035 | 1.15E-05 | young | BC |
| HIST3H2A | -0.743378397 | 0.021633699 | young | BC |
| CCDC85B | -0.743673617 | 0.001176783 | young | BC |
| RRP7A | -0.743771414 | 0.007803313 | young | BC |
| MRPL58 | -0.743957453 | 0.007212871 | young | BC |
| DNAJC15 | -0.745440749 | 0.013579074 | young | BC |
| PSMA4 | -0.746210874 | 0.003005574 | young | BC |
| PRELID1 | -0.747239526 | 0.002895764 | young | BC |
| VPS25 | -0.750862909 | 0.003309948 | young | BC |
| SLC39A3 | -0.751289533 | 0.016481101 | young | BC |
| NDUFAB1 | -0.752369921 | 0.002285319 | young | BC |
| MTRES1 | -0.752418614 | 0.001439926 | young | BC |
| CHTF18 | -0.753430085 | 0.012887152 | young | BC |
| ARL16 | -0.753471744 | 0.003867123 | young | BC |
| CHN2 | -0.754296737 | 0.006706793 | young | BC |
| MRPS33 | -0.755469769 | 0.001365472 | young | BC |
| THAP7 | -0.75555924 | 0.014662328 | young | BC |
| EDF1 | -0.756896089 | 0.00095508 | young | BC |
| HRAS | -0.760041717 | 0.00786318 | young | BC |
| SRSF3 | -0.762754242 | 0.002893128 | young | BC |
| MRTO4 | -0.764265424 | 0.013134833 | young | BC |
| HIGD2A | -0.764268885 | 0.003313967 | young | BC |
| MRPL21 | -0.765245771 | 0.00021313 | young | BC |
| LRRC24 | -0.766417136 | 0.024286835 | young | BC |
| JTB | -0.768407194 | 8.56E-05 | young | BC |
| NDUFA2 | -0.768706715 | 0.000264368 | young | BC |
| AL391650.1 | -0.768921391 | 0.002026145 | young | BC |
| PDHB | -0.773003591 | 0.003269694 | young | BC |
| SCAND1 | -0.774209808 | 0.000962744 | young | BC |
| MT-CO2 | -0.777395291 | 0.013129109 | young | BC |

| | | | | |
|---|---|---|---|---|
| CCT5 | -0.778165741 | 0.000114137 | young | BC |
| AURKAIP1 | -0.778710891 | 0.000510215 | young | BC |
| FAM222A-AS1 | -0.778752034 | 0.000993859 | young | BC |
| SNHG15 | -0.77967693 | 0.00072531 | young | BC |
| AL353759.1 | -0.781036739 | 0.018074014 | young | BC |
| ETFB | -0.782040888 | 0.002895764 | young | BC |
| SLC25A11 | -0.783029538 | 0.002895769 | young | BC |
| NDUFAF3 | -0.784283099 | 0.003149751 | young | BC |
| IMP3 | -0.788037747 | 0.002014262 | young | BC |
| CDR1 | -0.789882959 | 0.04958629 | young | BC |
| NDUFB1 | -0.790169493 | 0.002160912 | young | BC |
| MRPL36 | -0.790881755 | 0.000622572 | young | BC |
| HELLPAR | -0.792183084 | 0.000179866 | young | BC |
| MRPL27 | -0.792497944 | 0.002160912 | young | BC |
| MRPS34 | -0.794411645 | 0.002563044 | young | BC |
| HNRNPAB | -0.795199577 | 0.003288403 | young | BC |
| HHIP-AS1 | -0.79787938 | 0.043577473 | young | BC |
| MRPL54 | -0.798381798 | 0.000395499 | young | BC |
| LINC00672 | -0.79838401 | 0.006355745 | young | BC |
| MRPL34 | -0.799177555 | 0.002041224 | young | BC |
| NDUFB9 | -0.79982504 | 0.00072323 | young | BC |
| PSMB6 | -0.801106946 | 0.002085028 | young | BC |
| PAFAH1B3 | -0.801785094 | 0.000817273 | young | BC |
| EMC6 | -0.802318146 | 0.00272432 | young | BC |
| RNASEK | -0.805911111 | 0.002640247 | young | BC |
| TMEM44-AS1 | -0.808320227 | 0.012140708 | young | BC |
| NDUFB6 | -0.811773171 | 1.96E-07 | young | BC |
| GADD45GIP1 | -0.815726325 | 0.000297759 | young | BC |
| TRMT10C | -0.816343947 | 0.000112102 | young | BC |
| ALKBH7 | -0.817782912 | 0.000306291 | young | BC |
| BORCS8 | -0.819499048 | 0.005052674 | young | BC |
| NDUFB7 | -0.819716081 | 0.000144994 | young | BC |
| FDX2 | -0.820433219 | 0.001448009 | young | BC |
| PGP | -0.822038719 | 0.003349244 | young | BC |
| NDUFB11 | -0.822924753 | 0.00021313 | young | BC |
| TMEM141 | -0.82461363 | 0.014945557 | young | BC |
| MRPL32 | -0.826912285 | 0.000297988 | young | BC |
| PDCL3 | -0.827039068 | 0.005418813 | young | BC |
| GTF3C6 | -0.830853556 | 0.000192375 | young | BC |
| NDUFB2 | -0.832480947 | 0.001073339 | young | BC |
| MRM2 | -0.833148245 | 0.001065392 | young | BC |
| TTYH1 | -0.83419394 | 0.028706133 | young | BC |
| MRPL16 | -0.837031755 | 0.001599798 | young | BC |
| TIMM10 | -0.837498588 | 0.000263986 | young | BC |
| EFCAB2 | -0.839068357 | 0.001893064 | young | BC |
| NDUFS6 | -0.839739985 | 0.000201145 | young | BC |
| EID1 | -0.846473823 | 0.000962744 | young | BC |
| HIST1H2AC | -0.85089419 | 0.004104338 | young | BC |
| MRPS36 | -0.854229836 | 1.96E-07 | young | BC |
| FIBP | -0.859239281 | 0.001125047 | young | BC |
| USE1 | -0.86033448 | 0.000114137 | young | BC |
| MRPL15 | -0.860469378 | 0.000718901 | young | BC |
| ASPDH | -0.86146167 | 0.010911005 | young | BC |
| MRPL46 | -0.864395719 | 0.000306291 | young | BC |

| | | | | |
|---|---|---|---|---|
| POLR2L | -0.864717737 | 0.000817273 | young | BC |
| POP7 | -0.86638848 | 0.000114137 | young | BC |
| TMEM126A | -0.874073017 | 0.000559381 | young | BC |
| CXCL14 | -0.881009122 | 0.003313967 | young | BC |
| NAXE | -0.885610447 | 0.00043431 | young | BC |
| NCBP2AS2 | -0.891153282 | 0.000192333 | young | BC |
| NME1 | -0.898337844 | 3.57E-05 | young | BC |
| TRIAP1 | -0.899597927 | 0.000325995 | young | BC |
| PHB | -0.90750365 | 0.000112102 | young | BC |
| MRPL57 | -0.911236802 | 0.000104658 | young | BC |
| LENG1 | -0.911866469 | 0.000306291 | young | BC |
| TRAPPC5 | -0.914648863 | 1.23E-06 | young | BC |
| PCP2 | -0.918637439 | 0.002793481 | young | BC |
| PGLS | -0.919395807 | 0.000104658 | young | BC |
| CD320 | -0.920512968 | 0.000279593 | young | BC |
| NDUFA11 | -0.939847885 | 5.36E-08 | young | BC |
| TMEM208 | -0.940541836 | 5.41E-05 | young | BC |
| COA6 | -0.957897983 | 5.41E-05 | young | BC |
| RNASEH2C | -0.962575871 | 0.000849609 | young | BC |
| MRPS15 | -0.967503488 | 0.000168641 | young | BC |
| MRPS17 | -1.037315964 | 0.000299789 | young | BC |
| SMIM18 | -1.050796802 | 1.38E-05 | young | BC |
| MED11 | -1.134176136 | 1.23E-06 | young | BC |
| PGAM2 | -1.1547215 | 0.00022566 | young | BC |
| FAM138E | -1.287861222 | 1.23E-06 | young | BC |
| LINC02649 | 1.260087457 | 0.041782917 | old | Cone |
| NXPH4 | 1.238239224 | 0.046764874 | old | Cone |
| MAMDC2-AS1 | 1.175997153 | 0.041782917 | old | Cone |
| AL390957.1 | 1.133350996 | 0.041782917 | old | Cone |
| FMN2 | 1.05684405 | 0.041782917 | old | Cone |
| AP3B1 | 0.796909616 | 0.041782917 | old | Cone |
| MIR4422HG | 0.017156347 | 0.041782917 | old | Cone |
| IBA57-DT | 0.008073585 | 0.041782917 | old | Cone |
| AC021613.1 | 0.007859931 | 0.041782917 | old | Cone |
| IRF6 | -0.000273693 | 0.048150587 | young | Cone |
| BX255923.1 | -0.012188389 | 0.041782917 | young | Cone |
| FAM240C | -0.05732412 | 0.000380887 | young | Cone |
| S100A8 | -0.059485562 | 0.041782917 | young | Cone |
| NXNL1 | -0.918191357 | 0.041782917 | young | Cone |
| MED11 | -0.919513799 | 0.049421202 | young | Cone |
| AC084116.3 | -1.040885663 | 0.041782917 | young | Cone |
| DUSP6 | -1.265247957 | 0.005548921 | young | Cone |
| MDGA2 | 1.881641359 | 0.000361159 | old | HC |
| EGLN3 | 1.850789863 | 0.000197476 | old | HC |
| PFKFB4 | 1.709509952 | 0.000561488 | old | HC |
| VEGFA | 1.611214292 | 0.002266505 | old | HC |
| BHLHE40 | 1.541866644 | 0.000197476 | old | HC |
| DTNA | 1.523357117 | 0.00168535 | old | HC |
| OLFM3 | 1.497861681 | 0.0130934 | old | HC |
| SYCP2L | 1.430591654 | 0.013234321 | old | HC |
| KAZN | 1.373965457 | 0.001247318 | old | HC |
| HK2 | 1.365941355 | 0.027037602 | old | HC |
| ATF5 | 1.361543164 | 0.009735445 | old | HC |
| HIF1A-AS3 | 1.336681804 | 0.037228569 | old | HC |

| | | | | |
|---|---|---|---|---|
| RAPGEF4 | 1.332672653 | 0.020787803 | old | HC |
| AK4 | 1.328477816 | 0.013724383 | old | HC |
| DDIT4 | 1.309550325 | 0.020550398 | old | HC |
| KCNIP4 | 1.302929767 | 0.01398911 | old | HC |
| ERO1A | 1.287515547 | 0.004935046 | old | HC |
| SH3D21 | 1.27633573 | 0.041988299 | old | HC |
| C21orf62-AS1 | 1.258111768 | 0.032200288 | old | HC |
| PFKFB3 | 1.239994217 | 0.037228569 | old | HC |
| NXPH4 | 1.234507927 | 0.038666211 | old | HC |
| 1-Mar | 1.219431332 | 0.02008301 | old | HC |
| NEAT1 | 1.20933532 | 0.005897451 | old | HC |
| HIP1R | 1.199652543 | 0.031355641 | old | HC |
| CCNG2 | 1.193580495 | 0.021888192 | old | HC |
| DCLK1 | 1.191158603 | 0.004935046 | old | HC |
| MARCKSL1 | 1.188761026 | 0.013234321 | old | HC |
| DGCR9 | 1.169640693 | 0.032976623 | old | HC |
| MEG3 | 1.163637771 | 0.047855361 | old | HC |
| GRM7 | 1.137185367 | 0.04041591 | old | HC |
| PFKP | 1.133703426 | 0.019857737 | old | HC |
| YEATS2 | 1.123541035 | 0.007888316 | old | HC |
| MTERF1 | 1.122725487 | 0.003308029 | old | HC |
| TPD52 | 1.107113912 | 0.013234321 | old | HC |
| CACNA1D | 1.093150833 | 0.015364822 | old | HC |
| BNIP3L | 1.042779377 | 0.007776103 | old | HC |
| RNF165 | 1.035257412 | 0.009735445 | old | HC |
| STMN4 | 1.021275742 | 0.000544341 | old | HC |
| ZNF141 | 1.012067288 | 0.039417301 | old | HC |
| RHOBTB3 | 1.00831632 | 0.047587738 | old | HC |
| GLCE | 0.987932071 | 0.041988299 | old | HC |
| PDE9A | 0.968924737 | 0.025602826 | old | HC |
| PARP8 | 0.966401968 | 0.03127137 | old | HC |
| CADPS | 0.965404727 | 0.03169119 | old | HC |
| GTF2IRD1 | 0.950913562 | 0.021888192 | old | HC |
| NPAS2 | 0.935339656 | 0.049563567 | old | HC |
| TMEM87B | 0.92309594 | 0.047855361 | old | HC |
| AC025159.1 | 0.907116558 | 0.047587738 | old | HC |
| ESYT2 | 0.905952002 | 0.043735944 | old | HC |
| PRMT2 | 0.896701862 | 0.047855361 | old | HC |
| WSB1 | 0.896700076 | 0.036403432 | old | HC |
| TNIK | 0.894536878 | 0.041988299 | old | HC |
| SNAP25 | 0.885526008 | 0.013234321 | old | HC |
| RORA | 0.880264209 | 0.012012098 | old | HC |
| OGA | 0.860217481 | 0.002266505 | old | HC |
| ZSWIM5 | 0.821178578 | 0.044392269 | old | HC |
| MAPT | 0.81691301 | 0.005220669 | old | HC |
| ASPH | 0.810376588 | 0.019681653 | old | HC |
| NCOR2 | 0.797832919 | 0.025497503 | old | HC |
| KIF5C | 0.793065661 | 0.034939625 | old | HC |
| BIRC6 | 0.735511781 | 0.0060872 | old | HC |
| NEDD4L | 0.72039497 | 0.03169119 | old | HC |
| CPEB3 | 0.681766026 | 0.025065528 | old | HC |
| RLF | 0.679951641 | 0.04989141 | old | HC |
| EEF1A1 | 0.64739355 | 0.047855361 | old | HC |
| SETD5 | 0.63754291 | 0.023207174 | old | HC |

| | | | | |
|---|---|---|---|---|
| RERE | 0.604524515 | 0.037228569 | old | HC |
| KMT2E | 0.537492626 | 0.023207174 | old | HC |
| SRSF9 | -0.567677228 | 0.037228569 | young | HC |
| MPC1 | -0.668988857 | 0.04958518 | young | HC |
| NDUFAB1 | -0.684035601 | 0.021888192 | young | HC |
| TMBIM4 | -0.694720183 | 0.012884681 | young | HC |
| ATP5F1D | -0.699617373 | 0.037228569 | young | HC |
| SEC11C | -0.69999768 | 0.01398911 | young | HC |
| TALDO1 | -0.700104551 | 0.04958518 | young | HC |
| NDUFB7 | -0.706197775 | 0.020550398 | young | HC |
| PSMB6 | -0.713587905 | 0.023207174 | young | HC |
| PRELID1 | -0.722602109 | 0.049936557 | young | HC |
| JTB | -0.727579228 | 0.012884681 | young | HC |
| MRPL57 | -0.746114522 | 0.044392269 | young | HC |
| HIGD2A | -0.752203637 | 0.01398911 | young | HC |
| MRPS23 | -0.75850313 | 0.032976623 | young | HC |
| NDUFB1 | -0.765073905 | 0.036403432 | young | HC |
| NT5C | -0.786024145 | 0.047953715 | young | HC |
| IFT27 | -0.816730842 | 0.049936557 | young | HC |
| AL391650.1 | -0.816971923 | 0.049936557 | young | HC |
| LYSMD2 | -0.851383119 | 0.021888192 | young | HC |
| FIS1 | -0.867029264 | 0.022532378 | young | HC |
| AURKAIP1 | -0.871901604 | 0.038172923 | young | HC |
| MDP1 | -0.887733231 | 0.04989141 | young | HC |
| SOSTDC1 | -0.888604771 | 0.038085424 | young | HC |
| KRTCAP2 | -0.889891925 | 0.00168535 | young | HC |
| FZD1 | -0.899818022 | 0.02008301 | young | HC |
| BLOC1S1 | -0.906966911 | 0.022532378 | young | HC |
| MED11 | -0.926684807 | 0.04989141 | young | HC |
| MRPL36 | -0.932684848 | 0.037228569 | young | HC |
| ACER3 | -0.94043094 | 0.007888316 | young | HC |
| PAXX | -0.946967846 | 0.047855361 | young | HC |
| NAA38 | -0.950817688 | 0.001764032 | young | HC |
| NDUFA11 | -1.019260657 | 0.009570831 | young | HC |
| GKAP1 | -1.024126442 | 0.004132736 | young | HC |
| MRPL58 | -1.027412635 | 0.026541746 | young | HC |
| TMEM208 | -1.039672872 | 0.012884681 | young | HC |
| AC011525.1 | -1.051285597 | 0.041602935 | young | HC |
| NDUFAF3 | -1.084164107 | 0.014167643 | young | HC |
| RNF220 | -1.08595476 | 0.045153297 | young | HC |
| KITLG | -1.102301777 | 0.049351851 | young | HC |
| GPC6 | -1.119579976 | 0.013923311 | young | HC |
| GJA9 | -1.125362279 | 0.017121316 | young | HC |
| AL022332.1 | -1.199849688 | 0.032200288 | young | HC |
| PLCXD3 | -1.219594895 | 0.045153297 | young | HC |
| NUDT16 | -1.246409678 | 0.000544341 | young | HC |
| LINC02740 | -1.254643869 | 0.02008301 | young | HC |
| LENG1 | -1.280402619 | 0.009313719 | young | HC |
| HIST1H1E | -1.29701079 | 0.011251373 | young | HC |
| CARD16 | -1.351132078 | 0.01398911 | young | HC |
| PTPRT | -1.368985205 | 0.023207174 | young | HC |
| LINC01090 | -1.485341747 | 0.005301976 | young | HC |
| MLLT3 | 1.58791593 | 0.02017883 | old | Microglia |
| TTR | 1.577414185 | 0.019661551 | old | Microglia |

| | | | | |
|---|---|---|---|---|
| NKG7 | 1.550442219 | 0.008750244 | old | Microglia |
| ADSSL1 | 1.423558473 | 0.049471739 | old | Microglia |
| LIMD1 | 1.388457672 | 0.008750244 | old | Microglia |
| TOGARAM1 | 1.334012031 | 0.02017883 | old | Microglia |
| GLIS3 | 1.326995925 | 0.03711317 | old | Microglia |
| KCNMA1 | 1.192302566 | 0.041311965 | old | Microglia |
| C3AR1 | -0.811242359 | 0.006138883 | young | Microglia |
| MRPS28 | -0.841915126 | 0.031150087 | young | Microglia |
| DUSP6 | -1.108590207 | 0.004948488 | young | Microglia |
| DBI | -1.245765355 | 0.031150087 | young | Microglia |
| OAS1 | -1.247227023 | 0.004948488 | young | Microglia |
| IFI44L | -1.342781938 | 0.008361074 | young | Microglia |
| BATF3 | -1.343766885 | 0.041311965 | young | Microglia |
| AC046195.1 | -1.370205283 | 0.03128925 | young | Microglia |
| CCDC200 | -1.414310302 | 0.012548985 | young | Microglia |
| IFI6 | -1.419388695 | 0.008750244 | young | Microglia |
| IFIT3 | -1.4344366 | 0.03128925 | young | Microglia |
| OPRM1 | -1.45664924 | 0.027825307 | young | Microglia |
| IPCEF1 | -1.458437189 | 0.041311965 | young | Microglia |
| CCL3L1 | -1.460221163 | 0.035476492 | young | Microglia |
| IFIT1 | -1.643855084 | 0.008361074 | young | Microglia |
| PLAC8 | -1.6569985 | 0.004754857 | young | Microglia |
| ISG15 | -1.687445767 | 0.001093511 | young | Microglia |
| AP000439.2 | -1.722746132 | 0.008750244 | young | Microglia |
| SELL | -1.864638779 | 0.002293434 | young | Microglia |
| CCL4L2 | -1.867446266 | 0.001564975 | young | Microglia |
| AC007391.3 | 1.830014377 | 0.006361166 | old | Rod |
| SMIM3 | 1.71017122 | 0.02055282 | old | Rod |
| BAG3 | 1.668488937 | 0.023060027 | old | Rod |
| PLD1 | 1.657510657 | 0.010235996 | old | Rod |
| TNFAIP8 | 1.652158639 | 0.03248328 | old | Rod |
| RBM47 | 1.624163261 | 0.038198376 | old | Rod |
| AL645933.3 | 1.569153578 | 0.040463156 | old | Rod |
| NXPH4 | 1.53927465 | 0.049654487 | old | Rod |
| TMPRSS2 | 1.529161779 | 0.03248328 | old | Rod |
| CAVIN1 | 1.49153298 | 0.025455646 | old | Rod |
| TTR | 1.456275606 | 0.023060027 | old | Rod |
| RXRG | 1.439929476 | 0.036281133 | old | Rod |
| GPX3 | 1.377820105 | 0.023060027 | old | Rod |
| FRZB | 1.240077546 | 0.023060027 | old | Rod |
| PLCB3 | 1.144173047 | 0.02055282 | old | Rod |
| ARHGEF10 | 1.075073523 | 0.023060027 | old | Rod |
| PBXIP1 | 1.02002431 | 0.000172335 | old | Rod |
| HPS1 | 0.89638934 | 0.036980547 | old | Rod |
| PHKG1 | 0.800375231 | 0.023060027 | old | Rod |
| COL3A1 | 0.236008433 | 9.83E-12 | old | Rod |
| MNS1 | -0.59785599 | 0.023060027 | young | Rod |
| AC008040.1 | -0.648195831 | 0.023060027 | young | Rod |
| IFIT5 | -0.849409745 | 0.032848882 | young | Rod |
| ASTE1 | -0.875108092 | 0.049654487 | young | Rod |
| CCDC173 | -0.917430594 | 0.023060027 | young | Rod |
| AC010327.5 | -0.921736066 | 0.049654487 | young | Rod |
| CCDC39 | -1.032771894 | 0.049654487 | young | Rod |
| RN7SL832P | -1.060707003 | 0.023060027 | young | Rod |

| | | | | |
|---|---|---|---|---|
| AC034139.1 | -1.113753598 | 0.03248328 | young | Rod |
| FAM71F1 | -1.336039745 | 0.022779115 | young | Rod |
