## Supplementary material for "Interpretable Aging Signatures in Human Retinal Cell Types Revealed by Single-Cell RNA Sequencing and Sparse Logistic Regression": Table S8

Table S8: Gene symbol nomenclature reference. Provides standardized full names for all gen
manuscript.

| Gene symbol | Full standardized name |
|---|---|
| ADM | Adrenomedullin |
| AQP4 | Aquaporin 4 |
| BAG3 | BAG cochaperone 3 |
| BCO2 | Beta-carotene oxygenase 2 |
| BHLHE40 | Basic helix-loop-helix family member e40 |
| BHLHE41 | Basic helix-loop-helix family member e41 |
| BNIP3 | BCL2 interacting protein 3 |
| CALM1 | Calmodulin 1 |
| CCL2 | C-C motif chemokine ligand 2 |
| CEBPB | CCAAT/enhancer binding protein beta |
| CHD1 | Chromodomain helicase DNA binding protein 1 |
| COL4A3 | Collagen type IV alpha 3 chain |
| CRB1 | Crumbs cell polarity complex component 1 |
| CRYAB | Crystallin alpha B |
| CXCL1 | C-X-C motif chemokine ligand 1 |
| CXCL2 | C-X-C motif chemokine ligand 2 |
| DCLK1 | Doublecortin like kinase 1 |
| DDIT3 | DNA damage inducible transcript 3 |
| DNAJB1 | DnaJ heat shock protein family (Hsp40) member B1 |
| EGR1 | Early growth response 1 |
| ELOVL4 | ELOVL fatty acid elongase 4 |
| ERO1A | Endoplasmic reticulum oxidoreductase 1 alpha |
| FOSL1 | FOS like 1, AP-1 transcription factor subunit. |
| FOSL2 | FOS like 2, AP-1 transcription factor subunit. |
| FTH1 | Ferritin heavy chain 1 |
| FTL | Ferritin light chain |
| FTX | FTX transcript, XIST regulator (non-protein coding) |
| FUS | FUS RNA binding protein |
| GFAP | Glial fibrillary acidic protein |
| GNAT1 | G protein subunit alpha transducin 1 |
| GPX3 | Glutathione peroxidase 3 |
| GRIA1 | Glutamate ionotropic receptor AMPA type subunit 1 |
| GRIA4 | Glutamate ionotropic receptor AMPA type subunit 4 |
| HILPDA | Hypoxia inducible lipid droplet associated |
| HLA-A | Major histocompatibility complex, class I, A |
| HLA-C | Major histocompatibility complex, class I, C |
| HLA-DRA | Major histocompatibility complex, class II, DR alpha |
| HMGCS1 | 3-hydroxy-3-methylglutaryl-CoA synthase 1 |
| HOXA5 | Homeobox A5 |
| HP | Haptoglobin |
| HSP90AA1 | Heat shock protein 90 alpha family class A member 1 |
| HSP90AB1 | Heat shock protein 90 alpha family class B member 1 |
| HSPA1A | Heat shock protein family A (Hsp70) member 1A |
| HSPE1 | Heat shock protein family E (Hsp10) member 1 |
| HSPH1 | Heat shock protein family H (Hsp110) member 1 |
| IRF1 | Interferon regulatory factor 1 |
| JUND | JunD proto-oncogene, AP-1 transcription factor subunit |
| KIF2A | Kinesin family member 2A |
| LDHA | Lactate dehydrogenase A |
| LOX | Lysyl oxidase |
| MAFF | MAF bZIP transcription factor F |
| MAGI2 | Membrane associated guanylate kinase, WW and PDZ domain containing 2 |
| MT1E | Metallothionein 1E |

| | |
|---|---|
| MT1X | Metallothionein 1X |
| MT2A | Metallothionein 2A |
| MXI1 | MAX interactor 1, dimerization protein |
| MITF | Melanocyte inducing transcription factor |
| MYO9A | Myosin IXA |
| NCL | Nucleolin |
| NEUROD1 | Neuronal differentiation 1 |
| NFIA | Nuclear factor I A |
| NFKB1 | Nuclear factor kappa B subunit 1 |
| NLN | Neurolysin (metallopeptidase M3 family) |
| ONECUT1 | One cut homeobox 1 |
| P4HA1 | Prolyl 4-hydroxylase subunit alpha 1 |
| PDE4B | Phosphodiesterase 4B |
| PDE6B | Phosphodiesterase 6B |
| PDK1 | Pyruvate dehydrogenase kinase 1 |
| PFKFB4 | 6-phosphofructo-2-kinase/fructose-2,6-bisphosphatase 4 |
| PROX1 | Prospero homeobox 1 |
| RFXANK | Regulatory factor X associated ankyrin containing |
| RHO | Rhodopsin |
| RORA | RAR related orphan receptor A |
| RPS26 | Ribosomal protein S26 |
| RTN3 | Reticulon 3 |
| S100A6 | S100 calcium binding protein A6 |
| S100A10 | S100 calcium binding protein A10 |
| SEPTIN4 | Septin 4 |
| SERPINE1 | Serpin family E member 1 |
| SLC3A2 | Solute carrier family 3 member 2 |
| SLPI | Secretory leukocyte peptidase inhibitor |
| SNHG14 | Small nucleolar RNA host gene 14 |
| SREBF1 | Sterol regulatory element binding transcription factor 1 |
| STMN4 | Stathmin 4 |
| TCF4 | Transcription factor 4 |
| TF | Transferrin |
| THSD7B | Thrombospondin type 1 domain containing 7B |
| TMSB10 | Thymosin beta 10 |
| TTR | Transthyretin |
| TUBA1B | Tubulin alpha 1B |
| UBB | Ubiquitin B |
| UBC | Ubiquitin C |
| VEGFA | Vascular endothelial growth factor A |
| WIF1 | Wnt inhibitory factor 1 |
| WNT11 | Wnt family member 11 |
| WNT5B | Wnt family member 5B |
| XBP1 | X-box binding protein 1 |
| ZBTB21 | Zinc finger and BTB domain containing 21 |
| ZXDA | Zinc finger X-linked duplicated A |
| FZD3 | Frizzled class receptor 3 |
| ROM1 | Retinal outer segment membrane protein 1 |
| MT-ATP6 | Mitochondrially encoded ATP synthase membrane subunit 6 |
| MT-CO1 | Mitochondrially encoded cytochrome c oxidase I |
| MT-CO2 | Mitochondrially encoded cytochrome c oxidase II |
| MT-CO3 | Mitochondrially encoded cytochrome c oxidase III |
| MT-CYB | Mitochondrially encoded cytochrome b |
| MT-ND2 | Mitochondrially encoded NADH:ubiquinone oxidoreductase core subunit 2 |

| | |
|---|---|
| MT-ND3 | Mitochondrially encoded NADH:ubiquinone oxidoreductase core subunit 3 |
| MT-ND5 | Mitochondrially encoded NADH:ubiquinone oxidoreductase core subunit 5 |
| MT-ND6 | Mitochondrially encoded NADH:ubiquinone oxidoreductase core subunit 6 |
