## Supplementary material for "Interpretable Aging Signatures in Human Retinal Cell Types Revealed by Single-Cell RNA Sequencing and Sparse Logistic Regression": Table S9

Table S9: List of machine learning selected aging features for each cell type, ranked by importance.

| feature | cell_type |
|---|---|
| ZNF804A | AC |
| P4HA1 | AC |
| BNIP3 | AC |
| HMGCS1 | AC |
| PGK1 | AC |
| MT-ND6 | AC |
| NRXN1 | AC |
| AC002463.1 | AC |
| INSIG1 | AC |
| SSBP3 | AC |
| HLA-C | AC |
| VEGFA | AC |
| PGRMC1 | AC |
| GAPDH | AC |
| RNF165 | AC |
| UBE2B | AC |
| HLA-A | AC |
| H3F3B | AC |
| AC097534.2 | AC |
| NRG3 | AC |
| BEX1 | AC |
| MT-CO2 | AC |
| SLC2A3 | AC |
| SFPQ | AC |
| GPI | AC |
| STXBP5-AS1 | AC |
| NAPB | AC |
| MT-ND3 | AC |
| MT-ATP6 | AC |
| PREPL | AC |
| GABRG3-AS1 | AC |
| MAGI2 | AC |
| HTR5A | AC |
| SNCB | AC |
| VDAC1 | AC |
| ENO2 | AC |
| CACNA2D3 | AC |
| HSPA1A | AC |
| TCEAL7 | AC |
| AC096576.2 | AC |
| B2M | AC |
| PDK1 | AC |
| PKM | AC |
| PDE4A | AC |
| CRYAB | AC |
| SNAP25 | AC |
| COX7C | AC |
| CDH18 | AC |
| MAPT | AC |
| AC096576.3 | AC |
| DDX5 | AC |
| AL160254.1 | AC |
| AC024558.2 | AC |

| | |
|---|---|
| DYNLL2 | AC |
| KCNIP4 | AC |
| NDUFC1 | AC |
| FTX | AC |
| MT-CYB | AC |
| FAM155A | AC |
| DAAM1 | AC |
| H2AFY | AC |
| WSB1 | AC |
| INPP5F | AC |
| PAK3 | AC |
| UBC | AC |
| GRIK2 | AC |
| KBTBD2 | AC |
| C1QL1 | AC |
| PDE5A | AC |
| ATP5IF1 | AC |
| TIMP2 | AC |
| RTN3 | AC |
| LRCH2 | AC |
| HSP90AA1 | AC |
| MOCS2 | AC |
| ENO1 | AC |
| STX7 | AC |
| TCEAL6 | AC |
| CALM3 | AC |
| HS6ST3 | AC |
| NCDN | AC |
| LDHA | AC |
| PEG3 | AC |
| PPIA | AC |
| RTN4 | AC |
| CTSA | AC |
| NPEPPS | AC |
| HIST1H2BE | AC |
| LGALS1 | AC |
| PSMB8 | AC |
| BHLHE40 | AC |
| DAB1 | AC |
| NEDD4L | AC |
| GRIP2 | AC |
| PIKFYVE | AC |
| ZNF407 | AC |
| PDXP | AC |
| PRKAG2-AS1 | AC |
| IFIT1 | AC |
| ISCA1 | AC |
| FADS2 | AC |
| PFKL | AC |
| IKZF5 | AC |
| ARHGAP28 | AC |
| BTBD8 | AC |
| MT1X | Astrocyte |
| SLPI | Astrocyte |

| | |
|---|---|
| ANGPTL1 | Astrocyte |
| LDHA | Astrocyte |
| MT-CO2 | Astrocyte |
| HP | Astrocyte |
| SEMA6D | Astrocyte |
| MT-CO3 | Astrocyte |
| MT-ATP8 | Astrocyte |
| MT2A | Astrocyte |
| VEGFA | Astrocyte |
| ERO1A | Astrocyte |
| HILPDA | Astrocyte |
| TM7SF2 | Astrocyte |
| SELENOM | Astrocyte |
| MT-ND6 | BC |
| MT-CO2 | BC |
| MT-ND3 | BC |
| HSPA1A | BC |
| MT-CO3 | BC |
| MT-ND5 | BC |
| AC091938.1 | BC |
| NCL | BC |
| LRTM1 | BC |
| CDR1 | BC |
| H3F3A | BC |
| AC119868.2 | BC |
| MT-ATP6 | BC |
| CRYAB | BC |
| BHLHE40 | BC |
| AP001825.1 | BC |
| EID1 | BC |
| COX4I1 | BC |
| PCP2 | BC |
| ATP5MC1 | BC |
| NDUFAB1 | BC |
| TMSB4X | BC |
| GNG13 | BC |
| LINC02055 | BC |
| ANOS1 | BC |
| MT-CYB | BC |
| GNB3 | BC |
| HSPB1 | BC |
| P4HA1 | BC |
| KCNH5 | BC |
| NIF3L1 | BC |
| ATP6V0B | BC |
| CALM2 | BC |
| SEMA3E | BC |
| TRNP1 | BC |
| MT-ND2 | BC |
| NQO1 | BC |
| CNTN5 | BC |
| MT-ND1 | BC |
| HNRNPA2B1 | BC |
| CHCHD2 | BC |

| | |
|---|---|
| COX8A | BC |
| APOBEC2 | BC |
| AC092155.1 | BC |
| SRP14 | BC |
| CRY2 | BC |
| YBX1 | BC |
| PARK7 | BC |
| MT-ATP8 | BC |
| CA10 | BC |
| AC010478.1 | BC |
| MT-CO1 | BC |
| NME1 | BC |
| KRT222 | BC |
| SREBF2 | BC |
| LSM4 | BC |
| NDUFC2 | BC |
| ATP5F1C | BC |
| RPS19BP1 | BC |
| TTYH1 | BC |
| SNAP25-AS1 | BC |
| NDUFB4 | BC |
| CALM1 | BC |
| GRM5 | BC |
| MRPL57 | BC |
| CCDC136 | BC |
| C1QBP | BC |
| FKBP3 | BC |
| NTNG1 | BC |
| SAP18 | BC |
| C18orf32 | BC |
| SPCS1 | BC |
| LINC02649 | BC |
| CST3 | BC |
| TBCB | BC |
| PSMB6 | BC |
| SUMO2 | BC |
| PFKFB4 | BC |
| AC104117.3 | BC |
| ERH | BC |
| CLASP2 | BC |
| PEBP1 | BC |
| SET | BC |
| SH3GL3 | BC |
| CPD | BC |
| NDUFB9 | BC |
| COX7A2 | BC |
| SOD1 | BC |
| NDUFS6 | BC |
| PDK1 | BC |
| SRSF9 | BC |
| AL033504.1 | BC |
| NDUFB6 | BC |
| LRRTM4 | BC |
| NDUFB7 | BC |

| | |
|---|---|
| NDUFB1 | BC |
| TRPM1 | BC |
| COPS9 | BC |
| ATP5PF | BC |
| POLR2K | BC |
| DOK6 | BC |
| TF | BC |
| CXCL14 | BC |
| AK4 | BC |
| SDK1 | BC |
| MICOS10 | BC |
| STARD4 | BC |
| DTNA | BC |
| PPIA | BC |
| YWHAE | BC |
| ATP5MC3 | BC |
| ITM2C | BC |
| NDUFS5 | BC |
| RNASEK | BC |
| BEX1 | BC |
| CCNG2 | BC |
| FIS1 | BC |
| PRKCA-AS1 | BC |
| ZFHX2 | BC |
| PDZRN4 | BC |
| KMT2E | BC |
| CCNI | BC |
| NDUFA2 | BC |
| NLGN4X | BC |
| PSMB1 | BC |
| TAX1BP1 | BC |
| DDC | BC |
| AC117944.1 | BC |
| HSPA6 | BC |
| PNN | BC |
| RERE | BC |
| PSMB3 | BC |
| AP000857.2 | BC |
| SF3B5 | BC |
| PAIP2 | BC |
| ATP6V1G1 | BC |
| TUBA1B | BC |
| BHLHE41 | BC |
| PRDX5 | BC |
| TXN | BC |
| DST | BC |
| NPVF | BC |
| AC090825.1 | BC |
| GSTO1 | BC |
| TAFA3 | BC |
| COX6C | BC |
| AL513164.1 | BC |
| NEDD8 | BC |
| FMC1 | BC |

| | |
|---|---|
| CCT6A | BC |
| RPS17 | BC |
| COX6A1 | BC |
| CRYBG3 | BC |
| NDUFB11 | BC |
| NDUFA1 | BC |
| CACNA2D1 | BC |
| ASIC2 | BC |
| VBP1 | BC |
| C16orf74 | BC |
| FUS | BC |
| BTF3 | BC |
| SELENOW | BC |
| SLC6A6 | BC |
| VSTM2B | BC |
| PRKCA | BC |
| H2AFZ | BC |
| CCT5 | BC |
| FAM138C | BC |
| ANKS1B | BC |
| PSMA7 | BC |
| COX17 | BC |
| PNKD | BC |
| SLIRP | BC |
| SMDT1 | BC |
| TDRG1 | BC |
| CCT3 | BC |
| PSMA2 | BC |
| GPX3 | BC |
| MRFAP1 | BC |
| ANK2 | BC |
| TTR | Cone |
| LINGO2 | Cone |
| MT-ND6 | Cone |
| SLC35F1 | Cone |
| LINC02343 | Cone |
| UCMA | Cone |
| TMEM176B | Cone |
| MBP | Cone |
| HSPA1A | Cone |
| HSPA1B | Cone |
| CCSER1 | Cone |
| TMEM176A | Cone |
| CST3 | Cone |
| KIAA0825 | Cone |
| NAALADL2 | Cone |
| SGCD | Cone |
| PTPRD | Cone |
| EYS | Cone |
| RGS9 | Cone |
| KIF2A | Cone |
| VAX2 | Cone |
| GAS7 | Cone |
| AC007349.2 | Cone |

| | |
|---|---|
| THSD7B | Cone |
| MAGI2 | Cone |
| SSX2IP | Cone |
| PCP4 | Cone |
| AC116903.2 | Cone |
| WWOX | Cone |
| EEF1A2 | Cone |
| AC112206.2 | Cone |
| LINC00871 | Cone |
| CCNJL | Cone |
| RPS26 | Cone |
| MRLN | Cone |
| PDE5A | Cone |
| C1QTNF4 | Cone |
| AP000820.2 | Cone |
| LRFN5 | Cone |
| MAP3K7CL | Cone |
| GABRA2 | Cone |
| AL161716.1 | Cone |
| DRD4 | Cone |
| HLA-A | Cone |
| NCL | Cone |
| DLGAP2 | Cone |
| ST3GAL3 | Cone |
| PRUNE2 | Cone |
| SBSPON | Cone |
| AL357153.2 | Cone |
| HMGB2 | Cone |
| PRKDC | Cone |
| ARMC9 | Cone |
| NRN1L | Cone |
| SNAP91 | Cone |
| LRRFIP1 | Cone |
| BAG3 | Cone |
| USH2A | Cone |
| RRAD | Cone |
| AC117453.1 | Cone |
| CCNO | Cone |
| PRKN | Cone |
| H1FX | Cone |
| SLC38A1 | Cone |
| DPF3 | Cone |
| AC079467.1 | Cone |
| HSPH1 | Cone |
| BCO2 | Cone |
| PCAT1 | Cone |
| HNRNPH1 | Cone |
| AGBL4 | Cone |
| AL050403.2 | Cone |
| DLG2 | Cone |
| HPRT1 | Cone |
| ACTB | Cone |
| SYCE1L | Cone |
| PRR16 | Cone |

| | |
|---|---|
| COTL1 | Cone |
| SAMD7 | Cone |
| KLF7 | Cone |
| MIR2052HG | Cone |
| ATP2B4 | Cone |
| LGALS3BP | Cone |
| TF | Cone |
| PCAT4 | Cone |
| TMEM108 | Cone |
| PDE6C | Cone |
| MESP1 | Cone |
| AC137770.1 | Cone |
| ZBTB20 | Cone |
| MNDA | Cone |
| POLQ | Cone |
| AC110992.1 | Cone |
| AC106798.1 | Cone |
| RAB17 | Cone |
| RAX | Cone |
| PCDH15 | Cone |
| AKAP6 | Cone |
| C9orf16 | Cone |
| CRYAB | Cone |
| CREG2 | Cone |
| AC002460.2 | Cone |
| AHSA1 | Cone |
| NTM | Cone |
| TXK | Cone |
| SMYD3 | Cone |
| MAP2 | Cone |
| RCBTB1 | Cone |
| DSE | Cone |
| HLA-DRB5 | Cone |
| PEX5L-AS2 | Cone |
| AL137804.1 | Cone |
| ANTXR2 | Cone |
| STK24 | Cone |
| STK33 | Cone |
| ACTG1 | Cone |
| EFNA5 | Cone |
| SNHG14 | Cone |
| DMD | Cone |
| HAR1A | Cone |
| SLC22A17 | Cone |
| A2M | Cone |
| SOX2-OT | Cone |
| SLC4A8 | Cone |
| ME3 | Cone |
| UPK3BL1 | Cone |
| FAM135B | Cone |
| ADAMTSL1 | Cone |
| KCNB1 | Cone |
| LINC01184 | Cone |
| SDHAF3 | Cone |

| | |
|---|---|
| AL121821.2 | Cone |
| AL118516.1 | Cone |
| TCEAL9 | Cone |
| SNAP25-AS1 | Cone |
| PRSS51 | Cone |
| NUBPL | Cone |
| DNAJB1 | Cone |
| GLUL | Cone |
| FAM172A | Cone |
| MT2A | Cone |
| CLUL1 | Cone |
| WFDC11 | Cone |
| AC068051.1 | Cone |
| FSTL5 | Cone |
| TRAF3IP1 | Cone |
| CDH12 | Cone |
| IMMP2L | Cone |
| PGAM2 | Cone |
| FAM153A | Cone |
| AC092939.1 | Cone |
| TMEM141 | Cone |
| ZNF529 | Cone |
| IFITM3 | Cone |
| FTX | Cone |
| LDLRAD4 | Cone |
| EIF4G2 | Cone |
| FBXL17 | Cone |
| KIRREL1 | Cone |
| FGF12 | Cone |
| GNG12-AS1 | Cone |
| MAP1LC3A | Cone |
| LGALS3 | Cone |
| STPG2 | Cone |
| SNX29 | Cone |
| ZNF292 | Cone |
| AC007325.4 | Cone |
| GNGT1 | Cone |
| AC004540.2 | Cone |
| VPS53 | Cone |
| SERPINH1 | Cone |
| PLXDC1 | Cone |
| HLA-C | Cone |
| FAM166C | Cone |
| HSBP1 | Cone |
| SSBP4 | Cone |
| ROBO2 | Cone |
| STMN1 | Cone |
| EPCAM | Cone |
| ZNF225 | Cone |
| PRELID2 | Cone |
| POLH | Cone |
| BMPR1B | Cone |
| CADM1 | Cone |
| ROM1 | Cone |

| | |
|---|---|
| HCG17 | Cone |
| ARHGAP24 | Cone |
| DYRK4 | Cone |
| MFSD4B | Cone |
| SKAP2 | Cone |
| YPEL3 | Cone |
| WAKMAR2 | Cone |
| UPP2 | Cone |
| AL357172.1 | Cone |
| GPX3 | Cone |
| AC097662.1 | Cone |
| ZNF385D | HC |
| UBC | HC |
| HSPH1 | HC |
| GRIA4 | HC |
| MIR181A2HG | HC |
| NRG1 | HC |
| DOK6 | HC |
| GALNT13 | HC |
| KCNJ3 | HC |
| MT-ND6 | HC |
| FDFT1 | HC |
| AC093765.2 | HC |
| CRYAB | HC |
| AC092691.1 | HC |
| MDGA2 | HC |
| ADCY2 | HC |
| TAPT1-AS1 | HC |
| LHFPL6 | HC |
| TMSB10 | HC |
| BHLHE40 | HC |
| FUS | HC |
| SNTG2 | HC |
| STMN4 | HC |
| FRMPD4 | HC |
| PRR16 | HC |
| DTNA | HC |
| NDST3 | HC |
| PDE4B | HC |
| EPHA5 | HC |
| GPX3 | HC |
| SPP1 | HC |
| PLCB4 | HC |
| TUBA1B | HC |
| PGAM1 | HC |
| FHIT | HC |
| SNHG14 | HC |
| PCSK6 | HC |
| ENO2 | HC |
| TSPAN8 | HC |
| TRIM36 | HC |
| ARHGAP24 | HC |
| NCL | HC |
| MT2A | HC |

| | |
|---|---|
| MT3 | HC |
| CRPPA | HC |
| FAM162A | HC |
| FTX | HC |
| MARCKS | HC |
| SESTD1 | HC |
| DNAJB4 | HC |
| LRBA | HC |
| CCNG2 | HC |
| DOCK3 | HC |
| WDR37 | HC |
| MARCKSL1 | HC |
| RD3L | HC |
| GJA10 | HC |
| KMT5B | HC |
| HIST1H4C | HC |
| SEPTIN4 | HC |
| HSP90AB1 | HC |
| MAPRE2 | HC |
| PTN | HC |
| PTGFR | HC |
| ONECUT2 | HC |
| CUX2 | HC |
| TARS | HC |
| ELOVL5 | HC |
| SLC4A5 | HC |
| ADARB2 | HC |
| ERO1A | HC |
| NPIPB2 | HC |
| HSPE1 | HC |
| S100A10 | HC |
| PAM | HC |
| IER5L | HC |
| RALGAPA2 | HC |
| DCLK1 | HC |
| HMCES | HC |
| MT-ND6 | MGC |
| CRYAB | MGC |
| SOD2 | MGC |
| GADD45B | MGC |
| KHDRBS2 | MGC |
| GRID2 | MGC |
| DACH1 | MGC |
| JUNB | MGC |
| NAV3 | MGC |
| RARRES1 | MGC |
| GEM | MGC |
| UBC | MGC |
| MAFF | MGC |
| GDF15 | MGC |
| HLA-DRA | MGC |
| RORB | MGC |
| MT-CO2 | MGC |
| NAMPT | MGC |

| | |
|---|---|
| ABI3BP | MGC |
| MT-ND5 | MGC |
| GPX3 | MGC |
| MT-ATP8 | MGC |
| IFITM3 | MGC |
| KIAA1217 | MGC |
| HLA-A | MGC |
| DDIT3 | MGC |
| ID1 | MGC |
| LGALS3 | MGC |
| INHBA | MGC |
| NRXN3 | MGC |
| ILDR2 | MGC |
| LITAF | MGC |
| CD44 | MGC |
| PREX2 | MGC |
| ADGRL3 | MGC |
| CEBPB | MGC |
| FGFBP2 | MGC |
| C11orf96 | MGC |
| CDH2 | MGC |
| SPON1 | MGC |
| KLHL8 | MGC |
| GFAP | MGC |
| ID3 | MGC |
| HSPA1A | MGC |
| IFITM2 | MGC |
| SULF1 | MGC |
| SORBS2 | MGC |
| CD74 | MGC |
| LINC00461 | MGC |
| SOX6 | MGC |
| SERPING1 | MGC |
| GRIA4 | MGC |
| WTAP | MGC |
| MTTP | MGC |
| EPB41L2 | MGC |
| SOX5 | MGC |
| MYO10 | MGC |
| ATF3 | MGC |
| SIPA1L1 | MGC |
| KDR | MGC |
| YBX3 | MGC |
| PDLIM3 | MGC |
| MIR99AHG | MGC |
| FUT8 | MGC |
| CCDC107 | MGC |
| SPHK1 | MGC |
| CCN1 | MGC |
| AMER2 | MGC |
| CHST9 | MGC |
| ACTB | MGC |
| NFIA | MGC |
| JUND | MGC |

| | |
|---|---|
| PMEPA1 | MGC |
| MT2A | MGC |
| GPRC5A | MGC |
| SLC3A2 | MGC |
| HSPA1B | MGC |
| BHLHE40 | MGC |
| COL24A1 | MGC |
| CXCL2 | MGC |
| CPA6 | MGC |
| ZNF385D | MGC |
| ARL4D | MGC |
| CLRN1 | MGC |
| MT-ND3 | MGC |
| SYT11 | MGC |
| PCYT1B | MGC |
| GPM6A | MGC |
| AQP4-AS1 | MGC |
| ANGPTL1 | MGC |
| B2M | MGC |
| INSIG1 | MGC |
| PHYHIPL | MGC |
| DNAJB1 | MGC |
| LMCD1 | MGC |
| SDC4 | MGC |
| PMP22 | MGC |
| GRIA1 | MGC |
| PPP1R15A | MGC |
| ARHGAP21 | MGC |
| SERPINE1 | MGC |
| HLA-DRB1 | MGC |
| IER5L | MGC |
| LRRC4C | MGC |
| HECW2 | MGC |
| BHLHE41 | MGC |
| MT-ND4L | MGC |
| UTRN | MGC |
| SNHG16 | MGC |
| FBXO32 | MGC |
| MYO6 | MGC |
| ACTG1 | MGC |
| FTX | MGC |
| PTPRZ1 | MGC |
| SLCO4A1-AS1 | MGC |
| TFRC | MGC |
| MAGI2 | MGC |
| RHOU | MGC |
| PDLIM4 | MGC |
| S100A6 | MGC |
| ERO1A | MGC |
| PAX6 | MGC |
| IER3 | MGC |
| SERPINA3 | MGC |
| MKNK2 | MGC |
| ANXA1 | MGC |

| | |
|---|---|
| NRG3 | MGC |
| RBP1 | MGC |
| CHMP1B | MGC |
| COL4A3 | MGC |
| DUSP1 | MGC |
| WIF1 | MGC |
| ZNF331 | MGC |
| AMACR | MGC |
| MED12L | MGC |
| RORA | MGC |
| FOSL2 | MGC |
| HSPA6 | MGC |
| DYNC1I1 | MGC |
| PCSK2 | MGC |
| ADM | MGC |
| PIM1 | MGC |
| ZBTB21 | MGC |
| MAP2 | MGC |
| SPSB1 | MGC |
| CNTN1 | MGC |
| RPS26 | MGC |
| AL390957.1 | MGC |
| EIF1 | MGC |
| ANOS1 | MGC |
| CCL2 | MGC |
| RRAD | MGC |
| DENND11 | MGC |
| NR4A1 | MGC |
| KLKB1 | MGC |
| AC004264.1 | MGC |
| SLC19A2 | MGC |
| CYSTM1 | MGC |
| SRGAP3 | MGC |
| MEGF10 | MGC |
| SDS | MGC |
| RAB1A | MGC |
| S100A16 | MGC |
| NCKAP5 | MGC |
| MT-ND2 | MGC |
| EPCAM | MGC |
| MT-CO3 | MGC |
| SFXN5 | MGC |
| OSBPL10 | MGC |
| CXCL3 | MGC |
| CTTNBP2 | MGC |
| S100A10 | MGC |
| CALM1 | MGC |
| ID2 | MGC |
| SLC7A5 | MGC |
| COL4A4 | MGC |
| FAP | MGC |
| HSPE1 | MGC |
| KLHDC8A | MGC |
| AC092691.1 | MGC |

| | |
|---|---|
| EPHB6 | MGC |
| HLA-C | MGC |
| TUBB2B | MGC |
| BAIAP2 | MGC |
| HILPDA | MGC |
| NAALADL2 | MGC |
| TRH | MGC |
| ADD1 | MGC |
| ABCA8 | MGC |
| SCD | MGC |
| NFKBIA | MGC |
| CLDN1 | MGC |
| SFRP2 | MGC |
| PHLPP1 | MGC |
| AQP4 | MGC |
| COTL1 | MGC |
| MT-ATP6 | MGC |
| S100A11 | MGC |
| CYP4V2 | MGC |
| FTH1 | Microglia |
| MT2A | Microglia |
| GAPDH | Microglia |
| SLC35F1 | Microglia |
| IPCEF1 | Microglia |
| VIM | Microglia |
| ERO1A | Microglia |
| MIF | Microglia |
| CX3CR1 | Microglia |
| CCDC26 | Microglia |
| PLIN2 | Microglia |
| MT-ND6 | Microglia |
| APOC1 | Microglia |
| BASP1 | Microglia |
| ITPR2 | Microglia |
| SLC11A1 | Microglia |
| CD9 | Microglia |
| CDK6 | Microglia |
| IGSF6 | Microglia |
| GPNMB | Microglia |
| SPP1 | Microglia |
| SRGN | Microglia |
| EGR3 | Microglia |
| P4HA1 | Microglia |
| C4orf3 | Microglia |
| PLAUR | Microglia |
| ADAM8 | Microglia |
| SPRED1 | Microglia |
| LPCAT2 | Microglia |
| SLC16A10 | Microglia |
| PARP14 | Microglia |
| BNIP3L | Microglia |
| METRNL | Microglia |
| TBC1D5 | Microglia |
| ELMO1 | Microglia |

| | |
|---|---|
| HDAC9 | Microglia |
| NCKAP5 | Microglia |
| MT1X | Microglia |
| FTL | Microglia |
| TPI1 | Microglia |
| SAMHD1 | Microglia |
| MT3 | Microglia |
| BLNK | Microglia |
| SLC2A3 | Microglia |
| THBS1 | Microglia |
| RNASE4 | Microglia |
| SKAP2 | Microglia |
| FGF12 | Microglia |
| TUBA1B | Microglia |
| PGK1 | Microglia |
| ARL4C | Microglia |
| IFI16 | Microglia |
| ADM | Microglia |
| MS4A4E | Microglia |
| CSTB | Microglia |
| PMP22 | Microglia |
| CCL4L2 | Microglia |
| MT1G | Microglia |
| NR4A2 | Microglia |
| CYFIP1 | Microglia |
| FRMD4A | Microglia |
| JAK2 | Microglia |
| XAF1 | Microglia |
| CCL3L1 | Microglia |
| LGALS1 | Microglia |
| THADA | Microglia |
| 1-Mar | Microglia |
| GLUL | Microglia |
| MBNL1 | Microglia |
| QKI | Microglia |
| AC008574.1 | Microglia |
| CALM2 | Microglia |
| CH25H | Microglia |
| MT1E | Microglia |
| PFDN5 | Microglia |
| RGCC | Microglia |
| CCL2 | Microglia |
| ILDR1 | Microglia |
| ENSA | Microglia |
| TPT1 | Microglia |
| FPR3 | Microglia |
| EPSTI1 | Microglia |
| ZBTB20 | Microglia |
| FOSB | Microglia |
| CLEC2B | Microglia |
| APOE | Microglia |
| PSME2 | Microglia |
| IFI6 | Microglia |
| SHTN1 | Microglia |

| | |
|---|---|
| DOCK2 | Microglia |
| AL591518.1 | Microglia |
| NAV3 | Microglia |
| SLCO2B1 | Microglia |
| MGAT5 | Microglia |
| JDP2 | Microglia |
| SAMD9L | Microglia |
| FAU | Microglia |
| ENO1 | Microglia |
| CAMK1D | Microglia |
| BTG2 | Microglia |
| TBXAS1 | Microglia |
| ARHGAP15 | Microglia |
| BMP2K | Microglia |
| AC073352.2 | Microglia |
| IFI44L | Microglia |
| FCGR1B | Microglia |
| SOCS3 | Microglia |
| TNFRSF11B | Microglia |
| PLXDC2 | Microglia |
| HSPA1A | Rod |
| HSPA1B | Rod |
| HSP90AA1 | Rod |
| MT-ND6 | Rod |
| HSPH1 | Rod |
| EYS | Rod |
| SLC35F1 | Rod |
| METRNL | Rod |
| DNAJB1 | Rod |
| HMGB2 | Rod |
| HSPA6 | Rod |
| TRNP1 | Rod |
| CACYBP | Rod |
| A2M | Rod |
| MT-ND3 | Rod |
| CRYAB | Rod |
| PDE6G | Rod |
| DRD4 | Rod |
| TMEM176A | Rod |
| MT-ND5 | Rod |
| DMD | Rod |
| MT-CO2 | Rod |
| AL356804.1 | Rod |
| NTM | Rod |
| BAG3 | Rod |
| UBB | Rod |
| HSPD1 | Rod |
| EPHA3 | Rod |
| CADM1 | Rod |
| MT-ND2 | Rod |
| NLGN1 | Rod |
| FAM155A | Rod |
| MT-CO3 | Rod |
| NAALADL2 | Rod |

| | |
|---|---|
| EPB41L3 | Rod |
| WAPL | Rod |
| RALYL | Rod |
| NCL | Rod |
| HPRT1 | Rod |
| TMEM176B | Rod |
| OTX2-AS1 | Rod |
| AHSA1 | Rod |
| BMPR1B | Rod |
| HMGN1 | Rod |
| AC016723.1 | Rod |
| AL355835.1 | Rod |
| HSPB1 | Rod |
| RGS9 | Rod |
| PLCL2 | Rod |
| USH2A | Rod |
| POU2AF1 | Rod |
| AL392023.2 | Rod |
| LHFPL3 | Rod |
| PTPRM | Rod |
| MIAT | Rod |
| PARD6G-AS1 | Rod |
| CHORDC1 | Rod |
| MT-ND4 | Rod |
| AL137100.3 | Rod |
| ZNF532 | Rod |
| DNAJB4 | Rod |
| CEP112 | Rod |
| NRXN3 | Rod |
| MBP | Rod |
| DISC1 | Rod |
| PDE5A | Rod |
| WWOX | Rod |
| TMEM108 | Rod |
| PPDPF | Rod |
| DNAJA1 | Rod |
| CCSER1 | Rod |
| RPS26 | Rod |
| MT-ATP6 | Rod |
| BCO2 | Rod |
| KCNB1 | Rod |
| MT-CO1 | Rod |
| SRRM4 | Rod |
| CADM2 | Rod |
| AC100803.4 | Rod |
| MT-CYB | Rod |
| DSCAML1 | Rod |
| CDCA7L | Rod |
| CHCHD10 | Rod |
| HSP90AB1 | Rod |
| ARMC9 | Rod |
| RIMS2 | Rod |
| LRP2 | Rod |
| SERPINF1 | Rod |

| | |
|---|---|
| MACROD2 | Rod |
| MIR646HG | Rod |
| PRR16 | Rod |
| EPB41 | Rod |
| ZFAND2A | Rod |
| HSD17B7 | Rod |
| RD3 | Rod |
| TF | Rod |
| DPH6 | Rod |
| LTBP1 | Rod |
| DLG2 | Rod |
| OLFM3 | Rod |
| ANK3 | Rod |
| NPVF | Rod |
| P4HA2 | Rod |
| DNAJA4 | Rod |
| FBXL14 | Rod |
| RFX3-AS1 | Rod |
| LRRC39 | Rod |
| BBS9 | Rod |
| MT-ND1 | Rod |
| PCBP3 | Rod |
| HNRNPA2B1 | Rod |
| PRKN | Rod |
| AC130456.2 | Rod |
| FAM138D | Rod |
| PIK3R3 | Rod |
| YBX3 | Rod |
| IMPG1 | Rod |
| LHFPL3-AS1 | Rod |
| ABCA4 | Rod |
| ABHD3 | Rod |
| MT-ATP8 | Rod |
| NEGR1 | Rod |
| PEX14 | Rod |
| MT3 | Rod |
| CNKSR2 | Rod |
| PSD3 | Rod |
| AC073050.1 | Rod |
| AC130415.1 | Rod |
| S100A10 | Rod |
| ABLIM3 | Rod |
| SPAG16 | Rod |
| BTG4 | Rod |
| CAP2 | Rod |
| IL3RA | Rod |
| HNRNPA3 | Rod |
| KITLG | Rod |
| AL513166.1 | Rod |
| SLC25A24 | Rod |
| ZRANB2-AS2 | Rod |
| LUCAT1 | Rod |
| GLCCI1 | Rod |
| FTX | Rod |

| | |
|---|---|
| PLPP2 | Rod |
| SPDYE2 | Rod |
| PRSS51 | Rod |
| AL583808.1 | Rod |
| NORAD | Rod |
| SELENOW | Rod |
| AP1S2 | Rod |
| JARID2 | Rod |
| SNHG14 | Rod |
| RBP7 | Rod |
| MRPL18 | Rod |
| MAGI2 | Rod |
| LGALS3 | Rod |
| AC084361.1 | Rod |
| TMEM132D | Rod |
| PUM3 | Rod |
| KCTD8 | Rod |
| AL445250.1 | Rod |
| PROM1 | Rod |
| FHIT | Rod |
| P3H2 | Rod |
| GPX3 | Rod |
| MRLN | Rod |
| UNC5C | Rod |
| PLA2G4C-AS1 | Rod |
| GNAQ | Rod |
| WBP2 | Rod |
| AC012178.1 | Rod |
| AL022068.1 | Rod |
| FAM161A | Rod |
| ACSM6 | Rod |
| LYST | Rod |
| TTC3 | Rod |
| MT2A | Rod |
| SYNE2 | Rod |
| HNRNPDL | Rod |
| HNRNPD | Rod |
| DPF3 | Rod |
| DLEU2 | Rod |
| UQCRQ | Rod |
| GREM2 | Rod |
| AGAP1 | Rod |
| AL050403.2 | Rod |
| PRUNE2 | Rod |
| GABRR3 | Rod |
| AC093151.8 | Rod |
| DOCK3 | Rod |
| F13A1 | Rod |
| AJ011931.2 | Rod |
