## Supplementary material for "Interpretable Aging Signatures in Human Retinal Cell Types Revealed by Single-Cell RNA Sequencing and Sparse Logistic Regression": Table S10

Table S10: Machine learning model feature weights for age classification across retinal cell types.

| Gene | Weight | Class | CellTypes |
|---|---|---|---|
| MT-CO3 | -0.65572006 | young | rod |
| HSPA1A | -0.6125753 | young | rod |
| UBB | 0.5830505 | young | rod |
| WAPL | -0.51857185 | young | rod |
| MT-ND3 | 0.50077593 | young | rod |
| MT-CO2 | 0.49435583 | young | rod |
| MT-CO1 | -0.4841151 | young | rod |
| MT-ND2 | 0.46981257 | young | rod |
| DMD | 0.45779094 | young | rod |
| RPS26 | -0.4260229 | young | rod |
| RALYL | -0.39951456 | young | rod |
| PDE6G | -0.38354784 | young | rod |
| MT-CYB | 0.37999827 | young | rod |
| TMEM108 | 0.3738649 | young | rod |
| HSP90AB1 | 0.37171546 | young | rod |
| MT-ATP8 | -0.3710839 | young | rod |
| NLGN1 | -0.35940313 | young | rod |
| MT-ATP6 | -0.34169257 | young | rod |
| PLA2G4C-AS1 | 0.32653466 | young | rod |
| DNAJB1 | 0.3175668 | young | rod |
| AL050403.2 | 0.30588016 | young | rod |
| FAM155A | -0.3029041 | young | rod |
| FAM161A | 0.297915 | young | rod |
| AC130415.1 | -0.29744118 | young | rod |
| EYS | -0.29323292 | young | rod |
| HNRNPDL | 0.29240152 | young | rod |
| HNRNPA2B1 | 0.28544858 | young | rod |
| BAG3 | -0.27137655 | young | rod |
| MRLN | 0.2697056 | young | rod |
| PEX14 | -0.2634437 | young | rod |
| CADM1 | -0.25571698 | young | rod |
| YBX3 | 0.2535373 | young | rod |
| CADM2 | 0.25265405 | young | rod |
| DNAJA1 | 0.24999772 | young | rod |
| AL445250.1 | 0.24911797 | young | rod |
| LHFPL3 | 0.23112205 | young | rod |
| RFX3-AS1 | -0.22837944 | young | rod |
| PLPP2 | -0.22290768 | young | rod |
| MT3 | -0.2195469 | young | rod |
| RD3 | 0.21601416 | young | rod |
| FTX | -0.21119912 | young | rod |
| CNKSR2 | 0.20334913 | young | rod |
| PIK3R3 | -0.20214644 | young | rod |
| HSPA6 | -0.19986753 | young | rod |
| CRYAB | -0.18997478 | young | rod |
| DPF3 | -0.18764609 | young | rod |
| AC012178.1 | -0.18694018 | young | rod |
| NTM | -0.18644342 | young | rod |
| ARMC9 | -0.1843376 | young | rod |
| DRD4 | 0.18195163 | young | rod |
| UQCRQ | 0.18008757 | young | rod |
| GPX3 | -0.17645535 | young | rod |
| P4HA2 | -0.17576078 | young | rod |

| | | | |
|---|---|---|---|
| PCBP3 | 0.1751062 | young | rod |
| LRP2 | -0.17433168 | young | rod |
| KCNB1 | 0.17255145 | young | rod |
| GLCCI1 | -0.17074084 | young | rod |
| ABLIM3 | 0.16761762 | young | rod |
| AC093151.8 | 0.16371667 | young | rod |
| HSPA1B | -0.16229491 | young | rod |
| NCL | 0.15625553 | young | rod |
| FBXL14 | -0.15325196 | young | rod |
| MT-ND1 | -0.15013614 | young | rod |
| EPHA3 | -0.14646628 | young | rod |
| SELENOW | 0.14612588 | young | rod |
| SERPINF1 | -0.1431797 | young | rod |
| HSP90AA1 | 0.14308457 | young | rod |
| S100A10 | -0.14180781 | young | rod |
| TMEM176A | -0.14168042 | young | rod |
| AGAP1 | -0.14141968 | young | rod |
| MT2A | 0.13369673 | young | rod |
| CEP112 | -0.13267745 | young | rod |
| IMPG1 | 0.13036834 | young | rod |
| NPVF | -0.12772971 | young | rod |
| HSD17B7 | -0.12440054 | young | rod |
| MT-ND4 | -0.12389591 | young | rod |
| UNC5C | -0.12092754 | young | rod |
| MRPL18 | 0.11797118 | young | rod |
| ZFAND2A | 0.116240636 | young | rod |
| AC084361.1 | 0.1159983 | young | rod |
| PTPRM | -0.114940405 | young | rod |
| AL355835.1 | 0.11473295 | young | rod |
| AL356804.1 | -0.11330946 | young | rod |
| SRRM4 | 0.11153829 | young | rod |
| AL392023.2 | 0.111032784 | young | rod |
| GNAQ | 0.10807791 | young | rod |
| FAM138D | -0.10577808 | young | rod |
| ABCA4 | -0.104863495 | young | rod |
| HSPB1 | -0.103242844 | young | rod |
| TF | -0.09990001 | young | rod |
| CCSER1 | -0.09941152 | young | rod |
| LRRC39 | 0.09666906 | young | rod |
| PDE5A | 0.0965637 | young | rod |
| NEGR1 | 0.09456967 | young | rod |
| ZNF532 | 0.09438762 | young | rod |
| EPB41L3 | 0.09417473 | young | rod |
| KCTD8 | 0.0891678 | young | rod |
| SPDYE2 | 0.08785467 | young | rod |
| SLC25A24 | 0.087774634 | young | rod |
| PRKN | -0.08741324 | young | rod |
| USH2A | 0.08634904 | young | rod |
| JARID2 | -0.08291371 | young | rod |
| DSCAML1 | -0.08176999 | young | rod |
| AC100803.4 | -0.08171265 | young | rod |
| WBP2 | -0.08116384 | young | rod |
| MIR646HG | 0.08008407 | young | rod |
| PROM1 | -0.07899654 | young | rod |

| | | | |
|---|---|---|---|
| LGALS3 | -0.07802589 | young | rod |
| TMEM132D | -0.07769507 | young | rod |
| F13A1 | -0.07308318 | young | rod |
| A2M | -0.072736114 | young | rod |
| BMPR1B | -0.070853904 | young | rod |
| CDCA7L | -0.07071824 | young | rod |
| NORAD | 0.068979636 | young | rod |
| TRNP1 | -0.06825511 | young | rod |
| PARD6G-AS1 | -0.067621246 | young | rod |
| PPDPF | 0.067211494 | young | rod |
| LHFPL3-AS1 | 0.06624524 | young | rod |
| PLCL2 | 0.06621806 | young | rod |
| SLC35F1 | -0.06557447 | young | rod |
| SNHG14 | -0.065231845 | young | rod |
| EPB41 | 0.06170027 | young | rod |
| GABRR3 | -0.06076115 | young | rod |
| PRUNE2 | 0.060625393 | young | rod |
| NAALADL2 | -0.05899589 | young | rod |
| HNRNPD | 0.057420876 | young | rod |
| FHIT | 0.05694904 | young | rod |
| MIAT | 0.055500187 | young | rod |
| AC130456.2 | 0.05476493 | young | rod |
| MACROD2 | 0.05437553 | young | rod |
| HNRNPA3 | 0.054290894 | young | rod |
| TTC3 | -0.054227613 | young | rod |
| LYST | 0.053387843 | young | rod |
| ZRANB2-AS2 | 0.053062275 | young | rod |
| ANK3 | -0.052777313 | young | rod |
| HSPD1 | 0.05276094 | young | rod |
| CACYBP | -0.051553328 | young | rod |
| RIMS2 | -0.048776597 | young | rod |
| HMGB2 | 0.048316445 | young | rod |
| MT-ND6 | -0.045096993 | young | rod |
| SPAG16 | 0.04473288 | young | rod |
| HSPH1 | -0.044626184 | young | rod |
| BTG4 | -0.044478603 | young | rod |
| DPH6 | -0.041384537 | young | rod |
| TMEM176B | -0.04079119 | young | rod |
| HPRT1 | -0.039961174 | young | rod |
| AL022068.1 | -0.037738223 | young | rod |
| LTBP1 | 0.0361864 | young | rod |
| RGS9 | 0.03561156 | young | rod |
| MBP | -0.03525871 | young | rod |
| SYNE2 | 0.033622276 | young | rod |
| AJ011931.2 | 0.033479024 | young | rod |
| DISC1 | -0.033100575 | young | rod |
| RBP7 | -0.032737296 | young | rod |
| DLEU2 | -0.032416783 | young | rod |
| ACSM6 | 0.031751465 | young | rod |
| GREM2 | -0.03063083 | young | rod |
| AL583808.1 | -0.029885344 | young | rod |
| BCO2 | -0.029491547 | young | rod |
| DOCK3 | -0.028389653 | young | rod |
| BBS9 | 0.027993752 | young | rod |

| | | | |
|---|---|---|---|
| OLFM3 | -0.027628453 | young | rod |
| CHCHD10 | -0.027480833 | young | rod |
| CHORDC1 | 0.02632546 | young | rod |
| MAGI2 | 0.025204066 | young | rod |
| PRR16 | -0.024837127 | young | rod |
| AL137100.3 | 0.024631359 | young | rod |
| LUCAT1 | 0.021904798 | young | rod |
| IL3RA | 0.021215543 | young | rod |
| KITLG | -0.020823542 | young | rod |
| DNAJB4 | 0.020474125 | young | rod |
| PRSS51 | 0.020382246 | young | rod |
| P3H2 | -0.018326923 | young | rod |
| PSD3 | 0.015587448 | young | rod |
| HMGN1 | 0.015540654 | young | rod |
| WWOX | -0.013446365 | young | rod |
| AP1S2 | 0.013325091 | young | rod |
| AC016723.1 | -0.012844662 | young | rod |
| NRXN3 | -0.012269611 | young | rod |
| ABHD3 | -0.011895254 | young | rod |
| OTX2-AS1 | -0.01184875 | young | rod |
| AL513166.1 | -0.011498878 | young | rod |
| POU2AF1 | -0.010581056 | young | rod |
| METRNL | -0.009631348 | young | rod |
| CAP2 | -0.009474743 | young | rod |
| MT-ND5 | 0.009018756 | young | rod |
| AC073050.1 | 0.008613649 | young | rod |
| DNAJA4 | -0.006519981 | young | rod |
| AHSA1 | 0.006314519 | young | rod |
| DLG2 | 0.005982735 | young | rod |
| PUM3 | 0.00575358 | young | rod |
| MT-CO3 | 0.65572006 | old | rod |
| HSPA1A | 0.6125753 | old | rod |
| UBB | -0.5830505 | old | rod |
| WAPL | 0.51857185 | old | rod |
| MT-ND3 | -0.50077593 | old | rod |
| MT-CO2 | -0.49435583 | old | rod |
| MT-CO1 | 0.4841151 | old | rod |
| MT-ND2 | -0.46981257 | old | rod |
| DMD | -0.45779094 | old | rod |
| RPS26 | 0.4260229 | old | rod |
| RALYL | 0.39951456 | old | rod |
| PDE6G | 0.38354784 | old | rod |
| MT-CYB | -0.37999827 | old | rod |
| TMEM108 | -0.3738649 | old | rod |
| HSP90AB1 | -0.37171546 | old | rod |
| MT-ATP8 | 0.3710839 | old | rod |
| NLGN1 | 0.35940313 | old | rod |
| MT-ATP6 | 0.34169257 | old | rod |
| PLA2G4C-AS1 | -0.32653466 | old | rod |
| DNAJB1 | -0.3175668 | old | rod |
| AL050403.2 | -0.30588016 | old | rod |
| FAM155A | 0.3029041 | old | rod |
| FAM161A | -0.297915 | old | rod |
| AC130415.1 | 0.29744118 | old | rod |

| | | | |
|---|---|---|---|
| EYS | 0.29323292 | old | rod |
| HNRNPDL | -0.29240152 | old | rod |
| HNRNPA2B1 | -0.28544858 | old | rod |
| BAG3 | 0.27137655 | old | rod |
| MRLN | -0.2697056 | old | rod |
| PEX14 | 0.2634437 | old | rod |
| CADM1 | 0.25571698 | old | rod |
| YBX3 | -0.2535373 | old | rod |
| CADM2 | -0.25265405 | old | rod |
| DNAJA1 | -0.24999772 | old | rod |
| AL445250.1 | -0.24911797 | old | rod |
| LHFPL3 | -0.23112205 | old | rod |
| RFX3-AS1 | 0.22837944 | old | rod |
| PLPP2 | 0.22290768 | old | rod |
| MT3 | 0.2195469 | old | rod |
| RD3 | -0.21601416 | old | rod |
| FTX | 0.21119912 | old | rod |
| CNKSR2 | -0.20334913 | old | rod |
| PIK3R3 | 0.20214644 | old | rod |
| HSPA6 | 0.19986753 | old | rod |
| CRYAB | 0.18997478 | old | rod |
| DPF3 | 0.18764609 | old | rod |
| AC012178.1 | 0.18694018 | old | rod |
| NTM | 0.18644342 | old | rod |
| ARMC9 | 0.1843376 | old | rod |
| DRD4 | -0.18195163 | old | rod |
| UQCRQ | -0.18008757 | old | rod |
| GPX3 | 0.17645535 | old | rod |
| P4HA2 | 0.17576078 | old | rod |
| PCBP3 | -0.1751062 | old | rod |
| LRP2 | 0.17433168 | old | rod |
| KCNB1 | -0.17255145 | old | rod |
| GLCCI1 | 0.17074084 | old | rod |
| ABLIM3 | -0.16761762 | old | rod |
| AC093151.8 | -0.16371667 | old | rod |
| HSPA1B | 0.16229491 | old | rod |
| NCL | -0.15625553 | old | rod |
| FBXL14 | 0.15325196 | old | rod |
| MT-ND1 | 0.15013614 | old | rod |
| EPHA3 | 0.14646628 | old | rod |
| SELENOW | -0.14612588 | old | rod |
| SERPINF1 | 0.1431797 | old | rod |
| HSP90AA1 | -0.14308457 | old | rod |
| S100A10 | 0.14180781 | old | rod |
| TMEM176A | 0.14168042 | old | rod |
| AGAP1 | 0.14141968 | old | rod |
| MT2A | -0.13369673 | old | rod |
| CEP112 | 0.13267745 | old | rod |
| IMPG1 | -0.13036834 | old | rod |
| NPVF | 0.12772971 | old | rod |
| HSD17B7 | 0.12440054 | old | rod |
| MT-ND4 | 0.12389591 | old | rod |
| UNC5C | 0.12092754 | old | rod |
| MRPL18 | -0.11797118 | old | rod |

| | | | |
|---|---|---|---|
| ZFAND2A | -0.116240636 | old | rod |
| AC084361.1 | -0.1159983 | old | rod |
| PTPRM | 0.114940405 | old | rod |
| AL355835.1 | -0.11473295 | old | rod |
| AL356804.1 | 0.11330946 | old | rod |
| SRRM4 | -0.11153829 | old | rod |
| AL392023.2 | -0.111032784 | old | rod |
| GNAQ | -0.10807791 | old | rod |
| FAM138D | 0.10577808 | old | rod |
| ABCA4 | 0.104863495 | old | rod |
| HSPB1 | 0.103242844 | old | rod |
| TF | 0.09990001 | old | rod |
| CCSER1 | 0.09941152 | old | rod |
| LRRC39 | -0.09666906 | old | rod |
| PDE5A | -0.0965637 | old | rod |
| NEGR1 | -0.09456967 | old | rod |
| ZNF532 | -0.09438762 | old | rod |
| EPB41L3 | -0.09417473 | old | rod |
| KCTD8 | -0.0891678 | old | rod |
| SPDYE2 | -0.08785467 | old | rod |
| SLC25A24 | -0.087774634 | old | rod |
| PRKN | 0.08741324 | old | rod |
| USH2A | -0.08634904 | old | rod |
| JARID2 | 0.08291371 | old | rod |
| DSCAML1 | 0.08176999 | old | rod |
| AC100803.4 | 0.08171265 | old | rod |
| WBP2 | 0.08116384 | old | rod |
| MIR646HG | -0.08008407 | old | rod |
| PROM1 | 0.07899654 | old | rod |
| LGALS3 | 0.07802589 | old | rod |
| TMEM132D | 0.07769507 | old | rod |
| F13A1 | 0.07308318 | old | rod |
| A2M | 0.072736114 | old | rod |
| BMPR1B | 0.070853904 | old | rod |
| CDCA7L | 0.07071824 | old | rod |
| NORAD | -0.068979636 | old | rod |
| TRNP1 | 0.06825511 | old | rod |
| PARD6G-AS1 | 0.067621246 | old | rod |
| PPDPF | -0.067211494 | old | rod |
| LHFPL3-AS1 | -0.06624524 | old | rod |
| PLCL2 | -0.06621806 | old | rod |
| SLC35F1 | 0.06557447 | old | rod |
| SNHG14 | 0.065231845 | old | rod |
| EPB41 | -0.06170027 | old | rod |
| GABRR3 | 0.06076115 | old | rod |
| PRUNE2 | -0.060625393 | old | rod |
| NAALADL2 | 0.05899589 | old | rod |
| HNRNPD | -0.057420876 | old | rod |
| FHIT | -0.05694904 | old | rod |
| MIAT | -0.055500187 | old | rod |
| AC130456.2 | -0.05476493 | old | rod |
| MACROD2 | -0.05437553 | old | rod |
| HNRNPA3 | -0.054290894 | old | rod |
| TTC3 | 0.054227613 | old | rod |

| | | | |
|---|---|---|---|
| LYST | -0.053387843 | old | rod |
| ZRANB2-AS2 | -0.053062275 | old | rod |
| ANK3 | 0.052777313 | old | rod |
| HSPD1 | -0.05276094 | old | rod |
| CACYBP | 0.051553328 | old | rod |
| RIMS2 | 0.048776597 | old | rod |
| HMGB2 | -0.048316445 | old | rod |
| MT-ND6 | 0.045096993 | old | rod |
| SPAG16 | -0.04473288 | old | rod |
| HSPH1 | 0.044626184 | old | rod |
| BTG4 | 0.044478603 | old | rod |
| DPH6 | 0.041384537 | old | rod |
| TMEM176B | 0.04079119 | old | rod |
| HPRT1 | 0.039961174 | old | rod |
| AL022068.1 | 0.037738223 | old | rod |
| LTBP1 | -0.0361864 | old | rod |
| RGS9 | -0.03561156 | old | rod |
| MBP | 0.03525871 | old | rod |
| SYNE2 | -0.033622276 | old | rod |
| AJ011931.2 | -0.033479024 | old | rod |
| DISC1 | 0.033100575 | old | rod |
| RBP7 | 0.032737296 | old | rod |
| DLEU2 | 0.032416783 | old | rod |
| ACSM6 | -0.031751465 | old | rod |
| GREM2 | 0.03063083 | old | rod |
| AL583808.1 | 0.029885344 | old | rod |
| BCO2 | 0.029491547 | old | rod |
| DOCK3 | 0.028389653 | old | rod |
| BBS9 | -0.027993752 | old | rod |
| OLFM3 | 0.027628453 | old | rod |
| CHCHD10 | 0.027480833 | old | rod |
| CHORDC1 | -0.02632546 | old | rod |
| MAGI2 | -0.025204066 | old | rod |
| PRR16 | 0.024837127 | old | rod |
| AL137100.3 | -0.024631359 | old | rod |
| LUCAT1 | -0.021904798 | old | rod |
| IL3RA | -0.021215543 | old | rod |
| KITLG | 0.020823542 | old | rod |
| DNAJB4 | -0.020474125 | old | rod |
| PRSS51 | -0.020382246 | old | rod |
| P3H2 | 0.018326923 | old | rod |
| PSD3 | -0.015587448 | old | rod |
| HMGN1 | -0.015540654 | old | rod |
| WWOX | 0.013446365 | old | rod |
| AP1S2 | -0.013325091 | old | rod |
| AC016723.1 | 0.012844662 | old | rod |
| NRXN3 | 0.012269611 | old | rod |
| ABHD3 | 0.011895254 | old | rod |
| OTX2-AS1 | 0.01184875 | old | rod |
| AL513166.1 | 0.011498878 | old | rod |
| POU2AF1 | 0.010581056 | old | rod |
| METRNL | 0.009631348 | old | rod |
| CAP2 | 0.009474743 | old | rod |
| MT-ND5 | -0.009018756 | old | rod |

| | | | |
|---|---|---|---|
| AC073050.1 | -0.008613649 | old | rod |
| DNAJA4 | 0.006519981 | old | rod |
| AHSA1 | -0.006314519 | old | rod |
| DLG2 | -0.005982735 | old | rod |
| PUM3 | -0.00575358 | old | rod |
| SLC16A10 | -0.8719747 | young | microglia |
| MT1E | -0.72157484 | young | microglia |
| MT1X | -0.6839769 | young | microglia |
| APOC1 | 0.54511887 | young | microglia |
| PGK1 | -0.47926807 | young | microglia |
| P4HA1 | -0.46375445 | young | microglia |
| FTH1 | -0.42340624 | young | microglia |
| MT2A | -0.41094437 | young | microglia |
| MBNL1 | -0.35622734 | young | microglia |
| RGCC | 0.35424578 | young | microglia |
| PLAUR | 0.3278674 | young | microglia |
| THBS1 | -0.3100419 | young | microglia |
| IPCEF1 | 0.30916974 | young | microglia |
| IFI44L | 0.30911788 | young | microglia |
| GAPDH | -0.30510136 | young | microglia |
| PMP22 | -0.3034275 | young | microglia |
| FTL | 0.30026016 | young | microglia |
| IFI6 | 0.2992726 | young | microglia |
| NAV3 | 0.29474804 | young | microglia |
| CCL2 | -0.2904906 | young | microglia |
| IGSF6 | 0.29004958 | young | microglia |
| CH25H | 0.28429353 | young | microglia |
| TPT1 | -0.27340162 | young | microglia |
| NCKAP5 | -0.2732983 | young | microglia |
| CALM2 | 0.27087292 | young | microglia |
| TNFRSF11B | -0.26118505 | young | microglia |
| QKI | -0.24847297 | young | microglia |
| CX3CR1 | 0.23245806 | young | microglia |
| SLC2A3 | -0.22618306 | young | microglia |
| MT-ND6 | 0.21987256 | young | microglia |
| GPNMB | -0.21134143 | young | microglia |
| FGF12 | 0.2112857 | young | microglia |
| CCDC26 | 0.21023782 | young | microglia |
| BASP1 | 0.20334367 | young | microglia |
| SHTN1 | 0.19933265 | young | microglia |
| BTG2 | 0.19030076 | young | microglia |
| PARP14 | -0.18755955 | young | microglia |
| LPCAT2 | 0.18345101 | young | microglia |
| EPSTI1 | 0.1833577 | young | microglia |
| ADAM8 | -0.18244375 | young | microglia |
| BNIP3L | -0.1785519 | young | microglia |
| METRNL | 0.17640218 | young | microglia |
| CCL3L1 | 0.17302142 | young | microglia |
| EGR3 | -0.16459972 | young | microglia |
| HDAC9 | 0.16360408 | young | microglia |
| AL591518.1 | 0.16146979 | young | microglia |
| SLCO2B1 | -0.16122429 | young | microglia |
| MGAT5 | -0.15221803 | young | microglia |
| ITPR2 | -0.1489141 | young | microglia |

| | | | |
|---|---|---|---|
| SOCS3 | -0.14705306 | young | microglia |
| PLIN2 | 0.1395991 | young | microglia |
| SLC35F1 | -0.13567388 | young | microglia |
| SPP1 | -0.13346687 | young | microglia |
| CDK6 | 0.12556295 | young | microglia |
| SRGN | -0.12292234 | young | microglia |
| APOE | 0.11696667 | young | microglia |
| SPRED1 | -0.11621305 | young | microglia |
| CYFIP1 | -0.11616854 | young | microglia |
| THADA | 0.115350164 | young | microglia |
| JDP2 | 0.1150588 | young | microglia |
| TUBA1B | 0.111779734 | young | microglia |
| ENSA | 0.1089798 | young | microglia |
| CCL4L2 | 0.10872372 | young | microglia |
| MT3 | -0.108596936 | young | microglia |
| FOSB | 0.10635343 | young | microglia |
| CD9 | -0.104422346 | young | microglia |
| C4orf3 | 0.10138125 | young | microglia |
| FPR3 | 0.099783055 | young | microglia |
| ADM | -0.09923039 | young | microglia |
| XAF1 | 0.0980445 | young | microglia |
| ZBTB20 | -0.09574827 | young | microglia |
| TPI1 | -0.09489872 | young | microglia |
| MS4A4E | -0.09416489 | young | microglia |
| TBXAS1 | -0.09348125 | young | microglia |
| ERO1A | -0.09279746 | young | microglia |
| FRMD4A | -0.09257748 | young | microglia |
| FAU | 0.088505715 | young | microglia |
| SLC11A1 | -0.08477128 | young | microglia |
| 1-Mar | -0.083596304 | young | microglia |
| ARL4C | -0.082679346 | young | microglia |
| SAMHD1 | 0.08033611 | young | microglia |
| IFI16 | 0.07758081 | young | microglia |
| ARHGAP15 | -0.07452261 | young | microglia |
| ELMO1 | -0.07419221 | young | microglia |
| LGALS1 | 0.06977461 | young | microglia |
| CSTB | 0.0660962 | young | microglia |
| ILDR1 | 0.0642639 | young | microglia |
| TBC1D5 | -0.06381275 | young | microglia |
| JAK2 | 0.059250258 | young | microglia |
| VIM | 0.058982074 | young | microglia |
| NR4A2 | -0.058810644 | young | microglia |
| RNASE4 | -0.058007374 | young | microglia |
| CAMK1D | 0.056323253 | young | microglia |
| BLNK | 0.0553011 | young | microglia |
| BMP2K | -0.0550315 | young | microglia |
| AC073352.2 | 0.05091808 | young | microglia |
| PFDN5 | -0.046913445 | young | microglia |
| PLXDC2 | -0.043736108 | young | microglia |
| DOCK2 | 0.040294155 | young | microglia |
| FCGR1B | 0.03983237 | young | microglia |
| CLEC2B | 0.039719425 | young | microglia |
| SAMD9L | 0.03812679 | young | microglia |
| ENO1 | -0.03694582 | young | microglia |

| | | | |
|---|---|---|---|
| AC008574.1 | 0.03510002 | young | microglia |
| SKAP2 | 0.03448894 | young | microglia |
| GLUL | -0.03418973 | young | microglia |
| MIF | -0.03396509 | young | microglia |
| PSME2 | 0.03392204 | young | microglia |
| MT1G | -0.03351166 | young | microglia |
| SLC16A10 | 0.8719747 | old | microglia |
| MT1E | 0.72157484 | old | microglia |
| MT1X | 0.6839769 | old | microglia |
| APOC1 | -0.54511887 | old | microglia |
| PGK1 | 0.47926807 | old | microglia |
| P4HA1 | 0.46375445 | old | microglia |
| FTH1 | 0.42340624 | old | microglia |
| MT2A | 0.41094437 | old | microglia |
| MBNL1 | 0.35622734 | old | microglia |
| RGCC | -0.35424578 | old | microglia |
| PLAUR | -0.3278674 | old | microglia |
| THBS1 | 0.3100419 | old | microglia |
| IPCEF1 | -0.30916974 | old | microglia |
| IFI44L | -0.30911788 | old | microglia |
| GAPDH | 0.30510136 | old | microglia |
| PMP22 | 0.3034275 | old | microglia |
| FTL | -0.30026016 | old | microglia |
| IFI6 | -0.2992726 | old | microglia |
| NAV3 | -0.29474804 | old | microglia |
| CCL2 | 0.2904906 | old | microglia |
| IGSF6 | -0.29004958 | old | microglia |
| CH25H | -0.28429353 | old | microglia |
| TPT1 | 0.27340162 | old | microglia |
| NCKAP5 | 0.2732983 | old | microglia |
| CALM2 | -0.27087292 | old | microglia |
| TNFRSF11B | 0.26118505 | old | microglia |
| QKI | 0.24847297 | old | microglia |
| CX3CR1 | -0.23245806 | old | microglia |
| SLC2A3 | 0.22618306 | old | microglia |
| MT-ND6 | -0.21987256 | old | microglia |
| GPNMB | 0.21134143 | old | microglia |
| FGF12 | -0.2112857 | old | microglia |
| CCDC26 | -0.21023782 | old | microglia |
| BASP1 | -0.20334367 | old | microglia |
| SHTN1 | -0.19933265 | old | microglia |
| BTG2 | -0.19030076 | old | microglia |
| PARP14 | 0.18755955 | old | microglia |
| LPCAT2 | -0.18345101 | old | microglia |
| EPSTI1 | -0.1833577 | old | microglia |
| ADAM8 | 0.18244375 | old | microglia |
| BNIP3L | 0.1785519 | old | microglia |
| METRNL | -0.17640218 | old | microglia |
| CCL3L1 | -0.17302142 | old | microglia |
| EGR3 | 0.16459972 | old | microglia |
| HDAC9 | -0.16360408 | old | microglia |
| AL591518.1 | -0.16146979 | old | microglia |
| SLCO2B1 | 0.16122429 | old | microglia |
| MGAT5 | 0.15221803 | old | microglia |

| | | | |
|---|---|---|---|
| ITPR2 | 0.1489141 | old | microglia |
| SOCS3 | 0.14705306 | old | microglia |
| PLIN2 | -0.1395991 | old | microglia |
| SLC35F1 | 0.13567388 | old | microglia |
| SPP1 | 0.13346687 | old | microglia |
| CDK6 | -0.12556295 | old | microglia |
| SRGN | 0.12292234 | old | microglia |
| APOE | -0.11696667 | old | microglia |
| SPRED1 | 0.11621305 | old | microglia |
| CYFIP1 | 0.11616854 | old | microglia |
| THADA | -0.115350164 | old | microglia |
| JDP2 | -0.1150588 | old | microglia |
| TUBA1B | -0.111779734 | old | microglia |
| ENSA | -0.1089798 | old | microglia |
| CCL4L2 | -0.10872372 | old | microglia |
| MT3 | 0.108596936 | old | microglia |
| FOSB | -0.10635343 | old | microglia |
| CD9 | 0.104422346 | old | microglia |
| C4orf3 | -0.10138125 | old | microglia |
| FPR3 | -0.099783055 | old | microglia |
| ADM | 0.09923039 | old | microglia |
| XAF1 | -0.0980445 | old | microglia |
| ZBTB20 | 0.09574827 | old | microglia |
| TPI1 | 0.09489872 | old | microglia |
| MS4A4E | 0.09416489 | old | microglia |
| TBXAS1 | 0.09348125 | old | microglia |
| ERO1A | 0.09279746 | old | microglia |
| FRMD4A | 0.09257748 | old | microglia |
| FAU | -0.088505715 | old | microglia |
| SLC11A1 | 0.08477128 | old | microglia |
| 1-Mar | 0.083596304 | old | microglia |
| ARL4C | 0.082679346 | old | microglia |
| SAMHD1 | -0.08033611 | old | microglia |
| IFI16 | -0.07758081 | old | microglia |
| ARHGAP15 | 0.07452261 | old | microglia |
| ELMO1 | 0.07419221 | old | microglia |
| LGALS1 | -0.06977461 | old | microglia |
| CSTB | -0.0660962 | old | microglia |
| ILDR1 | -0.0642639 | old | microglia |
| TBC1D5 | 0.06381275 | old | microglia |
| JAK2 | -0.059250258 | old | microglia |
| VIM | -0.058982074 | old | microglia |
| NR4A2 | 0.058810644 | old | microglia |
| RNASE4 | 0.058007374 | old | microglia |
| CAMK1D | -0.056323253 | old | microglia |
| BLNK | -0.0553011 | old | microglia |
| BMP2K | 0.0550315 | old | microglia |
| AC073352.2 | -0.05091808 | old | microglia |
| PFDN5 | 0.046913445 | old | microglia |
| PLXDC2 | 0.043736108 | old | microglia |
| DOCK2 | -0.040294155 | old | microglia |
| FCGR1B | -0.03983237 | old | microglia |
| CLEC2B | -0.039719425 | old | microglia |
| SAMD9L | -0.03812679 | old | microglia |

| | | | |
|---|---|---|---|
| ENO1 | 0.03694582 | old | microglia |
| AC008574.1 | -0.03510002 | old | microglia |
| SKAP2 | -0.03448894 | old | microglia |
| GLUL | 0.03418973 | old | microglia |
| MIF | 0.03396509 | old | microglia |
| PSME2 | -0.03392204 | old | microglia |
| MT1G | 0.03351166 | old | microglia |
| MT-CO3 | -1.2107444 | young | MGC |
| MT-ND3 | 0.6808855 | young | MGC |
| HLA-DRA | -0.5290023 | young | MGC |
| GRIA4 | 0.47682288 | young | MGC |
| CALM1 | 0.46606606 | young | MGC |
| WIF1 | -0.45332193 | young | MGC |
| COL4A3 | -0.44625923 | young | MGC |
| RORA | -0.42573914 | young | MGC |
| ADM | -0.3927048 | young | MGC |
| ANXA1 | -0.3861575 | young | MGC |
| AQP4 | 0.3606024 | young | MGC |
| MT-ND6 | 0.35883197 | young | MGC |
| RPS26 | -0.35548684 | young | MGC |
| HILPDA | -0.34983432 | young | MGC |
| HSPA1A | -0.34578577 | young | MGC |
| MIR99AHG | 0.32789162 | young | MGC |
| JUNB | 0.3204837 | young | MGC |
| GPX3 | -0.31894124 | young | MGC |
| NRXN3 | -0.3009659 | young | MGC |
| GDF15 | -0.29497907 | young | MGC |
| LGALS3 | 0.29375204 | young | MGC |
| ERO1A | -0.2902341 | young | MGC |
| DACH1 | 0.2827653 | young | MGC |
| C11orf96 | -0.26930547 | young | MGC |
| ACTG1 | -0.25417632 | young | MGC |
| HSPA1B | -0.25410184 | young | MGC |
| CD74 | -0.24928865 | young | MGC |
| MT-ND2 | 0.24166396 | young | MGC |
| COL4A4 | -0.23683555 | young | MGC |
| ANGPTL1 | 0.23534185 | young | MGC |
| SYT11 | 0.23288287 | young | MGC |
| BHLHE41 | -0.23104113 | young | MGC |
| MT-CO2 | 0.22917655 | young | MGC |
| LINC00461 | 0.22602448 | young | MGC |
| SERPINA3 | -0.21759874 | young | MGC |
| UBC | -0.21445014 | young | MGC |
| CCN1 | 0.20908362 | young | MGC |
| MTTP | 0.20580158 | young | MGC |
| MED12L | -0.19843048 | young | MGC |
| YBX3 | -0.19595261 | young | MGC |
| RBP1 | 0.19334197 | young | MGC |
| DUSP1 | 0.19082882 | young | MGC |
| RAB1A | -0.1884729 | young | MGC |
| DDIT3 | 0.18457673 | young | MGC |
| GADD45B | 0.18266837 | young | MGC |
| IFITM3 | 0.1809538 | young | MGC |
| DYNC1I1 | -0.17779967 | young | MGC |

| | | | |
|---|---|---|---|
| KHDRBS2 | 0.1769813 | young | MGC |
| PMEPA1 | -0.17619266 | young | MGC |
| AL390957.1 | 0.17536826 | young | MGC |
| CDH2 | 0.17447378 | young | MGC |
| LITAF | -0.17322731 | young | MGC |
| MT-ND4L | 0.16778645 | young | MGC |
| SULF1 | 0.16582935 | young | MGC |
| SPON1 | 0.16434218 | young | MGC |
| NAMPT | -0.1639229 | young | MGC |
| JUND | 0.16124411 | young | MGC |
| SCD | 0.1603773 | young | MGC |
| NR4A1 | 0.16009846 | young | MGC |
| FTX | -0.15487336 | young | MGC |
| HLA-A | -0.1426314 | young | MGC |
| HSPA6 | -0.14204451 | young | MGC |
| CD44 | -0.14010204 | young | MGC |
| CLDN1 | -0.13852668 | young | MGC |
| NRG3 | -0.13434519 | young | MGC |
| HSPE1 | 0.13431026 | young | MGC |
| PMP22 | -0.13378167 | young | MGC |
| COL24A1 | 0.13266416 | young | MGC |
| DNAJB1 | 0.13020413 | young | MGC |
| ABI3BP | -0.12884644 | young | MGC |
| ID1 | -0.12825428 | young | MGC |
| GRID2 | 0.12735702 | young | MGC |
| CXCL2 | -0.12634225 | young | MGC |
| EPHB6 | 0.12547414 | young | MGC |
| SLC3A2 | -0.1243807 | young | MGC |
| ID2 | 0.123736925 | young | MGC |
| PAX6 | -0.123105414 | young | MGC |
| S100A11 | -0.1229473 | young | MGC |
| ILDR2 | 0.12263897 | young | MGC |
| KDR | 0.12138976 | young | MGC |
| SERPINE1 | -0.11999616 | young | MGC |
| NAALADL2 | -0.119713575 | young | MGC |
| MAFF | -0.11904408 | young | MGC |
| S100A10 | -0.11895286 | young | MGC |
| OSBPL10 | -0.117020026 | young | MGC |
| TFRC | -0.11339665 | young | MGC |
| RORB | 0.10712356 | young | MGC |
| MT-ND5 | 0.10708535 | young | MGC |
| GFAP | 0.10609929 | young | MGC |
| ARL4D | -0.106006734 | young | MGC |
| GPRC5A | -0.105204694 | young | MGC |
| EIF1 | 0.10494962 | young | MGC |
| TUBB2B | 0.103842124 | young | MGC |
| CLRN1 | 0.09668896 | young | MGC |
| SLCO4A1-AS1 | -0.094825655 | young | MGC |
| SPSB1 | -0.09440669 | young | MGC |
| PIM1 | -0.09305314 | young | MGC |
| SIPA1L1 | -0.0895263 | young | MGC |
| SDC4 | -0.08850088 | young | MGC |
| PCYT1B | 0.087945834 | young | MGC |
| PHYHIPL | 0.08705388 | young | MGC |

| | | | |
|---|---|---|---|
| ZBTB21 | 0.086828314 | young | MGC |
| CYP4V2 | 0.086109854 | young | MGC |
| HLA-C | -0.08343397 | young | MGC |
| MYO10 | 0.08280732 | young | MGC |
| COTL1 | -0.08093218 | young | MGC |
| MAGI2 | -0.08020736 | young | MGC |
| CCL2 | 0.080201976 | young | MGC |
| FOSL2 | -0.07924376 | young | MGC |
| ATF3 | 0.07581367 | young | MGC |
| FBXO32 | -0.07490357 | young | MGC |
| KLKB1 | 0.07413288 | young | MGC |
| INSIG1 | 0.07404623 | young | MGC |
| GPM6A | -0.07373863 | young | MGC |
| S100A6 | 0.06933736 | young | MGC |
| SLC19A2 | -0.06902955 | young | MGC |
| B2M | 0.06892461 | young | MGC |
| MKNK2 | 0.06882993 | young | MGC |
| SNHG16 | -0.067449555 | young | MGC |
| NFIA | -0.0672737 | young | MGC |
| TRH | -0.06331481 | young | MGC |
| CHMP1B | 0.056838542 | young | MGC |
| CYSTM1 | -0.05594083 | young | MGC |
| FGFBP2 | -0.055472642 | young | MGC |
| ADD1 | -0.055236623 | young | MGC |
| ACTB | 0.05483125 | young | MGC |
| GRIA1 | 0.05463142 | young | MGC |
| SLC7A5 | -0.05167719 | young | MGC |
| SERPING1 | -0.048895255 | young | MGC |
| KLHDC8A | 0.048420474 | young | MGC |
| SDS | -0.046732083 | young | MGC |
| KIAA1217 | -0.046358302 | young | MGC |
| MT-ATP6 | -0.045966312 | young | MGC |
| LRRC4C | -0.04273075 | young | MGC |
| SFXN5 | 0.04249099 | young | MGC |
| NCKAP5 | -0.039667048 | young | MGC |
| SPHK1 | -0.039028615 | young | MGC |
| CNTN1 | 0.03853646 | young | MGC |
| PTPRZ1 | 0.035539694 | young | MGC |
| SFRP2 | 0.034888223 | young | MGC |
| CCDC107 | 0.034450725 | young | MGC |
| AMER2 | -0.033743076 | young | MGC |
| AC092691.1 | -0.03361334 | young | MGC |
| AMACR | 0.033190534 | young | MGC |
| LMCD1 | -0.03167466 | young | MGC |
| ANOS1 | 0.031468146 | young | MGC |
| PDLIM4 | -0.03124394 | young | MGC |
| IER5L | -0.030797577 | young | MGC |
| ID3 | 0.030155065 | young | MGC |
| PPP1R15A | 0.028876815 | young | MGC |
| AQP4-AS1 | 0.028603118 | young | MGC |
| PDLIM3 | -0.024317887 | young | MGC |
| EPCAM | -0.023420919 | young | MGC |
| KLHL8 | -0.022997444 | young | MGC |
| ADGRL3 | 0.022990435 | young | MGC |

| | | | |
|---|---|---|---|
| MAP2 | 0.022749882 | young | MGC |
| SOD2 | 0.022414435 | young | MGC |
| MT2A | 0.022406006 | young | MGC |
| EPB41L2 | 0.021314073 | young | MGC |
| NFKBIA | 0.01956406 | young | MGC |
| SRGAP3 | 0.019167451 | young | MGC |
| PCSK2 | 0.016863769 | young | MGC |
| MEGF10 | -0.01676009 | young | MGC |
| PREX2 | 0.015625697 | young | MGC |
| RARRES1 | -0.014685992 | young | MGC |
| BAIAP2 | 0.013218169 | young | MGC |
| CPA6 | -0.012835086 | young | MGC |
| INHBA | 0.012519052 | young | MGC |
| FAP | 0.011083353 | young | MGC |
| HECW2 | 0.010070307 | young | MGC |
| SOX6 | 0.009834558 | young | MGC |
| BHLHE40 | -0.008354692 | young | MGC |
| HLA-DRB1 | 0.006560494 | young | MGC |
| RRAD | 0.006492203 | young | MGC |
| SORBS2 | -0.005820125 | young | MGC |
| ABCA8 | -0.004711746 | young | MGC |
| CHST9 | 0.003944216 | young | MGC |
| UTRN | 0.001391736 | young | MGC |
| S100A16 | -0.001254006 | young | MGC |
| FUT8 | -0.000754316 | young | MGC |
| RHOU | 0.000316126 | young | MGC |
| MT-ATP8 | 0.000294021 | young | MGC |
| CRYAB | 0 | young | MGC |
| NAV3 | 0 | young | MGC |
| GEM | 0 | young | MGC |
| PHLPP1 | 0 | young | MGC |
| WTAP | 0 | young | MGC |
| ZNF385D | 0 | young | MGC |
| MYO6 | 0 | young | MGC |
| CTTNBP2 | 0 | young | MGC |
| CXCL3 | 0 | young | MGC |
| IER3 | 0 | young | MGC |
| CEBPB | 0 | young | MGC |
| AC004264.1 | 0 | young | MGC |
| DENND11 | 0 | young | MGC |
| IFITM2 | 0 | young | MGC |
| ZNF331 | 0 | young | MGC |
| SOX5 | 0 | young | MGC |
| ARHGAP21 | 0 | young | MGC |
| MT-CO3 | 1.2107444 | old | MGC |
| MT-ND3 | -0.6808855 | old | MGC |
| HLA-DRA | 0.5290023 | old | MGC |
| GRIA4 | -0.47682288 | old | MGC |
| CALM1 | -0.46606606 | old | MGC |
| WIF1 | 0.45332193 | old | MGC |
| COL4A3 | 0.44625923 | old | MGC |
| RORA | 0.42573914 | old | MGC |
| ADM | 0.3927048 | old | MGC |
| ANXA1 | 0.3861575 | old | MGC |

| | | | |
|---|---|---|---|
| AQP4 | -0.3606024 | old | MGC |
| MT-ND6 | -0.35883197 | old | MGC |
| RPS26 | 0.35548684 | old | MGC |
| HILPDA | 0.34983432 | old | MGC |
| HSPA1A | 0.34578577 | old | MGC |
| MIR99AHG | -0.32789162 | old | MGC |
| JUNB | -0.3204837 | old | MGC |
| GPX3 | 0.31894124 | old | MGC |
| NRXN3 | 0.3009659 | old | MGC |
| GDF15 | 0.29497907 | old | MGC |
| LGALS3 | -0.29375204 | old | MGC |
| ERO1A | 0.2902341 | old | MGC |
| DACH1 | -0.2827653 | old | MGC |
| C11orf96 | 0.26930547 | old | MGC |
| ACTG1 | 0.25417632 | old | MGC |
| HSPA1B | 0.25410184 | old | MGC |
| CD74 | 0.24928865 | old | MGC |
| MT-ND2 | -0.24166396 | old | MGC |
| COL4A4 | 0.23683555 | old | MGC |
| ANGPTL1 | -0.23534185 | old | MGC |
| SYT11 | -0.23288287 | old | MGC |
| BHLHE41 | 0.23104113 | old | MGC |
| MT-CO2 | -0.22917655 | old | MGC |
| LINC00461 | -0.22602448 | old | MGC |
| SERPINA3 | 0.21759874 | old | MGC |
| UBC | 0.21445014 | old | MGC |
| CCN1 | -0.20908362 | old | MGC |
| MTTP | -0.20580158 | old | MGC |
| MED12L | 0.19843048 | old | MGC |
| YBX3 | 0.19595261 | old | MGC |
| RBP1 | -0.19334197 | old | MGC |
| DUSP1 | -0.19082882 | old | MGC |
| RAB1A | 0.1884729 | old | MGC |
| DDIT3 | -0.18457673 | old | MGC |
| GADD45B | -0.18266837 | old | MGC |
| IFITM3 | -0.1809538 | old | MGC |
| DYNC1I1 | 0.17779967 | old | MGC |
| KHDRBS2 | -0.1769813 | old | MGC |
| PMEPA1 | 0.17619266 | old | MGC |
| AL390957.1 | -0.17536826 | old | MGC |
| CDH2 | -0.17447378 | old | MGC |
| LITAF | 0.17322731 | old | MGC |
| MT-ND4L | -0.16778645 | old | MGC |
| SULF1 | -0.16582935 | old | MGC |
| SPON1 | -0.16434218 | old | MGC |
| NAMPT | 0.1639229 | old | MGC |
| JUND | -0.16124411 | old | MGC |
| SCD | -0.1603773 | old | MGC |
| NR4A1 | -0.16009846 | old | MGC |
| FTX | 0.15487336 | old | MGC |
| HLA-A | 0.1426314 | old | MGC |
| HSPA6 | 0.14204451 | old | MGC |
| CD44 | 0.14010204 | old | MGC |
| CLDN1 | 0.13852668 | old | MGC |

| | | | |
|---|---|---|---|
| NRG3 | 0.13434519 | old | MGC |
| HSPE1 | -0.13431026 | old | MGC |
| PMP22 | 0.13378167 | old | MGC |
| COL24A1 | -0.13266416 | old | MGC |
| DNAJB1 | -0.13020413 | old | MGC |
| ABI3BP | 0.12884644 | old | MGC |
| ID1 | 0.12825428 | old | MGC |
| GRID2 | -0.12735702 | old | MGC |
| CXCL2 | 0.12634225 | old | MGC |
| EPHB6 | -0.12547414 | old | MGC |
| SLC3A2 | 0.1243807 | old | MGC |
| ID2 | -0.123736925 | old | MGC |
| PAX6 | 0.123105414 | old | MGC |
| S100A11 | 0.1229473 | old | MGC |
| ILDR2 | -0.12263897 | old | MGC |
| KDR | -0.12138976 | old | MGC |
| SERPINE1 | 0.11999616 | old | MGC |
| NAALADL2 | 0.119713575 | old | MGC |
| MAFF | 0.11904408 | old | MGC |
| S100A10 | 0.11895286 | old | MGC |
| OSBPL10 | 0.117020026 | old | MGC |
| TFRC | 0.11339665 | old | MGC |
| RORB | -0.10712356 | old | MGC |
| MT-ND5 | -0.10708535 | old | MGC |
| GFAP | -0.10609929 | old | MGC |
| ARL4D | 0.106006734 | old | MGC |
| GPRC5A | 0.105204694 | old | MGC |
| EIF1 | -0.10494962 | old | MGC |
| TUBB2B | -0.103842124 | old | MGC |
| CLRN1 | -0.09668896 | old | MGC |
| SLCO4A1-AS1 | 0.094825655 | old | MGC |
| SPSB1 | 0.09440669 | old | MGC |
| PIM1 | 0.09305314 | old | MGC |
| SIPA1L1 | 0.0895263 | old | MGC |
| SDC4 | 0.08850088 | old | MGC |
| PCYT1B | -0.087945834 | old | MGC |
| PHYHIPL | -0.08705388 | old | MGC |
| ZBTB21 | -0.086828314 | old | MGC |
| CYP4V2 | -0.086109854 | old | MGC |
| HLA-C | 0.08343397 | old | MGC |
| MYO10 | -0.08280732 | old | MGC |
| COTL1 | 0.08093218 | old | MGC |
| MAGI2 | 0.08020736 | old | MGC |
| CCL2 | -0.080201976 | old | MGC |
| FOSL2 | 0.07924376 | old | MGC |
| ATF3 | -0.07581367 | old | MGC |
| FBXO32 | 0.07490357 | old | MGC |
| KLKB1 | -0.07413288 | old | MGC |
| INSIG1 | -0.07404623 | old | MGC |
| GPM6A | 0.07373863 | old | MGC |
| S100A6 | -0.06933736 | old | MGC |
| SLC19A2 | 0.06902955 | old | MGC |
| B2M | -0.06892461 | old | MGC |
| MKNK2 | -0.06882993 | old | MGC |

| | | | |
|---|---|---|---|
| SNHG16 | 0.067449555 | old | MGC |
| NFIA | 0.0672737 | old | MGC |
| TRH | 0.06331481 | old | MGC |
| CHMP1B | -0.056838542 | old | MGC |
| CYSTM1 | 0.05594083 | old | MGC |
| FGFBP2 | 0.055472642 | old | MGC |
| ADD1 | 0.055236623 | old | MGC |
| ACTB | -0.05483125 | old | MGC |
| GRIA1 | -0.05463142 | old | MGC |
| SLC7A5 | 0.05167719 | old | MGC |
| SERPING1 | 0.048895255 | old | MGC |
| KLHDC8A | -0.048420474 | old | MGC |
| SDS | 0.046732083 | old | MGC |
| KIAA1217 | 0.046358302 | old | MGC |
| MT-ATP6 | 0.045966312 | old | MGC |
| LRRC4C | 0.04273075 | old | MGC |
| SFXN5 | -0.04249099 | old | MGC |
| NCKAP5 | 0.039667048 | old | MGC |
| SPHK1 | 0.039028615 | old | MGC |
| CNTN1 | -0.03853646 | old | MGC |
| PTPRZ1 | -0.035539694 | old | MGC |
| SFRP2 | -0.034888223 | old | MGC |
| CCDC107 | -0.034450725 | old | MGC |
| AMER2 | 0.033743076 | old | MGC |
| AC092691.1 | 0.03361334 | old | MGC |
| AMACR | -0.033190534 | old | MGC |
| LMCD1 | 0.03167466 | old | MGC |
| ANOS1 | -0.031468146 | old | MGC |
| PDLIM4 | 0.03124394 | old | MGC |
| IER5L | 0.030797577 | old | MGC |
| ID3 | -0.030155065 | old | MGC |
| PPP1R15A | -0.028876815 | old | MGC |
| AQP4-AS1 | -0.028603118 | old | MGC |
| PDLIM3 | 0.024317887 | old | MGC |
| EPCAM | 0.023420919 | old | MGC |
| KLHL8 | 0.022997444 | old | MGC |
| ADGRL3 | -0.022990435 | old | MGC |
| MAP2 | -0.022749882 | old | MGC |
| SOD2 | -0.022414435 | old | MGC |
| MT2A | -0.022406006 | old | MGC |
| EPB41L2 | -0.021314073 | old | MGC |
| NFKBIA | -0.01956406 | old | MGC |
| SRGAP3 | -0.019167451 | old | MGC |
| PCSK2 | -0.016863769 | old | MGC |
| MEGF10 | 0.01676009 | old | MGC |
| PREX2 | -0.015625697 | old | MGC |
| RARRES1 | 0.014685992 | old | MGC |
| BAIAP2 | -0.013218169 | old | MGC |
| CPA6 | 0.012835086 | old | MGC |
| INHBA | -0.012519052 | old | MGC |
| FAP | -0.011083353 | old | MGC |
| HECW2 | -0.010070307 | old | MGC |
| SOX6 | -0.009834558 | old | MGC |
| BHLHE40 | 0.008354692 | old | MGC |

| | | | |
|---|---|---|---|
| HLA-DRB1 | -0.006560494 | old | MGC |
| RRAD | -0.006492203 | old | MGC |
| SORBS2 | 0.005820125 | old | MGC |
| ABCA8 | 0.004711746 | old | MGC |
| CHST9 | -0.003944216 | old | MGC |
| UTRN | -0.001391736 | old | MGC |
| S100A16 | 0.001254006 | old | MGC |
| FUT8 | 0.000754316 | old | MGC |
| RHOU | -0.000316126 | old | MGC |
| MT-ATP8 | -0.000294021 | old | MGC |
| CRYAB | 0 | old | MGC |
| NAV3 | 0 | old | MGC |
| GEM | 0 | old | MGC |
| PHLPP1 | 0 | old | MGC |
| WTAP | 0 | old | MGC |
| ZNF385D | 0 | old | MGC |
| MYO6 | 0 | old | MGC |
| CTTNBP2 | 0 | old | MGC |
| CXCL3 | 0 | old | MGC |
| IER3 | 0 | old | MGC |
| CEBPB | 0 | old | MGC |
| AC004264.1 | 0 | old | MGC |
| DENND11 | 0 | old | MGC |
| IFITM2 | 0 | old | MGC |
| ZNF331 | 0 | old | MGC |
| SOX5 | 0 | old | MGC |
| ARHGAP21 | 0 | old | MGC |
| SEPTIN4 | 0.747544 | young | HC |
| PCSK6 | -0.6100282 | young | HC |
| SNHG14 | -0.5053719 | young | HC |
| DTNA | -0.42230734 | young | HC |
| FTX | -0.42120534 | young | HC |
| ENO2 | -0.4144511 | young | HC |
| TUBA1B | -0.41114622 | young | HC |
| HIST1H4C | 0.40021858 | young | HC |
| SPP1 | 0.39990106 | young | HC |
| PDE4B | 0.30602932 | young | HC |
| UBC | -0.30283663 | young | HC |
| GRIA4 | 0.28392544 | young | HC |
| CRYAB | -0.26998407 | young | HC |
| FRMPD4 | 0.26400626 | young | HC |
| HSPE1 | 0.25955933 | young | HC |
| STMN4 | -0.25528976 | young | HC |
| SNTG2 | -0.23832142 | young | HC |
| FUS | 0.22693777 | young | HC |
| ZNF385D | 0.22377837 | young | HC |
| DCLK1 | -0.20738316 | young | HC |
| PTGFR | -0.20592818 | young | HC |
| RALGAPA2 | -0.19989783 | young | HC |
| PTN | 0.19775191 | young | HC |
| MDGA2 | -0.19137245 | young | HC |
| EPHA5 | -0.18933868 | young | HC |
| CRPPA | 0.17561218 | young | HC |
| NCL | 0.17386752 | young | HC |

| | | | |
|---|---|---|---|
| MT-ND6 | 0.17274284 | young | HC |
| AC093765.2 | 0.16797644 | young | HC |
| TMSB10 | -0.16193897 | young | HC |
| ERO1A | -0.15914741 | young | HC |
| BHLHE40 | -0.15743029 | young | HC |
| HSP90AB1 | -0.15248767 | young | HC |
| DOK6 | -0.14712444 | young | HC |
| KCNJ3 | 0.1461362 | young | HC |
| ONECUT2 | -0.14451206 | young | HC |
| GJA10 | 0.14200291 | young | HC |
| FDFT1 | -0.13949287 | young | HC |
| NPIPB2 | 0.12903506 | young | HC |
| MT2A | -0.12455315 | young | HC |
| HSPH1 | 0.1227409 | young | HC |
| ADARB2 | -0.11711002 | young | HC |
| PLCB4 | -0.11407545 | young | HC |
| GALNT13 | 0.109014496 | young | HC |
| ADCY2 | 0.105593316 | young | HC |
| DOCK3 | 0.10337688 | young | HC |
| TAPT1-AS1 | 0.102720015 | young | HC |
| CUX2 | -0.102070644 | young | HC |
| MARCKS | -0.09879172 | young | HC |
| WDR37 | -0.096067704 | young | HC |
| NDST3 | 0.09436044 | young | HC |
| NRG1 | 0.09332335 | young | HC |
| ARHGAP24 | 0.09300235 | young | HC |
| LRBA | -0.09264572 | young | HC |
| S100A10 | -0.083706796 | young | HC |
| SESTD1 | 0.08168463 | young | HC |
| AC092691.1 | 0.07544872 | young | HC |
| MARCKSL1 | -0.0737911 | young | HC |
| MAPRE2 | -0.07164435 | young | HC |
| PRR16 | 0.071254194 | young | HC |
| MIR181A2HG | -0.06970247 | young | HC |
| IER5L | -0.06953837 | young | HC |
| TARS | 0.069385596 | young | HC |
| GPX3 | -0.063141845 | young | HC |
| MT3 | -0.06295038 | young | HC |
| CCNG2 | -0.062227804 | young | HC |
| SLC4A5 | 0.05828805 | young | HC |
| RD3L | 0.057391565 | young | HC |
| PAM | -0.053825583 | young | HC |
| PGAM1 | -0.053775314 | young | HC |
| HMCES | -0.044254865 | young | HC |
| LHFPL6 | 0.043217544 | young | HC |
| KMT5B | 0.04124416 | young | HC |
| TSPAN8 | -0.039201465 | young | HC |
| DNAJB4 | -0.03873322 | young | HC |
| FAM162A | -0.037865914 | young | HC |
| TRIM36 | -0.034342013 | young | HC |
| ELOVL5 | -0.03230813 | young | HC |
| FHIT | 0.032038268 | young | HC |
| SEPTIN4 | -0.747544 | old | HC |
| PCSK6 | 0.6100282 | old | HC |

| | | | |
|---|---|---|---|
| SNHG14 | 0.5053719 | old | HC |
| DTNA | 0.42230734 | old | HC |
| FTX | 0.42120534 | old | HC |
| ENO2 | 0.4144511 | old | HC |
| TUBA1B | 0.41114622 | old | HC |
| HIST1H4C | -0.40021858 | old | HC |
| SPP1 | -0.39990106 | old | HC |
| PDE4B | -0.30602932 | old | HC |
| UBC | 0.30283663 | old | HC |
| GRIA4 | -0.28392544 | old | HC |
| CRYAB | 0.26998407 | old | HC |
| FRMPD4 | -0.26400626 | old | HC |
| HSPE1 | -0.25955933 | old | HC |
| STMN4 | 0.25528976 | old | HC |
| SNTG2 | 0.23832142 | old | HC |
| FUS | -0.22693777 | old | HC |
| ZNF385D | -0.22377837 | old | HC |
| DCLK1 | 0.20738316 | old | HC |
| PTGFR | 0.20592818 | old | HC |
| RALGAPA2 | 0.19989783 | old | HC |
| PTN | -0.19775191 | old | HC |
| MDGA2 | 0.19137245 | old | HC |
| EPHA5 | 0.18933868 | old | HC |
| CRPPA | -0.17561218 | old | HC |
| NCL | -0.17386752 | old | HC |
| MT-ND6 | -0.17274284 | old | HC |
| AC093765.2 | -0.16797644 | old | HC |
| TMSB10 | 0.16193897 | old | HC |
| ERO1A | 0.15914741 | old | HC |
| BHLHE40 | 0.15743029 | old | HC |
| HSP90AB1 | 0.15248767 | old | HC |
| DOK6 | 0.14712444 | old | HC |
| KCNJ3 | -0.1461362 | old | HC |
| ONECUT2 | 0.14451206 | old | HC |
| GJA10 | -0.14200291 | old | HC |
| FDFT1 | 0.13949287 | old | HC |
| NPIPB2 | -0.12903506 | old | HC |
| MT2A | 0.12455315 | old | HC |
| HSPH1 | -0.1227409 | old | HC |
| ADARB2 | 0.11711002 | old | HC |
| PLCB4 | 0.11407545 | old | HC |
| GALNT13 | -0.109014496 | old | HC |
| ADCY2 | -0.105593316 | old | HC |
| DOCK3 | -0.10337688 | old | HC |
| TAPT1-AS1 | -0.102720015 | old | HC |
| CUX2 | 0.102070644 | old | HC |
| MARCKS | 0.09879172 | old | HC |
| WDR37 | 0.096067704 | old | HC |
| NDST3 | -0.09436044 | old | HC |
| NRG1 | -0.09332335 | old | HC |
| ARHGAP24 | -0.09300235 | old | HC |
| LRBA | 0.09264572 | old | HC |
| S100A10 | 0.083706796 | old | HC |
| SESTD1 | -0.08168463 | old | HC |

| | | | |
|---|---|---|---|
| AC092691.1 | -0.07544872 | old | HC |
| MARCKSL1 | 0.0737911 | old | HC |
| MAPRE2 | 0.07164435 | old | HC |
| PRR16 | -0.071254194 | old | HC |
| MIR181A2HG | 0.06970247 | old | HC |
| IER5L | 0.06953837 | old | HC |
| TARS | -0.069385596 | old | HC |
| GPX3 | 0.063141845 | old | HC |
| MT3 | 0.06295038 | old | HC |
| CCNG2 | 0.062227804 | old | HC |
| SLC4A5 | -0.05828805 | old | HC |
| RD3L | -0.057391565 | old | HC |
| PAM | 0.053825583 | old | HC |
| PGAM1 | 0.053775314 | old | HC |
| HMCES | 0.044254865 | old | HC |
| LHFPL6 | -0.043217544 | old | HC |
| KMT5B | -0.04124416 | old | HC |
| TSPAN8 | 0.039201465 | old | HC |
| DNAJB4 | 0.03873322 | old | HC |
| FAM162A | 0.037865914 | old | HC |
| TRIM36 | 0.034342013 | old | HC |
| ELOVL5 | 0.03230813 | old | HC |
| FHIT | -0.032038268 | old | HC |
| TTR | -0.76097786 | young | cone |
| HSPH1 | 0.6713237 | young | cone |
| PCP4 | 0.5318151 | young | cone |
| RAB17 | -0.49236694 | young | cone |
| AC112206.2 | 0.42647642 | young | cone |
| AL050403.2 | 0.39811492 | young | cone |
| FSTL5 | -0.3872327 | young | cone |
| TXK | -0.37080723 | young | cone |
| ROM1 | 0.3512508 | young | cone |
| RPS26 | -0.3486466 | young | cone |
| RRAD | 0.3206233 | young | cone |
| GABRA2 | -0.31782165 | young | cone |
| AC002460.2 | -0.3169539 | young | cone |
| TMEM108 | 0.31599832 | young | cone |
| MAP2 | 0.29716104 | young | cone |
| BCO2 | -0.2867476 | young | cone |
| THSD7B | -0.28543863 | young | cone |
| ACTG1 | -0.28157365 | young | cone |
| CCSER1 | -0.27906695 | young | cone |
| DNAJB1 | 0.27645686 | young | cone |
| FGF12 | 0.26126692 | young | cone |
| RGS9 | 0.25713572 | young | cone |
| DRD4 | 0.22826447 | young | cone |
| PRKDC | 0.22548185 | young | cone |
| PRUNE2 | -0.20548671 | young | cone |
| FAM153A | 0.20313174 | young | cone |
| SNAP25-AS1 | 0.18867104 | young | cone |
| C9orf16 | 0.18686423 | young | cone |
| USH2A | -0.18134806 | young | cone |
| RAX | 0.18010873 | young | cone |
| BMPR1B | -0.1793643 | young | cone |

| | | | |
|---|---|---|---|
| TMEM141 | 0.17892712 | young | cone |
| SLC22A17 | -0.17699142 | young | cone |
| KCNB1 | 0.17627457 | young | cone |
| DPF3 | -0.17208685 | young | cone |
| AC097662.1 | 0.17013957 | young | cone |
| ARMC9 | -0.1696199 | young | cone |
| ACTB | -0.1670592 | young | cone |
| PDE6C | 0.16529278 | young | cone |
| BAG3 | -0.16413935 | young | cone |
| SERPINH1 | -0.15801029 | young | cone |
| AL357172.1 | -0.14599095 | young | cone |
| SKAP2 | 0.14456214 | young | cone |
| AKAP6 | -0.14389867 | young | cone |
| NCL | 0.14174506 | young | cone |
| TMEM176A | -0.13613337 | young | cone |
| SSX2IP | 0.13378002 | young | cone |
| SNHG14 | -0.1334578 | young | cone |
| TMEM176B | -0.1320176 | young | cone |
| ME3 | -0.13138978 | young | cone |
| AC079467.1 | 0.13114008 | young | cone |
| YPEL3 | -0.1290208 | young | cone |
| PGAM2 | 0.12892939 | young | cone |
| SNX29 | -0.12660876 | young | cone |
| CDH12 | 0.12566347 | young | cone |
| WWOX | 0.122128725 | young | cone |
| LDLRAD4 | -0.12179659 | young | cone |
| AC007349.2 | 0.11730705 | young | cone |
| LRFN5 | -0.10786745 | young | cone |
| PCDH15 | -0.10275979 | young | cone |
| FTX | -0.10127182 | young | cone |
| SLC35F1 | -0.10111235 | young | cone |
| KIF2A | 0.10068588 | young | cone |
| AC110992.1 | -0.10037729 | young | cone |
| AC116903.2 | -0.09887497 | young | cone |
| STPG2 | -0.09859419 | young | cone |
| AC137770.1 | 0.09282182 | young | cone |
| SAMD7 | 0.0875669 | young | cone |
| AC117453.1 | -0.08583759 | young | cone |
| PEX5L-AS2 | 0.08183348 | young | cone |
| AL121821.2 | -0.07677925 | young | cone |
| POLQ | 0.07626912 | young | cone |
| ROBO2 | 0.0718725 | young | cone |
| H1FX | -0.06858182 | young | cone |
| DLGAP2 | -0.06666222 | young | cone |
| FAM135B | 0.06027515 | young | cone |
| DSE | 0.05905722 | young | cone |
| AC092939.1 | 0.057163227 | young | cone |
| DLG2 | 0.05458807 | young | cone |
| PCAT4 | 0.053368706 | young | cone |
| AL357153.2 | -0.047865946 | young | cone |
| WFDC11 | 0.045934193 | young | cone |
| TRAF3IP1 | 0.03837493 | young | cone |
| PDE5A | -0.036505274 | young | cone |
| AC106798.1 | 0.03641248 | young | cone |

| | | | |
|---|---|---|---|
| CADM1 | -0.035196602 | young | cone |
| ADAMTSL1 | -0.033818424 | young | cone |
| CCNJL | 0.03350446 | young | cone |
| LRRFIP1 | -0.03321365 | young | cone |
| HNRNPH1 | -0.031212628 | young | cone |
| SLC38A1 | -0.030760417 | young | cone |
| AGBL4 | 0.026322225 | young | cone |
| AC068051.1 | 0.0248105 | young | cone |
| EEF1A2 | 0.018411772 | young | cone |
| MT-ND6 | 0.017312707 | young | cone |
| MT2A | -0.016923163 | young | cone |
| ZBTB20 | -0.008894019 | young | cone |
| KLF7 | -0.008120529 | young | cone |
| VAX2 | 0.002668804 | young | cone |
| AL118516.1 | -0.002140158 | young | cone |
| ZNF292 | -0.000660552 | young | cone |
| ZNF529 | -0.000233076 | young | cone |
| GPX3 | 0 | young | cone |
| C1QTNF4 | 0 | young | cone |
| GLUL | 0 | young | cone |
| NUBPL | 0 | young | cone |
| PRSS51 | 0 | young | cone |
| LINC00871 | 0 | young | cone |
| TCEAL9 | 0 | young | cone |
| MFSD4B | 0 | young | cone |
| MRLN | 0 | young | cone |
| AP000820.2 | 0 | young | cone |
| CLUL1 | 0 | young | cone |
| SDHAF3 | 0 | young | cone |
| MAP3K7CL | 0 | young | cone |
| DYRK4 | 0 | young | cone |
| AL161716.1 | 0 | young | cone |
| ARHGAP24 | 0 | young | cone |
| HLA-A | 0 | young | cone |
| LINC01184 | 0 | young | cone |
| AC007325.4 | 0 | young | cone |
| ST3GAL3 | 0 | young | cone |
| FAM172A | 0 | young | cone |
| WAKMAR2 | 0 | young | cone |
| MAGI2 | 0 | young | cone |
| MAP1LC3A | 0 | young | cone |
| LINGO2 | 0 | young | cone |
| FBXL17 | 0 | young | cone |
| EIF4G2 | 0 | young | cone |
| LINC02343 | 0 | young | cone |
| UCMA | 0 | young | cone |
| IFITM3 | 0 | young | cone |
| MBP | 0 | young | cone |
| HSPA1A | 0 | young | cone |
| HSPA1B | 0 | young | cone |
| CST3 | 0 | young | cone |
| HLA-C | 0 | young | cone |
| KIAA0825 | 0 | young | cone |
| NAALADL2 | 0 | young | cone |

| | | | |
|---|---|---|---|
| SGCD | 0 | young | cone |
| PTPRD | 0 | young | cone |
| EYS | 0 | young | cone |
| UPP2 | 0 | young | cone |
| LGALS3 | 0 | young | cone |
| GAS7 | 0 | young | cone |
| IMMP2L | 0 | young | cone |
| HCG17 | 0 | young | cone |
| SBSPON | 0 | young | cone |
| UPK3BL1 | 0 | young | cone |
| HMGB2 | 0 | young | cone |
| SYCE1L | 0 | young | cone |
| PRR16 | 0 | young | cone |
| COTL1 | 0 | young | cone |
| RCBTB1 | 0 | young | cone |
| KIRREL1 | 0 | young | cone |
| MIR2052HG | 0 | young | cone |
| ATP2B4 | 0 | young | cone |
| LGALS3BP | 0 | young | cone |
| TF | 0 | young | cone |
| VPS53 | 0 | young | cone |
| SSBP4 | 0 | young | cone |
| SMYD3 | 0 | young | cone |
| MESP1 | 0 | young | cone |
| NTM | 0 | young | cone |
| AHSA1 | 0 | young | cone |
| MNDA | 0 | young | cone |
| PLXDC1 | 0 | young | cone |
| CREG2 | 0 | young | cone |
| CRYAB | 0 | young | cone |
| HSBP1 | 0 | young | cone |
| FAM166C | 0 | young | cone |
| HLA-DRB5 | 0 | young | cone |
| HPRT1 | 0 | young | cone |
| AL137804.1 | 0 | young | cone |
| CCNO | 0 | young | cone |
| SLC4A8 | 0 | young | cone |
| SOX2-OT | 0 | young | cone |
| NRN1L | 0 | young | cone |
| SNAP91 | 0 | young | cone |
| A2M | 0 | young | cone |
| GNGT1 | 0 | young | cone |
| POLH | 0 | young | cone |
| PRELID2 | 0 | young | cone |
| HAR1A | 0 | young | cone |
| PRKN | 0 | young | cone |
| ANTXR2 | 0 | young | cone |
| DMD | 0 | young | cone |
| EFNA5 | 0 | young | cone |
| ZNF225 | 0 | young | cone |
| AC004540.2 | 0 | young | cone |
| EPCAM | 0 | young | cone |
| STMN1 | 0 | young | cone |
| PCAT1 | 0 | young | cone |

| | | | |
|---|---|---|---|
| STK33 | 0 | young | cone |
| STK24 | 0 | young | cone |
| GNG12-AS1 | 0 | young | cone |
| TTR | 0.76097786 | old | cone |
| HSPH1 | -0.6713237 | old | cone |
| PCP4 | -0.5318151 | old | cone |
| RAB17 | 0.49236694 | old | cone |
| AC112206.2 | -0.42647642 | old | cone |
| AL050403.2 | -0.39811492 | old | cone |
| FSTL5 | 0.3872327 | old | cone |
| TXK | 0.37080723 | old | cone |
| ROM1 | -0.3512508 | old | cone |
| RPS26 | 0.3486466 | old | cone |
| RRAD | -0.3206233 | old | cone |
| GABRA2 | 0.31782165 | old | cone |
| AC002460.2 | 0.3169539 | old | cone |
| TMEM108 | -0.31599832 | old | cone |
| MAP2 | -0.29716104 | old | cone |
| BCO2 | 0.2867476 | old | cone |
| THSD7B | 0.28543863 | old | cone |
| ACTG1 | 0.28157365 | old | cone |
| CCSER1 | 0.27906695 | old | cone |
| DNAJB1 | -0.27645686 | old | cone |
| FGF12 | -0.26126692 | old | cone |
| RGS9 | -0.25713572 | old | cone |
| DRD4 | -0.22826447 | old | cone |
| PRKDC | -0.22548185 | old | cone |
| PRUNE2 | 0.20548671 | old | cone |
| FAM153A | -0.20313174 | old | cone |
| SNAP25-AS1 | -0.18867104 | old | cone |
| C9orf16 | -0.18686423 | old | cone |
| USH2A | 0.18134806 | old | cone |
| RAX | -0.18010873 | old | cone |
| BMPR1B | 0.1793643 | old | cone |
| TMEM141 | -0.17892712 | old | cone |
| SLC22A17 | 0.17699142 | old | cone |
| KCNB1 | -0.17627457 | old | cone |
| DPF3 | 0.17208685 | old | cone |
| AC097662.1 | -0.17013957 | old | cone |
| ARMC9 | 0.1696199 | old | cone |
| ACTB | 0.1670592 | old | cone |
| PDE6C | -0.16529278 | old | cone |
| BAG3 | 0.16413935 | old | cone |
| SERPINH1 | 0.15801029 | old | cone |
| AL357172.1 | 0.14599095 | old | cone |
| SKAP2 | -0.14456214 | old | cone |
| AKAP6 | 0.14389867 | old | cone |
| NCL | -0.14174506 | old | cone |
| TMEM176A | 0.13613337 | old | cone |
| SSX2IP | -0.13378002 | old | cone |
| SNHG14 | 0.1334578 | old | cone |
| TMEM176B | 0.1320176 | old | cone |
| ME3 | 0.13138978 | old | cone |
| AC079467.1 | -0.13114008 | old | cone |

| | | | |
|---|---|---|---|
| YPEL3 | 0.1290208 | old | cone |
| PGAM2 | -0.12892939 | old | cone |
| SNX29 | 0.12660876 | old | cone |
| CDH12 | -0.12566347 | old | cone |
| WWOX | -0.122128725 | old | cone |
| LDLRAD4 | 0.12179659 | old | cone |
| AC007349.2 | -0.11730705 | old | cone |
| LRFN5 | 0.10786745 | old | cone |
| PCDH15 | 0.10275979 | old | cone |
| FTX | 0.10127182 | old | cone |
| SLC35F1 | 0.10111235 | old | cone |
| KIF2A | -0.10068588 | old | cone |
| AC110992.1 | 0.10037729 | old | cone |
| AC116903.2 | 0.09887497 | old | cone |
| STPG2 | 0.09859419 | old | cone |
| AC137770.1 | -0.09282182 | old | cone |
| SAMD7 | -0.0875669 | old | cone |
| AC117453.1 | 0.08583759 | old | cone |
| PEX5L-AS2 | -0.08183348 | old | cone |
| AL121821.2 | 0.07677925 | old | cone |
| POLQ | -0.07626912 | old | cone |
| ROBO2 | -0.0718725 | old | cone |
| H1FX | 0.06858182 | old | cone |
| DLGAP2 | 0.06666222 | old | cone |
| FAM135B | -0.06027515 | old | cone |
| DSE | -0.05905722 | old | cone |
| AC092939.1 | -0.057163227 | old | cone |
| DLG2 | -0.05458807 | old | cone |
| PCAT4 | -0.053368706 | old | cone |
| AL357153.2 | 0.047865946 | old | cone |
| WFDC11 | -0.045934193 | old | cone |
| TRAF3IP1 | -0.03837493 | old | cone |
| PDE5A | 0.036505274 | old | cone |
| AC106798.1 | -0.03641248 | old | cone |
| CADM1 | 0.035196602 | old | cone |
| ADAMTSL1 | 0.033818424 | old | cone |
| CCNJL | -0.03350446 | old | cone |
| LRRFIP1 | 0.03321365 | old | cone |
| HNRNPH1 | 0.031212628 | old | cone |
| SLC38A1 | 0.030760417 | old | cone |
| AGBL4 | -0.026322225 | old | cone |
| AC068051.1 | -0.0248105 | old | cone |
| EEF1A2 | -0.018411772 | old | cone |
| MT-ND6 | -0.017312707 | old | cone |
| MT2A | 0.016923163 | old | cone |
| ZBTB20 | 0.008894019 | old | cone |
| KLF7 | 0.008120529 | old | cone |
| VAX2 | -0.002668804 | old | cone |
| AL118516.1 | 0.002140158 | old | cone |
| ZNF292 | 0.000660552 | old | cone |
| ZNF529 | 0.000233076 | old | cone |
| GPX3 | 0 | old | cone |
| C1QTNF4 | 0 | old | cone |
| GLUL | 0 | old | cone |

| | | | |
|---|---|---|---|
| NUBPL | 0 | old | cone |
| PRSS51 | 0 | old | cone |
| LINC00871 | 0 | old | cone |
| TCEAL9 | 0 | old | cone |
| MFSD4B | 0 | old | cone |
| MRLN | 0 | old | cone |
| AP000820.2 | 0 | old | cone |
| CLUL1 | 0 | old | cone |
| SDHAF3 | 0 | old | cone |
| MAP3K7CL | 0 | old | cone |
| DYRK4 | 0 | old | cone |
| AL161716.1 | 0 | old | cone |
| ARHGAP24 | 0 | old | cone |
| HLA-A | 0 | old | cone |
| LINC01184 | 0 | old | cone |
| AC007325.4 | 0 | old | cone |
| ST3GAL3 | 0 | old | cone |
| FAM172A | 0 | old | cone |
| WAKMAR2 | 0 | old | cone |
| MAGI2 | 0 | old | cone |
| MAP1LC3A | 0 | old | cone |
| LINGO2 | 0 | old | cone |
| FBXL17 | 0 | old | cone |
| EIF4G2 | 0 | old | cone |
| LINC02343 | 0 | old | cone |
| UCMA | 0 | old | cone |
| IFITM3 | 0 | old | cone |
| MBP | 0 | old | cone |
| HSPA1A | 0 | old | cone |
| HSPA1B | 0 | old | cone |
| CST3 | 0 | old | cone |
| HLA-C | 0 | old | cone |
| KIAA0825 | 0 | old | cone |
| NAALADL2 | 0 | old | cone |
| SGCD | 0 | old | cone |
| PTPRD | 0 | old | cone |
| EYS | 0 | old | cone |
| UPP2 | 0 | old | cone |
| LGALS3 | 0 | old | cone |
| GAS7 | 0 | old | cone |
| IMMP2L | 0 | old | cone |
| HCG17 | 0 | old | cone |
| SBSPON | 0 | old | cone |
| UPK3BL1 | 0 | old | cone |
| HMGB2 | 0 | old | cone |
| SYCE1L | 0 | old | cone |
| PRR16 | 0 | old | cone |
| COTL1 | 0 | old | cone |
| RCBTB1 | 0 | old | cone |
| KIRREL1 | 0 | old | cone |
| MIR2052HG | 0 | old | cone |
| ATP2B4 | 0 | old | cone |
| LGALS3BP | 0 | old | cone |
| TF | 0 | old | cone |

| | | | |
|---|---|---|---|
| VPS53 | 0 | old | cone |
| SSBP4 | 0 | old | cone |
| SMYD3 | 0 | old | cone |
| MESP1 | 0 | old | cone |
| NTM | 0 | old | cone |
| AHSA1 | 0 | old | cone |
| MNDA | 0 | old | cone |
| PLXDC1 | 0 | old | cone |
| CREG2 | 0 | old | cone |
| CRYAB | 0 | old | cone |
| HSBP1 | 0 | old | cone |
| FAM166C | 0 | old | cone |
| HLA-DRB5 | 0 | old | cone |
| HPRT1 | 0 | old | cone |
| AL137804.1 | 0 | old | cone |
| CCNO | 0 | old | cone |
| SLC4A8 | 0 | old | cone |
| SOX2-OT | 0 | old | cone |
| NRN1L | 0 | old | cone |
| SNAP91 | 0 | old | cone |
| A2M | 0 | old | cone |
| GNGT1 | 0 | old | cone |
| POLH | 0 | old | cone |
| PRELID2 | 0 | old | cone |
| HAR1A | 0 | old | cone |
| PRKN | 0 | old | cone |
| ANTXR2 | 0 | old | cone |
| DMD | 0 | old | cone |
| EFNA5 | 0 | old | cone |
| ZNF225 | 0 | old | cone |
| AC004540.2 | 0 | old | cone |
| EPCAM | 0 | old | cone |
| STMN1 | 0 | old | cone |
| PCAT1 | 0 | old | cone |
| STK33 | 0 | old | cone |
| STK24 | 0 | old | cone |
| GNG12-AS1 | 0 | old | cone |
| MT-CO3 | -1.8344841 | young | BC |
| MT-CYB | 1.2354496 | young | BC |
| MT-CO2 | 0.869576 | young | BC |
| MT-ATP6 | -0.7251681 | young | BC |
| MT-ND2 | 0.7033883 | young | BC |
| MT-CO1 | -0.6645446 | young | BC |
| MT-ND3 | 0.6545998 | young | BC |
| MT-ND5 | 0.48423472 | young | BC |
| CDR1 | 0.4719102 | young | BC |
| TUBA1B | -0.45716298 | young | BC |
| CALM2 | -0.4475544 | young | BC |
| MT-ND6 | 0.44552824 | young | BC |
| CCNI | -0.4068531 | young | BC |
| AL033504.1 | -0.389989 | young | BC |
| PCP2 | 0.38390744 | young | BC |
| TMSB4X | -0.33285493 | young | BC |
| TRPM1 | -0.3287938 | young | BC |

| | | | |
|---|---|---|---|
| LINC02649 | -0.30318755 | young | BC |
| P4HA1 | -0.2999534 | young | BC |
| BHLHE40 | -0.28671256 | young | BC |
| MT-ND1 | 0.2859944 | young | BC |
| GPX3 | -0.28552735 | young | BC |
| EID1 | 0.28474548 | young | BC |
| DTNA | -0.24479589 | young | BC |
| MRPL57 | 0.23518929 | young | BC |
| CST3 | -0.2345133 | young | BC |
| CLASP2 | -0.2307754 | young | BC |
| ITM2C | -0.2306397 | young | BC |
| SUMO2 | -0.2297499 | young | BC |
| TF | -0.22827214 | young | BC |
| GNB3 | -0.22465678 | young | BC |
| SDK1 | -0.21957438 | young | BC |
| MT-ATP8 | 0.21926823 | young | BC |
| NPVF | -0.21827525 | young | BC |
| AK4 | -0.21564957 | young | BC |
| NTNG1 | 0.20230952 | young | BC |
| MRFAP1 | 0.20159455 | young | BC |
| NDUFB1 | 0.20131788 | young | BC |
| CHCHD2 | -0.19716503 | young | BC |
| LRTM1 | 0.18301827 | young | BC |
| TTYH1 | 0.18180606 | young | BC |
| NDUFB6 | 0.18136296 | young | BC |
| NCL | 0.17362662 | young | BC |
| KMT2E | -0.17047824 | young | BC |
| KCNH5 | 0.16986147 | young | BC |
| LRRTM4 | -0.16922367 | young | BC |
| COX4I1 | -0.16877745 | young | BC |
| AP001825.1 | 0.16158648 | young | BC |
| NDUFAB1 | 0.16083089 | young | BC |
| SAP18 | 0.1607517 | young | BC |
| SH3GL3 | -0.15863357 | young | BC |
| AC104117.3 | 0.15837052 | young | BC |
| SPCS1 | -0.15607378 | young | BC |
| POLR2K | 0.15418676 | young | BC |
| PSMB1 | -0.15134896 | young | BC |
| PSMA7 | 0.1490666 | young | BC |
| CRYAB | -0.14838085 | young | BC |
| GNG13 | -0.14513886 | young | BC |
| CCT5 | 0.14462315 | young | BC |
| SRP14 | -0.14226533 | young | BC |
| PRDX5 | 0.1399659 | young | BC |
| NDUFB7 | 0.13771911 | young | BC |
| BHLHE41 | -0.1375924 | young | BC |
| AC010478.1 | -0.1363852 | young | BC |
| SELENOW | 0.13597804 | young | BC |
| ERH | -0.13412856 | young | BC |
| PSMB6 | 0.13320608 | young | BC |
| VBP1 | 0.13293396 | young | BC |
| ANOS1 | 0.1314168 | young | BC |
| PRKCA | -0.13110001 | young | BC |
| NDUFA1 | 0.12987347 | young | BC |

| | | | |
|---|---|---|---|
| TRNP1 | -0.1287994 | young | BC |
| FMC1 | 0.12760101 | young | BC |
| CA10 | -0.12373522 | young | BC |
| NDUFB9 | 0.12355499 | young | BC |
| RPS19BP1 | 0.11861054 | young | BC |
| NDUFC2 | 0.11591643 | young | BC |
| AC092155.1 | 0.11578644 | young | BC |
| CRYBG3 | 0.115229055 | young | BC |
| CRY2 | -0.11348402 | young | BC |
| NDUFS6 | 0.11322921 | young | BC |
| COX17 | 0.11190877 | young | BC |
| VSTM2B | 0.11143477 | young | BC |
| PDK1 | -0.11082084 | young | BC |
| NDUFB11 | 0.10949134 | young | BC |
| C16orf74 | 0.10846216 | young | BC |
| C18orf32 | 0.10711721 | young | BC |
| AC090825.1 | -0.10340376 | young | BC |
| CCNG2 | -0.100648336 | young | BC |
| DOK6 | -0.10027259 | young | BC |
| CPD | -0.09715994 | young | BC |
| COPS9 | 0.096482046 | young | BC |
| SLC6A6 | -0.09429163 | young | BC |
| AP000857.2 | 0.093998514 | young | BC |
| HNRNPA2B1 | 0.09353084 | young | BC |
| KRT222 | 0.09291019 | young | BC |
| NME1 | 0.09247131 | young | BC |
| RERE | -0.09113323 | young | BC |
| PDZRN4 | -0.08906136 | young | BC |
| NDUFA2 | 0.08712696 | young | BC |
| PSMA2 | 0.08470141 | young | BC |
| NDUFS5 | 0.084497325 | young | BC |
| AC119868.2 | 0.081721984 | young | BC |
| SREBF2 | 0.08166971 | young | BC |
| C1QBP | 0.08158688 | young | BC |
| RNASEK | 0.081128664 | young | BC |
| AC091938.1 | 0.07986452 | young | BC |
| H3F3A | 0.077620275 | young | BC |
| PFKFB4 | -0.07734401 | young | BC |
| PPIA | -0.076919585 | young | BC |
| PRKCA-AS1 | -0.07595941 | young | BC |
| NLGN4X | 0.07503009 | young | BC |
| FIS1 | 0.073842146 | young | BC |
| ATP5MC3 | -0.07349491 | young | BC |
| SRSF9 | 0.07072271 | young | BC |
| SEMA3E | 0.068198726 | young | BC |
| LINC02055 | 0.0670296 | young | BC |
| SLIRP | 0.06436074 | young | BC |
| CALM1 | 0.06272212 | young | BC |
| CCDC136 | 0.061689433 | young | BC |
| SMDT1 | 0.06123644 | young | BC |
| COX6A1 | -0.061230354 | young | BC |
| COX6C | 0.06092706 | young | BC |
| GRM5 | 0.06058758 | young | BC |
| SOD1 | -0.060476977 | young | BC |

| | | | |
|---|---|---|---|
| NEDD8 | 0.060420144 | young | BC |
| GSTO1 | 0.05244259 | young | BC |
| FKBP3 | 0.050899208 | young | BC |
| ATP5PF | -0.049955036 | young | BC |
| ZFHX2 | 0.04992949 | young | BC |
| HSPB1 | 0.0497134 | young | BC |
| LSM4 | 0.04930681 | young | BC |
| TXN | -0.047603555 | young | BC |
| PSMB3 | -0.047379114 | young | BC |
| CCT6A | 0.047078602 | young | BC |
| STARD4 | 0.046234947 | young | BC |
| ASIC2 | -0.04612101 | young | BC |
| CCT3 | -0.045823395 | young | BC |
| YWHAE | -0.045302596 | young | BC |
| TAFA3 | 0.043405533 | young | BC |
| FUS | 0.041711558 | young | BC |
| NIF3L1 | -0.04130811 | young | BC |
| RPS17 | 0.04091356 | young | BC |
| SET | 0.040102486 | young | BC |
| PNN | -0.038143087 | young | BC |
| PARK7 | 0.038139038 | young | BC |
| MICOS10 | 0.03789793 | young | BC |
| COX8A | 0.037645843 | young | BC |
| DST | -0.037365332 | young | BC |
| DDC | -0.036788072 | young | BC |
| BTF3 | -0.03626301 | young | BC |
| COX7A2 | -0.035749175 | young | BC |
| HSPA1A | 0.035355892 | young | BC |
| SF3B5 | 0.034758724 | young | BC |
| CNTN5 | 0.032855757 | young | BC |
| H2AFZ | 0.03242473 | young | BC |
| TBCB | -0.03182141 | young | BC |
| AL513164.1 | 0.031782128 | young | BC |
| NDUFB4 | 0.024389332 | young | BC |
| ATP5MC1 | 0.023701573 | young | BC |
| PAIP2 | -0.018890085 | young | BC |
| NQO1 | -0.018377729 | young | BC |
| APOBEC2 | -0.017997427 | young | BC |
| YBX1 | -0.017969504 | young | BC |
| PEBP1 | -0.017427726 | young | BC |
| ATP6V1G1 | -0.01723942 | young | BC |
| SNAP25-AS1 | 0.015903149 | young | BC |
| TAX1BP1 | -0.015145406 | young | BC |
| CACNA2D1 | -0.01407451 | young | BC |
| HSPA6 | 0.013855033 | young | BC |
| CXCL14 | -0.013500354 | young | BC |
| TDRG1 | 0.013487465 | young | BC |
| FAM138C | 0.012059798 | young | BC |
| BEX1 | 0.011297806 | young | BC |
| PNKD | -0.010104606 | young | BC |
| ATP6V0B | -0.008934259 | young | BC |
| AC117944.1 | -0.007872626 | young | BC |
| ATP5F1C | -0.007657462 | young | BC |
| ANK2 | -0.00736347 | young | BC |

| | | | |
|---|---|---|---|
| ANKS1B | -0.006979622 | young | BC |
| MT-CO3 | 1.8344841 | old | BC |
| MT-CYB | -1.2354496 | old | BC |
| MT-CO2 | -0.869576 | old | BC |
| MT-ATP6 | 0.7251681 | old | BC |
| MT-ND2 | -0.7033883 | old | BC |
| MT-CO1 | 0.6645446 | old | BC |
| MT-ND3 | -0.6545998 | old | BC |
| MT-ND5 | -0.48423472 | old | BC |
| CDR1 | -0.4719102 | old | BC |
| TUBA1B | 0.45716298 | old | BC |
| CALM2 | 0.4475544 | old | BC |
| MT-ND6 | -0.44552824 | old | BC |
| CCNI | 0.4068531 | old | BC |
| AL033504.1 | 0.389989 | old | BC |
| PCP2 | -0.38390744 | old | BC |
| TMSB4X | 0.33285493 | old | BC |
| TRPM1 | 0.3287938 | old | BC |
| LINC02649 | 0.30318755 | old | BC |
| P4HA1 | 0.2999534 | old | BC |
| BHLHE40 | 0.28671256 | old | BC |
| MT-ND1 | -0.2859944 | old | BC |
| GPX3 | 0.28552735 | old | BC |
| EID1 | -0.28474548 | old | BC |
| DTNA | 0.24479589 | old | BC |
| MRPL57 | -0.23518929 | old | BC |
| CST3 | 0.2345133 | old | BC |
| CLASP2 | 0.2307754 | old | BC |
| ITM2C | 0.2306397 | old | BC |
| SUMO2 | 0.2297499 | old | BC |
| TF | 0.22827214 | old | BC |
| GNB3 | 0.22465678 | old | BC |
| SDK1 | 0.21957438 | old | BC |
| MT-ATP8 | -0.21926823 | old | BC |
| NPVF | 0.21827525 | old | BC |
| AK4 | 0.21564957 | old | BC |
| NTNG1 | -0.20230952 | old | BC |
| MRFAP1 | -0.20159455 | old | BC |
| NDUFB1 | -0.20131788 | old | BC |
| CHCHD2 | 0.19716503 | old | BC |
| LRTM1 | -0.18301827 | old | BC |
| TTYH1 | -0.18180606 | old | BC |
| NDUFB6 | -0.18136296 | old | BC |
| NCL | -0.17362662 | old | BC |
| KMT2E | 0.17047824 | old | BC |
| KCNH5 | -0.16986147 | old | BC |
| LRRTM4 | 0.16922367 | old | BC |
| COX4I1 | 0.16877745 | old | BC |
| AP001825.1 | -0.16158648 | old | BC |
| NDUFAB1 | -0.16083089 | old | BC |
| SAP18 | -0.1607517 | old | BC |
| SH3GL3 | 0.15863357 | old | BC |
| AC104117.3 | -0.15837052 | old | BC |
| SPCS1 | 0.15607378 | old | BC |

| | | | |
|---|---|---|---|
| POLR2K | -0.15418676 | old | BC |
| PSMB1 | 0.15134896 | old | BC |
| PSMA7 | -0.1490666 | old | BC |
| CRYAB | 0.14838085 | old | BC |
| GNG13 | 0.14513886 | old | BC |
| CCT5 | -0.14462315 | old | BC |
| SRP14 | 0.14226533 | old | BC |
| PRDX5 | -0.1399659 | old | BC |
| NDUFB7 | -0.13771911 | old | BC |
| BHLHE41 | 0.1375924 | old | BC |
| AC010478.1 | 0.1363852 | old | BC |
| SELENOW | -0.13597804 | old | BC |
| ERH | 0.13412856 | old | BC |
| PSMB6 | -0.13320608 | old | BC |
| VBP1 | -0.13293396 | old | BC |
| ANOS1 | -0.1314168 | old | BC |
| PRKCA | 0.13110001 | old | BC |
| NDUFA1 | -0.12987347 | old | BC |
| TRNP1 | 0.1287994 | old | BC |
| FMC1 | -0.12760101 | old | BC |
| CA10 | 0.12373522 | old | BC |
| NDUFB9 | -0.12355499 | old | BC |
| RPS19BP1 | -0.11861054 | old | BC |
| NDUFC2 | -0.11591643 | old | BC |
| AC092155.1 | -0.11578644 | old | BC |
| CRYBG3 | -0.115229055 | old | BC |
| CRY2 | 0.11348402 | old | BC |
| NDUFS6 | -0.11322921 | old | BC |
| COX17 | -0.11190877 | old | BC |
| VSTM2B | -0.11143477 | old | BC |
| PDK1 | 0.11082084 | old | BC |
| NDUFB11 | -0.10949134 | old | BC |
| C16orf74 | -0.10846216 | old | BC |
| C18orf32 | -0.10711721 | old | BC |
| AC090825.1 | 0.10340376 | old | BC |
| CCNG2 | 0.100648336 | old | BC |
| DOK6 | 0.10027259 | old | BC |
| CPD | 0.09715994 | old | BC |
| COPS9 | -0.096482046 | old | BC |
| SLC6A6 | 0.09429163 | old | BC |
| AP000857.2 | -0.093998514 | old | BC |
| HNRNPA2B1 | -0.09353084 | old | BC |
| KRT222 | -0.09291019 | old | BC |
| NME1 | -0.09247131 | old | BC |
| RERE | 0.09113323 | old | BC |
| PDZRN4 | 0.08906136 | old | BC |
| NDUFA2 | -0.08712696 | old | BC |
| PSMA2 | -0.08470141 | old | BC |
| NDUFS5 | -0.084497325 | old | BC |
| AC119868.2 | -0.081721984 | old | BC |
| SREBF2 | -0.08166971 | old | BC |
| C1QBP | -0.08158688 | old | BC |
| RNASEK | -0.081128664 | old | BC |
| AC091938.1 | -0.07986452 | old | BC |

| | | | |
|---|---|---|---|
| H3F3A | -0.077620275 | old | BC |
| PFKFB4 | 0.07734401 | old | BC |
| PPIA | 0.076919585 | old | BC |
| PRKCA-AS1 | 0.07595941 | old | BC |
| NLGN4X | -0.07503009 | old | BC |
| FIS1 | -0.073842146 | old | BC |
| ATP5MC3 | 0.07349491 | old | BC |
| SRSF9 | -0.07072271 | old | BC |
| SEMA3E | -0.068198726 | old | BC |
| LINC02055 | -0.0670296 | old | BC |
| SLIRP | -0.06436074 | old | BC |
| CALM1 | -0.06272212 | old | BC |
| CCDC136 | -0.061689433 | old | BC |
| SMDT1 | -0.06123644 | old | BC |
| COX6A1 | 0.061230354 | old | BC |
| COX6C | -0.06092706 | old | BC |
| GRM5 | -0.06058758 | old | BC |
| SOD1 | 0.060476977 | old | BC |
| NEDD8 | -0.060420144 | old | BC |
| GSTO1 | -0.05244259 | old | BC |
| FKBP3 | -0.050899208 | old | BC |
| ATP5PF | 0.049955036 | old | BC |
| ZFHX2 | -0.04992949 | old | BC |
| HSPB1 | -0.0497134 | old | BC |
| LSM4 | -0.04930681 | old | BC |
| TXN | 0.047603555 | old | BC |
| PSMB3 | 0.047379114 | old | BC |
| CCT6A | -0.047078602 | old | BC |
| STARD4 | -0.046234947 | old | BC |
| ASIC2 | 0.04612101 | old | BC |
| CCT3 | 0.045823395 | old | BC |
| YWHAE | 0.045302596 | old | BC |
| TAFA3 | -0.043405533 | old | BC |
| FUS | -0.041711558 | old | BC |
| NIF3L1 | 0.04130811 | old | BC |
| RPS17 | -0.04091356 | old | BC |
| SET | -0.040102486 | old | BC |
| PNN | 0.038143087 | old | BC |
| PARK7 | -0.038139038 | old | BC |
| MICOS10 | -0.03789793 | old | BC |
| COX8A | -0.037645843 | old | BC |
| DST | 0.037365332 | old | BC |
| DDC | 0.036788072 | old | BC |
| BTF3 | 0.03626301 | old | BC |
| COX7A2 | 0.035749175 | old | BC |
| HSPA1A | -0.035355892 | old | BC |
| SF3B5 | -0.034758724 | old | BC |
| CNTN5 | -0.032855757 | old | BC |
| H2AFZ | -0.03242473 | old | BC |
| TBCB | 0.03182141 | old | BC |
| AL513164.1 | -0.031782128 | old | BC |
| NDUFB4 | -0.024389332 | old | BC |
| ATP5MC1 | -0.023701573 | old | BC |
| PAIP2 | 0.018890085 | old | BC |

| | | | |
|---|---|---|---|
| NQO1 | 0.018377729 | old | BC |
| APOBEC2 | 0.017997427 | old | BC |
| YBX1 | 0.017969504 | old | BC |
| PEBP1 | 0.017427726 | old | BC |
| ATP6V1G1 | 0.01723942 | old | BC |
| SNAP25-AS1 | -0.015903149 | old | BC |
| TAX1BP1 | 0.015145406 | old | BC |
| CACNA2D1 | 0.01407451 | old | BC |
| HSPA6 | -0.013855033 | old | BC |
| CXCL14 | 0.013500354 | old | BC |
| TDRG1 | -0.013487465 | old | BC |
| FAM138C | -0.012059798 | old | BC |
| BEX1 | -0.011297806 | old | BC |
| PNKD | 0.010104606 | old | BC |
| ATP6V0B | 0.008934259 | old | BC |
| AC117944.1 | 0.007872626 | old | BC |
| ATP5F1C | 0.007657462 | old | BC |
| ANK2 | 0.00736347 | old | BC |
| ANKS1B | 0.006979622 | old | BC |
| HP | -1.8225088 | young | astrocyte |
| SLPI | -1.4090822 | young | astrocyte |
| MT1X | -1.3338946 | young | astrocyte |
| MT-ATP8 | 1.1117169 | young | astrocyte |
| MT-CO3 | -1.1048037 | young | astrocyte |
| MT-CO2 | 1.1037254 | young | astrocyte |
| ERO1A | -1.0387514 | young | astrocyte |
| MT2A | 0.8424649 | young | astrocyte |
| SELENOM | -0.7595936 | young | astrocyte |
| LDHA | -0.69253755 | young | astrocyte |
| HILPDA | -0.6796659 | young | astrocyte |
| ANGPTL1 | 0.6486343 | young | astrocyte |
| TM7SF2 | 0.59906185 | young | astrocyte |
| SEMA6D | 0.5521946 | young | astrocyte |
| VEGFA | 0.45605493 | young | astrocyte |
| HP | 1.8225088 | old | astrocyte |
| SLPI | 1.4090822 | old | astrocyte |
| MT1X | 1.3338946 | old | astrocyte |
| MT-ATP8 | -1.1117169 | old | astrocyte |
| MT-CO3 | 1.1048037 | old | astrocyte |
| MT-CO2 | -1.1037254 | old | astrocyte |
| ERO1A | 1.0387514 | old | astrocyte |
| MT2A | -0.8424649 | old | astrocyte |
| SELENOM | 0.7595936 | old | astrocyte |
| LDHA | 0.69253755 | old | astrocyte |
| HILPDA | 0.6796659 | old | astrocyte |
| ANGPTL1 | -0.6486343 | old | astrocyte |
| TM7SF2 | -0.59906185 | old | astrocyte |
| SEMA6D | -0.5521946 | old | astrocyte |
| VEGFA | -0.45605493 | old | astrocyte |
| MT-ATP6 | -0.78169566 | young | AC |
| BEX1 | 0.46964857 | young | AC |
| MT-ND6 | 0.3754455 | young | AC |
| MT-CO2 | 0.33762944 | young | AC |
| HLA-C | -0.32920003 | young | AC |

| | | | |
|---|---|---|---|
| ZNF804A | 0.31770003 | young | AC |
| HMGCS1 | 0.2925035 | young | AC |
| COX7C | 0.29245433 | young | AC |
| HSP90AA1 | 0.2784884 | young | AC |
| STXBP5-AS1 | -0.26677173 | young | AC |
| FTX | -0.26122153 | young | AC |
| BNIP3 | -0.23789343 | young | AC |
| RNF165 | -0.23714802 | young | AC |
| RTN3 | -0.23046067 | young | AC |
| H3F3B | -0.2253851 | young | AC |
| PAK3 | -0.2232989 | young | AC |
| C1QL1 | -0.21844393 | young | AC |
| P4HA1 | -0.21797794 | young | AC |
| KCNIP4 | -0.20155343 | young | AC |
| PDE5A | 0.19379929 | young | AC |
| PGK1 | -0.18878886 | young | AC |
| GPI | -0.1850431 | young | AC |
| LDHA | -0.1699042 | young | AC |
| FAM155A | -0.16936004 | young | AC |
| SSBP3 | -0.1691032 | young | AC |
| DDX5 | 0.16449985 | young | AC |
| SNAP25 | -0.15233451 | young | AC |
| HIST1H2BE | 0.1513024 | young | AC |
| DAB1 | -0.14974934 | young | AC |
| H2AFY | 0.14898762 | young | AC |
| MAPT | -0.14725044 | young | AC |
| HTR5A | -0.1451264 | young | AC |
| NEDD4L | -0.14193267 | young | AC |
| GABRG3-AS1 | 0.14045823 | young | AC |
| BTBD8 | 0.13752007 | young | AC |
| GAPDH | -0.13368645 | young | AC |
| NDUFC1 | 0.13066873 | young | AC |
| B2M | -0.12507972 | young | AC |
| PDK1 | -0.12441357 | young | AC |
| MT-ND3 | 0.12374308 | young | AC |
| SLC2A3 | -0.12209148 | young | AC |
| VEGFA | -0.11951253 | young | AC |
| INSIG1 | 0.11390379 | young | AC |
| SFPQ | 0.113423236 | young | AC |
| PGRMC1 | -0.10735928 | young | AC |
| UBC | -0.10670604 | young | AC |
| PIKFYVE | -0.105812036 | young | AC |
| PRKAG2-AS1 | 0.10391843 | young | AC |
| AC097534.2 | -0.10077661 | young | AC |
| ATP5IF1 | 0.09913887 | young | AC |
| FADS2 | 0.098710544 | young | AC |
| GRIP2 | 0.097649954 | young | AC |
| SNCB | -0.09520978 | young | AC |
| VDAC1 | -0.09078894 | young | AC |
| HLA-A | -0.089242846 | young | AC |
| PPIA | 0.088255614 | young | AC |
| PSMB8 | 0.08758153 | young | AC |
| DYNLL2 | 0.085310906 | young | AC |
| IFIT1 | 0.08524107 | young | AC |

| | | | |
|---|---|---|---|
| LGALS1 | -0.0827199 | young | AC |
| NPEPPS | -0.0773514 | young | AC |
| MOCS2 | 0.076294884 | young | AC |
| PDXP | -0.075859636 | young | AC |
| WSB1 | -0.06668006 | young | AC |
| CALM3 | -0.065712586 | young | AC |
| KBTBD2 | -0.0634139 | young | AC |
| STX7 | -0.06322383 | young | AC |
| CTSA | -0.06281593 | young | AC |
| PKM | -0.05973085 | young | AC |
| TCEAL7 | 0.059383694 | young | AC |
| AL160254.1 | 0.057205826 | young | AC |
| TIMP2 | -0.05508054 | young | AC |
| PEG3 | -0.055031277 | young | AC |
| NRG3 | -0.052802507 | young | AC |
| INPP5F | -0.051773254 | young | AC |
| NRXN1 | -0.051651917 | young | AC |
| MT-CYB | 0.050055865 | young | AC |
| BHLHE40 | -0.050015062 | young | AC |
| GRIK2 | 0.047063753 | young | AC |
| ZNF407 | 0.04656801 | young | AC |
| NAPB | -0.04619069 | young | AC |
| ENO1 | -0.046064787 | young | AC |
| AC002463.1 | 0.04278227 | young | AC |
| ENO2 | -0.04147884 | young | AC |
| CRYAB | -0.03588234 | young | AC |
| AC096576.2 | 0.035836108 | young | AC |
| PFKL | -0.03576373 | young | AC |
| ISCA1 | 0.0356126 | young | AC |
| DAAM1 | -0.035034727 | young | AC |
| NCDN | -0.031632997 | young | AC |
| IKZF5 | 0.030421283 | young | AC |
| CACNA2D3 | -0.029470475 | young | AC |
| RTN4 | 0.026953146 | young | AC |
| LRCH2 | -0.026059942 | young | AC |
| TCEAL6 | 0.02444727 | young | AC |
| PDE4A | -0.024395052 | young | AC |
| AC096576.3 | 0.021563347 | young | AC |
| CDH18 | -0.02034231 | young | AC |
| UBE2B | -0.019252572 | young | AC |
| HSPA1A | 0.01654863 | young | AC |
| MAGI2 | -0.016387826 | young | AC |
| PREPL | -0.016343068 | young | AC |
| HS6ST3 | -0.015863307 | young | AC |
| AC024558.2 | 0.014681264 | young | AC |
| ARHGAP28 | -0.013459981 | young | AC |
| MT-ATP6 | 0.78169566 | old | AC |
| BEX1 | -0.46964857 | old | AC |
| MT-ND6 | -0.3754455 | old | AC |
| MT-CO2 | -0.33762944 | old | AC |
| HLA-C | 0.32920003 | old | AC |
| ZNF804A | -0.31770003 | old | AC |
| HMGCS1 | -0.2925035 | old | AC |
| COX7C | -0.29245433 | old | AC |

| | | | |
|---|---|---|---|
| HSP90AA1 | -0.2784884 | old | AC |
| STXBP5-AS1 | 0.26677173 | old | AC |
| FTX | 0.26122153 | old | AC |
| BNIP3 | 0.23789343 | old | AC |
| RNF165 | 0.23714802 | old | AC |
| RTN3 | 0.23046067 | old | AC |
| H3F3B | 0.2253851 | old | AC |
| PAK3 | 0.2232989 | old | AC |
| C1QL1 | 0.21844393 | old | AC |
| P4HA1 | 0.21797794 | old | AC |
| KCNIP4 | 0.20155343 | old | AC |
| PDE5A | -0.19379929 | old | AC |
| PGK1 | 0.18878886 | old | AC |
| GPI | 0.1850431 | old | AC |
| LDHA | 0.1699042 | old | AC |
| FAM155A | 0.16936004 | old | AC |
| SSBP3 | 0.1691032 | old | AC |
| DDX5 | -0.16449985 | old | AC |
| SNAP25 | 0.15233451 | old | AC |
| HIST1H2BE | -0.1513024 | old | AC |
| DAB1 | 0.14974934 | old | AC |
| H2AFY | -0.14898762 | old | AC |
| MAPT | 0.14725044 | old | AC |
| HTR5A | 0.1451264 | old | AC |
| NEDD4L | 0.14193267 | old | AC |
| GABRG3-AS1 | -0.14045823 | old | AC |
| BTBD8 | -0.13752007 | old | AC |
| GAPDH | 0.13368645 | old | AC |
| NDUFC1 | -0.13066873 | old | AC |
| B2M | 0.12507972 | old | AC |
| PDK1 | 0.12441357 | old | AC |
| MT-ND3 | -0.12374308 | old | AC |
| SLC2A3 | 0.12209148 | old | AC |
| VEGFA | 0.11951253 | old | AC |
| INSIG1 | -0.11390379 | old | AC |
| SFPQ | -0.113423236 | old | AC |
| PGRMC1 | 0.10735928 | old | AC |
| UBC | 0.10670604 | old | AC |
| PIKFYVE | 0.105812036 | old | AC |
| PRKAG2-AS1 | -0.10391843 | old | AC |
| AC097534.2 | 0.10077661 | old | AC |
| ATP5IF1 | -0.09913887 | old | AC |
| FADS2 | -0.098710544 | old | AC |
| GRIP2 | -0.097649954 | old | AC |
| SNCB | 0.09520978 | old | AC |
| VDAC1 | 0.09078894 | old | AC |
| HLA-A | 0.089242846 | old | AC |
| PPIA | -0.088255614 | old | AC |
| PSMB8 | -0.08758153 | old | AC |
| DYNLL2 | -0.085310906 | old | AC |
| IFIT1 | -0.08524107 | old | AC |
| LGALS1 | 0.0827199 | old | AC |
| NPEPPS | 0.0773514 | old | AC |
| MOCS2 | -0.076294884 | old | AC |

| | | | |
|---|---|---|---|
| PDXP | 0.075859636 | old | AC |
| WSB1 | 0.06668006 | old | AC |
| CALM3 | 0.065712586 | old | AC |
| KBTBD2 | 0.0634139 | old | AC |
| STX7 | 0.06322383 | old | AC |
| CTSA | 0.06281593 | old | AC |
| PKM | 0.05973085 | old | AC |
| TCEAL7 | -0.059383694 | old | AC |
| AL160254.1 | -0.057205826 | old | AC |
| TIMP2 | 0.05508054 | old | AC |
| PEG3 | 0.055031277 | old | AC |
| NRG3 | 0.052802507 | old | AC |
| INPP5F | 0.051773254 | old | AC |
| NRXN1 | 0.051651917 | old | AC |
| MT-CYB | -0.050055865 | old | AC |
| BHLHE40 | 0.050015062 | old | AC |
| GRIK2 | -0.047063753 | old | AC |
| ZNF407 | -0.04656801 | old | AC |
| NAPB | 0.04619069 | old | AC |
| ENO1 | 0.046064787 | old | AC |
| AC002463.1 | -0.04278227 | old | AC |
| ENO2 | 0.04147884 | old | AC |
| CRYAB | 0.03588234 | old | AC |
| AC096576.2 | -0.035836108 | old | AC |
| PFKL | 0.03576373 | old | AC |
| ISCA1 | -0.0356126 | old | AC |
| DAAM1 | 0.035034727 | old | AC |
| NCDN | 0.031632997 | old | AC |
| IKZF5 | -0.030421283 | old | AC |
| CACNA2D3 | 0.029470475 | old | AC |
| RTN4 | -0.026953146 | old | AC |
| LRCH2 | 0.026059942 | old | AC |
| TCEAL6 | -0.02444727 | old | AC |
| PDE4A | 0.024395052 | old | AC |
| AC096576.3 | -0.021563347 | old | AC |
| CDH18 | 0.02034231 | old | AC |
| UBE2B | 0.019252572 | old | AC |
| HSPA1A | -0.01654863 | old | AC |
| MAGI2 | 0.016387826 | old | AC |
| PREPL | 0.016343068 | old | AC |
| HS6ST3 | 0.015863307 | old | AC |
| AC024558.2 | -0.014681264 | old | AC |
| ARHGAP28 | 0.013459981 | old | AC |
