## Supplementary material for "Interpretable Aging Signatures in Human Retinal Cell Types Revealed by Single-Cell RNA Sequencing and Sparse Logistic Regression": Table S12

Table S12: List of machine learning selected aging features recurring in more than two cel
with frequency counts.

| feature | occurrence_count |
|---|---|
| A2M | 2 |
| AC092691.1 | 2 |
| ACTB | 2 |
| ACTG1 | 2 |
| ADM | 2 |
| AHSA1 | 2 |
| AL050403.2 | 2 |
| ANGPTL1 | 2 |
| ANOS1 | 2 |
| ARHGAP24 | 2 |
| ARMC9 | 2 |
| B2M | 2 |
| BAG3 | 2 |
| BCO2 | 2 |
| BEX1 | 2 |
| BHLHE40 | 4 |
| BHLHE41 | 2 |
| BMPR1B | 2 |
| CADM1 | 2 |
| CALM1 | 2 |
| CALM2 | 2 |
| CCL2 | 2 |
| CCNG2 | 2 |
| CCSER1 | 2 |
| COTL1 | 2 |
| CRYAB | 6 |
| CST3 | 2 |
| DLG2 | 2 |
| DMD | 2 |
| DNAJB1 | 3 |
| DNAJB4 | 2 |
| DOCK3 | 2 |
| DOK6 | 2 |
| DPF3 | 2 |
| DRD4 | 2 |
| DTNA | 2 |
| ENO1 | 2 |
| ENO2 | 2 |
| EPCAM | 2 |
| ERO1A | 4 |
| EYS | 2 |
| FAM155A | 2 |
| FGF12 | 2 |
| FHIT | 2 |
| FTX | 5 |
| FUS | 2 |
| GAPDH | 2 |
| GLUL | 2 |
| GPX3 | 5 |
| GRIA4 | 2 |
| HILPDA | 2 |
| HLA-A | 3 |
| HLA-C | 3 |

| | |
|---|---|
| HMGB2 | 2 |
| HNRNPA2B1 | 2 |
| HPRT1 | 2 |
| HSP90AA1 | 2 |
| HSP90AB1 | 2 |
| HSPA1A | 5 |
| HSPA1B | 3 |
| HSPA6 | 3 |
| HSPB1 | 2 |
| HSPE1 | 2 |
| HSPH1 | 3 |
| IER5L | 2 |
| IFITM3 | 2 |
| INSIG1 | 2 |
| KCNB1 | 2 |
| LDHA | 2 |
| LGALS1 | 2 |
| LGALS3 | 3 |
| MAGI2 | 4 |
| MAP2 | 2 |
| MBP | 2 |
| METRNL | 2 |
| MRLN | 2 |
| MT-ATP6 | 4 |
| MT-ATP8 | 4 |
| MT-CO1 | 2 |
| MT-CO2 | 5 |
| MT-CO3 | 4 |
| MT-CYB | 3 |
| MT-ND1 | 2 |
| MT-ND2 | 3 |
| MT-ND3 | 4 |
| MT-ND5 | 3 |
| MT-ND6 | 7 |
| MT1X | 2 |
| MT2A | 6 |
| MT3 | 3 |
| NAALADL2 | 3 |
| NAV3 | 2 |
| NCKAP5 | 2 |
| NCL | 4 |
| NPVF | 2 |
| NRG3 | 2 |
| NRXN3 | 2 |
| NTM | 2 |
| P4HA1 | 3 |
| PDE5A | 3 |
| PDK1 | 2 |
| PGK1 | 2 |
| PMP22 | 2 |
| PPIA | 2 |
| PRKN | 2 |
| PRR16 | 3 |
| PRSS51 | 2 |

| | |
|---|---|
| PRUNE2 | 2 |
| RGS9 | 2 |
| RPS26 | 3 |
| RRAD | 2 |
| S100A10 | 3 |
| SELENOW | 2 |
| SKAP2 | 2 |
| SLC2A3 | 2 |
| SLC35F1 | 3 |
| SNAP25-AS1 | 2 |
| SNHG14 | 3 |
| SPP1 | 2 |
| TF | 3 |
| TMEM108 | 2 |
| TMEM176A | 2 |
| TMEM176B | 2 |
| TRNP1 | 2 |
| TUBA1B | 3 |
| UBC | 3 |
| USH2A | 2 |
| VEGFA | 2 |
| WWOX | 2 |
| YBX3 | 2 |
| ZBTB20 | 2 |
| ZNF385D | 2 |
