## Supplementary material for "Interpretable Aging Signatures in Human Retinal Cell Types Revealed by Single-Cell RNA Sequencing and Sparse Logistic Regression": Table S13

Table S13: Transcription factor regulon activity across retinal interneuron populations during aging.

| Topic | cellType | RSS | Z | CellType |
|---|---|---|---|---|
| ATF3(-) | old | 0.563818883 | 1.125142602 | AC |
| ATF3(-) | young | 0.311519957 | 0.797873129 | AC |
| ATF3(+) | old | 0.526456782 | 0.881383892 | AC |
| ATF3(+) | young | 0.33662015 | 1.100524801 | AC |
| ATF4(+) | old | 0.499719901 | 0.706946497 | AC |
| ATF4(+) | young | 0.34998851 | 1.261717052 | AC |
| BACH2(+) | old | 0.567539247 | 1.149415089 | AC |
| BACH2(+) | young | 0.307650279 | 0.751213552 | AC |
| BCLAF1(+) | old | 0.540438445 | 0.97260339 | AC |
| BCLAF1(+) | young | 0.329593557 | 1.015799955 | AC |
| BHLHE40(+) | old | 0.57407324 | 1.192044321 | AC |
| BHLHE40(+) | young | 0.29849832 | 0.640861587 | AC |
| BHLHE41(+) | old | 0.558812844 | 1.092482081 | AC |
| BHLHE41(+) | young | 0.316375248 | 0.856416987 | AC |
| CEBPB(+) | old | 0.533109554 | 0.924788066 | AC |
| CEBPB(+) | young | 0.328590271 | 1.00370259 | AC |
| CEBPD(-) | old | 0.546236986 | 1.010434367 | AC |
| CEBPD(-) | young | 0.322397127 | 0.929027257 | AC |
| CHD1(+) | old | 0.555849563 | 1.073148975 | AC |
| CHD1(+) | young | 0.307111538 | 0.744717553 | AC |
| CRX(-) | old | 0.532706528 | 0.922158636 | AC |
| CRX(-) | young | 0.333724355 | 1.06560806 | AC |
| DBP(+) | old | 0.550928203 | 1.041040918 | AC |
| DBP(+) | young | 0.304299645 | 0.710812466 | AC |
| EGR1(+) | old | 0.541811671 | 0.981562623 | AC |
| EGR1(+) | young | 0.331302203 | 1.03640237 | AC |
| EGR2(+) | old | 0.545827539 | 1.007763046 | AC |
| EGR2(+) | young | 0.323115166 | 0.937685181 | AC |
| EGR3(-) | old | 0.545804161 | 1.007610519 | AC |
| EGR3(-) | young | 0.308369018 | 0.759879918 | AC |
| ELF1(-) | old | 0.549281535 | 1.030297691 | AC |
| ELF1(-) | young | 0.318653771 | 0.88389083 | AC |
| ETS2(+) | old | 0.553738036 | 1.059372895 | AC |
| ETS2(+) | young | 0.318376945 | 0.880552938 | AC |
| FOS(-) | old | 0.514532244 | 0.80358554 | AC |
| FOS(-) | young | 0.349856281 | 1.260122668 | AC |
| FOS(+) | old | 0.516556317 | 0.816791044 | AC |
| FOS(+) | young | 0.357132604 | 1.347858703 | AC |
| FOSB(-) | old | 0.539189257 | 0.964453406 | AC |
| FOSB(-) | young | 0.337444111 | 1.11045992 | AC |
| FOSB(+) | old | 0.51603777 | 0.813407927 | AC |
| FOSB(+) | young | 0.341261184 | 1.156485197 | AC |
| FOSL2(-) | old | 0.545709069 | 1.006990118 | AC |
| FOSL2(-) | young | 0.325763272 | 0.969615368 | AC |
| GBX2(+) | old | 0.547257766 | 1.017094167 | AC |
| GBX2(+) | young | 0.243062375 | 0 | AC |
| GLIS3(-) | old | 0.570000464 | 1.165472617 | AC |
| GLIS3(-) | young | 0.299399 | 0.65172175 | AC |
| HMGA1(-) | old | 0.499877731 | 0.707976217 | AC |
| HMGA1(-) | young | 0.32849681 | 1.002575665 | AC |
| JUN(-) | old | 0.50395454 | 0.734574232 | AC |
| JUN(-) | young | 0.353565554 | 1.304848125 | AC |
| JUN(+) | old | 0.539795534 | 0.968408893 | AC |

| | | | | |
|---|---|---|---|---|
| JUN(+) | young | 0.340276152 | 1.144607941 | AC |
| JUNB(+) | old | 0.519934305 | 0.838829791 | AC |
| JUNB(+) | young | 0.352844781 | 1.296157231 | AC |
| JUND(-) | old | 0.470933235 | 0.519135852 | AC |
| JUND(-) | young | 0.329471085 | 1.014323216 | AC |
| JUND(+) | old | 0.521081542 | 0.846314625 | AC |
| JUND(+) | young | 0.349026846 | 1.250121558 | AC |
| KLF6(-) | old | 0.534996812 | 0.937100962 | AC |
| KLF6(-) | young | 0.335037992 | 1.08144755 | AC |
| KLF6(+) | old | 0.548469237 | 1.024998075 | AC |
| KLF6(+) | young | 0.328789988 | 1.006110726 | AC |
| MXI1(+) | old | 0.566196805 | 1.140656693 | AC |
| MXI1(+) | young | 0.304298988 | 0.710804551 | AC |
| NEUROD1(+) | old | 0.537135593 | 0.951054848 | AC |
| NEUROD1(+) | young | 0.334846767 | 1.079141805 | AC |
| NFIA(+) | old | 0.569679483 | 1.163378462 | AC |
| NFIA(+) | young | 0.302048849 | 0.683672954 | AC |
| NFKB1(+) | old | 0.561288639 | 1.108634722 | AC |
| NFKB1(+) | young | 0.28475115 | 0.475101739 | AC |
| PROX1(+) | old | 0.565741944 | 1.137689081 | AC |
| PROX1(+) | young | 0.287181991 | 0.504412194 | AC |
| RXRG(-) | old | 0.556900227 | 1.080003742 | AC |
| RXRG(-) | young | 0.291588219 | 0.55754136 | AC |
| SMAD1(+) | old | 0.54480569 | 1.001096272 | AC |
| SMAD1(+) | young | 0.317913784 | 0.874968261 | AC |
| SPI1(-) | old | 0.511782091 | 0.785642928 | AC |
| SPI1(-) | young | 0.332339914 | 1.048914821 | AC |
| SPI1(+) | old | 0.509587252 | 0.771323307 | AC |
| SPI1(+) | young | 0.329161836 | 1.010594378 | AC |
| TCF4(+) | old | 0.562354511 | 1.115588706 | AC |
| TCF4(+) | young | 0.309679573 | 0.775682263 | AC |
| TFAP2B(+) | old | 0.559847698 | 1.099233702 | AC |
| TFAP2B(+) | young | 0.297728197 | 0.631575644 | AC |
| THRB(+) | old | 0.55518439 | 1.068809237 | AC |
| THRB(+) | young | 0.300729737 | 0.667767435 | AC |
| XBP1(-) | old | 0.451190767 | 0.390331569 | AC |
| XBP1(-) | young | 0.34266213 | 1.173377449 | AC |
| XBP1(+) | old | 0.516171757 | 0.814282088 | AC |
| XBP1(+) | young | 0.337535068 | 1.111556654 | AC |
| YBX1(+) | old | 0.516386237 | 0.815681404 | AC |
| YBX1(+) | young | 0.352480553 | 1.291765467 | AC |
| ZBTB21(-) | old | 0.546497753 | 1.012135674 | AC |
| ZBTB21(-) | young | 0.267625085 | 0.268600051 | AC |
| ZNF267(-) | old | 0.555196413 | 1.068887674 | AC |
| ZNF267(-) | young | 0.307588305 | 0.750466286 | AC |
| ATF3(-) | old | 0.520414392 | 1.102509364 | BC |
| ATF3(-) | young | 0.356457661 | 1.064021218 | BC |
| ATF3(+) | old | 0.506299716 | 1.005921098 | BC |
| ATF3(+) | young | 0.36851626 | 1.183451287 | BC |
| ATF4(+) | old | 0.470322818 | 0.759727262 | BC |
| ATF4(+) | young | 0.392349791 | 1.419501963 | BC |
| BACH2(+) | old | 0.526948861 | 1.147225453 | BC |
| BACH2(+) | young | 0.351119639 | 1.011152686 | BC |
| BCLAF1(+) | old | 0.511287807 | 1.040055149 | BC |

| | | | | |
|---|---|---|---|---|
| BCLAF1(+) | young | 0.364136795 | 1.140076446 | BC |
| BHLHE40(+) | old | 0.533223432 | 1.190163026 | BC |
| BHLHE40(+) | young | 0.341152502 | 0.912436747 | BC |
| BHLHE41(+) | old | 0.527164178 | 1.148698888 | BC |
| BHLHE41(+) | young | 0.350118915 | 1.001241369 | BC |
| CEBPB(+) | old | 0.501489631 | 0.973005165 | BC |
| CEBPB(+) | young | 0.370945747 | 1.207513275 | BC |
| CEBPD(-) | old | 0.521007586 | 1.106568658 | BC |
| CEBPD(-) | young | 0.348753224 | 0.987715378 | BC |
| CEBPD(+) | old | 0.487066318 | 0.87430486 | BC |
| CEBPD(+) | young | 0.361003299 | 1.109041863 | BC |
| CHD1(+) | old | 0.530450346 | 1.171186497 | BC |
| CHD1(+) | young | 0.334861452 | 0.850129293 | BC |
| CREM(+) | old | 0.51446855 | 1.06182132 | BC |
| CREM(+) | young | 0.347926984 | 0.979532174 | BC |
| CRX(-) | old | 0.498302961 | 0.951198436 | BC |
| CRX(-) | young | 0.361433009 | 1.113297768 | BC |
| CRX(+) | old | 0.50555634 | 1.000834095 | BC |
| CRX(+) | young | 0.318430647 | 0.68739627 | BC |
| DBP(+) | old | 0.505352152 | 0.999436817 | BC |
| DBP(+) | young | 0.361850613 | 1.117433777 | BC |
| DDIT3(+) | old | 0.476809276 | 0.804114801 | BC |
| DDIT3(+) | young | 0.37075374 | 1.205611608 | BC |
| EGR1(+) | old | 0.513918521 | 1.058057413 | BC |
| EGR1(+) | young | 0.361069079 | 1.109693357 | BC |
| EGR2(+) | old | 0.513921095 | 1.058075029 | BC |
| EGR2(+) | young | 0.36010317 | 1.100126849 | BC |
| EGR3(-) | old | 0.528497824 | 1.157825176 | BC |
| EGR3(-) | young | 0.326608786 | 0.768393718 | BC |
| ELF1(-) | old | 0.528684892 | 1.159105303 | BC |
| ELF1(-) | young | 0.338399386 | 0.885169489 | BC |
| ETS2(+) | old | 0.52312646 | 1.121068345 | BC |
| ETS2(+) | young | 0.349470995 | 0.994824286 | BC |
| FOS(-) | old | 0.503777383 | 0.9886605 | BC |
| FOS(-) | young | 0.37344124 | 1.23222899 | BC |
| FOS(+) | old | 0.495407798 | 0.931386517 | BC |
| FOS(+) | young | 0.383179287 | 1.328675995 | BC |
| FOSB(-) | old | 0.509983548 | 1.031129961 | BC |
| FOSB(-) | young | 0.368594504 | 1.184226233 | BC |
| FOSB(+) | old | 0.495691057 | 0.933324886 | BC |
| FOSB(+) | young | 0.3716873 | 1.214857724 | BC |
| FOSL2(-) | old | 0.514245522 | 1.060295114 | BC |
| FOSL2(-) | young | 0.360822946 | 1.107255615 | BC |
| GLIS3(-) | old | 0.527547724 | 1.151323539 | BC |
| GLIS3(-) | young | 0.345614876 | 0.956632736 | BC |
| JUN(-) | old | 0.504237913 | 0.991811958 | BC |
| JUN(-) | young | 0.359237528 | 1.091553409 | BC |
| JUN(+) | old | 0.512158365 | 1.046012471 | BC |
| JUN(+) | young | 0.369124904 | 1.189479384 | BC |
| JUNB(+) | old | 0.496130573 | 0.936332548 | BC |
| JUNB(+) | young | 0.381933656 | 1.316339087 | BC |
| JUND(+) | old | 0.49474153 | 0.926827171 | BC |
| JUND(+) | young | 0.382457746 | 1.321529746 | BC |
| KLF6(-) | old | 0.516182623 | 1.073550912 | BC |

| | | | | |
|---|---|---|---|---|
| KLF6(-) | young | 0.356458835 | 1.06403284 | BC |
| KLF6(+) | old | 0.517693608 | 1.083890743 | BC |
| KLF6(+) | young | 0.361841853 | 1.117347017 | BC |
| MAFF(-) | old | 0.489781371 | 0.89288426 | BC |
| MAFF(-) | young | 0.357140688 | 1.070786008 | BC |
| MITF(+) | old | 0.507845537 | 1.016499324 | BC |
| MITF(+) | young | 0.309320118 | 0.597164295 | BC |
| MXI1(+) | old | 0.528101604 | 1.155113794 | BC |
| MXI1(+) | young | 0.341715334 | 0.918011111 | BC |
| NEUROD1(+) | old | 0.511502346 | 1.041523264 | BC |
| NEUROD1(+) | young | 0.368063108 | 1.17896321 | BC |
| NFIA(+) | old | 0.526639466 | 1.14510823 | BC |
| NFIA(+) | young | 0.34887979 | 0.988968908 | BC |
| NFKB1(+) | old | 0.518733811 | 1.091008968 | BC |
| NFKB1(+) | young | 0.321466912 | 0.717467864 | BC |
| PAX6(-) | old | 0.477395792 | 0.808128392 | BC |
| PAX6(-) | young | 0.353373386 | 1.033474115 | BC |
| PPARD(-) | old | 0.507360933 | 1.013183126 | BC |
| PPARD(-) | young | 0.3320586 | 0.822369449 | BC |
| PROX1(+) | old | 0.498361732 | 0.951600609 | BC |
| PROX1(+) | young | 0.358263239 | 1.081903918 | BC |
| RXRG(-) | old | 0.508852134 | 1.023387572 | BC |
| RXRG(-) | young | 0.337358805 | 0.874863431 | BC |
| SMAD1(+) | old | 0.510373074 | 1.033795534 | BC |
| SMAD1(+) | young | 0.360442687 | 1.103489479 | BC |
| SPI1(-) | old | 0.496557724 | 0.939255588 | BC |
| SPI1(-) | young | 0.364026414 | 1.138983216 | BC |
| SPI1(+) | old | 0.485085484 | 0.860749798 | BC |
| SPI1(+) | young | 0.37147158 | 1.212721195 | BC |
| TCF4(+) | old | 0.52366496 | 1.124753359 | BC |
| TCF4(+) | young | 0.352786628 | 1.02766278 | BC |
| TFAP2B(+) | old | 0.518021305 | 1.086133208 | BC |
| TFAP2B(+) | young | 0.340363507 | 0.904622428 | BC |
| THRB(+) | old | 0.515443041 | 1.068489871 | BC |
| THRB(+) | young | 0.351996112 | 1.019833399 | BC |
| XBP1(+) | old | 0.490834767 | 0.900092768 | BC |
| XBP1(+) | young | 0.376075033 | 1.258314451 | BC |
| YBX1(+) | old | 0.487387618 | 0.876503548 | BC |
| YBX1(+) | young | 0.38778122 | 1.374254194 | BC |
| ZNF267(-) | old | 0.520359784 | 1.10213568 | BC |
| ZNF267(-) | young | 0.341982774 | 0.920659874 | BC |
| ATF3(-) | old | 0.515501983 | 1.012260693 | HC |
| ATF3(-) | young | 0.363374241 | 0.865250885 | HC |
| ATF4(+) | old | 0.454098981 | 0.603173607 | HC |
| ATF4(+) | young | 0.407284025 | 1.277382117 | HC |
| ATF6(+) | old | 0.46895326 | 0.702137718 | HC |
| ATF6(+) | young | 0.38095295 | 1.030242245 | HC |
| BHLHE40(+) | old | 0.535693644 | 1.146784207 | HC |
| BHLHE40(+) | young | 0.339440782 | 0.640614708 | HC |
| BHLHE41(+) | old | 0.524779923 | 1.074073389 | HC |
| BHLHE41(+) | young | 0.348313795 | 0.723895598 | HC |
| CHD1(+) | old | 0.516193242 | 1.01686609 | HC |
| CHD1(+) | young | 0.339024236 | 0.636705063 | HC |
| CRX(-) | old | 0.459250177 | 0.637492576 | HC |

| | | | | |
|---|---|---|---|---|
| CRX(-) | young | 0.397715486 | 1.187573117 | HC |
| DBP(+) | old | 0.527642753 | 1.093146509 | HC |
| DBP(+) | young | 0.341652449 | 0.661373112 | HC |
| DDIT3(+) | old | 0.477131074 | 0.756621022 | HC |
| DDIT3(+) | young | 0.381377308 | 1.034225214 | HC |
| FOS(-) | old | 0.452484007 | 0.59241412 | HC |
| FOS(-) | young | 0.405425067 | 1.259934195 | HC |
| FOS(+) | old | 0.492391152 | 0.858288699 | HC |
| FOS(+) | young | 0.383854441 | 1.057475247 | HC |
| FOSB(-) | old | 0.500337021 | 0.911226702 | HC |
| FOSB(-) | young | 0.379306702 | 1.014790784 | HC |
| FOSL1(-) | old | 0.514650321 | 1.006586635 | HC |
| FOSL1(-) | young | 0.313382222 | 0.396032632 | HC |
| GABPB1(-) | old | 0.45918541 | 0.637061076 | HC |
| GABPB1(-) | young | 0.38033144 | 1.02440884 | HC |
| HMGA1(-) | old | 0.455230768 | 0.610713945 | HC |
| HMGA1(-) | young | 0.385420046 | 1.072169794 | HC |
| IRF1(+) | old | 0.522126592 | 1.056396022 | HC |
| IRF1(+) | young | 0.224730946 | 0 | HC |
| JUN(-) | old | 0.451604692 | 0.586555827 | HC |
| JUN(-) | young | 0.399842862 | 1.20754038 | HC |
| JUNB(+) | old | 0.494278539 | 0.870863093 | HC |
| JUNB(+) | young | 0.381865148 | 1.038804014 | HC |
| JUND(+) | old | 0.489607215 | 0.839741191 | HC |
| JUND(+) | young | 0.386473928 | 1.082061388 | HC |
| MXI1(+) | old | 0.520400727 | 1.044897738 | HC |
| MXI1(+) | young | 0.351203866 | 0.751021408 | HC |
| NEUROD1(+) | old | 0.498981803 | 0.902197792 | HC |
| NEUROD1(+) | young | 0.379049309 | 1.012374932 | HC |
| NR2F1(-) | old | 0.40348171 | 0.265944628 | HC |
| NR2F1(-) | young | 0.392900617 | 1.142381412 | HC |
| ONECUT1(+) | old | 0.479008212 | 0.769127134 | HC |
| ONECUT1(+) | young | 0.38492274 | 1.067502156 | HC |
| SPI1(+) | old | 0.471932324 | 0.721985231 | HC |
| SPI1(+) | young | 0.387462562 | 1.091340573 | HC |
| USF1(-) | old | 0.358365339 | 0 | HC |
| USF1(-) | young | 0.39407952 | 1.153446437 | HC |
| YBX1(+) | old | 0.47138184 | 0.718317722 | HC |
| YBX1(+) | young | 0.40306099 | 1.237745283 | HC |
| ZNF675(-) | old | 0.439028451 | 0.502768757 | HC |
| ZNF675(-) | young | 0.387379088 | 1.090557103 | HC |
