## Supplementary material for "Interpretable Aging Signatures in Human Retinal Cell Types Revealed by Single-Cell RNA Sequencing and Sparse Logistic Regression": Table S14

Table S14: Transcription factor regulon activity across retinal glial populations during aging.

| Topic | cellType | RSS | Z | CellType |
|---|---|---|---|---|
| ATF3(+) | old | 0.483806336 | 1.241653454 | MGC |
| ATF3(+) | young | 0.383098892 | 0.916845894 | MGC |
| ATF4(+) | old | 0.480440686 | 1.216975554 | MGC |
| ATF4(+) | young | 0.376070986 | 0.856650517 | MGC |
| BACH1(+) | old | 0.472407387 | 1.158073123 | MGC |
| BACH1(+) | young | 0.380928851 | 0.898259076 | MGC |
| BCLAF1(+) | old | 0.442138539 | 0.936133349 | MGC |
| BCLAF1(+) | young | 0.407654089 | 1.127165926 | MGC |
| BHLHE40(+) | old | 0.490575041 | 1.291283515 | MGC |
| BHLHE40(+) | young | 0.377523274 | 0.869089642 | MGC |
| BHLHE41(+) | old | 0.499313546 | 1.355356711 | MGC |
| BHLHE41(+) | young | 0.366366133 | 0.773526571 | MGC |
| CEBPA(+) | old | 0.478328865 | 1.201491079 | MGC |
| CEBPA(+) | young | 0.377912016 | 0.872419292 | MGC |
| CEBPB(+) | old | 0.483392469 | 1.238618862 | MGC |
| CEBPB(+) | young | 0.371124206 | 0.814280384 | MGC |
| CEBPD(+) | old | 0.48543941 | 1.25362761 | MGC |
| CEBPD(+) | young | 0.369507678 | 0.800434507 | MGC |
| CHD1(+) | old | 0.473679923 | 1.16740372 | MGC |
| CHD1(+) | young | 0.362755957 | 0.74260471 | MGC |
| CREB3(+) | old | 0.469703838 | 1.13824994 | MGC |
| CREB3(+) | young | 0.388471078 | 0.962859712 | MGC |
| CREM(+) | old | 0.480468658 | 1.21718065 | MGC |
| CREM(+) | young | 0.392355667 | 0.996131973 | MGC |
| E2F1(-) | old | 0.375401016 | 0.446794918 | MGC |
| E2F1(-) | young | 0.425807549 | 1.282653832 | MGC |
| E2F1(+) | old | 0.432473451 | 0.865266189 | MGC |
| E2F1(+) | young | 0.410469538 | 1.151280791 | MGC |
| E2F6(-) | old | 0.391695215 | 0.566268601 | MGC |
| E2F6(-) | young | 0.397979577 | 1.044301848 | MGC |
| E2F8(-) | old | 0.432335789 | 0.864256812 | MGC |
| E2F8(-) | young | 0.399178363 | 1.054569685 | MGC |
| E2F8(+) | old | 0.444148987 | 0.950874529 | MGC |
| E2F8(+) | young | 0.422587842 | 1.25507642 | MGC |
| ELF1(-) | old | 0.437154726 | 0.899590619 | MGC |
| ELF1(-) | young | 0.43202829 | 1.335935689 | MGC |
| ELF1(+) | old | 0.476519224 | 1.188222279 | MGC |
| ELF1(+) | young | 0.388106144 | 0.959733978 | MGC |
| ELK1(+) | old | 0.401786235 | 0.64025882 | MGC |
| ELK1(+) | young | 0.400024098 | 1.061813573 | MGC |
| ETS1(+) | old | 0.484527772 | 1.246943222 | MGC |
| ETS1(+) | young | 0.33661778 | 0.518726147 | MGC |
| ETS2(-) | old | 0.426999446 | 0.825129232 | MGC |
| ETS2(-) | young | 0.439402214 | 1.399094781 | MGC |
| ETS2(+) | old | 0.491419223 | 1.297473297 | MGC |
| ETS2(+) | young | 0.379845825 | 0.888982745 | MGC |
| ETV1(+) | old | 0.37920855 | 0.474712838 | MGC |
| ETV1(+) | young | 0.438793039 | 1.393877074 | MGC |
| ETV5(+) | old | 0.423315574 | 0.798118039 | MGC |
| ETV5(+) | young | 0.407625847 | 1.126924029 | MGC |
| FLI1(+) | old | 0.455729493 | 1.035786081 | MGC |
| FLI1(+) | young | 0.376981708 | 0.864451027 | MGC |
| FOS(+) | old | 0.457061802 | 1.045554947 | MGC |

| | | | | |
|---|---|---|---|---|
| FOS(+) | young | 0.413386535 | 1.176265437 | MGC |
| FOSB(+) | old | 0.453427 | 1.018903546 | MGC |
| FOSB(+) | young | 0.403046442 | 1.087700535 | MGC |
| FOSL1(+) | old | 0.470263602 | 1.142354289 | MGC |
| FOSL1(+) | young | 0.383100969 | 0.916863686 | MGC |
| FOSL2(+) | old | 0.500785181 | 1.366147154 | MGC |
| FOSL2(+) | young | 0.360659641 | 0.724649354 | MGC |
| GABPB1(-) | old | 0.364752705 | 0.368718484 | MGC |
| GABPB1(-) | young | 0.404189902 | 1.097494487 | MGC |
| HES2(+) | old | 0.50281443 | 1.381026182 | MGC |
| HES2(+) | young | 0.340276014 | 0.550059631 | MGC |
| HES7(-) | old | 0.473319363 | 1.164759991 | MGC |
| HES7(-) | young | 0.386659155 | 0.947340237 | MGC |
| HES7(+) | old | 0.458013564 | 1.052533532 | MGC |
| HES7(+) | young | 0.351133339 | 0.643054734 | MGC |
| HMGA1(+) | old | 0.48301978 | 1.235886201 | MGC |
| HMGA1(+) | young | 0.364637854 | 0.758723524 | MGC |
| HMX1(+) | old | 0.437131604 | 0.899421088 | MGC |
| HMX1(+) | young | 0.42944244 | 1.313787382 | MGC |
| IKZF1(-) | old | 0.469140007 | 1.134115767 | MGC |
| IKZF1(-) | young | 0.394180342 | 1.011760665 | MGC |
| IKZF1(+) | old | 0.456037044 | 1.038041132 | MGC |
| IKZF1(+) | young | 0.388125044 | 0.959895862 | MGC |
| ILF2(+) | old | 0.452200525 | 1.009910686 | MGC |
| ILF2(+) | young | 0.405652353 | 1.110020669 | MGC |
| IRF5(-) | old | 0.439501711 | 0.916799384 | MGC |
| IRF5(-) | young | 0.393745208 | 1.00803366 | MGC |
| IRF8(+) | old | 0.471048762 | 1.148111299 | MGC |
| IRF8(+) | young | 0.349358159 | 0.627849965 | MGC |
| JUNB(+) | old | 0.463867166 | 1.095453807 | MGC |
| JUNB(+) | young | 0.405397099 | 1.107834369 | MGC |
| JUND(+) | old | 0.471318943 | 1.150092346 | MGC |
| JUND(+) | young | 0.400668067 | 1.067329291 | MGC |
| KLF4(-) | old | 0.483452986 | 1.239062587 | MGC |
| KLF4(-) | young | 0.362197898 | 0.737824825 | MGC |
| KLF6(+) | old | 0.482868936 | 1.234780165 | MGC |
| KLF6(+) | young | 0.392199325 | 0.99479287 | MGC |
| MAFB(-) | old | 0.435926764 | 0.890586855 | MGC |
| MAFB(-) | young | 0.428291578 | 1.303930025 | MGC |
| MAFF(+) | old | 0.495489182 | 1.327315388 | MGC |
| MAFF(+) | young | 0.345484855 | 0.594674369 | MGC |
| MAX(+) | old | 0.458789113 | 1.058220077 | MGC |
| MAX(+) | young | 0.395994075 | 1.027295637 | MGC |
| MEIS1(+) | old | 0.424222169 | 0.804765451 | MGC |
| MEIS1(+) | young | 0.394991864 | 1.018711511 | MGC |
| MITF(+) | old | 0.473161408 | 1.163601823 | MGC |
| MITF(+) | young | 0.369457874 | 0.800007932 | MGC |
| MXI1(+) | old | 0.472209862 | 1.156624814 | MGC |
| MXI1(+) | young | 0.391652924 | 0.990112842 | MGC |
| MYBL2(+) | old | 0.435043326 | 0.884109241 | MGC |
| MYBL2(+) | young | 0.406346269 | 1.11596419 | MGC |
| NFATC2(+) | old | 0.453440456 | 1.01900221 | MGC |
| NFATC2(+) | young | 0.366543553 | 0.775046201 | MGC |
| NFIA(+) | old | 0.381775759 | 0.493536336 | MGC |

| | | | | |
|---|---|---|---|---|
| NFIA(+) | young | 0.428684771 | 1.307297797 | MGC |
| NFIB(+) | old | 0.395755024 | 0.596036271 | MGC |
| NFIB(+) | young | 0.434494255 | 1.35705716 | MGC |
| NFKB1(+) | old | 0.484161094 | 1.244254636 | MGC |
| NFKB1(+) | young | 0.373080423 | 0.831035766 | MGC |
| NR3C1(+) | old | 0.453007512 | 1.015827741 | MGC |
| NR3C1(+) | young | 0.384548902 | 0.929265512 | MGC |
| PBX1(+) | old | 0.417220613 | 0.753428055 | MGC |
| PBX1(+) | young | 0.424349696 | 1.270167038 | MGC |
| PRDM1(+) | old | 0.472503098 | 1.158774903 | MGC |
| PRDM1(+) | young | 0.382325444 | 0.910221161 | MGC |
| RARB(+) | old | 0.406499663 | 0.674819013 | MGC |
| RARB(+) | young | 0.422104444 | 1.250936023 | MGC |
| RAX(+) | old | 0.405816172 | 0.669807458 | MGC |
| RAX(+) | young | 0.451045479 | 1.498821606 | MGC |
| REL(+) | old | 0.473004519 | 1.162451468 | MGC |
| REL(+) | young | 0.361682721 | 0.733412234 | MGC |
| RELB(+) | old | 0.490656807 | 1.291883046 | MGC |
| RELB(+) | young | 0.371485846 | 0.817377906 | MGC |
| RUNX1(-) | old | 0.424400808 | 0.806075278 | MGC |
| RUNX1(-) | young | 0.418798413 | 1.222619226 | MGC |
| RXRA(+) | old | 0.453980531 | 1.022962191 | MGC |
| RXRA(+) | young | 0.353687771 | 0.664933932 | MGC |
| SOX2(+) | old | 0.384333814 | 0.512292722 | MGC |
| SOX2(+) | young | 0.461372125 | 1.58727133 | MGC |
| SPI1(-) | old | 0.443944545 | 0.949375497 | MGC |
| SPI1(-) | young | 0.417574891 | 1.212139515 | MGC |
| SPI1(+) | old | 0.473791793 | 1.168223979 | MGC |
| SPI1(+) | young | 0.39316069 | 1.003027146 | MGC |
| STAT1(+) | old | 0.433506634 | 0.872841777 | MGC |
| STAT1(+) | young | 0.414836099 | 1.188681235 | MGC |
| XBP1(+) | old | 0.477234468 | 1.193466649 | MGC |
| XBP1(+) | young | 0.392666504 | 0.998794347 | MGC |
| YY1(+) | old | 0.457838103 | 1.051247002 | MGC |
| YY1(+) | young | 0.410124328 | 1.148324003 | MGC |
| ZBTB21(+) | old | 0.505094524 | 1.397744481 | MGC |
| ZBTB21(+) | young | 0.338167754 | 0.532001977 | MGC |
| ZBTB43(+) | old | 0.470772548 | 1.146086019 | MGC |
| ZBTB43(+) | young | 0.384627054 | 0.929934896 | MGC |
| ZNF148(+) | old | 0.446478221 | 0.967953129 | MGC |
| ZNF148(+) | young | 0.400563208 | 1.066431156 | MGC |
| ZNF267(+) | old | 0.475250923 | 1.178922739 | MGC |
| ZNF267(+) | young | 0.385867742 | 0.940561634 | MGC |
| ZNF480(+) | old | 0.39101888 | 0.561309525 | MGC |
| ZNF480(+) | young | 0.419112968 | 1.225313447 | MGC |
| ZNF771(+) | old | 0.379942278 | 0.480092736 | MGC |
| ZNF771(+) | young | 0.399251556 | 1.055196599 | MGC |
| ZXDA(-) | old | 0.470439678 | 1.143645323 | MGC |
| ZXDA(-) | young | 0.262349193 | 0 | MGC |
| ATF3(+) | old | 0.616106209 | 1.105315144 | Microglia |
| ATF3(+) | young | 0.257614666 | 1.191734189 | Microglia |
| ATF4(+) | old | 0.641054895 | 1.229831582 | Microglia |
| ATF4(+) | young | 0.217588691 | 0.641019919 | Microglia |
| BACH1(+) | old | 0.643042973 | 1.239753883 | Microglia |

| | | | | |
|---|---|---|---|---|
| BACH1(+) | young | 0.215706891 | 0.615128371 | Microglia |
| BATF3(-) | old | 0.546778089 | 0.759305321 | Microglia |
| BATF3(-) | young | 0.251902165 | 1.113136332 | Microglia |
| BCL11A(-) | old | 0.549330557 | 0.772044439 | Microglia |
| BCL11A(-) | young | 0.250808939 | 1.098094725 | Microglia |
| BCLAF1(+) | old | 0.630042718 | 1.17487089 | Microglia |
| BCLAF1(+) | young | 0.223755279 | 0.725865516 | Microglia |
| BHLHE40(+) | old | 0.645505044 | 1.252041838 | Microglia |
| BHLHE40(+) | young | 0.236541216 | 0.901786233 | Microglia |
| BHLHE41(+) | old | 0.639393432 | 1.221539381 | Microglia |
| BHLHE41(+) | young | 0.241272406 | 0.966882302 | Microglia |
| CEBPA(+) | old | 0.613870818 | 1.094158528 | Microglia |
| CEBPA(+) | young | 0.257631304 | 1.191963105 | Microglia |
| CEBPB(+) | old | 0.635689972 | 1.20305578 | Microglia |
| CEBPB(+) | young | 0.234358945 | 0.87176053 | Microglia |
| CEBPD(+) | old | 0.60566798 | 1.053218972 | Microglia |
| CEBPD(+) | young | 0.268381275 | 1.33987112 | Microglia |
| CHD1(+) | old | 0.636606993 | 1.207632541 | Microglia |
| CHD1(+) | young | 0.217696657 | 0.642505408 | Microglia |
| CREB3(+) | old | 0.619576975 | 1.122637397 | Microglia |
| CREB3(+) | young | 0.244767453 | 1.014970379 | Microglia |
| CREM(+) | old | 0.631974624 | 1.184512844 | Microglia |
| CREM(+) | young | 0.243628869 | 0.999304692 | Microglia |
| E2F1(+) | old | 0.597824235 | 1.014071611 | Microglia |
| E2F1(+) | young | 0.258701309 | 1.206685225 | Microglia |
| E2F8(-) | old | 0.611061461 | 1.080137305 | Microglia |
| E2F8(-) | young | 0.232220166 | 0.842333238 | Microglia |
| E2F8(+) | old | 0.595597085 | 1.002956128 | Microglia |
| E2F8(+) | young | 0.260612283 | 1.232978163 | Microglia |
| ELF1(-) | old | 0.609141845 | 1.070556692 | Microglia |
| ELF1(-) | young | 0.247485592 | 1.052369044 | Microglia |
| ELF1(+) | old | 0.632670809 | 1.187987431 | Microglia |
| ELF1(+) | young | 0.232035432 | 0.839791502 | Microglia |
| EP300(+) | old | 0.617235328 | 1.110950466 | Microglia |
| EP300(+) | young | 0.227235577 | 0.773750674 | Microglia |
| ETS1(-) | old | 0.602637059 | 1.038091943 | Microglia |
| ETS1(-) | young | 0.216999089 | 0.632907625 | Microglia |
| ETS1(+) | old | 0.607407042 | 1.061898459 | Microglia |
| ETS1(+) | young | 0.22213789 | 0.703611989 | Microglia |
| ETS2(-) | old | 0.600914509 | 1.029494865 | Microglia |
| ETS2(-) | young | 0.252707963 | 1.124223239 | Microglia |
| ETS2(+) | old | 0.637111842 | 1.21015219 | Microglia |
| ETS2(+) | young | 0.236884286 | 0.906506502 | Microglia |
| ETV5(+) | old | 0.5942036 | 0.996001382 | Microglia |
| ETV5(+) | young | 0.256718748 | 1.179407321 | Microglia |
| ETV6(+) | old | 0.603628281 | 1.043039036 | Microglia |
| ETV6(+) | young | 0.215942698 | 0.618372829 | Microglia |
| FLI1(-) | old | 0.624243201 | 1.145926073 | Microglia |
| FLI1(-) | young | 0.243118663 | 0.992284808 | Microglia |
| FLI1(+) | old | 0.625779216 | 1.153592173 | Microglia |
| FLI1(+) | young | 0.225701743 | 0.752646763 | Microglia |
| FOS(+) | old | 0.602951873 | 1.039663149 | Microglia |
| FOS(+) | young | 0.269495938 | 1.355207679 | Microglia |
| FOSB(+) | old | 0.586113987 | 0.95562692 | Microglia |

| | | | | |
|---|---|---|---|---|
| FOSB(+) | young | 0.267944813 | 1.333865875 | Microglia |
| FOSL1(+) | old | 0.637196206 | 1.210573244 | Microglia |
| FOSL1(+) | young | 0.207452975 | 0.501563391 | Microglia |
| FOSL2(+) | old | 0.646700905 | 1.25801026 | Microglia |
| FOSL2(+) | young | 0.225696948 | 0.752580795 | Microglia |
| FOXC1(-) | old | 0.567158092 | 0.86101991 | Microglia |
| FOXC1(-) | young | 0.256112815 | 1.171070338 | Microglia |
| FOXM1(+) | old | 0.344696768 | 0 | Microglia |
| FOXM1(+) | young | 0.270615521 | 1.370611944 | Microglia |
| GABPB1(-) | old | 0.464521436 | 0.348770465 | Microglia |
| GABPB1(-) | young | 0.255719835 | 1.165663354 | Microglia |
| GATA3(-) | old | 0.617622692 | 1.112883764 | Microglia |
| GATA3(-) | young | 0.186001016 | 0.206407547 | Microglia |
| HES2(+) | old | 0.596249761 | 1.006213567 | Microglia |
| HES2(+) | young | 0.235263573 | 0.884207236 | Microglia |
| HES7(-) | old | 0.628006997 | 1.164710807 | Microglia |
| HES7(-) | young | 0.241974631 | 0.976544168 | Microglia |
| HMGA1(-) | old | 0.577464236 | 0.912456863 | Microglia |
| HMGA1(-) | young | 0.246622947 | 1.040499977 | Microglia |
| HMGA1(+) | old | 0.615708176 | 1.1033286 | Microglia |
| HMGA1(+) | young | 0.238885563 | 0.934041923 | Microglia |
| HOXA5(+) | old | 0.616754193 | 1.10854917 | Microglia |
| HOXA5(+) | young | 0.239448051 | 0.941781144 | Microglia |
| HOXB2(-) | old | 0.607680909 | 1.063265302 | Microglia |
| HOXB2(-) | young | 0.234519227 | 0.873965848 | Microglia |
| IKZF1(-) | old | 0.622904293 | 1.139243714 | Microglia |
| IKZF1(-) | young | 0.244300563 | 1.008546475 | Microglia |
| IKZF1(+) | old | 0.628565074 | 1.167496115 | Microglia |
| IKZF1(+) | young | 0.21496079 | 0.604862829 | Microglia |
| ILF2(+) | old | 0.609129522 | 1.070495185 | Microglia |
| ILF2(+) | young | 0.258921016 | 1.209708157 | Microglia |
| ING4(+) | old | 0.597166312 | 1.010787982 | Microglia |
| ING4(+) | young | 0.239452226 | 0.941838591 | Microglia |
| IRF5(-) | old | 0.600449147 | 1.027172288 | Microglia |
| IRF5(-) | young | 0.242771604 | 0.987509651 | Microglia |
| IRF8(+) | old | 0.62339454 | 1.141690486 | Microglia |
| IRF8(+) | young | 0.243100621 | 0.992036579 | Microglia |
| JUNB(+) | old | 0.603430526 | 1.042052058 | Microglia |
| JUNB(+) | young | 0.271025681 | 1.376255299 | Microglia |
| JUND(+) | old | 0.62212459 | 1.135352291 | Microglia |
| JUND(+) | young | 0.258920599 | 1.209702425 | Microglia |
| KLF4(-) | old | 0.622997325 | 1.139708028 | Microglia |
| KLF4(-) | young | 0.246626366 | 1.040547023 | Microglia |
| KLF6(+) | old | 0.632374375 | 1.186507958 | Microglia |
| KLF6(+) | young | 0.239313264 | 0.939926622 | Microglia |
| MAFB(-) | old | 0.498741177 | 0.519557826 | Microglia |
| MAFB(-) | young | 0.262929708 | 1.264863434 | Microglia |
| MAFB(+) | old | 0.631568123 | 1.182484034 | Microglia |
| MAFB(+) | young | 0.185718467 | 0.202519985 | Microglia |
| MAFF(+) | old | 0.625054345 | 1.149974411 | Microglia |
| MAFF(+) | young | 0.239079924 | 0.936716111 | Microglia |
| MAX(+) | old | 0.640596918 | 1.227545865 | Microglia |
| MAX(+) | young | 0.228052136 | 0.784985642 | Microglia |
| MEIS1(+) | old | 0.597134504 | 1.010629233 | Microglia |

| | | | | |
|---|---|---|---|---|
| MEIS1(+) | young | 0.215516065 | 0.612502813 | Microglia |
| MITF(+) | old | 0.647572403 | 1.262359821 | Microglia |
| MITF(+) | young | 0.198286389 | 0.375441049 | Microglia |
| MSX2(-) | old | 0.524471023 | 0.647972953 | Microglia |
| MSX2(-) | young | 0.254650877 | 1.150955646 | Microglia |
| MXI1(+) | old | 0.649173351 | 1.270349997 | Microglia |
| MXI1(+) | young | 0.221398272 | 0.693435646 | Microglia |
| MYBL2(+) | old | 0.580118141 | 0.925702241 | Microglia |
| MYBL2(+) | young | 0.268717557 | 1.344498008 | Microglia |
| NEUROD1(+) | old | 0.607195867 | 1.060844508 | Microglia |
| NEUROD1(+) | young | 0.219207112 | 0.663287641 | Microglia |
| NFATC2(+) | old | 0.597810332 | 1.014002223 | Microglia |
| NFATC2(+) | young | 0.263986159 | 1.279399059 | Microglia |
| NFE2L2(+) | old | 0.540439651 | 0.727670805 | Microglia |
| NFE2L2(+) | young | 0.2464057 | 1.037510892 | Microglia |
| NFIA(+) | old | 0.600952899 | 1.029686465 | Microglia |
| NFIA(+) | young | 0.214857456 | 0.603441071 | Microglia |
| NFKB1(+) | old | 0.620172645 | 1.12561033 | Microglia |
| NFKB1(+) | young | 0.235784772 | 0.891378374 | Microglia |
| NR3C1(+) | old | 0.629525274 | 1.172288379 | Microglia |
| NR3C1(+) | young | 0.224193038 | 0.731888614 | Microglia |
| PBX1(+) | old | 0.609328211 | 1.071486825 | Microglia |
| PBX1(+) | young | 0.242748474 | 0.987191404 | Microglia |
| POU2F2(-) | old | 0.399120467 | 0.02236067 | Microglia |
| POU2F2(-) | young | 0.276128614 | 1.446466164 | Microglia |
| PRDM1(+) | old | 0.617282189 | 1.111184348 | Microglia |
| PRDM1(+) | young | 0.241172707 | 0.965510558 | Microglia |
| RARB(+) | old | 0.622478245 | 1.137117352 | Microglia |
| RARB(+) | young | 0.225955457 | 0.756137602 | Microglia |
| RAX(+) | old | 0.504187055 | 0.546737667 | Microglia |
| RAX(+) | young | 0.255307004 | 1.159983243 | Microglia |
| REL(+) | old | 0.612611886 | 1.087875321 | Microglia |
| REL(+) | young | 0.230281984 | 0.81566595 | Microglia |
| RELB(+) | old | 0.631221791 | 1.180755527 | Microglia |
| RELB(+) | young | 0.238005414 | 0.921932022 | Microglia |
| RFXANK(+) | old | 0.324826874 | 0 | Microglia |
| RFXANK(+) | young | 0.258950441 | 1.210113016 | Microglia |
| RUNX1(-) | old | 0.602263547 | 1.036227783 | Microglia |
| RUNX1(-) | young | 0.242581537 | 0.984894542 | Microglia |
| SOX2(+) | old | 0.61317644 | 1.090692958 | Microglia |
| SOX2(+) | young | 0.219123449 | 0.662136536 | Microglia |
| SPI1(-) | old | 0.595286789 | 1.001407472 | Microglia |
| SPI1(-) | young | 0.261870854 | 1.250294747 | Microglia |
| SPI1(+) | old | 0.627179285 | 1.160579779 | Microglia |
| SPI1(+) | young | 0.250915582 | 1.099562016 | Microglia |
| STAT1(+) | old | 0.59660451 | 1.007984084 | Microglia |
| STAT1(+) | young | 0.25736124 | 1.188247316 | Microglia |
| USF1(+) | old | 0.581445862 | 0.932328763 | Microglia |
| USF1(+) | young | 0.24919091 | 1.075832381 | Microglia |
| VDR(+) | old | 0.456450847 | 0.308490949 | Microglia |
| VDR(+) | young | 0.255006737 | 1.155851891 | Microglia |
| XBP1(+) | old | 0.631055661 | 1.17992639 | Microglia |
| XBP1(+) | young | 0.250015222 | 1.087174036 | Microglia |
| YY1(+) | old | 0.634452177 | 1.196878067 | Microglia |

| | | | | |
|---|---|---|---|---|
| YY1(+) | young | 0.23375256 | 0.863417333 | Microglia |
| ZBTB21(+) | old | 0.64521318 | 1.250585172 | Microglia |
| ZBTB21(+) | young | 0.220591734 | 0.68233855 | Microglia |
| ZBTB43(+) | old | 0.637394677 | 1.211563793 | Microglia |
| ZBTB43(+) | young | 0.214442659 | 0.597733909 | Microglia |
| ZNF148(+) | old | 0.629979074 | 1.17455325 | Microglia |
| ZNF148(+) | young | 0.218923635 | 0.659387302 | Microglia |
| ZNF200(-) | old | 0.586250289 | 0.956307186 | Microglia |
| ZNF200(-) | young | 0.262589887 | 1.26018786 | Microglia |
| ZNF267(+) | old | 0.626098545 | 1.155185911 | Microglia |
| ZNF267(+) | young | 0.230113618 | 0.813349406 | Microglia |
| ATF3(+) | old | 0.578996831 | 1.358837417 | Astrocyte |
| ATF3(+) | young | 0.288548199 | 0.462430004 | Astrocyte |
| ATF4(+) | old | 0.568590582 | 1.29332241 | Astrocyte |
| ATF4(+) | young | 0.300310817 | 0.586417619 | Astrocyte |
| BACH1(+) | old | 0.522166069 | 1.001045875 | Astrocyte |
| BACH1(+) | young | 0.332915317 | 0.930095721 | Astrocyte |
| BCLAF1(+) | old | 0.497986687 | 0.848818836 | Astrocyte |
| BCLAF1(+) | young | 0.359057127 | 1.20565178 | Astrocyte |
| BHLHE40(+) | old | 0.565298976 | 1.272599324 | Astrocyte |
| BHLHE40(+) | young | 0.299867714 | 0.581746961 | Astrocyte |
| BHLHE41(+) | old | 0.570982478 | 1.308381156 | Astrocyte |
| BHLHE41(+) | young | 0.29210444 | 0.4999157 | Astrocyte |
| CEBPA(+) | old | 0.572499445 | 1.317931584 | Astrocyte |
| CEBPA(+) | young | 0.285550572 | 0.430832564 | Astrocyte |
| CEBPB(+) | old | 0.583054771 | 1.384385141 | Astrocyte |
| CEBPB(+) | young | 0.270852837 | 0.275906415 | Astrocyte |
| CEBPD(+) | old | 0.576634191 | 1.343962858 | Astrocyte |
| CEBPD(+) | young | 0.285832818 | 0.43380767 | Astrocyte |
| CEBPG(+) | old | 0.344570091 | 0 | Astrocyte |
| CEBPG(+) | young | 0.340416595 | 1.009165329 | Astrocyte |
| CREB3(+) | old | 0.534339884 | 1.077689014 | Astrocyte |
| CREB3(+) | young | 0.328868695 | 0.887441013 | Astrocyte |
| CREM(+) | old | 0.559313035 | 1.234913412 | Astrocyte |
| CREM(+) | young | 0.310404602 | 0.692814374 | Astrocyte |
| E2F8(-) | old | 0.49553964 | 0.833412874 | Astrocyte |
| E2F8(-) | young | 0.343913013 | 1.046020435 | Astrocyte |
| ELF1(-) | old | 0.522438542 | 1.002761296 | Astrocyte |
| ELF1(-) | young | 0.343089074 | 1.037335439 | Astrocyte |
| ELF1(+) | old | 0.54002722 | 1.113494987 | Astrocyte |
| ELF1(+) | young | 0.323386363 | 0.829652754 | Astrocyte |
| ELK3(+) | old | 0.532589391 | 1.066668373 | Astrocyte |
| ELK3(+) | young | 0.305532785 | 0.641461436 | Astrocyte |
| EP300(+) | old | 0.484336099 | 0.762878324 | Astrocyte |
| EP300(+) | young | 0.357313616 | 1.18727375 | Astrocyte |
| ERF(+) | old | 0.315941161 | 0 | Astrocyte |
| ERF(+) | young | 0.339693912 | 1.001547655 | Astrocyte |
| ETS1(-) | old | 0.552003789 | 1.188896319 | Astrocyte |
| ETS1(-) | young | 0.280703496 | 0.379740421 | Astrocyte |
| ETS2(-) | old | 0.520295174 | 0.98926721 | Astrocyte |
| ETS2(-) | young | 0.351297546 | 1.123859451 | Astrocyte |
| ETS2(+) | old | 0.570564897 | 1.305752176 | Astrocyte |
| ETS2(+) | young | 0.300518785 | 0.58860978 | Astrocyte |
| ETV1(+) | old | 0.406363371 | 0.271982517 | Astrocyte |

| | | | | |
|---|---|---|---|---|
| ETV1(+) | young | 0.397103084 | 1.606687286 | Astrocyte |
| ETV5(+) | old | 0.475975636 | 0.710243046 | Astrocyte |
| ETV5(+) | young | 0.353996431 | 1.152307912 | Astrocyte |
| FLI1(-) | old | 0.548832256 | 1.168929179 | Astrocyte |
| FLI1(-) | young | 0.286092187 | 0.436541631 | Astrocyte |
| FLI1(+) | old | 0.492603453 | 0.814927413 | Astrocyte |
| FLI1(+) | young | 0.351313229 | 1.124024763 | Astrocyte |
| FOS(+) | old | 0.555301875 | 1.209660201 | Astrocyte |
| FOS(+) | young | 0.317322628 | 0.765736029 | Astrocyte |
| FOSB(+) | old | 0.558030991 | 1.226841998 | Astrocyte |
| FOSB(+) | young | 0.298179172 | 0.563948338 | Astrocyte |
| FOSL1(+) | old | 0.542667943 | 1.130120287 | Astrocyte |
| FOSL1(+) | young | 0.304891758 | 0.634704494 | Astrocyte |
| FOSL2(+) | old | 0.576363725 | 1.342260075 | Astrocyte |
| FOSL2(+) | young | 0.28604856 | 0.436081773 | Astrocyte |
| FOXC1(-) | old | 0.409644391 | 0.292638956 | Astrocyte |
| FOXC1(-) | young | 0.356004319 | 1.173472693 | Astrocyte |
| HES2(+) | old | 0.551779963 | 1.18748717 | Astrocyte |
| HES2(+) | young | 0.298222605 | 0.564406158 | Astrocyte |
| HES7(-) | old | 0.542513979 | 1.129150969 | Astrocyte |
| HES7(-) | young | 0.306119202 | 0.647642758 | Astrocyte |
| HMBOX1(+) | old | 0.354075612 | 0 | Astrocyte |
| HMBOX1(+) | young | 0.345780917 | 1.065709669 | Astrocyte |
| HMGA1(+) | old | 0.560228579 | 1.240677437 | Astrocyte |
| HMGA1(+) | young | 0.281390125 | 0.386978053 | Astrocyte |
| HMGA2(+) | old | 0.427202344 | 0.403179212 | Astrocyte |
| HMGA2(+) | young | 0.342741858 | 1.033675502 | Astrocyte |
| HMX1(+) | old | 0.524042624 | 1.01286017 | Astrocyte |
| HMX1(+) | young | 0.342523658 | 1.031375494 | Astrocyte |
| HOXA5(+) | old | 0.589439618 | 1.42458246 | Astrocyte |
| HOXA5(+) | young | 0.226967749 | 0 | Astrocyte |
| HOXB2(-) | old | 0.528113535 | 1.038489555 | Astrocyte |
| HOXB2(-) | young | 0.293912123 | 0.51897016 | Astrocyte |
| IKZF1(-) | old | 0.536429155 | 1.090842517 | Astrocyte |
| IKZF1(-) | young | 0.328329316 | 0.881755523 | Astrocyte |
| ILF2(+) | old | 0.538996204 | 1.107003983 | Astrocyte |
| ILF2(+) | young | 0.322709261 | 0.822515545 | Astrocyte |
| ING4(+) | old | 0.540193265 | 1.114540366 | Astrocyte |
| ING4(+) | young | 0.285587955 | 0.43122662 | Astrocyte |
| JUNB(+) | old | 0.569138284 | 1.2967706 | Astrocyte |
| JUNB(+) | young | 0.304477315 | 0.630335925 | Astrocyte |
| JUND(+) | old | 0.563366556 | 1.260433313 | Astrocyte |
| JUND(+) | young | 0.308219956 | 0.669786414 | Astrocyte |
| KLF4(-) | old | 0.54115033 | 1.120565794 | Astrocyte |
| KLF4(-) | young | 0.320917753 | 0.803631581 | Astrocyte |
| KLF6(+) | old | 0.553315854 | 1.197156734 | Astrocyte |
| KLF6(+) | young | 0.320409973 | 0.798279173 | Astrocyte |
| LCORL(+) | old | 0.537481094 | 1.097465248 | Astrocyte |
| LCORL(+) | young | 0.234726138 | 0 | Astrocyte |
| MAFB(-) | old | 0.537140457 | 1.095320688 | Astrocyte |
| MAFB(-) | young | 0.33104285 | 0.91035839 | Astrocyte |
| MAFF(+) | old | 0.561941272 | 1.251460103 | Astrocyte |
| MAFF(+) | young | 0.293466527 | 0.514273208 | Astrocyte |
| MAFG(+) | old | 0.536442942 | 1.090929312 | Astrocyte |

| | | | | |
|---|---|---|---|---|
| MAFG(+) | young | 0.267383102 | 0.239332575 | Astrocyte |
| MAX(+) | old | 0.53367205 | 1.073484506 | Astrocyte |
| MAX(+) | young | 0.323906732 | 0.835137872 | Astrocyte |
| MEIS1(+) | old | 0.479914543 | 0.73504137 | Astrocyte |
| MEIS1(+) | young | 0.350551745 | 1.115998099 | Astrocyte |
| MXI1(+) | old | 0.538093538 | 1.101321033 | Astrocyte |
| MXI1(+) | young | 0.323418939 | 0.82999613 | Astrocyte |
| MYB(-) | old | 0.25097167 | 0 | Astrocyte |
| MYB(-) | young | 0.373658204 | 1.359559083 | Astrocyte |
| MYBL2(+) | old | 0.531225232 | 1.058079985 | Astrocyte |
| MYBL2(+) | young | 0.290913242 | 0.487359493 | Astrocyte |
| MYC(+) | old | 0.400454194 | 0.23477989 | Astrocyte |
| MYC(+) | young | 0.364206061 | 1.259925755 | Astrocyte |
| NEUROD1(+) | old | 0.496118695 | 0.837058454 | Astrocyte |
| NEUROD1(+) | young | 0.347937445 | 1.088441235 | Astrocyte |
| NFIA(+) | old | 0.445374571 | 0.517586777 | Astrocyte |
| NFIA(+) | young | 0.385195146 | 1.48116788 | Astrocyte |
| NFIB(+) | old | 0.471016835 | 0.679023736 | Astrocyte |
| NFIB(+) | young | 0.37996318 | 1.426018686 | Astrocyte |
| NFKB1(+) | old | 0.547893998 | 1.163022157 | Astrocyte |
| NFKB1(+) | young | 0.311170761 | 0.700890315 | Astrocyte |
| NR2F1(-) | old | 0.204798883 | 0 | Astrocyte |
| NR2F1(-) | young | 0.353446009 | 1.146506007 | Astrocyte |
| PBX1(+) | old | 0.462046543 | 0.622549137 | Astrocyte |
| PBX1(+) | young | 0.384562807 | 1.474502515 | Astrocyte |
| PLAGL2(+) | old | 0.42412413 | 0.383799587 | Astrocyte |
| PLAGL2(+) | young | 0.355278552 | 1.165822516 | Astrocyte |
| PRDM1(+) | old | 0.527786551 | 1.036430951 | Astrocyte |
| PRDM1(+) | young | 0.328434344 | 0.882862605 | Astrocyte |
| RARB(+) | old | 0.473139698 | 0.692388723 | Astrocyte |
| RARB(+) | young | 0.37653032 | 1.389833533 | Astrocyte |
| RAX(+) | old | 0.481854654 | 0.747255794 | Astrocyte |
| RAX(+) | young | 0.369799262 | 1.31888267 | Astrocyte |
| REL(+) | old | 0.528266898 | 1.039455089 | Astrocyte |
| REL(+) | young | 0.30268166 | 0.611408246 | Astrocyte |
| RELB(+) | old | 0.552831158 | 1.194105216 | Astrocyte |
| RELB(+) | young | 0.292680387 | 0.505986654 | Astrocyte |
| RFX1(-) | old | 0.391200201 | 0.176519183 | Astrocyte |
| RFX1(-) | young | 0.351296422 | 1.123847602 | Astrocyte |
| RUNX1(-) | old | 0.498942465 | 0.854836166 | Astrocyte |
| RUNX1(-) | young | 0.35376017 | 1.149817523 | Astrocyte |
| RXRA(+) | old | 0.54135119 | 1.121830357 | Astrocyte |
| RXRA(+) | young | 0.292487185 | 0.503950143 | Astrocyte |
| SOX10(+) | old | 0.342288387 | 0 | Astrocyte |
| SOX10(+) | young | 0.366595672 | 1.285114214 | Astrocyte |
| SOX15(+) | old | 0.489823078 | 0.797422898 | Astrocyte |
| SOX15(+) | young | 0.343819139 | 1.045030922 | Astrocyte |
| SOX2(+) | old | 0.465004352 | 0.641170722 | Astrocyte |
| SOX2(+) | young | 0.387246767 | 1.50279365 | Astrocyte |
| SPI1(-) | old | 0.530131925 | 1.051196813 | Astrocyte |
| SPI1(-) | young | 0.324142676 | 0.837624916 | Astrocyte |
| SPI1(+) | old | 0.573965474 | 1.32716132 | Astrocyte |
| SPI1(+) | young | 0.293454753 | 0.514149106 | Astrocyte |
| SREBF1(+) | old | 0.533113687 | 1.069969202 | Astrocyte |

| | | | | |
|---|---|---|---|---|
| SREBF1(+) | young | 0.235823685 | 0 | Astrocyte |
| STAT1(+) | old | 0.469993718 | 0.67258246 | Astrocyte |
| STAT1(+) | young | 0.367493152 | 1.294574385 | Astrocyte |
| TBL1XR1(+) | old | 0.430469146 | 0.423746141 | Astrocyte |
| TBL1XR1(+) | young | 0.344581309 | 1.053064821 | Astrocyte |
| USF1(-) | old | 0.349660055 | 0 | Astrocyte |
| USF1(-) | young | 0.341451792 | 1.020077147 | Astrocyte |
| XBP1(+) | old | 0.570792327 | 1.307184017 | Astrocyte |
| XBP1(+) | young | 0.30477781 | 0.633503389 | Astrocyte |
| YY1(+) | old | 0.531022037 | 1.056800724 | Astrocyte |
| YY1(+) | young | 0.335073358 | 0.952843243 | Astrocyte |
| YY2(-) | old | 0.437226283 | 0.4662873 | Astrocyte |
| YY2(-) | young | 0.385953182 | 1.489158205 | Astrocyte |
| ZBTB21(+) | old | 0.579228911 | 1.360298534 | Astrocyte |
| ZBTB21(+) | young | 0.2609558 | 0.171583559 | Astrocyte |
| ZBTB43(+) | old | 0.54249539 | 1.129033939 | Astrocyte |
| ZBTB43(+) | young | 0.309020722 | 0.678227144 | Astrocyte |
| ZNF148(+) | old | 0.502374683 | 0.876444505 | Astrocyte |
| ZNF148(+) | young | 0.345309839 | 1.060744122 | Astrocyte |
| ZNF16(+) | old | 0.084374125 | 0 | Astrocyte |
| ZNF16(+) | young | 0.355597282 | 1.169182185 | Astrocyte |
| ZNF267(+) | old | 0.532189979 | 1.064153778 | Astrocyte |
| ZNF267(+) | young | 0.324984393 | 0.846497294 | Astrocyte |
| ZNF528(-) | old | 0.453507017 | 0.568786525 | Astrocyte |
| ZNF528(-) | young | 0.357919268 | 1.193657819 | Astrocyte |
| ZNF771(+) | old | 0.410279619 | 0.296638185 | Astrocyte |
| ZNF771(+) | young | 0.34993589 | 1.109506486 | Astrocyte |
