## Supplementary material for "Interpretable Aging Signatures in Human Retinal Cell Types Revealed by Single-Cell RNA Sequencing and Sparse Logistic Regression": Table S15

Table S15: Transcription factor regulon activity across retinal photoreceptor populations during aging.

| Topic | cellType | RSS | Z | CellType |
|---|---|---|---|---|
| ATF4(+) | old | 0.375867845 | 1.299198981 | Rod |
| ATF4(+) | young | 0.491980334 | 1.875714707 | Rod |
| BACH2(+) | old | 0.424471266 | 1.675828113 | Rod |
| BACH2(+) | young | 0.441175769 | 1.541595518 | Rod |
| CEBPB(+) | old | 0.3527919 | 1.120382905 | Rod |
| CEBPB(+) | young | 0.405662949 | 1.308043392 | Rod |
| CEBPD(+) | old | 0.42730474 | 1.697784769 | Rod |
| CEBPD(+) | young | 0.397063083 | 1.251485866 | Rod |
| CHD2(+) | old | 0.424457747 | 1.67572335 | Rod |
| CHD2(+) | young | 0.443400934 | 1.556229451 | Rod |
| CREM(+) | old | 0.391639075 | 1.421410636 | Rod |
| CREM(+) | young | 0.356570895 | 0.985186629 | Rod |
| CRX(+) | old | 0.404734236 | 1.522885371 | Rod |
| CRX(+) | young | 0.46240361 | 1.681201656 | Rod |
| DDIT3(+) | old | 0.336824662 | 0.996652377 | Rod |
| DDIT3(+) | young | 0.515053211 | 2.027454826 | Rod |
| ELF1(+) | old | 0.411784988 | 1.577521825 | Rod |
| ELF1(+) | young | 0.407047947 | 1.317151908 | Rod |
| ETV1(+) | old | 0.346901955 | 1.074741573 | Rod |
| ETV1(+) | young | 0.303082743 | 0.633418679 | Rod |
| ETV2(+) | old | 0.352223904 | 1.115981491 | Rod |
| ETV2(+) | young | 0.353156788 | 0.962733556 | Rod |
| FIGLA(+) | old | 0.369991687 | 1.253664486 | Rod |
| FIGLA(+) | young | 0.454614072 | 1.629973308 | Rod |
| FOS(+) | old | 0.387655005 | 1.390537983 | Rod |
| FOS(+) | young | 0.492831715 | 1.881313866 | Rod |
| FOSB(+) | old | 0.36905261 | 1.246387555 | Rod |
| FOSB(+) | young | 0.504757377 | 1.959743676 | Rod |
| HOXC8(+) | old | 0.372438723 | 1.272626632 | Rod |
| HOXC8(+) | young | 0.208593915 | 0.012007391 | Rod |
| IRF1(+) | old | 0.363211034 | 1.201121037 | Rod |
| IRF1(+) | young | 0.443255134 | 1.555270588 | Rod |
| IRF7(+) | old | 0.296492444 | 0.684117004 | Rod |
| IRF7(+) | young | 0.501668633 | 1.939430369 | Rod |
| JUN(+) | old | 0.387156746 | 1.386676963 | Rod |
| JUN(+) | young | 0.487422718 | 1.845741279 | Rod |
| JUNB(+) | old | 0.382288694 | 1.348954299 | Rod |
| JUNB(+) | young | 0.496972668 | 1.908547082 | Rod |
| JUND(+) | old | 0.383878042 | 1.361270198 | Rod |
| JUND(+) | young | 0.495587989 | 1.899440659 | Rod |
| LHX4(-) | old | 0.341098565 | 1.02977096 | Rod |
| LHX4(-) | young | 0.248863482 | 0.276842546 | Rod |
| MAF(+) | old | 0.346305977 | 1.070123321 | Rod |
| MAF(+) | young | 0.268134701 | 0.40358084 | Rod |
| MAFF(+) | old | 0.402228021 | 1.503464648 | Rod |
| MAFF(+) | young | 0.388449993 | 1.194841383 | Rod |
| MEIS1(+) | old | 0.395808832 | 1.453722192 | Rod |
| MEIS1(+) | young | 0.457174509 | 1.646812168 | Rod |
| NEUROD1(+) | old | 0.39916217 | 1.479707293 | Rod |
| NEUROD1(+) | young | 0.481536655 | 1.807031239 | Rod |
| NR2E3(+) | old | 0.402797933 | 1.507880913 | Rod |
| NR2E3(+) | young | 0.458585153 | 1.656089353 | Rod |
| PAX6(+) | old | 0.413531907 | 1.591058745 | Rod |

| | | | | |
|---|---|---|---|---|
| PAX6(+) | young | 0.317537004 | 0.728477968 | Rod |
| POU2F1(+) | old | 0.416314734 | 1.612622941 | Rod |
| POU2F1(+) | young | 0.450003135 | 1.599649209 | Rod |
| RAX(+) | old | 0.414864163 | 1.601382433 | Rod |
| RAX(+) | young | 0.44263307 | 1.551179545 | Rod |
| RXRG(-) | old | 0.397194227 | 1.464457654 | Rod |
| RXRG(-) | young | 0.471940175 | 1.743919433 | Rod |
| RXRG(+) | old | 0.413962868 | 1.594398274 | Rod |
| RXRG(+) | young | 0.462913406 | 1.684554361 | Rod |
| SPI1(+) | old | 0.352080603 | 1.114871046 | Rod |
| SPI1(+) | young | 0.423722467 | 1.426812862 | Rod |
| TCF12(+) | old | 0.405977512 | 1.532519548 | Rod |
| TCF12(+) | young | 0.431112397 | 1.475413168 | Rod |
| TEAD4(-) | old | 0.347037294 | 1.075790318 | Rod |
| TEAD4(-) | young | 0.308853323 | 0.671369235 | Rod |
| YBX1(+) | old | 0.381884734 | 1.345824008 | Rod |
| YBX1(+) | young | 0.486469011 | 1.83946917 | Rod |
| ZNF200(+) | old | 0.240670355 | 0.251550237 | Rod |
| ZNF200(+) | young | 0.35963128 | 1.005313433 | Rod |
| ATF4(+) | old | 0.516637471 | 1.444212209 | Cone |
| ATF4(+) | young | 0.344184433 | 1.847219552 | Cone |
| BACH2(+) | old | 0.5405288 | 1.579250587 | Cone |
| BACH2(+) | young | 0.32188611 | 1.637868395 | Cone |
| CEBPB(+) | old | 0.489020093 | 1.288113482 | Cone |
| CEBPB(+) | young | 0.233899205 | 0.811790104 | Cone |
| CEBPD(+) | old | 0.55270838 | 1.648091913 | Cone |
| CEBPD(+) | young | 0.27490745 | 1.19680218 | Cone |
| CHD2(+) | old | 0.550662372 | 1.636527483 | Cone |
| CHD2(+) | young | 0.302847689 | 1.459123311 | Cone |
| CREM(+) | old | 0.474903034 | 1.208321149 | Cone |
| CREM(+) | young | 0.217857519 | 0.661180326 | Cone |
| CRX(+) | old | 0.525663355 | 1.495228239 | Cone |
| CRX(+) | young | 0.327841496 | 1.69378143 | Cone |
| DDIT3(+) | old | 0.505750578 | 1.382677395 | Cone |
| DDIT3(+) | young | 0.339702204 | 1.805137465 | Cone |
| ELF1(+) | old | 0.506440991 | 1.386579741 | Cone |
| ELF1(+) | young | 0.236840814 | 0.839407842 | Cone |
| ETV1(-) | old | 0.400635568 | 0.788547162 | Cone |
| ETV1(-) | young | 0.297044986 | 1.404643761 | Cone |
| ETV4(-) | old | 0.483779166 | 1.258490755 | Cone |
| ETV4(-) | young | 0.321367288 | 1.632997358 | Cone |
| FIGLA(+) | old | 0.502185296 | 1.362525735 | Cone |
| FIGLA(+) | young | 0.330376859 | 1.717585069 | Cone |
| FOS(+) | old | 0.5419483 | 1.587273874 | Cone |
| FOS(+) | young | 0.339792435 | 1.805984617 | Cone |
| FOSB(+) | old | 0.530727017 | 1.523849025 | Cone |
| FOSB(+) | young | 0.341004151 | 1.817360995 | Cone |
| IRF1(+) | old | 0.52077138 | 1.467577854 | Cone |
| IRF1(+) | young | 0.246103791 | 0.926374699 | Cone |
| IRF7(+) | old | 0.493365003 | 1.312671748 | Cone |
| IRF7(+) | young | 0.28533748 | 1.294726089 | Cone |
| JUN(+) | old | 0.542912815 | 1.592725501 | Cone |
| JUN(+) | young | 0.328513435 | 1.700090033 | Cone |
| JUNB(+) | old | 0.537247739 | 1.560705399 | Cone |

| | | | | |
|---|---|---|---|---|
| JUNB(+) | young | 0.342175452 | 1.828357929 | Cone |
| JUND(+) | old | 0.535715639 | 1.552045675 | Cone |
| JUND(+) | young | 0.343642179 | 1.842128514 | Cone |
| LHX4(-) | old | 0.508278691 | 1.396966774 | Cone |
| LHX4(-) | young | 0.262754173 | 1.082699307 | Cone |
| LHX9(-) | old | 0.476872921 | 1.219455332 | Cone |
| LHX9(-) | young | 0.240726048 | 0.875884952 | Cone |
| MAFF(+) | old | 0.514484216 | 1.432041593 | Cone |
| MAFF(+) | young | 0.235949375 | 0.831038438 | Cone |
| MEIS1(+) | old | 0.527877125 | 1.507740891 | Cone |
| MEIS1(+) | young | 0.318953775 | 1.610337726 | Cone |
| NEUROD1(+) | old | 0.548351921 | 1.623468372 | Cone |
| NEUROD1(+) | young | 0.331223593 | 1.725534759 | Cone |
| NHLH1(+) | old | 0.475849572 | 1.213671167 | Cone |
| NHLH1(+) | young | 0.235172489 | 0.823744529 | Cone |
| NR2E3(+) | old | 0.429330473 | 0.950736281 | Cone |
| NR2E3(+) | young | 0.319700033 | 1.617344077 | Cone |
| NR2F1(-) | old | 0.501195887 | 1.356933407 | Cone |
| NR2F1(-) | young | 0.245959682 | 0.925021706 | Cone |
| PAX6(+) | old | 0.425318511 | 0.928059901 | Cone |
| PAX6(+) | young | 0.255146133 | 1.01127008 | Cone |
| POU2F1(+) | old | 0.540981276 | 1.581808069 | Cone |
| POU2F1(+) | young | 0.313282054 | 1.557087919 | Cone |
| RAX(+) | old | 0.530449578 | 1.522280888 | Cone |
| RAX(+) | young | 0.335617409 | 1.76678676 | Cone |
| RXRG(+) | old | 0.538508064 | 1.567829 | Cone |
| RXRG(+) | young | 0.334007823 | 1.751674921 | Cone |
| SPI1(+) | old | 0.438476137 | 1.002429329 | Cone |
| SPI1(+) | young | 0.222997903 | 0.709441596 | Cone |
| TCF12(+) | old | 0.522932017 | 1.479790191 | Cone |
| TCF12(+) | young | 0.30271812 | 1.457906837 | Cone |
| YBX1(+) | old | 0.522800576 | 1.479047262 | Cone |
| YBX1(+) | young | 0.342627592 | 1.832602915 | Cone |
| ZNF175(-) | old | 0.29206134 | 0.174864769 | Cone |
| ZNF175(-) | young | 0.298076687 | 1.414330042 | Cone |
